# Peptidome Analysis of Western Blots Identifies Natural Bispecific Antibody-Bound *Corynebacterium* and Phage B-cell Epitopes with Potential Relevance to Psoriasis

**DOI:** 10.64898/2026.03.02.708956

**Authors:** Jens-Michael Schröder

## Abstract

The high prevalence of *Corynebacterium (C.) simulans* in lesional and non-lesional psoriatic skin, and its correlation with disease severity, suggest a potential role in psoriasis pathophysiology. Previous exploratory Western blot and peptidome analyses of *C. simulans* extracts using IgG from psoriasis patients identified B-cell epitopes bound by natural bispecific antibodies (nBsAbs, presumably IgG4). These epitopes were primarily derived from intrinsically disordered proteins, autoantigens, and bacteriophage proteins. Subsequent analyses using pooled psoriasis serum unexpectedly revealed antigenic peptides from numerous *Corynebacterium* species. Taxonomic filtering identified several thousand *Corynebacterium*-derived epitopes across 40 species, including *C. simulans*, *C. striatum*, *C. diphtheriae*, common skin commensals, environmental and food-associated strains, the plant pathogen *Clavibacter michiganensis*, and the zoonotic pathogens *C. pseudotuberculosis* and *C. ulcerans*. Additional epitopes originated from related genera (*Prescottella*, *Tsukamurella*, *Mycobacterium*, *Gordonia*, *Nocardia*, and *Rhodococcus*) as well as bacteriophages of the order *Caudovirales*. Among the identified peptides, 183 epitopes from 46 antigens mapped to 30S and 50S ribosomal proteins. Numerous additional epitopes derived from proteins involved in transcription, translation, aminoacyl-tRNA ligases (covering 17 amino acids), transcriptional regulation, RNA processing and degradation, sigma factors, and ribosome-associated proteins. Notably, 40 epitopes originated from highly conserved FoF1-ATP synthase subunits (α, β, γ, δ). One peptide containing the catalytic Walker A nucleotide-binding motif showed sequence identity with mitochondrial FoF1-ATP synthase, suggesting potential autoimmune cross-reactivity and implicating this enzyme complex as a psoriasis-associated autoantigen candidate. Thousands of further epitopes were identified from proteins involved in respiratory chain function, stress responses, bacterial immunity, membrane transport, chaperones, cell wall biosynthesis, proteases, and peptidases, particularly ATP-dependent Clp proteases. Antigens from biosynthetic and metabolic pathways represented the most abundant nBsAb targets. In addition, 368 DECOY peptides were assigned to bacteriophage proteins. The detection of epitopes from toxin-producing species such as *C. diphtheriae*, *C. pseudotuberculosis*, and *C. ulcerans* further supports a potential microbial contribution to psoriasis pathogenesis. This exploratory study presents a streamlined strategy for B-cell epitope mapping of *Corynebacterium* antigens using IDP-enriched antigen-IgG4 complexes. The approach holds promise for the development of B-cell epitope-based vaccines targeting microbial, viral, tumor, and allergen-associated diseases.

## 1 INTRODUCTION

Psoriasis is a common chronic inflammatory skin disease characterized by a wide range of clinical manifestations. Although its exact etiology and pathogenesis remain incompletely understood, current evidence indicates significant overlap with other inflammatory and autoimmune disorders. One emerging hypothesis suggests that commensal microbes may contribute to autoimmunity, with growing interest in the role of skin microbiota in disease initiation and progression.

Recent studies have consistently demonstrated a marked dysbiosis in psoriatic skin, including a reduction in microbiome diversity. Certain bacterial taxa, such as *Corynebacterium simulans*, *Corynebacterium kroppenstedtii*, *Finegoldia* spp., and *Neisseria* spp., are found in increased abundance in psoriatic lesions, whereas beneficial taxa such as *Burkholderia* spp., *Cutibacterium*, and *Lactobacilli* are significantly depleted^1,2,3,4,5,6,7,8^.

Particularly notable are species of the genus *Corynebacterium*, which have been shown to promote, in a mouse model and independently of other microbes, IL-23-dependent expansion and activation of a γδ T cell subset^9^. This immunomodulatory effect is conserved across multiple *Corynebacterium* species and is dependent on the expression of mycolic acid - an essential component of the cell envelope in the *Mycobacteriales* order. These findings implicate *C. simulans*, a newly identified skin-colonizing but not mucosa-associated species^10^, *C. kroppenstedtii*, and possibly other yet unidentified *Corynebacterium* species as potential contributors to psoriasis pathophysiology.

*Corynebacterium* species are increasingly recognized as opportunistic pathogens of clinical relevance^11,12,13^. Unlike the lipophilic *Corynebacterium* taxa that dominate healthy skin microbiota, *C. simulans* is non-lipophilic, facultatively anaerobic, and fermentative, displaying a diphtheroid morphology^10^. It is a rare resident of healthy skin, in contrast to its elevated prevalence in psoriatic conditions.

Biochemically, *C. simulans* contains high levels of meso-diaminopimelic acid (DAP) in its peptidoglycan, a hallmark feature of the genus and shared with other *Mycobacteriales*^10,14^. DAP-containing peptidoglycans are specifically recognized by the intracellular receptor NOD1, while muramyl dipeptide structures are recognized by NOD2. NOD1-stimulatory activity has been shown to be present in the supernatant of *Corynebacterium* cultures^15^ and remains stable even under extreme conditions^15^, suggesting persistent stimulation of NOD1 signaling at colonized skin sites. Slc46a2, a highly expressed transporter in mammalian epidermal keratinocytes, was shown to be critical for the delivery of DAP-muropeptides and activation of NOD1 in keratinocytes^16^. In a mouse model, this transporter- and NOD-1-deficiency strongly suppressed psoriatic inflammation, whereas methotrexate, a commonly used psoriasis therapeutic, inhibited Slc46a2-dependent transport of DAP-muropeptides^17^.

NOD1 activation in keratinocytes induces type I interferon (IFN-β) responses, upregulates chemokines such as CXCL8-11, and activates NF-κB, leading to increased expression of key inflammatory mediators (such as IL-1β, IL-8, IL-18, IL-36, S100A7, S100A8, S100A9, and hBD-2) - all of which are prominently expressed in psoriatic lesions^18,19,20,21,22^. Experimental models show that NOD1 ligand administration leads to chemokine production and neutrophil recruitment^23^, effects abolished in NOD1-deficient mice^24^. This pathway is essential for neutrophil recruitment in response to bacterial infections such as *Clostridium difficile*, further highlighting NOD1’s role in epithelial immune responses^15^. Moreover, IL-36 cytokines, which are elevated in psoriatic lesions^25,26^ and are indirectly induce hBD-2^27^ - abundantly present in lesional psoriatic scales^28^ - are critical regulators of epithelial immune responses. They act as gatekeepers by distinguishing harmless commensals from invasive pathogens^29^.

Together, these findings suggest that *C. simulans*, despite being a rare commensal of healthy skin, may act as a potent immunological trigger in psoriasis through sustained activation of NOD1 and IL-36-mediated pathways. Its increased abundance in both lesional and non-lesional psoriatic skin^5,30^ correlates with disease severity and supports the hypothesis that in addition, other selected *Corynebacterium* species may contribute to the etiology and progression of psoriasis.

## 2. MATERIALS AND METHODS

### 2.1 Psoriasis Serum

A pooled surplus serum from hospitalized psoriasis patients was used in this exploratory study. Serum samples had originally been collected for chemokine determination in a clinical study (1994 -1998), which was approved by the local ethics committee^31^. The pooled serum was stored below -70 °C until use.

### 2.2 Culture of *Corynebacterium simulans*

*C. simulans* (DSM 44415) was cultivated for 24 h under planktonic conditions (shaking at 37 °C, 170 rpm) in PYG medium 104 (pH 7.2), according to the guidelines of the Leibniz Institute DSMZ-German Collection of Microorganisms and Cell Cultures.

### 2.3 Culture of *Corynebacterium kroppenstedtii*

*C. kroppenstedtii* (DSM 44385) was cultured for 24 h under planktonic conditions (shaking at 37 °C, 170 rpm) in PYG medium 104 (pH 7.2), as recommended by DSMZ.

### 2.4 Bacterial Lysis

Bacterial cells were harvested by centrifugation (5000 rpm, 10 min). After removal of the supernatant, the pellets were frozen at -70 °C, resuspended in 1 ml MQ-grade water, and lysed with 1% Triton X-100.

### 2.5 Heat Lysis

For heat lysis, bacterial cells were resuspended in 1% Triton X-100 and incubated at 95 °C for 10 min, followed by cooling on ice for 10 min. Cell debris was pelleted (10,000 × g, 30 min), and heat lysates were used immediately.

### 2.6 Western Blot Analysis

Bacterial lysates (untreated, heat-treated, or sonicated using three 10-s bursts at high intensity) were prepared in loading buffer containing 50 mM DTT. Samples (14 µl each) were separated on a 12% (37.5:1 acrylamide:bisacrylamide) gel and transferred onto a nitrocellulose membrane (0.2 µm pore size; Schleicher & Schuell BioScience, Dassel, Germany) using alkaline transfer buffer (48 mM Tris, 39 mM glycine, 0.0375% SDS, 20% ethanol; pH 9.2).

An alkaline buffer was essential for efficient transfer of cationic proteins. Membranes were blocked for 1 h in 5% (w/v) BSA in PBS/Tris (pH 7.4, 0.05% Tween), washed, and incubated with pooled psoriasis serum (1:100 in PBS/Tris) for 3 h. After washing, membranes were incubated with goat anti-human IgG-HRP (Jackson; 1:10,000) overnight at 4 °C. Following six additional washes with PBS/Tris (pH 7.4), membranes were developed with Lumilight substrate (Roche, Cat. No. 12015196001) and imaged using a Diana III Digital CCD Imaging System.

### 2.7 Proteomic and Peptidomic Analyses

Proteomic and peptidomic analyses for this exploratory study were conducted in 2014 at the Proteomics Facility, King’s College London. For in-gel digestion, the 16 kDa band detected in *C. simulans* Western blots was excised from nitrocellulose membranes, washed with triethylammonium bicarbonate (TEAB), reduced with dithiothreitol (DTT), alkylated with iodoacetamide, and digested on-membrane using sequencing-grade bovine trypsin (Sigma). The resulting peptides were analyzed by LC/ESI mass spectrometry.

As the primary objective of this exploratory study was the identification of the 16 kDa *C. simulans* antigen by proteomic analysis rather than comprehensive peptidomic profiling, all identified peptides originated from tryptic digestion. Consequently, these peptides may not represent the full-length sequences of antibody-bound B-cell epitopes, which would be expected from a true peptidomic approach that omits enzymatic digestion.

### 2.8 Data Analysis

LC/MS/MS data were processed using Scaffold Software™ (v5.3.0; Proteome Software, Portland, OR). Raw spectra (Mascot v2.2.06) were searched against a *Corynebacterium* protein database (corynebacterium.fasta; 258,295 entries; version not specified). Because all samples were digested with trypsin, naturally occurring peptides from the antibody-bound *Corynebacterium* immunopeptidome may have been truncated at the N- and/or C-termini. Identified peptide sequences were subsequently aligned against protein databases using BLASTp.

### 2.9 Protein Basic Local Alignment Search Tool (BLASTp) Analyses

Peptides were queried against non-redundant protein sequences within the family *Corynebacteriaceae* (taxid: 1653). If no 100% sequence match was identified, the search was expanded to the phylum *Actinomycetota* (taxid: 201174). BLASTp searches were performed using default parameters, with a maximum of 100 target sequences and automatic parameter adjustment for short query sequences. When all 100 target sequences showed 100% identity to the epitopic peptides, selected antigenic peptides were further analyzed for their presence in specific *Actinomycetota* species. DECOY peptides were searched exclusively against proteins belonging to bacteriophages of the class *Caudoviricetes* (taxid: 2731619).

In selected cases, protein sequences from different species were aligned for comparative analysis.

## 3 RESULTS AND SPECIFIC DISCUSSION

### 3.1 Western Blot Analysis of *C. simulans* Intrinsically Disordered Antigenic Proteins

Based on earlier findings, it was hypothesized that *C. simulans* may trigger immune responses in psoriasis. This was examined in a previous exploratory study using Western blot (WB) analysis and purified IgG from pooled psoriasis serum to assess immunoreactivity against *C. simulans* antigens. Interestingly, immunoreactive WB bands were detected only in boiled *C. simulans* extracts^32^. This observation supports the hypothesis that the thermostable antigens of *C. simulans* are intrinsically disordered proteins (IDPs) - proteins that lack a stable tertiary structure, tend to oligomerize, and are released as monomers upon boiling^33^.

To eliminate serum protein contamination in proteomic analyses of WB bands, purified IgG was used in the initial study^32^. Efforts to characterize the *C. simulans* 16 kDa antigen through proteomic peptide mapping unexpectedly identified individual 8- to 36-mer peptides of microbial origin. These peptides derived from *C. simulans*, *C. kroppenstedtii*, autoantigens, and putative bacteriophage-derived "DECOY" peptides^32^. The most plausible explanation for these findings is the presence of natural bispecific IgG antibodies (nBsAb) that bind epitopic peptides from bacterial, viral, and autoantigenic sources present in psoriasis serum.

Analysis of WB/peptidome data using pooled psoriasis serum and *C. simulans* extracts revealed numerous serum-derived contaminants alongside distinct peptides from various *Corynebacterium* species. This led to the hypothesis that nBsAbs, directed against *C. simulans* antigens, may also bind epitopes from other members of the *Mycobacteriales* order. To test this, WB was repeated with boiled *C. simulans* extracts and pooled psoriasis serum. A prominent 16 kDa band reappeared, along with additional bands at 40, 45, and 90 kDa (**Figure 1**), consistent with previous results using purified IgG^32^. Sonication, which similarly disrupts IDPs^34^, produced a comparable banding pattern (**Figure1**).

**Figure 1.**
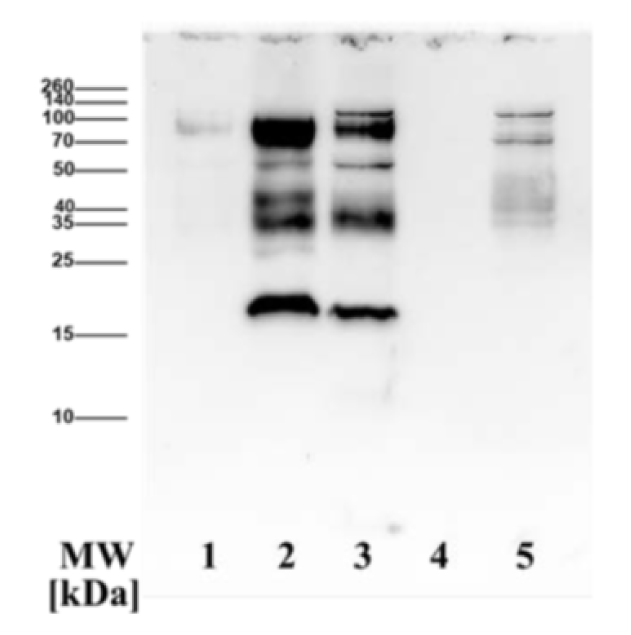
Western Blot Analysis of *Corynebacterium simulans* Protein Extracts Probed with Pooled Psoriasis Serum. *C. simulans* (DSM 44415) and *C. kroppenstedtii* (DSM 44385) were cultured under planktonic conditions, lysed using bacterial lysis buffer, and protein extracts were prepared either untreated or subjected to heating (95 °C) or sonication. Samples were mixed with DTT-containing loading buffer and separated by SDS-PAGE on a 12% acrylamide:bisacrylamide gel. **Lane MW** (Spectra™ Protein Ladder); **lane 1**, *C. simulans* extract without heat treatment; **lane 2**, heat-treated *C. simulans* extract; **lane 3**, sonicated *C. simulans* extract; **lane 4**, heat-treated *C. kroppenstedtii* extract; **lane 5**, sonicated *C. kroppenstedtii* extract. Western blotting was performed using pooled, non-specified psoriasis serum. Antigen-bound psoriasis IgG was detected using horseradish peroxidase-conjugated goat anti-human IgG and visualized with LumiLight (Roche). Immunoreactive bands were observed exclusively in heat-treated or sonicated samples of *C. simulans* extracts, consistent with conditions known to disrupt complexes of intrinsically disordered proteins or regions (IDP(R)s).

Notably, boiled extracts of *C. kroppenstedtii* - another abundant *Corynebacterium* species in psoriatic lesions^30^ - showed no WB bands **(Figure 1)**, suggesting a species-specific immune response to *C. simulans*. The 16 kDa band was excised and subjected to trypsin digestion and liquid chromatography/mass spectrometry (LC/MS) analysis. As in the previous study, proteomic analysis revealed a high number of *C. simulans* proteins but failed to identify a single definitive antigen. To reduce serum-derived peptide interference, data were filtered for peptides matching proteins from the *Corynebacterium* genus. Thousands of peptide hits were subsequently identified and characterized by BLASTp for their microbial origin.

### 3.2 Psoriasis Serum Contains Natural Bispecific Antibody-Bound Epitopic Peptides from Numerous *Corynebacterium* Species

BLASTp analysis of the filtered peptide data revealed species-specific epitopic peptides originating from 40 different *Corynebacterium* species and nine additional members of the Actinomycetota phylum. In addition to *C. simulans*, the genetically very close *C. striatum*^35^ emerged as a major source of these epitopic peptides **(Table 1, Table S1)**. *C. striatum* is a non-lipophilic, fermentative Gram-positive bacillus with a striated appearance on Gram stain^36^. While commonly part of the healthy skin microbiota^37^, it is increasingly recognized as an opportunistic pathogen in both community-acquired and nosocomial infections^38,39^, including prosthetic joint infections^40^ and endocarditis^41^. The recent demonstration of its robust intracellular invasion and cytotoxic activity in human airway epithelial cells^42^ highlights an important virulence trait that enables intracellular survival in epithelial cells and underscores its potential role in the pathophysiology of psoriasis.

**Table 1.** Corynebacterium species-specific Antigens and Epitopes*.

| *Corynebacterium* Species and Other Actinomycetota | Number of Antigens | Number of Epitopes | Comments |
| --- | --- | --- | --- |
| *C. accolens* | 14 | 17 | It is considered as an inhabitant of skin and the upper respiratory tract and has been isolated from a variety of human clinical specimens. |
| *C. ammoniagenes* | 6 | 6 | A soil bacterium that produces amino acids and was isolated from stool of an infant. |
| *C. amycolatium* | 8 | 8 | A common component of the natural microflora found on human skin and mucous membranes. |
| *C. argenteoratense* | 8 | 8 | Isolated from throat specimens of patients with tonsillitis. |
| *C. aurimucosum* | 7 | 7 | Isolated from healthy female urogenital tract samples and is considered part of the normal microbial community. |
| *C. bovis* | 1 | 1 | A natural commensal in the bovine udder, and as such, is usually isolated from bovine mastitis. |
| *C. callunae* | 8 | 8 | The most likely habitat of *C. callunae* is soil around heather plants. |
| *C. casei* | 6 | 7 | Isolated from cheese. |
| *C. diphtheriae* | 19 | 24 | Infection is spread by droplets, secretions, or direct contact solely among humans. It may carry the diphtheria toxin gene (*tox*). |
| *C. durum* | 11 | 11 | Isolated from human sputum. |
| *C. efficiens* | 8 | 8 | Isolated from soil and vegetables. |
| *C. falsenii* | 14 | 15 | Isolated from the respiratory tracts of eagles and black storks and from bioaerosols sampled in duck houses. |
| *C. genitalium* | 6 | 6 | Isolated from human semen from patient with prostatitis. |
| *C. glucuronolyticum* | 13 | 13 | Isolated from human semen from patient with prostatitis. |
| *C. glutamicum* | 23 | 23 | This glutamate-producing *Corynebacterium* has been found in the avian feces-contaminated soil. |
| *C. glyciniphilum* | 13 | 13 | This *Corynebacterium* was isolated from putrefied bananas. |
| *C. halotolerans* | 16 | 16 | This halotolerant Actinobacterium was isolated from a soil sample from the west of China. |
| *C. jeikeium* | 11 | 11 | Is found in soil and water and is part of normal human skin flora. |
| *C. kefirresidentii* | 2 | 2 | Isolated from kefir grains. |
| *C. kroppenstedtii* | 7 | 7 | Isolated from a case of recurrent granulomatous mastitis. |
| *C. lipophiloflavum* | 5 | 5 | Isolated from a patient with bacterial vaginosis. |
| *C. maris* | 14 | 16 | A salt-tolerant species originally isolated from the mucus of a coral. |
| *C. matruchotii* | 20 | 21 | A bacterium of the oral cavity and comprises the central filament of "corn-cob formations" (formations in which *Streptococcus sanguinis* bacteria bind to and surround *C. matruchotii* to create a corn-cob appearance). |
| *C. otitidis* | 1 | 1 | Isolated from middle ear fluid from otitis media patient. |
| *C. pseudodiphtheriticum* | 13 | 13 | Forming part of the upper respiratory tract flora. |
| *C. pseudotuberculosis* | 17 | 17 | A zoonotic pathogen, forming part of the upper respiratory tract flora, causes lymphadenitis in humans, livestock, and wildlife. It may carry the diphtheria toxin gene (*tox*). |
| *C. pyruviciproducens* | 4 | 4 | Isolated from clinical sources (groin abscess). |
| *C. resistens* | 5 | 5 | Isolated from clinical sources. |
| *C. sanguinis* | 1 | 1 | Isolated from clinical and environmental sources. |
| *C. simulans* | 4 | 9 | A rare commensal of the skin, which is absent in mucous membranes. |
| *C. striatum* | 48 | 146 | Commonly occupies the normal flora of the skin and oropharynx. |
| *C. terpenotabidum* | 7 | 7 | Was isolated from soil based on its ability to degrade squalene. |
| *C. tuberculostearicum* | 2 | 2 | Is the predominant species found on human skin. |
| *C. ulcerans* | 7 | 7 | Isolated from a wide host of domestic and wild animals. It may carry the diphtheria toxin gene (*tox*). |
| *C. urealyticum* | 12 | 12 | An important isolate when found in conjunction with a urinary tract infection. |
| *C. variabile* | 14 | 14 | Is part of the complex microflora on the surface of smear-ripened cheeses. |
| *C. vitaeruminis* | 12 | 12 | Was isolated from rumen. |
| *Actinobaculum massiliense* | 1 | 1 | Isolated from urine and skin. |
| *Brevibacterium flavum* | 1 | 1 | Associated with the surface of bacterial ripened cheeses. |
| *Clavibacter michiganensis subsp.* | 16 | 16 | Cause the disease called bacterial wilt, canker of tomato and ring rot of potato. |
| *Corynebacterium* sp. KPL1814 | 1 | 1 | ? |
| *Corynebacterium* sp. KPL1818 | 2 | 2 | ? |
| *Corynebacterium* sp. CCUG 65737 | 1 | 1 | ? |
| *Gordonia malaquae* | 1 | 1 | Isolated from sludge of a wastewater treatment plant. |
| *Mycobacterium tuberculosis* | 2 | 2 | Pathogenic bacterium and the causative agent of tuberculosis. |
| *Nocardia zapadnavensis* | 1 | 1 | Isolated from soil. |
| *Prescottella equi* | 36 | 36 | An intracellular pathogen that causes pyogranulomatous pneumonia in thoroughbred foals. It is commonly found in dry and dusty soil. |
| *Rhodococcus* sp. AG1013 | 1 | 1 | ? |
| *Tsukamurella paurometabola* | 23 | 23 | saprophytic bacterium found in soil and water. |

\* Comprehensive data are provided in Table S1.

Other abundant epitopic peptides originated from *C. accolens*, a nasal commensal known to inhibit *Staphylococcus aureus*-induced mucosal barrier disruption^43^. Additional peptides came from the skin-associated commensals *C. amycolatum*, *C. halotolerans*, and *C. jeikeium*, which have all been implicated in hospital-acquired infections^44^. Less abundant but still notable peptides were identified from *C. kefirresidentii*, *C. kroppenstedtii*, and *C. tuberculostearicum*, three predominant *Corynebacterium* species found on human skin^45^.

Epitopic peptides were also derived from oral microbiota members such as *C. durum* and *C. matruchotii*. These bacteria inhabit the diverse environments of the oral cavity, including the gingival sulcus and tongue^46^. Of particular interest, *C. matruchotii* is a calcifying bacterium typically isolated from dental calculus, where it promotes mineralization^47^. It has been negatively associated with severe periodontitis and inflammatory markers^48^. Psoriasis patients have been reported to exhibit more severe gingivitis, bone loss, dental caries, and tooth loss compared to non-psoriatic individuals, with disease severity correlating with both psoriasis and psoriatic arthritis^49,50^.

Additional peptides originated from *C. argentoratense*, originally isolated from throat swabs in tonsillitis patients, and genetically clustered with toxigenic species such as *C. diphtheriae*, *C. ulcerans*, *C. pseudotuberculosis*, and *C. kutscheri*^51^. Operational taxonomic units showing 99% nucleotide sequence identity to the 16S rRNA gene of *C. argentoratense* were obtained from several distinct skin sites of healthy humans in the course of the Human Microbiome Project^52^ and in a study of the skin microbiome associated with atopic dermatitis in children^53^.

Interestingly, a peptide from *C. bovis* - a bovine commensal but also a major mastitis pathogen^54^ - was also identified. In animal models, *C. bovis* has been shown to induce a psoriasis-like dermatitis with hyperkeratosis, acanthosis, and mononuclear infiltration in nude mice^55,56(p20),57^, making it a compelling subject for future psoriasis research.

Peptides from *C. aurimucosum*, *C. genitalium*, *C. glucuronolyticum*, *C. lipophiloflavum*, and *C. urealyticum* reflect immune recognition of genitourinary commensals. Moreover, abundant epitopic peptides from *Corynebacterium* spp. such as *C. ammoniagenes*, *C. callunae*, *C. efficiens*, *C. falsenii*, *C. glutamicum*, *C. jeikeium*, *C. maris*, *C. sanguinis*, and *C. terpenotabidum*, as well as from other Actinomycetota including *Gordonia malaquae*, *Nocardia zapadnayensis*, *Prescottella* (*P*.) *equi* (syn. *Rhodococcus equi*), and *Tsukamurella* (*T*.) *paurometabola*, suggest immune responses to environmental soil bacteria, even in the absence of known infections.

*C. falsenii*, typically associated with birds and rarely found in humans^58^, was isolated from duck house aerosols^59^. Among the most abundant peptides - aside from those from *C. striatum* - were from *P*. *equi* and *T*. *paurometabola* **(Table 1, Table S1)**. *P. equi*, a facultative intracellular coccobacillus that can survive within human alveolar epithelial cells^60^, is a well-known veterinary pathogen that can infect immunocompetent humans^61,62^, raising the possibility of a zoonotic link to psoriasis.

*Tsukamurella* species, environmental saprophytes related to *Nocardia*, *Rhodococcus*, and *Mycobacterium*, are emerging opportunistic pathogens, especially in immunocompromised hosts or those with indwelling medical devices^63,64^. Though rare, cutaneous infections with *T. paurometabola* have been reported even in immunocompetent individuals^65,66^, suggesting a potential, yet unproven, role in psoriasis.

Numerous peptides were also attributed to *Actinomyces* spp. **(Table 1, Table S1)**, which may have originated from dietary sources. Non-pathogenic species such as *C. casei*, *C. variabile*, and *Brevibacterium flavum* are commonly found on the surfaces of smear-ripened cheeses^67^.

*C. glyciniphilum*, initially isolated from decayed bananas, is known to metabolize serine into glycine^67^.

Of particular note, a high number of epitopic peptides originated from *Clavibacter michiganensis* proteins **(Table 1, Table S1)**, a group of phytopathogenic Actinomycetes known to infect economically significant crops such as tomatoes (*C. michiganensis* subsp. *michiganensis*) and potatoes (*C. michiganensis* subsp. *sepedonicus*)^68,69^. These likely represent food-derived peptides rather than human pathogens.

Most strikingly, abundant epitopic peptides were identified from the human pathogens *C. diphtheriae* (24 epitopes) and the zoonotic pathogens *C. ulcerans*^70^ (7 epitopes) and *C. pseudotuberculosis*^71^ (17 epitopes), - all known diphtheria toxin (DT)-producing, facultative intracellular pathogenic bacteria^72^ **(Table 1, Table S1)**. DT is encoded by a *tox* gene acquired via a prophage and integrated into the *Corynebacterium* genome through site-specific recombination^73^. While DT-producing strains are vaccine-preventable, non-toxigenic *C. diphtheriae* and related species are emerging pathogens capable of causing severe disease^74(p20),75^.

Among *C. diphtheriae* biotypes, *belfantii* can be toxigenic^76,77^. The biotypes *gravis*, *mitis*, and *intermedius* are associated with disease severity^73^. It is conceivable that these toxigenic *Corynebacterium* species contribute to psoriasis pathophysiology. Upon prophage activation, DT may be produced and bind to heparin-binding epidermal growth factor-like growth factor (HB-EGF), which is highly expressed in keratinocytes^78,79^ and implicated in psoriasis^80^. DT induces ribotoxic stress and activates the NLRP1 inflammasome in keratinocytes^81^, a mechanism increasingly recognized in psoriasis^82,83^. NLRP1 activation in human keratinocytes within organotypic skin cultures has been shown to induce a psoriasis-like phenotype^84^. Furthermore, DT selectively kills stratum granulosum keratinocytes by inhibiting protein synthesis through ADP-ribosylation^85^. This destruction may release precursors of cationic intrinsically disordered antimicrobial peptides (CIDAMPs), such as HRNR repeats and FLG spacer regions - both previously identified as nBsAb-bound autoantigens in psoriasis^32^.

Collectively, these findings support the hypothesis that toxigenic *Corynebacterium* species in psoriatic skin, potentially activated by prophages, produce a yet-unidentified DT-like “psoriasis toxin.” This proposed toxin may be encoded by phages infecting *C. simulans*^32^, offering a novel perspective on the pathogenesis of psoriasis.

### 3.3 Ribosomal Proteins of *Corynebacterium* Taxa Are Major Targets of nBsAbs in Psoriasis

BLASTp analysis of epitopic peptides identified by WB/peptidome experiments revealed 183 peptides derived from both the small (30S) and large (50S) ribosomal subunits of *Corynebacterium* taxa **(Table 2, 3; Table S2, S3)**. Notably, 46 out of the 53 ribosomal proteins typically expressed in prokaryotes^86^ were identified as antigens. Among the *Corynebacterium* species, *C. striatum* stood out, with ribosomal proteins representing the dominant nBsAb targets - 88 epitopic peptides mapped to six ribosomal protein antigens from both the 30S and 50S subunits **(Table 2, 3; Table S2, S3)**.

**Table 2.** C*o*rynebacterium 30S-Ribosomal Proteins are Antigens*.

| Antigen | Epitopes conserved in *Corynebacterium* species | Number of Epitopes |
| --- | --- | --- |
| 30S ribosomal protein S1 | 20 | 21 |
| 30S ribosomal protein S2 | 4 | 8 |
| 30S ribosomal protein S3 | 2 | 12 |
| 30S ribosomal protein S4 | 2 | 15 |
| 30S ribosomal protein S5 | 2 | 7 |
| 30S ribosomal protein S6 | 2 | 3 |
| 30S ribosomal protein S7 | 1 | 9 |
| 30S ribosomal protein S8 | 4+1 | 5 |
| 30S ribosomal protein S9 | 3 | 4 |
| 30S ribosomal protein S10 | 14 | 3 |
| 30S ribosomal protein S11 | 11 | 3 |
| 30S ribosomal protein S12 | 18 | 4 |
| 30S ribosomal protein S13 | 3 | 3 |
| 30S ribosomal protein S14 | 6 | 2 |
| 30S ribosomal protein S15 | 2+2 | 2 |
| 30S ribosomal protein S16 | 4 | 5 |
| 30S ribosomal protein S17 | 2+3+14 | 4 |
| 30S ribosomal protein S18 | 4 | 1 |

\* Comprehensive data are provided in Table S2.

**Table 3.**
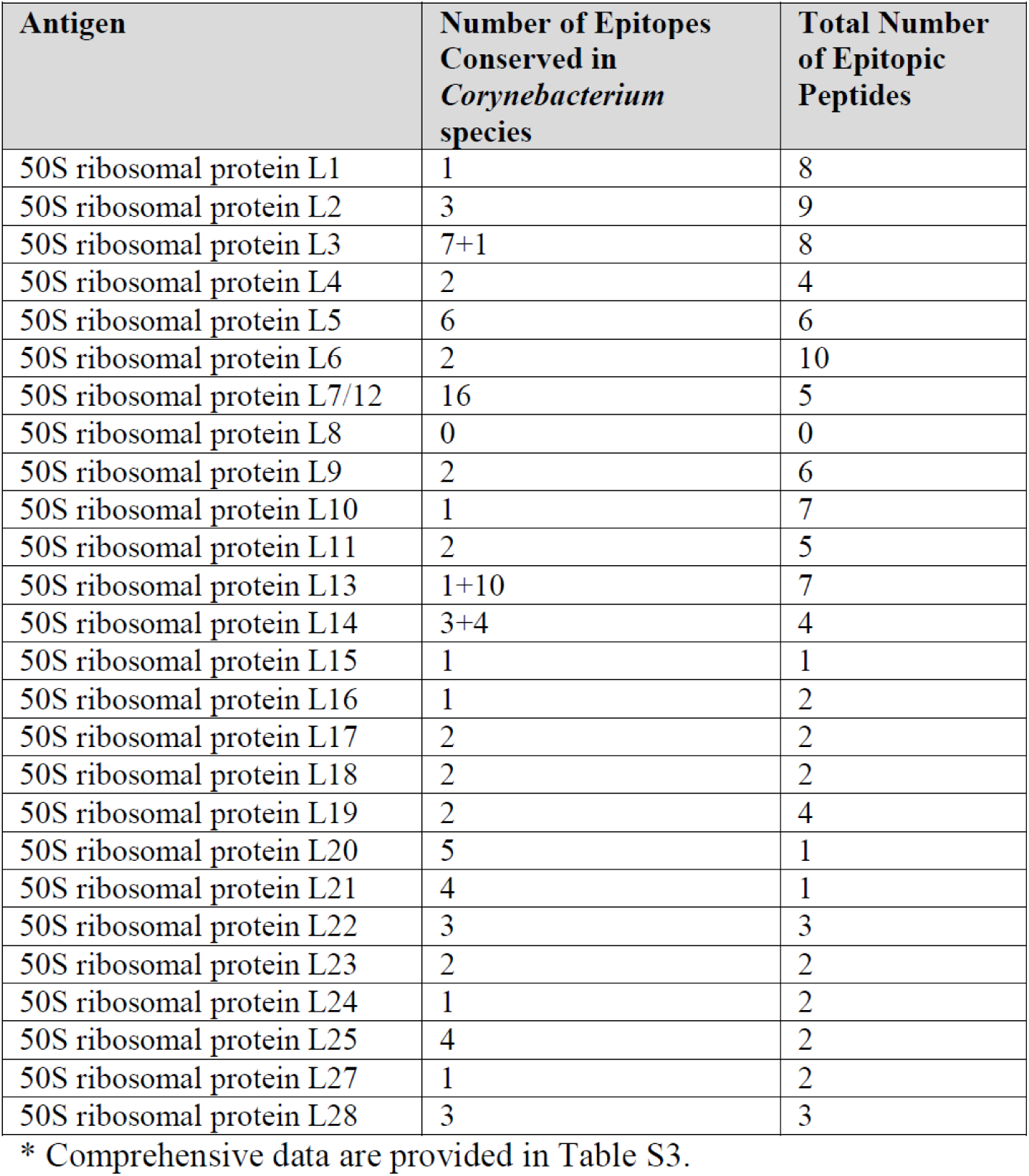
Corynebacterium 50S-Ribosomal Proteins are Antigen.

Interestingly, 21 epitopic peptides originated from the 30S ribosomal protein S1 and were conserved across 20 *Corynebacterium* species **(Table S2)**. Ribosomal protein S1, the largest protein of the 30S subunit in *E. coli*^87^, has a high affinity for mRNA^88^ and is implicated in translation initiation^87^. Its strong immunogenicity and cross-species conservation suggest that S1 functions as an immunodominant antigen within *Corynebacterium*.

Additional immunodominant candidates included 10 peptides from the 50S ribosomal protein L6 and 5 peptides from the 50S ribosomal protein L7/L12, the latter being conserved across 18 *Corynebacterium* species (**Table** S2). The L7/L12 protein plays a critical role in translational fidelity by facilitating proofreading steps during protein synthesis^89^

Immunoproteomic studies in *Nocardia seriolae* - a pathogen responsible for fish nocardiosis - have also identified the L7/L12 protein as a common immunodominant antigen across three fish-pathogenic *Nocardia* species, underscoring its potential as a DNA vaccine target^90^. Furthermore, the L7/L12 protein from *Mycobacterium tuberculosis* functions as a TLR4 agonist and enhances CTL and HTL activation through dendritic cell maturation, positioning it as a promising adjuvant in vaccine development^91^.

A particularly striking finding in this study was that epitopes of *Corynebacterium* ribosomal proteins L5, L7/L12, and L14 share 70–78% sequence identity with those of *Akkermansia muciniphila* **(Table S3)**, a gut commensal known for its beneficial roles in host-microbiota interactions and gut health^92^. Given that psoriasis shares immunological features with inflammatory bowel diseases - such as Th17-driven immune responses and inflammatory cytokine profiles - and that *A. muciniphila* abundance is significantly reduced in psoriasis patients^93^, this unexpected molecular overlap may provide novel insights into the pathophysiology of psoriasis.

### 3.4 Multiple *Corynebacterium* Transcription and Translation Proteins Are Antigens

BLASTp analysis of nBsAb-bound peptides identified 183 epitopic peptides derived from 55 transcription and translation proteins across multiple *Corynebacterium* species. These proteins include DNA-directed RNA polymerases, translation initiation factors, topoisomerases, primases, helicases, methyltransferases, DNA polymerase III, and others **(Table S4)**. Notably, the highest numbers of epitopic peptides were associated with DNA-directed RNA polymerases: 25 originated from the RNA polymerase β subunit (*rpoB*) and 33 from the β’ subunit (*rpoC*). BLASTp searches revealed that *rpoB*-derived peptides were conserved in up to 88 *Corynebacterium* species and *Mycobacterium tuberculosis*, while *rpoC*-derived peptides were found in up to 81 species **(Table S4)**. These peptides are largely conserved across the *Corynebacterium* genus and partially in other members of the order Mycobacteriales.

Bacterial RNA polymerase is the target of the broad-spectrum antibiotic rifampicin, which binds the β subunit of *rpoB*, inhibiting DNA transcription and serving as a frontline treatment for tuberculosis^94,95^. Mutations in *rpoB* confer rifampicin resistance^96^. Given these parallels, antibodies targeting *rpoB* could potentially mimic rifampicin’s inhibitory mechanism, making *rpoB*-derived epitopic peptides promising candidates for peptide-based tuberculosis vaccines^97^. Indeed, 7 *rpoB*- and 9 *rpoC*-derived peptides showed sequence identity with *M. tuberculosis* homologs **(Table S4)**. Another noteworthy candidate is the *Mycobacterium*-specific peptide **EALFDAFR**, derived from the DNA repair helicase XPB (antigen no. 15), which plays a role in nucleotide excision repair^98^.

Additional RNA-related enzymes were identified as antigens. Twenty-six antigens comprising 28 epitopic peptides originated from RNA methyltransferases, RNA helicases, and other enzymes **(Table S5)**. Of these, ten peptides were species-specific. The epitope peptide **VDDLSKR** from a DEAD/DEAH-box helicase, potentially derived from *Gordonia malaquae*^99^, is conserved in many Actinomycetes. While early *Gordonia* isolates were considered opportunistic pathogens^100^, recent environmental strains play key roles in bioremediation.

Seventeen aminoacyl-tRNA synthetases (aaRSs), recognizing all canonical amino acids except His, Gln, and Asn, were also identified as antigens. These enzymes are universally conserved, yet exhibit specific variations among kingdoms. Notably, *Corynebacterium* species appear to possess non-discriminating aaRSs to compensate for missing Asn- or Gln-RSs^101^. While only 1 - 3 epitopic peptides were found for Ala-, Phe-, and Tyr-RSs, Asp-RSs and Glu-RSs yielded 21 and 32 peptides, respectively, spanning up to 50 and 42 species **(Table S6)**. Given their essential role in protein biosynthesis and kingdom-specific sequences, aaRSs are viable antimicrobial targets - such as Ile-RS is the target of mupirocin^102^ - and therefore promising components for broad-spectrum, serotype-independent peptide vaccines.

BLASTp searches also revealed nBsAb-bound peptides from translation elongation factors EF-G, EF-Tu, EF-P, and Ts, as well as transcription elongation factor GreA **(Table S7)**. EF-G and EF-Tu, catalyzing core translation steps^103^, were major antigens, recognized by 17 and 14 peptides respectively. These highly conserved factors are essential for protein synthesis and represent attractive drug targets. EF-G is targeted by fusidic acid, active even against *M. tuberculosis*^104,105^. EF-Tu, targeted by four unrelated antibiotic families^106^, was recognized by conserved peptides in both *Corynebacterium* and *M. tuberculosis*. Thus, EF-G and EF-Tu peptides, including those mapping to functional domains (such as *83***DAPGH***87*, *138***NKCD***141*), offer potential for multivalent vaccine design against diphtheria and tuberculosis.

CRISPR-associated endonuclease Cas9 and RNA polymerase sigma factors were also identified as species-specific antigens **(Table S8, S9)**. Sigma factors, essential for transcription initiation, were represented by 7 antigens with predominantly single, species-specific epitopic peptides.

Enzymes of the RNA degradosome complex, such as RNase J, RNA helicases, and other ribonucleases, were also identified **(Table S10)**. In *C. diphtheriae*, RNase J modulates virulence gene expression^107^, highlighting the immunogenic potential of its epitope **SMQGADLTQR** (peptide no. 4, **Table S10**).

Several ribosome-associated proteins also elicited immune responses **(Table S11)**. The ribosome recycling factor RsfS, with 6 peptides from up to 37 species, plays a critical role in translation termination and cellular adaptation under nutrient stress^108,109^. The hibernation-promoting factor (HPF) was represented by 5 peptides from 35 species, functioning as a ribosome inactivator under stress^110^. HPF has been proposed as a potential drug target^111^. Similarly, RsfA and RbfA, both associated with translational regulation and stress response, were recognized by nBsAb-bound peptides^109,112^.

Lastly, 57 *Corynebacterium* transcriptional regulators were identified as antigens, representing diverse families such as TetR, MarR, GntR, MerR, IclR, and others **(Table S12)**. These regulators are key mediators of bacterial responses to antibiotics, toxic compounds, stress, and virulence, typically via conserved helix-turn-helix motifs^113^. Notably, the TetR family alone accounted for 13 antigens. Particularly significant was the identification of metal-dependent regulators, including the iron-responsive diphtheria toxin repressor (DtxR)^114^ (antigen no. 39, **Table S12**). DtxR-like regulators are highly conserved across both pathogenic and saprophytic *Corynebacterium* species^115^. Strikingly, the identified epitopes (antigen no. 39, **Table S12**) map close to residues required for binding of the activated repressor to the diphtheria tox operator^116^, suggesting that these nBsAbs may disrupt regulation of iron homeostasis in *Corynebacterium* species. Additional regulators identified as antigens include NusB, CarD, Rho, GlxR, and WYL domain-containing proteins.

This comprehensive analysis reveals that multiple transcription and translation-related proteins from *Corynebacterium* species serve as immunogenic antigens. Notably, highly conserved peptides from RNA polymerase subunits, elongation factors, aminoacyl-tRNA synthetases, and ribosomal regulators were frequently targeted by nBsAb. Several of these peptides also show homology with *M*. *tuberculosis*, supporting their potential as cross-protective, multi-epitope vaccine candidates.

### 3.5 *Corynebacterium* FoF1-ATP Synthase Is a Major nBsAb Target

A BLASTp search of nBsAb-bound peptides identified 40 epitopic peptides corresponding to the alpha, beta, gamma, and delta subunits of the membrane-bound FoF1-ATP synthase from over 86 *Corynebacterium* species, *M*. *tuberculosis* and *M. leprae* (**Table S13**). FoF1-ATP synthase is the final enzyme in the oxidative phosphorylation pathway, utilizing electrochemical energy to catalyze ATP synthesis. As a key component of cellular bioenergetics, ATP synthases are present in all domains of life, including the plasma membrane of bacteria, thylakoid membranes of chloroplasts, and the inner mitochondrial membrane in eukaryotes^117^.

The enzyme comprises two major sectors: the cytoplasmic F1 sector, responsible for ATP synthesis, and the membrane-embedded Fo sector, responsible for proton translocation and torque generation^118^. The catalytic F1 "head" includes three alpha and three beta subunits arranged around a central gamma subunit forming the rotor shaft, while the delta and epsilon subunits connect the stator and rotor components^119^ (**Figure 2**).

**Figure 2.**
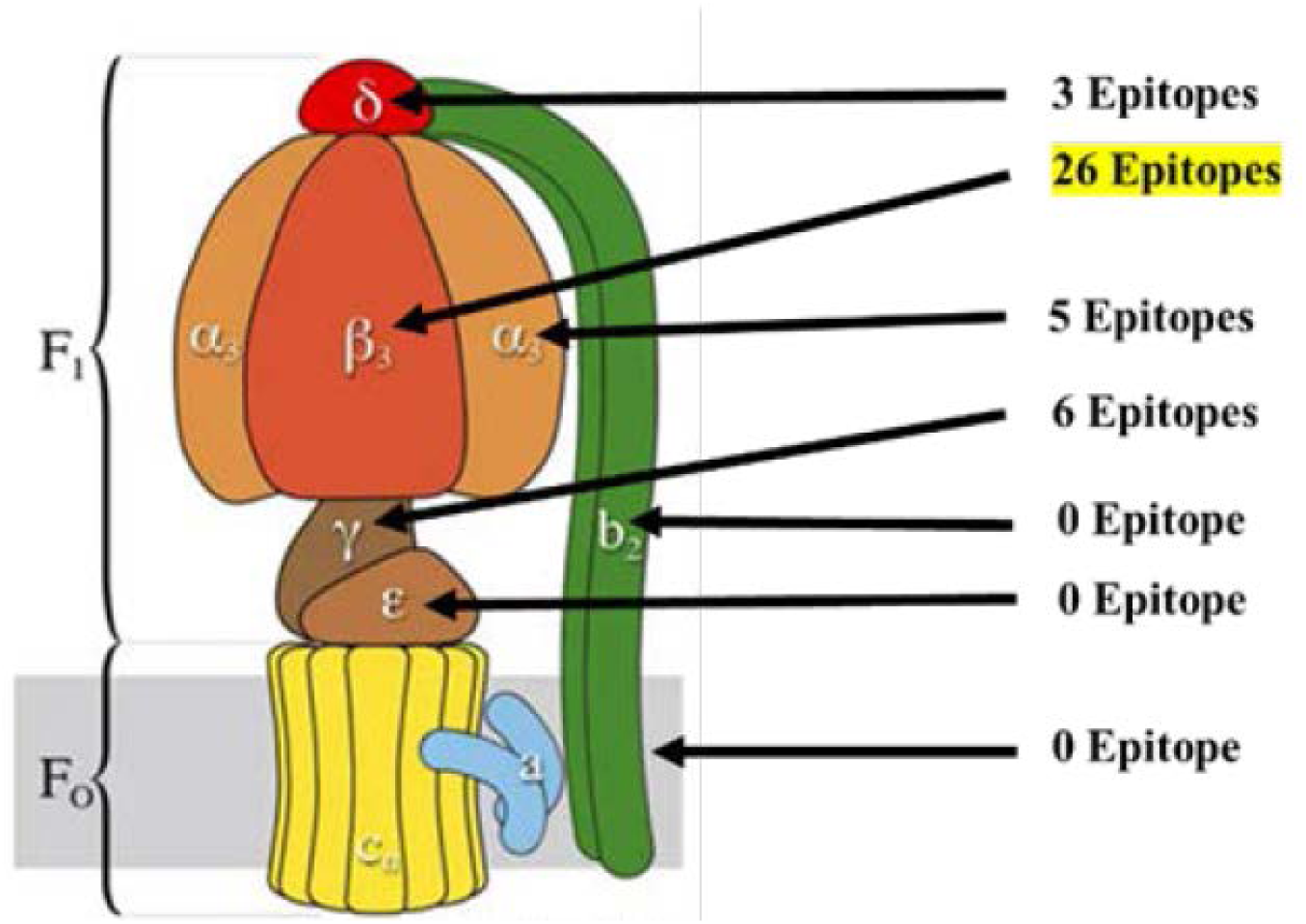
Schematic Representation of the Structural Organization of the F1Fo-ATP Synthase and Localization of Identified *Corynebacterium* B-cell Epitopes. Numerous epitopes of *Corynebacterium* F1Fo-ATP synthase were identified in the catalytic F1 domain, particularly in the β-subunit (**Table S13**), whereas none were detected in the membrane-embedded Fo subunits. The schematic was adapted from https://www.biophys.mpg.de/2051971 with permission.

The F1 sector emerged as the principal target of the immune response, with 26 epitopic peptides identified from the beta subunit and 5 from the alpha subunit (**Figure 2, Table S13**). The gamma subunit carried 5 epitopes, while the delta subunit contained 4. Notably, no epitopes were detected from the epsilon subunit or from proteins of the Fo sector (**Figure 2**, **Table S13**).

Many of these epitopic peptides are highly conserved, with beta subunit-derived peptides present in at least 63 *Corynebacterium* species, and several also found in *M. tuberculosis* and *M. leprae*. Alpha subunit peptides are conserved in 53 *Corynebacterium* species, with a subset shared with *M. tuberculosis* as well. Gamma subunit epitopes are found in at least 28 *Corynebacterium* species and *M. tuberculosis* (**Table S13**).

Strikingly, the beta subunit of *C. simulans* and *C. striatum* contains 26 epitopes, and that of *C. diphtheriae* harbors 12, suggesting the catalytic site of F1-ATP synthase is a major target of immune recognition. This aligns with structural data, as the catalytic site - located on the beta subunit at its interface with the alpha subunit - contains three nucleotide-binding sites^120^. One identified peptide (**IGLFGGAGVGKT**; peptide no. 9 and 13, **Table S13**) includes the Walker-A motif (**GXXXXGKT**), a highly conserved ATP-binding motif^121,122^. Another peptide (antigen no. 2, peptide no. 3; **Table S13**) contains residues involved in ATP and Mg² binding^123^.

Importantly, *M. tuberculosis* F1-ATP synthase is also recognized by nBsAb. Five beta subunit epitopes, including the Walker-A motif, along with two alpha subunit and one gamma subunit epitopes, were identified (**Table S13**). The F1 domain of ATP synthase is essential for the survival of *M. tuberculosis* and nontuberculous mycobacteria (NTM), particularly under hypoxic stress, and has been proposed as a high-value drug target for therapeutic development^124,125,126^.

The identification of conserved epitopic peptides from the catalytic subunits of FoF1-ATP synthase in both, *Corynebacterium* species and *M. tuberculosis*, highlights this enzyme as a dominant antigenic target of natural bispecific antibodies. Given its essential role in bacterial energy metabolism, structural accessibility, and evolutionary conservation, especially of functionally critical motifs like Walker-A, F1-domain-derived peptides are promising candidates for the development of multi-epitope peptide vaccines targeting *Corynebacterium* infections, tuberculosis and leprosy.

### 3.51 Epitopic Peptides Identify FoF1-ATP Synthase Alpha and Beta Chains as Autoantigens

FoF1-ATP synthases represent a highly conserved family of enzymes found across biological kingdoms, including in bacterial cytoplasmic membranes, chloroplast thylakoid membranes, and the inner mitochondrial membrane of eukaryotes^117^. In this exploratory study, analysis of epitopic peptides derived from various *Corynebacterium* FoF1-ATP synthases identified a highly conserved sequence, **IGLFGGAGVGK**, in the beta chain (peptide no. 9, **Table S13**). This sequence, including Mg²⁺ - and ATP-binding sites, shares significant homology with the mitochondrial FoF1-ATP synthase of *Homo sapiens* (**Figure 3**). Importantly, this region corresponds to the catalytic nucleotide-binding Walker motif A^121^, indicating that the detected nBsAb targets a fundamental component of cellular bioenergetics - possibly contributing to psoriasis pathophysiology.

**Figure 3.**
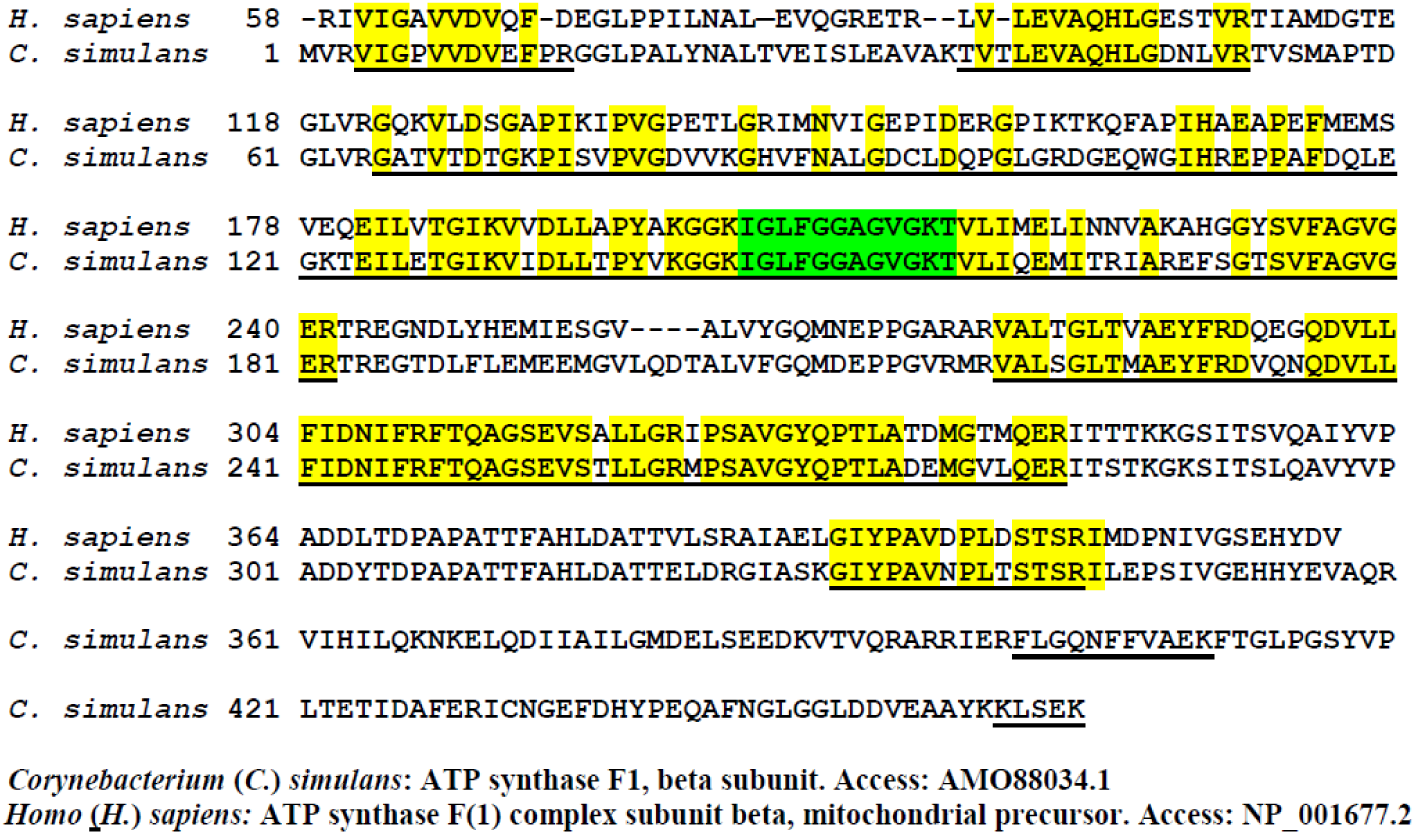
Sequence Alignment of *H. sapiens* and *C. simulans* ATP Synthase F1 β Subunit Identifies Autoantigenic Epitopes. A sequence alignment between the *C. simulans* ATP synthase β subunit and the human mitochondrial ATP synthase F1 β subunit is shown. Conserved amino acid residues between both sequences are highlighted in yellow or green. Sequences corresponding to nBsAb-bound *C. simulans* epitopic peptides (see **Table S13**) are underlined. The catalytic Walker-A motif-containing epitope is highlighted in green. Note the high degree of sequence conservation within several epitopic regions, representing potential autoantigenic epitopes.

Although historically thought to reside exclusively in mitochondria, FoF1-ATP synthase has also been identified on the plasma membrane of keratinocytes^127^. In normal human skin, its localization is layer-specific, with mitochondrial expression in basal keratinocytes and cell surface localization in more differentiated layers^128^. Notably, antibodies against the beta chain of ATP synthase have been shown to impair keratinocyte differentiation in vitro^128^, suggesting that this autoantibody, detected in psoriasis patient sera, may contribute to the altered differentiation phenotype observed in psoriatic lesions^129^.

Cell surface expression of FoF1-ATP synthase has been reported in various cell types, including keratinocytes^127^ and endothelial cells^130^. While mitochondrial ATP synthase functions in ATP production via oxidative phosphorylation, the surface-expressed form is implicated in alternative roles such as regulation of intracellular pH, response to anti-angiogenic factors, and cholesterol metabolism. A monoclonal antibody targeting the beta subunit (MAb3D5AB1) has been shown to inhibit the F1 domain of cell surface ATP synthase in endothelial cells^131^.

Autoantibodies against ATP synthase have also been detected in sera from patients with vasculitides^132^, implicating a pathogenic role in vascular inflammation - and potentially in psoriasis. Moreover, similar autoantibodies have been reported in both serum and cerebrospinal fluid of patients with Alzheimer’s disease^133^. In mouse models, anti-ATP synthase antibodies purified from Alzheimer’s patients caused cognitive deficits and hippocampal cell damage, indicating a potential contribution to disease progression^134^.

Beyond the Walker motif A, several additional nBsAb-bound epitopic peptides of the *Corynebacterium* FoF1-ATP synthase beta chain (Table S13) showed 60-90% sequence identity with the human mitochondrial beta chain (**Figure 3**). Likewise, among the five identified epitopes in the alpha chain (Table S13), peptide no. 3 (**EAYPGDVFYLHSR**) was conserved in the human mitochondrial ATP synthase (**Figure S1**), indicating that both alpha and beta subunits may act as autoantigens.

These findings provide compelling evidence that FoF1-ATP synthase, particularly its alpha and beta subunits, represents a potential autoantigenic target in psoriasis. Further studies are warranted to elucidate its precise role in disease pathophysiology and related comorbidities.

### 3.6 *Corynebacterium* Proteins of the Electron Transport Chain are Antigens

The electron transport chain (ETC) comprises a series of enzyme complexes that sequentially transfer electrons derived from the oxidation of NADH and FADH2 (produced in the Krebs cycle), regenerating NAD^+^ and FAD. This process generates an electrochemical gradient across the membrane, ultimately driving ATP synthesis via oxidative phosphorylation. The electrons originate from the catabolism of organic molecules, releasing energy used to power cellular processes.

BLASTp analysis of the nBsAb-bound peptides identified 90 epitopes corresponding to 20 ETC-associated antigens. These include NADP-dependent isocitrate dehydrogenase (IDH), fumarate reductase/succinate dehydrogenase, ubiquinol-cytochrome *c* reductase, cytochrome *bc* complex, cytochrome *c* oxidase subunits I and II, the cytochrome *b* subunit of succinate dehydrogenase, cytochrome *c* oxidase assembly protein, succinate dehydrogenase subunit, NAD-dependent succinate-semialdehyde dehydrogenase, Fe-S cluster assembly proteins SufB and SufD, and NAD(P)H-quinone dehydrogenase (**Table S14**).

Remarkably, 25 epitopic peptides were identified for IDH, originating from at least 84 *Corynebacterium* species (**Table S14**). IDH plays a critical role in energy metabolism, amino acid biosynthesis, and vitamin production. It catalyzes the oxidative dehydrogenation of isocitrate to form oxalosuccinate, followed by oxidative decarboxylation to yield 2-oxoglutarate^135^. As the rate-limiting enzyme of the tricarboxylic acid (TCA) cycle, IDH significantly influences fatty acid synthesis^136^ and thus represents a major antigenic target of nBsAbs in *Corynebacterium*.

Fumarate reductase, another key antigen with 11 identified epitopes, catalyzes the reduction of fumarate to succinate. This reaction is vital not only for the TCA cycle but also for anaerobic respiration, where fumarate serves as the terminal electron acceptor in many facultative anaerobes^137^. Notably, this enzyme plays an essential role in the energy metabolism of *Helicobacter pylori*, being critical for its colonization and survival in the acidic environment of the stomach^137^.

### 3.7 *Corynebacterium* Glycolytic Enzymes are Antigens

In bacteria, glycolysis represents one of several pathways for the catabolic breakdown of glucose. It is most commonly associated with anaerobic or fermentative metabolism, particularly in bacteria and yeasts. As a fundamental and evolutionarily ancient metabolic pathway^138^, glycolysis plays a central role in cellular energy production and biosynthetic precursor generation.

The primary functions of the glycolytic pathway are the provision of intermediates for anabolic processes and the conservation of energy required to support vital cellular activities. BLASTp analysis of nBsAb-bound peptides identified 107 epitopes corresponding to 28 distinct glycolytic enzymes in *Corynebacterium* species (**Table S15**).

Among these, the ATP-dependent 6-phosphofructokinase (Pfk) emerged as a major antigen, with 17 identified epitopes (**Table S15**). Pfk catalyzes the phosphorylation of fructose-6-phosphate to fructose-1,6-bisphosphate using ATP as the phosphoryl donor. This irreversible reaction is a key regulatory step in glycolysis and is conserved across all domains of life, underscoring its metabolic significance.

Additional glycolytic antigens identified include NAD(P)H-dependent glycerol-3-phosphate dehydrogenase, various gluco- and mannomutases, glycerol kinases, diacylglycerol kinases, dihydroxyacetone kinase subunits, and 2,3-bisphosphoglycerate-dependent phosphoglycerate mutase (**Table S15**). These enzymes are involved in energy generation, carbohydrate metabolism, and lipid precursor biosynthesis, highlighting their potential as antigenic targets in *Corynebacterium*.

### 3.8 Reverse Citric Acid Cycle Proteins of *Corynebacterium* are nBsAb Targets

The reverse (or reductive) citric acid cycle is a series of biochemical reactions utilized by certain bacteria and archaea to fix carbon dioxide and water into organic compounds using energy-rich reducing agents as electron donors^139^. This pathway is considered an ancestral carbon fixation mechanism and is characterized by three key enzymes: citrate lyase, fumarate reductase, and α-ketoglutarate synthase.

BLASTp analysis of nBsAb-bound peptides identified several antigens associated with the reductive citric acid cycle in *Corynebacterium* species, including fumarate reductase, a citrate lyase family protein, fumarate hydratase, succinate-semialdehyde dehydrogenase, the cytochrome *b* subunit of succinate dehydrogenase, and the succinate dehydrogenase iron-sulfur subunit (**Table S16**). The highest number of epitopes was found for fumarate hydratase (15 epitopes), followed by fumarate reductase (11 epitopes). Notably, several of these epitope-containing peptides are conserved across up to 26 *Corynebacterium* species (**Table S16**), indicating their potential functional importance and broad immunogenicity.

Succinate dehydrogenase has previously been implicated as a regulator of cellular respiration, particularly under low-oxygen conditions^140^. In *M*. *tuberculosis*, disruption of respiratory control via this enzyme compromises recovery from stationary phase, suggesting its central role in pathogen persistence. One conserved epitope, **DLQPGSSIMPGK** - derived from the succinate dehydrogenase enzyme - is present in both, *Corynebacterium* species and *M. tuberculosis*, highlighting its potential as a broadly cross-reactive target. Additionally, the succinate dehydrogenase iron-sulfur subunit from *M. tuberculosis*, identified as a distinct antigenic target (**Table S16**), represents a promising candidate for peptide-based *M. tuberculosis* vaccine development.

### 3.9 Biosynthetic and Metabolic Pathway Enzymes are *Corynebacterium* Antigens

BLASTp analysis identified 3,367 nBsAb-bound epitopic peptides corresponding to 634 biosynthetic and metabolic pathway enzymes, primarily from *Corynebacterium simulans*, *C. striatum*, and several other *Corynebacterium* species (**Table S17**). Seventeen of these antigens

- catalase, polyphosphate kinase 2 (Ppk2), phosphopyruvate hydratase (enolase), formate C-acetyltransferase, ATP synthase subunit beta, aldehyde dehydrogenase (NAD) family protein, pyruvate kinase, pyruvate carboxylase, pyruvate dehydrogenase (acetyl-transferring), adenylosuccinate lyase, dihydrolipoyl dehydrogenase, glutamine-hydrolyzing GMP synthase, citrate synthase, type I polyketide synthase, serine hydroxymethyltransferase, fumarate reductase/succinate dehydrogenase flavoprotein subunit, and aconitate hydratase - harbored between 11 and 23 epitopic peptides each (**Table S17**). In contrast, the majority of *Corynebacterium* antigens featured only one or two epitopes.

Due to the large number of identified antigens, select candidates of particular interest are discussed in more detail. Many of the multi-epitope antigens are essential enzymes involved in core metabolic pathways such as the Krebs cycle and the purine nucleotide cycle.

Two type I polyketide synthases were identified as antigens: Pks13 (170 kDa; antigen no. 103; 5 epitopes), and a pks13-related antigen (310 kDa; antigens nos. 86 (17 epitopes), 343(1epitope), 368(1 epitope), and 426(2 epitopes); **Table S17**). Type I polyketide synthases are large, long-chain fatty acids synthesizing multidomain enzymes that are essential for synthesizing mycolic acid components of the cell wall, enabling formation of the lipid-rich outer layer that confers structural integrity, low permeability, and stress resistance in *Corynebacteriaceae*^141^, *M*. *tuberculosis*^142^ and *M. leprae*^143^. Pks 13 functions as a condensase, catalyzing the final condensation of two long-chain acyl precursors during the biosynthesis of mycolic acids. Owing to its pivotal role in the *M. tuberculosis* cell wall, Pks13 represents a promising drug target for the development of novel anti-tuberculosis therapies^142^. Most of the identified epitopes are highly conserved across many species within the genus *Corynebacterium* (**Table S17**). Several of them (such as antigen no. 86, epitopes no. 2, 4, 5, 10, 13–15) share more than 80% sequence identity with homologs in *C. diphtheriae*, *M. tuberculosis*, or *M. leprae*, making them attractive candidates for peptide-based vaccines against diphtheria, tuberculosis and leprosy^143^.

Argininosuccinate synthase (antigens no. 158, 291, 311) catalyzes the formation of argininosuccinate from citrulline and aspartate, a key step in arginine biosynthesis. Argininosuccinate lyase (antigen no. 155) then converts argininosuccinate into L-arginine and fumarate^144^.

Peroxiredoxins (antigen no. 179, 10 epitopic peptides) play a vital role in bacterial oxidative stress response by using reactive cysteine thiols to reduce peroxides with electrons from reduced pyridine dinucleotides^145^. Notably, the epitope **DFTFVCPTEIAAFGK** contains the active site **Cys60** (shaded grey, **Table S17**).

Catalases (antigen no. 22) also protect against oxidative damage by decomposing hydrogen peroxide^146^.

Two enzymes involved in peptidoglycan biosynthesis - UDP-N-acetylmuramoyl-L-alanyl-D-glutamate-2,6-diaminopimelate ligase (antigens no. 118, 337, 363) and DAP epimerase (antigen no. 241) - represent promising antimicrobial targets^147^.

Polyphosphate kinase 2 (Ppk2, antigen no. 19) is a conserved enzyme that uses inorganic polyphosphate (polyP) to convert GDP into GTP. In *Pseudomonas aeruginosa*, it appears during stationary phase and may contribute to the synthesis of alginate, a key exopolysaccharide for virulence^148^.

Phosphopyruvate hydratase (enolase,) catalyzes the reversible conversion of 2-phosphoglycerate to phosphoenolpyruvate, a critical glycolytic step^149,150^. Enolase from *C. striatum* (MF_00318) has two active sites at **Glu204** and **Lys335**; both are present in two nBsAb-bound epitopic peptides (antigen no. 24, Table S17, shaded grey).

Formate C-acetyltransferase (antigen no. 25) regulates anaerobic glucose metabolism by catalyzing the reversible conversion of pyruvate and CoA into formate and acetyl-CoA via a unique radical mechanism^151^. The epitope motif **AIACCVSPM** contains catalytic cysteines **Cys415** and **Cys416** (shaded grey)^152^.

Dihydroorotase (antigen no. 53), a zinc metalloenzyme involved in pyrimidine nucleotide biosynthesis, was targeted by nine nBsAb-bound epitopic peptides (**Table S17**).

Orotate phosphoribosyltransferase (OPRT; antigen no. 76, six epitopic peptides) is a key enzyme in the *de novo* pyrimidine biosynthesis pathway, catalyzing the conversion of orotate to orotidine-5′-monophosphate. This reaction is indispensable for bacterial growth and survival, particularly in *M*. *tuberculosis*^153^. Notably, four of the six identified OPRT epitopes are highly conserved across the genus *Corynebacterium* as well as in *M. tuberculosis*. Thus, targeting OPRT with an epitope-based peptide vaccine, rather than conventional enzyme inhibition, represents an alternative therapeutic strategy with potential efficacy against both active and latent tuberculosis^154^. Structurally, OPRTs display a highly conserved architecture characterized by a Rossmann fold^155^ and a dynamic “hood” domain undergoing conformational rearrangements during catalysis^153^. Intriguingly, the epitope **EADYYVDLR** maps to this “hood” domain, suggesting that immune recognition may interfere with catalytically relevant conformational transitions. During the catalytic reaction, OPRTases undergo large conformational changes switching from an open to a closed state. In this transition, the hood domain and the flexible loop show the most prominent structural change.

Type I glyceraldehyde-3-phosphate dehydrogenase (antigen no.189) featured eight epitopic peptides. One peptide (**HNVISAASCTTNCLAPMAK**) includes the active site **Cys151** (shaded grey).

Adenylate kinase (antigen no. 233), a ubiquitous enzyme, was represented by eight epitopic peptides. **LNQDDAANGFLLDGFPR** contains an AMP-binding site, while **AEGTVEEINER** includes an ATP-binding site (both shaded grey).

Fumarate reductase/succinate dehydrogenase flavoprotein subunit (antigen no. 258) was targeted by 11 epitopic peptides. The peptide **YPAFGNLVPR** contains the active site residue **Arg361** (shaded grey).

Citrate synthase (antigen no. 67 with 13 epitopes) is central to energy production via the TCA cycle and links to the electron transport chain^156^. The epitopic peptides **GPLHGGANEAVMK** and **LMGFGHR** contain **His228** and **His317** - two residues from the enzyme’s catalytic triad^157^.

Aconitate hydratase (antigen no. 267), with 23 epitopic peptides, catalyzes the isomerization of citrate to isocitrate in the TCA cycle and acts as an iron regulatory protein^158^. All epitopes were derived from *C. simulans* and *C. striatum* (**Table S17**).

### 3.10 Bacterial Immune Proteins are *Corynebacterium* Antigens

BLASTp analysis identified six *Corynebacterium* CRISPR–Cas system proteins (antigens no. 1-6; 16 epitopes, **Table S18**) and four restriction endonucleases (antigens no. 7-10; 4 epitopes, **Table S18**) as antigenic targets. Components of the Type I-E CRISPR-Cas system (antigens no. 2, 5, and 6; **Table S18**), which is present in several *Corynebacterium* species including *C. diphtheriae*, *C. bovis*, *C. pseudotuberculosis*, and *C. striatum*, form a multi-protein effector complex responsible for nucleic acid interference^159^. CRISPR-Cas systems provide adaptive immunity in bacteria and archaea by targeting invading nucleic acids through sequence-specific CRISPR RNAs and often require a protospacer-adjacent motif (PAM) for DNA recognition^160^.

Restriction endonucleases represent an additional bacterial defense mechanism and cleave foreign DNA; they are classified into Type I and Type II enzymes. Type II restriction endonucleases are widely used in molecular biology because of their ability to cleave DNA at defined recognition sequences^161^.

Notably, two epitopic peptides derived from antigens no. 3 and 7 (**Table S18**) were specific to *C. diphtheriae*, suggesting activation of a bacterial immune response against the toxin-encoding bacteriophage Corynephage β^162^. Similarly, epitopes derived from antigens no. 1 (eight epitopes) and no. 6 (one epitope) were specific to *C. striatum*, indicating phage-related immune activity in this species as well. These findings point to a yet unexplored role of *C. striatum*-specific bacteriophages in the pathophysiology of psoriasis. Consistently, a recent Western blot/peptidome study demonstrated that nBsAb-bound peptides share sequence similarity with multiple bacteriophage proteins as well as diphtheria toxin^32^.

In line with these observations, single-molecule real-time (SMRT) sequencing has revealed the complete genome of *Corynebacterium simulans* together with its associated bacteriophage from a skin metagenomic sample^163^. The phage genome encodes a comprehensive set of structural (head and tail) proteins, integrases, a type VI secretion system, and lytic enzymes, including chitinase-like hydrolases with lysozyme-like properties-features that may be linked to bacterial virulence and contribute to the pathophysiology of psoriasis^163^.

### 3.11 Chaperones and Chaperonins are *Corynebacterium* Antigens and Psoriasis Autoantigens

BLASTp analysis of nBsAb-bound epitopic peptides identified several *Corynebacterium*-derived molecular chaperones (DnaK, GroES, ClpB, DnaJ, PCu(A)C) and chaperonins (GroEL) as antigens (**Table S19**). These proteins are essential for protein folding, especially under stress conditions. In *Escherichia coli*, well-characterized ATP-dependent systems such as DnaK (Hsp70)/DnaJ (Hsp40)/GrpE and GroEL/GroES maintain protein homeostasis by assisting in the folding of nascent and misfolded proteins^164^. Hsp70 proteins act as central hubs in the cellular quality control network and are involved in numerous survival pathways^165^.

In this exploratory study, 21 epitopes were identified on the DnaK antigen (**Table S19**), found across at least 72 *Corynebacterium* species and *M*. *tuberculosis*. However, only *C. simulans* and *C. striatum* harbored the full set of 21 DnaK epitopes, suggesting these species are key immune targets of psoriasis-associated nBsAbs. DnaK, a bona fide Hsp70, contains an N-terminal nucleotide-binding domain (NBD, ∼385 amino acids) linked to a substrate-binding domain (SBD, ∼240 amino acids). Notably, 15 of the 21 identified epitopes were located in the NBD (**Table S19**).

Interestingly, similar to several epitopes of *Corynebacterium* ribosomal proteins (**Table S3**), *Corynebacterium* DnaK epitopes also exhibit 70–90% sequence identity with those of *Akkermansia muciniphila* and *Faecalibacterium prausnitzii* (**Table S19**), two gut commensals known for their beneficial roles in host-microbiota interactions and gut health^92,166^. Given that psoriasis shares immunological features with inflammatory bowel diseases and that the abundance of *A. muciniphila* and *F. prausnitzii* is reduced in psoriasis patients^93,167,168,169^, this unexpected molecular overlap may provide new insights into the pathophysiology of psoriasis.

Hsp70 chaperones operate via ATP-dependent conformational cycling: ATP binding promotes a low-affinity state for substrate capture, while hydrolysis to ADP stabilizes the high-affinity client-bound form^165^. Although intrinsic ATPase activity is low, it can be significantly enhanced by the conserved interdomain linker. For example, adding the linker sequence (*386***VKDVLLLD***393*) to the NBD increases ATPase activity by 40-fold^170^. This sequence appears in epitope peptide no. 5 (**Table S19**), which is conserved in all 73 *Corynebacterium* species, *M. tuberculosis*, *M. leprae*, and human Hsp70 - highlighting its potential as an autoantigen.

Several other epitopic peptides (such as peptides no. 6, 11, 18; **Table S19**) exhibit ≥85% sequence similarity with human Hsp70 (**Figure 4**), classifying DnaK as a multi-epitope autoantigen. Elevated levels of anti-Hsp autoantibodies have been observed in various inflammatory and autoimmune conditions, including rheumatoid arthritis^171^, juvenile idiopathic arthritis^172^, autoimmune myasthenia gravis^173^, dermatitis herpetiformis^174^, psoriasis^175^, systemic lupus erythematosus^176^, epidermolysis bullosa acquisita^177^, celiac disease^178^, and atopic dermatitis^179^. Despite their prevalence, the pathological roles and mechanisms of anti-Hsp antibodies remain poorly understood.

The presence of anti-Hsp antibodies in healthy individuals^180^ has led to the hypothesis that bacterial infection-induced antibodies may cross-react with human Hsps via molecular mimicry^181^. This study supports the broader theory that infections may initiate autoimmunity^182^.

Notably, Hsp70-derived autoepitope sequences identified here may fulfill an unmet need for antigen-specific immunotherapies. The Hsp70-derived peptide **VLRVINEPTAAALAY** (peptides no. 6 and 11, **Table S19**) is currently being investigated in a Phase I/II clinical trial as part of an autologous tolerogenic dendritic cell therapy aimed at inducing antigen-specific tolerance in rheumatoid arthritis^183^.

### 3.12 Stress Proteins are *Corynebacterium* Antigens

BLASTp analysis of nBsAb-bound epitopic peptides identified several stress response proteins from *Corynebacterium* species as antigens. These proteins are crucial for bacterial survival under adverse environmental conditions. Specifically, 11 epitopic peptides were derived from the universal stress protein (Usp) antigen, originating from multiple *Corynebacterium* species. In addition, two epitopes mapped to the 50S ribosomal protein L25/general stress protein Ctc, and three to the Asp23/Gls24 family envelope stress response protein (Table S20).

The universal stress protein A (UspA) superfamily is an ancient and evolutionarily conserved group found across bacteria, archaea, fungi, flies, and plants^184^. In *E. coli*, UspA is produced in response to a wide range of environmental stresses and becomes one of the most abundant proteins in growth-arrested cells^185^. While the regulation of *E. coli* UspA expression has been studied, the exact physiological roles of Usp proteins remain incompletely understood. They are thought to contribute to resistance against DNA-damaging agents and respiratory uncouplers^185^.

Asp23, also known as alkaline shock protein 23, is a membrane-associated protein that contributes to the cell envelope stress response. Its upregulation is associated with maintenance of cell envelope homeostasis^186^. Gls24, a general stress protein in *Enterococcus faecalis*, is linked to both stress adaptation and virulence. Notably, anti-Gls24 rabbit serum conferred protection in a mouse peritonitis model of *E. faecalis* infection, suggesting Gls24-derived peptides may be promising candidates for peptide-based vaccines^187^.

The protein annotated as "50S ribosomal protein L25/general stress protein Ctc" serves dual functions - contributing to the structure of the 50S ribosomal subunit and acting as a general stress protein in certain bacterial species.

These findings highlight that stress-related proteins from *Corynebacterium* not only play key roles in bacterial resilience but also serve as immune targets, expanding the landscape of potential diagnostic markers and therapeutic peptides for immune-mediated diseases.

### 3.13 Metalloproteins are *Corynebacterium* Antigens

A BLASTp search of nBsAb-bound peptides identified 53 metalloproteins as antigens, collectively containing 101 epitopes (**Table S21**). While most antigens harbored only a single epitope, metallohydrolases and metallopeptidases carried between 4 and 10 epitopic peptides. Peptides from M3-, M13-, and M20-family metallohydrolases showed 100% sequence identity with metallohydrolases from both *C. simulans* and *C. striatum* (**Table S21**).

Six antigens (nos. 9, 12, 15, 16, 17, and 20; **Table S21**) were identified as metallohydrolases of the metallo-β-lactamase (MBL) superfamily^188^, a group of proteins present across all domains of life. These proteins feature a characteristic αββα-fold and metal-binding sites - typically coordinating Zn(II), Fe(II)/Fe(III), or Mn(II) - positioned within a cleft that accommodates various substrates. Zinc-β-lactamases inactivate β-lactam antibiotics by hydrolyzing the β-lactam ring^189^.

Five antigens (comprising six epitopic peptides) were associated with molybdenum cofactor (Moco) biosynthesis proteins (**Table S21**). Moco is essential for the function of enzymes such as sulfite oxidase, xanthine dehydrogenase, and aldehyde oxidase^190(p20)^.

Antigen no. 47 (**Table S21**) was identified as part of the cytochrome bc1 complex, harboring three epitopes from at least 15 *Corynebacterium* species. The cytochrome bc1 complex is a ubiquitous energy-transducing electron transfer complex found in the membranes of diverse photosynthetic and respiring bacteria and in the inner mitochondrial membrane of eukaryotes^191^.

Additionally, 13 antigens containing 1–2 epitopes featured iron-sulfur (Fe-S) clusters (**Table S21**), essential for key biochemical processes including respiration, photosynthesis, and nitrogen fixation. The biosynthesis of these cofactors is tightly regulated to prevent cellular toxicity caused by free iron and sulfide^192^.

### 3.14 Surface-Associated Proteins are *Corynebacterium* and *Prescottella equi* Antigens

Twenty-six nBsAb-bound epitopic peptides from 21 antigens originated from *Corynebacterium* species and *Prescottella equi* (**Table S22**), with 11 peptides being species-specific.

Three antigens (3 epitopes) were components of Type VII secretion systems (T7SS), used by bacteria to secrete proteins into their environment^193^. For instance, *Mycobacterium* spp. employ T7SS to translocate proteins across their complex cell envelope^194^.

Two antigens with three epitopes were associated with DoxX family membrane proteins, integral membrane components found in bacteria, archaea, and animals. In *M. tuberculosis*, DoxX is part of the membrane-associated oxidoreductase complex (MRC), linking oxidative stress responses with cytosolic thiol homeostasis^195^.

Four antigens with four epitopes originated from LPXTG cell wall anchor domain-containing proteins, specifically the 1,4-alpha-glucan branching enzyme GlgB. GlgB catalyzes alpha-1,6-glucosidic linkages during glycogen biosynthesis and participates in the metabolism of storage polysaccharides^196^.

Pilin and fimbrial proteins were also detected as antigens (**Table S22**). In Gram-positive bacteria, pili are formed by covalent polymerization of pilin subunits via sortases^197^. These pili, such as the SpaA-type in *C. diphtheriae*, play key roles in host adhesion, with gene clusters encoding SpaA, SpaB, SpaC, and sortase A orchestrating their assembly^198,199^. Notably, sortases themselves were also identified as antigens (antigen no. 17; **Table S22**).

### 3.15 Proteases and Peptidases are *Corynebacterium* and *Tsukamurella* Antigens

A total of 92 nBsAb-bound epitopic peptides from 31 antigens were associated with proteases and peptidases from *Corynebacterium* spp. and *Tsukamurella paurometabola* (**Table S23**).

Among these, the ATP-binding and proteolytic subunits of ATP-dependent caseinolytic proteases (Clp) were prominent, comprising antigens 1 - 5 with 25 epitopes found in >48 *Corynebacterium* species. Clp proteases (ClpP), conserved across bacteria and eukaryotes, are involved in degrading misfolded or short-lived proteins and maintaining proteostasis^200^. Their critical role in virulence has made ClpP attractive antimicrobial targets, leading to the development of acyldepsipeptides (ADEPs), which activate ClpP uncontrollably, disrupting regulated proteolysis and leading to bacterial cell death^201,202^.

Interestingly, the ATP-binding site of *Corynebacterium* ClpP emerged as a major target of the nBsAbs. Four epitopes (peptides 5, 8, 13, and 16) contain the Walker-A motif **GEPGVGK**(**T/S**), a highly conserved ATP-binding signature motif^121,122^ (**Table S23**). In addition, epitope peptide 2 of ClpP antigen no. 4 (**Table S23**) encompasses **His123**, a residue of the catalytically essential **Ser-His-Asp** triad^203^. This suggests that nBsAbs may directly interfere with Clp activity across 27 *Corynebacterium* species. Together, these findings underscore the potential of Clp-derived epitopes as vaccine candidates, particularly against pathogens such as *M*. *tuberculosis*, for which ADEPs are currently being explored as novel therapeutic agents^204^.

Additionally, 59 epitopic peptides were derived from 25 antigens, including trypsin-like serine protease 2, various aminopeptidases, and metallopeptidases (**Table S23**).

### 3.16 Domains of Unknown Function (DUFs) are *Corynebacterium* Antigens

Domains of unknown function (DUFs) represent protein families with no characterized roles, cataloged in the Pfam database^205^. In total, 87 nBsAb-bound epitopic peptides from 60 DUF-containing antigens were identified (**Table S24**). Most DUF-containing antigens had 1-3 epitopes; however, DUF349 (antigen no. 6) exhibited 12 epitopes.

DUF349 typically occurs as a single domain or in tandem repeats of up to five within bacterial proteins, and is predicted to comprise two or three alpha-helices, with possible beta strands^206^. All 12 epitopes of DUF349 matched sequences from *C. simulans* and *C. striatum*, while epitopes 6 and 8 also matched *C. aurimucosum* and *C. faecipullorum*. No additional functional data are currently available for other DUFs in this context.

### 3.17 Bacterial Transporters and Exporters are *Corynebacterium* Antigens

Analysis of nBsAb-bound peptides revealed 66 transporter and exporter proteins as antigens, comprising 92 epitopic peptides (**Table S25**). While most (62) contained only one epitope, three antigens carried between 7 and 12 epitopes. Forty-one peptides originated from ATP-binding cassette (ABC) transporters, six from the major facilitator superfamily (MFS), and three from exporter family proteins.

ABC transporters utilize ATP hydrolysis to drive the uptake and efflux of solutes across membranes in both prokaryotes and eukaryotes, and contribute significantly to bacterial virulence^207^. These systems transport diverse substrates including ions, amino acids, sugars, and peptides.

A Zn/Mn-type ABC transporter solute-binding protein (antigen no. 8) carried seven epitopes, matching sequences from *C. simulans* and *C. striatum*. Likewise, maltose/maltodextrin transport system substrate-binding proteins were prominent antigens, with 18 epitopes derived from up to 75 *Corynebacterium* species. These epitopes matched sequences from *C. simulans*, *C. striatum*, and *C. accolens* - suggesting they are the primary sources of these peptides. These periplasmic binding protein-dependent systems import nutrients across the membrane against concentration gradients^208^.

Among the numerous single-epitope antigens, one epitope was derived from a queuosine precursor transporter (Table S25). Queuosine, a modified nucleoside found in certain tRNAs^209,210(p2)^, plays a role in translational accuracy.

Notably, antigen no. 14’s epitope peptide (Table S25) originates from the daunorubicin resistance ABC transporter ATPase subunit of *M. tuberculosis*. Anthracyclines like doxorubicin are potent inhibitors of Mtb DnaG^211^. Overexpression of ABC transporters contributes to multidrug resistance and treatment failure^211^, making these epitopes - such as **AEGLEKR** - potential candidates for peptide vaccine development against tuberculosis.

### 3.18 Bacteriophage Proteins Are Targets of *C. simulans*-Directed nBsAbs

Peptidomic analysis of *C*. *simulans* WB bands identified 368 DECOY-peptides (**Table S26**). These peptides, although derived from a DECOY database designed to estimate false discovery rates (FDR), may in some cases reflect actual biological sequences. This is particularly true for palindromic peptides or those that, despite being classified as DECOYs, match sequences in the target database. Such overlap can result in inflated FDR estimates due to the mistaken classification of true positives as false positives^212^. Another explanation is that these peptides represent genuine sequences from yet unannotated proteins, possibly from unsequenced viral genomes^213^.

The presence of *C. simulans* phages in the human skin microbiome^163^ supports the hypothesis that some DECOY peptides may originate from phage proteins not yet catalogued in reference databases. In a recent exploratory study, BLASTp analysis limited to viral taxa (taxid:10239) revealed that many DECOY peptides displayed significant similarity to bacteriophage and viral proteins, suggesting that phages infecting *C. simulans* may induce immune responses in psoriatic lesions^32^. When the search was restricted to the class *Caudoviricetes* (taxid:2731619), all 368 peptides showed high similarity - many even complete identity - to proteins from annotated members of this taxonomic group (**Table S26**).

The identified epitopes mapped to a wide variety of phage proteins, including structural and enzymatic components such as integrases, tail and capsid proteins, large terminases, DNA polymerases and helicases, holins, portal proteins, lysozyme-like enzymes, endolysins and diphteria toxin-like proteins, as well as numerous hypothetical proteins. Most peptides originated from unclassified or previously undescribed *Caudoviricetes*, while a smaller number corresponded to known phage taxa, including members of the *Caudovirales* and *Crassvirales* orders, and the families *Herelleviridae* and *Siphoviridae*.

A subset of DECOY peptides matched annotated phage proteins with plausible environmental or dietary origins. For instance, one peptide (peptide no.9, **Table S26**) aligned with a NAD-dependent DNA ligase from *Ralstonia* phage phiRSL1, a virus infecting *Ralstonia solanacearum*, a plant pathogen known to cause bacterial wilt in tomatoes and potatoes^214,215^, suggesting a potential food-derived source. Another peptide (peptide no.125, **Table S26**) matched a helicase from *Gordonia* phage Gmala1; *Gordonia* species are known to inhabit wastewater treatment plants, implying an environmental origin^216^. A terminase peptide (peptide no.165, **Table S26**) showed similarity to a protein from *Aeromonas* phage ZPAH14, pointing toward a potential aquacultural source, as *Aeromonas hydrophila* is a pathogen in fish farming and a known vector of antibiotic resistance genes^217(p2),218^. Additional peptides resembled proteins from phages infecting *Gordonia terrae*, *Moraxella catarrhalis*, and *Bacillus* species, as well as from phages targeting clinically relevant bacteria such as *Klebsiella pneumoniae* and uropathogenic *Escherichia coli*.

The sequence of peptide no. 255 (**Table S26**) shows high similarity to the head scaffolding protein of *Bacillus* phage Thornton. Peptide no. 275 exhibits sequence identity with a capsid maturation protease and a MuF-like fusion protein from *Mycobacterium* phage Zonia, while peptide no. 280 matches a bifunctional NMN adenylyltransferase/Nudix hydrolase from *Klebsiella* phage 150040. *Klebsiella* phages primarily target *Klebsiella* species, particularly *K. pneumoniae*, a major cause of nosocomial infections. Recent research has focused on these phages as promising agents for phage therapy, offering a potential alternative to antibiotics in the treatment of multidrug-resistant infections^219^.

Peptide no. 326 shows high similarity to an endolysin from *Escherichia* phage UE-S5a, a podovirus known to infect uropathogenic *E. coli* (UPEC). UE-S5a and related phages have demonstrated strong lytic activity against UPEC strains, including PSU-5266 (UE-17)^220^. Peptide no. 342 closely matches the hypothetical protein SEA_RUNHAAR_10 from *Gordonia* phage Runhaar, which infects *Gordonia terrae*, an emerging and diagnostically challenging pathogen^221,222^. Peptide no. 360 shows a high degree of sequence overlap with the hypothetical protein SEA_URZA_55 of *Streptomyces* phage Urza; however, no additional information on this phage is currently available.

Three peptides (nos. 366, 367, and 368; **Table S26**) showed high sequence similarity to diphtheria toxin (DT) from *C*. *diphtheriae* phage and *C. ulcerans*. These findings support the hypothesis that, in psoriasis, a recently proposed DT-related “psoriasis toxin” may be generated^72^. Notably, *C. ulcerans* carries the DT-encoding *tox* gene on a prophage distinct from that of *C. diphtheriae*^223^. Moreover, sequencing of the *tox* gene from two *C. ulcerans* isolates derived from skin infections revealed differences compared with *C. diphtheriae* DT sequences, and *C. ulcerans* culture supernatants were markedly less potent than those from *C. diphtheriae*^224^. Together, these observations favor the hypothesis that DT variants are released in psoriatic lesions that are structurally distinct and less toxic than classical DT. Future studies should aim to isolate toxigenic *Corynebacterium* species from lesional psoriatic skin and to further characterize these variants.

The data suggest that many of these nBsAb-targeted peptides originate from previously uncharacterized bacteriophage proteins, some of which may be derived from dietary, environmental, or microbial sources commonly encountered by humans. Collectively, the findings support a model in which *C. simulans*-associated phages, including *C. striatum*-associated phages^159,225,226^ and those of the *Caudoviricetes* class, contribute to the immunogenic peptidome observed in inflammatory skin conditions such as psoriasis.

## 4 GENERAL DISCUSSION

This study, in line with earlier exploratory investigations, identified a set of peptides unexpectedly detected within a 16 kDa Western blot (WB) band through peptidome analysis. Since WB procedures separate peptides and small molecules from proteins, the source of these peptides can only be serum-derived immunoglobulin G (IgG).

I propose that during WB analysis of a boiled *C*. *simulans* extract incubated with psoriasis patient serum, one Fab domain of naturally bispecific IgG antibodies (nBsAbs) recognized a bacterial intrinsically disordered protein (IDP) or IDP-rich (IDPR) 16 kDa antigen, while the other Fab domain simultaneously bound unrelated epitopes^32^. The harsh extraction conditions likely facilitated the release of epitopic peptides bound to one Fab arm of an nBsAb, enabling their identification by peptidome analysis. Although the primary aim was to identify the *C. simulans* antigen via tryptic proteomic mapping, exploratory peptidome analysis of these datasets revealed truncated epitopes generated by trypsin digestion - an artifact not observed in native peptidome analyses^227^.

Although not directly tested here, it is plausible that the detected epitopes were bound by IgG4, the only bispecific IgG-antibody found in human plasma^228^. Unlike other IgG subclasses, IgG4 undergoes Fab-arm exchange - a stochastic, concentration-driven process independent of antigen binding - resulting in bispecific antibodies^229,230,231,232^. IgG4, though typically the least abundant subclass, reflects chronic or repeated antigen exposure^228,233,234,235^. Long regarded as non-pathogenic due to its inability to activate complement and its weak Fc receptor binding^236^, IgG4 is now implicated in autoimmune diseases such as systemic lupus erythematosus^237^ and psoriasis, where anti-gliadin IgG4 autoantibodies have been detected^238^. Whether nBsAb (presumed IgG4)-bound autoantigen epitopes - such as those in HRNR, FLG, and other epidermal IDPs^32^ - are protective or pathogenic in psoriasis remains unclear.

A striking finding was that only boiled bacterial extracts yielded WB-detectable bands, suggesting that the antigens are heat-stable and likely represent IDPs or IDPR-containing proteins. While structural data on the *C. simulans* 16 kDa antigen remain scarce, our results, supported by earlier work^32^, suggest these epitopes derive from IDP(R)s of *C. simulans*, potentially in complexes with other *Corynebacterium* or host proteins. Such complexes dissociate upon boiling^239^ - a rarely used WB condition - possibly explaining why these antigens were overlooked previously. Keratinocytes of the stratum granulosum and stratum corneum are rich in IDPs and IDPR-containing proteins^33,240,241^.

Another provocative observation is that most identified epitopes originate from cytosolic bacterial proteins - challenging the assumption that primary bacterial antigens are secreted or membrane-bound. These cytosolic proteins likely became antigenic following bacterial lysis in the outer epidermal layers. I hypothesize that lysis was mediated by cationic intrinsically disordered antimicrobial peptides (CIDAMPs), derived from precursors such as hornerin (HRNR), filaggrin (FLG), and late cornified envelope proteins (LCEs), which are highly expressed in differentiated epidermal layers^242,243,244^. Unlike conventional AMPs that form pores^245^, CIDAMPs penetrate bacterial membranes in an energy-dependent manner, accumulate intracellularly, and cause protein misfolding or aggregation^246^. HRNR-derived CIDAMPs, for example, selectively bind bacterial ribosomal proteins, disrupting protein synthesis followed by bacterial lysis and mimicking multiple antibiotic mechanisms^246^.

Supporting this, 16 HRNR- and 19 FLG-derived CIDAMPs were previously identified as autoantigens in complexes with *C. simulans*-reactive nBsAbs, along with multiple *C. simulans* ribosomal proteins^32^ (**Table S2**). Together, these findings suggest that B-cell epitopes arise from CIDAMP-ribosomal protein complexes.

Notably, the potent *C. simulans*-targeting CIDAMP **HRNR232–294**^243^ overlaps with the nBsAb-bound autoantigen **HRNR236–263**^32^. This supports a model in which CIDAMP activity and subsequent bacterial lysis involve HRNR-ribosomal protein complexes acting as antigens^246^. These antigen complexes may engage B-cell receptors, thereby activating B-cells to secrete IL-10^247^ and promoting the generation of antigen-specific IL-10-producing regulatory B-cells (B_regs_)^248^. Chronic *C. simulans* infection and sustained antigen exposure in psoriatic skin^5^ could drive IgG4 production in B_regs_^249^ as part of an IL-10–mediated tolerogenic response in psoriasis. A proposed mechanism illustrating CIDAMP-mediated *Corynebacterium* lysis, the subsequent release of IDP-complexes as cytosolic protein antigens, their binding to B-cell receptors, and the generation of antigen-specific bispecific IgG4 antibodies is depicted in **Figure 5**.

**Figure 5.**
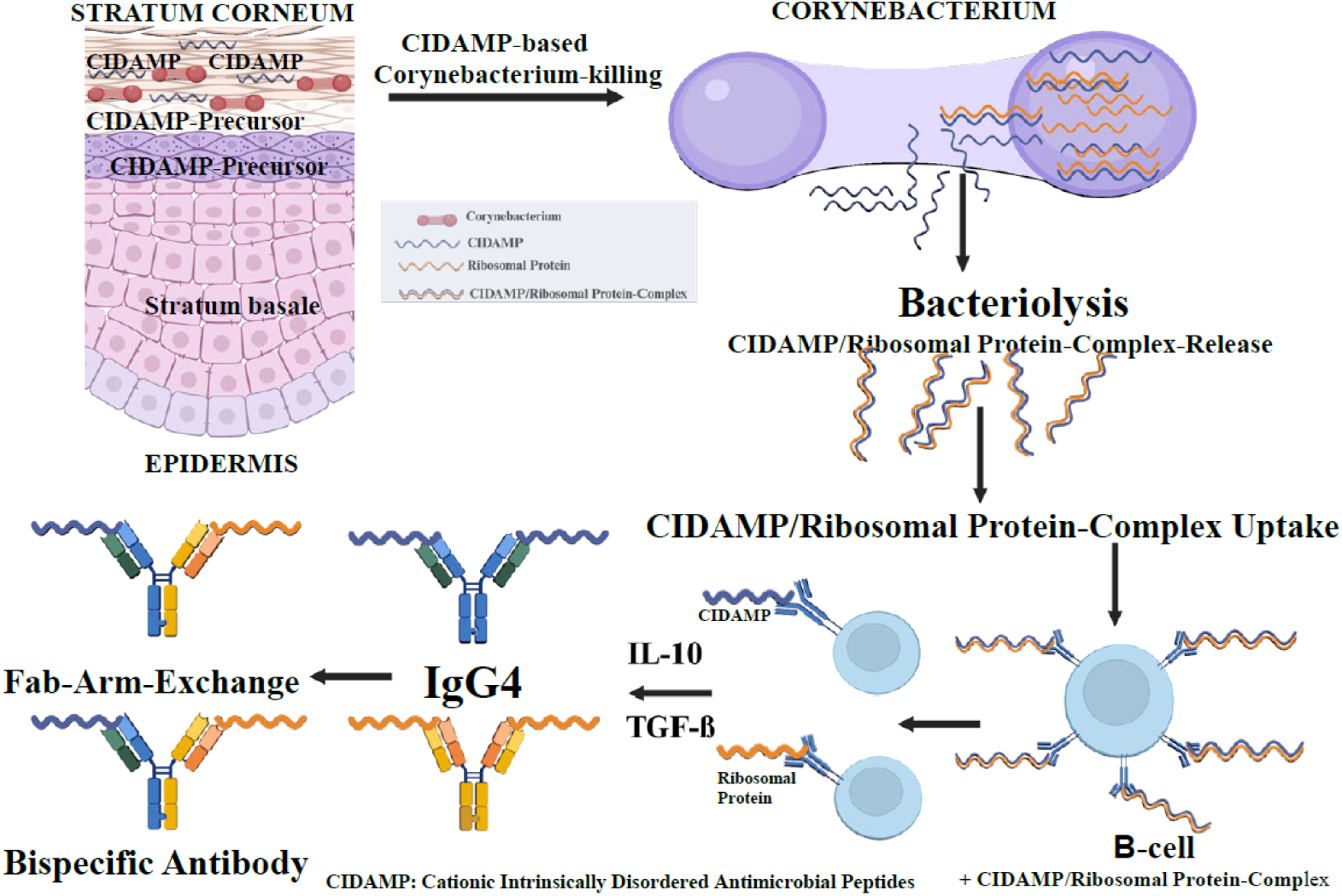
Proposed Mechanism of CIDAMP-mediated Killing of *Corynebacterium* and Subsequent Immune Responses Leading to Bispecific Antibody Formation. Fully differentiated epidermal keratinocytes in the stratum granulosum contain high levels of CIDAMP precursors, such as hornerin (HRNR)²² and profilaggrin (FLG)²³³. Limited proteolysis of HRNR and FLG by skin proteases²³ and bacterial proteases²³ within the stratum corneum generates large numbers of CIDAMPs with varying degrees of bacterial target specificity²². *Corynebacterium*-specific CIDAMPs are taken up by the bacteria, where they bind to ribosomal proteins, forming ribosomal protein/CIDAMP-complexes that may assemble into amyloid-like fibers and inhibit protein synthesis²³. This leads to bacterial lysis and the release of cytosolic proteins as well as CIDAMP/ribosomal protein-complexes²³. These released antigen-CIDAMP-complexes are bound by B-cell receptors and internalized via endocytosis. Persistent exposure to CIDAMP-*Corynebacterium* antigen-complexes promotes the generation of regulatory B cells (B_regs_) producing IL-10²³¹. IL-10 and TGF-β drive IgG4 class-switch recombination, resulting in the production of IgG4 antibodies^249^. Fab-arm exchange between IgG4 molecules subsequently leads to the formation of bispecific antibodies capable of simultaneously recognizing CIDAMPs and *Corynebacterium* ribosomal proteins and other antigens. Created in BioRender. Schröder, J. (2026) https://BioRender.com/v7qj58i.

Since nearly all identified nBsAb-bound epitopes derive from cytosolic antigens - and antibodies act predominantly extracellularly - I further propose that bispecific IgG4 may exert antimicrobial effects within lesional psoriatic skin also intracellularly. Synthetic peptides derived from antibody complementarity-determining regions (CDRs), particularly the IDP-like CDR-H3, have shown antimicrobial peptide-like activity, including membrane disruption, protein binding, enzyme inhibition, and interference with protein-protein interactions^250,251^. Limited proteolysis of nBsAbs in the epidermis, potentially mediated by bacterial^252^ or stratum corneum proteases^253^, could release paratopic peptides capable of penetrating bacterial membranes and acting like CIDAMPs.

An additional unexpected observation was the detection of species-specific epitopic peptides from 40 different *Corynebacterium* species and nine additional Actinomycetota (**Table 1, Table S1**). Notably, the by far most abundant nBsAb-bound epitopes originated from *C. striatum* antigens (**Table S2 - S25**), suggesting that immunity towards this airway epithelial cells invading^42^ skin commensal including its corynebacteriophages^159^ (antigen no. 1, **Table S18**) may also contribute to psoriasis pathology.

The data in **Table 1** and **Table S1** reveal immune responses to multiple *Corynebacterium* and other Actinomycetota species, including commensal, pathogenic, and environmental strains from barrier tissues, soil, food, and plants. Because this exploratory study used pooled serum from patients with psoriasis, the findings reflect collective responses from an undefined number of individuals. Applying the same WB/peptidome approach to single-subject sera could provide individualized molecular-level immune profiles.

Finally, highly conserved epitopes across *Corynebacterium* species - such as those in rpoB and rpoC proteins, shared across 80+ species (**Table S4**) - highlight key immunological principles of cross-reactivity, tolerance, and commensal-immune homeostasis. Since commensal- and pathogen-derived epitopes can be identical, immune outcomes likely depend on prior exposure and antigen context. Chronic recognition of commensal-derived epitopes may establish tolerance, whereas pathogenic encounters - depending on inflammatory cues - could trigger protective or pathogenic responses.

## 5 CONCLUDING REMARKS AND PERSPECTIVES

This exploratory study demonstrates that natural bispecific IgG antibodies - most likely of the IgG4 subtype - detected in pooled serum from psoriasis patients can recognize IDP/IDPR antigens from *C. simulans*, the closely related *C. striatum*, and several other *Corynebacterium* and Actinomycetota species, including toxigenic *C. diphtheriae*, *C. ulcerans* and *C.pseudotuberculosis*. These interactions generate immunogenic IDP-derived epitopes, detectable by Western blotting and peptidome analysis with taxonomic filtering. BLASTp analysis of epitopic peptides revealed a high degree of antigen overlap between *C. simulans* and *C. striatum*, suggesting that these two species are dominant immune targets in psoriasis. The identification of numerous epitopes derived from toxigenic *C. diphtheriae*, *C. ulcerans* and *C. pseudotuberculosis*, and corynebacteriophages highlights the need to further investigate their role in psoriasis pathophysiology.

A subset of identified *Corynebacterium* epitopes exhibits molecular mimicry with human ATP synthase and HSP70, thereby implicating these proteins as autoantigens in psoriasis and reinforcing the link between infection and autoimmunity. Together, these findings support a model in which chronic exposure to *Corynebacterium*-derived antigens may drive either IgG4-mediated immune tolerance or cross-reactive memory, depending on the individual’s immune history and context. This exploratory study further underscores the immunological significance of microbial IDP/IDPR complexes and the potential role of epidermis-derived CIDAMPs in modulating host-microbe interactions at epithelial barriers such as the skin.

The combination of WB/peptidome analyses with taxonomic filtering and BLASTp comparison provides a straightforward strategy for epitope mapping. This approach directly identifies high-affinity B-cell epitopes from serum IgG, in contrast to current state-of-the-art techniques that require labor-intensive mutational scanning, protein display, and high-throughput screening^254,255,256^.

While pooled psoriasis serum was analyzed here, applying this method to serum from individual donors would enable a personalized, epitope-based molecular characterization of the presumed IgG4-mediated immune response to IDP/IDPR antigens. Extending this approach to IgM-, IgA-, IgG1-antigen complexes could provide insights into antigen-specific antibody maturation, including the progressive increase in affinity that underlies effective humoral immunity^257^.

Notably, many epitopes identified in this study are conserved across *Corynebacterium* taxa, including commensals and pathogens. Such shared epitopes may confer cross-protection and should be analyzed in detail to test the “hygiene hypothesis”^258^ in relation to the “epithelial barrier hypothesis”^259^.

The >6000 B-cell epitopes identified here provide a valuable resource for developing epitope-based vaccines targeting pathogenic *Corynebacterium* and related Actinomycetota. B-cell epitope-based peptide vaccines, increasingly considered “vaccines of the future”^260,261,262^ offer precision by including only key epitopes from microbial or viral antigens or tumor (neo)antigens, thereby avoiding potential risks associated with whole-cell vaccines. Their ability to activate targeted immune responses makes them particularly promising for personalized immunotherapy^97,263,264,265,266,267^. A critical bottleneck in therapeutic vaccine and antibody development is B-cell epitope mapping, as most predictive models assume predominantly conformational epitopes^268^. WB/peptidome analyses, as demonstrated here, represent a technically simple method for generating thousands of IDP-derived linear B-cell epitopes thus overcoming this limitation.

The unexpectedly high abundance of cytosolic epitopic peptides from *C. diphtheriae* raises the prospect of developing peptide-based vaccines targeting essential cytosolic proteins rather than solely the diphtheria toxin. This is particularly relevant as non-toxigenic *C. diphtheriae* strains have recently been shown to cause severe invasive disease^269^. Similarly, abundant epitopes from *C. pseudotuberculosis*, an intracellular pathogenic bacterium that causes chronic, debilitating and currently incurable infectious diseases in small ruminants^270^, from *C. ulcerans*, an emerging human pathogen^271^ causing also cutaneous diphtheria^272^ and from the facultative intracellular pathogen *Prescottella (Rhodococcus) equi*, responsible for pyogranulomatous pneumonia in foals and immunocompromised patients^62,273,274^, provide promising candidates for urgently needed vaccines.

Importantly, several epitopes identified here are shared with *M. tuberculosis* and *M. leprae*, the causative agents of tuberculosis and leprosy, respectively. These include RNA polymerases (rpoB, rpoC), aminoacyl-tRNA ligases, elongation factors, ATP synthase subunits, fumarate hydratases, catalase, enolase, orotate phosphoribosyltransferase, molecular chaperones such as DnaK, ClpB, GroEL, and polyketide synthase 13, an essential enzyme in the mycolic acid biosynthesis pathway. Such overlap suggests that WB/peptidome analyses using extracts of *M. tuberculosis* and *M. leprae*, and patient sera may yield novel epitopic peptides with potential for multi-epitope vaccine development, addressing the urgent need for effective prevention of TB^275,276^ and leprosy^277^.

In conclusion, this exploratory study highlights the promise of Western blot and taxonomically filtered peptidome analyses of IDP-antigens - or antigenic regions - from a broad spectrum of infectious agents (bacteria, fungi, viruses, protozoa), as well as tumor antigens, allergens, and other proteins. When combined with patient sera, WB/peptidome analysis of an infectious agent extract with serum may reveal epitopes from both the target antigen and unrelated antigens. Applying taxonomic filters to peptidome datasets, as shown here, enables the identification of a rich repertoire of B-cell epitope sequences of specific immunological relevance. Such insights are pivotal for advancing the development of IDP-derived epitopic peptide vaccines^278^, therapeutic antibodies across diverse diseases, and for deepening our understanding of immune tolerance, autoimmunity, and the complex interplay between host immunity and microbial or viral exposure.

## ABBREVIATIONS

WB: (Western Blot),
NOD1: (Nucleotide-binding Oligomerization Domain 1),
CIDAMPs: (Cationic Intrinsically Disordered Antimicrobial Peptides),
BsAbs: (Bispecific Antibodies),
HRNR: (Hornerin),
DT: (Diphtheria Toxin),
IDP(R)s: (Intrinsically Disordered Proteins or Regions),

## 7 ACKNOWLEDGEMENTS

I have to give credit to Davide Pennino, Ulrich Gerstel, and Britta Hansmann, who helped with pilot experiments and greatly thank Anke Rose and Jutta Quitzau for technical assistance.

D. P. was supported by the MAARS (Microbes in Allergy and Autoimmunity Related to the Skin) project and U. G. and B. H. were supported by the author’s high-risk Reinhart Koselleck-DFG-project “Resistance-avoiding Antimicrobial Peptides of Healthy Human Skin”.

