## Supplemental Figure S1 for "Peptidome Analysis of Western Blots Identifies Natural Bispecific Antibody-Bound *Corynebacterium* and Phage B-cell Epitopes with Potential Relevance to Psoriasis"

```
H. sapiens    41  THLQKTGTAEMSSILEERILGADTSVDLEETGRVLSIGDGIARVHGLRNVQAEEMVEFSS
C. simulans    1  MAELTISSDEIRSAIANYTSSYSAEASREEVGVVISAADGIAQVSGLPSVMANELLEFPG

H. sapiens   101  GLKGMSLNLEPDNVGVVVFGNDKLIKEGDIVKRTGAIVDVPVGEELLGRVVDALGNAIDG
C. simulans   61  GVIGVAQNLDTNSIGVVVLGNFESLTEGDEVKRTGEVLSIPVGEEFLGRVINPLGQPIDG

H. sapiens   161  KGPIGSKTRRRVGLKAPGIIPRISVREPMQTGIKAVDSLVPIGRGQRELIIGDRQTGKTS
C. simulans  121  MGPIAAEEDRVLELQAPSVLQRQPVEEPMQTGIKAIDAMTPIGRGQRQLVIGDRKTGKTA

H. sapiens   221  IAIDTIINQKRFNDGSDEKKKLYCIYVAIGQKRSTVAQLVKRLTDADAMKYTIVVSATAS
C. simulans  181  VCIDTILNQKANWESGDKDKQVRCIYVAIGQKGSTIAGVRHTLEQHGALEYTTIVAAPAS

H. sapiens   281  DAAPLQYLAPYSGCSMGEYFRDNGKHALIIYDDLSKQAVAYRQMSLLLRRPPGREAYPGD
C. simulans  241  DAAGFKWLAPFSGAALGQHWMYQGNHVLVIYDDLTKQAEAYRAISLLLRRPPGREAYPGD

H. sapiens   341  VFYLHSRLLERAAKMNDAFGGGSLTALPVIETQAGDVSAYIPTNVISITDGQIFLETELF
C. simulans  301  VFYLHSRLLERAAKLSDDMGAGSMTALPIIETKANDVSAFIPTNVISITDGQVFLESDLF

H. sapiens   401  YKGIRPAINVGLSVSRVGSAAQTRAMKQVAGTMKLELAQYREVAAFAQFGSDLDAATQQL
C. simulans  361  NQGVRPAINVGVSVSRVGGAAQTKGMKKVAGNLRLELAAYRDLQAFAAFASDLDPASKAQ

H. sapiens   461  LSRGVRLTELLKQGQYSPMAIEEQVAVIYAGVRGYLDKLEPSKITKFENAFLSHVVSQHQ
C. simulans  421  LERGERLVELLKQAESSPQPVEFQMVSIYLADQGIFDVVPVEDVRRFEAEVHEYLHSNTP

H. sapiens   521  ALLGTIRADGKISEQSDAKLKEIVTNFLAGFEA
C. simulans  481  QVFEQIAGGQPLSDESKDALVKAAKDFTPTFRTTEGHNLGSEAEVEALDAVEVKKTELSV

C. simulans  541  SRKTAK
```

***Corynebacterium simulans***: ATP synthase F1, alpha subunit. Access: WP_284841268.1
***Homo sapiens:*** ATP synthase F(1) complex subunit alpha, mitochondrial isoform a precursor. Access: NP_001001937.1

**Figure S1. Sequence Alignment of *H. sapiens* and *C. simulans* ATP Synthase F1 α Subunit Identifies Autoantigenic Epitopes.**
A sequence alignment between the *C. simulans* ATP synthase α-subunit and the human mitochondrial ATP synthase F1 α subunit is shown. Conserved amino acid residues between both sequences are highlighted in yellow. Sequences corresponding to nBsAb-bound *C. simulans* epitopic peptides (see Table S13) are underlined. Note the high degree of sequence conservation within several epitopic regions, representing potential autoantigenic epitopes.
