## Supplemental Table S1 for "Peptidome Analysis of Western Blots Identifies Natural Bispecific Antibody-Bound *Corynebacterium* and Phage B-cell Epitopes with Potential Relevance to Psoriasis"

### Table S1. nBsAbs-bound *Corynebacterium* Species-Specific Epitopes.

| *Corynebacterium* (C.), Other Actinomycetota | Identified Protein | Accession Number | Peptide Sequence | Peptide Start/Stop | Molecular Weight |
|---|---|---|---|---|---|
| C. accolens | Phosphoenolpyruvate hydrolase family protein | WP_237800462.1 | VQSLADAAR | 204-212 | 29281 |
| C. accolens | GDSL-type esterase/lipase family protein | WP_302518661.1 | GVPGATDNYPQREGCLQAPNNWPR | 53-76 | 29931 |
| C. accolens | Trypsin-like serine protease | WP_302503567.1 | EWAEGVMSGK | 332-341 | 40648 |
| C. accolens | DUF1906 domain-containing protein (Tat pathway signal sequence domain protein) | WP_284637643.1 | AGHIGAIR | 56-63 | 32066 |
| C. accolens | ABC transporter substrate-binding protein | WP_302502935.1 | AVAGVLGLK | 88-96 | 31560 |
| C. accolens | Class II fumarate hydratase | WP_284609611.1 | AKEFADVVK<br>DGLVEFSGAMR<br>DLQPGSSIMPGK<br>DSGSLDAEKADAIIAAAK | 168-176<br>266-276<br>307-318<br>61-78 | 49819 |
| C. accolens | Hypothetical protein | WP_284624671.1 | IGAEVSTYR | 174-182 | 23338 |
| C. accolens | DUF6286 domain-containing protein | WP_005276893.1 | AEMNMPKHFGQQPK | 2-15 | 20223 |
| C. accolens | MetQ/NlpA family ABC transporter substrate-binding protein | WP_284639927.1 | SHKSIDDVDK | 125-134 | 30747 |
| C. accolens | Glutamate--tRNA ligase | EFM43688.1 | GSTTMVAMTEVR | 4-15 | 59579 |
| C. accolens | Glutamate ABC transporter substrate-binding protein | WP_284832274.1 | MLVLTSSGMNSIDDADGK | 188-205 | 36746 |
| C. accolens | Hypothetical protein HMPREF0276_1823 | EEI14244.1 | MVFVTEGLEETMTISNR | 1-17 | 7780 |
| C. accolens | Nucleoside hydrolase | WP_284637326.1 | EHGDAVAGCGSLTIMGGAVNYR | 139-160 | 35026 |
| C. accolens | heterodisulfide reductase-related iron-sulfur binding cluster, partial | WP_139027181.1 | MESMTIHPTCSAFQLGMMGDVEK | 2-23 | 11512 |
| C. ammoniagenes | Hypothetical protein CAMM_12040 | APT83592.1 | KLEGQATVK | 35-43 | 8492 |
| C. ammoniagenes | NCS2 family permease | WP_205908312.1 | GPYVNDVLDR | 4-13 | 50741 |
| C. ammoniagenes | Hypothetical protein | WP_168938770.1 | MTQQGAAGTPDPHFIDVQR | 1-19 | 10886 |
| C. ammoniagenes | amidohydrolase | WP_236163494.1 | EFGDSMAEKIGVER | 414-427 | 58668 |
| C. ammoniagenes | Substrate-binding domain-containing protein | WP_003846409.1 | RLEGQDPASVRLVPR | 316-330 | 37255 |
| C. ammoniagenes | Signal recognition particle protein | WP_236163646.1 | GGGMPQMPGMPGMPGGGGGMPSMEELQK | 478-505 | 57885 |
| C. amycolatum | Glycosyltransferase | WP_253291995.1 | NSITLGR | 39-45 | 71677 |
| C. amycolatum | Virulence RhuM family protein | WP_256881576.1 | NFLAEDELK | 241-249 | 39816 |
| C. amycolatum | DUF4247 domain-containing protein | WP_005510135.1 | DPIEVGR | 63-69 | 16096 |
| C. amycolatum | Hypothetical protein | WP_244261990.1 | AGDVVNLK | 252-259 | 37902 |
| C. amycolatu | N-acetylmuramoyl-L-alanine amidase | WP_284839858.1 | INAPSMGR | 7-14 | 52335 |
| C. amycolatum | Hypothetical protein | WP_005511015.1 | MPTGETPTR | 1-9 | 17895 |
| C. amycolatum | ParB/RepB/Spo0J family partition protein | WP_256887192.1 | VIGKSSDVSR | 102-111 | 46090 |
| C. amycolatum | C4-type zinc ribbon domain-containing protein | WP_154843811.1 | LNDAEVDLIAR | 150-160 | 27582 |
| C. argentoratense | Phosphoserine transaminase | WP_234653563.1 | FIPSFLNLQTAVDNSR | 226-241 | 40040 |
| C. argentoratense | Molecular chaperone DnaJ | WP_314929504.1 | FREISLAQEVLSDPNKR | 49-65 | 40313 |
| C. argentoratense | 2-oxoglutarate dehydrogenase, E2 component, dihydrolipoamide succinyltransferase | WP_314929111.1 | AEEPKAEAK | 95-103 | 69519 |
| C. argentoratense | ABC transporter ATP-binding protein | WP_169733206.1 | GLTVLSPR | 378-385 | 53417 |
| C. argentoratense | Ribonuclease J | WP_234870613.1 | GRGGNSNGSSNR | 81-92 | 76327 |
| C. argentoratense | Bifunctional riboflavin kinase/FAD synthetase | WP_315040446.1 | ASGTTETLR | 132-140 | 37040 |
| C. argentoratense | Phosphoglucomutase (alpha-D-glucose-1,6-bisphosphate-dependent) | WP_234858678.1 | MDCSSPDSMASLVANR | 285-300 | 58129 |
| C. argentoratense | Imidazolonepropionase | WP_278762704.1 | MAGEDYAAGGIAVTMNATR | 81-99 | 41308 |
| C. argentoratense | HtaA domain-containing protein | WP_234873405.1 | GQCTGSSSGGGSGSGSGGGSNNDAKEMK | 216-243 | 64219 |

| | | | | | |
|---|---|---|---|---|---|
| C. aurimucosum | GntR family transcriptional regulator | WP_193634551.1 | MLADSVASALR | 1-11 | 22176 |
| C. aurimucosum | Phosphopyruvate hydratase | WP_010187392.1 | LTETLGDKVQIVGDDFFVTNPAR | 296-318 | 45297 |
| C. aurimucosum | Hypothetical protein cauri_0135 | ACP31734.1 | IDLPSFDCGGPALVDLSDK | 5-23 | 3702 |
| C. aurimucosum | Hypothetical protein | WP_012715151.1 | MGIPVQPGVVEVGPEGNIEHK | 169-189 | 29858 |
| C. aurimucosum | Single-stranded DNA-binding protein | WP_010191118.1 | AGGTEYGPTKFIK | 44-56 | 17019 |
| C. aurimucosum | DUF3427 domain-containing protein | WP_046648887.1 | SELEEAPK | | 118011 |
| C. aurimucosum | Acyl-CoA dehydrogenase family protein | WP_010188978.1 | ADLSMAEQR | 76-84 | 76400 |
| C. bovis | Phosphoribosylformylglycinamidine synthase subunit PurL | WP_125187423.1 | LAGALTR | 597-603 | 83909 |
| C. callunae | ATP-dependent helicase | WP_015650695.1 | SFGASITPK | 1478-1486 | 164394 |
| C. callunae | Hypothetical protein | WP_015651613.1 | AEDLISSGRKAR | 422-433 | 66382 |
| C. callunae | Diaminopimelate epimerase | WP_015651503.1 | KTVIPFAK | 2-9 | 29190 |
| C. callunae | RNA polymerase sigma factor | WP_015650134.1 | TLIDALPVER | 126-135 | 21357 |
| C. callunae | Riboflavin synthase | WP_247775370.1 | FSLPENLAR | 122-130 | 21828 |
| C.callunae | Hypothetical protein | WP_015652257.1 | KDDLEFQK | 270-277 | 42098 |
| C. callunae | Hypothetical protein | WP_015651227.1 | NLNNKPLR | 2-9 | 20403 |
| C. callunae | Hypothetical protein | WP_015651958.1 | MSEPGSVGVKQK | 1-12 | 16784 |
| C. casei | MDR family MFS transporter | WP_006822801.1 | GKKTADNVGIVFAALMLTMLMSSLGQMIFGSALPTIV<br>GELGGVDQMSWVISAFMVTMTIAMPLAGQLGDRMG<br>R | 40-112 | 57941 |
| C. casei | Disulfide bond formation protein DsbA | WP_098072481.1 | DITVEFVPMSLAVLNDGR<br>LGDTAFFGPVITR | 31-48<br>156-168 | 22890 |
| C. otitides<br>*C. casei* | 16S rRNA (cytosine(1402)-N(4))-methyltransferase RsmH<br>*iron ABC transporter permease* | WP_046644458.1<br>*WP_006822721.1* | LAHFGGR | 145-151 | 35688 |
| C. casei | Penicillin-binding protein | CCE54652.1 | WPWIVLIVVAVCIAIPGGMFAYAYSQYEVPEPEELAN<br>NQISSIYASDNSTQIAR | 24-77 | 79276 |
| C. casei | DUF6286 domain-containing protein | WP_301436480.1 | DMWVTYSESTR | 133-143 | 31634 |
| C. casei | VWA domain-containing protein | WP_006823125.1 | VNAAEVSNHMDGSDCRADESMTVWR | 340-364 | 73720 |
| C. diphtheriae | Type I restriction enzyme, S subunit | WP_014304115.1 | SGVLELGDGYR | 11-21 | 45368 |
| C. diphtheriae | Pup-protein ligase | WP_047928670.1 | AGDKIVDQLAQR | 99-110 | 53658 |
| C. diphtheriae | Hypothetical protein NY049_11340 | WP_196975733.1 | TLIAYAKPAITYGELGSTPIR | 412-432 | 49733 |
| C. diphtheriae | 2-oxoglutarate dehydrogenase, E1 subunit | WP_199326401.1 | DLFEKQGAPSTPRTEAKNTSSQPAAPAK | 34-61 | 136659 |
| C. diphtheriae | Glutamine--fructose-6-phosphate transaminase (isomerizing) | WP_106361598.1 | VIAICNTNGSSIPRESDACLYTHAGPEIAVASTKAFLA<br>QITATYLLGLYLAQLRGNMFADEVNAVLGELR | 381-450 | 67635 |
| C. diphtheriae | Glutamate-5-semialdehyde dehydrogenase | WP_196979889.1 | CSVCNATETVLIDSALDSAYQLAIITALQEAGVTVHG<br>DVAQLEAVGASGIVPAEEHDWAEEYLSLDIACALVD<br>GVDAAMEHIRTYSTK | 265-352 | 45133 |
| C. diphtheriae | Orotate phosphoribosyltransferase | WP_014319414.1 | ATGAADVIAAEGLEYR<br>DIDAFVVR<br>EADYYVDLR<br>EAGAEVVGVATVVDR<br>ELAVVHGK<br>RATLQHEASR | 155-170<br>89-96<br>32-40<br>140-154<br>17-24<br>41-50 | 19234 |
| C. diphtheriae | DNA-binding protein WhiA; sporulation protein | WP_010934973.1 | LVDLGPMATRR | 73-83 | 35299 |
| C. diphtheriae | 3'-5' exonuclease | WP_014317772.1 | RSDTAGQDK | 10-18 | 30881 |
| C. diphtheriae | Inositol monophosphatase family protein | WP_014308398.1 | MAKTSLK | 1-7 | 31462 |
| C. diphtheriae | DNA primase | WP_014302173.1 | AIVVAGGCSDAIR | 522-534 | 71056 |
| C. diphtheriae | TIGR03773 family transporter-associated surface protein | WP_199337754.1 | DGIADTEGGIAEFK | 510-523 | 117348 |

| | | | | | |
|---|---|---|---|---|---|
| C. diphtheriae | Phospho-sugar mutase | WP_235697053.1 | VYLGGRIAQGAAEGVQLISPADK | 161-183 | 58076 |
| C. diphtheriae | RdgB/HAM1 family non-canonical purine NTP pyrophosphatase | WP_106202317.1 | TFADNALIK | 48-56 | 21601 |
| C. diphtheriae | Hypothetical protein FRC0477_00118 | CAB0935687.1 | LIPQGGPR | 101-108 | 11582 |
| C. diphtheriae | Type II CRISPR RNA-guided endonuclease Cas9 | WP_088265531.1 | EMDGDMR | 510-516 | 121519 |
| C. diphtheriae | DUF294 nucleotidyltransferase-like domain-containing protein | WP_235697141.1 | MLATPCADFMSR | 124-135 | 43258 |
| C. diphtheriae | DUF5129 domain-containing protein | WP_014308841.1 | LENGDAVIRVR | 341-351 | 53682 |
| C. diphtheriae | SpaH/EbpB family LPXTG-anchored major pilin | WP_014301239.1 | TYWGNLNFKK | 347-356 | 56727 |
| C. diphtheriae bv. gravis | PII uridylyl-transferase | KLN44026.1 | SMMPFSATPADIRER | 2-16 | 79233 |
| C. durum | Branched-chain-amino-acid transaminase | WP_315559093.1 | FAGNYAASLVAQAQAAEK | 205-222 | 39931 |
| C. durum | Hypothetical protein | WP_315158668.1 | AIDIAGAR | 46-53 | 11174 |
| C. durum | Salicylate synthase | WP_315159939.1 | TVYQHAGK | 395-402 | 47268 |
| C. durum | Hypothetical protein | WP_315186013.1 | AGDDRDPR | 256-263 | 34800 |
| C. durum | Accessory Sec system translocase SecA2 | WP_040359935.1 | LAKLSDGALAK | 39-49 | 83654 |
| C. durum | Acyl-CoA carboxylase subunit beta | WP_179418910.1 | AEATYPMGR | 31-39 | 59067 |
| C. durum | DUF3068 domain-containing protein | WP_273112225.1 | IGMTLSK | 86-92 | 35831 |
| C. durum | Sigma-70 family RNA polymerase sigma factor | WP_231287024.1 | NTAFADR | 103-109 | 21665 |
| C. durum | Hypothetical protein | WP_315525496.1 | IPIPDTR | 76-82 | 33559 |
| C. durum | Linear gramicidin synthase subunit D | WJY83809.1 | VQVVTTDAGPR | 887-897 | 160212 |
| C. durum | Hypothetical protein | WP_314939114.1 | SFASTRQEVSASGMDFSCAEGNNTEEK | 172-198 | 27174 |
| C. efficiens | Putative vanillate O-demethylase oxygenase subunit B | WP_006769686.1 | RITLRPEHPHK | 28-38 | 41874 |
| C. efficiens | Biotin--[acetyl-CoA-carboxylase] ligase | WP_006769578.1 | QPLDITR | 13-19 | 29717 |
| C. efficiens | Hypothetical protein | WP_006769469.1 | MSAIAIK | 1-7 | 12745 |
| C. efficiens | AAA family ATPase | WP_011076137.1 | ISNEGLWK | 122-129 | 48319 |
| C. efficiens | Cytochrome c oxidase assembly protein | WP_006768357.1 | TVDGVNDLR | 76-84 | 81283 |
| C. efficiens | Glycerate kinase | WP_006767981.1 | SNDVAAQLR | 344-352 | 37099 |
| C. efficiens | Pantoate--beta-alanine ligase | WP_041628383.1 | MSFEHDQGR | 1-9 | 19720 |
| C. efficiens | Putative cyclopropane-fatty-acyl-phospholipid synthase | BAC18404.1 | MPPGGAPKR | 2-10 | 48631 |
| C. falsenii | ATP-dependent DNA helicase | WP_119664081.1 | AIRAAAGESSPR | 150-161 | 120947 |
| C. falsenii | FMN-binding glutamate synthase family protein | WP_025403717.1 | LAPHMLR | 453-459 | 57267 |
| C. falsenii | NUDIX hydrolase | WP_039910949.1 | DTLSGIEVYVQERASTMAFCAEMTVFPGGGVDSR | 17-50 | 33569 |
| C. falsenii | FMN-binding glutamate synthase family protein | WP_025403717.1 | LAPHMLR | 469-475 | 57267 |
| C. falsenii | Hypothetical protein | WP_144084474.1 | VLRLDSSRACGGKGEWAR | 334-351 | 47520 |
| C. falsenii | TPM domain-containing protein | WP_276784545.1 | EGISKSNIK | 70-78 | 70114 |
| C. falsenii | Hypothetical protein | WP_263437054.1 | YESGESLEQIGGR | 5-17 | 4801 |
| C. falsenii | Transaldolase | WP_025402711.1 | GTPAHSAVFDMAADDVR<br>VSLEVDPR | 72-88<br>108-115 | 40193 |
| C. falsenii | YebC/PmpR family DNA-binding transcriptional regulator | WP_025402776.1 | GDLTEDDLLMAVLDAGAEEVNDLGEK | 146-171 | 26824 |
| C. falsenii | 2-C-methyl-D-erythritol 4-phosphate cytidylyltransferase | WP_119665254.1 | LGFSTPK | 20-26 | 30829 |
| C. falsenii | Arabinosyltransferase domain-containing protein | WP_272713711.1 | NVVPLSIPK | 132-140 | 128071 |
| C. falsenii | SbcC/MukB-like Walker B domain-containing protein | WP_025402615.1 | LQLAIDSIK | 358-366 | 125420 |
| C. falsenii | Cation:proton antiporter | WP_224208616.1 | ALVADDSRPR | 241-250 | 43471 |
| C. falsenii | L-serine ammonia-lyase, iron-sulfur-dependent, subunit alpha, partial | WP_272712594.1 | RDHDSEAMPGAK | 203-214 | 48405 |
| C. genitalium | Primosomal protein N' | WP_040423685.1 | KQEVPLR | 639-645 | 72491 |
| C. genitalium | Beta-galactosidase | WP_005287622.1 | LLQATYPK | 217-224 | 56587 |
| Corynebacterium sp. Marseille-P3884; Periplasmic binding protein; | ABC transporter substrate-Binding protein | WP_210575019.1 | IGLYTAEDGR | 205-214 | 33735 |

| | | | | | |
|---|---|---|---|---|---|
| Corynebacterium genitalium ATCC 33030 GN=HMPREF0291_10252 PE=4 SV=1 | | | | | |
| C. genitalium | Hypothetical protein HMPREF0291_11524 | EFK53867.1 | ATVTDAVDK | 122-130 | 17545 |
| C. genitalium | Efflux RND transporter periplasmic adaptor subunit | WP_156774821.1 | ALGEAHAQGR | 181-190 | 47778 |
| C. glucuronolyticum | Nucleoside triphosphate pyrophosphatase | WP_201839608.1 | ERVAALAQAK | 49-58 | 22146 |
| C. glucuronolyticum | tRNA dihydrouridine synthase DusB | WP_276542633.1 | GVSGTDIPVTVK | 141-152 | 43656 |
| C. glucuronolyticum | Porin PorA family protein | WP_201839729.1 | ATDLDGLLK | 116-124 | 35101 |
| C. glucuronolyticum | hypothetical protein HMPREF0293_0753 | EEI63688.1 | TVLLSGLDEK | 50-59 | 7025 |
| C. glucuronolyticum | Anchored repeat ABC transporter, substrate-binding protein | WP_259815086.1 | EAIAGADAR | 343-351 | 50925 |
| C. glucuronolyticum | AAA family ATPase | WP_005394624.1 | IEAQIELWR | 6-14 | 48195 |
| C. glucuronolyticum | Regulatory protein RecX | WP_201841492.1 | GSGLVDDALFAQEWVRVR | 85-102 | 22751 |
| C. glucuronolyticum | Aminotransferase class I/II-fold pyridoxal phosphate-dependent enzyme | WP_232621892.1 | EIAAVDGISIVTPACNAIVTFTTGSEK | 377-403 | 49538 |
| C. glucuronolyticum | Dihydroxyacetone kinase subunit DhaK | WP_005393004.1 | SLDDVAAIAKK | 181-191 | 32023 |
| C. glucuronolyticum | 2-Isopropylmalate synthase | WP_259815016.1 | DLGRTYEAVIR | 396-406 | 63469 |
| C. glucuronolyticum | 2-Isopropylmalate synthase | WP_259815016.1 | KDKEGVK | 172-178 | 60839 |
| C. glucuronolyticum | NAD(P)-binding domain-containing protein | WP_005395145.1 | AALNSSPTTDR | 2-12 | 37999 |
| C. glucuronolyticum | Hypothetical protein HMPREF0293_0294 | EEI64207.1 | MGAVAVTLNER | 1-11 | 6401 |
| C. glutamicum | Metalloregulator ArsR/SmtB family transcription factor | WP_038583602.1 | ALSIGTQWAHVMGIK | 126-140 | 25246 |
| C. glutamicum | Ydcf family protein | WP_256856422.1 | NAESFCMDPYPVITSGEAR | 98-116 | 21115 |
| C. glutamicum | Multidrug efflux MFS transporter Cmr | WP_077313408.1 | MSTFHKVLINTMISNVTTGFLFFAVVFWMYLSTGDV ALTGIVSGIYMGLIAVCSIFFGTVVDHNRK | 1-66 | 49463 |
| C. glutamicum | DUF456 domain-containing protein | WP_011014634.1 | LMVGLFIGLCILIVGGPALGLMLGGFIGVLIGLCLGTS AAMIVGPFGLMRMKAYPPCMDSPWVYMSQEQWR | 96-166 | 22671 |
| C. glutamicum | Helicase-associated domain-containing protein | WP_074508311.1 | GQSEEQVR | 147-154 | 78472 |
| C. glutamicum | TetR/AcrR family transcriptional regulator [Corynebacterium glutamicum] | WP_003855609.1 | AGLSSTVLQR | 49-58 | 10184 |
| C. glutamicum | Succinate CoA transferase | WP_143854664.1 | MAVASATSFALSPEYAEK | 307-324 | 55124 |
| C. glutamicum | Hypothetical protein | WP_011013460.1 | TVEAQYR | 94-100 | 16428 |
| C. glutamicum | DNA-binding protein WhiA | WP_074508156.1 | MGAQKTR | 199-205 | 35618 |
| C. glutamicum | alpha/beta hydrolase-fold protein | WP_216312922.1 | TRILMAALTVVALINTAAVINVIYQPYPTLGSFNPVPT AVSMSYADFESQTTTPTMDNR | 96-154 | 45707 |
| C. glutamicum | HD domain-containing protein | WP_011897161.1 | LGFNELNK | 167-174 | 21373 |
| C. glutamicum | NTD biosynthesis operon regulator NtdR | BAV24098.1 | APLMAKPCELPTR | 127-139 | 13979 |
| C. glutamicum | ABC transporter ATP-binding protein | WP_087062649.1 | LTAQDLLLK | 166-174 | 26015 |
| C. glutamicum | Metalloregulator ArsR/SmtB family transcription factor | WP_038583602.1 | ALSIGTQWAHVMGIK | 126-140 | 25246 |
| C. glutamicum | ABC-type transporter, ATPase component and permease component | CCH24135.1 | MGGLVDK | 1-7 | 56266 |
| C. glutamicum | Hypothetical protein | WP_044027526.1 | LMSFQEWR | 139-146 | 12558 |
| C. glutamicum | Substrate-binding domain-containing protein | WP_012917662.1 | NELNSHMGIGGR | 34-45 | 38929 |
| C. glutamicum | Alpha/beta hydrolase | WP_003857213.1 | DCYLTKDTMER | 193-203 | 31600 |
| C. glutamicum | Hypothetical protein | BAB98399.1 | SCHEEDGTHQSASAR | 184-198 | 24907 |
| C. glutamicum | signal recognition particle protein | WP_011014851.1 | MPMGGMPGMPGMPGMGGAGMPDLAELQK | 481-508 | 58337 |
| C. glutamicum | Hypothetical protein | WP_211439437.1 | MQAGDVSNDSSLDSEASAVRSALGR | 120-144 | 35180 |
| C. glutamicum | Hypothetical protein | WP_020309549.1 | MEPGGIR | 1-7 | 11312 |
| C. glutamicum | Class I SAM-dependent DNA methyltransferase | WP_216312896.1 | EWKTTEDATQQSGPLTR | 39-55 | 171706 |
| C. glyciniphilum | Histidine phosphatase family protein | WP_304032479.1 | QTAAGIALGGSGR | 75-87 | 32468 |

| | | | | | |
|---|---|---|---|---|---|
| C. glyciniphilum | MerR family transcriptional regulator | WP_038547313.1 | EQVRIEK | 90-96 | 13759 |
| C. glyciniphilum | Hypothetical protein | WP_038545399.1 | MKLPNMLNARVTR | 1-13 | 7582 |
| C. glyciniphilum | ATP-dependent DNA helicase | WP_052539681.1 | AVAQEDQAAR | 160-169 | 120455 |
| C. glyciniphilum | TetR/AcrR family transcriptional regulator | WP_038547077.1 | QDAVSSLAR | 86-94 | 39144 |
| C. glyciniphilum | Catechol 1,2-dioxygenase | WP_304031649.1 | ANAIYKDLLGAIGDVAR | 35-51 | 32023 |
| C. glyciniphilum | DEAD/DEAH box helicase | WP_304032435.1 | TFTFGLPVITR | 69-79 | 43126 |
| C. glyciniphilum | Endonuclease/exonuclease/phosphatase family protein | WP_052539768.1 | SSGATYSR | 174-181 | 26621 |
| C. glyciniphilum | Hypothetical protein | WP_041628665.1 | ATEMTKPMR | 2-10 | 23826 |
| C. glyciniphilum | Hypothetical protein | WP_038544983.1 | AEEAKEKAAEVK | 477-488 | 56993 |
| C. glyciniphilum | Mycothiol synthase | WP_144313691.1 | MSVDCTDPDRR | 144-154 | 35877 |
| C. glyciniphilum | Transcriptional regulator, TetR-family | AHW62610.1 | MEDSTGMNHPRR | 1-12 | 25680 |
| C. glyciniphilum | GMC family oxidoreductase N-terminal domain-containing protein | WP_052540052.1 | DSTTSAEHTSTMIGEPL | 503-519 | 55720 |
| C. glyciniphilum | Glutamyl-tRNA reductase | WP_081803755.1 | DIDDAVVDVASGTGAASGADVR | 299-320 | 51674 |
| C. glyciniphilum | DUF6615 family protein | WP_052541097.1 | DWYRCSVDHQSGHCPGCEKR | 133-152 | 32704 |
| C. glyciniphilum | RNA polymerase sigma factor | WP_038548374.1 | MAAHTSSTSGNDAEGIDAPGTSASSAGR | 1-28 | 60299 |
| C. glyciniphilum | TetR-like C-terminal domain-containing protein | WP_038545210.1 | MAASGSTSTTQR | 1-12 | 21277 |
| C. halotolerans | Sensor histidine kinase | WP_015402006.1 | DRTDIVGLSER | 312-322 | 58695 |
| C. halotolerans | Hypothetical protein | WP_015399991.1 | LAGYPGVPHATR | 100-111 | 22257 |
| C. halotolerans | FAD-dependent monooxygenase | WP_015401872.1 | VAATLGLELK | 526-531 | 68304 |
| C. halotolerans | ABC transporter ATP-binding protein | WP_034990807.1 | ALNEFNLSVEK | 16-26 | 32266 |
| C. halotolerans | Acetoacetate--CoA ligase | WP_027004178.1 | DNPAIMQVEEDGRR | 87-100 | 71816 |
| C. halotolerans | Cation-translocating P-type ATPase | WP_015401738.1 | LPHALGLAR | 535-543 | 64781 |
| C. halotolerans | Ferritin | WP_034991252.1 | LVGDDGSGLLR | 144-154 | 19301 |
| C. halotolerans | Peptidylprolyl isomerase | WP_015399458.1 | GPFYDGAIFHR | 57-67 | 18821 |
| C. halotolerans | TetR/AcrR family transcriptional regulator | WP_015400291.1 | SLAAEIVGEDDPR | 144-156 | 24277 |
| C. halotolerans | Mannose-6-phosphate isomerase, class I | WP_015400169.1 | MDQLTGMLR | 1-9 | 41736 |
| C. halotolerans | RNA 2',3'-cyclic phosphodiesterase | WP_027004290.1 | YETVSEIR | 170-177 | 20297 |
| C. halotolerans | TetR/AcrR family transcriptional regulator | WP_015400417.1 | AEPAEALAVR | 122-131 | 21615 |
| C. halotolerans | Maleylpyruvate isomerase N-terminal domain-containing protein | WP_015399730.1 | EQSQRAVEPR | 120-129 | 30118 |
| C. halotolerans | PfkB family carbohydrate kinase | WP_015399904.1 | VVLDGAVTDPEK | 160-171 | 30591 |
| C. halotolerans | GTPase ObgE | WP_015401604.1 | GLIDEYDYGDDRPADR | 484-499 | 53679 |
| C. halotolerans | Putative Apolipoprotein N-acyltransferase | AGF73868.1 | LFRGIAGDNWRRLAGPTALMMGAAVVWWLALPPR | 6-39 | 54827 |
| C. jeikeium | Hypotheical protein | WP_111711727.1 | EDRFPLRPLFAVWDSALPSSALSALHSISMSAISDFGIP SVSDIEAISACERSVNAYEQDEPWASSVK | 79-146 | 24174 |
| C. jeikeium | Pyruvate dehydrogenase [ubiquinone] | WCZ53282.1 | TVTLFCGEGVK | 203-213 | 63088 |
| C jeikeium | Multidrug ABC transporter ATP-binding protein | OOD32146.1 | MFEAIQSVDEGTAR | 94-107 | 127712 |
| C. jeikeium | FBP domain-containing protein | WP_011272809.1 | TPDGTTQMAETLTVEEK | 143-159 | 18156 |
| C. jeikeium | IMP dehydrogenase | WP_041626416.1 | SEAGMITDPVTASPDMTIQEVDDLCAR | 103-129 | 54043 |
| C. jeikeium | Restriction endonuclease subunit S | WP_011273488.1 | LKVAGPGTVLFAMYGATLGAVSR | 264-286 | 44305 |
| C. jeikeium | Copper chaperone PCu(A)C | WP_172457282.1 | MSLTSLNNSK | 1-10 | 23325 |
| C. jeikeium | dUTP diphosphatase | WP_035011199.1 | RNGNDAPLR | 5-13 | 16458 |
| C. jeikeium | Hypothetical protein | WP_005294427.1 | GASQKIGDEVRR | 288-299 | 32370 |
| C. jeikeium | SDR family oxidoreductase | WP_011272816.1 | LEQVAQAITQEGGK | 63-76 | 28641 |
| C. jeikeium | Translational GTPase TypA | WP_011273863.1 | MESMDNTGTGWVR | 454-466 | 69160 |
| C. kefirresidentii | Alpha/beta hydrolase | WP_239207579.1 | SRRSGQSWHYVSDLAFYFADLTAALDAIPNDEVIFIA HSTGGLIAPLWMDHLRR | 99-152 | 38146 |

| | | | | | |
|---|---|---|---|---|---|
| C. kefirresidentii | Acyl-CoA thioesterase II | WP_239204103.1 | NGDCGPQHQDPMPEVPGPEETAR | 107-129 | 32261 |
| C. kroppenstedtii | SDR family oxidoreductase | WP_012730858.1 | AAHPYLSR | 121-128 | 28738 |
| Corynebacterium kroppenstedtii DSM 44385 | Mannose specific PTS system component | ACR16958.1 | CIASESKSNRK | 2-12 | 77935 |
| C. kroppenstedtii | Hypothetical protein | WP_012730760.1 | APLFGDDAQCQRHR | 38-51 | 48884 |
| C. kroppenstedtii | Hypothetical protein | WP_012730918.1 | SSASSSSASSK | 89-99 | 22249 |
| C. kroppenstedtii | Thiamine phosphate synthase | WP_012732318.1 | VAAAGAGVVQVR | 28-39 | 22923 |
| C. kroppenstedtii | Putative membrane protein | ACR17060.1 | MNACRVNAMSRAR | 1-13 | 22703 |
| C. kroppenstedtii | LLM class flavin-dependent oxidoreductase | WP_012731253.1 | ATDGNGYTAR | 16-25 | 37580 |
| C. lipophiloflavum | ATP-binding cassette domain-containing protein | WP_006841224.1 | VLGATHQPSR | 213-222 | 56754 |
| C. lipophiloflavum | Sulfite exporter TauE/SafE family protein | WP_304041736.1 | KTAAADGAPASATR | 124-137 | 32006 |
| C. lipophiloflavum | 30S ribosomal protein S8 | WP_006839458.1 | EASIAGLR | 71-78 | 14359 |
| C. lipophiloflavum | Depupylase/deamidase Dop | WP_006840327.1 | VVASSDVDK | 324-332 | 450410 |
| C. lipophiloflavum | DEAD/DEAH box helicase | WP_006840129.1 | EAGNVFADGGR | 1143-1153 | 183520 |
| C. maris | Glyceraldehyde-3-phosphate dehydrogenase | WP_052337728.1 | DREALGQHLQSK<br>EALGQHLQSK<br>LTGNAIR | 237-248<br>239-248<br>364-370 | 53618 |
| C. maris | Glycosyltransferase 87 family protein | WP_020934607.1 | TVPTSAGSVAVER | 416-428 | 10433 |
| C. maris | Hytoene/squalene synthase family protein | WP_020934521.1 | AVLAAPGK | 84-91 | 32455 |
| C. maris | Malate synthase G | WP_020934049.1 | YALNAANAR | 129-137 | 79730 |
| C. maris | DoxX family protein | WP_020934673.1 | AQDRADDALSK | 266-276 | 33685 |
| C. maris | C4-type zinc ribbon domain-containing protein | WP_020935231.1 | DVHAAHVDENLR | 130-141 | 26799 |
| C. maris | PspA/IM30 family protein | WP_020936510.1 | MGEIAAAGTDMKASAK | 216-231 | 30728 |
| C. maris | DivIVA domain-containing protein | WP_020935155.1 | AEAEAKSAK | 100-108 | 38972 |
| C. maris | DUF3499 domain-containing protein | WP_156844687.1 | VTTGLVDEK | 69-77 | 13562 |
| C. maris | Aromatic acid exporter family protein | WP_020933460.1 | MSSPAPMQALAR | 1-12 | 41713 |
| C. maris | Hypothetical protein B841_12355 | AGS35943.1 | RSSPAMEEPLAR | 23-34 | 56897 |
| C. maris | DNA methyltransferase | WP_020935841.1 | VLKNERER | 927-934 | 106292 |
| C. maris | Chromosome segregation protein SMC | WP_020935076.1 | LAGELAAAVEK | 824-834 | 125546 |
| C. maris | Type I pantothenate kinase | WP_020934321.1 | KGFPEAYDR | 143-151 | 35160 |
| C. maris | ABC transporter--like protein | AGS33501.1 | MVRQGK | 1-6 | 26144 |
| C. matruchotii | Putative dihydrodipicolinate reductase domain protein | WP_311166904.1 | WTAISPAANFLPDAPAR | 140-156 | 21938 |
| C. matruchotii | Proline dehydrogenase family protein | WP_314822490.1 | SLKVGKGTEIDTTMNGIIEAPGEK | 783-806 | 125436 |
| C. matruchotii | Drug resistance MFS transporter, drug:H+ antiporter-2 family | WP_315118707.1 | YKWMPIASMVVLAISLWLLSTLTVETPLWVLLSYMF<br>LLGAGIGLGMQILVLVVQNSFSDHEVGMATASNNFF<br>R | 350-422 | 56305 |
| C. matruchotii | Hypothetical protein | WP_005520586.1 | DLDERQLANVR | 69-79 | 18895 |
| C. matruchotii | DUF1003 domain-containing protein | WP_314974200.1 | IEAKIADEAAAR | 151-162 | 21559 |
| C. matruchotii | Aminotransferase class I/II-fold pyridoxal phosphate-dependent enzyme | WP_005524031.1 | ELAAMDTAAPDFR<br>WLCPVPGYDR | 216-228<br>147-156 | 46146 |
| C. matruchotii | DUF6474 family protein | WP_005522019.1 | KGRLNSGK | 78-85 | 23478 |
| C. matruchotii | NAD(P)H-quinone oxidoreductase | WP_005524062.1 | IVTDTIK | 265-272 | 33643 |
| C. matruchotii | DEAD/DEAH box helicase | WP_311324260.1 | HAVDAIR | 464-470 | 100912 |
| C. matruchotii | DNA primase | WP_005525931.1 | NIASSRQCVVVEGYTDVMAMHAAGVATAVATCGTA<br>FGDDHLQMIR | 257-301 | 70564 |
| C. matruchotii | Type II CRISPR RNA-guided endonuclease Cas9 | WP_315120567.1 | CAYCGAEISFK | 557-567 | 123355 |
| C. matruchotii | Hypothetical protein | WP_311323893.1 | VQKCSAGVMTKR | 186-197 | 59901 |

| | | | | | |
|---|---|---|---|---|---|
| C. matruchotii | DUF2550 domain-containing protein | WP_278718476.1 | LDKMSPSEAR | 132-141 | 17802 |
| C. matruchotii | Hypothetical protein CORMATOL_01952 | EEG26516.1 | LPLTTNGGHR | 12-21 | 5135 |
| C. matruchotii | Nitrate reductase subunit alpha | WP_314973696.1 | STADTLIEAMK | 649-659 | 139355 |
| C. matruchotii | Hypothetical protein | WP_311323466.1 | EAVVADYPGVTR | 182-193 | 25557 |
| C. matruchotii | Hypothetical protein HMPREF0299_6387 | EFM48950.1 | FRMKITR | 2-8 | 14503 |
| C. matruchotii | Hypothetical protein CORMATOL_01952 | EEG26516.1 | LPLTTNGGHR | 12-21 | 5135 |
| C. matruchotii | Phosphoserine transaminase | WP_314822380.1 | EEVSCFVSDPNK | 293-304 | 40106 |
| C. matruchotii | Trigger factor | WP_126299819.1 | SYNMSPEEFMSQVTK | 375-389 | 49615 |
| C. otitides<br>*C. casei* | 16S rRNA (cytosine(1402)-N(4))-methyltransferase RsmH<br>*iron ABC transporter permease* | WP_046644458.1<br>*WP_006822721.1* | LAHFGGR | 145-151 | 35688 |
| C. pseudodiphthereticum | N-acetylmuramoyl-L-alanine amidase | WP_284573452.1 | RISTSGSLR | 6-14 | 74075 |
| C. pseudodiphtheriticum | Aldose epimerase | WP_284596435.1 | NADGHGAILWADEHFSWCQVYTSPESAPSIGRAVAV EPMTCPPNALRSGESLLQLASASSMR | 254-315 | 34917 |
| C. pseudodiphtheriticum | Malate synthase G | WP_284814861.1 | VAFINTGFLDR | 453-463 | 82269 |
| C. pseudodiphtheriticum | Alanine dehydrogenase | WP_284585175.1 | AVAAGGVDQAIER | 322-334 | 37968 |
| C. pseudodiphtheriticum | Phosphatidylinositol mannoside acyltransferase | WP_272696498.1 | LPLWLTRR | 32-39 | 46435 |
| C. pseudodiphtheriticum | Type I-E CRISPR-associated protein Cas5/CasD | WP_284586519.1 | SGIIGLLAAAEGRR | 34-47 | 26796 |
| C. pseudodiphthereticum | IclR family transcriptional regulator | WP_272730776.1 | AIAIMKATATTPMNLSELCTDTGIPR | 16-41 | 24801 |
| C. pseudodiphtheriticum | Sodium:alanine symporter family protein | WP_249617946.1 | HGFIDR | 56-61 | 50843 |
| C. pseudodiphtheriticum | Hypothetical protein | WP_284859974.1 | QVIVDQPR | 44-51 | 38821 |
| C. pseudodiphtheriticum | PDZ domain-containing protein | RUP96078.1 | NSMNKHGSSGR | 2-12 | 47733 |
| C. pseudodiphtheriticum | Aldo/keto reductase | WP_284586931.1 | GLGCTGMTPHSYSDSPR | 37-53 | 35213 |
| C. pseudodiphthereticum | Glycosyltransferase family 2 protein | WP_284606252.1 | QGEDMETEREAGWLSGSCLLVR | 178-199 | 32512 |
| C. pseudodiphtheriticum | DNA-directed RNA polymerase subunit beta | WP_272696996.1 | EMQSPEGQHLMLDTDDIDHFGNR | 342-364 | 128433 |
| C. pseudotuberculosis | Permease, uncharacterized plasma membrane protein | WP_013241187.1 | DWFLVLGDKPVLACLAMAVLAVLLALCSEADAFIAA SFTMMPPTAQLVFLVVGPMVDVKLMAMQR | 447 | 33767 |
| C. pseudotuberculosis | Multicopper oxidase domain-containing protein | WP_145951886.1 | GLAGMILLEDEVSK | 179-192 | 60173 |
| C. pseudotuberculosis | 16S rRNA (cytosine(1402)-N(4))-methyltransferase RsmH | WP_014800655.1 | MAELIAAPVIK | 27-37 | 38580 |
| C. pseudotuberculosis | ComF family protein | WP_014522513.1 | MWEFLFPQACMGCGKPGLR | 2-20 | 21937 |
| C. pseudotuberculosis | Amino acid ABC transporter permease | WP_213173218.1 | ESATFLADNNFR | 87-98 | 24475 |
| C. pseudotuberculosis | Oxoglutarate dehydrogenase inhibitor Odhl | WP_013241856.1 | EPKNSEVLSSGDEIQIGK | 115-132 | 15397 |
| C. pseudotuberculosis | Multicopper oxidase domain-containing protein | WP_145951886.1 | GLAGMILLEDEVSK | 179-192 | 60173 |
| C. pseudotuberculosis | 16S rRNA (cytosine(1402)-N(4))-methyltransferase RsmH | WP_014800655.1 | MAELIAAPVIK | 27-37 | 38580 |
| C. pseudotuberculosis | ComF family protein | WP_014522513.1 | MWEFLFPQACMGCGKPGLR | 2-20 | 21937 |
| C. pseudotuberculosis | Amino acid ABC transporter permease | WP_213173218.1 | ESATFLADNNFR | 87-98 | 24475 |
| C. pseudotuberculosis | Oxoglutarate dehydrogenase inhibitor Odhl | WP_013241856.1 | EPKNSEVLSSGDEIQIGK | 115-132 | 15397 |
| C. pseudotuberculosis | Carboxylesterase/lipase family protein | WP_014300614.1 | LGPLELER | 529-536 | 59482 |
| C. pseudotuberculosis | N-acetyl-gamma-glutamyl-phosphate reductase | WP_014366976.1 | YVSGELK | 28-34 | 40384 |
| C. pseudotuberculosis | D-alanyl-D-alanine carboxypeptidase/D-alanyl-D-alanine-endopeptidase | WP_058832193.1 | ATLDAMAHDR | 56-65 | 40521 |
| C. pseudotuberculosis | RNA pseudouridine synthase B | AEK92306.1 | GSEGSRAGENTRESYGFR | 65-82 | 41826 |
| C. pseudotuberculosis | Hypothetical protein | WP_013240914.1 | MEHREGMDGK | 71-80 | 10474 |
| C. pyruviciproducens | GNAT family N-acetyltransferase | WP_016458261.1 | LVGCARVYK | 60-68 | 16521 |
| C: pyruviciproducens | Prolipoprotein diacylglyceryl transferase | WP_276921981.1 | RFNMGR | 215-220 | 32097 |
| C. pyruviciproducens | Amidohydrolase | WP_284840332.1 | NGEGPVIGMR | 112-121 | 47747 |
| C. pyruviciproducens | FeoC-like transcriptional regulator | WP_016458897.1 | SCAGNCSSCAISSRCGRGR | 52-70 | 8443 |

| | | | | | |
|---|---|---|---|---|---|
| C. resistens | Aminotransferase class V-fold PLP-dependent enzyme | WP_273352814.1 | AFDAVDR | 154-160 | 45366 |
| C. resistens | Hypothetical protein | WP_273351864.1 | DALAAAVGVAGR | 60-71 | 19313 |
| C. resistens | Universal stress protein | WP_013887886.1 | QYPEVRVEEVVERDRPINALRDAAADAQLLITGSHGR | 233-269 | 32799 |
| C. resistens | Hsp70 family protein | WP_273352345.1 | EEFNEVIRPSIDRAIDLTR | 293-311 | 77789 |
| C. resistens | UDP-N-acetylmuramate-L-alanine ligase | AEI10413.1 | TATSVCTNEGGDNMDAGVISALLAGR | 92-117 | 50646 |
| C. sanguinis | Signal recognition particle protein | WP_259810719.1 | MQEQLGGGMPGMPGMPGAGK | 500-519 | 58202 |
| C. simulans | Carboxyl transferase domain-containing protein | WP_248092398.1 | APQGEVPAAR<br>AQEGIVSSAMQAR<br>TVGIIANR | 23-32<br>129-141<br>276-283 | 46640 |
| C. simulans | Nucleoside-diphosphate kinase | WP_062043559.1 | GDFALTVGENIVHGSDSPESAER<br>QLAGGTDPVSK<br>TLILIKPDGVK | 108-130<br>90-100<br>8-18 | 14958 |
| C. simulans | Hypothetical protein | WP_275437264.1 | DVTIQVEPDSGETSLK<br>VAAVDGSSAATEDGAVAVR | 58-73<br>39-57 | 20714 |
| C. simulans | IMP dehydrogenase | WP_046646763.1 | ASMGYTGSATLEELK | 462-476 | 53421 |
| C. striatum | Aldehyde dehydrogenase (NAD) family protein | WP_086891871.1 | AAPADAQGENTYSCLSSR<br>AVLELGGSDPYILLDTDDVK<br>AVLELGGSDPYILLDTDDVKEAAK<br>DAEICPR<br>ELGPLGMDEFVNK<br>FVGPNLVLGNTIVLK<br>IQGVSLTGSER<br>KAVLELGGSDPYILLDTDDVK<br>MYNTGQACNSNK<br>MYNTGQACNSNKR<br>RLEVGMANVNTPAGEGEELPFGGVK<br>YYADNGEEFTK | 294-311<br>228-247<br>228-251<br>158-164<br>439-451<br>143-157<br>202-212<br>227-247<br>258-269<br>258-270<br>408-432<br>95-105 | 49262 |
| C. striatum | DNA-directed RNA polymerase subunit alpha | WP_046646783.1 | DALASAGGTLVELFGLAR<br>EGPGVVTAGDIQPPAGVEIHNPDLHIASLNDTAK<br>GLVLSSDSDEPVVMYLSK<br>GYVPAAPTSGEIGR<br>IPVDQIYSPVLQVSYK<br>LDMELVVER<br>RQEIHTVGELAECTESDLLDIR<br>SYNCLK<br>TLLSSIPGAAVTSIK<br>VEATRVEQR | 203-220<br>100-133<br>82-99<br>145-158<br>159-174<br>134-142<br>263-284<br>257-262<br>41-55<br>175-183 | 36542 |
| C. striatum | Choline dehydrogenase | WP_110333688.1 | AIGVEYEWK<br>EFSPGPDVQTDEEILEWVR<br>EWVEAVR<br>GNPMDYEK<br>KPLAPLNDVTWYK<br>NDGETALHPSCTTK<br>QEGFAPFDR<br>SLLDTEAMAEVGAR | 247-255<br>460-478<br>436-442<br>114-121<br>551-563<br>479-492<br>197-205<br>446-459 | 65900 |
| C. striatum | Recombinase RecA | WP_166684297.1 | EGGIAAFIDAEHALDPEYAR<br>LMSQALR<br>SGSWFTYEGDQLGQGK | 97-116<br>181-187<br>298-313 | 41621 |

| | | | | | |
|---|---|---|---|---|---|
| C. striatum | Serine--tRNA ligase | WP_005529450.1 | DHLELGESLGLIDVK<br>DMLAAMELPYR<br>ELTSTSNCTTFQAR<br>ENPDVVR<br>IGQASPEERPALLEGSNELK<br>IGQASPEERPALLEGSNELKEK<br>KAEAELAEAEAK<br>KFDNEAWVPTQGCYR<br>SAILAADELR<br>TIDIAGGDLGSSAAR<br>YTGWSSCFR | 135-149<br>303-313<br>344-357<br>9-15<br>59-78<br>59-80<br>83-94<br>329-343<br>40-49<br>314-328<br>249-257 | 46201 |
| C. striatum | Type I polyketide synthase | WP_284765642.1 | AAVMVPGIDVPFHSR<br>AGAALFQTGGIIDVFR<br>CHVVLPGSPNR<br>FRDEVVAR<br>GGFLEGEGGGTVLLVR<br>GLGLTPNDVSVLSK<br>GQQYSVAGTK<br>GSAMGSLVPR<br>GSIDLTTVFLDR<br>GTFGGDGAYGEVK<br>HDTSTNANDPNESELHSLLWPAIGR<br>IDGEVVSR<br>IPANVSLDCVDPLIAPK<br>LLLWSVER<br>TAQFNTMFFDR<br>TRFEAEYGLQR<br>VMIDGYGLVDADK | 1475-1489<br>2817-2832<br>2167-2177<br>2414-2421<br>2690-2705<br>2759-2772<br>1449-1458<br>1391-1400<br>2437-2448<br>2178-2190<br>2773-2797<br>1241-1248<br>2836-2852<br>2143-2150<br>419-429<br>2355-2365<br>2954-2966 | 317703 |
| C. striatum | Malate dehydrogenase | WP_284810350.1 | AAFLVGARPR<br>ADLLEANGAIFTVQGK<br>GAEIIEVR<br>IAAGDVYGK<br>IQANVEELR<br>TAAVVSDGSYGVDEGLIFGFPTVAK<br>TLSQLSLK | 113-122<br>129-144<br>249-256<br>54-62<br>328-336<br>284-308<br>193-200 | 37652 |
| C. striatum | DoxX family membrane protein | HAT6550504.1 | EYVDDNKDDWQATGK<br>VGAGSTLALGK | 195-209<br>71-81 | 33955 |
| C. striatum | F0F1 ATP synthase subunit B | WP_086891558.1 | SDIGQNSINLAEK<br>YNAQLADAR | 146-158<br>85-93 | 19035 |
| C. striatum | Purine-nucleoside phosphorylase | WP_309221894.1 | FIADTFLEDVVQFNEVR<br>GAEIAETILLPGDPLR<br>QTAFHTMMK | 41-57<br>23-38<br>232-240 | 26957 |
| C. striatum | Cysteine synthase A | WP_049147014.1 | AVYNNILETIGNTPLVR<br>AYGAEIVLTPGAAGMK<br>IQGIGANFVPEVLDR<br>VESFNPANSVK<br>VILTMPETMSNER<br>YVSTVLFEDIR | 2-18<br>108-123<br>220-234<br>34-44<br>90-102<br>299-309 | 32886 |

| | | | | | |
|---|---|---|---|---|---|
| C. striatum | LutB/LldF family L-lactate oxidation iron-sulfur protein | WP_284786984.1 | DAGSAIKEDVAAR<br>TAALSSPIGHQALK | 68-80<br>314-327 | 55253 |
| C. striatum | DUF885 domain-containing protein | WP_306592399.1 | DTLLLMPQDTPEQLDAIR<br>QINEVFVQCER<br>YVLEGTDALVEWMQATADK | 126-143<br>174-184<br>294-312 | 62002 |
| C. striatum | 50S ribosomal protein L10 | WP_284789638.1 | LAGAESIVLTEYR | 17-29 | 18150 |
| C. striatum | Methyltransferase domain-containing protein | WP_086891631.1 | LPLQDDSVDAISVVFAPR | 155-172 | 30527 |
| C. striatum | TetR/AcrR family transcriptional regulator | WP_086890869.1 | TLVDWLFK | 100-107 | 27822 |
| C. striatum | Class I SAM-dependent methyltransferase | WP_070420299.1 | EVTAANPAHSENFAR | 6-20 | 21690 |
| C. striatum | Anaerobic C4-dicarboxylate transporter | WP_188311039.1 | MKDPEFAK | 222-229 | 48574 |
| C. striatum | GNAT family N-acetyltransferase | WP_306496603.1 | AALEDESTAGKR | 61-72 | 10433 |
| C. striatum | DEAD/DEAH box helicase family protein | WP_165482998.1 | EEAELRER | 879-886 | 133299 |
| C. striatum | Hypothetical protein | WP_005531174.1 | SDNTHAVADHASETYPEFSGK | 2-22 | 13214 |
| C. striatum | DUF402 domain-containing protein | WP_306495913.1 | YGDDAMAWLR<br>ANIFHFR | 151-160<br>66-72 | 19926 |
| C. striatum | Phosphoglycerate kinase | WP_049145845.1 | ANGLNDGDVMLIENVR<br>AQASVYDVAK<br>FGDKIVLPVDLIAATEFNADAENK<br>GYNVQNSLLQEDQIDNCK<br>ITASLPTIK<br>IVLPVDLIAATEFNADAENK<br>TLDDLIAEGVESR<br>TVFWNGPMGVFEMEAFSK<br>VVALDGIPEGWMSLDIGPESVK<br>YSLAPVAEALSEALGQYVALAGDVTGEDAHER | 107-122<br>163-172<br>264-287<br>241-258<br>41-49<br>268-287<br>5-17<br>320-337<br>288-309<br>75-106 | 42468 |
| C. striatum | 30S ribosomal protein S5 | WP_114975949.1 | QLVRPEEVAAR<br>SLEEVAPAQMLR<br>VPMIAGTITHPVEGR<br>AKEVPAAIQK<br>GKSLEEVAPAQMLR<br>NQQDNERDKYIER<br>SLGSDNALNVVR | 176-186<br>190-201<br>104-118<br>83-92<br>188-201<br>32-44<br>157-168 | 22259 |
| C. striatum | Universal stress protein | WP_049147615.1 | IVVGTDGSK<br>KIVVGTDGSK<br>LLGSVPADVAR | 7-15<br>6-15<br>123-133 | 15203 |
| C. striatum | NAD(P)/FAD-dependent oxidoreductase | WP_046645997.1 | TLRDQYSNYGTTSAK | 194-208 | 48838 |
| C. striatum | Long-chain-fatty-acid--CoA ligase | WP_284773384.1 | MQFPEISK | 205-212 | 65893 |
| C. striatum | 50S ribosomal protein L3 | WP_110092488.1 | MDDTSAYEVGQEVTADIFEGITFVDVTGTTK<br>TAETDGYNAIQIAFGEIDPR<br>IDAESNLLLIK<br>VGGIGACATPAR<br>VVPVTVVEAGPCVVTQIR | 59-70<br>44-63<br>185-195<br>148-159<br>26-43 | 22958 |
| C. striatum | Carbon-nitrogen hydrolase family protein | WP_005529738.1 | LVAAVQIQTGGDIEENLELAVEKIRVAAEGGAQLIVL<br>PEATSQAFGAGRLDK | 2-53 | 28302 |
| C. striatum | Holliday junction branch migration protein RuvA | WP_201807188.1 | MAERLALELR | 111-120 | 20183 |
| C. striatum | Oxoglutarate dehydrogenase inhibitor Odhl | WP_005531457.1 | HPEADIFLDDVTVSR<br>NSQVLHVGDEIQIGK<br>RGPNAGAR | 75-89<br>120-134<br>55-62 | 15535 |

| | | | | | |
|---|---|---|---|---|---|
| C. striatum | DUF4229 domain-containing protein | HCG3140840.1 | AWVQAELAGR<br>VEATETLAEYSAQR | 100-109<br>82-95 | 12117 |
| C. striatum | Cell division protein SepF | WP_049063104.1 | GADISTLELER | 130-140 | 15957 |
| C. striatum | RNA-binding S4 domain-containing protein | WP_306497072.1 | VAAADDDDDDYFDEATADDDFDPEKWR | 67-93 | 10127 |
| C. striatum | DUF4307 domain-containing protein | WP_284764964.1 | IAVDVPTNHR | 122-131 | 13429 |
| C. striatum | TetR/AcrR family transcriptional regulator | WP_110092290.1 | DQIATAALDLFDQR | 8-21 | 15043 |
| C. striatum | Carbohydrate ABC transporter permease | WP_100087946.1 | QLSNDSTEPVTVAIAR | 211-226 | 29736 |
| C. striatum | CarD family transcriptional regulator | WP_049166643.1 | EIDVEEAGNWSR<br>QILVGELALASPVDEK | 76-87<br>133-148 | 21937 |
| C. striatum | Phenylalanine--tRNA ligase beta subunit-related protein, partial | WP_218023517.1 | AVGENLIDELR | 104-114 | 14771 |
| C. striatum | MogA/MoaB family molybdenum cofactor biosynthesis protein | WP_100087247.1 | SVLDQMVPGIAQALR | 128-142 | 21043 |
| C. striatum | 3'-5' exonuclease | WP_284790400.1 | EMLADPGVEIPEAAAK<br>MLSFDLETTSVNPK | 42-57<br>8-21 | 25046 |
| C. striatum | MarR family transcriptional regulator | WP_131771608.1 | AGLNQNEAAEELGISK | 143-158 | 20131 |
| C. striatum | Type I-E CRISPR-associated protein Cas5/CasD | WP_100086675.1 | ALLENIESALR | 108-118 | 25688 |
| C. striatum | sigma-70 family RNA polymerase sigma factor | WP_049160475.1 | ALREAMLQGANT | 173-184 | 21090 |
| C. striatum | Hypothetical protein | WP_279109222.1 | QMAEMEK | 26-32 | 6334 |
| C. striatum | tRNA dihydrouridine synthase DusB | WP_114976561.1 | VESVDNLR | 319-326 | 41238 |
| C. striatum | Hypothetical protein | WP_204083840.1 | DRVAELNR | 191-198 | 25535 |
| C. striatum | Transglycosylase domain-containing protein | WP_168713047.1 | DAWMIGSTPQLATAVWVGTADNTSAIFNSYGGIMYG<br>SDAPTRIWKQILDTSLVNSEYQSFPTAYPIR | 550-616 | 75011 |
| C. terpenotabidum | DNA-directed RNA polymerase subunit beta (rpoB) | WP_041631175.1 | RQAQAEEGDR | 52-61 | 128747 |
| C. terpenotabidum | Glutamate synthase-related protein | WP_020441182.1 | LTGTVDAADPR | 1751-1761 | 200354 |
| C. terpenotabidum | Sucrose-6-phosphate hydrolase | AGP31162.1 | VLATVRHSGDR | 372-382 | 54368 |
| C. terpenotabidum | DUF732 domain-containing protein | WP_020440221.1 | DPETTAAAIK | 151-160 | 17030 |
| C. terpenotabidum | Deoxycytidylate deaminase | WP_020440770.1 | RAIELSEESR | 14-23 | 12280 |
| C. terpenotabidum | NAD(P)H-dependent glycerol-3-phosphate dehydrogenase | WP_041631453.1 | VCADAGCDVTLWAR | 18-31 | 34420 |
| C. terpenotabidum | 3-Dehydroquinate synthase | WP_020441204.1 | GLPCGEVFR | 43-51 | 59830 |
| C. tuberculostearicum | biotin/lipoyl-containing protein | WP_301714491.1 | IYAPFAGIVRCHVNVGDTVDTGVPLATVEATKLEAPV<br>ESPGPGKVHR | 3-49 | 21170 |
| C. tuberculostearicum | Hypothetical protein | WP_284821633.1 | VGSSAPMGGVR | 89-99 | 14194 |
| C. ulcerans | Lysophospholipid acyltransferase family protein | WP_197911364.1 | IAFTTNDPVIPVAMIGSREANPIGSWIPRPYKVRMK | 141-176 | 26999 |
| C. ulcerans | Pyruvate dehydrogenase | WP_046096294.1 | EQVLALAQK | 217-225 | 62233 |
| C. ulcerans | VIT1/CCC1 transporter family protein | WP_253698248.1 | GMNHEEAENKASEVFR | 221-236 | 38921 |
| C. ulcerans | Adhesin, putative phage tail fiber protein | WP_151812868.1 | AENAAATAAASATEAVAR | 134-151 | 48136 |
| C. ulcerans | Neocarzinostatin apoprotein domain-containing protein | WP_046693714.1 | SGGMGTTPLNADGTATATLHAASTGQGVDCK | 135-165 | 19515 |
| C. ulcerans | Transposase | WP_014836948.1 | LMLKMMSSIR | 20-29 | 13460 |
| C. ulcerans | Hypothetical protein | WP_111727762.1 | TGSSNGGRREGYGR | 48-61 | 45924 |
| C. urealyticum | Rrf2 family transcriptional regulator | WP_197913865.1 | QLTTFADLGLR | 1-11 | 17130 |
| C. urealyticum | Putative uncharacterized protein | WP_148791960.1 | ASYENVRDVGR | 110-120 | 47111 |
| C- urealyticum | Cadmium-translocating P-type ATPase | TYR16327.1 | AEMSAVVDSIR | 766-777 | 86554 |
| C. urealyticum | Abi-like protein | WP_012360942.1 | ATGEVEVYLR | 42-51 | 25398 |
| C. urealyticum | Signal recognition particle-docking protein FtsY | WP_317210142.1 | VLVGMGK | 384-390 | 57809 |
| C. urealyticum | Metal-sensitive transcriptional regulator | WP_012360560.1 | MDNAKPATK | 1-9 | 13375 |
| C. urealyticum | Hypothetical protein | WP_149121714.1 | EFTLTFED | 310-317 | 6525 |

| | | | | | |
|---|---|---|---|---|---|
| C. urealyticum | Pyruvate kinase | WP_148792403.1 | GLIDAGMNVAR | 23-33 | 50514 |
| C. urealyticum | 3-Methyl-2-oxobutanoate hydroxymethyltransferase | WP_149121667.1 | QDSPRGYMVPTK | 4-15 | 29554 |
| C. urealyticum | Molecular chaperone DnaJ | WP_149121529.1 | DLGVSDSASAEEIK | 13-26 | 41991 |
| C. urealyticum | Recombination regulator RecX | WP_012360124.1 | GNTADHVTTAGDTPDNR | 4-20 | 23808 |
| C. urealyticum | NUDIX hydrolase | WP_187470384.1 | MNTNDSATTSSGSPRSTR | 39-56 | 34469 |
| C. variabile | DNA cytosine methyltransferase | WP_148263290.1 | LISEVRPR | 157-164 | 50961 |
| C. variabile | Long-chain fatty acid--CoA ligase | WP_303944325.1 | GLCPVASQMVVIGNNRK | 506-522 | 68041 |
| C. variabile | polyketide synthase | WP_312775706.1 | LIQEIGSNLEAVSARR | 691-706 | 329271 |
| C. variabile | Type I methionyl aminopeptidase | WP_312776128.1 | SPAELDAMQAAGEIVGK | 14-30 | 27858 |
| C. variabile | Aspartate/tyrosine/aromatic aminotransferase | CUU67319.1 | AAALEREGTEILK | 32-44 | 46258 |
| C. variabile | IS3 family transposase | WP_148263200.1 | WKTATPAR | 133-140 | 35050 |
| C. variabile | Lrp/AsnC family transcriptional regulator | WP_312775823.1 | SVGLSEGAAR | 29-38 | 16077 |
| C. variabile | Hypothetical protein | WP_301518764.1 | ELTTDDLVSR | 57-66 | 18646 |
| C. variabile | 16S rRNA (uracil(1498)-N(3))-methyltransferase | WP_313007511.1 | WDGKPGKAEKAR | 119-130 | 26362 |
| C. variabile | Hypothetical protein | WP_234946299.1 | ECACAGTSHDAPGR | 548-561 | 74153 |
| C. variabile | Replicative DNA helicase | WP_141657038.1 | MDDGQWDKLTR | 345-355 | 58064 |
| C. variabile | WXG100 family type VII secretion target | WP_313358398.1 | MSFRTTTDVMHATAGK | 1-16 | 10324 |
| C. variabile | GntR-family transcription regulator | AEK37031.1 | MPDAGDAWEGGDAGAWAQR | 151-169 | 24346 |
| C. variabile | Hypothetical protein CVAR_0354 | AEK35702.1 | YVQPCGCMLIERKCCGK | 13-29 | 18015 |
| C. vitaeruminis | MalY/PatB family protein | WP_025253321.1 | MNIATSREILER | 352-363 | 41471 |
| C. vitaeruminis | Multidrug ABC transporter permease | WP_276652900.1 | AEPLLAGSLR | 373-382 | 55171 |
| C. vitaeruminis | Fructose-specific PTS transporter subunit EIIC | WP_276651473.1 | LDADLGADK | 13-21 | 70249 |
| C. vitaeruminis | GH32 C-terminal domain-containing protein | WP_025252491.1 | LVIAQSHNGR | 177-186 | 51164 |
| C. vitaeruminis | ATP-dependent DNA helicase | WP_313286629.1 | EAADAEGRNGFMEVAATHAALLMAQGAGRLLRR | 581-613 | 71575 |
| C. vitaeruminis | Molybdenum cofactor biosynthesis protein MoaE | WP_025251633.1 | EVAARVSAAHPK | 242-253 | 32567 |
| C. vitaeruminis | ATP-binding protein | WP_025253786.1 | YAGSYFR | 125-131 | 54312 |
| C. vitaeruminis | Sugar ABC transporter permease | WP_051645655.1 | ETATVGVDK | 345-353 | 36500 |
| C. vitaeruminis | MMPL family transporter | WP_025253758.1 | FSSWGDFAYR | 2-11 | 87643 |
| C. vitaeruminis | Sugar ABC transporter permease | WP_051483541.1 | ETATVGVDK | 348-356 | 36500 |
| C. vitaeruminis | Cation:dicarboxylase symporter family transporter | WP_277102341.1 | VNIFKLCK | 268-275 | 50429 |
| C. vitaeruminis | Holliday junction branch migration protein RuvA | WP_025252955.1 | SSLNALGSKR | 193-202 | 21029 |
| Actinobaculum massiliense | Glycogen synthase | WP_007001875.1 | AKEMGDAGRR | 377-386 | 45093 |
| Brevibacterium flavum | RpiR family transcriptional regulator | AKF27435.1 | CIEGLDTARR | 133-142 | 30559 |
| Clavibacter michiganensis | DNA topoisomerase IB | WP_045529628.1 | LASGETELR | 309-317 | 36094 |
| Clavibacter michiganensis | Preprotein translocase subunit SecA, partial | WP_316293024.1 | NETVELRER | 44-52 | 104589 |
| Clavibacter michiganensis | Carbohydrate ABC transporter permease | WP_094116250.1 | GGLTDGAVK | 313-321 | 34757 |
| Clavibacter michiganensis | Iron-sulfur cluster insertion protein ErpA | OUE30845.1 | VKSLLQQEGREDLRLR | 26-41 | 12935 |
| Clavibacter michiganensis | Aminomethyl-transferring glycine dehydrogenase | WP_204575344.1 | FIAAMIGIK | 904-912 | 105112 |
| Clavibacter michiganensis | UDP-N-acetylmuramoyl-L-alanine--D-glutamate ligase | WP_094113683.1 | TTTTQLTAALLQEGGVR | 149-165 | 54953 |
| Clavibacter michiganensis | Long-chain-fatty-acid--CoA ligase | WP_094116470.1 | VVDRTKDMILR | 424-434 | 56763 |
| Clavibacter michiganensis | Acyl-CoA dehydrogenase family protein | WP_080939243.1 | LSASGSVR | 213-220 | 43716 |
| Clavibacter michiganensis | DNA repair protein RecO | WP_094130231.1 | SLGLVDRSDAR | 234-244 | 32836 |
| Clavibacter michiganensis subsp. michiganensis | ATP-dependent RNA helicase DeaD | OUD99502.1 | GERRARPAR | 477-485 | 64092 |
| Clavibacter michiganensis | Mur ligase family protein | WP_104343076.1 | FGYEGLEVGR | 368-377 | 46163 |
| Clavibacter michiganensis | Right-handed parallel beta-helix repeat-containing protein | WP_079532018.1 | DVDIAQGTRR | 707-716 | 88998 |

| | | | | | |
|---|---|---|---|---|---|
| Clavibacter michiganensis | Phosphomannomutase/phosphoglucomutase | WP_209657882.1 | DQMAATGAVFGGEHSAHYYFR | 331-351 | 48868 |
| Clavibacter michiganensis | Alpha/beta hydrolase, partial | WP_259340805.1 | AGLGDSDPDPAPR | 66-78 | 30139 |
| Clavibacter michiganensis | ATP-dependent DNA helicase RecG | WP_104280092.1 | VMVIEDADR | 112-120 | 44554 |
| Clavibacter sepedonicus (Clavibacter michiganensis subsp. sepedonicus) | Hypothetical protein CMS2311 | CAQ02398.1 | MRPTAAINDAR | 1-11 | 25341 |
| Corynebacterium sp. KPL1995 | ATP-dependent Clp protease adapter protein ClpS | ERS42111.1 | LAVMNTRK | 11-18 | 15635 |
| Corynebacterium sp. Marseille-P3884; Periplasmic binding protein; Corynebacterium genitalium ATCC 33030 GN=HMPREF0291_10252 PE=4 SV=1 | ABC transporter substrate-binding protein | WP_210575019.1 | IGLYTAEDGR | 205-214 | 33735 |
| Corynebacterium sp. KPL1814 | Hypothetical protein HMPREF1257_01810 | ERS62660.1 | MSGVRDFPR | 1-9 | 27485 |
| Corynebacterium sp. KPL1818 | Bifunctional indole-3-glycerol-phosphate synthase TrpC/phosphoribosylanthranilate isomerase Trp | WP_023029750.1 | LLNQPEKKD | 467-475 | 51771 |
| Corynebacterium sp. KPL1818 | Hypothetical protein | WP_023030849.1 | CRCSSPPIAQGSSAPTW | 142-158 | 16041 |
| Corynebacterium sp. CCUG 65737 | M23 family metallopeptidase | WP_256003305.1 | SMKPAEGTFTSGFGMR | 111-126 | 24645 |
| Gordonia malaquae | Superfamily II RNA helicase | SEC55612.1 | VDDLSKR | 127-133 | 17493 |
| Mycobacterium tuberculosis | Daunorubicin resistance ABC transporter ATPase subunit | CND67133.1 | AEGLEKR | 8-14 | 8849 |
| Mycobacterium tuberculosis | DNA repair helicase XPB | WP_348790221.1 | EALFDAFR | 446-453 | 61287 |
| Mycobacteriales bacterium (soil metagenome)<br>Tsukamurella | Argininosuccinate synthase | NUS72237.1<br><br>WP_146488150.1 | IGQLTMR | 403-409 | 51581 |
| Nocardia zapadnayensis | Replication-associated recombination protein A | WP_265808822.1 | VPLHLRDAHYPGAK | 414-427 | 51972 |
| Tsukamurella paurometabola<br>Nocardia wallacei | Non-ribosomal peptide synthase/polyketide synthase | WP_013127570.1<br>WP_280333217.1 | AGELVLR | 338-344 | 42249 |
| Nocardia<br>Rhodococcus<br>Prescottella | Amino acid permease | WP_098086689.1<br>WP_033235792.1<br>RVW11666.1 | RGLTARHIR | 10-18 | 49262 |
| Prescottella equi | Short-chain fatty acyl-CoA regulator family protein | WP_084843967.1 | ICERTNCAQR | 440-449 | 52258,4 |
| Prescottella equi | Zinc binding alcohol dehydrogenase | WP_084960953.1 | CDACREGAIIR | 106-116 | 39992 |
| Prescottella equi | Hypotheical protein | WP_081206175.1 | MSPQEKMSLLWKR | 1-13 | 17001 |
| Prescottella equi | Ribosome recycling factor | WP_013415676.1 | IDEALFEAEEKMEK | 2-15 | 20765 |
| Prescottella equi | Nicotinate-nucleotide adenylyltransferase | WP_221282419.1 | HPAADPSQSSVQAEPGEH | 229-246 | 25842 |
| Prescottella equi<br>(formerly Rhodococcus equi) | TetR/AcrR family transcriptional regulator | WP_268965648.1 | RLPPEER | 11-17 | 22053 |
| Prescottella equi | DUF3499 domain-containing protein | WP_084968582.1 | EAGLGERPR | 87-95 | 12018 |
| Rhodococcus<br>Prescottella equi | LLM class F420-dependent oxidoreductase | NKR92484.1<br>WP_213573252.1 | YDAPLGR | 107-113 | 37344 |
| Prescottella equi | TPM domain-containing protein | WP_084987093.1 | AAAEAALHNAQSAK | 476-489 | 67785 |
| Prescottella equi | TraM recognition domain-containing protein | WP_143524795.1 | LADTMVDPR | 110-118 | 72821 |
| Rhodococcus<br>Prescottella equi<br>Nocardia sp | SDR family oxidoreductase | WP_317747646.1<br>MCA1005516.1<br>WP_297621523.1 | HGLVGLTK | 174-181 | 28691 |
| Prescottella equi | Type VII secretion target | WP_005514188.1 | ALLSDNAAAIGR | 59-70 | 10354 |
| Prescottella equi<br>Nocardia | AMP-binding protein | NKZ70514.1<br>WP_109529638.1 | GEVLEAFVVLR | 477-487 | 58188 |

| | | | | | |
|---|---|---|---|---|---|
| Prescottella equi | SDR family oxidoreductase | WP_013415346.1 | FMSVNMDGALNVCRAVVPHMR | 113-133 | 26613 |
| Prescottella equi | NAD(P)-dependent oxidoreductase | WP_084987694.1 | ILVAGAGGVVGLPLTR | 3-18 | 29331 |
| Nocardia<br>Rhodococcus<br>Prescottella | Amino acid permease | WP_098086689.1<br>WP_033235792.1<br>RVW11666.1 | RGLTARHIR | 10-18 | 49262 |
| Prescottella equi | TetR/AcrR family transcriptional regulator | WP_084985939.1 | ARAELPA | 211-217 | 36758 |
| Prescottella equi | Bifunctional phosphoribosylaminoimidazolecarboxamide formyltransferase/IMP cyclohydrolase | WP_069856157.1 | IAEAGIPVTK | 44-53 | 55572 |
| Prescottella equi | Glycosyltransferase | WP_084848591.1 | SLSVTLGR | 437-444 | 68982 |
| Prescottella equi | PaaI family thioesterase | WP_080561082.1 | KPTPLGK | 167-173 | 24489 |
| Prescottella equi | Enoyl-CoA hydratase-related protein | WP_221282843.1 | ELLVTGRTLSAER | 149-161 | 26831 |
| Prescottella equi | Hypothetical protein | WP_221286210.1 | TSSEAGFTR | 28-36 | 20934 |
| Prescottella equi | ABC transporter substrate-binding protein | WP_064059497.1 | KDGYMPAAK | 270-278 | 34869 |
| Prescottella equi | Glycosyltransferase family 87 protein | WP_286461333.1 | LASTIGGPVGR | 46-56 | 58446 |
| Prescottella equi | Cytochrome P450 | WP_221278048.1 | DPWGMYAALR | 21-30 | 44397 |
| Prescottella equi | APC family permease | WP_221281422.1 | AETDPTQRRR | 426-435 | 68073 |
| Prescottella equi | Glyoxalase | BCN68035.1 | EQVFDMSGATR | 6-16 | 14821 |
| Prescottella equi | Cellulose biosynthesis cyclic di-GMP-binding regulatory protein BcsB | WP_238772026.1 | MARTGAPSRAPR | 1-12 | 67672 |
| Prescottella equi | Tubulin-like doman-containing protein | WP_084848448.1 | GLAGMMMSGDATR | 302-314 | 128586 |
| Prescottella equi | Aldehyde dehydrogenase | WP_221282760.1 | MTDYDKLFIGGR | 1-12 | 50913 |
| Prescottella equi | Urease subunit alpha | WP_084848376.1 | GFLSAADDSDGDCAADRR | 125-142 | 76127 |
| Prescottella equi | Single-stranded DNA-binding protein | WP_238840838.1 | TRGGATDGSEGEAPSESEGR | 122-141 | 17205 |
| Prescottella equi | DUF11 domain-containing protein | WP_081205740.1 | DVTVNEGDFTGTGTMSAITCPDESK | 1385-1409 | 161162 |
| Prescottella equi | Alpha/beta hydrolase family protein | WP_205316481.1 | IAMLDAGGYNSDAMWGPPWDGRWMR | 205-229 | 37361 |
| Prescottella equi | Hypothetical protein REQ_34500 | CBH49444.1 | STGVSMTGQTPTSEGCSECRPLGASPATR | 22-50 | 18209 |
| Prescottella equi | Helix-turn-helix domain-containing GNAT family N-acetyltransferase | WP_022593503.1 | MRDLPK | 1-6 | 33162 |
| Prescottella equi | Helix-turn-helix domain-containing protein | WP_005513350.1 | AQYEAGASIR | 27-36 | 8133 |
| Rhodococcus sp. AG1013 | UDP-N-acetylglucosamine--N-acetylmuramyl-(pentapeptide) pyrophosphoryl-undecaprenol N-acetylglucosamine transferase | WP_114720055.1 | MLAGEPAPR | 1-9 | 39984 |
| Tsukamurella paurometabola | Carboxylesterase/lipase family protein | WP_013125362.1 | ATLVIDDR | 478-485 | 54645 |
| Tsukamurella paurometabola | Type II secretion system F family protein | WP_126196048.1 | AGMPTADAAR | 67-76 | 19978 |
| Tsukamurella<br>Mycobacteriales bacterium (soil metagenome) | Argininosuccinate synthase | WP_146488150.1<br>NUS72237.1 | IGQLTMR | 403-409 | 51581 |
| Tsukamurella paurometabola | Hypothetical protein | WP_176579865.1 | APYPRRSGGR | 676-685 | 75418 |
| Tsukamurella paurometabola | Hypothetical protein | WP_013125555.1 | QAADQAAADAAR | 1823-1834 | 220884 |
| Tsukamurella paurometabola | ABC transporter related protein | WP_041944652.1 | AEALIDSLGMTR | 125-136 | 28997 |
| Tsukamurella paurometabola<br>Nocardia wallacei | Non-ribosomal peptide synthase/polyketide synthase | WP_013127570.1<br>WP_280333217.1 | AGELVLR | 338-344 | 42249 |
| Tsukamurella paurometabola | LysR substrate-binding domain-containing protein | WP_041944519.1 | GVAPTAAATELAR | 63-75 | 31518 |
| Tsukamurella paurometabola | aminotransferase class I/II-fold Pyridoxal phosphate-dependent enzyme | WP_013125514.1 | ALAQGPSAAIIQPR | 228-241 | 49535 |
| Tsukamurella paurometabola | Glycosyltransferase family 39 protein | WP_013128542.1 | TETTTSATADR | 2-12 | 54912 |
| Tsukamurella paurometabola | DEAD/DEAH box helicase | WP_086012733.1 | VLASGMGR | 221-228 | 50057 |
| Tsukamurella paurometabola | Zinc-dependent metalloprotease | WP_126196731.1 | LVNPVAEK | 136-143 | 48395 |
| Tsukamurella paurometabola | PIG-L family deacetylase | WP_013126697.1 | LIVRLPPR | 224-231 | 27491 |
| Tsukamurella paurometabola | Conserved hypothetical protein | ADG77360.1 | MCHAVPCR | 1-8 | 5947 |
| Tsukamurella paurometabola | Iron ABC transporter permease | WP_013128332.1 | GLGLNVNLSR | 258-267 | 37272 |

| | | | | | |
|---|---|---|---|---|---|
| Tsukamurella paurometabola | TIGR04338 family metallohydrolase | WP_013125427.1 | VMGPEAGLALR | 139-149 | 16954 |
| Tsukamurella paurometabola | FdhF/YdeP family oxidoreductase | WP_013126973.1 | LNRSHLVHGR | 518-527 | 85189 |
| Tsukamurella paurometabola | Urea amidolyase associated protein UAAP1 | WP_013126244.1 | YGAGEAHSDSPAGR | 151-164 | 29752 |
| Tsukamurella paurometabola | Non-ribosomal peptide synthetase | WP_013128267.1 | FGAGDDICIGTPVAGR | 1252-1267 | 241289 |
| Tsukamurella paurometabola | Cellulase family glycosylhydrolase | WP_115329732.1 | ANNDSKKIWVESLCCHR | 231-247 | 43112 |
| Tsukamurella paurometabola | Hypothetical protein | WP_013126412.1 | DETIDFTGAESADDSPCER | 51-69 | 19864 |
| Tsukamurella paurometabola | Amino acid permease | WP_013126206.1 | TQTDYDHSALAHEEEGYHK | 2-20 | 52668 |
| Tsukamurella paurometabola | Acyl-CoA dehydrogenase family protein | WP_013128406.1 | ELWCQGYSEPGAGSDLANVGTTAR | 123-146 | 41867 |
