## Supplemental Table S2 for "Peptidome Analysis of Western Blots Identifies Natural Bispecific Antibody-Bound *Corynebacterium* and Phage B-cell Epitopes with Potential Relevance to Psoriasis"

### Table S2. 30S Ribosomal Proteins Are *Corynebacterium* Antigens.

| Antigen Number | Identified Antigen | Corynebacterium, Actinomycetota | Accession Number | Peptide Sequence | Peptide Number | Peptide Start/Stop | Molecular Weight |
|---|---|---|---|---|---|---|---|
| 1 | 30S ribosomal protein S1 | C. striatum | WP_086891322.1 | AFLEQTQSEVR | 1 | 187-197 | 53907 |
| | | C. ammoniagenes | WP_003848920.1 | ATQEDPWR | 2 | 276-283 | |
| | | C. stationis | WP_313679817.1 | DLEPYIGQELEAK | 3 | 158-170 | |
| | | C. casei | WP_301437303.1 | FNLHTAQIER | 4 | 423-432 | |
| | | C. pseudodiphtheriticum | WP_284587930.1 | GFLPASLVEMR | 5 | 144-154 | |
| | | C. propinquum | WP_239211683.1 | GGLILDIGLR | 6 | 134-143 | |
| | | C. phocae | WP_075734247.1 | HDVDPDEVVEVGDQIDALVLTK | 7 | 70-91 | |
| | | C. pyruviciproducens | WP_016457374.1 | IIELDKQR | 8 | 171-178 | |
| | | C. matruchotii | WP_314907517.1 | LVPFGAFVR | 9 | 302-310 | |
| | | C. glutamicum | WP_074495440.1 | NNVVLSR | 10 | 179-185 | |
| | | C. incognita | WP_185175853.1 | PTSNTPQVAINDIGTAEDFLAAVDATIK | 11 | 2-29 | |
| | | C. glucuronolyticum | MCI6205331.1 | QADEDYTEEFDPSK | 12 | 362-375 | |
| | | C. marquesiae | WP_337888753.1 | QAWEAR | 13 | 410-415 | |
| | | C. belfantii | WP_196977390.1 | RAFLEQTQSEVR | 14 | 186-197 | |
| | | C. tuberculostearicum | WP_301714479.1 | SEFLHQLQK | 15 | 198-206 | |
| | | C. callunae | WP_131385988.1 | TEGVIPSR | 16 | 57-64 | |
| | | C. kefirresidentii | WP_239205594.1 | THAVGQIVPGK | 17 | 288-298 | |
| | | C. kutscheri | WP_126369275.1 | VEEGIEGLVHISELAQR | 18 | 311-327 | |
| | | C. diphtheria | RKX01556.1 | VIDIDLER | 19 | 346-353 | |
| | | C. marinum | WP_042621326.1 | VIDIDLERR | 20 | 346-354 | |
| | | | | VRDLEPYIGQELEAK | 21 | 156-170 | |
| 2 | 30S ribosomal protein S2 | C. simulans | WP_062038471.1 | APSALWIVDTNK | 1 | 158-169 | 29593 |
| | | C. striatum | WP_005528910.1 | DAAEAAAASDPASAESAE | 2 | 251-268 | |
| | | C. phoceense | WP_257034878.1 | ELQAMDAAEDGYK | 3 | 116-128 | |
| | | C. endometrii | WP_136141404.1 | KQAQEAVAEEATR | 4 | 74-86 | |
| | | | | LLSGIIATAVEEGK | 5 | 213-226 | |
| | | | | LLSGIIATAVEEGKK | 6 | 213-227 | |
| | | | | MKELQAMDAAEDGYK | 7 | 114-128 | |
| | | | | QAQEAVAEEATR | 8 | 75-86 | |
| 3 | 30S ribosomal protein S3 | C. striatum | WP_086892080.1 | AEIDYGTAEAHTTFGR | 1 | 180-195 | 27970 |
| | | C. simulans | WP_248093772.1 | AGIADVVIER | 2 | 49-58 | |
| | | | | LGITSDWK | 3 | 12-19 | |
| | | | | MVALNILEVK | 4 | 97-106 | |
| | | | | QVDANATLVAQSVAEQLVNR | 5 | 107-126 | |
| | | | | VPLHTLR | 6 | 173-179 | |
| | | | | AEIDYGTAEAHTTFGR | 7 | 180-195 | |
| | | | | AGIADVVIER | 8 | 49-58 | |
| | | | | LGITSDWK | 9 | 12-19 | |
| | | | | MVALNILEVK | 10 | 97-106 | |
| 4 | 30S ribosomal protein S4 | C. striatum | WP_284822790.1 | AKYTYGVLER | 1 | 56-65 | 23408 |
| | | C. flavescens | WP_301515615.1 | AQIDVPLQEQLIVELYSK | 2 | 184-201 | |
| | | | | ESEYLLQLQEK | 3 | 43-53 | |
| | | | | IKESEYLLQLQEK | 4 | 41-53 | |
| | | | | IKESEYLLQLQEKQK | 5 | 41-55 | |

| | | | | | | | |
|---|---|---|---|---|---|---|---|
| | | | | ILVHQLPER<br>LDNVVYR<br>LRVDLVGGDMAFER<br>MEWFEDAQDR<br>SQKMEWFEDAQDR<br>TGDNLVVLLESR<br>VDLVGGDMAFER<br>YTYGVLER<br>YYAEANR<br>YYAEANRRPGK | 6<br>7<br>8<br>9<br>10<br>11<br>12<br>13<br>14<br>15 | 175-183<br>93-99<br>15-28<br>147-156<br>144-156<br>81-92<br>17-28<br>58-65<br>70-76<br>70-80 | |
| 5 | 30S ribosomal protein S5 | C. striatum<br>C. minutissimum | WP_114975949.1<br>WP_239186704.1 | QLVRPEEVAAR<br>SLEEVAPAQMLR<br>VPMIAGTITHPVEGR<br>AKEVPAAIQK<br>GKSLEEVAPAQMLR<br>NQQDNERDKYIER<br>SLGSDNALNVVR | 1<br>2<br>3<br>4<br>5<br>6<br>7 | 176-186<br>190-201<br>104-118<br>83-92<br>188-201<br>32-44<br>157-168 | 22259 |
| 6 | 30S ribosomal protein S6 | C. simulans<br>C. striatum | WP_061925220.1<br>WP_284773277.1 | DGGKVENVDVWGK<br>LNLNDTILR<br>TVSPSLDKFLEVVR | 1<br>2<br>3 | 36-48<br>82-90<br>21-34 | 11461 |
| 7 | 30S ribosomal protein S7 | Tsukamurella paurometabola | WP_126196419.1 | ANTLALR | 1 | 96-102 | 117472 |
| 8 | 30S ribosomal protein S7 | C. striatum | WP_272706990.1 | AFAHYR<br>ALGNIRPDLEVR<br>ANTLALR<br>DPVYNSEQVTMLVNK<br>EKTGTDPVGTLEK<br>IVYGALEACR<br>LANEILDASNGLGASVK<br>TGTDPVGTLEK<br>VGGATYQVPVEVKPAR | 1<br>2<br>3<br>4<br>5<br>6<br>7<br>8<br>9 | 149-154<br>64-75<br>95-101<br>14-28<br>51-63<br>41-50<br>119-135<br>53-63<br>79-94 | 17472 |
| 9 | 30S ribosomal protein S8 | C. camporealensis<br>C. ammoniagenes<br>C. minutissimum<br>C. aurimucosum | WP_046453190.1<br>WP_003848323.1<br>WP_039673739.1<br>WP_039673739.1 | GVGGEVLAYVW<br>QEGYIADYSVEGR<br>TMTDPIADMLSR<br>VSKPGLR | 1<br>2<br>3<br>4 | 117-127<br>42-54<br>2-13<br>75-81 | 13952 |
| 10 | 30S ribosomal protein S8 | C. lipophiloflavum | WP_006839458.1 | EASIAGLR | 1 | 71-78 | 14359 |
| 11 | 30S ribosomal protein S9 | C. striatum<br>C. simulans<br>C. accolens | WP_239299705.1<br>WP_248092707.1<br>PCC84031.1 | LHQQDILLPLTLLER<br>ALNVVYNPADR<br>EGQFDIK<br>KAGLLTR | 1<br>2<br>3<br>4 | 93-107<br>135-144<br>107-113<br>149-155 | 19031 |
| 12 | 30S ribosomal protein S10 | C. glutamicum<br>C. bovis<br>C. accolens<br>C. endometrii<br>C. urealyticum<br>C. auris<br>C. phoceense<br>C. belfantii<br>C. appendicis | AGT04507.1<br>WP_284833350.1<br>WP_005281226.1<br>WP_136140522.1<br>AGE35903.1<br>WP_290342546.1<br>WP_303936013.1<br>WP_197695650.1<br>WJY60247.1 | AYDHEAIDASAK<br>LIDILDPTPK<br>VVGPVPLPTEK | 1<br>2<br>3 | 12-23<br>73-82<br>36-46 | 11474 |

| | | | | | | | |
|---|---|---|---|---|---|---|---|
| | | C. resistens<br>C. striatum<br>C. diphtheria<br>C. jeikeium<br>C. urealyticum | AEI10242.1<br>WP_166684475.1<br>WP_199343317.1<br>WP_035012119.1<br>AGE35903.1 | | | | |
| 13 | 30S ribosomal protein S11 | C. glutamicum<br>C. camporealensis<br>C. halotolerans<br>C. striatum<br>C. callunae<br>C. minutissimum<br>C. appendicis<br>C. pyruviproducens<br>C. breve<br>C. cystitides<br>C. aurimucosum | WP_216312884.1<br>WP_046453194.1<br>WP_015399974.1<br>WP_306496458.1<br>WP_015650421.1<br>WP_239186691.1<br>WP_284835017.1<br>WP_016458936.1<br>WP_284825466.1<br>WP_257159300.1<br>MTD91744.1 | KSTPFAAQMAAENAAR<br>KVDVFVK<br>STPFAAQMAAENAAR | 1<br>2<br>3 | 62-77<br>86-92<br>63-77 | 14286 |
| 14 | 30S ribosomal protein S12 | C. pseudotuberculosis<br>C. confusum<br>C. diphtheriae<br>C. otitides<br>C. camporealensis<br>C. glutamicum<br>C. faecale<br>C. halotolerans<br>C. callunae<br>C. lipophiloflavum<br>C. efficiens<br>C. gallinarum<br>C. incognita<br>C. phoceense<br>C. breve<br>C. striatum<br>C. cystitides<br>C. urealyticum | AEK91670.1<br>WP_290224662.1<br>WP_014309991.1<br>WP_004601458.1<br>WP_035106521.1<br>WP_040072528.1<br>WP_290278344.1<br>WP_015399918.1<br>WP_015650380.1<br>WP_006839420.1<br>WP_006769815.1<br>WP_191732737.1<br>WP_185175283.1<br>WP_257035939.1<br>WP_284825393.1<br>WP_005527715.1<br>WP_092256507.1<br>WP_148792186.1 | PTIQQLVR<br>GALDTQGVKDR<br>VKDLPGVR<br>VYTTTPK | 1<br>2<br>3<br>4 | 2-9<br>100-110<br>87-94<br>37-43 | 13687 |
| 15 | 30S ribosomal protein S13 | C. striatum<br>C. ammoniagenes<br>C. stationis | WP_110332755.1<br>WP_003848284.1<br>WP_301201589.1 | TDNLDDDQLSALR<br>LAGVDLPR<br>MEVALTYIYGIGPTR | 1<br>2<br>3 | 45-57<br>4-11<br>15-29 | 13838 |
| 16 | 30S ribosomal protein S14 | C. striatum<br>C. incognita<br>C. halotolerans<br>C. phoceense<br>C. rouxii<br>C. diphtheriae | MDU3174082.1<br>WP_185175573.1<br>WP_015400294.1<br>WP_155872089.1<br>WP_257035430.1<br>CAB0947083.1 | LDAQFELNR<br>NPNTSDEDRLDAQFELNR | 1<br>2 | 42-50<br>33-50 | 11886 |
| 17 | 30S ribosomal protein S15 | C. propinquum<br>C. guangdongense | WP_302524571.1<br>WP_290194409.1 | ITNLTEHLK | 1 | 36-44 | 10444 |
| | | C. striatum<br>C. tuberculostearicum | WP_284822316.1<br>WP_302257953.1 | IGFADTAR | 2 | 55-62 | 10292 |
| 18 | 30S ribosomal protein S16 | C. simulans | WP_061923259.1 | AIENIGIYQPK | 1 | 33-43 | 18530 |

| | | | | | | | |
|---|---|---|---|---|---|---|---|
| | | C. aurimucosum<br>C. minutissimum<br>C. endometrii | WP_010190428.1<br>WP_115022740.1<br>WP_136141421.1 | FKGLPGAEGTLK<br>LDLFNAALAEANNGPTIEAITEK<br>QEPSLIQIDSER<br>VAEEKPSKLDLFNAALAEANNGPTIEAITEK | 2<br>3<br>4<br>5 | 83-94<br>103-125<br>44-55<br>95-125 | |
| 19 | 30S ribosomal protein S17 | C. striatum<br>C. lizhenjunii | WP_239299517.1<br>WP_165010665.1 | QHALYGK<br>SEANVNEVKK | 1<br>2 | 41-47<br>2-11 | 10725 |
| 20 | 30S ribosomal protein S17 | C. simulans<br>C. urogenitale<br>C. urealyticum | KXU18851.1<br>WP_151903394.1<br>WP_148820883.1 | AHDENETAGVGDR | 1 | 52-64 | 10602 |
| 21 | 30S ribosomal protein S17 | C. incognita<br>C. tuberculostearicum<br>C. accolens<br>C. massiliense<br>C. camporealensis<br>C. maris<br>C. lizhenjunii<br>C. striatum<br>C. phocae<br>C. halotolerans<br>C. guaraldiae<br>C. renale<br>C. mastitides<br>C. simulans | WP_185175302.1<br>WP_316986466.1<br>WP_284838521.1<br>WP_022863466.1<br>WP_035106541.1<br>WP_041631708.1<br>WP_165010665.1<br>WP_239299517.1<br>WP_075735611.1<br>WP_015399938.1<br>WP_144268734.1<br>WP_048379923.1<br>WP_026166028.1<br>KXU18851.1 | IAETRPLSK | 1 | 80-88 | 11513 |
| 22 | 30S ribosomal protein S18 | C. simulans<br>C. mendelii<br>C. guangdongense<br>C. massiliense | WP_005530160.1<br>WP_207279190.1<br>WP_290197819.1<br>WP_022862872.1 | EMALLPFTAR | 1 | 75-84 | 9825 |
