## Supplemental Table S3 for "Peptidome Analysis of Western Blots Identifies Natural Bispecific Antibody-Bound *Corynebacterium* and Phage B-cell Epitopes with Potential Relevance to Psoriasis"

### Table S3. 50S Ribosomal Proteins Are *Corynebacterium* Antigens.

| Antigen number | Identified Antigen | Corynebacterium, Actinomycetota | Accession number | Peptide Sequence | Peptide Number | Peptide Start/Stop | Molecular Weight |
|---|---|---|---|---|---|---|---|
| 1 | 50S ribosomal protein L1 | C. striatum | WP_101504662.1 | AAAELVDK<br>ASNLHAILGK<br>IAENYGALYDEILR<br>KADQLVR<br>KVTISTTSGPGIPVDASVQK<br>TGTVTADVAK<br>VAVFAEGEKAEAAK<br>VTISTTSGPGIPVDASVQK | 1<br>2<br>3<br>4<br>5<br>6<br>7<br>8 | 10-17<br>168-177<br>185-198<br>54-60<br>211-230<br>142-151<br>75-88<br>212-230 | 24583 |
| 2 | 50S ribosomal protein L2 | C. striatum<br>C. auris<br>C. aurimucosum | WP_284765305.1<br>WP_290342551.1<br>WP_158381161.1 | ASSVSEFEEITR<br>ATVGEVGNADQMNIR<br>NIPTGTTIHAVELKPGAGAK<br>QGTIVEAGANADIK<br>SAGASIQLLGK<br>TANIALLHYADGEK<br>TANIALLHYADGEKR<br>VAHIEYDPNR<br>YIIAPK | 1<br>2<br>3<br>4<br>5<br>6<br>7<br>8<br>9 | 15-26<br>191-205<br>135-154<br>113-126<br>158-168<br>89-102<br>89-103<br>79-88<br>104-109 | 30840 |
| 3 | 50S ribosomal protein L3 | C. simulans<br>C. diphtheria<br>C. deserti<br>C. vitaeruminis<br>C. terpenotabidum<br>C. falsenii<br>C. variabile | WP_284841061.1<br>CAB0536266.1<br>WP_053544144.1<br>WP_025251889.1<br>WP_020441963.1<br>WP_025403584.1<br>WP_313095095.1 | KAGVTPR<br>KLGMTQIFDEENR<br>LGMTQIFDEENR | 1<br>2<br>3 | 75-81<br>13-25<br>14-25 | 23154 |
| 4 | 50S ribosomal protein L3 | C. striatum | WP_110092488.1 | MDDTSAYEVGQEVTADIFEGITFVDVTGTTK<br>TAETDGYNAIQIAFGEIDPR<br>IDAESNLLLIK<br>VGGIGACATPAR<br>VVPVTVVEAGPCVVTQIR | 1<br>2<br>3<br>4<br>5 | 89-119<br>44-63<br>185-195<br>148-159<br>26-43 | 22958 |
| 5 | 50S ribosomal protein L4 | C. simulans<br>C. striatum | WP_284841060.1<br>HCG3139600.1 | AALFGALSDR<br>IHVIEELVPGQTPSTK<br>LDVQTAEGK<br>NVLLVVGR | 1<br>2<br>3<br>4 | 110-119<br>125-140<br>6-14<br>154-161 | 24032 |
| 6 | 50S ribosomal protein L5 | C. diphtheria<br>C. striatum<br>C. ammoniagenes<br>C. accolens<br>C. stationis<br>C. aurimucosum | WP_196975176.1<br>WP_005527723.1<br>WP_003848105.1<br>WP_237799349.1<br>WP_305939146.1<br>HCT9179808.1 | EGMPIGAR<br>GMDITVVTTATNDEEGR<br>GMDITVVTTATNDEEGRR<br>MWEFLDR<br>SENYTPR<br>VVVNMGVGDAAR<br>Akkermansia muciniphila (WP_102714965.1)<br>Query 1 GMDITVVTTATNDEEGR 17 | 1<br>2<br>3<br>4<br>5<br>6 | 37-48<br>59-64<br>59-65<br>85-92<br>85-93<br>71-78 | 14791 |

| | | | | | | | |
|---|---|---|---|---|---|---|---|
| | | | | G DI VTTA D+EGR<br>Sbjct 153 GFDIIFVTTAPTDDEGR 169 | | | |
| 7 | 50S ribosomal protein L6 | C. simulans<br>C. striatum | WP_062042260.1<br>WP_201817903.1 | ALHGLSR<br>GIRYEGEQVR<br>IGLAPIAVPSGVEIK<br>LSVSGIDK<br>LSVSGIDKQK<br>MEIFGVGYR<br>VGQVAANIR<br>VNGQEVEVK<br>YEGEQVR<br>YEGEQVRR | 1<br>2<br>3<br>4<br>5<br>6<br>7<br>8<br>9<br>10 | 63-69<br>161-170<br>4-18<br>131-138<br>131-140<br>87-95<br>141-149<br>19-27<br>164-170<br>164-171 | 19247 |
| 7 | 50S ribosomal protein L7/L12 | C. simulans<br>C. ammoniagenes<br>C. aurimucosum<br>C. maris<br>C. diphtheria<br>C. humireducens<br>C. confusum<br>C. tuberculostearicum<br>C. accolens<br>C. incognita<br>C. macginleyi<br>C. striatum<br>C. argentoratense<br>C. phoceense<br>C. matruchotii<br>C. urogenitale | AMO91403.1<br>EFG81066.1<br>WP_049358668.1<br>WP_020933892.1<br>WP_181998572.1<br>WP_040084814.1<br>WP_290224629.1<br>WP_284820937.1<br>WP_284831030.1<br>WP_185175272.1<br>WP_121927332.1<br>HCD1919209.1<br>WP_278764011.1<br>WP_257055087.1<br>WP_005519536.1<br>WP_151903420.1 | AILEGASKDDAEAAK<br>DELIEAFK<br>EIVSGLGLK<br>EMTLIELSEFVK<br>LEEAGATANLK<br><br>*Akkermansia muciniphila* (WP_215444414.1)<br><br>C. simulans 2 ILEGASKDDAEAAK 15<br>ILE SKD+A AAK<br>M. muciniphila 98 ILEAVSKDEANAAK 111 | 1<br>2<br>3<br>4<br>5 | 101-115<br>7-14<br>81-89<br>15-26<br>120-130 | 13400 |
| 8 | 50S ribosomal protein L9 | C. accolens<br>C. striatum | PCC83560.1<br>HBC8576744.1 | DLDHAR<br>GLAIVATR<br>KTGNYQVELK<br>NQLEQLSGVQVAMK<br>NYLLPR<br>TGNYQVELK | 1<br>2<br>3<br>4<br>5<br>6 | 61-66<br>34-41<br>125-134<br>70-83<br>28-33<br>126-134 | 16039 |
| 9 | 50S ribosomal protein L10 | C. striatum | HCG2987865.1 | GLTVSQLQELR<br>IAANEAGIEGLDDK<br>LAGAESIVLTEYR<br>LTGPTAIAFIK<br>NNLGFDVEYSVAK<br>NTADLAELK<br>NTADLAELKEK<br>TAAALQDKK | 1<br>2<br>3<br>4<br>5<br>6<br>7<br>8 | 30-40<br>59-72<br>17-29<br>73-83<br>41-53<br>6-14<br>6-16<br>161-169 | 18150 |
| 10 | 50S ribosomal protein L11 | C. striatum<br>C. flavescens | WP_049192497.1<br>MDN6460911.1 | EIAQTKFEDLNAR<br>FEDLNAR<br>GNVVPVEITVYEDR<br>IIAGTAR<br>SMGITVDGIPAK | 1<br>2<br>3<br>4<br>5 | 109-121<br>115-121<br>53-66<br>129-135<br>136-147 | 15530 |

| | | | | | | | |
|---|---|---|---|---|---|---|---|
| 11 | 50S ribosomal protein L13 | C. striatum | WP_110092867.1 | HSGYPGGLK<br>IHVSSNK<br>LASTVADMLR<br>LSDASIK<br>SMTLGQSLDTNPVR<br>VIEEAVKGMMPHNK | 1<br>2<br>3<br>4<br>5<br>6 | 117-125<br>102-108<br>68-77<br>154-160<br>126-139<br>140-153 | 20823 |
| 12 | 50S ribosomal protein L13 | C. accolens<br>C. endometrii<br>C. casei<br>C. cystitides<br>C. glaucum<br>C. tuberculostearicum<br>C. appendicis<br>C. incognita<br>C. camporealensis<br>C. propinquum | WP_284636650.1<br>WP_136140567.1<br>WP_025387096.1<br>WP_257159316.1<br>WP_095659270.1<br>WP_316992952.1<br>WP_076599157.1<br>WP_185175343.1<br>WP_035106601.1<br>WP_284797058.1 | LASTVADLLR | 1 | 28-37 | 16325 |
| 13 | 50S ribosomal protein L14 | C. diphtheria<br>C. sphenisci<br>C. mendelii | WP_071574145.1<br>WP_075691186.1<br>WP_207118774.1 | FAGIGDVIVATVK<br>FDENAAVLIK<br>IFGPVAR | 1<br>2<br>3 | 32-44<br>79-88<br>98-104 | 13303 |
| 14 | 50S ribosomal protein L14 | C. falsenii<br>C. argentoratense<br>C. durum<br>C. matruchotii | WP_025403573.1<br>WP_020975587.1<br>WP_006062960.1<br>WP_005519735.1 | RPDGSYIR FDENAAVIIK<br>*Akkermansia muciniphila* (MFR4436513.1)<br>Query 3 DGSYIRFDENAAVII 17<br>DGS RFD NA VII<br>Sbjct 71 DGSVLRFDGNAVVII 85 | 1 | 71-88 | 13353 |
| 15 | 50S ribosomal protein L15 | Candidatus Corynebacterium faecipullorum | HIX78999.1 | IEAAGGTATATK | 1 | 115-126 | 15509 |
| 16 | 50S ribosomal protein L16 | C. aurimucosum | WP_201828699.1 | WVANVKPGR<br>IIKKEDQF | 1<br>2 | 93-101<br>131-138 | 15749 |
| 17 | 50S ribosomal protein L17 | C. striatum<br>C. simulans | WP_046646782.1<br>WP_248092734.1 | MLRPYAEK<br>SGTLADRR | 1<br>2 | 43-50<br>57-64 | 18700 |
| 18 | 50S ribosomal protein L18 | C. striatum<br>C. simulans | WP_046646786.1<br>WP_248092739.1 | VAALAEAAR<br>AAGIEAVVFDR | 1<br>2 | 119-127<br>100-110 | 14430 |
| 19 | 50S ribosomal protein L19 | C. simulans<br>C. striatum | WP_061923234.1<br>MBD0857342.1 | ANLIDKVDAAQLR<br>DDVPDFRPGDTLDVNVK<br>RQGAGIR<br>TFPVHSPNIESIQVVR | 1<br>2<br>3<br>4 | 2-14<br>15-31<br>50-56<br>73-88 | 13146 |
| 20 | 50S ribosomal protein L20 | C. diphtheria<br>C. accolens<br>C. pseudodiphthereticum<br>C. striatum<br>C. simulans | WP_088251251.1<br>WP_302518868.1<br>WP_302517286.1<br>HAT1213399.1<br>WP_062040644.1 | LAEIEVDRK | 1 | 85-93 | 14653 |
| 21 | 50S ribosomal protein L21 | C. pseudodiphtheriticum<br>C. auris<br>C. durum<br>C. accolens | WP_284575292.1<br>WP_290341644.1<br>WP_303132148.1<br>WP_284639488.1 | VAEGDLVKVEK | 1 | 14-24 | 10846 |

| | | | | | | | |
|---|---|---|---|---|---|---|---|
| 22 | 50S ribosomal protein L22 | C. striatum<br>C. casei | WP_306592414.1<br>WP_006823226.1 | RVIDLVR<br>SETITSASATAR | 1<br>2 | 24-30<br>2-13 | 12955 |
| 23 | 50S ribosomal protein L22 | C. casei<br>C. striatum<br>C. singulare | WP_006823226.1<br>WP_306592414.1<br>WP_239180415.1 | SVAEALAILK<br>VVASAAANAENNFGLDPR<br>YAPQGAAKPVAK | 1<br>2<br>3 | 33-42<br>55-72<br>43-54 | 12955 |
| 24 | 50S ribosomal protein L23 | C. simulans<br>C. striatum | WP_062042289.1<br>HCG2966061.1 | DAIEQIFGVK<br>VDSVNTVNR | 1<br>2 | 44-53<br>54-62 | 11090 |
| 25 | 50S ribosomal protein L24 | C. diphtheriae | WP_072574660.1 | DKVLVEGVNR<br>GAESGGIVTQEAPIHVSNVAIVDSEGNPTR | 1<br>2 | 31-40<br>53-82 | 11156 |
| 26 | 50S ribosomal protein L25 | C. aurimucosum<br>C. simulans<br>C. accolens<br>C. striatum | WP_010187470.1<br>WP_062035544.1<br>PCC83094.1<br>WP_347004832.1 | ANTRPVLK<br>IELTALVR | 1<br>2 | 2-9<br>49-56 | 23370 |
| 27 | 50S ribosomal protein L27 | Corynebacterium<br>(unknown species) | WP_075730276.1 | FGGQQVK<br>IVNIVPAEQAAVEASA | 1<br>2 | 26-32<br>77-92 | 9774 |
| 28 | 50S ribosomal protein L28 | C. massiliense<br>C. phocae<br>C. stationis | WP_022862869.1<br>WP_075735033.1<br>WP_075723188.1 | IIDRDGIESVVAK<br>SAICQVTGR<br>YYLPSEGR | 1<br>2<br>3 | 58-70<br>2-10<br>38-45 | 8950 |
