## Supplemental Table S4 for "Peptidome Analysis of Western Blots Identifies Natural Bispecific Antibody-Bound *Corynebacterium* and Phage B-cell Epitopes with Potential Relevance to Psoriasis"

### Table S4. DNA-Transcription and Translation Proteins Are *Corynebacterium* Antigens.

| Antigen number | Identified Antigen | *Corynebacterium* (C.), Actinomycetota | Accession number | Peptide Number | Peptide Sequence | Peptide Start/Stop | Molecular Weight |
|---|---|---|---|---|---|---|---|
| 1 | DNA-directed RNA polymerase subunit beta (rpoB) | C. terpenotabidum | WP_041631175.1 | 1 | RQAQAEEGDR | 52-61 | 128747 |
| 2 | DNA polymerase III alpha subunit | C. variabile | WP_141329024.1 | 1 | VTDSLIKAGAFDSLGHPRK | 909-927 | 132246 |
| | | C. falsenii | WP_119664632.1 | | | | |
| | | C. nuruki | WP_010122200.1 | | | | |
| | | C. provencense | WP_066588926.1 | | | | |
| 3 | DNA-directed RNA polymerase subunit beta (rpoB) | C. simulans (1-12,14-20,22-25) | WP_339019060.1 | 1 | AAYDAGDLIITPK | 613-625 | 117597 |
| | | C. phoceense (1,6,22) | WP_257045771.1 | 2 | AGDILVGK | 777-784 | |
| | | C. striatum (1,2,6-12,14-20,22-25) | WP_347004530.1 | 3 | AGIVENVTADLITIMDDEGQR | 626-646 | |
| | | C. aurimucosum (1,2,6,7,8,24) | WP_216382059.1 | 4 | AGIVENVTADLITIMDDEGQRDTYMLR | 626-652 | |
| | | C. singulare (1,6,7) | WP_144793647.1 | 5 | ALGWSEEQIR | 220-229 | |
| | | C. minutissimum (1,6,7) | WP_239186749.1 | 6 | DGDVMVDEHGK | 977-987 | |
| | | C. intestinale (1,6,7) | WP_250224080.1 | 7 | EIPNVSDDVLR | 751-761 | |
| | | C. guaraldiae (1,6,7) | WP_144269193.1 | 8 | ELAQSLLDNSFFR | 270-282 | |
| | | C. hesseae (1,6,7) | MFG6303322.1 | 9 | FSREDDDDLAPGVNEMIR | 832-849 | |
| | | C. caspium (2,8,17) | WKD59862.1 | 10 | GETELTPEER | 789-798 | |
| | | C. testudinoris (2,10-15,17,18,20,21,25) | WP_201774826.1 | 11 | LGAEEITR | 743-750 | |
| | | C. atrinae (2,11-17,18,20,21,25) | WP_290219190.1 | 12 | LSALGPGGLSR | 434-444 | |
| | | C. flavescens (2,7,8) | WP_301506945.1 | 13 | MTTQDAESITPTSLINVRPVSAAIR | 383-407 | |
| | | C. atypicum (2) | AIG63631.1 | 14 | QAVPLVR | 593-599 | |
| | | C. suedecumii (2,9,11,14,15,17,18,23) | WP_284876167.1 | 15 | SPGVYFDQTIDK | 162-173 | |
| | | C. humireducens (2,11,14,15,23) | WP_082028335.1 | 16 | SQTVFIGDFPMMTDK | 129-143 | |
| | | C. marinum (2,11,14,15,18) | WP_188656879.1 | 17 | STERPLHSVK | 174-183 | |
| | | C. incognita (2,9,11,15,23) | WP_185175274.1 | 18 | STGPYSMITQQPLGGK | 1025-1040 | |
| | | C. hylobatis (2) | WP_206436357.1 | 19 | TLPEELYDVPAGSLTATPVFDGASNEEIGR | 938-967 | |
| | | C. pollutisoli (2,11,14,15,18,23) | WP_205410452.1 | 20 | TNQGTNYNQTPLVELGQR | 657-674 | |
| | | C. comes (2,14,15) | WP_197085778.1 | 21 | TVGELIQNQVR | 359-369 | |
| | | C. nasicanis (2) | WP_377001948.1 | 22 | VDVDDPANAELLK | 925-937 | |
| | | C. alimapuense (2) | WP_197714729.1 | 23 | VEAGQVLADGPGTHNGEMSLGR | 675-696 | |
| | | C. accolens (2,5,8) | WP_284628546.1 | 24 | VNAFGFIETPYR | 485-496 | |
| | | C. marquesiae (2,5,8,9,24) | WP_284821753.1 | 25 | VVVSQLVR | 154-161 | |
| | | C. tuberculostearicum (2,5,8,24) | WP_316992046.1 | | | | |
| | | C. lowii (2) | WP_055175527.1 | | | | |
| | | C. renale (2) | WP_111725639.1 | | | | |
| | | C. atypicum (2) | WP_038607514.1 | | | | |
| | | C. caspium (2,8) | WP_018340281.1 | | | | |
| | | C. mastitidis (2) | WP_337890029.1 | | | | |
| | | C. kefirresidentii (2,5,8) | WP_086589060.1 | | | | |
| | | C. frankenforstense (2) | WP_075663206.1 | | | | |
| | | C. confusum (5,8,9,22) | WP_290224634.1 | | | | |
| | | C. yonathiae (5,8,9) | WP_238800425.1 | | | | |
| | | C. segmentosum (5) | WP_126318251.1 | | | | |
| | | C. lizhenjunii (5,26) | WP_165008708.1 | | | | |
| | | C. imitans (6) | UWV67219.1 | | | | |

| | | | |
|---|---|---|---|
| | | C. pseudogenitalium (6) | AAT96666.1 |
| | | C. endometrii (7,10,14-16,18,20,21,23,25) | WP_136142094.1 |
| | | C. yudongzhengii (7) | WP_108430823.1 |
| | | C. macginleyi (8) | WP_200445465.1 |
| | | C. felinum (8) | WP_277104557.1 |
| | | C. curiae (8) | WP_269946993.1 |
| | | C. camporealensis (8) | WJY91269.1 |
| | | C. faecale (9-16,18,20,21,25) | WP_290279780.1 |
| | | C. efficiens (9-16,18,20,21,25) | WP_035110049.1 |
| | | C. gerontici (9-18,20,21,24,25) | AIG06683.1 |
| | | C. pseudotuberculosis (9-18,20,21,23,25) | WP_172815612.1 |
| | | C. silvaticum (9-15,17,18,20,21,23,25) | WP_171483329.1 |
| | | C. ulcerans (9-15,17,18,20,21,23,25) | PME07283.1 |
| | | C. flavescens (9-15,18,20,21,23,24,25) | GEB98300.1 |
| | | C. durum (9,11,14,15) | WP_196794122.1 |
| | | C. epidermidicanis (9,11,14) | WP_158408008.1 |
| | | C. occultum (9,11,24) | WP_156229963.1 |
| | | C. halotolerans (9,11,15,23,24) | WP_211208981.1 |
| | | C. glutamicum (9,11) | WP_063967310.1 |
| | | C. casei (9,22,24) | WP_193024522.1 |
| | | C. deserti (9) | WP_053544124.1 |
| | | C. gallinarum (9) | WP_191732865.1 |
| | | C. hindlerae (9) | WP_182386996.1 |
| | | C. callunae (9) | WP_247775984.1 |
| | | C. kutscheri (9) | WP_172595864.1 |
| | | C. matruchotii (10-18,20,21,25) | EEG27920.1 |
| | | C. faecigallinarum (10,12,13,14,16,18,25) | HJC84208.1 |
| | | C. argentoratense (10-18,20,21,25) | AGU14430.1 |
| | | C. diphtheria (10,12-18,20,21,23,24,25) | AEX41129.1 |
| | | C. rouxii (10,12-15,17,18,20,21,23,24,25) | QIN91254.1 |
| | | C. riegelii (10,11,12,13,17,18,20,21,25) | GAA0210583.1 |
| | | C. belfantii (10,12-18,20,21,24,25) | SNW30590.1 |
| | | C. jeikeium (10,12,13,16-18,20,21,25) | WP_035012124.1 |
| | | C. appendicis (10,11,12,17,18) | WJY60230.1 |
| | | C. massiliense (2,8,10,11,12,14,17,18,23) | WCZ31773.1 |
| | | C. glyciniphilum (10,12,13,16,18,25) | WP_304041269.1 |
| | | C. bovis (10,12,13,16-18,20,21,25) | RRO83958.1 |
| | | C. falsenii (10,12,16-18,20,21,25) | WP_276784400.1 |
| | | C. dentalis (10,13,16,18,20,21,25) | WP_312097320.1 |
| | | C. urealyticum (10,12,13,16,18,20,21,25) | WP_012359561.1 |
| | | C. resistens (10,12,13,16,18,20,21,25) | WP_273351542.1 |
| | | C. urogenitale (10,13,16,20,25) | WP_151903418.1 |
| | | C. pyruviciproducens (11) | EPD67850.1 |
| | | C. glucuronolyticum (11) | WKD62731.1 |
| | | C. canis (11) | WP_146325663.1 |
| | | C. nasicanis (11) | WP_377001948.1 |
| | | C. uterequi (12,13,14,18,20,21,25) | AKK10318.1 |
| | | C. gallistercoris (12,16,20,21) | HIW96393.1 |
| | | C. glaucum (17) | WP_095659190.1 |

| | | | | | | | |
|---|---|---|---|---|---|---|---|
| | | C. kroppenstedtii (17)<br>M. tuberculosis (2,10,12,14,18,21,25) | HJD68837.1<br>WP_343358947.1 | | | | |
| 4 | DNA-directed RNA polymerase subunit beta' (rpoC) | C. striatum (1-12,13,16-27,29,30,31,32) | WP_110333590.1 | 1 | AAKLEQDLAELEAAGAK | 180-196 | 148211 |
| | | C. simulans (1-11,13,16-27,29,30,31,32) | WP_284841070.1 | 2 | AVEDLYPDDNPIPMIVK | 805-821 | |
| | | C. accolens (4,5,8,9,10,11,16-21,23,24,25,27-32,33) | WP_284897177.1 | 3 | DRGVLGLQAPIK | 633-644 | |
| | | C. phocae (1,4,5,7,13,16-19,21,24,26,29) | WP_075735677.1 | 4 | EGLSVLEYFNNSHGSR | 859-874 | |
| | | C. guaraldiae (1,13,29) | WP_144015140.1 | 5 | GAADPHDVLEVLGR | 1136-1149 | |
| | | C. aurimucosum (1,3,13,29) | WP_193633751.1 | 6 | GTVIDSGSTDYLPGVLVDLSEAK | 1187-1209 | |
| | | C. singulare (1,3,13,24,29) | WP_144793643.1 | 7 | HLIDEVQAVYR | 1155-1165 | |
| | | C. hesseae (1) | WP_269949020.1 | 8 | IGLATADDIRR | 18-28 | |
| | | C. minutissimum (1,4,8-10,13,16,17,20,21,23-25,28-32) | WP_115021140.1 | 9 | KPETINYR | 36-43 | |
| | | C. hiratae (1,13,29) | WP_144014110.1 | 10 | KYWEQGALTER | 775-785 | |
| | | C. riegelii (2,5,7,8,9,14,15,18-20,23,25,27,28,30,31,32) | GAA0210575.1 | 11 | LDEIWNTFLK | 225-234 | |
| | | C. glutamicum (2,7,8,9,15,16,20, 23,25,27,30) | WP_087497655.1 | 12 | LDMVTGLYYLTMDKAESEIGGEGR | 585-608 | |
| | | C. massiliense (2,4,5,7,9,15-17,20,21,24,25,27,29,31.32) | WCZ31774.1 | 13 | LEQDLAELEAAGAK | 183-196 | |
| | | C. ureicelerivorans (2,5,7-9,14,15,18-20,23,24,25,27-32) | WP_301694878.1 | 14 | LGIQAFEPK | 513-521 | |
| | | C. fournieri (2,5,14,18,19,20,24,25,27,30,31,32) | WP_085956825.1 | 15 | LGYLLDLAPK | 120-129 | |
| | | C. gerontici (2,4,8,16,18,19,21,25) | WP_123935332.1 | 16 | LVELWK | 791-796 | |
| | | C. comes (2,4,7,8,9,14,17,20,23,28,30,31,32) | WP_156226866.1 | 17 | LVENDYAQNIK | 466-476 | |
| | | C. suedecumii (2,4,7,8,28,30,31,32) | WP_284875859.1 | 18 | MIDLGAPEIIVNNEK | 363-377 | |
| | | C. pollutisoli (2,4,7,8,27,30,31,32) | WP_085550527.1 | 19 | MIDLGAPEIIVNNEKR | 363-378 | |
| | | C. marinum (2) | WP_042620696.1 | 20 | MLQESVDALFDNGR | 379-392 | |
| | | C. imitans (2,3,5,7,24) | WP_038588203.1 | 21 | NISVKPTEAAR | 1291-1301 | |
| | | C. mucifaciens (2,5,24) | WP_168684069.1 | 22 | NSVLLGDVINDLAAK | 705-719 | |
| | | C. guangdongense (2) | WP_290197322.1 | 23 | QIWTLAGMK | 830-838 | |
| | | C. glaucum (2,5,24, | WP_095659191.1 | 24 | RPMESNPDAMIER | 1107-1119 | |
| | | C. cystitides (2,5,24) | WP_257161937.1 | 25 | SEIMGITK | 1227-1234 | |
| | | C. mastitides (2,3,24) | WP_284829705.1 | 26 | SLRDGDHVTTGER | 1120-1132 | |
| | | C. faecale (2,3) | WP_290278340.1 | 27 | SLSDLLK | 407-413 | |
| | | C. urinipleomorphum (2,3,24) | WP_087117120.1 | 28 | SVIIVGPQLK | 434-443 | |
| | | C. deserti (2,5) | WP_053544125.1 | 29 | SVLTCQTPAGVCAK | 980-993 | |
| | | C. appendicis (2,5,24) | WP_076598034.1 | 30 | TADSGYLTR | 884-892 | |
| | | C. breve (2) | WP_284825391.1 | 31 | TFHQGGVGGDITGGLPR | 1030-1046 | |
| | | C. sanguinis (2,3,5) | WP_144739993.1 | 32 | VLTDAAINKR | 1254-1263 | |
| | | C. auris (2,5,24) | WP_290342526.1 | 33 | DVEDETESEIAER | 161-173 | |
| | | C. uterequi (3-5,7,8,9,14,15,17,20,23,25,27,28,30-32) | WP_047258935.1 | | | | |
| | | C. durum (3,7-9,15,16,18-21,23,25,27,30,31,32) | WP_179418789.1 | | | | |
| | | C. oculi (3,4,7-9,14,15,18,19,20,23-25,27,28,30-32) | WP_055122282.1 | | | | |
| | | C. ammoniagenes (3,4,5,8,9,15,16,17-21,23,25,27-32) | EFG81071.1 | | | | |
| | | C. stationis (3,4,5,7,8,9,14-21,23,27,28-32) | AAZ91716.1 | | | | |
| | | C. otitides (3,4,8,14,16,18,19,25) | WP_004601464.1 | | | | |
| | | C. genitalium (3,5,7,18,19,21,25) | WP_005286797.1 | | | | |
| | | C. canis (3,7) | WP_146325662.1 | | | | |
| | | C. gallinarum (3) | WP_191732739.1 | | | | |
| | | C. camporealensis (3,5,22,29) | WP_321115343.1 | | | | |
| | | C. lipophiloflavum (3) | WP_006839418.1 | | | | |
| | | C. pseudodiphtheriticum (3,24) | WP_027017514.1 | | | | |
| | | C. propinquum (3,29) | WP_049148394.1 | | | | |
| | | C. pseudotuberculosis (4,8-10,14-21,23,25,27,28,30-32) | WP_014558579.1 | | | | |

| | | | | | | | |
|---|---|---|---|---|---|---|---|
| | | C. matruchotii (4,8,9,14-20,23,25,27,28,30,31,32)<br>C. ulcerans (4,8,9,10,14-21,23,25,27,28,30,31,32,<br>C. kroppenstedtii (4,8,9,15,18-20,23,27,28,30,31,32)<br>C. pseudogenitalium (4,5,8,10,14-17,20,21,24,25,28-32)<br>C. lactis (4,8,9,15,18,19,27,28)<br>C. diphtheria (4,8,10,14-17,20,21,23,25,27,28,30-32)<br>C. humireducens (4,7,17)<br>C. belfantii (4,8-10,14-17,20,21,23,25,27,28,30-32)<br>C. testudinoris (4,7,18,19,21)<br>C. rouxii (4,8-10,14-17,20,21,23,25,28,30-32)<br>C. silvaticum (4,10,16,21)<br>C. halotolerans (4,7,21,29)<br>C. atrinae (4,7,8,9,14,15,18,20,21,23,27,28,30,31,32)<br>C. marinum (4,7,17)<br>C. nasicanis (4,7,8,9,14,15,17-20,23,27,28,30,31)<br>C. afermentans (5,7-9,14,18-21,24,25,27,28,30-32)<br>C. renale (5)<br>C. urinipleomorphum (5)<br>C. glucuronolytikum (7,9,14-16,18-21,27,28,30-32)<br>C. pyruviciproducens (7,14-16,18-21,27,28,30-32)<br>C. maris (7)<br>C. occultum (7)<br>C. ulceribovis (9,15,20,27,28,30-32)<br>C. felinum (10)<br>C. parakroppenstedtii (11)<br>C. macginleyi (11,29,33)<br>C. incognita (11)<br>C. segmentosum (11)<br>C. atypicum (15,18-21,23,25,27-29,30-32)<br>C. casei (29)<br>C. kefirresidentii (29)<br>C. tuberculostearicum (29)<br>C. flavescens (26,29)<br>C. phoceense (26)<br>M. tuberculosis (8,9,14-17,20,27,30)<br>M. leprae (8,9,14,15,20,27,30) | EEG27921.1<br>WP_053825302.1<br>KXB49578.1<br>EFQ81722.1<br>ALA66682.1<br>WP_106361354.1<br>WP_040084818.1<br>WP_374064188.1<br>WP_047252295.1<br>WP_155871610.1<br>WP_087453364.1<br>WP_015399915.1<br>WP_290219193.1<br>WP_042620696.1<br>WP_377001949.1<br>WP_126857343.1<br>WP_087117120.1<br>WP_087117120.1<br>WP_005394804.1<br>WP_280193574.1<br>WP_020933895.1<br>WP_156229964.1<br>WP_018023258.1<br>WP_277104556.1<br>WP_340051502.1<br>WP_121953163.1<br>WP_185175275.1<br>WP_126318253.1<br>WP_038604371.1<br>WP_193024523.1<br>WP_086589061.1<br>WP_316992980.1<br>WP_301501430.1<br>WP_257052677.1<br>WP_031720909.1<br>WP_010908593.1 | | | | |
| 5 | DNA-directed RNA polymerase subunit alpha | C. striatum | WP_046646783.1 | 1<br>2<br>3<br>4<br>5<br>6<br>7<br>8<br>9<br>10 | DALASAGGTLVELFGLAR<br>EGPGVVTAGDIQPPAGVEIHNPDLHIASLNDTAK<br>GLVLSSDSDEPVVMYLSK<br>GYVPAAPTSGEIGR<br>IPVDQIYSPVLQVSYK<br>LDMELVVER<br>RQEIHTVGELAECTESDLLDIR<br>SYNCLK<br>TLLSSIPGAAVTSIK<br>VEATRVEQR | 203-220<br>100-133<br>82-99<br>145-158<br>159-174<br>134-142<br>263-284<br>257-262<br>41-55<br>175-183 | 36542 |
| 6 | DNA gyrase subunit A | C. striatum<br>C. simulans | WP_100619148.1<br>WP_248092847.1 | 1<br>2<br>3 | ENTVDFSPNYDGK<br>GAELKQDDIVK<br>GQHVANLLEFQPEEK | 144-156<br>542-552<br>589-603 | 76034 |

| | | | | | | | |
|---|---|---|---|---|---|---|---|
| | | | | 4<br>5<br>6<br>7<br>8<br>9<br>10<br>11<br>12<br>13 | IIDELAEIEEIIADLK<br>MTPLAMEMVR<br>QIVHDELAEIVEK<br>SVEDVQQEHLK<br>TKVDAYK<br>TQIVAASGDVTDEDLIAR<br>TGEYLSR<br>VEHTEPGLQGDR<br>VETATVDRR<br>VTDKTFNAGVK | 447-462<br>131-140<br>473-485<br>832-842<br>525-531<br>492-509<br>178-184<br>133-144<br>60-68<br>49-59 | |
| 7 | DNA topoisomerase (ATP-hydrolyzing) subunit B | C. simulans<br>C. striatum<br>C. phoceense<br>C. tuberculostearicum<br>C. marquesiae<br>C. aurimucosum<br>C.kefirresidentii<br>C. guaraldiae<br>C. pseudogenitalium<br>C. curieae<br>C. flavescens<br>C. singulare<br>C. minutissimum | WP_062036505.1<br>WP_201806516.1<br>WP_257052411.1<br>WP_293821919.1<br>WP_340419488.1<br>WP_201828904.1<br>WP_284836029.1<br>WP_144268805.1<br>WP_005325988.1<br>WP_269946652.1<br>WP_312716258.1<br>WP_239179689.1<br>WP_115020873.1 | 1<br>2<br>3<br>4 | ADELFSILMGDDVSAR<br>DSMFQAILPLR<br>ISELYIVEGDSAGGSAK<br>RVDLEDAQR | 653-668<br>481-491<br>461-477<br>644-652 | 76131 |
| 8 | Class I SAM-dependent DNA methyltransferase | C. tuberculostearicum<br>C. striatum<br>C. marquesiae<br>C. kefirresidentii<br>C. simulans<br>C. pseudogenitalium<br>C. phoceense<br>C. diphtheriae | WP_250413034.1<br>WP_100087494.1<br>WP_340419467.1<br>WP_248099961.1<br>WP_239240330.1<br>WP_256001363.1<br>WP_257070430.1<br>WP_047939529.1 | 1<br>2<br>3 | DLLGEVYEYFLEK<br>FLETTEKDR<br>NVFWVDPIAR | 183-195<br>245-253<br>94-103 | 61331 |
| 9 | Type I DNA topoisomerase | C. simulans<br>C. accolens<br>C. striatum | WP_061920230.1<br>PCC81749.1<br>WP_049147252.1 | 1<br>2<br>3<br>4<br>5 | ELDENLIDAQETR<br>FGPYVTDGTTNASLR<br>NSQEAHEAIRPAGEK<br>RGDDPATLTDAR<br>SLIDINLEAIDAR | 144-156<br>892-906<br>368-382<br>907-918<br>610-622 | 108384 |
| 10 | DNA primase | C. striatum<br>C. flavescens<br>C. macginleyi<br>C.accolens<br>C. tuberculostearicum<br>C. marquesiae<br>C. kefirresidentii<br>C. aurimucosum<br>C. simulans | WP_201807366.1<br>WP_301504448.1<br>WP_200441644.1<br>WP_302527295.1<br>WP_301714600.1<br>WP_284810591.1<br>WP_284835711.1<br>WP_049361438.1<br>WP_339019257.1 | 1<br>2 | HIATDQVVTVLR<br>SFFEDIASALDAEGIESR | 84-95<br>2-19 | 31510 |
| 11 | DNA-binding protein WhiA; sporulation protein | C. diphtheriae | WP_014306925.1 | 1 | LVDLGPMATRR | 73-83 | 35299 |
| 12 | ATP-dependent DNA helicase | C. falsenii | WP_119664081.1 | 1 | AIRAAAGESSPR | 150-161 | 120947 |

| | | | | | | | |
|---|---|---|---|---|---|---|---|
| 13 | Replicative DNA helicase | C. amycolatum<br>C. vitaeruminis<br>C. ulceribovis<br>C. lactis<br>C. endometrii<br>C. phoceense<br>C. massiliense<br>C. uterequi<br>C. phocae<br>C. lizhenjunii<br>C. minutissimum<br>C. simulans<br>C. striatum<br>C. macginleyei<br>C. tuberculostearicum<br>C. curieae<br>C. kefirresidentii<br>C. flavescens | WP_005511988.1<br>WP_048760061.1<br>WP_040428584.1<br>WP_053413105.1<br>WP_210726537.1<br>WP_257052576.1<br>WP_273369625.1<br>WP_047260393.1<br>WP_075736084.1<br>WP_165009871.1<br>WP_039672719.1<br>WP_248092786.1<br>WP_368918751.1<br>WP_121912383.1<br>WP_316993863.1<br>WP_269946706.1<br>WP_086588341.1<br>WP_312714667.1 | 1 | ELEVPLIAISQLNR | 386-399 | 52999 |
| 14 | Single-stranded DNA-binding protein | C. striatum<br>C. simulans<br>C. aurimucosum<br>C. minutissimum<br>C. accolens | WP_272707505.1<br>WP_248092787.1<br>WP_158381945.1<br>WP_039672723.1<br>WP_284902446.1 | 1 | QAAENVAETLTK | 63-74 | 20725 |
| 15 | DNA repair helicase XPB | Mycobacterium tuberculosis<br>Tsukamurella paurometabola | WP_348790221.1<br>WP_013125486.1 | 1 | EALFDAFR | 446-453 | 61287 |
| 16 | ATP-dependent DNA helicase | C. glyciniphilum | WP_052539681.1 | 1 | AVAQEDQAAR | 160-169 | 120455 |
| 17 | DNA-binding protein WhiA | C. glutamicum | WP_074508156.1 | 1 | MGAQKTR | 199-205 | 35618 |
| 18 | DNA topoisomerase IB | Clavibacter michiganensis | WP_045529628.1 | 1 | LASGETELR | 309-317 | 36094 |
| 19 | ATP-dependent DNA helicase | C. vitaeruminis | WP_313286629.1 | 1 | EAADAEGRNGFMEVAATHAALLMAQGAGRLLRR | 581-613 | 71575 |
| 20 | ATP-dependent DNA helicase<br><br><br><br><br><br>Anthranilate synthase component I family protein | C. glyciniphilum<br>C. imitans<br>C. riegelii<br>C. cystitides<br>C. glaucum<br>C. deserti<br><br>C. jeikeium<br>C. macclintockiae | WP_038549871.1<br>WP_239244666.1<br>WP_239244666.1<br>WP_257162585.1<br>WP_290184939.1<br>WP_053546101.1<br><br>WP_231913344.1<br>WP_269955237.1 | 1 | YGAFLR | 664-669 | 73471 |
| 21 | DNA primase | C. diphtheriae | WP_014302173.1 | 1 | AIVVAGGCSDAIR | 522-534 | 71056 |
| 22 | DNA primase | C. matruchotii | WP_005525931.1 | 1 | NIASSRQCVVVEGYTDVMAMHAAGVATAVATCGT AFGDDHLQMIR | 257-301 | 70564 |
| 23 | Single-stranded DNA-binding protein | C. aurimucosum | WP_010191118.1 | 1 | AGGTEYGPTKFIK | 44-56 | 17019 |
| 24 | SAM-dependent DNA methyltransferase | C. glyciniphilum<br>C. nuruki | WP_038550587.1<br>WP_010121668.1 | 1 | AETRQGR | 85-91 | 104554 |
| 25 | DNA polymerase III subunit gamma and tau | C. accolens<br>C. macginleyi<br>C. tuberculostearicum | WP_284640050.1<br>WP_200450210.1<br>WP_301714980.1 | 1 | TAGFDAPR | 802-809 | 89253 |

| | | | | | | | |
|---|---|---|---|---|---|---|---|
| | | C. aurimucosum<br>C. pseudogenitalium<br>C. marquesiae | WP_049358863.1<br>WP_005323163.1<br>WP_284836698.1 | | | | |
| 26 | YebC/PmpR family DNA-binding transcriptional regulator | C. falsenii | WP_025402776.1 | 1 | GDLTEDDLLMAVLDAGAEEVNDLGEK | 146-171 | 26824 |
| 27 | DNA polymerase III subunit alpha | C. aurimucosum<br>C. minutissimum<br>C. guaraldiae<br>C. hesseae<br>C. striatum<br>C. simulans<br>C. accolens<br>C. camporealensis<br>C. stationis | WP_193633973.1<br>WP_039676049.1<br>WP_154761809.1<br>WP_101735735.1<br>WP_166684572.1<br>WP_062043992.1<br>PCC82148.1<br>WP_046453447.1<br>WP_278821617.1 | 1 | LKQVLVNNK | 1137-1145 | 132204 |
| 28 | DNA repair protein RecO | Clavibacter michiganensis | WP_094130231.1 | 1 | SLGLVDRSDAR | 234-244 | 32836 |
| 29 | Recombinase family protein; mismatch-specific DNA-glycosylase | C. kutscheri<br>C. argentoratensis<br>C. nuruki | WP_126316755.1<br>WP_052331889.1<br>WP_010120108.1 | 1 | AGARPTK | 170-176 | 14257 |
| 30 | NAD-dependent DNA ligase LigA | C. bovis<br>C. urealyticum<br>C. dentalis<br>C. glyciniphilum<br>C. terpenotabidum<br>C. jeikeium<br>C. variabile<br>C. macclintockiae | WP_125186080.1<br>WP_012359981.1<br>WP_099297251.1<br>WP_304034101.1<br>AGP31105.1<br>WP_011273805.1<br>WP_141327709.1<br>WP_274706402.1 | 1 | MFANPRNAAAGAMR | 217-230 | 80598 |
| 31 | DNA methyltransferase | C. maris | WP_020935841.1 | 1 | VLKNERER | 927-934 | 106292 |
| 32 | DNA translocase FtsK | C. efficiens<br>C. gallinarum | WP_231295015.1<br>WP_225213312.1 | 1 | SGEIETPLGSK | 742-752 | 100974 |
| 33 | Replicative DNA helicase | C. variabile | WP_141657038.1 | 1 | MDDGQWDKLTR | 345-355 | 58064 |
| 34 | NAD-dependent DNA ligase LigA | C. bovis<br>C. urealyticum<br>C. dentalis<br>C. glyciniphilum<br>C. jeikeium<br>C. variabile<br>C. macclintockiae<br>C. falsenii | WP_125172725.1<br>WP_012359981.1<br>WP_099297251.1<br>WP_304034101.1<br>WP_011273805.1<br>WP_141327709.1<br>WP_274706402.1<br>WP_276784333.1 | 1 | MFANPRNAAAGAMR | 217-230 | 80598 |
| 35 | Bifunctional DNA-formamidopyrimidine glycosylase/DNA-(apurinic or apyrimidinic site) lyase | C. aurimucosum | WP_010190448.1 | 1 | GTHYCPSCQNRGVGR | 263-277 | 30701 |
| 36 | Single-stranded DNA-binding protein | Prescottella equi | WP_238840838.1 | 1 | TRGGATDGSEGEAPSESEGR | 122-141 | 17205 |
| 37 | DNA-directed RNA polymerase subunit beta | C. pseudodiphtheriticum | WP_272696996.1 | 1 | EMQSPEGQHLMLDTDDIDHFGNR | 342-364 | 128433 |
| 38 | ATP-dependent DNA helicase | Clavibacter michiganensis | WP_104280092.1 | 1 | VMVIEDADR | 112-120 | 44554 |

| | | | | | | | |
|---|---|---|---|---|---|---|---|
| | RecG | | | | | | |
| 39 | ATP-dependent DNA helicase | C. glyciniphilum<br>C. imitans<br>C. riegelii<br>C. cystitides<br>C. deserti<br>C. jeikeium<br>C. macclintockiae | WP_038549871.1<br>WP_239244666.1<br>WP_311357376.1<br>WP_257162585.1<br>WP_053546101.1<br>WP_231913344.1<br>WP_269955237.1 | 1 | YGAFLR | 664-669 | 73471 |
| 40 | Class I SAM-dependent DNA methyltransferase | C. glutamicum | WP_216312896.1 | 1 | EWKTTEDATQQSGPLTR | 39-55 | 171706 |
| 41 | ATP-dependent DNA ligase | Prescottella equi | WP_037084524.1 | 1 | DDKSPNDVRIE | 736-746 | 81072 |
| 42 | DNA helicase PcrA | C. tuberculostearicum<br>C. kefirresidentii<br>C. pseudogenitalium | WP_316991966.1<br>WP_284835665.1<br>WP_005323803.1 | 1 | SYGRDWSGWGSGGSR | 743-747 | 92066 |
| 43 | DNA polymerase III subunit delta' | C. falsenii | WP_224209184.1 | 1 | DAMMQAVGAVDDDGAPIGGYQHPDMAATSR | 327-356 | 43643 |
| 44 | Translation initiation factor IF-3 | C. argentoratense<br>C. stationis<br>C. striatum | WP_234858154.1<br>WP_168970467.1<br>HAT1276202.1 | 1 | LVGPGGEQVGIVR | 25-37 | 21353 |
| 45 | Energy-dependent translational throttle protein EttA | C. striatum<br>C. simulans | WP_110378799.1<br>WP_239239657.1 | 1<br>2<br>3<br>4 | CPPSDAPVNNLSGGER<br>EGGNLILLDEPTNDLDVETLGSLENALEK<br>LLLSEPDLLLLDEPTNHLDAESVEWLEK<br>LSYVDQGR | 150-165<br>459-487<br>173-200<br>386-393 | 62020 |
| 46 | Energy-dependent translational throttle protein EttA | C. afermentans<br>C. cystitides<br>C. fournieri<br>C. auris<br>C. urinipleomorphum<br>C. sanguinis<br>C. urealyticum<br>C. appendicis<br>C. simulans | WP_228496298.1<br>WP_257162271.1<br>WP_085956736.1<br>WP_290341692.1<br>WP_087117630.1<br>WP_136652864.1<br>WP_148812377.1<br>WP_076599618.1<br>WP_239239657.1 | 1 | YEEMAAEAEQYR | 291-302 | 62350 |
| 47 | Translation initiation factor IF-2 | C. simulans<br>C. striatum | WP_248092216.1<br>WP_086891847.1 | 1<br>2<br>3<br>4<br>5 | AEAKEDSFYAALETALAK<br>EDSFYAALETALAK<br>NVEVPDHFQER<br>SQPTNAQVTITGR<br>YIDPYAETVEDVRPGQIVR | 75-92<br>79-92<br>15-25<br>2-14<br>138-156 | 24329 |
| 48 | Translation initiation factor IF-2 | C. tuberculostearicum<br>C. kefirresidentii<br>C. aurimucosum<br>C. marquesiae | WP_316986386.1<br>WP_239204207.1<br>MCG7446990.1<br>WP_269953074.1 | 1 | KPAKGAAK | 68-75 | 95635 |
| 49 | Translation initiation factor IF-2 N-terminal domain-containing protein | C. amycolatum<br>C. vitaeruminis | WP_275435322.1<br>WP_048761261.1 | 1 | KIEHALK | 618-624 | 150280 |
| 50 | Translational GTPase TypA | C. jeikeium | WP_011273863.1 | 1 | MESMDNTGTGWVR | 454-466 | 69160 |
| 51 | Transcription termination/antitermination protein NusG | C. simulans<br>C. striatum | WP_248093691.1<br>WP_086892101.1 | 1<br>2<br>3 | AQTLEVEDSIFEVVVPVEQILENKDGK<br>ETPGVTSFVGNEGNATPVK<br>MELNDAAWSVVR | 109-135<br>164-182<br>152-163 | 30583 |
