## Supplemental Table S5 for "Peptidome Analysis of Western Blots Identifies Natural Bispecific Antibody-Bound *Corynebacterium* and Phage B-cell Epitopes with Potential Relevance to Psoriasis"

### Table S5. RNA-directed Transcription-Translation Enzymes are *Corynebacterium* Antigens.

| Antigen Number | Identified Antigen | Corynebacterium, Actinomycetota | Accession Number | Peptide Sequence | Peptide Number | Peptide Start/Stop | Molecular Weight |
|---|---|---|---|---|---|---|---|
| 1 | tRNA (adenine-N1)-methyltransferase | C. striatum<br>C. simulans<br>C. aurimucosum | WP_284765339.1<br>WP_062040796.1<br>WP_102233936.1 | STLGSDFLLFR<br>VILDMLEPWEHLEK | 56-66<br>177-190 | 31468 | 31468 |
| 2 | DEAD/DEAH box RNA helicase | Gordonia malaquae | WP_336791511.1 | VDDLSKR | 127-133 | 17493 | 17493 |
| 3 | tRNA (N6-isopentenyl adenosine(37)-C2)-methylthiotransferase MiaB | C. striatum<br>C. minutissimum<br>C. hiratae<br>C. aurimucosum<br>C. guaraldiae<br>C. faecipullorum<br>C. hesseae<br>C. simulans | WP_368917809.1<br>WP_082013940.1<br>WP_158396538.1<br>WP_227486157.1<br>WP_154737008.1<br>HIX80213.1<br>WP_269948084.1<br>WP_345791625.1 | LVALQDSIQAEENAK | 400-415 | 56262 | 56262 |
| 4 | tRNA (guanosine(46)-N7)-methyltransferase TrmB | C. pseudodiphtheriticum<br>C. propinquum<br>C. camporealensis<br>C. striatum<br>C. macginleyi<br>C. kefirresidentii<br>C. marquesiae<br>C. simulans<br>C. hesseae<br>C. guaraldiae<br>C. minutissimum<br>C. aurimucosum<br>C. faecipullorum<br>C. hiratae | WP_272722424.1<br>WP_284790565.1<br>WP_046453580.1<br>WP_100087774.1<br>WP_121912209.1<br>WP_248099996.1<br>WP_408927422.1<br>WP_062036873.1<br>WP_339016979.1<br>WP_143336110.1<br>WP_115023857.1<br>WP_158381442.1<br>HIX79839.1<br>WP_144013355.1 | MFAPESLDGIR | 148-158 | 29940 | 29940 |
| 5 | 16S rRNA (cytosine(1402)-N(4))-methyltransferase RsmH | C. pseudotuberculosis | WP_014800655.1 | MAELIAAPVIK | 27-37 | 38580 | 38580 |
| 6 | tRNA dihydrouridine synthase DusB | C. glucuronolyticum | WP_276542633.1 | GVSGTDIPVTVK | 141-152 | 43656 | 43656 |
| 7 | tRNA adenosine deaminase-associated protein | C. striatum<br>C. simulans | WP_086892281.1<br>WP_239240418.1 | MLISDATYADEDDFAADFLELR<br>NEGQWVVR | 70-91<br>15-22 | 17808 | 17808 |
| 8 | 16S rRNA (cytosine(1402)-N(4))-methyltransferase RsmH<br>*iron ABC transporter permease* | C. otitides<br>*C. casei* | WP_046644458.1<br>*WP_006822721.1* | LAHFGGR | 145-151 | 35688 | 35688 |
| 9 | tRNA guanosine(34) transglycosylase Tgt | C. diphtheriticum<br>C. striatum<br>C. simulans | WP_284584383.1<br>MDU3174759.1<br>WP_284841675.1 | AYQEESVAR | 186-194 | 46647 | 46647 |
| 10 | RNA methyltransferase | C. striatum<br>C. simulans<br>C. endometrii | WP_049159484.1<br>WP_239240721.1<br>WP_168707160.1 | NAAGMDVDAVLLGSGAADPLYR | 132-153 | 29261 | 29261 |
| 11 | RNase adapter RapZ | C. striatum<br>C. tuberculostearicum<br>C. kefirresidentii<br>C. accolens<br>C. yonathiae<br>C. camporealensis<br>C. marquesiae<br>C. flavescens<br>C. aurimucosum<br>C. curieae | WP_272706836.1<br>WP_316987251.1<br>WP_208612441.1<br>WP_284897590.1<br>WP_288794165.1<br>WP_035104714.1<br>WP_284836339.1<br>WP_174775506.1<br>WP_193387598.1<br>WP_269945266.1 | KVAAVTDVR | 61-69 | 32896 | 32896 |

| | | | | | | | |
|---|---|---|---|---|---|---|---|
| | | C. macginleyi<br>C. simulans | WP_121910719.1<br>WP_239238967.1 | | | | |
| 12 | ATP-dependent RNA helicase HrpA | C. tuberculostearicum<br>C.kefirresidentii | WKS54074.1<br>WP_259824647.1 | LPMHLR | 963-968 | | |
| 13 | RNA-binding S4 domain-containing protein | C. striatum | WP_306497072.1 | VAAADDDDDDYFDEATADDDFDPEKWR | 67-93 | 10127 | 10127 |
| 14 | tRNA (adenosine(37)-N6)-threonylcarbamoyltransferase complex ATPase subunit type 1 TsaE | C. diphtheria<br>C. confusum<br>C. phceense<br>C. callunae<br>C. striatum | CAB0550252.1<br>WP_290224813.1<br>WP_257046221.1<br>WP_247776017.1<br>WP_272707442.1 | VTSPTFVIAR | 63-72 | 17748 | 17748 |
| 15 | tRNA (adenine-N1)-methyltransferase | C. simulans<br>C. striatum<br>C. aurimucosum | WP_062040796.1<br>WP_086891396.1<br>WP_102233936.1 | LGDLADVTVEELGGPVDR | 159-176 | 31392 | 31392 |
| 16 | RNA methyltransferase | C. striatum<br>C. simulans<br>C. camporealensis<br>C. flavescens | WP_166684518.1<br>WP_096337125.1<br>WP_321113870.1<br>WP_301505940.1 | NAAGMDVDAVLLGSGAADPLYR | 132-153 | 29261 | 29261 |
| 17 | 16S rRNA (guanine(966)-N(2))-methyltransferase RsmD | C. striatum<br>C. guaraldiae<br>C. aurimucosum<br>C. hesseae<br>C. faecipullorum | WP_086891473.1<br>WP_144269098.1<br>WP_158381122.1<br>WP_269948218.1<br>HIX78688.1 | EGLFSSLNVR | 30-39 | 21908 | 21908 |
| 18 | tRNA (adenine-N1)-methyltransferase | C. striatum<br>C. simulans | WP_284765339.1<br>WP_062040796.1 | STLGSDFLLFR | 56-66 | 31468 | 31468 |
| 19 | CCA tRNA nucleotidyltransferase | C. pseudodiphtheriticum<br>C. propinquum | WP_284862401.1<br>WP_284603917.1 | MNSSDTQQIALLARATEAVR | 1-20 | 54284 | 54284 |
| 20 | RNA 2',3'-cyclic phosphodiesterase | C. halotolerans | WP_027004290.1 | YETVSEIR | 170-177 | 20297 | 20297 |
| 21 | tRNA dihydrouridine synthase DusB | C. striatum | WP_114976561.1 | VESVDNLR | 319-326 | 41238 | 41238 |
| 22 | ATP-dependent RNA helicase DeaD | Clavibacter michiganensis subsp. michiganensis | OUD99502.1 | GERRARPAR | 477-485 | 64092 | 64092 |
| 23 | RNA pseudouridine synthase B | C. pseudotuberculosis | AEK92306.1 | GSEGSRAGENTRESYGFR | 65-82 | 41826 | 41826 |
| 24 | 16S rRNA (uracil(1498)-N(3))-methyltransferase | C. variabile | WP_313007511.1 | WDGKPGKAEKAR | 119-130 | 26362 | 26362 |
| 25 | tRNA lysidine(34) synthetase TilS | C. casei | WP_098072589.1 | DSPAGMEAEAR | 126-136 | 39218 | 39218 |
| 26 | ATP-dependent RNA helicase HrpA | C. genitalium | WP_040423649.1 | TATATDDHDDSRDRDPK | 621-637 | 147368 | 147368 |
