## Supplemental Table S6 for "Peptidome Analysis of Western Blots Identifies Natural Bispecific Antibody-Bound *Corynebacterium* and Phage B-cell Epitopes with Potential Relevance to Psoriasis"

### Table S6. Aminoacyl-tRNA-Ligases are *Corynebacterium* Antigens.

| Antigen Number | Identified Antigen | *Corynebacterium* (C.), *Mycobacterium* (M.) *tuberculosis* (Peptide numbers) | Accession Number | Peptide Number | Peptide Sequence | Peptide Start/Stop | Molecular Weight |
|---|---|---|---|---|---|---|---|
| 1 | Alanine--tRNA ligase | C. striatum (1,2) | WP_306496304.1 | 1 | GNFEILGPLPK | 217-227 | 97495 |
| | | C. simulans (1) | WP_239238690.1 | 2 | LFDEAVEALK | 378-387 | |
| | | C. accolens (1,3) | PCC82825.1 | 3 | LGMEDNYWSMGVPGPCGPCSEIYYDR | 155-180 | |
| | | C. aurimucosum (1) | WP_158381258.1 | | | | |
| | | C. lizhenjunii (1,3) | WP_165008249.1 | | | | |
| | | C. phocae (1) | WP_075733917.1 | | | | |
| | | C. hiratae (1) | WP_328287743.1 | | | | |
| | | C. oculi (1) | WP_055121813.1 | | | | |
| | | C. camporealensis (3) | WP_321113903.1 | | | | |
| | | C. segmentosum (3) | WP_126319135.1 | | | | |
| 2 | Arginine--tRNA ligase | C. striatum (1,2,3,4,5,6,7,8,9,10,12) | WP_005532004.1 | 1 | AGEETQPIHIAR | 516-527 | 60193 |
| | | C. simulans (2,3,4,5,6,7,8,10,12) | WP_239238812.1 | 2 | AGTVITLDDLVEAIGVDGAR | 379-398 | |
| | | C. lizhenjunii (5,9) | WP_165241798.1 | 3 | AVAHLKDNEK | 267-276 | |
| | | C. phoceense (5,7) | WP_257070704.1 | 4 | ETAVAVLSEHELDTAVLPETVTVERPR | 12-38 | |
| | | C. lactis (5) | WP_053411866.1 | 5 | FSNSLVAAAK | 177-186 | |
| | | C. vitaeruminis (5) | WP_025252568.1 | 6 | INLEFVSANPTGPIHLGGTR | 123-142 | |
| | | C. singular (2,10) | WP_239180679.1 | 7 | LYFEDGAWWLK | 277-287 | |
| | | C. aurimucosum (2) | WP_395419686.1 | 8 | MNPVDLSVLIK | 1-11 | |
| | | C. minutissimum (2) | WP_039674952.1 | 9 | QVLANALTMVGVSAPEKM | 536-553 | |
| | | C. urealyticum (2) | PZP00401.1 | 10 | SDGDAAYIAGDIAYVADKFDR | 302-322 | |
| | | C. faecipullorum (2) | HIX79552.1 | 11 | TLGEFPEVVSTAATLR | 472-487 | |
| | | C. guaraldiae (2) | WP_154737118.1 | 12 | WAAVGDSLGR | 143-152 | |
| | | C. hiratae (2) | WP_158395294.1 | | | | |
| | | C. bovis (6) | WP_125173180.1 | | | | |
| | | C. kroppenstedtii (6) | ACR18037.1 | | | | |
| | | C. jeikeium (6) | WP_034985617.1 | | | | |
| | | C. genitalium (6,10) | EFK54197.1 | | | | |
| | | C. fourneri (6,10) | WJY97349.1 | | | | |
| | | C. afermentas (6) | WJY56534.1 | | | | |
| | | C. marinum (6) | AJK68599.1 | | | | |
| | | C. urealyticum (6) | AGE36275.1 | | | | |
| | | C. glyciniphilum (6) | WP_145941540.1 | | | | |
| | | C. faecigallinarum (6) | HJC85263.1 | | | | |
| | | C. camporealensis (6) | AKE38939.1 | | | | |
| | | C. glaucum (6,10) | AQQ15018.1 | | | | |
| | | C. variabile (6) | WP_301522345.1 | | | | |
| | | C. flavescens (6) | GEB96627.1 | | | | |
| | | C. terpenotabidum (6) | WP_020441500.1 | | | | |
| | | C. argentoratense (6) | WP_314929684.1 | | | | |
| | | C. macclintockii (6) | WP_284819915.1 | | | | |
| | | C. falsenii (6) | WP_272713644.1 | | | | |
| | | C. accolens (7,10) | PCC82930.1 | | | | |
| | | C. appendicis (10) | WP_076598374.1 | | | | |

| | | | | | | | |
|---|---|---|---|---|---|---|---|
| | | C. diphtheria (10) | WP_010934757.1 | | | | |
| | | C. aurimucosum (12) | TVU57667.1 | | | | |
| 3 | Aspartate--tRNA ligase | C. aurimucosum (1-14,16,17,18,19) | WP_010190145.1 | 1 | GLAYITIAEDGTLGGPVAK | 346-364 | 66879 |
| | | C. simulans (1-20) | WP_062040573.1 | 2 | IVECTEFFK | 296-304 | |
| | | C. striatum (1-19) | WP_269152651.1 | 3 | IVECTEFFKDTTFR | 296-309 | |
| | | C. minutissimum (1-12,14,16,17, 18) | WP_239188640.1 | 4 | IVSLLGGFDSIR | 543-554 | |
| | | C. tuberculostearicum (19,11,12,13,14,16-19) | WP_302257823.1 | 5 | MTYADAMK | 274-281 | |
| | | C. kefirresidentii (1-5,7,9,11,12,13,14,16) | WP_394287099.1 | 6 | QFDAWQEWAK | 331-340 | |
| | | C. glucuronolyticum (1-4,7,9,14) | WP_201841571.1 | 7 | QLLMVAGMER | 203-212 | |
| | | C. marquesiae (1-5,9,11,14,16,19) | WP_284838861.1 | 8 | RQFDAWQEWAK | 330-340 | |
| | | C. curiae (1-5,7,9,12,14,16,19) | WP_269945302.1 | 9 | THLAGELR | 4-11 | |
| | | C. yonathiae (1-9,11,12,14,16-19) | WP_288794186.1 | 10 | VFDVMGIGEEEAQEK | 504-518 | |
| | | C. confusum (1,2,3,4,7,12,14,16,20) | WP_290221938.1 | 11 | VFQNEYVGAVVMEGGASQPR | 310-329 | |
| | | C. macginleyi (1-5,9,10,12,16,19) | WP_121910766.1 | 12 | YYGSDKPDLR | 282-291 | |
| | | C. accolens (1-3,5,6,8-10,12,13,15,16,19,20) | WP_284637310.1 | 13 | IHQQDVQKR | 490-498 | |
| | | C. faecale (4,14,19) | WJY92387.1 | 14 | IVSLLGGFDSIR | 538-549 | |
| | | C. glaucum (4,7,14) | WP_290187075.1 | 15 | LIGYEITTPIPR | 257-268 | |
| | | C. genitalium (4,7,11,14) | EFK53849.1 | 16 | MTYADAMK | 269-276 | |
| | | C. casei (4,6-9,11,12,14,15,17, 18,20) | WP_193024854.1 | 17 | QFDAWQEWAK | 326-335 | |
| | | C. ammoniagenes (4-9,11,14,15,16,17, 18,20) | EFG82285.1 | 18 | RQFDAWQEWAK | 325-335 | |
| | | C. efficiens (4,14) | EEW49631.1 | 19 | SGIAQVVFR | 38-46 | |
| | | C. falsenii (4,7,9,14) | WP_025402759.1 | 20 | SAALPFQIDDPSTSGEVGEEAR | 119-140 | |
| | | C. appendicis (4,11) | WP_259798215.1 | 21 | SGGGVDPLTDAPAPIPTAQR | 561-580 | |
| | | C. frankenforstense (4,7,9,14) | WP_304324741.1 | | | | |
| | | C. occultum (4,14,15) | WP_156231015.1 | | | | |
| | | C. bovis (4,7,9,14) | WP_342003955.1 | | | | |
| | | C. stationis (5,6,8,15,16,17, 18) | WP_278755516.1 | | | | |
| | | C. guaraldiae (5,6,8,10,12,16,17, 18) | WP_143335995.1 | | | | |
| | | C. singular (5,6,8,10,15,16,17, 18) | WP_144790055.1 | | | | |
| | | C. hindlerae (5,12,16,19) | WP_208767412.1 | | | | |
| | | C. resistens (7,11) | WP_193024854.1 | | | | |
| | | C. cystitidis (7) | WP_257158583.1 | | | | |
| | | C. dentalis (7,9) | WP_288800877.1 | | | | |
| | | C. propinquum (7) | RUP80582.1 | | | | |
| | | C. imitans (7) | SNV73621.1 | | | | |
| | | C. amycolatum (7) | WP_284827000.1 | | | | |
| | | C. diphtheriae (7) | WP_046095240.1 | | | | |
| | | C. pseudotuberculosis (7,11) | WP_014523357.1 | | | | |
| | | M. tuberculosis (7) | WP_010950740.1 | | | | |
| | | M. leprae (7) | WP_162618225.1 | | | | |
| | | C. phocae (10,20) | WP_075733915.1 | | | | |
| | | C. lizhenjunii (10,20) | WP_165008255.1 | | | | |
| | | C. faecipullorum (10) | HIX79070.1 | | | | |
| | | C. endometrii (14, 15,19) | WP_136141263.1 | | | | |
| | | C. humirecucens (15) | WP_040085950.1 | | | | |
| | | C. camporealensis (15,20) | WP_035104785.1 | | | | |
| | | C. auris (15) | WJY68165.1 | | | | |
| | | C. phoceense (15,20) | WP_257045095.1 | | | | |
| | | C. incognita (15) | WP_185176018.1 | | | | |

| | | | | | | | |
|---|---|---|---|---|---|---|---|
| | | C. renale (19)<br>C. riegelii (19)<br>C. ulcerans (11)<br>C. sanguinis (21)<br>C. lipophiloflavum (21) | WP_115243631.1<br>WP_311356469.1<br>AKA96776.1<br>WP_259812518.1<br>WP_136651265.1 | | | | |
| 4 | Cysteine--tRNA ligase | C. phoceense (1,2,4,5)<br>C. simulans (1,2,3,4,5)<br>C. striatum (1,2,3,4,5)<br>C. accolens (1,2,3,4,5)<br>C. flavescens (4)<br>C. aurimucosum (2,4)<br>C. guaraldiae (2,4)<br>C. argentoratense (6)<br>C. variabile (1)<br>C. nuruki (1)<br>C. singulare (1,4)<br>C. kroppenstedtii (1)<br>C. pseudokroppenstedtii (1)<br>C. parakroppenstedii (1)<br>C. minutissimum (1,2)<br>C. faecipullorum (1)<br>C. endometrii (2)<br>C. appendicis (2)<br>C. urinipleomorphum (2)<br>C. efficiens (2,3)<br>C. ammoniagenes (2)<br>C. frankenforstense (4)<br>C. camporealense (3)<br>C. glutamicum (3)<br>C. tuberculostearicum (3)<br>C. deserti (3)<br>C. breve (3)<br>C. callunae (3)<br>C. massiliense (5)<br>C. pseudodiphtheriticum (5)<br>C. propinquum (5)<br>C. urogenitale (5)<br>C. macginleyi (5)<br>C. jeikeium (5)<br>C. lipophiloflavum (5)<br>C. diphtheriae (5)<br>C. pseudotuberculosis (5)<br>M. tuberculosis (5)<br>M. leprae (5) | WP_257071465.1<br>WP_239240082.1<br>WP_166684765.1<br>WP_302503406.1<br>WP_301504764.1<br>WP_012715309.1<br>WP_144014586.1<br>WP_041747787.1<br>WP_312778126.1<br>WP_273052937.1<br>WP_239181084.1<br>WP_406605552.1<br>WP_204087991.1<br>WP_340055101.1<br>WP_239189044.1<br>HIX78523.1<br>QCB29197.1<br>WP_076598968.1<br>WP_087117512.1<br>EEW50971.1<br>EFG81287.1 (2)<br>WP_075664533.1<br>WP_105360408.1<br>WP_011015271.1<br>WP_302258629.1<br>WP_053545652.1<br>WP_284824365.1<br>WP_247776184.1<br>WP_022863085.1<br>WP_284860691.1<br>WP_239211183.1<br>WP_201738916.1<br>WP_200437520.1<br>EEW16681.1<br>EEI17911.1<br>WP_196971765.1<br>WP_014401374.1<br>WP_044090845.1<br>WP_010907674.1 | 1<br>2<br>3<br>4<br>5<br>6 | AEKDWATADAVR<br>AYNTLGVLPPSVEPR<br>GPQDFALWK<br>SVLEYSEAALTEAAAGYR<br>YYLGSAHYR<br>SLGNVLSVENITKIVRPVELRYYLGSAHYRS<br>MIEYSEGALLEAAAGYR | 444-455<br>104-118<br>196-204<br>318-335<br>309-317<br>267-314 | 52527 |
| 5 | Glutamate--tRNA ligase | C. striatum (1-11,13-31)<br>C. accolens (1-6,8-23,25-27,29-32)<br>C. simulans (1-11,13-26,28-31)<br>C. incognita (2,11,23)<br>C. pseudogenitalium (2,4,10) | WP_172454484.1<br>PCC81867.1<br>WP_239238905.1<br>WP_185175823.1<br>WP_256008513.1 | 1<br>2<br>3<br>4<br>5 | AALAVTPFQAPAQQ<br>ALIEELELKPR<br>DPQSNLFNHR<br>DSEESYQAIIDSLK<br>EDAVQPLEVSIAR | 489-502<br>442-452<br>266-275<br>57-70<br>411-423 | 56685 |

| | | | | | | | |
|---|---|---|---|---|---|---|---|
| | | C. renale (2,23) | WP_111725945.1 | 6 | EGGYVYPAYSTAEEVEER | 107-124 | |
| | | C. maris (2) | WP_020934626.1 | 7 | FAFVADLVQTR | 367-377 | |
| | | C. breve (2) | WP_284826405.1 | 8 | FCPSPTGTPHVGMVR | 15-29 | |
| | | C. tuberculostearicum (3,8,9,12,27,28,29) | EET77800.1 | 9 | GEDLLSSTPR | 219-228 | |
| | | C. singular (3,8,9,11,12,25-28,30) | WP_391498207.1 | 10 | IEDTDAARDSEESYQAIIDSLK | 49-70 | |
| | | C. endometrii (3,8,9,11,12,23,27,29) | QCB28314.1 | 11 | KAYGALR | 453-459 | |
| | | C. phoceense (3,8,9,11,12,27,28,29) | WP_257055141.1 | 12 | KLEAINADHIR | 327-337 | |
| | | C. guaraldiae (3,25,26,27,28) | WP_144014399.1 | 13 | LEELEEWKTDAIESVLSK | 424-441 | |
| | | C. hiratae (3,5,27,28,30) | WP_144013560.1 | 14 | LGYDNYDRDLTDEQIAAFEAEGR | 134-156 | |
| | | C. aurimucosum (3,5,25,26,27,28,30) | WP_080975116.1 | 15 | LKEGGYVYPAYSTAEEVEER | 105-124 | |
| | | C. hesseae (3,5,25,26,27,28) | WP_339017493.1 | 16 | LLEPEVFAQR | 338-347 | |
| | | C. jeikeium (4,5,11,19,29) | WP_071056913.1 | 17 | LRDYLTEYTDFPADYDEQK | 348-366 | |
| | | C. macclintockiae (4,19) | WP_269954417.1 | 18 | LRMPEQDWTWK | 162-172 | |
| | | C. diphtheria (4,10,25,26,27) | WP_010934815.1 | 19 | MDIYADVLEK | 95-104 | |
| | | C. belfantii (4,10) | WP_374071733.1 | 20 | MLGDAYGLMSFLVTPDAELELDEK | 380-403 | |
| | | C. rouxii (4,10) | WP_342330779.1 | 21 | MPEQDWTWK | 164-172 | |
| | | C. minutissimum (5,25,26,30) | WP_239188017.1 | 22 | NLKEDAVQPLEVSIAR | 408-423 | |
| | | C. auris (8,9,12,28) | WJY67900.1 | 23 | QLALYEALK | 229-237 | |
| | | C. riegelii (8,9,12,23) | GAA0199541.1 | 24 | QLALYEALKELGIAK | 229-243 | |
| | | C. sanguinis (8,9,12,23) | WP_259810097.1 | 25 | QTPEFGHLPFVMGEGNK | 244-260 | |
| | | C. glaucum (8,9,12,) | WJZ07612.1 | 26 | QTPEFGHLPFVMGEGNKK | 244-261 | |
| | | C. lipophiloflavum (8,9,12,) | WP_050758393.1 | 27 | RDPQSNLFNHR | 265-275 | |
| | | C. otitides (9,11,29) | CCI83419.1 | 28 | SETQPDYVVAR | 183-193 | |
| | | C. propinquum (9,11,29) | WP_302500921.1 | 29 | TALFNWAYAR | 30-39 | |
| | | C. falsenii (13,14) | WP_272712839.1 | 30 | VAISGQQVSPPLFESMELLGK | 460-480 | |
| | | C. nuruki (13,14) | WP_010119086.1 | 31 | WLGMDWDEGVVTGGPHEPYR | 71-90 | |
| | | C. camporealensis (19) | WP_321113260.1 | 32 | GSTTMVAMTEVR | 4-15 | |
| | | C. marinum (19) | WP_084602806.1 | | | | |
| | | C. matruchotii (19) | WP_314823046.1 | | | | |
| | | C. glutamicum (23,29) | BCB35081.1 | | | | |
| | | C. pseudotuberculosis (23,25,26,27) | BCB35081.1 | | | | |
| | | C. glucuronolyticum (24) | WP_259815813.1 | | | | |
| | | C. ulceribovis (29) | WP_018023718.1 | | | | |
| | | C. efficiens (29) | BAC18199.1 | | | | |
| | | C. incognita (29) | WP_185175823.1 | | | | |
| | | C. epidermidicanis (31) | WP_047240066.1 | | | | |
| | | C. felinum (31) | WP_277105438.1 | | | | |
| 6 | Glycine--tRNA ligase | C. striatum (1,2,3,4,6,7,8,9,10) | WP_284810341.1 | 1 | ADHLLEAYEEK | 99-109 | 52643 |
| | | C. simulans (1,3,4,7,8,9,10) | WP_248093957.1 | 2 | AFSGLLK | 137-143 | |
| | | C. aurimucosum (2,3,6,7,8,9,10) | WP_193634054.1 | 3 | ASVIDTVVSLCK | 2-13 | |
| | | C. minutissimum (3,7,8,10) | WP_039676344.1 | 4 | EDVVGVDTSVIQPR | 58-71 | |
| | | C. resistens (1,2,5) | AEI09050.1 | 5 | ERDTMEQER | 434-442 | |
| | | C. jeikeium (1,2,5,9,10) | EEW16902.1 | 6 | GLVYQAGEIYGGSR | 16-29 | |
| | | C. variabile (1,10) | WP_313007517.1 | 7 | GSGEDLSYYDQTTEER | 298-313 | |
| | | C. efficiens (1,2,10) | BAC18988.1 | 8 | HGHEPANGLADINDPETGQPGDWTEPK | 110-136 | |
| | | C. deserti (1,2) | WP_053545376.1 | 9 | LYEHPKEK | 249-256 | |
| | | C. mastitides (1,2,5,9,10) | WP_101172968.1 | 10 | NEITPGNFIFR | 194-204 | |
| | | C. callunae (1,9,10) | WP_015651785.1 | 11 | SAWDYGPLGVELK | 30-42 | |
| | | C. durum (1) | WP_315560655.1 | 12 | TDYDLSVHAK | 288-297 | |

| | | | | | | |
|---|---|---|---|---|---|---|
| | | C. kutscheri (1,2,10) | WP_046439067.1 | 13 | TVDVEYAFGFTGSK | 264-277 |
| | | C. occultum (1,2,9,10) | WP_156231350.1 | 14 | VAVLPLSK | 366-373 |
| | | C. pyruviciproducens (1,9,10) | WP_101678393.1 | 15 | VKIDELEAYLAAR | 443-455 |
| | | C. otitides (1,6,9) | WP_004600325.1 | 16 | WGELEGVANR | 278-287 |
| | | C. halotolerans (1,2) | WP_015401488.1 | 17 | WVPYVIEPAAGLGR | 314-327 |
| | | C. glucuronolyticum (1,9) | MCI6205664.1 | 18 | KEELSTPAR | 374-382 |
| | | C. vitaeruminis (1,2,9,10) | WP_034649598.1 | | | |
| | | C. falsenii (1,2,5,9,10) | WP_272713155.1 | | | |
| | | C. gallinarum (1,2) | WP_191732284.1 | | | |
| | | C. cystitides (1) | WP_313007517.1 | | | |
| | | C. nuruki (1,2,9) | WP_010122379.1 | | | |
| | | C. lubricantis (1) | WP_018296043.1 | | | |
| | | C. appendicis (1,5) | WP_273412240.1 | | | |
| | | C. ureicelerivorans (1) | WP_301693509.1 | | | |
| | | C. faecigallinarum (1,9) | HJC86162.1 | | | |
| | | C. afermentans (1,2,5) | WP_286205833.1 | | | |
| | | C. bovis (1,9) | WP_125186994.1 | | | |
| | | C. oculi (2,5) | WP_055121493.1 | | | |
| | | C. glutamicum (2) | WP_006287484.1 | | | |
| | | C. faecale (2) | WP_290276418.1 | | | |
| | | C. frankenforstense (2,9) | WP_075664299.1 | | | |
| | | C. terpenotabidum (2) | WP_020440850.1 | | | |
| | | C. stationis (2,6) | WP_278821587.1 | | | |
| | | C. auriscanis (2) | WP_290183325.1 | | | |
| | | C. accolens (2,6,7,9) | WP_302525054.1 | | | |
| | | C. urogenitale (2) | WP_151902461.1 | | | |
| | | C. suedecumii (2,6) | WP_284874829.1 | | | |
| | | C. marquesiae (2,6,7,9) | WP_340418867.1 | | | |
| | | C. lizhenjunii (2,4,6) | WP_165241699.1 | | | |
| | | C. phoceense (3,6) | WP_257069528.1 | | | |
| | | C. hiratae (3,6,7,8) | WP_257069528.1 | | | |
| | | C. endometrii (3,6) | WP_136141608.1 | | | |
| | | C. hesseae (3,7,8) | WP_136141608.1 | | | |
| | | C. kroppenstedtii (5) | WP_081429389.1 | | | |
| | | C. xerosis (5,10) | WP_120982172.1 | | | |
| | | C. renale (5,10) | WP_111726473.1 | | | |
| | | C. imitans (5) | WP_284784187.1 | | | |
| | | C. pseudokroppenstedtii (5) | WP_337448591.1 | | | |
| | | C. riegelii (5) | WP_276906920.1 | | | |
| | | C. fournieri (5) | WP_085958317.1 | | | |
| | | C. otitidis (6) | WP_004600325.1 | | | |
| | | C. flavescens | WP_276621642.1 | | | |
| | | C. casei | WP_301478747.1 | | | |
| | | C. tuberculostearicum | WP_317000265.1 | | | |
| | | C. pseudodiphtheriticum | WP_284861332.1 | | | |
| | | C. incognita (6) | WP_185176346.1 | | | |
| | | C. sanguinis (9) | WP_259884473.1 | | | |
| | | C. pseudogenitalium | WP_256007991.1 | | | |
| | | C. lipophilum | WP_252931283.1 | | | |

| | | | | | | | |
|---|---|---|---|---|---|---|---|
| | | C. lipophiloflavum (9)<br>C. pseudotuberculosis (10)<br>C. ulcerans (10)<br>C. matruchotii (10)<br>C. belfantii (10)<br>C. ulceribovis (10)<br>C. durum (10)<br>C. canis (10)<br>C. amycolatum (10)<br>C. diphtheria (10)<br>C. glutamicum (10)<br>M. tuberculosis (10)<br>M. leprae (10) | WP_006839838.1<br>WP_072577887.1<br>WP_101515478.1 (<br>WP_278719816.1<br>WP_371891214.1<br>WP_018024503.1<br>WP_315560655.1<br>WP_146323626.1<br>WP_377780882.1<br>WP_235697491.1<br>WP_074495386.1<br>WP_031697458.1<br>WP_010907951.1 | | | | |
| 7 | Isoleucine--tRNA ligase | C. accolens (3)<br>C. camporealensis (1,2, 3,4)<br>C. striatum (1,2,3,4,5)<br>C. phoceense (1,2,3,4,5)<br>C. singular (1,2,4,5)<br>C. lizhenjunii (1,2,4,5)<br>C. minutissimum (1,2,4,5)<br>C. faecipullorum (1,2,4,5)<br>C. aurimucosum (1,2,4,5)<br>C. hesseae (1,2,4,5)<br>C. confusum (1,2,3,4)<br>C. hiratae (1,2,4,5)<br>C. guaraldiae (1,2,4,5)<br>C. intestinale (1,2,4,5)<br>C. flavescens (1,2,3,4)<br>C. massiliense (3)<br>C. endometrii (3)<br>C. simulans (3)<br>C. incognita (3,5)<br>C. stationis (3)<br>C. casei (3)<br>C. pseudodiphtheriticum (4,6)<br>C. propinquum (4)<br>C. otitides (6)<br>C. xerosis (6)<br>C. efficiens (6)<br>C. jeikeium (6)<br>C. epidermicanis (6)<br>C. amycolatum (6)<br>C. macclintockiae (6)<br>C. lactis (6)<br>C. kroppenstedtii (6)<br>C. parakroppenstedtii (6)<br>M. tuberculosis (6)<br>M. leprae (6) | WP_302527128.1<br>WP_105360350.1<br>WP_114976422.1<br>WP_257045480.1<br>WP_042531528.1<br>WP_165010183.1<br>WP_115022870.1<br>HIX79172.1<br>WP_010190515.1<br>WP_269948028.1<br>WP_290226351.1<br>WP_158396074.1<br>WP_143334913.1<br>WP_250224159.1<br>WP_312714816.1<br>WP_022862307.1<br>WP_281276120.1<br>WP_248091406.1<br>WP_246389108.1<br>WP_301205826.1<br>WP_035095843.1<br>ERJ46177.1<br>WP_284603912.1<br>CCI83019.1<br>QGS33676.1<br>BAC18853.1<br>WP_115737483.1<br>AKK03510.1<br>WP_040361169.1<br>WP_284803396.1<br>ALA67631.1<br>HJD69440.1<br>WP_340055196.1<br>MBZ4296137.1<br>WP_010908192.1 | 1<br>2<br>3<br>4<br>5<br>6 | AVASAASSVR<br>AVASAASSVRK<br>SGEPLIYK<br>VPDVLDCWFESGSMPFAQK<br>VVAPLLPHVAEVIYR<br>WFLMSSPILR | 833-842<br>833-843<br>427-434<br>550-568<br>785-799<br>655-664 | 119066 |
| 8 | Leucine--tRNA ligase | C. striatum (1,2,3,4,5,6,7,8,9,10,11) | WP_149721470.1 | 1 | AAVEPIAQLVSPIAPHIAEELWK | 843-865 | 107027 |

| | | | | | | | |
|---|---|---|---|---|---|---|---|
| | | C. simulans (1,2,3,4, 6,7,8,9,10,11)<br>C. accolens (1,2,3,9,11)<br>C. phoceense (1,2)<br>C. pseudogenitalium (4,5)<br>C. endometrii (4,5,6,7,10)<br>C. confusum (4)<br>C. cystitides (5,6,7)<br>C. ulcerans (5,10)<br>C. epididermidicanis (5)<br>C. ramonii (5,10)<br>C. pseudotuberculosis (5)<br>C. spheniscorum (5)<br>C. phocae (5)<br>C. appendicis (5)<br>C. ulceribovis (5)<br>C. vitaeruminis (5)<br>C. nuruki (5)<br>C. terpenotabidum (5)<br>C. pyruviciproducens (5)<br>C. kutscheri (5)<br>C. atypicum (6,7)<br>C. propinquum (6,7)<br>C. pseudodiphtheriticum (6,7)<br>C. uropygiale (6,7)<br>C. faecale (9)<br>C. aurimucosum (8,9)<br>C. hesseae (8)<br>C. minutissimum (8)<br>C. guaraldiae (8,9)<br>C. faecipullorum (8)<br>C. callunae (9)<br>C. flavescens (9) | WP_284841493.1<br>PCC83598.1<br>WP_257036476.1<br>EFQ80722.1<br>WP_136142034.1<br>WP_290223792.1<br>WP_257202530.1<br>WP_095076254.1<br>AKK04310.1<br>AIU33695.1<br>WP_014401472.1<br>WP_092286616.1<br>WP_075736140.1<br>WP_284834948.1<br>WP_018025044.1<br>WP_276651300.1<br>WP_010122511.1<br>WP_020442348.1<br>WP_284840762.1<br>WP_126316509.1<br>WP_038607155.1<br>WP_302531450.1<br>WP_239264978.1<br>WP_236119975.1<br>WP_290277508.1<br>WP_395419763.1<br>WP_101736547.1<br>WP_239187446.1<br>WP_143336349.1<br>HIX79192.1<br>WP_015652373.1<br>WP_312714751.1 | 2<br>3<br>4<br>5<br>6<br>7<br>8<br>9<br>10<br>11<br>12<br>13 | AIAGVRDDYENLR<br>DYEFATEFGLPITEVVSGGDISK<br>FYYNGEEVNQEYGK<br>GVYVPAAEVEEK<br>LLDDLELLDWPEK<br>LLDDLELLDWPEKVK<br>QLGLLGLGHDPR<br>SVASTDPEFYK<br>TTTMANIENMR<br>TYGNSAEGAPR<br>YIDPQNSEEFCSLENER<br>YYSDLTAAEQAAALDEFR | 802-814<br>429-451<br>705-718<br>690-701<br>272-284<br>272-286<br>133-144<br>146-156<br>121-131<br>832-842<br>601-617<br>199-216 | |
| 9 | Lysine--tRNA ligase | C. simulans (1,2,5,6,7,8,9)<br>C. phoceense (2,5,6,7,9)<br>C. accolens (1,5,8,9)<br>C. aurimucosum (1,5)<br>C. striatum (1,2,4,5,6,7,8,9)<br>C. tuberculostearicum (5)<br>C. guaraldiae (1)<br>C. variabile (1,2,10)<br>C. maris (1,2,10)<br>C. ulcerans (3)<br>C. diphtheria (3)<br>C. pseudotuberculosis (3)<br>C. nuruki (2,3)<br>C. falsenii (1,2,3,6)<br>C. jeikeium (1,2,3,6,7,9)<br>C. halotolerans (2,7)<br>C. matruchotii (2,6) | WP_282440143.1<br>WP_068801344.1<br>PCC82344.1<br>WP_158381531.1<br>WP_049191856.1<br>WP_251068499.1<br>WP_154737176.1<br>WP_312778140.1<br>WP_020935643.1<br>WP_216605889.1<br>WP_283718230.1<br>WP_014523605.1<br>WP_010120215.1<br>WP_025402027.1<br>WP_035011004.1<br>WP_027004403.1<br>WP_278720683.1 | 1<br>2<br>3<br>4<br>5<br>6<br>7<br>8<br>9<br>10 | ETVLFPIVKPEK<br>GFELATGYSELVDPVIQR<br>IAPELYLK<br>LLDSGVEAYPVTVDR<br>LVEEIWEVLCEDQLEGPIFVK<br>NFPVETSPLTR<br>RAVVGGIDR<br>RGELSIMATK<br>YNDLIVR<br>DLPEQLK | 522-533<br>449-466<br>258-265<br>35-49<br>400-420<br>421-431<br>266-274<br>154-163<br>191-197<br>20-26 | 59728 |

| | | | | | | | |
|---|---|---|---|---|---|---|---|
| | | C. stationis (2)<br>C. marinum (2)<br>C. hesseae (1)<br>C. auriscanis (2,3)<br>C. resistens (3)<br>C. lactis (3)<br>C. endometrii (3,5,9)<br>C. macclintockiae (2,3,6,7,9)<br>C. dentalis (3)<br>C. propinquum (1,7,9)<br>C. phocae (1,5,8)<br>C. urogenitale (1,2)<br>C. pseudodiphtheriticum (1,7,8)<br>C. incognita (2,6,8,9)<br>C. lizhenjunii (1,6,9)<br>C. felinum (2,6)<br>C. glyciniphilum (1)<br>C. faecigallinarum (1)<br>C. renale (2)<br>C. efficiens (2)<br>C. terpenotabidum (2,10)<br>C. confusum (5,7)<br>C. massiliense (5,7)<br>C. macginleyi (5,6,7)<br>C. kefirresidentii (5)<br>C. curieae (5)<br>C. marquesiae (5)<br>C. yonathiae (5)<br>C. amycolatum (8)<br>C. vitaeruminis (8)<br>C. occultum (8)<br>C. suedekumii (8)<br>C. casei (9)<br>C. kroppenstedtii (6)<br>C. pseudokroppenstedtii (6)<br>C. parakroppenstedtii (6)<br>C. oculi (6)<br>C. uterequi (6)<br>C. durum (6)<br>C. argentoratense (10)<br>C. sphenisci (10)<br>M. tuberculosis (2,3)<br>M. leprae (2) | WP_278829009.1<br>AJK69750.1<br>WP_339016918.1<br>WP_339016918.1<br>WP_273353752.1<br>WP_082312922.1<br>WP_136141837.1<br>WP_284819823.1<br>WP_312099083.1<br>WP_049150446.1<br>WP_075732789.1<br>WP_151902216.1<br>WP_284850702.1<br>QNE90162.1<br>WP_165008821.1<br>WJY95991.1<br>WP_038550149.1<br>HJC85593.1<br>WP_048380829.1<br>BAC19343.1<br>WP_020440424.1<br>WJY90483.1<br>WCZ33209.1<br>WP_121911514.1<br>WP_259825279.1<br>WP_269946870.1<br>WP_408934594.1<br>WP_340418550.1<br>WP_256886994.1<br>WP_048760415.1<br>WP_156231755.1<br>WP_284875242.1<br>WP_276686843.1<br>WP_406604991.1<br>WP_284866906.1<br>WP_221923110.1<br>KQB83177.1<br>WP_407919165.1<br>WP_315560426.1<br>WP_281707215.1<br>WP_075693089.1<br>WP_031705037.1<br>WP_081439395.1 | | | | |
| 10 | Methionine--tRNA ligase | C. striatum (1,2,3,4,5,6,7)<br>C. simulans (1,2,3,4,7)<br>C. accolens (1,2,3,4,7)<br>C. pseudogenitalium (1,2,3,5,6)<br>C. tuberculostearicum (1,2,3,5,6)<br>C. uterequi (1,6) | WP_166684504.1<br>WP_062039073.1<br>PCC83265.1<br>WP_083798074.1<br>WP_301988834.1<br>WP_047259250.1 | 1<br>2<br>3<br>4<br>5<br>6 | DFLAEFGPDALR<br>DIDWGIPIPVEGWEENSAK<br>FSLNLLEDMRPR<br>INGETPEFIETEHFLLDLPSVK<br>INNELANGWGNLVNR<br>VHETLGR | 382-393<br>254-272<br>238-249<br>197-218<br>417-431<br>541-547 | 70239 |

| | | | | | | | |
|---|---|---|---|---|---|---|---|
| | | C. aurimucosum (1,2,3,5)<br>C. kefirresidentii (1)<br>C. marquesiae (1)<br>C. curieae (1)<br>C. yonathiae (1)<br>C. faecale (1)<br>C. confusum (1,5,6)<br>C. genitalium (1,2,3,5,6)<br>C. auris (2,3,6)<br>C. appendicis (2,3)<br>C. urinipleomorphum (2,3,6)<br>C. afermentas (2,3,6)<br>C. lipophilum (2,3,6)<br>C. pilosum (2,3)<br>C. cystitides (2,3,6)<br>C. lizhenjunii (2,3,5)<br>C. glaucum (2,3)<br>C. coyelae (2,3,6)<br>C. camporealensis (2,4,5,6)<br>C. renale (5,6)<br>C. matruchotii (5,6)<br>C. casei (5,6)<br>C. ammoniagenes (5,6)<br>C. pseudodiphtheriticum (5)<br>C. maris (5)<br>C. phoceense (5)<br>C. stationis (5)<br>C. breve (5)<br>C. propinquum (5)<br>C. xerosis (6)<br>C. macginleyi (7)<br>C. pyruviciproducens (7)<br>C. glucuronolyticum (7) | WP_080971210.1<br>WP_086588663.1<br>WP_337888465.1<br>WP_269946143.1<br>WP_238801803.1<br>WP_290279037.1<br>WP_290225462.1<br>EFK54434.1<br>WP_290342867.1<br>WP_273412517.1<br>WP_204211794.1<br>WP_286361247.1<br>MCO6394205.1<br>STC69993.1<br>WP_373369971.1<br>WP_165007843.1<br>WP_290186399.1<br>WP_257964897.1<br>WP_368530563.1<br>PFG26969.1<br>WP_315536783.1<br>WP_301437668.1<br>WP_050759841.1<br>ERJ45537.1<br>AGS34311.1<br>WP_257076659.1<br>WP_313678355.1<br>WP_284825964.1<br>WP_284576069.1<br>WP_406913808.1<br>WP_200439953.1<br>WP_016457008.1<br>MDY5834768.1 | 7 | GDQCDNCGNQLDPVDLIDPVSK | 154-175 | |
| 11 | Phenylalanine--tRNA ligase subunit alpha | C. simulans (1,3)<br>C. kefirresidentii (3)<br>C. striatum (1,3)<br>C. tuberculostearicum (3)<br>C. aurimucosum (3)<br>C. massiliense (3)<br>C. marquesiae (1,3)<br>C. curieae (3)<br>C. endometrii (3)<br>C. diphtheriae (1)<br>C. ammoniagenes (1,2)<br>C. casei (1,2)<br>C. pseudogenitalium (1,2,3)<br>C. appendicis (1,2)<br>C. accolens (1,2)<br>C. urinipleomorphum (1,2) | WP_062040652.1<br>WP_301732491.1<br>WP_166683955.1<br>WP_301431877.1<br>WP_049358443.1<br>WP_022863667.1<br>WP_340419563.1<br>WP_269947124.1<br>WP_210726576.1<br>WP_196976164.1<br>GJN43009.1<br>AHI20136.1<br>EFQ80087.1<br>WP_259796797.1<br>WP_039885592.1<br>WP_087117863.1 | 1<br>2<br>3 | FTLPFGVQA<br>THTSPVQVR<br>TQTGALHPITALSER | 340-348<br>181-189<br>111-125 | 38484 |

| | | | | | | | |
|---|---|---|---|---|---|---|---|
| | | C. halotolerans (1,2)<br>C. sanguinis<br>C. ulcerans (1)<br>C. rouxii (1)<br>C. pseudotuberculosis (1)<br>C. belfantii (1)<br>C. camporealensis (1,3)<br>C. silvaticum (1)<br>C. sanguinis (1)<br>C. lipophiloflavum (1)<br>C. faecipullorum (1,3)<br>C. yonathiae (1,3)<br>C. phoceense (1)<br>C. glutamicum (2)<br>C. incognita (1)<br>C. glaucum (2)<br>C. coyleae (2)<br>C. mucifaciens (2)<br>C. afermentans (2)<br>C. ureicelerivorans (2)<br>C. faecium (2)<br>C. lactis (2)<br>C. pseudodiphtheriticum (2,3)<br>C. cystitides (2)<br>C. propinquum (2,3)<br>C. renale (2)<br>C. endometrii (3)<br>C. minutissimum (3)<br>C. guaraldiae (3)<br>C. intestinale (3)<br>C. lizhenjunii (3)<br>C. confusum (3) | WP_027004571.1<br>MCT2154894.1 (1)<br>WP_046095848.1<br>WP_155872648.1<br>WP_014800376.1<br>WP_197690070.1<br>WP_105360319.1<br>WP_087453886.1<br>WP_375545438.1<br>WP_006840934.1<br>HIX78435.1<br>WP_340418582.1<br>WP_068801748.1<br>WP_051861055.1<br>WP_185177013.1<br>WJZ07667.1 (2)<br>WP_284780127.1<br>WP_343033586.1<br>WP_290172343.1<br>WP_301694118.1<br>WP_269511710.1<br>WP_053412282.1<br>WP_284584849.1<br>WJY82325.1<br>WP_249606488.1<br>WP_098389239.1<br>QCB28367.1<br>WP_239188682.1<br>WP_144014977.1<br>WP_250224733.1<br>WP_165010911.1<br>WP_290226113.1 | | | | |
| 12 | Phenylalanine--tRNA ligase beta subunit-related protein, partial | C. striatum (1) | WP_218023517.1 | 1 | AVGENLIDELR | 104-114 | 14771 |
| 13 | Proline--tRNA ligase | C. striatum (1,2,3,4,5,6)<br>C. simulans (1,2,3,4,5)<br>C. falsenii (1)<br>C. tuberculostearicum (1)<br>C. marquesiae (1)<br>C. kefirresidentii (1)<br>C. phoceense (1)<br>C. pyruviproducens (1)<br>C. singular (1)<br>C. aurimucosum (1)<br>C. curiae (1)<br>C. hiratae (2)<br>C. hesseae (2)<br>C. intestinale (2) | WP_086891171.1<br>WP_248091668.1<br>WP_276785429.1<br>WP_316992206.1<br>WP_408932914.1<br>WP_301731966.1<br>WP_068801856.1<br>WP_311354157.1<br>WP_144790244.1<br>WP_049361175.1<br>WP_269945412.1<br>WP_158396503.1<br>WP_339016578.1<br>WP_250224110.1 | 1<br>2<br>3<br>4<br>5<br>6 | AYQNIFNR<br>DAAALEAGQK<br>LSNLFLR<br>LVEDLDAAGIEVLFDDRPK<br>SFADGIIELR<br>KIENIVR | 182-189<br>506-515<br>5-11<br>516-534<br>558-567<br>54-60 | 65390 |

| | | | | | | | |
|---|---|---|---|---|---|---|---|
| | | C. guaraldiae (2) | WP_143335416.1 | | | | |
| | | C. aurimucosum (2) | WP_216379689.1 | | | | |
| | | C. minutissimum (2) | KKO80322.1 | | | | |
| | | C. faecipullorum (2) | HIX79127.1 | | | | |
| | | C. genitalium (5) | EFK53727.1 | | | | |
| | | C. endometrii (5) | WP_136141386.1 | | | | |
| | | C. glucuronolyticum (6) | WP_084037021.1 | | | | |
| | | C. ulcerans (6) | WP_101507035.1 | | | | |
| | | C. pseudotuberculosis (6) | WP_048589024.1 | | | | |
| | | C. sylvaticum (6) | WP_263059967.1 | | | | |
| 14 | Serine--tRNA ligase | C. striatum (1,2,5,6,7,8,9,10,11) | WP_005529450.1 | 1 | DHLELGESLGLIDVK | 135-149 | 46201 |
| | | C. singulare (1,2,3,4,5,6,10) | AJI79989.1 | 2 | DMLAAMELPYR | 303-313 | |
| | | C. aurimucosum (2,3,4,5,6,10) | WP_102233427.1 | 3 | ELTSTSNCTTFQAR | 344-357 | |
| | | C. endometrii (1,2,4,11) | WP_136141976.1 | 4 | ENPDVVR | 9-15 | |
| | | C. camporealensis (1,2,3,4,9,11) | WP_035106303.1 | 5 | IGQASPEERPALLEGSNELK | 59-78 | |
| | | C. minutissimum (1,2) | WP_039672610.1 | 6 | IGQASPEERPALLEGSNELKEK | 59-80 | |
| | | C. urealyticum (2,3,4,10) | PZO98914.1 | 7 | KAEAELAEAEAK | 83-94 | |
| | | C. flavescens (2,4,10) | WP_075730709.1 | 8 | KFDNEAWVPTQGCYR | 329-343 | |
| | | C. faecipullorum (2,5,6,10) | HIX79237.1 | 9 | SAILAADELR | 40-49 | |
| | | C. hiratae (2,4,5,6,10) | WP_144013370.1 | 10 | TIDIAGGDLGSSAAR | 314-328 | |
| | | C. curieae (1,2) | WP_269946751.1 | 11 | YTGWSSCFR | 249-257 | |
| | | C. marquesiae (1,2,10) | WP_198492203.1 | | | | |
| | | C. tuberculostearicum (1,2,10) | WP_408927931.1 | | | | |
| | | C. yonathiae (1,2) | WP_238801916.1 | | | | |
| | | C. lizhenjenii (1,3,4,5,6,10) | WP_408920680.1 | | | | |
| | | C. accolens (1,5,6,9,10,11) | WP_284609831.1 | | | | |
| | | C. macginleyi (1) | WP_200438689.1 | | | | |
| | | C. otitides (3) | CCI83901.1 | | | | |
| | | C. diphtheriae (3,4) | CAB0526331.1 | | | | |
| | | C. xerosis (3) | NMF10115.1 | | | | |
| | | C. variabile (3,4,11) | WP_301529747.1 | | | | |
| | | C. glutamicum (3,5,6) | BAV24516.1 | | | | |
| | | C. renale (3) | SQG63498.1 | | | | |
| | | C. ulcerans (3) | AEG82583.1 | | | | |
| | | C. pseudotuberculosis (3) | AKS14320.1 | | | | |
| | | C. felinum (3,4) | WJY96240.1 | | | | |
| | | C. nuruki (3) | WP_334144462.1 | | | | |
| | | C. falsenii (3) | WP_025401838.1 | | | | |
| | | C. glaucum (3) | GAA1165419.1 | | | | |
| | | C. resistens (3,4) | WP_273352519.1 | | | | |
| | | C. terpenotabidum (3,4) | WP_020440208.1 | | | | |
| | | C. bovis (3) | WP_301714001.1 | | | | |
| | | C. auriscanis (3) | WP_265915246.1 | | | | |
| | | C. faecigallinarum (3,4) | HJC84745.1 | | | | |
| | | C. kroppenstedtii (3) | WP_012730894.1 | | | | |
| | | C. genitalium (4) | EFK55431.1 | | | | |
| | | C. durum (4) | WP_006062214.1 | | | | |
| | | C. glyciniphilum (4) | GAA1419423.1 | | | | |
| | | C. minutissimum (4,5,6,10) | WP_039672610.1 | | | | |

| | | | | | | | |
|---|---|---|---|---|---|---|---|
| | | C. epidermidicanis (4)<br>C. stationis (5,6,9)<br>C. casei (5,6,9)<br>C. hindlerae (5,6)<br>C. mastitidis (5,6)<br>C. phocae (10)<br>C. phoceense (10,11)<br>C. ammoniagenes (9)<br>C. occultum (9,11)<br>C. suedekumii (11)<br>C. humireducens<br>C. nasicanis (11)<br>C. maris (11)<br>C. comes (11)<br>C. marinum (11) | AKK04171.1<br>WP_300917282.1<br>WP_098072537.1<br>WP_208768217.1<br>WP_018117787.1<br>WP_075736013.1<br>WP_368527387.1<br>WP_003847929.1<br>QGU08552.1<br>WP_284875442.1<br>WP_040087733.1<br>WP_377001273.1<br>WP_020935816.1<br>WP_156229142.1<br>WP_042622256.1 | | | | |
| 15 | Threonine--tRNA ligase | C. simulans (1,2,3,4,5,6,7,8)<br>C. striatum (1,2,3,4,5,6,7,8)<br>C. incognita (1,3)<br>C. minutissimum (2,6,7)<br>C. stationis (3)<br>C. casei (3)<br>C. pseudodiphtheriticum (3)<br>C. camporealensis (3)<br>C. phocae (3)<br>C. endometrii (3,8)<br>C. lizhenjunii (3,6)<br>C. confusum (3)<br>C. tuberculostearicum (5,6,8)<br>C. marquesiae (5,6,8)<br>C. kefirresidentii (5,6,8)<br>C. yonathiae (5,6,8)<br>C. aurimucosum (5,6,7,8)<br>C. atypicum (6)<br>C. guaraldiae (6,7)<br>C. hesseae (6,7,8)<br>C. intestinale (6,7)<br>C. parakroppenstedtii (6)<br>C. hiratae (6,7,8)<br>C. curieae (6)<br>C. glaucum (7)<br>C. riegelii (7)<br>C. afermentans (7)<br>C. ureicelerivorans (7)<br>C. fournieri (7)<br>C. mucifaciens (7)<br>C. freneyi (8)<br>C. gallinarum (8)<br>C. stationis (8)<br>C. ammoniagenes (8) | WP_062038780.1<br>WP_049151806.1<br>WP_185176045.1<br>WP_039676886.1<br>WP_278755484.1<br>WP_006823586.1<br>WP_249619538.1<br>WP_368530171.1<br>WP_075733859.1<br>WP_136141286.1<br>WP_165006292.1<br>WP_290222023.1<br>WP_316980731.1<br>WP_408932896.1<br>WP_086587565.1<br>WP_238799957.1<br>WP_049360663.1<br>WP_038605235.1<br>WP_154736857.1<br>WP_101735680.1<br>WP_250224684.1<br>PMC66519.1<br>WP_158396584.1<br>WP_269945329.1<br>WP_095661101.1<br>WP_311356439.1<br>WP_063936800.1<br>WP_273408748.1<br>WP_085958077.1<br>WP_168683751.1<br>WP_284822152.1<br>WP_225213646.1<br>WP_278755484.1<br>WP_003845623.1 | 1<br>2<br>3<br>4<br>5<br>6<br>7<br>8 | ELELPNKGPEAVVVVR<br>GNVDPDSDEAAEVGAGDLTHYDNINPR<br>GPHLPTTK<br>INEQPNEELISAR<br>STSILQSVAEK<br>TGEVEWFDLCR<br>VPFMLLAGER<br>YIPAFTLTR | 29-44<br>169-195<br>207-214<br>674-686<br>475-485<br>196-206<br>628-637<br>215-223 | 77045 |

| | | | | | | | |
|---|---|---|---|---|---|---|---|
| | | C. flavescens (8)<br>C. occultum (8) | WP_301501862.1<br>WP_156231046.1 | | | | |
| 16 | Tryptophan--tRNA ligase | C. simulans (1,2,3,4,5)<br>C. aurimucosum (1,2,3,6)<br>C. minutissimum (1,2,3)<br>C. singular (1,2,3)<br>C. flavescens (2,3)<br>C. appendicis (1)<br>C. phocae (1)<br>C. diphtheria (1)<br>C. pseudokroppenstedtii (1)<br>C. parakroppenstedtii (1)<br>C. kroppenstedtii (1)<br>C. ulcerans (1)<br>C. intestinale (1,2,3)<br>C. hiratae (1,2,3)<br>C. faecipullorum (1,2)<br>C. lizhenjunii (2)<br>C. confusum (2)<br>C. striatum (2)<br>C. pseudodiphtheriticum (4)<br>C. otitides (4)<br>C. lubricantis (5)<br>C. variabile (5)<br>C. accolens (6)<br>C. macginleyi (6)<br>C. curieae (6)<br>C. kefirresidentii (6)<br>C. tuberculostearicum (6)<br>C. marquesiae (6)<br>C. camporealensis (6) | WP_248092517.1<br>WP_049157255.1<br>WP_115021235.1<br>WP_239180133.1<br>WP_301501895.1<br>WP_284834601.1<br>WP_075735391.1<br>WP_041626869.1<br>WP_337447662.1<br>WP_221926126.1<br>WP_012732174.1<br>WP_041476981.1<br>WP_250224619.1<br>WP_158397699.1<br>HIX78505.1<br>WP_165241561.1<br>WP_290224942.1<br>WP_284765171.1<br>WP_249617596.1<br>WP_004601070.1<br>WP_026196333.1<br>WP_301519901.1<br>WP_302524850.1<br>WP_200437379.1<br>WP_269946368.1<br>WP_284835541.1<br>WP_316986227.1<br>WP_284788121.1<br>WP_035106676.1 | 1<br>2<br>3<br>4<br>5<br>6 | IYDLQEPTSK<br>SAVTDDLGVVAFDR<br>SAVTDDLGVVAFDREK<br>GLINLLDPPK<br>VDTADALEAFTTPLK<br>VPEPFIPEGAAK | 196-205<br>232-245<br>232-247<br>214-223<br>278-292<br>181-192 | 37730 |
| 17 | Tyrosine--tRNA ligase | C. simulans (1)<br>C. striatum (1)<br>C. glucuronolyticum (1)<br>C. glutamicum (1)<br>C. accolens (1) | WP_284841614.1<br>WP_086891343.1<br>MCI6205359.1<br>WP_011897210.1<br>PCC82778.1 | 1 | MNIIDELSWR | 1-10 | 46397 |
| 18 | Valine--tRNA ligase | C. simulans (1,2,3,4,5,6,7,8,9,10,11)<br>C. striatum (1,2,3,4,5,6,7,8,9,10,11)<br>C. diphtheria (1,2)<br>C. falsenii (1,2,9)<br>C. jeikeium (1)<br>C. dentalis (1,2)<br>C. urogenitale (1)<br>C. urealyticum (1,2)<br>C. minutissimum (1)<br>C. pseudotuberculosis (1)<br>C. accolens (2,3,9,11)<br>C. mucifaciens (2)<br>C. xerosis (2) | WP_284841779.1<br>WP_236592620.1<br>AEX77203.1<br>WP_039910879.1<br>WP_071056164.1<br>WP_312589224.1<br>WP_151902404.1<br>WP_148812268.1<br>WP_039675222.1<br>WP_014522455.1<br>EFM42991.1<br>NKY69062.1<br>WP_377782709.1 | 1<br>2<br>3<br>4<br>5<br>6<br>7<br>8<br>9<br>10<br>11<br>12 | AIGDSVDWSR<br>FTLDEGLSR<br>GANPGVDLPLGSDAAAASR<br>HNLDMPTIMDSTGHIANTGTK<br>IADADKLITELR<br>IPIWYGPETTDAEGNTTR<br>LDFASVDLAGQENLVR<br>QQVAQEEVDR<br>SLGNGIDPMDWVR<br>VIADDYVDMEFGTGAVK<br>YGSLNDDEPHLIVATTR<br>LEKDLAEANR | 167-176<br>179-187<br>612-630<br>325-345<br>794-805<br>450-467<br>823-838<br>923-932<br>586-598<br>293-309<br>238-254<br>876-885 | 105845 |

| | | | | | | | |
|---|---|---|---|---|---|---|---|
| | | C. terbenotabidum (2)<br>C. kroppenstedtii (2)<br>C. bovis (2)<br>C. auriscanis (2,9)<br>C. macginleyi (2)<br>C. phoceense (5,10)<br>C. lizhenjunii (5)<br>C. resistens (9)<br>C. flavescens (10)<br>C. aurimucosum (11)<br>C. glyciniphilum (12) | WP_020440725.1<br>ACR18108.1<br>QQC48132.1<br>WP_265914808.1<br>WP_375231833.1<br>WP_257045019.1<br>WP_196823557.1<br>WP_013888011.1<br>KAA8720443.1<br>WJY71149.1<br>WP_038549557.1 | | | | |
| 20 | Methionyl-tRNA formyltransferase | C. minutissimum (1,2)<br>C. simulans (1,2)<br>C. tuberculostearicum (1,2)<br>C. kefirresidentii (1)<br>C. curieae (1)<br>C. pseudogenitalium (1)<br>C. marquesiae (1,2)<br>C. aurimucosum (1,2)<br>C. accolens (1)<br>C. macginleyi (1) | WP_039675222.1<br>WP_248090393.1<br>WP_316993257.1<br>WP_259824576.1<br>WP_269945275.1<br>WP_005324883.1<br>WP_269953003.1<br>WP_049360763.1<br>WP_284636430.1<br>WP_200438051.1 | 1<br>2 | IIFAGTPEPAVVALEK<br>GAAPVQAAIAAGDER | 3-18<br>121-135 | 33780 |
| 21 | CCA tRNA nucleotidyltransferase | C. pseudodiphtheriticum (1)<br>C. propinquum (1) | WP_284862401.1<br>WP_284603917.1 | 1 | MNSSDTQQIALLARATEAVR | 1-20 | 55216 |
