## Supplemental Table S7 for "Peptidome Analysis of Western Blots Identifies Natural Bispecific Antibody-Bound *Corynebacterium* and Phage B-cell Epitopes with Potential Relevance to Psoriasis"

### Table S7. Elongation Factors are *Corynebacterium* Antigens.

| Antigen Number | Identified Antigen | Corynebacterium (C.), *Mycobacterium* (M.) *tuberculosis* | Accession Number | Peptide Number | Peptide Sequence | Peptide Start/Stop | Molecular Weight |
|---|---|---|---|---|---|---|---|
| 1 | Elongation factor G | C. striatum (1,2,3,4,6,7,8,9,10,11,12,13,14,15,16,17) | WP_284806442.1 | 1 | ATLEDGAYHDVDSSEMAFK | 574-592 | 77548 |
| | | C. simulans (1,2,3,4,5,6,7,8,9,10,11,12,13,14,15,16,17) | WP_061920474.1 | 2 | FENAVTGGR | 534-542 | |
| | | C. riegelii (13) | WP_201822638.1 | 3 | GKVETGAEPTIEEIPADLQEK | 186-206 | |
| | | C. diphtheria (2,5,12,14,17) | WP_072574199.1 | 4 | GKVETGAEPTIEEIPADLQEKAEEYR | 186-211 | |
| | | C. pseudotuberculosis (2,5,12,14,17) | WP_278072391.1 | 5 | ILFYTGINR | 28-36 | |
| | | C. faecale (1,5,14,17) | WJY91274.1 | 6 | KTVQSLDYTHK | 488-498 | |
| | | C. efficiens (1,2,5,14,17) | BAC17326.1 | 7 | LAGSQVLK | 593-600 | |
| | | C. flavescens (1,5,6,7,10,12,14,15,16,17) | WP_075729096.1 | 8 | LLEAVAESDEELMEK | 214-228 | |
| | | C. pseudodiphtheriticum (1,2,5,7,11,14,17) | WP_284873128.1 | 9 | LLEAVAESDEELMEKYFGGEELTLDEIK | 214-241 | |
| | | C. accolens (1,2,5,11,14,15,17) | WP_302520668.1 | 10 | TVQSLDYTHK | 489-498 | |
| | | C. propinquum (1,2,5,7,11,14,17) | WP_018121317.1 | 11 | VEANIGNPQVAYR | 471-483 | |
| | | C. phoceense (1,5,11,12) | WP_068801243.1 | 12 | VGETHDGASTTDWMEQEKER | 38-57 | |
| | | C. callunae (1,2,5,12) | WP_411206865.1 | 13 | VIVTIEPYNPDPETLEEGESATYK | 510-533 | |
| | | C. marquesiae (1,5,11,15) | WP_284811125.1 | 14 | VLDGAVAVFDGK | 96-107 | |
| | | C. macginleyi (1,2,5,11,15) | WP_200439524.1 | 15 | VPLSEMFGYIGDLR | 656-669 | |
| | | C. lizhenjunii (1,3,4,5,11,12,15) | WP_165008661.1 | 16 | YFGGEELTLDEIK | 229-241 | |
| | | C. glutamicum (1,2,5) | WP_145832257.1 | 17 | EGVEPQSEQVWR | 130-141 | |
| | | C. curiae (1) | WP_269946453.1 | | | | |
| | | C. endometrii (1,3,4,7,15) | WP_136140509.1 | | | | |
| | | C. kefirresidentii (1,12,15) | WP_408911796.1 | | | | |
| | | C. casei (1,2,5,6,7,10,11,12) | WP_301438352.1 | | | | |
| | | C. aurimucosum (1,2,7,11,12,15) | WP_193628947.1 | | | | |
| | | C. belfantii (2,5,12,14,17) | SNW30611.1 | | | | |
| | | C. renale (1,2,5,14,17) | STD03615.1 | | | | |
| | | C. kroppenstedtii (2,12,14,17) | KXB49586.1 | | | | |
| | | C. mastitidis (2,5,12,14,17) | MCH6197781.1 | | | | |
| | | C. oculi (2,5,12,14,15) | KQB84337.1 | | | | |
| | | C. tuberculostearicum (2,5,11,15) | WP_301715404.1 | | | | |
| | | C. vitaeruminis (2,11,12,17) | WP_313286197.1 | | | | |
| | | C. faecigallinarum (2,5,12) | HJC84212.1 | | | | |
| | | C. urinipleomorphum (3,4) | WP_087117141.1 | | | | |
| | | C. xerosis (5,12,14,17) | WP_155869056.1 | | | | |
| | | C. lactis (5,12,13) | WP_082313017.1 | | | | |
| | | C. freneyi (5,8,9,11,14,17) | WP_035121343.1 | | | | |
| | | C. phocae (6,7) | WP_075735647.1 | | | | |
| | | C. confusum (7,15) | WP_290224666.1 | | | | |
| | | C. hesseae (7,15) | WP_269949025.1 | | | | |
| | | C. stationis (7) | WP_168969502.1 | | | | |
| | | C. kutscheri (7,12,15) | WP_269471347.1 | | | | |
| | | C. incognita (7,15) | WP_185175285.1 | | | | |
| | | C. ammoniagenes (7) | WP_003848062.1 | | | | |
| | | C. sanguinis (8) | WP_259790532.1 | | | | |
| | | C. matruchotii (8) | WP_278720817.1 | | | | |
| | | C. uropygiale (11,12,14,17) | WP_338026117.1 | | | | |

| | | | | | | | |
|---|---|---|---|---|---|---|---|
| | | C. amycolatum (11,12,14,17) | WP_115598015.1 | | | | |
| | | C. massiliense (11) | WP_027018760.1 | | | | |
| | | C. argentoratense (12) | WP_278763771.1 | | | | |
| | | C. otitidis (14,17) | WP_046643844.1 | | | | |
| | | C. phocense (15) | WP_068801243.1 | | | | |
| | | C. camporealensis (15) | WP_368530631.1 | | | | |
| | | C. curieae (15) | WP_269946453.1 | | | | |
| | | C. yonathiae (15) | WP_238800414.1 | | | | |
| | | C. bovis (17) | WP_125207324.1 | | | | |
| | | M. tuberculosis (14,15,17) | WP_031679490.1 | | | | |
| | | M. leprae (14,15,17) | WP_010908589.1 | | | | |
| 2 | Elongation factor Tu | C. simulans (2,3,4,5,6,7,8,9,10,11,12,13) | WP_248093753.1 | 1 | CDMVDDEEIIELVEMEVR | 140-157 | 43968 |
| | | C. auris (1,5,13) | WP_290342538.1 | 2 | EHVLLAR | 120-126 | |
| | | C. efficiens (1) | WP_006769810.1 | 3 | GITINISHVEYSTPK | 62-76 | |
| | | C. stationis (1,3,6,7,9,10) | NME89210.1 | 4 | GTVVTGRVER | 226-235 | |
| | | C. striatum (3,6,7,12,13,14) | WP_272706989.1 | 5 | HYAHVDAPGHADYIK | 78-92 | |
| | | C. glucuronolyticum | WP_260322383.1 | 6 | KMMDYTEAGDNCGLLLR | 266-282 | |
| | | C. casei (1,3,4,6,7,8,9,10,11,14) | WP_006823212.1 | 7 | MMDYTEAGDNCGLLLR | 267-282 | |
| | | C. flavescens (1,14) | WP_075729097.1 | 8 | QVGVPYILVALNK | 127-139 | |
| | | C. vitaeruminis (1) | WP_034649074.1 | 9 | SLSTTVTGIEMFR | 253-265 | |
| | | C. mucifaciens (1) | WP_168684063.1 | 10 | SLSTTVTGIEMFRK | 253-266 | |
| | | C. coyleae (1) | WP_101740556.1 | 11 | TTTTAAITK | 26-34 | |
| | | C. hindlerae (1,2,4,5,8,11) | WP_182386981.1 | 12 | CDMVDDEEIIELVEMEIR | 140-157 | |
| | | C. faecale (1,2,5,8,11) | WP_290278349.1 | 13 | ELLAEQDYDEEAPIVHISALK | 158-178 | |
| | | C. suedecumii (1,2,4,5,8,11) | WP_284875869.1 | 14 | SISTTVTGIEMFR | 253-265 | |
| | | C. durum (1) | WP_315183756.1 | | | | |
| | | C. comes (1) | WP_156226871.1 | | | | |
| | | C. glaucum (1,2,4,5,8) | WP_290185955.1 | | | | |
| | | C. afermentans (1,6,7,12) | WP_194561138.1 | | | | |
| | | C. ureicelerivorans (1) | WP_301694876.1 | | | | |
| | | C. xerosis (1) | WP_155869058.1 | | | | |
| | | C. pollutisoli (1) | NLP38935.1 | | | | |
| | | C. callunae (1,2,4,5,8,11) | WP_247775987.1 | | | | |
| | | C. imitans (1) | WP_239245999.1 | | | | |
| | | C. fournieri (1,2,4,5,8,11,12) | WP_085956816.1 | | | | |
| | | C. breve (1,2,3,5,8,11) | WP_284825395.1 | | | | |
| | | C. cystitides (1,2,4,5,7,8,11) | WP_257159257.1 | | | | |
| | | C. gallinarum (1) | WP_191732734.1 | | | | |
| | | C. canis (1) | WP_146325690.1 | | | | |
| | | C. nasicanis (1) | WP_377001956.1 | | | | |
| | | C. humireducens (1,4) | NLA56465.1 | | | | |
| | | C. diphtheria (2,4,5,8,11,12) | WP_304680562.1 | | | | |
| | | C. pseudotuberculosis (2,4,5,8,11,12) | WP_322823421.1 | | | | |
| | | C. ulcerans (2,4,5,8,11,12) | KPJ25107.1 | | | | |
| | | C. argentoratense (2,4,5,8) | WP_234858297.1 | | | | |
| | | C. otitides (2,3,4,5,8,11) | WP_004601455.1 | | | | |
| | | C. matruchotii (2,4,5,8,11) | WP_315535724.1 | | | | |
| | | C. urogenitale (2,4,5,8,11) | WP_151903405.1 | | | | |
| | | C. ulceribovis (2,5,8,11) | WP_018023279.1 | | | | |

| | | | | | | | |
|---|---|---|---|---|---|---|---|
| | | C. lizhenjunii (2,3,4,5,6,7,8,11)<br>C. marquesiae (2,3,4,5,6,7,11,12,13)<br>C. accolens (2,3,4,5,6,7,8,11,13)<br>C. kefirresidentii (2,3,4,5,6,7,8,11,12,13)<br>C. jeikeium (2,4,5,8,11)<br>C. tuberculostearicum (2,3,4,5,6,7,11,12,13)<br>C. yonathiae (2,3,4,5,6,7,8,11,12,13)<br>C. confusum (2,3,4,5,6,7,8,11,12)<br>C. urinipleomorphum (2,4,5,8,11,12)<br>C. glutamicum (2,4,5,8,11,13)<br>C. dentalis (2,4,5,8,11)<br>C. faecium (2,4,5,8,11)<br>C. kroppenstedtii (2,4,5,8,11)<br>C. massiliense (3,4,5,6,7,8,11,12)<br>C. camporealensis (3,6,7,12)<br>C. phoceense (3,6,7,12,13)<br>C. macginleyi (3,6,7)<br>C. propinquum (3)<br>C. pseudodiphtheriticum (3)<br>C. curieae (3,6,7,12,13)<br>C. oculi (3,6,7,12)<br>C. variabile (4,5,8,11)<br>C. endometrii (6,7,12)<br>C. incognita (6,7,9,10)<br>C. minutissimum (6,7,12,13)<br>C. amycolatum (12,13)<br>C. occultum (12)<br>C. renale (12)<br>C. appendicis (12)<br>M. tuberculosis (2,4,5,8)<br>M. leprae (2,4,5,8) | WP_165008659.1<br>WP_408926491.1<br>WP_237799364.1<br>WP_394285903.1<br>WP_011274172.1<br>WP_301988991.1<br>WP_340418451.1<br>WP_290224668.1<br>WP_087117142.1<br>WP_211439940.1<br>WP_312097304.1<br>WP_269508393.1<br>WP_303733989.1<br>WP_022863448.1<br>WP_368530630.1<br>WP_257052649.1<br>WP_200445462.1<br>WP_018121316.1<br>WP_027017517.1<br>WP_269946452.1<br>WP_055122279.1<br>AEK37747.1<br>WP_136140510.1<br>WP_185175286.1<br>WP_039673679.1<br>WP_076773081.1<br>WP_156229970.1<br>WP_048379947.1<br>WP_076598029.1<br>WP_057357813.1<br>WP_010908588.1 | | | | |
| 3 | Transcription elongation factor GreA | C. striatum<br>C. renale<br>C. guaraldiae<br>C. maris<br>C. phoceense<br>C. callunae<br>C. diphtheria<br>C. vitaeruminis<br>C. rouxiii<br>C. pseudotuberculosis<br>C. flavescens<br>C. faecale<br>C. mastitides | MDU3174057.1<br>WP_115243562.1<br>WP_143335638.1<br>WP_020934313.1<br>WP_257053763.1<br>WP_015650808.1<br>WP_014311354.1<br>WP_025252419.1<br>WP_155872206.1<br>WP_013241577.1<br>WP_075729517.1<br>WP_290279155.1<br>WP_018119018.1 | 1<br>2<br>3 | QISEVLANSTTER<br>QYITPETK<br>REEGDLKENAGYDAAREMQDQEEAR | 66-78<br>7-14<br>39-63 | 18711 |
| 4 | Transcription elongation factor GreA | C. stationis<br>C. massiliense<br>C. camporealensis<br>C. renale<br>C. kutscheri | WP_301200234.1<br>WP_027018614.1<br>WP_035105785.1<br>WP_111725846.1<br>WP_046440214.1 | 1<br>2<br>3 | AAASDNKDLETYSEQSPLGAAILGAQEGETR<br>QYITPETK<br>REEGDLKENAGYDAAREMQDQEEAR | 111-141<br>7-14<br>39-63 | 18898 |

| | | | | | | | |
|---|---|---|---|---|---|---|---|
| | | C. kefirresidentii<br>C. ammoniagenes | WP_284835692.1<br>WP_168938798.1 | | | | |
| 5 | Elongation factor P | C. gerontici<br>C. aurimucosum | WP_123934172.1<br>WP_010190126.1 | 1<br>2<br>3<br>4<br>5 | DMTYLYNDGQNYVVMDDK<br>LKDVVSGK<br>LQQIVEFQHVKPGK<br>TGEYLSR<br>VTDKTFNAGVK | 69-86<br>41-48<br>19-32<br>178-184<br>49-59 | 20894 |
| 6 | Elongation factor P | C. pseudodiphthereticum<br>C. striatum | WP_284818356.1<br>WP_049192614.1 | 1<br>2<br>3<br>4 | TGEYLSR<br>VEHTEPGLQGDR<br>VETATVDRR<br>VTDKTFNAGVK | 178-184<br>133-144<br>60-68<br>49-59 | 20876 |
| 7 | Translation elongation factor Ts | C. pseudotuberculosis<br>C. pseudodiphthereticum<br>C. kroppenstedtii<br>C. diphtheriae<br>C. glutamicum<br>C. simulans | ATV80418.1<br>WP_284575352.1<br>WP_012731800.1<br>WP_016829767.1<br>WP_003857558.1<br>WP_062038474.1 | 1<br>2<br>3<br>4<br>5<br>6<br>7<br>8<br>9<br>10<br>11<br>12<br>13<br>14<br>15 | ALDESNGDYDKAVEFLR<br>ANYTAADVK<br>AVTLEGDNVAVYLHQR<br>EATGSGMLDCK<br>EATGSGMLDCKK<br>EDVPAEVVEK<br>EDVPAEVVEKER<br>EEGKPEAALPK<br>EIAEATTREEGKPEAALPK<br>KVSEIVDEESAK<br>KVSEIVDEESAKTGEK<br>SVVLLEQAALSDSK<br>SVVLLEQAALSDSKK<br>VSEIVDEESAK<br>VSEIVDEESAKTGEK | 26-42<br>2-10<br>137-152<br>14-24<br>14-25<br>191-200<br>191-202<br>211-221<br>203-221<br>116-127<br>116-131<br>233-246<br>233-247<br>117-127<br>117-131 | 28940 |
| 8 | Translation elongation factor Ts | C. aurimucosum<br>C. minutissimum<br>C. hesseae | WP_010190381.1<br>WP_239188378.1<br>WP_269948059.1 | 1<br>2<br>3<br>4<br>5<br>6 | ANYTAADVK<br>AVTIEGDNVAVYLHQR<br>EEGKPEAALPK<br>EIAEATTREEGKPEAALPK<br>VNSGEELNNLEIDGK<br>VNSGEELNNLEIDGKK | 2-10<br>137-152<br>211-221<br>203-221<br>101-115<br>101-116 | 28960 |
