## Supplemental Table S8 for "Peptidome Analysis of Western Blots Identifies Natural Bispecific Antibody-Bound *Corynebacterium* and Phage B-cell Epitopes with Potential Relevance to Psoriasis"

### Table S8. Endonucleases are *Corynebacterium* Antigens.

| Antigen Number | Identified Antigen | *Corynebacterium* (C.), *Clavibacter* species | Accession Number | Peptide Sequence | Peptide Start/Stop | Molecular Weight |
|---|---|---|---|---|---|---|
| 1 | Type II CRISPR RNA-guided endonuclease Cas9 | C. matruchotii | WP_315120567.1 | CAYCGAEISFK | 557-567 | 123355 |
| 2 | Restriction endonuclease subunit S | C. jeikeium | WP_011273488.1 | LKVAGPGTVLFAMYGATLGAVSR | 264-286 | 44305 |
| 3 | Endonuclease/exonuclease/ phosphatase family protein | C. glyciniphilum | WP_052539768.1 | SSGATYSR | 174-181 | 26621 |
| 4 | Restriction endonuclease | C. striatum<br>C. simulans<br>C. aurimucosum<br>C. minutissimum<br>C. guaraldiae | WP_062042895.1<br>WP_114976733.1<br>WP_201828944.1<br>KKO78158.1<br>WP_144014664.1 | NNEFVAGELLTR | 156-167 | 14537 |
| 5 | Type II CRISPR RNA-guided endonuclease Cas9 | C. diphtheriae | WP_088265531.1 | EMDGDMR | 510-516 | 121519 |
| 6 | Endonuclease/exonuclease/ phosphatase family protein | Clavibacter species | WP_012299902.1 | TWGNMVPR | 132-139 | 31379 |
| 7 | HNH endonuclease signature motif containing protein | C. amycolatum<br>C. vitaeruminis<br>C. jeikeium | WP_197915149.1<br>WP_048759202.1<br>AYX80873.1 | KNGGATTMENMVMTCKEHNAANDDDR | 318-343 | 43433 |
| 8 | HNH Endonuclease signature motif containing protein | C. glyciniphilum | WP_038546597.1 | LQAAACMERSR | 48-58 | 51643 |
| 9 | HNH endonuclease signature motif containing protein | C. callunae | WP_247776224.1 | TTLDDSADLCQHHHNMKTDGR | 285-305 | 42185 |
| 10 | Endonuclease III | C. efficiens<br>C. gallinarum<br>C. pacaense | WP_006770227.1<br>WP_191734198.1<br>WP_080793107.1 | AACGACMLAADCPSFGLEGPADPMEAQK | 221-248 | 29225 |
