## Supplemental Table S9 for "Peptidome Analysis of Western Blots Identifies Natural Bispecific Antibody-Bound *Corynebacterium* and Phage B-cell Epitopes with Potential Relevance to Psoriasis"

### Table S9. RNA Polymerase Sigma Factor are *Corynebacterium* Antigens.

| Antigen Number | Identified Antigen | *Corynebacterium* (C.) | Accession Number | Peptide Sequence | Peptide Start/Stop | Molecular Weight |
|---|---|---|---|---|---|---|
| 1 | RNA polymerase sigma factor | C. matruchotii<br>C. glucuronolyticum<br>C. falsenii<br>C. otitides<br>C. resistens | WP_314823290.1<br>WP_005394145.1<br>WP_276783450.1<br>WP_004599926.1<br>WP_273353525.1 | VALLNAEQEVSLAK | 198-211 | 60299 |
| 2 | RNA polymerase sigma factor | C. resistens | WP_013889503.1 | MELCGVEK | 1-8 | 24608 |
| 3 | RNA polymerase sigma factor | C. callunae | WP_015650134.1 | TLIDALPVER | 126-135 | 21357 |
| 4 | Sigma-70 family RNA polymerase sigma factor | C. striatum | WP_049160475.1 | ALREAMLQGANT | 173-184 | 21090 |
| 5 | Sigma-70 family RNA polymerase sigma factor | C. durum | WP_231287024.1 | NTAFADR | 103-109 | 21665 |
| 6 | RNA polymerase sigma factor | C. glyciniphilum | WP_038548374.1 | MAAHTSSTSGNDAEGIDAPGTSASSAGR | 1-28 | 60299 |
| 7 | RNA polymerase sigma factor, partial | C. pyruviciproducens | WP_276849450.1 | AASASEPTEDSSEKATAPAK | 104-123 | 64511 |
