## Supplemental Table S10 for "Peptidome Analysis of Western Blots Identifies Natural Bispecific Antibody-Bound *Corynebacterium* and Phage B-cell Epitopes with Potential Relevance to Psoriasis"

### Table S10. RNA-Degradosome Enzymes are Corynebacterium Antigens.

| Antigen Number | Identified Antigen | Corynebacterium (C.), Actinomycetota | Accession Number | Peptide Sequence | Peptide Start/Stop | Molecular Weight |
|---|---|---|---|---|---|---|
| 1 | Ribonuclease J | C. falsenii<br>C. jeikeium<br>C. maccklintockiae | WP_272712888.1<br>WP_111712058.1<br>WP_284803748.1 | AMQGADLTQR | 170-179 | 80008 |
| 2 | Ribonuclease J | C. argentoratense | WP_234870613.1 | GRGGNSNGSSNR | 81-92 | 76327 |
| 3 | Ribonuclease J | C. callunae<br>C. glutamicum<br>C. xerosis<br>C. kroppenstedtii<br>C. phoceense<br>C. argentoratense<br>C. bovis<br>C. deserti<br>C. pseudokroppenstedtii<br>C. parakroppenstedtii<br>C. freneyi<br>C. faecale<br>C. gallinarum<br>C. efficiens<br>C. amycolatum<br>C. vitaeruminis<br>C. diphtheriae | WP_015651527.1<br>WP_143854996.1<br>WP_155870415.1<br>WP_303735538.1<br>WP_181734549.1<br>WP_234870613.1<br>WP_125187070.1<br>WP_053545137.1<br>WP_337447958.1<br>WP_340054851.1<br>WP_035120901.1<br>WP_290275899.1<br>WP_191732025.1<br>WP_011075647.1<br>WP_011075647.1<br>WP_048758409.1<br>WP_071572121.1 | SMQGADLTQR | 134-143 | 76327 |
| 4 | Superfamily II RNA helicase | Gordonia malaquae | SEC55612.1 | VDDLSKR | 127-133 | 17493 |
| 5 | ATP-dependent RNA helicase HrpA | C. tuberculostearicum<br>C.kefirresidentii | WKS54074.1<br>WP_259824647.1 | LPMHLR | 963-968 | |
| 6 | ATP-dependent RNA helicase DeaD | Clavibacter michiganensis subsp. michiganensis | OUD99502.1 | GERRARPAR | 477-485 | 64092 |
| 7 | ATP-dependent RNA helicase HrpA | C. genitalium | WP_040423649.1 | TATATDDHDDSRDRDPK | 621-637 | 147368 |
| 8 | CRISPR system CASCADE complex protein CasC | C. striatum | WP_049063132.1 | AIIDKDGEK<br>AIIDKDGEKFK<br>GLDGYFTEFADSVTIPELEER<br>LAQEAIETQR<br>LVGELICK<br>RFPAEVMGK<br>SVSLVNAFEEPVEAGAAGR<br>TAVDSAAAFVDSFVK | 145-153<br>145-155<br>349-369<br>324-333<br>67-74<br>55-63<br>298-316<br>256-270 | 41921 |
| 9 | Type I-E CRISPR-associated protein Cas5/CasD | C. pseudodiphtheriticum | WP_284586519.1 | SGIIGLLAAAEGRR | 34-47 | 26796 |
| 10 | Type II CRISPR RNA-guided endonuclease Cas9 | C. matruchotii | WP_315120567.1 | CAYCGAEISFK | 557-567 | 123355 |
| 11 | Type I-E CRISPR-associated protein Cas6/Cse3/CasE | C. striatum<br>C. simulans<br>C. resistens | WP_100086674.1<br>WP_062043657.1<br>WP_042379763.1 | AYGCGLLTLAPAE<br>FLEQIVNGQK<br>GAFPPDIDESQSR | 207-219<br>90-99<br>32-44 | 24213 |

| | | | | | | |
|---|---|---|---|---|---|---|
| | | | | LLANPQAMHAAVR | 19-31 | |
| 12 | Type I-E CRISPR-associated protein Cas2/CasB | C. simulans<br>C. striatum | AMO92348.1<br>WP_049192140.1 | NIGFDYGAFAQDLK | 159-172 | 22653 |
| 13 | Type I-E CRISPR-associated protein Cas5/CasD | C. striatum | WP_100086675.1 | ALLENIESALR | 108-118 | 25688 |
| 14 | Type I-E CRISPR-associated protein Cas6/Cse3/CasE | C. accolens<br>C. macginleyi<br>C. frankenforstense | WP_284639393.1<br>WP_121911426.1<br>WP_075663064.1 | ILINPQR | 7-13 | 26302 |
| 15 | Type II CRISPR RNA-guided endonuclease Cas9 | C. diphtheriae | WP_088265531.1 | EMDGDMR | 510-516 | 121519 |
