## Supplemental Table S11 for "Peptidome Analysis of Western Blots Identifies Natural Bispecific Antibody-Bound *Corynebacterium* and Phage B-cell Epitopes with Potential Relevance to Psoriasis"

### Table S11. Ribosome-Associated Proteins are *Corynebacterium* Antigens.

| Antigen Number | Identified Antigen | *Corynebacterium* (C.), *Prescottella equi* (Peptide numbers) | Accession Number | Peptide Number | Peptide Sequence | Peptide Start/Stop | Molecular Weight |
|---|---|---|---|---|---|---|---|
| 1 | Ribosome recycling factor | Prescottella equi | WP_013415676.1 | 1 | IDEALFEAEEKMEK | 2-15 | 20765 |
| 2 | Ribosome recycling factor | C. striatum (1,2,3,4,5) | WP_239298168.1 | 1 | DGDAGEDEVVAAEKEMEK | 145-162 | 20845 |
| | | C. simulans (1,3,5,6) | WP_239238430.1 | 2 | TTAGYIDQVDK | 163-173 | |
| | | C. confusum (1) | WP_290222258.1 | 3 | VTIPQLTEER | 100-109 | |
| | | C. flavescens (1,3) | WP_075730029.1 | 4 | TTAGYIDQVDKLVANKENELMEV | 163-185 | |
| | | C. tuberculostearicum (1) | WP_296179831.1 | 5 | LVANKENELMEV | 174-185 | |
| | | C. appendicis (3) | WP_273412111.1 | 6 | NSDLGVNPTNDGHVLR | 84-99 | |
| | | C. matruchotii (3,6) | WP_005526295.1 | | | | |
| | | C. canis (3) | WP_146325426.1 | | | | |
| | | C. faecale (3) | WP_290276001.1 | | | | |
| | | C. faecipullorum (3) | HIX79434.1 | | | | |
| | | C. amycolatum (3) | WP_258225198.1 | | | | |
| | | C. fournieri (3) | WP_085958178.1 | | | | |
| | | C. durum (3) | WP_303131072.1 | | | | |
| | | C. urealyticum (3) | WP_148791855.1 | | | | |
| | | C. propinquum (3) | WP_144736238.1 | | | | |
| | | C. vitaeruminis (3) | WP_048761839.1 | | | | |
| | | C. pseudodiphtheriticum (3,6) | WP_284571084.1 | | | | |
| | | C. stationis (3) | WP_066840924.1 | | | | |
| | | C. suedekumii (3) | WP_284874592.1 | | | | |
| | | C. ulceribovis (3,6) | WP_018023793.1 | | | | |
| | | C. variabile (3) | WP_303946240.1 | | | | |
| | | C. glaucum (3) | WP_290184417.1 | | | | |
| | | C. diphtheria (3) | WP_088270290.1 | | | | |
| | | C. cystitidis (3) | WP_257158449.1 | | | | |
| | | C. nasicanis (3) | WP_376998987.1 | | | | |
| | | C. amycolatum (3) | WP_011014829.1 | | | | |
| | | C. incognita (3) | WP_185176147.1 | | | | |
| | | C. jeikeium (3) | WP_005295269.1 | | | | |
| | | C. felinum (3) | WP_277103542.1 | | | | |
| | | C. dentalis (3) | WP_099297089.1 | | | | |
| | | C. efficiens (3) | WP_006767912.1 | | | | |
| | | C. macginleyi (6) | WP_200441245.1 | | | | |
| | | C. mastitides (6) | WP_018117405.1 | | | | |
| | | C. oculi (6) | WP_055121681.1 | | | | |
| | | C. glyciniphilum (6) | WP_038548111.1 | | | | |
| | | C. xerosis (6) | WP_102214614.1 | | | | |
| | | C. otitides (6) | WP_004601811.1 | | | | |
| 3 | Ribosome hibernation-promoting | C. simulans (1,2,3,4,5) | WP_248092216.1 | 1 | AEAKEDSFYAALETALAK | 75-92 | 24329 |
| | | C. striatum (1,2,3,4,5) | WP_368920326.1 | 2 | EDSFYAALETALAK | 79-92 | |
| | | C. efficiens (1,2) | WP_035109635.1 | 3 | NVEVPDHFQER | 15-25 | |

| | | | | | | | |
|---|---|---|---|---|---|---|---|
| | factor,<br>HPF/YfiA<br>family | C. confusum (1,2,6)<br>C. callunae (1,2)<br>C. cystitides (1,2)<br>C. faecale (1,2)<br>C. incognita (1,2)<br>C. renale (1,2)<br>C. halotolerans (1,2)<br>C. marinum (1,2)<br>C. glutamicum (1,2)<br>C. endometrii (1,2)<br>C. camporealensis (1,2,4)<br>C. minutissimum (5)<br>C. singulare (5)<br>C. aurimucosum (3)<br>C. tuberculostearicum (3)<br>C. kefirresidentii (3)<br>C. atypicum (1,2)<br>C. gallinarum (1,2)<br>C. frankenforsztense (1,2)<br>C. ciconiae (1,2)<br>C. pilosum (1.2)<br>C. suedecumii (1,2)<br>C. comes (1,2)<br>C. humireducens (1,2)<br>C. deserti (1,2)<br>C. uropygiale (1,2)<br>C. nasicanis (1,2)<br>C. phoceense (1,2)<br>C. lizhenjunii (1,2)<br>C. occultum (1,2)<br>C. casei (5)<br>C. urealyticum (5) | WP_290225149.1<br>WP_015650572.1<br>WP_257162233.1<br>WP_290278775.1<br>WP_185176954.1<br>WP_048381417.1<br>WP_015400178.1<br>WP_042620857.1<br>WP_059288879.1<br>WP_136140694.1<br>WP_035106884.1<br>WP_239189693.1<br>WP_144793575.1<br>WP_049359388.1<br>WP_316986078.1<br>WP_394286346.1<br>WP_038604683.1<br>WP_191732530.1<br>WP_075663394.1<br>WP_026161524.1<br>WP_018581467.1<br>WP_284876084.1<br>WP_156227052.1<br>WP_040085086.1<br>WP_053544326.1<br>WP_236117843.1<br>WP_377002181.1<br>WP_257069041.1<br>WP_165008520.1<br>WP_156230221.1<br>WP_006821993.1<br>PZO98124.1 | 4<br>5 | SQPTNAQVTITGR<br>YIDPYAETVEDVRPGQIVR | 2-14<br>138-156 | |
| 4 | Ribosome<br>silencing<br>factor | C. striatum (1,2,3)<br>C. simulans(1,2,3)<br>C. variabile (1)<br>C. terpenotabidum (1)<br>C. atypicum (1)<br>C. nasicanis (1)<br>C. marinum(1)<br>C. comes (1)<br>C. efficiens (1)<br>C. faecale (1)<br>C. halotolerans (1)<br>C. deserti (1)<br>C. mucifaciens (1)<br>C. atrinae (1)<br>C. appemndicis (1)<br>C. pyruviciproducens (1) | WP_049151643.1<br>WP_239239470.1<br>WP_014009471.1<br>WP_020440741.1<br>WP_038606393.1<br>WP_376999600.1<br>WP_042621914.1<br>WP_156228730.1<br>WP_006768260.1<br>WP_290276557.1<br>WP_015401596.1<br>WP_053545445.1<br>WP_168685029.1<br>WP_290217031.1<br>WP_284834411.1<br>WP_245554399.1 | 1<br>2<br>3 | EFYGLDR<br>EVESIDEIPLAQPEPEEDEEY<br>MAETAAHAADEK | 100-106<br>136-156<br>10-21 | 17848 |
| 5 | 30S | C. striatum (1,2,3,4,5,6) | WP_086891178.1 | 1 | DIDSEPDFDKAAEALNR | 49-65 | 15312 |

| | | | | | | | |
|---|---|---|---|---|---|---|---|
| | ribosome-binding factor RbfA | C. accolens (2,3,4,6)<br>C. phocae (3)<br>C. yonathiae (3)<br>C. marquesiae (3)<br>C. curieae (3)<br>C. phoceense (3,6)<br>C. amycolatum (3)<br>C. bovis (4)<br>C. matruchotii (4)<br>C. falsenii (4)<br>C. glucuronolyticum (4)<br>C. variabile (4)<br>C. terpenotabidum (4)<br>C. imitans (4)<br>C. dentalis (4)<br>C. glyciniphilum (4)<br>C. diphtheria (4,6)<br>C. macginleyi (4)<br>C. tuberculostearicum<br>C. belfantii (4,6)<br>C. ulcerans (4,6)<br>C. pseudotuberculosis (4,6)<br>C. durum (6)<br>C. maris (6)<br>C. imitans (6)<br>C. oculi (6)<br>C. guaraldiae (6)<br>C. aurimucosum (6)<br>C. intestinale (6)<br>C. hiratae (6)<br>C. hesseae (6)<br>C. mucifaciens (6)<br>C. cystitides (6)<br>C. breve (6)<br>C. genitalium (6)<br>C. fournieri (6) | WP_284643164.1<br>WP_075733704.1<br>WP_288794068.1<br>WP_239273073.1<br>WP_269945404.1<br>WP_257051996.1<br>WP_197914336.1<br>WP_280066360.1<br>EEG27284.1<br>WP_025402861.1<br>EEI26861.1<br>WP_301519352.1<br>WP_020441326.1<br>WP_038591174.1<br>WP_312097840.1<br>WP_304032792.1<br>WP_010935093.1<br>WP_200445254.1<br>WP_005328680.1<br>WP_088266261.1<br>WP_368266558.1<br>WP_014367230.1<br>WP_303130926.1<br>WP_020935003.1<br>WP_038591174.1<br>WP_055122010.1<br>WP_143335792.1<br>WP_158381023.1<br>WP_250224108.1<br>WP_144012981.1<br>WP_269948070.1<br>WP_168684332.1<br>WP_257161277.1<br>WP_284823498.1<br>WP_040423637.1<br>WP_085958154.1 | 2<br>3<br>4<br>5<br>6 | GKDIDSEPDFDKAAEALNR<br>IQEIVASAIER<br>IVGDQLSVR<br>RLEMVTITDTR<br>VTGDLHDATVYYTVR | 47-65<br>5-15<br>73-81<br>21-31<br>32-46 | |
| 6 | Ribosome maturation factor RimP | C. amycolatum<br>C. vitaeruminis<br>C. striatum<br>C. simulans<br>C. minutissimum | WP_076773757.1<br>WP_048761789.1<br>WP_306496345.1<br>WP_062038526.1<br>WP_039675877.1 | 1 | GLDVEDVK | 21-28 | 20823 |
| 7 | Ribosome small subunit-dependent GTPase A | C. jeikeium | WP_07105 7358.1 | 1 | EVTEDCPRGCTHMGPPADPECALD TLTGASAR | 284-315 | 36259 |
