## Supplemental Table S12 for "Peptidome Analysis of Western Blots Identifies Natural Bispecific Antibody-Bound *Corynebacterium* and Phage B-cell Epitopes with Potential Relevance to Psoriasis"

### Table S12. Transcriptional Regulators are *Corynebacterium* Antigens.

| Antigen Number | Identified Antigen | Corynebacterium (C.), Actinomycetota | Accession Number | Peptide Sequence | Peptide Start/Stop | Molecular Weight |
|---|---|---|---|---|---|---|
| 1 | Rrf2 family transcriptional regulator | C. urealyticum | WP_197913865.1 | QLTTFADLGLR | 1-11 | 17130 |
| 2 | Metalloregulator ArsR/SmtB family transcription factor | C. glutamicum | WP_038583602.1 | ALSIGTQWAHVMGIK | 126-140 | 25246 |
| 3 | Response regulator transcription factor | C. simulans<br>C. striatum<br>C. accolens<br>C. marinum | WP_284841897.1<br>WP_284789432.1<br>PCC82549.1<br>WP_042622093.1 | FQGFEVFSANSGTEALR<br>VLVVDDEPNIVELLTVSLK | 9-27<br>19-29 | 26246 |
| 4 | YebC/PmpR family DNA-binding transcriptional regulator | C. durum<br>C. mendelii<br>C. phoceense<br>C. renale<br>C. accolens<br>C. casei<br>C. striatum<br>C. stationis<br>C. aurimucosum<br>C. tuberculostearicum<br>C. terpenotabidum<br>C. simulans<br>C. phocae<br>C. macginleyi<br>C. marquesiae<br>C. faecipullorum<br>C. casei<br>C. curieae<br>C. deserti<br>C. minutissimum<br>C. ammoniagenes<br>C. diphtheria<br>C. pseudotuberculosis | EKX90093.1<br>WP_207118009.1<br>WP_257069670.1<br>WP_098389183.1<br>WP_005278990.1<br>WP_006823595.1<br>WP_244080562.1<br>HCM81013.1<br>WP_201828084.1<br>WP_005328529.1<br>WP_020441237.1<br>WP_239238655.1<br>WP_075733873.1<br>WP_121910785.1<br>WP_284838854.1<br>HIX78750.1<br>MDN6130701.1<br>WP_269945322.1<br>WP_053545061.1<br>WP_039676897.1<br>WP_003845641.1<br>WP_085664464.1<br>WP_014523938.1 | KASVPNDNIER<br>TGGGDPAANPTLDDMIK<br>TGGGDPAANPTLDDMIKK<br>TGLVLVNK | 58-68<br>38-54<br>38-55<br>138-145 | 26966 |
| 5 | Transcription termination/antitermination protein NusG | C. simulans<br>C. striatum | WP_248093691.1<br>WP_086892101.1 | AQTLEVEDSIFEVVVPVEQILENKDGK<br>ETPGVTSFVGNEGNATPVK<br>MELNDAAWSVVR | 109-135<br>164-182<br>152-163 | 30583 |
| 6 | MarR (Multiple antibiotic resistance regulator) family winged helix-turn-helix transcriptional regulator | C. striatum<br>C. simulans<br>C. camporealensis<br>C. urealyticum<br>C. aurimucosum<br>C. hesseae<br>C. guaraldiae<br>C. minutissimum<br>C. accolens<br>C. tuberculostearicum | WP_258552506.1<br>WP_062042989.1<br>WP_321112847.1<br>PZP01687.1<br>WP_201828965.1<br>WP_101736539.1<br>WP_143336343.1<br>WP_115024034.1<br>WP_237801656.1<br>WP_316986356.1 | LMLIFQR<br>MPTASNALYQLER<br>VSTIAQAELIR | 29-35<br>72-84<br>61-71 | 18291 |

| | | | | | | |
|---|---|---|---|---|---|---|
| | | C. singulare | AJI80042.1 | | | |
| 7 | TetR/AcrR family transcriptional regulator | C. striatum | WP_086890869.1 | TLVDWLFK | 100-107 | 27822 |
| 8 | MULTISPECIES: TetR/AcrR family transcriptional regulator | Prescottella equi (formerly Rhodococcus (equi) | WP_268965648.1 | RLPPEER | 11-17 | 22053 |
| 9 | TetR/AcrR family transcriptional regulator [Corynebacterium glutamicum] | C. glutamicum | WP_003855609.1 | AGLSSTVLQR | 49-58 | 10184 |
| 10 | GntR family transcriptional regulator | C. aurimucosum | WP_193634551.1 | MLADSVASALR | 1-11 | 22176 |
| 11 | MULTISPECIES: CRP-like cAMP-activated global transcriptional regulator GlxR | Corynebacterium | WP_287136018.1 | GTTIFDEGEPGDR<br>LYIITSGK<br>MEGVQDILSR<br>TLLQLANR<br>TNANLADLIFTDVPGR<br>VNHDLTQEEIAQLVGASR | 37-49<br>50-57<br>1-10<br>156-163<br>137-152<br>174-191 | 25126 |
| 12 | MULTISPECIES: response regulator transcription factor | Corynebacterium | WP_062044319.1 | TEFALLQLLMENPR | 160-173 | 25892 |
| 13 | MerR family transcriptional regulator | C. glyciniphilum | WP_038547313.1 | EQVRIEK | 90-96 | 13759 |
| 14 | MerR family transcriptional regulator | C tuberculostearicum<br>C. marquesiae | WP_005327608.1<br>WP_239272892.1 | QKESLLAQR | 88-96 | 29668 |
| 15 | MULTISPECIES: bifunctional pyr operon transcriptional regulator/uracil phosphoribosyltransferase PyrR | Corynebacterium | WP_204581723.1 | IAHQIIEK | 24-31 | 20899 |
| 16 | Crp/Fnr family transcriptional regulator | C. variabile<br>C. terpenotabidum | HIW92058.1<br>WP_014010971.1<br>WP_020442086.1 | FGVQEGGALR<br>GTTIFDEGEPGDR<br>TNNALADLIFTDVPGR | 164-173<br>37-49<br>137-152 | 25024 |
| 17 | MULTISPECIES: response regulator transcription factor | Corynebacterium | WP_061924579.1 | VLVVDDEPNIVELLTVSLK | 154-169 | 26205 |
| 18 | MULTISPECIES: CRP-like cAMP-activated global transcriptional regulator GlxR | Corynebacterium (C. striatum, C. simulans) | WP_046645713.1 | AGIFQGVDPVAVQNLIEQMETVR<br>FGVQEGGALR<br>LYIITSGK<br>MEGVQDILSR<br>SVLIVDTEHLAK<br>TLLQLANR<br>TNANLADLIFTDVPGR<br>VNHDLTQEEIAQLVGASR | 11-33<br>164-173<br>50-57<br>1-10<br>213-224<br>156-163<br>137-152<br>174-191 | 25032 |
| 19 | RpiR family transcriptional regulator | Brevibacterium flavum | AKF27435.1 | CIEGLDTARR | 133-142 | 30559 |
| 20 | MULTISPECIES: TetR/AcrR family transcriptional regulator | Corynebacterium | WP_061921270.1 | LWSLVAAESR<br>VPDNAEETTLELLK<br>LLLTSGWNVGR | 96-105<br>204-217<br>106-116 | 25936 |

| | | | | | | |
|---|---|---|---|---|---|---|
| 21 | TetR/AcrR family transcriptional regulator | C. glyciniphilum | WP_038547077.1 | QDAVSSLAR | 86-94 | 39144 |
| 22 | IclR family transcriptional regulator | C. pseudodiphthereticum | WP_272730776.1 | AIAIMKATATTPMNLSELCTDTGIPR | 16-41 | 24801 |
| 23 | Metalloregulator ArsR/SmtB family transcription factor | C. glutamicum | WP_038583602.1 | ALSIGTQWAHVMGIK | 126-140 | 25246 |
| 24 | MULTISPECIES: division/cell wall cluster transcriptional repressor MraZ | Corynebacterium<br>C. striatum | WP_021353831.1<br>WP_086891054.1 | GQDHSLAVYPR<br>KAAAVSR<br>NLAASADEQRPDGHGR | 42-52<br>61-67<br>78-93 | 12381 |
| 25 | TetR family transcriptional regulator | C. variabile<br>C. riegelii | WP_312810083.1<br>WP_284858858.1 | VEMLLK | 203-208 | ? |
| 26 | MarR family winged helix-turn-helix transcriptional regulator | C. striatum<br>C. simulans | WP_046645631.1<br>WP_062035378.1 | AVPEHVESVR<br>ELCADLEWDR<br>LILGSIR<br>WLNEEEQELWR | 121-130<br>69-78<br>24-30<br>7-17 | 20509 |
| 27 | MarR family transcriptional regulator | C. striatum<br>C. flavescens | WP_005530667.1<br>WP_075728785.1 | ILAAIESDER<br>TTLAQLTDAGWDK<br>VVATAPGHAAQAR | 147-156<br>100-112<br>113-125 | 18312 |
| 28 | MULTISPECIES: transcription antitermination factor NusB | Corynebacterium | WP_239192056.1 | AADILYEAENR | 25-35 | 23546 |
| 29 | Transcription elongation factor GreA | C. striatum<br>C. renale<br>C. guaraldiae<br>C. maris<br>C. phoceense<br>C. callunae | MDU3174057.1<br>WP_115243562.1<br>WP_143335638.1<br>WP_020934313.1<br>WP_257053763.1<br>WP_015650808.1 | QISEVLANSTTER<br>QYITPETK<br>REEGDLKENAGYDAAREMQDQEEAR | 66-78<br>7-14<br>39-63 | 18711 |
| 30 | TetR/AcrR family transcriptional regulator | Prescottella equi | WP_084985939.1 | ARAELPA | 211-217 | 36758 |
| 31 | Response regulator transcription factor | C. striatum<br>C. simulans | WP_049064300.1<br>WP_275437323.1 | EIGQDLMLSEATVK | 171-184 | 22110 |
| 32 | TetR/AcrR family transcriptional regulator | C. halotolerans | WP_015400291.1 | SLAAEIVGEDDPR | 144-156 | 24277 |
| 33 | TetR/AcrR family transcriptional regulator | C. striatum | WP_110092290.1 | DQIATAALDLFDQR | 8-21 | 15043 |
| 34 | Transcription elongation factor GreA | C. stationis<br>C. massiliense<br>C. camporealensis<br>C. renale<br>C. kutscheri<br>C. kefirresidentii<br>C. ammoniagenes | WP_301200234.1<br>WP_027018614.1<br>WP_035105785.1<br>WP_111725846.1<br>WP_046440214.1<br>WP_284835692.1<br>WP_168938798.1 | AAASDNKDLETYSEQSPLGAAILGAQEGETR<br>QYITPETK<br>REEGDLKENAGYDAAREMQDQEEAR | 111-141<br>7-14<br>39-63 | 18898 |
| 35 | TetR/AcrR family transcriptional regulator | C. striatum<br>C. simulans<br>C. aurimucosum | WP_086890568.1<br>WP_248093794.1<br>WP_193629509.1 | GIVAACVEHR<br>GSHFLNAAGEYPRPETDAER | 127-136<br>107-126 | 22450 |
| 36 | CarD family transcriptional regulator | C. striatum | WP_049166643.1 | EIDVEEAGNWSR<br>QILVGELALASPVDEK | 76-87<br>133-148 | 21937 |

| | | | | | | |
|---|---|---|---|---|---|---|
| 37 | TetR/AcrR family transcriptional regulator | C. striatum<br>C. striatum | WP_049191721.1<br>WP_062036744.1 | DALVIAYVESLDEK<br>ELAIQLLSAPPADYSI<br>GSHFLNAAGEYPRPETDAER<br>LLESATNLFTTEGIR<br>VLAFFDK | 57-70<br>187-202<br>107-126<br>17-31<br>89-95 | 22540 |
| 38 | Response regulator transcription factor | C. striatum<br>C. simulans<br>C. accolens | WP_204083817.1<br>WP_248092860.1<br>PCC83691.1 | VFLVDDDPLVR | 5-15 | 23167 |
| 39 | Metal-dependent transcriptional regulator (Chain A, DIPHTHERIA TOXIN REPRESSOR) | C. simulans<br>C. aurimucosum<br>C. marquesiae<br>C. tuberculostearicum<br>C. macginleyi<br>C. accolens<br>C. flavescens<br>C. minutissimum<br>C. pseudodiphtheriticum<br>C. propinquum<br>C. striatum<br>C. diphtheriae | WP_248091815.1<br>WP_049361228.1<br>WP_349819523.1<br>WP_296179643.1<br>WP_200447185.1<br>WP_237791826.1<br>WP_075729948.1<br>WP_115022480.1<br>WP_249617266.1<br>WP_018119694.1<br>WP_166684278.1<br>WP_014316797.1 | DGLLHVR<br>LEQSGPTVSQTVAR<br>TIYELEEEGITPLR | 66-72<br>49-62<br>29-42 | 24975 |
| 40 | MarR family transcriptional regulator | C. striatum | WP_131771608.1 | AGLNQNEAAEELGISK | 143-158 | 20131 |
| 41 | YebC/PmpR family DNA-binding transcriptional regulator | C. falsenii | WP_025402776.1 | GDLTEDDLLMAVLDAGAEEVNDLGEK | 146-171 | 26824 |
| 42 | Response regulator transcription factor | C. phocae<br>C. propinquum<br>C. pseudodiphtheriticum<br>C. incognita<br>C. accolens<br>C. occultum<br>C. curieae<br>C. aurimucosum<br>C. kefirresidentii | WP_075732879.1<br>WP_284585581.1<br>WP_021354004.1<br>WP_185176514.1<br>WP_284641814.1<br>QGU08255.1<br>WP_269946840.1<br>WP_049360102.1<br>WP_200288656.1 | GGELIELSPTEFNLLR | 159-174 | 26205 |
| 43 | Helix-turn-helix transcriptional regulator | C. xerosis<br>C. amycolatum<br>C. vitaeruminis<br>C. pseudotuberculosis<br>C. ulcerans<br>C. lactis<br>C. diphtheriae | WP_120984396.1<br>WP_197914978.1<br>WP_025253717.1<br>WP_100210446.1<br>WP_095075753.1<br>WP_053412964.1<br>WP_014320511.1 | LGLVTPQR | 41-48 | 15681 |
| 44 | Lrp/AsnC family transcriptional regulator | C. variabile | WP_312775823.1 | SVGLSEGAAR | 29-38 | 16077 |
| 45 | Metal-sensitive transcriptional regulator | C. urealyticum | WP_012360560.1 | MDNAKPATK | 1-9 | 13375 |
| 46 | TetR/AcrR family transcriptional regulator | C. halotolerans | WP_015400417.1 | AEPAEALAVR | 122-131 | 21615 |
| 47 | MULTISPECIES: | Corynebacterium | WP_286955090.1 | IAPEVLDALR | 138-147 | 55463 |

| | | | | | | |
|---|---|---|---|---|---|---|
| | transcription antitermination factor NusB | | | | | |
| 48 | Transcriptional regulator, TetR-family | C. glyciniphilum | AHW62610.1 | MEDSTGMNHPRR | 1-12 | 25680 |
| 49 | FeoC-like transcriptional regulator | C. pyruviciproducens | WP_016458897.1 | SCAGNCSSCAISSRCGRGR | 52-70 | 8443 |
| 50 | IclR family transcriptional regulator | C. amycolatum | HJE84982.1 | MLPDADFNEQDLDDCR | 161-176 | 25715 |
| 51 | GntR-family transcription regulator | C. variabile | AEK37031.1 | MPDAGDAWEGGDAGAWAQR | 151-169 | 24346 |
| 52 | Transcription termination factor Rho | C. aurimucosum<br>C. kefirresidentii | WP_049361388.1<br>WP_301716185.1 | GDNKQRGNRHDDNNNHER | 175-192 | 68400 |
| 53 | WYL domain-containing protein, transcriptional regulator | C. ammoniagenes | WP_003845804.1 | LTNVLPYFR | 368-376 | 74489 |
| 54 | MarR family winged helix-turn-helix transcriptional regulator | C. halotolerans | WP_015400618.1 | DLCAQLEWDR | 63-72 | 19425 |
| 55 | Rv2640c family ArsR-like transcriptional regulator | Tsukamurella paurometabola | WP_013126866.1 | HGMSMHHSPRR | 92-102 | 12278 |
| 56 | TetR family transcriptional regulator | C. falsenii | WP_052337535.1 | ERLNPAEEADPAGDADDSTDR | 157-177 | 19944 |
| 57 | Helix-turn-helix transcriptional regulator | C. tuberculostearicum<br>C. kefirresidentii<br>C. aurimucosum<br>C. marquesiae<br>C. curieae | WP_316970898.1<br>WP_244169500.1<br>WP_049361230.1<br>WP_337887421.1<br>WP_269945351.1 | QGDAAAIPMMTGYRAVNAEGKICR | 73-96 | 28609 |
