## Supplemental Table S13 for "Peptidome Analysis of Western Blots Identifies Natural Bispecific Antibody-Bound *Corynebacterium* and Phage B-cell Epitopes with Potential Relevance to Psoriasis"

### Table S13. ATP Synthases Are *Corynebacterium* Antigens.

| Antigen Number | Identified Antigen | *Corynebacterium* (C.), *Mycobacterium* (M.) | Accession Number | Peptide Number | Peptide Sequence | Peptide Start/Stop | Molecular Weight |
|---|---|---|---|---|---|---|---|
| 1 | F0F1 ATP synthase subunit gamma | C. kefirresidentii | WP_259824363.1 | 1 | GMCGGYNNNVFK | 86-97 | 35937 |
| | | C. flavescens | WP_301505200.1 | | | | |
| | | C. confusum | WP_290225893.1 | | | | |
| | | C. casei | WP_006823371.1 | | | | |
| | | C. macginleyi | WP_200448729.1 | | | | |
| | | C. marquesiae | WP_269953383.1 | | | | |
| | | C. tuberculostearicum | WP_179386383.1 | | | | |
| | | C. accolens | WP_284899747.1 | | | | |
| | | C. pseudogenitalium | WP_005323379.1 | | | | |
| | | C. aurimucosum | WP_049361378.1 | | | | |
| | | C. curiae | WP_269945956.1 | | | | |
| | | C. endometrii | WP_136141012.1 | | | | |
| | | C. camporealensis | WP_046453334.1 | | | | |
| | | C. striatum | WP_086891555.1 | | | | |
| | | C. stationis | WP_283116714.1 | | | | |
| | | C. ammoniagenes | WP_003848681.1 | | | | |
| | | C. minutissimum | WP_115021733.1 | | | | |
| | | C. urogenitale | WP_151903035.1 | | | | |
| | | C. jeikeium | WP_005292974.1 | | | | |
| | | C. falsenii | WP_224207498.1 | | | | |
| | | C. resistens | WP_273351786.1 | | | | |
| | | C. bovis | WP_337891600.1 | | | | |
| | | C. auriscanis | WP_282941044.1 | | | | |
| | | C. dentalis | WP_312096626.1 | | | | |
| 2 | ATP synthase subunit beta | C. aurimucosum (1,4,6,7,8,9,10,14,15,17) | WP_193634369.1 | 1 | DGEQWGIHREPPAFDQLEGK | 121-140 | 52192 |
| | | C. minutissimum (1,4,6,7,8,9,14,15,16,17) | WP_115021734.1 | 2 | DVQNQDVLLFIDNIFR | 250-265 | |
| | | C. jeikeium (1,4,9) | WP_034985588.1 | 3 | EFSGTSVFAGVGER | 187-200 | |
| | | C. striatum (1,2,3,4,5,6,7,8,9,11,12,13,14,15,17) | WP_420693562.1 | 4 | FLGQNFFVAEK | 418-428 | |
| | | C. accolens (1,8,9,14,17) | WP_302519069.1 | 5 | FTQAGSEVSTLLGR Homo sapiens 310-322 | 266-279 | |
| | | C. simulans (1,2,3,5,6,7,9,11,12,13,14,15,16) | WP_062035306.1 | 6 | GAAVTDTGKPISVPVGDVVK | 83-102 | |
| | | C. singulare (1,4,6,7,8,9,14,17) | WP_239180726.1 | 7 | GHVFNALGDCLDQPGLGR | 103-120 | |
| | | C. guaraldiae (1,4,6,7,8,9,10,14,17) | WP_143335146.1 | 8 | GIYPAVNPLTSTSR | 348-361 | |
| | | C. argentoratense (1,4,8, 9,10,14,17) | WP_020976561.1 | 9 | IGLFGGAGVGK Homo sapiens 202-212 | 163-173 | |
| | | C. phoceense (1,4,7, 9,10,14,17) | WP_141628783.1 | 10 | KLTEKK | 477-482 | |
| | | C. lizhenjunii (1,4,7, 9,10,14,17) | WP_165009097.1 | 11 | MPSAVGYQPTLADEMGVLQER | 280-300 | |
| | | C. resistens (1,4,8, 9,14,17) | WP_013888779.1 | 12 | TEILETGIK | 141-149 | |
| | | C. marquesiae (1,8, 9,17) | WP_408920239.1 | 13 | TVLIQEMITR | 174-183 | |
| | | C. incognita (1,7,9,) | WP_185175769.1 | 14 | TVTLEVAQHLGDNLVR | 55-70 | |
| | | C. camporealensis (1,6,9,14) | WP_035105630.1 | 15 | VALSGLTMAEYFR | 237-249 | |
| | | C. endometrii (1,4,6,9,17) | WP_136141013.1 | 16 | VIDLLTPYVK | 150-159 | |
| | | C. stationis (1,4,9,17) | WP_066794684.1 | 17 | VIGPVVDVEFPR | 22-33 | |
| | | C. macginleyi (1,9,14) | WP_121910989.1 | | | | |
| | | C. urogenitale (1,4,9,17) | WP_151903034.1 | | | | |
| | | C. humireducens (1,9) | WP_040085615.1 | | | | |

| | | | |
|---|---|---|---|
| | | C. casei (1,9) | WP_276687390.1 |
| | | C. ammoniagenes (1,9,17) | WP_003848682.1 |
| | | C. curieae (1,9,14) | WP_269945955.1 |
| | | C. falsenii (1,9) | WP_025403050.1 |
| | | C. vitaeruminis (1) | WP_048758778.1 |
| | | C. amycolatum (1,9) | WP_256881411.1 |
| | | C. xerosis (1,6,9) | WP_102212289.1 |
| | | C. appendicis (2,3,5,9,10,12,13,15) | WP_234958702.1 |
| | | C. urinipleomorphum (2,3,5,9,10,12,13,15) | WP_235839392.1 |
| | | C. ureicelerivorans (2,3,5,9,15) | WP_273407041.1 |
| | | C. mucifaciens (2,3,5,9,10,12,13,15) | WP_246233731.1 |
| | | C. afermentans (2,3,5,9,11,12,13,15) | WP_228496370.1 |
| | | C. lipophiloflavum (2,3,5,9,11,12,13,15) | WP_006840807.1 |
| | | C. sanguinis (2,3,5,9,11,12,13,15) | WP_261369346.1 |
| | | C. lipophilum (2,3,5,9,12,13,15) | WP_252931753.1 |
| | | C. glaucum (2,3,5,9,11,12,13,15) | WP_290187036.1 |
| | | C. bovis (2,3,4,9,11,12,13,15) | WP_010267175.1 |
| | | C. massiliense (2,3,5,9,10,11,12,13,17) | WP_022862464.1 |
| | | C. callunae (2,3,4,6,8,9) | WP_015651002.1 |
| | | C. phocae (2,4,8,9,17) | WP_075734587.1 |
| | | C. glutamicum (2,3,4,5,8,9,11,12,13,15) | WP_044029907.1 |
| | | C. deserti (2,4,6,8,9) | WP_053544660.1 |
| | | C. propinquum (2,5) | WP_302500938.1 |
| | | C. pseudodiphtheriticum (2,9) | WP_249618623.1 |
| | | M. tuberculosis (2,5,9,11,13) | MBP0573160.1 |
| | | M. leprae (2,5,9,11,13) | WP_010908163.1 |
| | | C. auris (2,3,5,9,11,12,13,15) | WP_290343051.1 |
| | | C. otitidis (3,5,9,12,13,15) | WP_046643558.1 |
| | | C. mastitides (3,5,8,9,12,13,15) | WP_337890130.1 |
| | | C. diphtheriae (3,4,5,8,9,10,11,12,13,15,16,17) | WP_191652101.1 |
| | | C. cystitidis (4,5,8,9,11,13,15) | WP_257158664.1 |
| | | C. renale (4,8,9) | WP_048380569.1 |
| | | C. epidermidicanis (4,8,9,14,17) | WP_047240008.1 |
| | | C. urealyticum (4,8,9,14,17) | WP_149121548.1 |
| | | C. matruchotii (4,8,9,17) | WP_314973954.1 |
| | | C. canis (4,9,17) | WP_146323311.1 |
| | | C. incognita (4,9,17) | WP_185175769.1 |
| | | C. dentalis (4,9,17) | WP_312096624.1 |
| | | C. halotolerans (4,8,9) | WP_027004461.1 |
| | | C. pseudotuberculosis (4,5,8,9,11,12,13,15,16) | WP_369336359.1 |
| | | C. nasicanis (4,9) | WP_377000993.1 |
| | | C. breve (4,9) | WP_284826305.1 |
| | | C. comes (4,17) | WP_156227717.1 |
| | | C. occultum (4,6,9,17) | WP_156230629.1 |
| | | C. imitans (5,9,11,13) | WP_239244269.1 |
| | | C. oculi (8,9,14) | WP_055122938.1 |
| | | C. variabile (8,9,17) | WP_313094903.1 |
| | | C. terpenotabidum (8,9,17) | WP_041631147.1 |
| | | C. efficiens (8,9) | WP_006769276.1 |

| | | | | | | | |
|---|---|---|---|---|---|---|---|
| | | C. fournieri (2,9,11,12,13,15)<br>C. atypicum (9,14)<br>C. hindlerae (9,14,17)<br>C. spheniscorum (9,17)<br>C. nuruki (9,17)<br>C. glyciniphilum (9,17) | WP_230471720.1<br>WP_038605820.1<br>WP_038605820.1<br>WP_092285318.1<br>WP_010119239.1<br>WP_038547745.1 | | | | |
| 3 | F0F1 ATP synthase subunit gamma | C. simulans (1,2,3,4)<br>C. striatum (1,2,3,4)<br>C. confusum (1,2,5)<br>C. endometri (1,2,5)<br>C. singulare (1,5)<br>C. minutissimum (1,5)<br>C. phoceense (1)<br>C. kefirresidentii (2)<br>C. flavescens (2,5)<br>C. pseudotuberculosis (2)<br>C. marquesiae (2)<br>C. tuberculostearicum (2)<br>C. casei (2,5)<br>C. macginleyi (2)<br>C. yonathiae (2)<br>C. accolens (2)<br>C. pseudogenitalium (2)<br>C. aurimucosum (2)<br>C. curieae (2)<br>C. camporealensis (2,5)<br>C. matruchotii (2)<br>C. stationis (2,5)<br>C. durum (2)<br>C. diphtheria (2)<br>M. tuberculosis (2)<br>C. incognita (5)<br>C. ammoniagenes (5)<br>C. kutscheri (5)<br>C. vitaeruminis (5) | WP_062041127.1<br>WP_046645539.1<br>WP_290225893.1<br>WP_136141012.1<br>WP_239180724.1<br>WP_115021733.1<br>WP_257035171.1<br>WP_259824363.1<br>WP_301505200.1<br>WP_137431671.1<br>WP_198492897.1<br>WP_005327646.1<br>WP_006823371.1<br>WP_200448729.1<br>WP_238801024.1<br>WP_284899747.1<br>WP_005323379.1<br>WP_049361378.1<br>WP_269945956.1<br>WP_368530440.1<br>WP_311166298.1<br>WP_191372946.1<br>WP_315497725.1<br>WP_088245939.1<br>WP_128239442.1<br>WP_185175768.1<br>WP_003848681.1<br>WP_126369418.1<br>WP_025252589.1 | 1<br>2<br>3<br>4<br>5 | AAELEALLK<br>AQELIATSR<br>EADVAGSWAGFSQDPSWETTHDVR<br>KQGYETVR<br>QAQITQEITEIVGGAGALAESAESD | 99-107<br>23-31<br>130-153<br>108-115<br>303-327 | 35986 |
| 4 | F0F1 ATP synthase subunit delta | C. striatum (1,2,3)<br>C. accolens (1,2,3)<br>C. simulans (1,2,3) | WP_239297927.1<br>PCC82908.1<br>WP_284841267.1 | 1<br>2<br>3 | GAEAQGQLGQVEDELFR<br>NEAELTQLLSDR<br>VGDEVIDGSTAGK | 112-128<br>135-146<br>250-262 | 29036 |
| 5 | F0F1 ATP synthase subunit alpha | C. accolens (1,2,3,4)<br>C. macginleyi (1,2,3)<br>C. argentoratense (1,2,3,4)<br>C. phocae (1,2,3)<br>C. canis (1,2,3,4)<br>C. kutscheri (1,2,3,4)<br>C. diphtheria (1,2,3,4)<br>C. mastitidis (1,2,3,4)<br>C. belfantii (1,2,3,4)<br>C. rouxii (1,2,3,4) | WP_035108618.1<br>WP_200449084.1<br>WP_234870741.1<br>WP_075734591.1<br>WP_146323309.1<br>WP_046440010.1<br>WP_196982430.1<br>WP_018118675.1<br>WP_197694888.1<br>WP_342295177.1 | 1<br>2<br>3<br>4<br>5 | AELTISSDEIR<br>AIDAMTPIGR<br>EAYPGDVFYLHSR<br>LSDDMGAGSMTALPIIETK<br>QPVEEPLQTGMK | 2-12<br>155-164<br>295-307<br>315-333<br>143-154 | 59594 |

| | | | | | | | |
|---|---|---|---|---|---|---|---|
| | | C. renale (1,2,3,4)<br>C. endometrii (1,2,3,4)<br>C. striatum (1,2,3,4)<br>C. durum (1,2,3,4)<br>C. guaraldiae (1,2,3,4)<br>C. glucuronolyticum (1,2,3,4,5)<br>C. simulans (1,2,3,4)<br>M. tuberculosis (2,3)<br>C. urinipleomorphum (1,2,3)<br>C. appendicis (1,2,3)<br>C. glutamicum (1,2,3)<br>C. pilosum (1,3)<br>C. kroppenstedtii (1,2,3,5)<br>C. cystitidis (1,2,3)<br>C. bovis (1,2,3,5)<br>C. breve (1,2,3)<br>C. pseudodiphtheriticum (1.2,3)<br>C. uterequi (1,2,3)<br>C. pseudogenitalium (1,2,3,4)<br>C. ammoniagenes (1,2,3,5)<br>C. resistens (1,2,3)<br>C. propinquum (1,2,3)<br>C. auris (1,2,3,5)<br>C. maris (1,3)<br>C. jeikeium (1,2,3)<br>C. sanguinis (1,2,3)<br>C. riegelii (1,2,3)<br>C. glaucum (1,2,3)<br>C. pseudotuberculosis (1,2,3)<br>C. efficiens (2,3)<br>C. matruchotii (3)<br>C. incognita (3,4)<br>C. kefirresidentii (3,4)<br>C. confusum (3,4)<br>C. marquesiae (3,4)<br>C. lizhenjunii (3,4)<br>C. flavescens (3,4)<br>C. tuberculostearicum (3,4)<br>C. aurimucosum (3,4)<br>C. afermentans (3,5)<br>C. ureicelerivorans (3,5)<br>C. fourneri (3,5)<br>C. imitans (3,5)<br>C. pyruviproducens (3,5) | WP_048380565.1<br>WP_136141011.1<br>WP_049062705.1<br>WP_315497731.1<br>WP_154762087.1<br>MCI6206902.1<br>WP_339019359.1<br>WP_031683680.1<br>WP_087116794.1<br>WP_076598388.1<br>NII87562.1<br>WP_018580842.1<br>WP_303974423.1<br>WP_092257116.1<br>WP_010267177.1<br>WP_284826921.1<br>WP_284845552.1<br>WP_082121263.1<br>WP_256006507.1<br>EFG80680.1<br>AEI09771.1<br>WP_081604909.1<br>WP_290343049.1<br>WP_084481981.1<br>EEW17517.1<br>WP_259812643.1<br>WP_311342121.1<br>WP_290186723.1<br>WP_323965334.1<br>WP_303735018.1<br>MFC2726246.1<br>WP_185175767.1<br>WP_086588745.1<br>WP_290226272.1<br>WP_408921153.1<br>WP_165009093.1<br>WP_312713863.1<br>WP_316999904.1<br>WP_102234375.1<br>WP_063937388.1<br>WP_038611100.1<br>WP_085956613.1<br>WP_038589986.1<br>WP_016457263.1 | | | | |
| 6 | F0F1 ATP synthase subunit B | C. flavescens (2,3,4,5,6,7)<br>C. striatum (1,2,3,4,6,7)<br>C. ammoniagense (1,2,3,4,6,7)<br>C. casei (1,2,3,4,5,6,7)<br>C. kefirresidentii(1,2,3,4,5,6) | WP_276623457.1<br>WP_005532053.1<br>WP_003848678.1<br>WP_276687397.1<br>WP_200287689.1 | 1<br>2<br>3<br>4<br>5 | AALEKYNAQLADAR<br>AEAAEIR<br>GKQIEAEAK<br>RAEAQQAEAK<br>SDIGQNSINLAEK | 80-93<br>94-100<br>107-115<br>70-79<br>146-158 | 19035 |

| | | | | | | | |
|---|---|---|---|---|---|---|---|
| | | C. marquesiae(1,2,3,4,5,6)<br>C. tuberculostearicum(1,2,3,4,5,6)<br>C. stationis(1,2,3,4,6,7)<br>C. phoceense(1,2,4,6)<br>C. simulans (1,2,3,4,5,6,7) | WP_049378381.1<br>WP_301431952.1<br>WP_066794696.1<br>WP_257035174.1<br>WP_061921968.1 | 6<br>7 | YNAQLADAR<br>YNAQLADARAEAAEIR | 85-93<br>85-100 | |
| 7 | F0F1 ATP synthase subunit beta | C. urealyticum(1,2)<br>C. accolens(1,2)<br>C. stationis(1,2)<br>C. striatum(1,2)<br>C. kefirresidentii(1,2)<br>C. simulans(1,2) | WP_149121548.1<br>WP_284902074.1<br>WP_066794684.1<br>WP_306495854.1<br>WP_239205706.1<br>AMO90719.1 | 1<br>2 | TVSMAPTDGLVR<br>VIGPVVDVEFPR | 71-82<br>22-33 | 52280 |

Peptide sequences highlighted in green are conserved and identical in *M. tuberculosis* and *M. leprae.*
