## Supplemental Table S14 for "Peptidome Analysis of Western Blots Identifies Natural Bispecific Antibody-Bound *Corynebacterium* and Phage B-cell Epitopes with Potential Relevance to Psoriasis"

### Table S14. Respiratory Chain Proteins are *Corynebacterium* Antigens.

| Antigen Number | Identified Antigen | *Corynebacterium* (C.), *Actinomycetota* (Peptide numbers) | Accession Number | Peptide Number | Peptide Sequence | Peptide Start/Stop | Molecular Weight |
|---|---|---|---|---|---|---|---|
| 1 | NADP-dependent isocitrate dehydrogenase | C. simulans (1-23) | WP_062042093.1 | 1 | AAFDAALENGPDLAMVNSAK | 298-317 | 78434 |
| | | C. striatum (1-23) | WP_239299420.1 | 2 | AGVLADALDK | 607-616 | |
| | | C. uropygiale (5,12) | WP_236117776.1 | 3 | AGVLADALDKATETLLDEGNSPSR | 607-630 | |
| | | C. ammoniagenes (5) | WP_040356380.1 | 4 | AGVLADALDKATETLLDEGNSPSRK | 607-631 | |
| | | C. phocae (5) | WP_075735397.1 | 5 | ALDEFLK | 214-220 | |
| | | C. singulare (5,13) | WP_144791235.1 | 6 | ANGAFDPSTMGTVPNVGLMAQK | 376-397 | |
| | | C. guaraldiae (5,13) | WP_144014208.1 | 7 | ATETLLDEGNSPSR | 617-630 | |
| | | C. hiratae (5) | WP_158397697.1 | 8 | ATETLLDEGNSPSRK | 617-631 | |
| | | C. aurimucosum (5,11,13,17,20) | WP_193629289.1 | 9 | DAPIQDWVK | 444-452 | |
| | | C. faecipullorum (5,13) | HIX78501.1 | 10 | DYNTDLFPILELGTSAK | 529-545 | |
| | | C. minutissimum (5) | WP_239189144.1 | 11 | HVQQVEEENHLR | 570-581 | |
| | | C. intestinale (5) | WP_250224616.1 | 12 | ILAQFPER | 46-53 | |
| | | C. hesseae (5,13) | WP_339017218.1 | 13 | KGEDTISVTGNVLR | 515-528 | |
| | | C. stationis (5) | WP_279192432.1 | 14 | LPNISASLVQLK | 80-91 | |
| | | C. casei (5) | WP_098072660.1 | 15 | MLSVVPLMAGGGLFETGAGGSAPK | 546-569 | |
| | | C. durum (9,10,15,16,19,24,25) | WP_179418879.1 | 16 | RAPIAVK | 134-140 | |
| | | C. felinum (9,10,16,19,25) | WP_277104959.1 | 17 | TIFWLDPER | 465-473 | |
| | | C. pseudotuberculosis (9,10,15,16,19,21,22) | ANK55852.1 | 18 | TNVATMSDNDFR | 159-170 | |
| | | C. ulcerans (9,10,15,16,19,21,22) | AKA96058.1 | 19 | TPEANIIK | 72-79 | |
| | | C. kutscheri (9,10,16,21,22,25) | WP_126369586.1 | 20 | TPEANIIKLPNISASLVQLK | 72-91 | |
| | | C. efficiens (9,10) | BAC17492.1 | 21 | TSGHMWNK | 340-347 | |
| | | C. confusum (9,11,14,17,21,23) | WP_290224916.1 | 22 | VSDPIIFGHVVR | 244-255 | |
| | | C. glutamicum (9,11,12,16,19,21,22,23) | WP_011013800.1 | 23 | WDSLGEFLALAESFR | 582-596 | |
| | | C. phoceense (9,11,13,14,16,17,19,20,21,23) | WP_303938491.1 | 24 | EGKDTISVTGNVLR | 577-590 | |
| | | C. deserti (9,12,16,19,21,22,23) | WP_053544252.1 | 25 | GITNLHVPSDVIVDASMPAMIR | 380-401 | |
| | | C. pollutisoli (9,15,19) | WP_085549516.1 | | | | |
| | | C. callunae (9,12,16,18,19,21,23) | WP_015650502.1 | | | | |
| | | C. epidermidicanis (9,16,19,22,25) | WP_047239640.1 | | | | |
| | | C. crudilactis (9,16,19,21,22) | WP_066564793.1 | | | | |
| | | C. suedecumii (9,19) | WP_284875995.1 | | | | |
| | | C. humireducens (9,19) | WP_040084995.1 | | | | |
| | | C. hindlerae (9,16,19,22) | WP_208768755.1 | | | | |
| | | C. massiliense (9,14,19,21,22) | WP_022862660.1 | | | | |
| | | C. vitaeruminis (9,12,15,16,19) | WP_025252008.1 | | | | |
| | | C. occultum (9,16,19) | WP_156230125.1 | | | | |
| | | C. halotolerans (9,19) | WP_015400059.1 | | | | |
| | | C. matruchotii (9) | WP_315556010.1 | | | | |
| | | C. flavescens (9,11,19,20,21) | WP_276623521.1 | | | | |
| | | C. marquesiae (9,11,17,19,20) | WP_340419316.1 | | | | |
| | | C. sanguinis (10,14,22) | WP_259811241.1 | | | | |
| | | C. lipophiloflavum (10,14) | WP_006839541.1 | | | | |
| | | C. genitalium (10,14,15) | WP_005286416.1 | | | | |
| | | C. amycolatum (12,15) | WP_284825822.1 | | | | |

| | | | | | | | |
|---|---|---|---|---|---|---|---|
| | | C. lactis (11,12,15) | WP_053411537.1 | | | | |
| | | C. mastitidis (12,19,21,22) | WP_101173567.1 | | | | |
| | | C. atypicum (12,15) | WP_038604576.1 | | | | |
| | | C. oculi (12,19,21,22) | WP_055122176.1 | | | | |
| | | C. macginleyi (12,17,20) | WP_121953198.1 | | | | |
| | | C. incognita (12) | WP_185175400.1 | | | | |
| | | C. dentalis (13,25) | WP_185175400.1 | | | | |
| | | C. propinquum (13,14,15,23) | WP_249606005.1 | | | | |
| | | C. kroppenstedtii (13,23) | WP_303734223.1 | | | | |
| | | C. pseudokroppenstedtii (13,23) | WP_204087418.1 | | | | |
| | | C. parakroppenstedtii (13,23) | WP_340055899.1 | | | | |
| | | C. argentoratense (13,16,18,19,22) | WP_021012075.1 | | | | |
| | | C. singulare (13) | WP_144791235.1 | | | | |
| | | C. pseudodiphtheriticum (14,23) | WP_027017573.1 | | | | |
| | | C. auris (14) | WP_290342669.1 | | | | |
| | | C. canis (15,23) | WJY74380.1 | | | | |
| | | C. xerosis (11,15) | WP_406913038.1 | | | | |
| | | C. ulceribovis (11,15) | WP_018023403.1 | | | | |
| | | C. efficiens (15,19,23) | BAC17492.1 | | | | |
| | | C. frankenforstensis (15, | WP_075663334.1 | | | | |
| | | C. anserum (19,24,25) | WP_185770262.1 | | | | |
| | | C. lizhenjunii (6) | WP_165009828.1 | | | | |
| | | C. tuberculostearicum (11,17,20) | WP_316991337.1 | | | | |
| | | C. kefirresidentii (11,17,20) | WP_301732308.1 | | | | |
| | | C. curieae (11) | WP_269946372.1 | | | | |
| | | C. diphtheria (9,10,15,16,19,22) | WP_259605522.1 | | | | |
| | | C. belfantii (16) | WP_410494605.1 | | | | |
| | | C. rouxii (9,16) | WP_315643693.1 | | | | |
| | | C. accolens (17,20) | WP_284610022.1 | | | | |
| | | C. pseudogenitalium (20) | WP_256006196.1 | | | | |
| | | C. variabile (21,22,25) | WP_303944548.1 | | | | |
| | | C. uropygiale (18) | WP_236117776.1 | | | | |
| | | C. faecigallinarum (22) | HJC85449.1 | | | | |
| | | C. glyciniphilum (22) | WP_145943122.1 | | | | |
| | | C. auriscanis (25) | WP_282940993.1 | | | | |
| | | C. terpenotabidum (25) | WP_020441818.1 | | | | |
| | | C. urealyticum (25) | WP_012359678.1 | | | | |
| | | C. resistence (25) | AEI10135.1 | | | | |
| | | C. macclintockiae (25) | WP_269955084.1 | | | | |
| | | C. jeikeium (25) | WP_005291732.1 | | | | |
| | | C. bovis (25) | WP_125173488.1 | | | | |
| 2 | Ubiquinol-cytochrome c reductase iron-sulfur subunit | C. simulans (1,2,3,4,5,6) | WP_061923549.1 | 1 | ALPQLPIGVDEEGYLVAK | 373-390 | 45034 |
| | | C. striatum | WP_166684805.1 | 2 | GNFIEPVGPAFWER | 391-404 | |
| | | C. diphtheria (2,3,5) | WP_010935205.1 | 3 | ICTHIGCPTSLYEAQTNR | 330-347 | |
| | | C. ulcerans | AKA97057.1 | 4 | MSNDELAALGTELDDVTVAFR | 16-36 | |
| | | C. pseudotuberculosis | AKS13762.1 | 5 | NAVMLIR | 295-301 | |
| | | C. ramonii | AIU33083.1 | 6 | TLTALLNDSWK | 142-152 | |
| | | C. glutamicum | WP_074506856.1 | | | | |
| | | C. callunae | WP_015651716.1 | | | | |

| | | | | | | | |
|---|---|---|---|---|---|---|---|
| | | C. gallinarum<br>C. efficiens<br>C. deserti<br>C. sanguinis<br>C. auris<br>C. vitaeruminis<br>C. marquesiae | WP_191732212.1<br>WP_006768084.1<br>WP_053545314.1<br>WP_259810789.1<br>WP_290341456.1<br>WP_034648487.1<br>WP_337888181.1 | | | | |
| 3 | Cytochrome bc complex cytochrome b subunit | C. simulans (1,2,3)<br>C. halotolerans<br>C. renale<br>C. efficiens<br>C. kutscheri<br>C. diphtheria<br>C. striatum<br>C. marquesiae<br>C. stationis<br>C. casei<br>C. aurimucosum<br>C. pseudotuberculosis<br>C. matruchotii<br>C. lizhenjunii<br>C. minutissimum | WP_062043913.1<br>WP_027004078.1<br>WP_048379400.1<br>WP_011075748.1<br>WP_046439943.1<br>WP_010935204.1<br>WP_306592346.1<br>WP_337888180.1<br>WP_313678661.1<br>WP_276686157.1<br>WP_049361052.1<br>WP_160186677.1<br>WP_315555662.1<br>WP_165010261.1<br>WP_115022928.1 | 1<br>2<br>3 | AENNVVGVR<br>EVLEHGIETGVIK<br>YSAAGVLR | 239-247<br>434-446<br>16-23 | 60093 |
| 4 | Cytochrome c oxidase subunit II | C. simulans<br>C. kefirresidentii<br>C. striatum<br>C. tuberculostearicum<br>C. marquesiae<br>C. curiae<br>C. macginleyi<br>C. accolens | WP_282440012.1<br>WP_282438215.1<br>WP_086891036.1<br>WP_301980248.1<br>WP_284786815.1<br>WP_269945552.1<br>WP_121952995.1<br>WP_284640541.1 | 1<br>2 | DVYAHPEANAQER<br>RDVYAHPEANAQER | 254-266<br>253-266 | 39764 |
| 5 | Cytochrome c oxidase subunit I | C. striatum<br>C. simulans<br>C. aurimucosum<br>C. singulare<br>C. minutissimum<br>C. intestinale<br>C. guaraldiae | WP_201816471.1<br>WP_061924416.1<br>WP_010191271.1<br>WP_239179294.1<br>WP_039676739.1<br>WP_250224189.1<br>WP_143335362.1 | 1<br>2 | HNFVSLPR<br>LDDYVAPTRPEPTGNAK | 529-536<br>8-24 | 62790 |
| 6 | Succinate dehydrogenase cytochrome b subunit | C. glutamicum<br>C. minutissimum<br>C. tuberculostearicum<br>C. kefirresidentii | WP_172768109.1<br>WP_115021072.1<br>WP_316986120.1<br>WP_086589111.1 | 1 | YGAFLR | 664-669 | 73471 |
| 7 | Succinate dehydrogenase cytochrome b subunit | C. simulans (1,2,5)<br>C. minutissimum<br>C. striatum (1,2,3,4,5)<br>C. aurimucosum<br>C. singulare | WP_239239278.1<br>WP_115021072.1<br>WP_306588472.1<br>WP_012714792.1<br>WP_239180312.1 | 1<br>2<br>3<br>4<br>5 | ANMIATFSR<br>LAASDLGITGAK<br>NPDREALAHGR<br>SVKNPDREALAHGR<br>TNLMGGLDSFATK | 175-183<br>207-218<br>5-15<br>2-15<br>124-136 | 27304 |
| 8 | Cytochrome c oxidase assembly protein | C. efficiens | WP_006768357.1 | 1 | TVDGVNDLR | 76-84 | 81283 |

| | | | | | | | |
|---|---|---|---|---|---|---|---|
| 9 | Cytochrome c oxidase subunit I | C. striatum<br>C. minutissimum<br>C. simulans<br>C. aurimucosum<br>C. singulare | WP_086890794.1<br>WP_039676739.1<br>WP_061924416.1<br>WP_010191271.1<br>WP_042532128.1 | 1<br>2 | HNFVSLPR<br>LDDYVAPTRPEPTGNAK | 529-536<br>8-24 | 62790 |
| 10 | Cytochrome c oxidase subunit II | C. jeikeium<br>C. macclintockiae<br>C. renale<br>C. efficiens<br>C. durum<br>C. glutamicum<br>C. breve<br>C. comes<br>C. marinum<br>C. humireducens<br>C. confusum<br>C. maris<br>C. halotolerans<br>C. mastitides | WP_034965192.1<br>WP_284803414.1<br>PFG28370.1<br>BAC18897.1<br>EKX89802.1<br>WP_208400630.1<br>WP_284823783.1<br>WP_156228525.1<br>WP_042621783.1<br>WP_040086260.1<br>WP_290222508.1<br>WP_041631850.1<br>WP_015401347.1<br>WP_101172919.1 | 1 | EGAFVGRCAEMCGTYHAMMNFELR | 285-308 | 36902 |
| 11 | Cytochrome c oxidase assembly protein | C. halotolerans | WP_149029434.1 | 1 | VARGEDDEFESYNQMLARMNAGGE QR | 670-695 | 75011 |
| 12 | Iron-sulfur succinate dehydrogenase subunit | Micrococcus<br>Brevibacterium<br>Clavibacter michiganensis<br>M.tuberculosis<br>M.leprae<br>C. cyclohexanicum | WP_224355525.1<br>WP_069600613.1<br>WP_104294766.1<br>MBZ4296222.1<br>WP_010907879.1<br>WP_229232118.1 | 1 | SCAHGVCGSDAMRINGRNR | 83-101<br><br><br><br>54-72 | 26417 |
| 13 | Fumarate reductase/succinate dehydrogenase flavoprotein subunit | C. casei<br>C. stationis<br>C. ammoniagenes<br>C. minutissimum<br>C. singulare<br>C. aurimucosum<br>C. intestinale<br>C. guaraldiae<br>C. flavescens<br>C. marquesiae<br>C. striatum<br>C. phocae<br>C. simulans<br>C. camporealensis<br>C. confusum | WP_301437437.1<br>WP_075722276.1<br>WP_003846793.1<br>WP_115021073.1<br>WP_239180314.1<br>WP_216382020.1<br>WP_250224029.1<br>WP_154736750.1<br>WP_312714870.1<br>WP_284791618.1<br>WP_284790122.1<br>WP_075732558.1<br>WP_248093517.1<br>WP_321115301.1<br>WP_290224415.1 | 1<br>2<br>3<br>4<br>5<br>6<br>7<br>8<br>9<br>10<br>11 | EYGGTLATR<br>GQTGQQLQLSTTSALYR<br>HAEPLYFDSIPLMTR<br>ILESHEPHGVPMK<br>ITGNANDMNQVLEYGLR<br>KVDNDSAYR<br>NLITGELK<br>NSNAGAMMR<br>VIDHMNAIGAPFAR<br>YPAFGNLVPR<br>YSNLIEMYEEAIGESAYETPMR | 173-181<br>196-212<br>663-677<br>37-49<br>585-601<br>123-131<br>248-255<br>281-289<br>159-172<br>361-370<br>414-435 | 75852 |
| 14 | NAD-dependent succinate-semialdehyde dehydrogenase | C. simulans<br>C. striatum | WP_284841480.1<br>WP_284806951.1 | 1<br>2<br>3<br>4<br>5<br>6 | AFELVTER<br>AQDNVAALVEDAVAK<br>GEGAGYFYEPTVLTNVSR<br>KVSFTGSTPVGK<br>NIGEACTAANR<br>SASAISEPVMADSR | 82-89<br>340-354<br>365-382<br>232-243<br>288-298<br>216-229 | 52196 |

| | | | | | | | |
|---|---|---|---|---|---|---|---|
| | | | | 7<br>8<br>9<br>10 | SLPAPEGTLQMVTR<br>TINVEQLLAK<br>VGNGMDEGVTCGPLIEK<br>WFSEEAVR | 133-146<br>2-11<br>322-338<br>121-128 | |
| 15 | NAD-dependent succinate-semialdehyde dehydrogenase | C. tuberculostearicum<br>C. aurimucosum<br>C. kefirresidentii<br>C. marquesiae | WP_200435610.1<br>WP_049359949.1<br>WP_239206630.1<br>WP_284788802.1 | 1<br>2<br>3<br>4<br>5<br>6 | AEVQIVADIYR<br>FAAPNLLLGNTIVLK<br>HASICPLSSQACQDILEEAGLPK<br>LIVLEDFYDR<br>MYNTGQACNAPK<br>WGVGEFVNEHLYR | 81-91<br>140-154<br>155-177<br>268-277<br>255-266<br>441-453 | 51229 |
| 16 | NAD-dependent succinate-semialdehyde dehydrogenase | Tsukamurella paurometabola | WP_245537894.1 | 1 | DAARVQRVGAGLRSGMVGVNR | 422-442 | 49706 |
| 17 | succinic semialdehyde dehydrogenase | C. maris | WP_020933498.1 | 1 | QDELMDVIQDESGKNR | 63-78 | 52144 |
| 18 | Fe-S cluster assembly protein SufB | C. amycolatum<br>C.diphtheriae | WP_005510784.1<br>WP_014308346.1 | 1<br>2 | GLAEEEAMAMIVR<br>INTENMGQFER | 438-450<br>220-230 | 53863 |
| 19 | Fe-S cluster assembly protein SufD | C. accolens<br>C. tuberculostearicum | WP_284894033.1<br>WP_316980693.1 | 1 | FDDIELFYLMSR | 328-339 | 42822 |
| 20 | NAD(P)H-quinone dehydrogenase | C. felinum (1,2,5)<br>C. durum (1,5)<br>C. diphtheria (1,5)<br>C. phoceense (1,2,3,4,5,7)<br>C. striatum (1,2,3,4,5,6,7)<br>C. lizhenjunii (1,5,6,7)<br>C. simulans (1,2,3,4,5,6,7)<br>C. flavescens (1,5)<br>C. efficiens (1,5)<br>C. singulare (1,3,5,7)<br>C. minutissimum (1,5,7)<br>C. guaraldiae (1,3,5,7)<br>C. hiratae (1,3,5,7)<br>C. faecipullorum (1,5,7)<br>C. aurimucosum (1,3,4,5,6,7)<br>C. incognita (1,2)<br>C. matruchotii (2)<br>C. confusum (2,4,6)<br>C. accolens (2,3,4,6,7)<br>C. pseudotuberculosis (4)<br>C. ulcerans (4)<br>C. matruchotii (4)<br>C. frankenforstense (4)<br>C. marquesiae (4,6)<br>C. tuberculostearicum (4,6)<br>C. propinquum (4)<br>C. camporealense (6)<br>C. pseudogenitalium (6)<br>C. casei (6)<br>C. stationis (6) | WJY94203.1<br>GAA1417466.1<br>WP_041627805.1<br>WP_371132873.1<br>WP_309221918.1<br>WP_196823561.1<br>WP_284841020.1<br>WP_075729235.1<br>BAC17518.1<br>WP_144791219.1<br>WP_039673950.1<br>WP_348522051.1<br>WP_328287847.1<br>HIX78515.1<br>WP_216380018.1<br>WP_185175420.1<br>WP_005519938.1<br>WP_290224968.1<br>PCC83540.1<br>AEK91806.1<br>WP_196296466.1<br>WP_005519938.1<br>WP_075663352.1<br>WP_337891323.1<br>WP_301438124.1<br>PZQ26887.1<br>WP_368531136.1<br>WP_005322584.1<br>WP_301525513.1<br>WP_278829194.1 | 1<br>2<br>3<br>4<br>5<br>6<br>7 | DRILPHDDADAADVLETVLAER<br>IAMYHALGEGVSPLR<br>IGVNVIDGR<br>ILPGAQPDGER<br>ILPHDDADAADVLETVLAER<br>RLVAHDDLE<br>SFIAGANLR | 225-246<br>345-359<br>116-124<br>168-178<br>227-246<br>474-482<br>61-69 | 50754 |

| | | | |
|---|---|---|---|
| | | C. massiliense (6)<br>C. ammoniagenes (6) | WP_022862643.1<br>WP_168939080.1 |
