## Supplemental Table S15 for "Peptidome Analysis of Western Blots Identifies Natural Bispecific Antibody-Bound *Corynebacterium* and Phage B-cell Epitopes with Potential Relevance to Psoriasis"

### Table S15. Glycolysis Enzymes are *Corynebacterium* Antigens.

| Antigen Number | Identified Antigen | *Corynebacterium* (C.), other *Actinomycetota* | Accession Number | Peptide Sequence | Peptide Start/Stop | Molecular Weight |
|---|---|---|---|---|---|---|
| 1 | Phosphomannomutase/phosphogluco mutase | C. aurimucosum<br>C. deserti<br>C. macginleyi<br>C. striatum<br>C. accolens<br>C. minutissimum<br>C. simulans<br>C. stationis<br>C. flavescens<br>C. hiratae<br>C. singulare<br>C. guaraldiae<br>C. casei<br>C. faecipullorum | WP_193629999.1<br>WP_053544317.1<br>WP_200437479.1<br>WP_049146850.1<br>WP_284809265.1<br>WP_039674179.1<br>WP_248092270.1<br>WP_301205178.1<br>WP_312714387.1<br>WP_144013121.1<br>WP_239180564.1<br>WP_154762259.1<br>MDO5507798.1<br>HIX78477.1 | ADIGLAFDGDADR<br>ASNTEPLLR<br>GTDAWFNVR<br>GVVGEDIDENFVR | 237-249<br>423-431<br>414-422<br>18-30 | 48710 |
| 2 | 6-Phosphogluconolactonase | C. ammoniagenes<br>C. stationis<br>C. simulans<br>C. guaraldiae<br>C. accolens<br>C. striatum<br>C. hesseae<br>C. aurimucosum<br>C. intestinale<br>C. singulare<br>C. faecipullorum | WP_003845902.1<br>WP_066794095.1<br>WP_248090355.1<br>WP_143335552.1<br>WP_284901940.1<br>WP_101504612.1<br>WP_101736373.1<br>WP_010186760.1<br>WP_250223851.1<br>WP_239179046.1<br>HIX78565.1 | NVPVSDPDSNEGQAR | 91-105 | 26973 |
| 3 | Phosphomannomutase/phosphogluco mutase | Clavibacter michiganensis | WP_209657882.1 | DQMAATGAVFGGEHSAHYYFR | 331-351 | 48868 |
| 4 | Phosphoglucomutase (alpha-D-glucose-1,6-bisphosphate-dependent) | C. argentoratense | WP_234858678.1 | MDCSSPDSMASLVANR | 285-300 | 58129 |
| 5 | Phosphoglucomutase (alpha-D-glucose-1,6-bisphosphate-dependent) | C. maris<br>C. xerosis<br>C. freneyi<br>C. lactis | WP_020935557.1<br>WP_046650327.1<br>WP_035123894.1<br>WP_120491165.1 | MDCSSPDAMASLIGNR | 285-300 | 58457 |
| 6 | MULTISPECIES: ATP-dependent 6-phosphofructokinase | C. striatum<br>Corynebacterium | CQD13738.1<br>WP_061922085.1 | GGTPTAYDR<br>ILIVEVMGR<br>LAVLTSGGDCPGLNAVIR<br>TASNEFGSTVVGYEDGWVGLMEDR<br>TTVLGHIQR<br>WLSDNGIPVIGVPK<br>YGIICVAEGALPK<br>YGVHAAR | 289-297<br>176-184<br>15-32<br>37-60<br>280-288<br>124-137<br>228-240<br>303-309 | 38080 |
| 7 | ATP-dependent 6-phosphofructokinase | C. simulans<br>C. minutissimum<br>C. faecipullorum | WP_062041068.1<br>WP_039675088.1<br>HIX78424.1 | EGTMDFEEGGVDQFGHQTFNGIGQVIGDEIK<br>GGTPTAYDR<br>ILIVEVMGR | 229-259<br>277-285<br>164-172 | 36729 |

| | | | | | | |
|---|---|---|---|---|---|---|
| | | | | LAVLTSGGDCPGLNAVIR<br>TTVLGHIQR | 3-20<br>268-276 | |
| 8 | ATP-dependent 6-phosphofructokinase | C. simulans<br>C. minutissimum<br>C. aurimucosum | WP_062041068.1<br>WP_039675088.1<br>GAA1472382.1 | DLYDDAEIDR<br>EGTMDFEEGGVDQFGHQTFNGIGQVIGDEIK<br>ILIVEVMGR<br>LAVLTSGGDCPGLNAVIR | 51-60<br>229-259<br>164-172<br>3-20 | 36729 |
| 9 | Phosphoglycerate kinase | C. simulans<br>C. aurimucosum<br>C. accolens | WP_248090357.1<br>WP_102233912.1<br>PCC81818.1 | AQASVYDVAK<br>EVETLAAVAEKPEHPYVVVLGGAK<br>ITASLPTIK<br>TVFWNGPMGVFEMEAFSK | 163-172<br>185-208<br>41-49<br>320-337 | 42525 |
| 10 | Type I Glyceraldehyde-3-phosphate dehydrogenase | C. striatum<br>C. tuberculostearicum<br>C. simulans | WP_005531614.1<br>WP_251067654.1<br>WP_284841327.1 | AAAVNMVPTSTGAAK<br>HNVISAASCTTNCLAPMAK<br>IAVSAERDPK<br>KVIISAPGK<br>LDGYAMR<br>TTSLVASKL<br>VGINGFGR<br>YDSVLGR | 201-215<br>145-163<br>73-82<br>117-125<br>228-234<br>327-335<br>5-12<br>48-54 | 36061 |
| 11 | MULTISPECIES: Glyceraldehyde-3-phosphate dehydrogenase | Corynebacterium<br>(C. striatum, C. simulans) | WP_167599039.1 | AAGLNMVLTETGAAK<br>EAAYGGVR<br>ELSTAETLPILK<br>LTGNAIR<br>NVSLHSDLR<br>VLLTAPGKGDLK | 336-350<br>149-156<br>66-77<br>363-369<br>399-407<br>254-265 | 53277 |
| 12 | MULTISPECIES: NAD(P)H-dependent glycerol-3-phosphate dehydrogenase | Corynebacterium | WP_147760641.1 | GLGNNTLATIITR | 205-217 | 34440 |
| 13 | Type I Glyceraldehyde-3-phosphate dehydrogenase | C. striatum<br>C. simulans<br>C. accolens<br>C. flavescens | WP_005531614.1<br>WP_284841327.1<br>PCC81819.1<br>WP_075729858.1 | AAAVNMVPTSTGAAK<br>AHIEAGAK<br>AVALVLPELEGK<br>GLMTTVHSYTGDQR<br>IAVSAERDPK<br>TTSLVASKL<br>VLNDKFGIEK<br>YDSVLGR | 218-232<br>126-133<br>233-244<br>191-204<br>90-99<br>344-352<br>181-190<br>65-71 | 37791 |
| 14 | NAD(P)H-dependent glycerol-3-phosphate dehydrogenase | C. terpenotabidum | WP_041631453.1 | VCADAGCDVTLWAR | 18-31 | 34420 |
| 15 | Type I glyceraldehyde-3-phosphate dehydrogenase | C. pyruviciproducens | WP_016457562.1 | LSDDITYDDESITVDGK | 55-71 | 36311 |
| 16 | Glyceraldehyde-3-phosphate dehydrogenase | C. genitalium | WP_005291253.1 | TEAAGQPQNLTEDWNNK | 2-18 | 54176 |
| 17 | MULTISPECIES: Phosphoglycerate kinase | Corynebacterium | WP_061922322.1 | GAFADELAALAADNGAFVSDGFGVVHR | 136-162 | 43078 |
| 18 | MULTISPECIES: Phosphoglyceromutase | Corynebacterium | WP_070563655.1 | DKYGEEQFMAWR<br>HWIPVVR<br>HYGALQGLNK<br>TANIALNAADR<br>TECLKDVVER | 109-120<br>81-87<br>95-104<br>70-80<br>153-162 | 27922 |

| | | | | | | |
|---|---|---|---|---|---|---|
| | | | | YGEEQFMAWR | 111-120 | |
| 19 | MULTISPECIES: Glycerol kinase GlpK | Corynebacterium | WP_075730700.1 | AVLEATAYQTR<br>DVADAMVADSGVEITELR<br>ETTVVWDK<br>ETTVVWDKNTGEPVYNAIVWQDTR<br>FSENGLLTTVCFQR<br>GSLAGVPIR<br>GVIVGLTR<br>NTGEPVYNAIVWQDTR<br>NTYGTGLFLLLNTGTTPK<br>TLLMDIEKLEWDEELCK | 388-398<br>399-416<br>85-92<br>85-108<br>289-302<br>240-248<br>371-378<br>93-108<br>271-288<br>198-214 | 57012 |
| 20 | Phosphoglycerate kinase | C. striatum | WP_049145845.1 | ANGLNDGDVMLIENVR<br>AQASVYDVAK<br>FGDKIVLPVDLIAATEFNADAENK<br>GYNVQNSLLQEDQIDNCK<br>ITASLPTIK<br>IVLPVDLIAATEFNADAENK<br>TLDDLIAEGVESR<br>TVFWNGPMGVFEMEAFSK<br>VVALDGIPEGWMSLDIGPESVK<br>YSLAPVAEALSEALGQYVALAGDVTGEDAHER | 107-122<br>163-172<br>264-287<br>241-258<br>41-49<br>268-287<br>5-17<br>320-337<br>288-309<br>75-106 | 42468 |
| 21 | Glycerol kinase | C. kutscheri<br>C. casei<br>C. stationis<br>C. halotolerans<br>C. simulans<br>C. striatum | VEH05893.1<br>WP_301438465.1<br>WP_066796784.1<br>WP_015402044.1<br>WP_062036799.1<br>WP_110301885.1 | AVLEATAYQTR<br>FVAAIDQGTTSTR | 388-398<br>6-18 | 56837 |
| 22 | Glycerate kinase | C. efficiens | WP_006767981.1 | SNDVAAQLR | 344-352 | 37099 |
| 23 | Diacylglycerol kinase | C. variabile<br>C. nuruki<br>C. kalidii<br>C. glyciniphilum | WP_303948348.1<br>WP_010121418.1<br>WP_244803269.1<br>WP_145943038.1 | EHWFGTIACAGFDSLVSDRTNR | 155-176 | 35641 |
| 24 | Dihydroxyacetone kinase subunit DhaK | C. simulans<br>C. striatum<br>C. accolens | WP_284841216.1<br>WP_239299452.1<br>PCC82883.1 | EIPLVSADDVTDHLMEPILADLK<br>FTEPLFVAR<br>GVAGTLLVEK<br>LAGAAAERGDDLAAVTAVAK<br>RGVAGTLLVEK | 226-248<br>30-38<br>150-159<br>160-179<br>149-159 | 44663 |
| 25 | Dihydroxyacetone kinase subunit DhaK | C. glucuronolyticum | WP_005393004.1 | SLDDVAAIAKK | 181-191 | 32023 |
| 26 | Dihydroxyacetone kinase subunit DhaL | C. striatum<br>C. aurimucosum<br>C. singulare<br>C. simulans | WP_100619471.1<br>WP_193629369.1<br>WP_239178779.1<br>WP_284841233.1 | AAEGEKTMVDAWAPAAR<br>ASYLGER<br>EQADAGAGVADVLR | 111-127<br>167-173<br>131-144 | 20295 |
| 27 | Dihydroxyacetone kinase phosphoryl donor subunit DhaM | C. simulans<br>C. striatum | WP_284841217.1<br>WP_239299454.1 | AIEVFVPK<br>LAEGLAELAGQMAADVR | 125-132<br>17-33 | 23067 |

| | | | | | | |
|---|---|---|---|---|---|---|
| 28 | 2,3-Bisphosphoglycerate-dependent phosphoglycerate mutase | C. striatum<br>C. simulans | GKH16036.1<br>WP_239239238.1 | HWIPVVR<br>HYGALQGLNK<br>RSYGTPPPELEDSSEYSQSNDPR<br>SYGTPPPELEDSSEYSQSNDPR<br>TANIALNAADR<br>TECLKDVVER<br>YADLDQVPR<br>YGEEQFMAWR | 81-87<br>95-104<br>121-143<br>122-143<br>70-80<br>153-162<br>144-152<br>111-120 | 27878 |
