## Supplemental Table S16 for "Peptidome Analysis of Western Blots Identifies Natural Bispecific Antibody-Bound *Corynebacterium* and Phage B-cell Epitopes with Potential Relevance to Psoriasis"

### Table S16. Reverse Citric Acid Cycle Enzymes are *Corynebacterium* Targets.

| Antigen Number | Identified Antigen | *Corynebacterium* (C.), *Actinomycetota* | Accession Number | Peptide Sequence | Peptide Start/Stop | Molecular Weight |
|---|---|---|---|---|---|---|
| 1 | Fumarate reductase/succinate dehydrogenase flavoprotein subunit | C. macginleyi<br>C. halotolerans<br>C. propinquum<br>C. kefirresidentii<br>C. tuberculostearicum<br>C. accolens<br>C. aurimucosum<br>C. ureicelerivorans<br>C. mucifaciens<br>C. striatum<br>C. marquesiae<br>C. casei<br>C. efficiens<br>C. camporealensis<br>C. renale<br>C. intestinale<br>C. stationis<br>C. gallinarum<br>C. guaraldiae<br>C. confusum<br>C. lizhenjunii<br>C. simulans<br>C. minutissimum<br>C. diphtheria<br>C. pseudotuberculosis<br>C. argentoratense | WP_200439599.1<br>WP_015399819.1<br>WP_201810760.1<br>WP_301732893.1<br>WP_005327184.1<br>WP_284628487.1<br>WP_049358776.1<br>WP_038609360.1<br>WP_168684200.1<br>WP_049062448.1<br>WP_337888422.1<br>WP_301465913.1<br>WP_006770371.1<br>WP_035107261.1<br>WP_111725560.1<br>WP_250224029.1<br>WP_075722276.1<br>WP_191732836.1<br>WP_154736750.1<br>WP_290224415.1<br>WP_165011045.1<br>WP_248093517.1<br>WP_052319632.1<br>WP_082257455.1<br>WP_014366412.1<br>WP_020975481.1 | EYGGTLATR<br>GQTGQQLQLSTTSALYR<br>HAEPLYFDSIPLMTR<br>ILESHEPHGVPMK<br>ITGNANDMNQVLEYGLR<br>KVDNDSAYR<br>NLITGELK<br>NSNAGAMMR<br>VIDHMNAIGAPFAR<br>YPAFGNLVPR<br>YSNLIEMYEEAIGESAYETPMR | 173-181<br>196-212<br>663-677<br>37-49<br>585-601<br>123-131<br>248-255<br>281-289<br>159-172<br>361-370<br>414-435 | 75852 |
| 2 | Class II fumarate hydratase | C. aurimucosum<br>C. striatum<br>C. simulans<br>C. guaraldiae<br>M. tuberculosis<br>M. leprae | WP_010187338.1<br>WP_046645452.1<br>WP_239239995.1<br>WP_143335287.1<br>WP_031753822.1<br>WP_010908634.1 | AKEFADVVK<br>CVDGIEANEER<br>DGLVEFSGAMR<br>DLQPGSSIMPGK<br>EAENHFEAQAAR<br>FAESSTSIVTPLNSK<br>GLETAQIR<br>IGYENAAK<br>KFAESSTSIVTPLNSK<br>LLANVSR<br>LNVLSMTNR<br>QIELGIER<br>THLMDATPITLGQEFGGYAR<br>TVAVSLNK<br>VPVDALWR | 168-176<br>413-423<br>298-308<br>339-350<br>286-297<br>427-441<br>71-78<br>442-449<br>426-441<br>401-407<br>485-493<br>232-239<br>212-231<br>309-316<br>48-55 | 53693 |
| 3 | NAD-dependent succinate-semialdehyde dehydrogenase | C. tuberculostearicum<br>C. aurimucosum<br>C. kefirresidentii | WP_200435610.1<br>WP_049359949.1<br>WP_239206630.1 | AEVQIVADIYR<br>FAAPNLLLGNTIVLK<br>HASICPLSSQACQDILEEAGLPK | 81-91<br>140-154<br>155-177 | 51229 |

| | | | | | | |
|---|---|---|---|---|---|---|
| | | C. marquesiae | WP_284788802.1 | LIVLEDFYDR<br>MYNTGQACNAPK<br>WGVGEFVNEHLYR | 268-277<br>255-266<br>441-453 | |
| 4 | Succinate dehydrogenase cytochrome b subunit | C. minutissimum<br>C. simulans<br>C. singulare<br>C. striatum<br>C. lizhenjunii | WP_115021072.1<br>WP_239239278.1<br>WP_042529061.1<br>HCG3010610.1<br>WP_165011043.1 | ANMIATFSR<br>LAASDLGITGAK<br>NPDREALAHGR<br>SVKNPDREALAHGR<br>TNLMGGLDSFATK | 175-183<br>207-218<br>5-15<br>2-15<br>124-136 | 27304 |
| 5 | NAD-dependent succinate-semialdehyde dehydrogenase | Tsukamurella paurometabola | WP_245537894.1 | DAARVQRVGAGLRSGMVGVNR | 422-442 | 49706 |
| 6 | Succinate dehydrogenase iron-sulfur subunit | Clavibacter michiganensis<br>Mycobacterium tuberculosis | WP_104294766.1<br>MBZ4296222.1 | SCAHGVCGSDAMRINGRNR | 83-101<br>54-72 | 26417 |
| 7 | Succinate dehydrogenase cytochrome b subunit | C. glutamicum<br>C. minutissimum<br>C. tuberculostearicum<br>C. kefirresidentii | WP_172768109.1<br>WP_115021072.1<br>WP_316986120.1<br>WP_086589111.1 | YGAFLR | 71-76 | 73471 |
| 8 | MULTISPECIES: HpcH/HpaI aldolase/citrate lyase family protein | Corynebacterium | WP_049152030.1 | AADKPVGVNAFNLEQAQR<br>ATDAYGPTR | 209-226<br>64-72 | 27366 |
