## Supplemental Table S17 for "Peptidome Analysis of Western Blots Identifies Natural Bispecific Antibody-Bound *Corynebacterium* and Phage B-cell Epitopes with Potential Relevance to Psoriasis"

### Table S17. Biosynthetic and Metabolic Pathway Enzymes are *Corynebacterium* Antigens.

| Antigen Number | Identified Antigen | *Corynebacterium* (C.), *Actinomycetota* | Accession Number | Peptide Sequence | Peptide Start/Stop | Molecular Weight |
|---|---|---|---|---|---|---|
| 1 | Thiamine-phosphate synthase | C. striatum<br>C. simulans<br>C. pseudodiphtheriticum<br>C. afermentas<br>C. auris<br>C. ureicelerivorans<br>C.dentalis<br>C. genitalium<br>C. sanguinis<br>C. lipophiloflavum<br>C. urealyticum<br>C. singulare<br>C. phoceense<br>C. aurimucosum<br>C. maris<br>C. intestinale<br>C. guaraldiae<br>C. faecium | WP_257212250.1<br>WP_338079183.1<br>RUQ00782.1<br>WP_286362417.1<br>WP_290342767.1<br>WP_273407388.1<br>WP_099298446.1<br>WP_005291530.1<br>WP_259896483.1<br>WP_006841182.1<br>PZO97559.1<br>WP_144790276.1<br>WP_257045931.1<br>WP_158396145.1<br>WP_020935052.1<br>WP_250224127.1<br>WP_154736443.1<br>WP_302470858.1 | ATPTKDSGR | 136-144 | 64977 |
| 2 | Short-chain fatty acyl-CoA regulator family protein | Prescottella equi | WP_084843967.1 | ICERTNCAQR | 440-449 | 52258,4 |
| 3 | Ribose-phosphate diphosphokinase | C. accolens<br>C. endometrii<br>C. confusum<br>C. aurimucosum<br>C. tuberculostearicum<br>C. curieae<br>C. casei<br>C. striatum<br>C. phocae<br>C. macginleyi<br>C. camporealensis<br>C. ammoniagenes<br>C. phoceense<br>C. kefirresidentii<br>C. simulans | WP_284631182.1<br>WP_136140879.1<br>WP_290225532.1<br>WP_049361516.1<br>WP_250411071.1<br>WP_269946105.1<br>WP_025387851.1<br>WP_086891709.1<br>WP_075734956.1<br>WP_200450876.1<br>WP_035105818.1<br>WP_003846333.1<br>WP_257053725.1<br>WP_259824438.1<br>WP_248090973.1 | DCILLDDMIDTGGTIAGAVR<br>DFANGEIFIR<br>FEESVR<br>ITAILPFYPYAR<br>SVVIACTHGVFSDPAR<br>TDLVPTTAR | 225-244<br>42-51<br>52-57<br>93-104<br>253-268<br>33-41 | 35304 |
| 4 | Putative vanillate O-demethylase oxygenase subunit B | C. efficiens | WP_006769686.1 | RITLRPEHPHK | 28-38 | 41874 |
| 5 | Precorrin-3B C(17)-methyltransferase | C. pseudotuberculosis<br>C. diphtheria<br>C. ulcerans<br>C. silvaticum<br>C. rouxii | WP_014367025.1<br>WP_088247766.1<br>WP_197906693.1<br>WP_087453956.1<br>WP_155872765.1 | VEAERAAMALDMAK | 311-324 | 53900 |

| | | | | | | |
|---|---|---|---|---|---|---|
| | | C. belfantii<br>C. vitaeruminis | WP_088267031.1<br>WP_276652215.1 | | | |
| 6 | Thioredoxin-disulphide reductase | C. aurimucosum<br>C. hesseae<br>C. guaraldiae<br>C. camporealensis<br>C. endometrii | WP_193628707.1<br>WP_269948678.1<br>WP_144014691.1<br>WP_035106127.1<br>WP_136142060.1 | GIMGPELMEEMRAQAIR<br>GIMGPELMEEMR<br>LHVGDEVFEAR<br>TVILATGAAPR | 56-72<br>56-67<br>94-104<br>105-115 | 33109 |
| 7 | Putative dihydrodipicolinate reductase domain protein | Corynebacterium matruchotii | WP_311166904.1 | WTAISPAANFLPDAPAR | 140-156 | 21938 |
| 8 | Nicotinate-nucleotide adenylyltransferase | Prescottella equi | WP_221282419.1 | HPAADPSQSSVQAEPGEH | 229-246 | 25842 |
| 9 | Proline dehydrogenase family protein | C. matruchotii | WP_314822490.1 | SLKVGKGTEIDTTMNGIIEAPGEK | 783-806 | 125436 |
| 10 | GDSL-type esterase/lipase family protein | C. accolens | WP_302518661.1 | GVPGATDNYPQREGCLQAPNNWPR | 53-76 | 29931 |
| 11 | 2-Oxoglutarate dehydrogenase, E1 subunit | C. diphtheriae | WP_199326401.1 | DLFEKQGAPSTPRTEAKNTSSQPAAPAK | 34-61 | 136659 |
| 12 | Glutamine--fructose-6-phosphate aminotransferase [isomerizing] | C. diphtheriae | WP_106361598.1 | VIAICNTNGSSIPRESDACLYTHAGPEIAVASTKAFLAQITATYLLGLYLAQLRGNMFADE<br>VNAVLGELR | 381-450 | 67635 |
| 13 | Lysophospholipid acyltransferase family protein | C. ulcerans | WP_197911364.1 | IAFTTNDPVIPVAMIGSREANPIGSWIPRPYKVRMK | 141-176 | 26999 |
| 14 | Aldose epimerase | C. pseudodiphtheriticum | WP_284596435.1 | NADGHGAILWADEHFSWCQVYTSPESAPSIGRAVAVEPMTCPPNALRSGESLLQLASASS<br>MR | 254-315 | 34917 |
| 15 | Glutamine--fructose-6-phosphate transaminase (isomerizing) | C. diphtheriae | WP_106361598.1 | VIAICNTNGSSIPRESDACLYTHAGPEIAVASTKAFLAQITATYLLGLYLAQLRGNMFADE<br>VNAVLGELR | 381-450 | 67635 |
| 16 | Glutamate-5-semialdehyde dehydrogenase | C. diphtheriae | WP_196979889.1 | CSVCNATETVLIDSALDSAYQLAIITALQEAGVTVHGDVAQLEAVGASGIVPAEEHDWAE<br>EYLSLDIACALVDGVDAAMEHIRTYSTK | 265-352 | 45133 |
| 17 | Aminotransferase class I/II-fold Pyridoxal phosphate-dependent enzyme | C. simulans<br>C. striatum | WP_062042630.1<br>CQD15012.1 | NYGNLEGIKELR<br>DAGIVLTK<br>ELVKDPQVK<br>ENLDWYLSHAK<br>FWAMSSTSK<br>NYGNLEGIK<br>VWELAK<br>WICPVPGYDR | 69-80 | 46497 |
| 18 | Type I glutamate--ammonia ligase | C. simulans<br>C. striatum | WP_062043873.1W<br>P_188311178.1 | ALEEDNEFLTEGDVFTEDLIETYLQYK<br>DGKPLFYDEAGYGGLSDIAR<br>FFVNDPFTLEPFSR | 428-454<br>283-302<br>93-106 | 53380 |

| | | | | | | |
|---|---|---|---|---|---|---|
| | | | | FNSLLHAADDIQTFK<br>GGYFPTAPVDK<br>IPITGSNPK<br>LVPGFEAPINLVYSQR | 231-245<br>184-194<br>352-360<br>329-344 | |
| 19 | Polyphosphate kinase 2 | C. striatum<br>C. glutamicum<br>C. lipophiloflavum<br>C. sanguinis<br>C. ulcerans<br>C. pseudotuberculosis<br>C. vitaeruminis<br>C. cystitides<br>C. accolens<br>C. tuberculostearicum<br>C. incognita<br>C. dentalis<br>C. auris | WP_306592800.1<br>WP_077313360.1<br>WP_040422825.1<br>WP_144773718.1<br>WP_013912375.1<br>AEK93127.1<br>WP_048760381.1<br>WP_092259910.1<br>PCC82371.1<br>WP_316986494.1<br>WP_185176554.1<br>WP_099298129.1<br>WP_290341878.1 | ADIKDDELPEIDLAK<br>ALQIELLK<br>ENYPYEER<br>ESTSWYFQR<br>EVPMLENMILGSGISLTK<br>FNEHLNPR<br>HIILFEGR<br>QIDPVR<br>TVALEKPSPR<br>VMGFCTESQHAEFLR<br>YTDTEESPWITIK<br>VMGFCSQQEYVR | 2-16<br>64-71<br>45-52<br>123-131<br>173-190<br>102-109<br>83-90<br>208-213<br>113-122<br>158-172<br>241-253<br>120-131 | 35172 |
| 20 | Homoserine dehydrogenase | C. aurimucosum<br>C. curiae<br>C. kefirresidentii<br>C. tuberculostearicum<br>C. simulans<br>C. striatum | WP_049361402.1<br>WP_269945980.1<br>WP_301716586.1<br>WP_301431973.1<br>WP_062041180.1<br>WP_100086839.1 | HNVSLR<br>IGGPAEIR<br>LLGENADAFAHR | 392-397<br>51-58<br>39-50 | 46987 |
| 21 | Homoserine dehydrogenase | C. argentoratense<br>C. guaraldiae<br>C. aurimucosum<br>C. minutissimum | WP_314928990.1<br>WP_144014733.1<br>WP_201828368.1<br>WP_115021714.1 | LGYAEADPTADVEGHDAASK<br>LLAICER | 202-221<br>264-270 | 46987 |
| 22 | Catalase | C. simulans<br>C. striatum<br>M. tuberculosis | WP_339019011.1<br>WP_284806901.1<br>PLV46239.1 | DADMQWDFWTR<br>DVHGFALR<br>DYPLVDVGYFVLNR<br>EDLYNAIEEGDYPIWDVK<br>FNPFDLTK<br>FPDFIHSQK<br>FSTVAGEQGSPDTWR<br>FYTEDGNYDIVGNNTPVFFLR<br>NADYHREDLYNAIEEGDYPIWDVK<br>NPESTHQVTYLMGDR<br>VFAYADQQR<br>VQIMPLAEAAEYR<br>VRELFAEKN | 172-182<br>122-129<br>301-314<br>257-274<br>288-295<br>155-163<br>107-121<br>130-150<br>251-274<br>183-197<br>349-357<br>275-287<br>507-515 | 58221 |
| 23 | Adenylosuccinate synthetase | C. striatum<br>C. simulans<br>C. accolens | WP_131771446.1<br>WP_061924846.1<br>PCC82397.1 | LAPMVIDAELELNK<br>LDVLTGIGEIPICVAYDVDGK<br>MSYIGVGPGR<br>QKVESALDIK<br>TFEELPQK | 196-209<br>331-351<br>408-417<br>158-167<br>385-392 | 46942 |

| | | | | | | |
|---|---|---|---|---|---|---|
| | | | | TSLGIIK<br>VQDIFDESILR<br>GIGPAYSDK | 260-266<br>147-157<br>131-139 | |
| 24 | Phosphopyruvate hydratase (enolase) | C. simulans<br>C. aurimucosum<br>C. diphtheria<br>C. pseudotuberculosis<br>C. ulcerans<br>C. minutissimum<br>C. accolens<br>M. tuberculosis<br>C. striatum | WP_284841357.1<br>WP_010187392.1<br>WP_106202524.1<br>WP_058832493.1<br>WP_046693914.1<br>WP_115021570.1<br>PCC83167.1<br>WP_052612112.1<br>WP_284806307.1 | AAAESAGLPLYR<br>AANALLVK<br>DGGDRYLGK<br>DGKYHFEGGEHTAEEMAK<br>GLSTGLGDEGGFAPEAESTK<br>GNPTVEAEVFLDDGAR<br>GVAGVPSGASTGVHEAHELR<br>KAANALLVK<br>LTETLGDK<br>SGETEDTTIADLAVALNCGQIK<br>SKGLSTGLGDEGGFAPEAESTK<br>TMMSHR<br>VNQIGTLTETFDAVELAHR<br>VQIVGDDFFVTNPAR<br>YHFEGGEHTAEEMAK<br>AALDLIVEAIK<br>AALDLIVEAIKK<br>ADIIHVFAR<br>AGFEPGKDIALALDVASSEFFK<br>LIDQALIALDGTENK<br>MGAEVYHSLK<br>SGETEDTTIADLAVALNCGQIK<br>TMMSHR | 120-131 327-334<br>53-61<br>250-267<br>196-215<br>17-32<br>33-52<br>327-335<br>296-303<br>364-385<br>195-216<br>358-363<br>335-353<br>304-318<br>253-267<br>216-226<br>216-227<br>2-10<br>228-249<br>250-267<br>180-189<br>364-385<br>358-363 | 45297 |
| 25 | Formate C-acetyltransferase | C. simulans<br>C. striatum<br>C. aurimucosum<br>M. tuberculosis | WP_275437400.1<br>WP_275432387.1<br>WP_193629694.1<br>SGC81874.1 | AGTPFAPGANPENGADSHGMVASMLSVGK<br>ALLYAINGGRDEVTGK<br>ATGAFPSGHK<br>DALDGISLTNTITPSGLGR<br>EQWGDDAAIACCVSPMEVGK<br>IIGDYR<br>LSAFFDCYFER<br>MVEQAIR<br>NAIPTQSVLTITSNVVYGK<br>NYTPYDGDASFLAGPTEK<br>QMQFFGAR<br>QVVEGYDAIQGDGPLDFDEVWK<br>QVVEGYDAIQGDGPLDFDEVWKK<br>SMGCGIAGLSIVADSLSAIK<br>SQDGAAMSIGR<br>SSHIITGLPDAYGR<br>TEDYDQIFSGDPYWATWSDAGFGNDGR<br>THNDAVFDIYTPR<br>TWDTLEKDYLSVER<br>VALYGVDYLIAEK<br>YREEHAEQIK | 623-651<br>438-453<br>613-622<br>656-674<br>405-424<br>179-184<br>277-287<br>119-125<br>594-612<br>34-51<br>425-432<br>454-475<br>454-476<br>519-538<br>266-276<br>163-176<br>318-344<br>145-157<br>55-68<br>186-198<br>217-226 | 77524 |
| 26 | Glutamine--fructose-6- | C. striatum | WP_306496452.1 | EIHDQPAAVR | 247-256 | 65268 |

| | | | | | | |
|---|---|---|---|---|---|---|
| | phosphate aminotransferase [isomerizing] | C, simulans | WP_248092683.1 | GGFNSFMEK<br>GNMFADEVNAVLQELR<br>GYDVDQPR<br>IPTEVELAHEFR<br>VIAICNTQGSSIPR<br>VVSNIQEIR<br>YRDPIVNEK | 238-246<br>415-430<br>588-595<br>312-323<br>361-374<br>525-533<br>324-332 | |
| 27 | Glyceraldehyde-3-phosphate dehydrogenase | C. maris | WP_052337728.1 | DREALGQHLQSK<br>EALGQHLQSK<br>LTGNAIR | 237-248<br>239-248<br>364-370 | 53618 |
| 28 | ATP synthase subunit beta | C. aurimucosum<br>C. minutissimum<br>C. jeikeium<br>C. striatum<br>C. accolens<br>C. simulans | WP_193634369.1<br>WP_115021734.1<br>WP_034985588.1<br>WP_284772821.1<br>WP_302519069.1<br>WP_062035306.1 | DGEQWGIHREPPAFDQLEGK<br>DVQNQDVLLFIDNIFR<br>EFSGTSVFAGVGER<br>FLGQNFFVAEK<br>FTQAGSEVSTLLGR<br>GAAVTDTGKPISVPVGDVVK<br>GHVFNALGDCLDQPGLGR<br>GIYPAVNPLTSTSR<br>IGLFGGAGVGK<br>KLTEKK<br>MPSAVGYQPTLADEMGVLQER<br>TEILETGIK<br>TVLIQEMITR<br>TVTLEVAQHLGDNLVR<br>VALSGLTMAEYFR<br>VIDLLTPYVK<br>VIGPVVDVEFPR | 121-140<br>250-265<br>187-200<br>418-428<br>266-279<br>83-102<br>103-120<br>348-361<br>163-173<br>477-482<br>280-300<br>141-149<br>174-183<br>55-70<br>237-249<br>150-159<br>22-33 | 52192 |
| 29 | Phosphoribosylformyl glycinamidine synthase | C. striatum<br>C. simulans<br>C. accolens | WP_240504751.1<br>WP_248093135.1<br>PCC82541.1 | FDSTVGAGTVLMPFGGTR<br>ITAGESEWER<br>MAAALDR<br>NYFASEDREPTITELK<br>SQMLFLPGGFSGGDEPDGSAK<br>TVTELLR<br>VIDTYWSDHCR | 636-653<br>1138-1147<br>1009-1015<br>185-200<br>1016-1036<br>828-834<br>201-211 | 129845 |
| 30 | Aldehyde dehydrogenase (NAD) family protein | C. striatum | WP_086891871.1 | AAPADAQGENTYSCLSSR<br>AVLELGGSDPYILLDTDDVK<br>AVLELGGSDPYILLDTDDVKEAAK<br>DAEICPR<br>ELGPLGMDEFVNK<br>FVGPNLVLGNTIVLK<br>IQGVSLTGSER<br>KAVLELGGSDPYILLDTDDVK<br>MYNTGQACNSNK<br>MYNTGQACNSNKR<br>RLEVGMANVNTPAGEGEELPFGGVK<br>YYADNGEEFTK | 294-311<br>228-247<br>228-251<br>158-164<br>439-451<br>143-157<br>202-212<br>227-247<br>258-269<br>258-270<br>408-432<br>95-105 | 49262 |
| 31 | L-serine ammonia-lyase | C. striatum<br>C.diphtheriae | HAT6434148.1<br>WP_014316377.1 | IGIGPSSSHTLGPMK<br>LMLGGTR | 11-25<br>97-103 | 49579 |

| | | | | | | |
|---|---|---|---|---|---|---|
| | | C. pseudotuberculosis<br>C. minutissimum<br>C. ulcerans<br>C. simulans | WP_104992939.1<br>WP_039675753.1<br>WP_014525335.1<br>WP_248091825.1 | NAVAAVDSITSAR | 395-407 | |
| 32 | Carbamoyl-phosphate synthase large chain | C. simulans<br>C. striatum<br>C. accolens | WP_284841318.1<br>WP_306594062.1<br>PCC81839.1 | EVGVDTGGCNIQFAVNPVDGR<br>FAFEKFPGADDTLTTTMK<br>IAGGGLAASPEANVLIEESILGWK<br>MYDVELAFR<br>VIILGSGPNR | 279-299<br>366-383<br>196-219<br>444-452<br>576-585 | 119636 |
| 33 | 3'(2'),5'-Bisphosphate nucleotidase CysQ | C. simulans<br>C. striatum | WP_239240696.1<br>WP_205690651.1 | MGFETVGIGSAGAK<br>VWIVDPLDGTK | 169-182<br>90-100 | 28305 |
| 34 | FadD32-like long-chain-fatty-acid--AMP ligase | C. striatum<br>C. simulans<br>C. accolens | WP_005529491.1<br>WP_062036842.1<br>PCC82444.1 | AVFGDSNPSIVLTNNVSAAAVR<br>DFVQQPK<br>LKDLIVVAGR<br>TPAGVLLTNR | 130-151<br>277-283<br>507-516<br>215-224 | 67100 |
| 35 | 2-Oxoglutarate dehydrogenase, E2 component, dihydrolipoamide succinyltransferase | C. striatum<br>C. minutissimum<br>C. simulans | WP_110292140.1<br>WP_039676153.1<br>WP_062038148.1 | ATVEALVSHPNVNASYNPETK<br>GLLTPVIHK<br>HGVDLSTVSGTGVGGR<br>QMCYLPFTYDHQVVDGADAGR<br>SVDPEKQELIGTTQK | 395-415<br>434-442<br>270-285<br>526-546<br>320-334 | 60233 |
| 36 | IMP dehydrogenase | C. striatum<br>C. tuberculostearicum<br>C. accolens<br>C. simulans | WP_272707438.1<br>WP_301979514.1<br>WP_302532021.1<br>WP_062035817.1 | FVQITAAGLK<br>GMGSMGAMQGR<br>ISGLPVVDKDGTLLGICTNR<br>LGVPLASAAMDTVTEAR<br>MAIAMAR<br>QGGIGVLHR<br>RFVQITAAGLK<br>SEDKLVPEGVEGR<br>VGIGPGSICTTR<br>VLEMVSR | 480-489<br>399-409<br>130-149<br>52-68<br>69-75<br>76-84<br>479-489<br>430-442<br>309-320<br>268-274 | 53460 |
| 37 | Ribose-phosphate diphosphokinase | C. striatum<br>C. simulans | WP_086891709.1<br>WP_248090973.1 | AHPELAEAVAK<br>DFANGEIFIR<br>FEESVR<br>GADCFVMQSHSQPLNK<br>ITAILPFYPYAR | 19-29<br>42-51<br>52-57<br>58-73<br>93-104 | 35377 |
| 38 | Formate--tetrahydrofolate ligase | C. striatum<br>C. camporealensis<br>C. tuberculostearicum<br>C. singulare<br>C. accolens<br>C. simulans<br>C. kefirresidentii<br>C. marquesiae | WP_188311025.1<br>WP_321114360.1<br>WP_301988631.1<br>WP_239179142.1<br>WP_302525859.1<br>WP_248090238.1<br>WP_086587676.1<br>WP_284810701.1 | AIVALREPSQGPVMGIK<br>CLDVNDR<br>EDLTTENLDALR<br>EDLTTENLDALRK<br>GIVNLER<br>KGIVNLER<br>LVLVTGVSPTPAGEGK | 90-106<br>167-173<br>344-355<br>344-356<br>357-363<br>356-363<br>57-72 | 59167 |
| 39 | Multifunctional oxoglutarate decarboxylase/oxoglutarate dehydrogenase thiamine | C. simulans<br>C. striatum | WP_201057398.1<br>WP_306588608.1 | IGEAPEAGER<br>RAQSSTATGIAK<br>VIQGAESGEFLR<br>VNAAEAFENFLHTK | 111-120<br>1217-1228<br>333-344 | 137168 |

| | | | | | | |
|---|---|---|---|---|---|---|
| | pyrophosphate-binding subunit/dihydrolipoyllysine-residue succinyltransferase subunit] | | | | | |
| 40 | MULTISPECIES: S-(hydroxymethyl)mycothiol dehydrogenase | unclassified Corynebacterium | WP_070521258.1 | DGDIEDAFPFLLGHEAAGVVER<br>NGQFPLDKFVSER<br>TLVHEGQCTK<br>VQACGVCHTDLAYR<br>YCFNTHNASAK | 51-72<br>333-345<br>144-153<br>37-50<br>107-117 | 39156 |
| 41 | 2,3-Bisphosphoglycerate-dependent phosphoglycerate mutase | C. striatum<br>C. simulans | GKH16036.1<br>WP_239239238.1 | HWIPVVR<br>HYGALQGLNK<br>RSYGTPPPELEDSSEYSQSNDPR<br>SYGTPPPELEDSSEYSQSNDPR<br>TANIALNAADR<br>TECLKDVVER<br>YADLDQVPR<br>YGEEQFMAWR | 81-87<br>95-104<br>121-143<br>122-143<br>70-80<br>153-162<br>144-152<br>111-120 | 27878 |
| 42 | MULTISPECIES: ATP-dependent 6-phosphofructokinase | C. striatum<br>Corynebacterium | CQD13738.1<br>WP_061922085.1 | GGTPTAYDR<br>ILIVEVMGR<br>LAVLTSGGDCPGLNAVIR<br>TASNEFGSTVVGYEDGWVGLMEDR<br>TTVLGHIQR<br>WLSDNGIPVIGVPK<br>YGIICVAEGALPK<br>YGVHAAR | 289-297<br>176-184<br>15-32<br>37-60<br>280-288<br>124-137<br>228-240<br>303-309 | 38080 |
| 43 | Pyruvate kinase | C. simulans<br>C. striatum | AMO92590.1<br>WP_086892379.1 | AEASDVANAVLDGADAVMLSGETSVGIDPQNVVR<br>AIVAFTTSGDTAK<br>AVGILADLQGPK<br>EVDGNDVVCEVTEGGPVSNNK<br>GDLGVEVPLEQVPLFQK<br>GVSLPGMDISVPALSEKDIADLR<br>ITVDDVEGTHDR<br>IVCTLGPAVASK<br>LLIDDGK<br>RGVVSYSAR<br>SPADVDLVHAIMDEEGR | 292-325<br>369-381<br>62-73<br>136-156<br>244-260<br>157-179<br>94-105<br>8-19<br>123-129<br>351-359<br>196-212 | 50592 |
| 44 | Adenylosuccinate lyase | C. striatum<br>C. phoceense<br>C. phocae<br>M. tuberculosis<br>C. bovis<br>C. kroppenstedtii<br>C. lizhenjenii<br>C. glaucum<br>C. pseudogenitalium<br>C. pseudodiphthériticum<br>C. urealyticum | WP_049062730.1<br>WP_306430196.1<br>WP_075732885.1<br>WP_003403952.1<br>WP_342015339.1<br>WP_303733782.1<br>WP_413227991.1<br>WP_095660653.1<br>WP_256008640.1<br>WP_284586376.1<br>WP_012360865.1 | DLTENVEQLQIHSSLR<br>EGQVGSSAMPHK<br>ETAHEVIK<br>ILMAAVR<br>INNLVSEHPK<br>LMAGNETVTEGFK<br>NISNVLSSR<br>SHNVAAQATTLGK<br>SLDFDAVSALVELGAGPSSLATTIR<br>VCGFQVILR<br>YLPFLATTR | 109-124<br>279-290<br>392-399<br>380-386<br>460-469<br>266-278<br>9-17<br>153-165<br>241-265<br>299-307<br>371-379 | 52426 |

| | | | | | | |
|---|---|---|---|---|---|---|
| | | C. appendicis<br>C. urinipleomorphum<br>C. intestinavium<br>C. stationis<br>C. faecipullorum<br>C. pollutisoli<br>C. falsenii<br>C. faecigallinarum<br>C. durum | WP_273415838.1<br>WP_087117521.1<br>HJD48853.1<br>WP_347284881.1<br>HIX78635.1<br>HJD77659.1<br>WP_276785164.1<br>HJC85745.1<br>WP_315160218.1 | | | |
| 45 | Pyruvate carboxylase | C. striatum<br>C. accolens<br>C. simulans<br>C. faecipullorum<br>C. aurimucosum<br>C. minutissimum<br>C. hiratae<br>C. guaraldiae<br>C. hesseae<br>C. intestinale | WP_284806246.1<br>PCC83475.1<br>WP_062042037.1<br>HIX79884.1<br>WP_102234259.1<br>WP_039673967.1<br>WP_328287848.1<br>WP_348522052.1<br>WP_269948377.1<br>WP_256486006.1 | APPAVDESGR<br>GSRDELLELGPK<br>GTDLVTAAEAVAR<br>IGTEGQPVK<br>QLYRPFEK<br>SGVDVFR<br>VQVEHTVTEEVTGVDIVK<br>ALLREPDFVNTR<br>APPAVDESGR<br>SGVDVFR<br>VDTSFIPEHPDLLKAPPAVDESGR<br>VQVEHTVTEEVTGVDIVK<br>VVEIAPAPTLDPELR | 460-469<br>508-519<br>552-564<br>58-66<br>811-818<br>637-643<br>300-317<br>434-445<br>460-469<br>637-643<br>446-469<br>300-317<br>245-259 | |
| 46 | FAD-binding oxidoreductase | C. diphtheriae<br>C. ulcerans<br>C. pseudotuberculosis<br>C. striatum<br>C. deserti<br>C. belfantii<br>C. rouxii | WP_003850144.1<br>WP_029975597.1<br>WP_104992924.1<br>MDU3173950.1<br>WP_053543885.1<br>WP_088268052.1<br>WP_155871212.1 | VMEFGGR<br>GSLATLAQLEELAPK<br>IHSIDPETAIVDVDGGVTLDQLMK<br>NAIDPTGVFASDMSR<br>QVTIGGAIGPDIHGK | 434-440<br>272-286<br>100-123<br>469-483<br>140-154 | 53211 |
| 47 | Bifunctional UDP-N-acetylglucosamine diphosphorylase/glucosamine-1-phosphate N-acetyltransferase GlmU | C. striatum<br>C. kefirresidentii<br>C. marquesiae<br>C. tuberculostearicum<br>C. curieae<br>C. aurimucosum<br>C. pseudogenitalium<br>C. simulans<br>C. camporealensis | WP_110333425.1<br>WP_324291861.1<br>WP_284792203.1<br>WP_296181254.1<br>WP_316987100.1<br>WP_080971258.1<br>WP_005323654.1<br>WP_248090969.1<br>WP_035105816.1 | LDSNNAQGELYITDVLEIAR<br>RPGTPAAEAAAR | 445-456<br>179-198 | 50086 |
| 48 | Type I methionyl aminopeptidase | C. striatum<br>C. simulans | WP_086891163.1<br>WP_096337185.1 | AVAPGVTTDEIDR<br>EVNVIGR<br>FGYNVVR<br>IAANALQEAGK<br>LTPQKPTPIR<br>THEAMMR<br>TVPDSIERPEYVWK<br>VIESYANR | 68-80<br>180-186<br>195-201<br>57-67<br>7-16<br>164-170<br>17-30<br>187-194 | 31887 |
| 49 | NAD(P)H-quinone oxidoreductase | C. striatum<br>C. phocae | WP_100086813.1<br>WP_075732361.1 | LTLQGTTLR<br>NQCDVILDIIGAK | 250-258<br>200-212 | 33366 |

| | | | | | | |
|---|---|---|---|---|---|---|
| | | C. simulans | WP_062036272.1 | | | |
| 50 | Dihydrolipoyl dehydrogenase | C. simulans<br>C. striatum<br>C. guaraldiae<br>C. aurimucosum<br>C. intestinale<br>C. hesseae<br>C. hiratae | WP_096336201.1<br>WP_284765155.1<br>TRX32002.1<br>WP_010189871.1<br>MCL8493303.1<br>WP_101735756.1<br>TRX64535.1 | ATFCNPQVASFGYTEEQAR<br>DGSKTDTLTVDR<br>EAAEGIGGHMINL<br>FDLTAEEIGR<br>GVHYLMK<br>KVAVIEK<br>KVSEGIVK<br>NAEVAHTFNHEAK<br>TIEITEGDDKGK<br>TNVPGIYAIGDVTAK<br>VEGFGLENTGVELTER<br>VLPNEDKDVSK | 352-370<br>255-266<br>458-470<br>435-444<br>98-104<br>30-36<br>90-97<br>57-69<br>122-133<br>303-317<br>277-292<br>208-218 | 50291 |
| 51 | Aspartate ammonia-lyase | C. accolens<br>C. macginleyi<br>C. simulans | WP_237791907.1<br>WP_200448931.1<br>WP_339019349.1 | AEAIIWACDQILDEGR<br>AGLGEINLPAR<br>QAGSSIMPGK<br>TQLQDAVPMTLGDEFK<br>VLSVENLMHPEFR | 136-151<br>370-380<br>381-390<br>253-268 | 56665 |
| 52 | Acetyl-CoA hydrolase/transferase family protein | C. simulans<br>C. striatum | WP_284841984.1<br>WP_005531380.1 | AAHAAGEEFK<br>IAANFIEFLEGEVAAGR<br>IGTPYIELPTEK<br>ILEDGNIIPSSAVGNNLEYIEAADK<br>IMNGTGGSGDFTR<br>NAPFKAPDEVSEK<br>VVAVVETNAPDR | 54-63<br>240-256<br>203-214<br>139-163<br>380-392<br>227-239<br>215-226 | 54723 |
| 53 | Dihydroorotase | C. striatum<br>C. simulans<br>C. aurimucosum<br>C. hiratae | WP_284772562.1<br>WP_339019330.1<br>WP_049156574.1<br>WP_158395427.1 | CVQDPQLMR<br>GLDVLLAQHAEDHR<br>MFSDDGK<br>MHICHASTEGTVELLK<br>QALLDGLIDVVATDHAPHGSEDK<br>TLTEFGMMAR<br>VMSERPAEITK<br>VNPPLREER<br>VTHTILR | 170-178<br>186-199<br>163-169<br>243-258<br>294-316<br>147-156<br>368-378<br>292-300<br>428-434 | 48517 |
| 54 | Alpha/beta hydrolase family protein | C. striatum | WP_205690679.1 | CDVWSPAMGR<br>HPDQFR<br>NGGNAGLYLLDGLR<br>NIPVQIQPAK<br>SKPQVLNVMNAW<br>TNAWVNDVNAAR | 62-71<br>192-197<br>82-95<br>72-81<br>328-339<br>98-109 | 37059 |
| 55 | Transketolase | C. striatum<br>C. simulans | WP_168926468.1<br>WP_284841340.1 | EETERPSFIR<br>ESVLPSAVR<br>LGNLIVFWDDNR<br>NLHFGIR<br>RYPADWTDVDTR<br>TVIGYPAPNK<br>VINHDPNDVK<br>VLAADAVQK | 246-255<br>620-628<br>188-199<br>424-430<br>5-16<br>258-267<br>54-63<br>23-31 | 74425 |

| | | | | | | |
|---|---|---|---|---|---|---|
| | | | | VVSAPCLEWFDEQDDAYR<br>VVSAPCLEWFDEQDDAYRESVLPSAVR<br>YPADWTDVDTR | 602-619<br>602-628<br>6-16 | |
| 56 | NADP-specific glutamate dehydrogenase | C. striatum<br>C. riegelii<br>C. appendicis<br>C. ureicelerivorans<br>C. mucifaciens<br>C. camporealensis<br>C. curieae<br>C. otitides<br>C. phoceense<br>C. simulans | WP_018297022.1<br>WP_276907305.1<br>WP_273412178.1<br>WP_273406399.1<br>WP_168684417.1<br>WP_321113165.1<br>WP_316986904.1<br>WP_004600094.1<br>WP_257051932.1<br>WP_248091539.1 | AANAGGVATSALEMQQNASR<br>FLGFEQIFK<br>GLSWGGSLVR<br>HESGVLTGK<br>LCEPER<br>NSLTGLPIGGGK<br>THGESFDGAK<br>VQFNSALGPYK | 371-390<br>108-116<br>199-208<br>190-198<br>54-59<br>117-128<br>226-235<br>82-92 | 48735 |
| 57 | Glutamine-hydrolyzing GMP synthase | C. simulans<br>C. striatum | WP_062035815.1<br>WP_172453545.1 | AIGAEFIR<br>DFVASTGAK<br>EANIYSEVVPNSATVEEIK<br>ELGLPEEIVAR<br>EQVEKDFVASTGAK<br>FLTEVAGLEQNWTADNIAEQLIADIQK<br>IIGEVTEER<br>IIGEVTEERLETLR<br>KAIGAEFIR<br>LAGVTEPEAK<br>LLFKDEVR<br>LPYDVLEK<br>LTCVFVDHGLLR<br>LVTADER<br>QPFPGPGLGIR<br>SHHNVGGLPDDVEFELVEPLR<br>TDVNVSGGVLHQGLEATHK<br>VVLDCTSKPPGTIEWE | 290-297<br>256-264<br>17-35<br>375-385<br>251-264<br>176-202<br>397-405<br>397-410<br>289-297<br>278-287<br>363-370<br>474-481<br>235-246<br>265-271<br>386-396<br>342-362<br>99-117<br>496-511 | 54502 |
| 58 | Histidinol-phosphate transaminase | C. striatum | WP_100086816.1 | AMEAAAADANRYPDMGAIELR<br>AYGLAGVR<br>LSSNEATQPPLPEALK | 40-60<br>214-221<br>24-39 | 37538 |
| 59 | Aminotransferase class I/II-fold pyridoxal phosphate-dependent enzyme | C. striatum<br>C. simulans | WP_100087349.1<br>WP_339018754.1 | DGTGTLGVLAAEAAYR<br>DWLVEELPK<br>ELLEEALDRLEK<br>LNFATSR | 244-259<br>277-285<br>351-362<br>344-350 | 40478 |
| 60 | Phosphoribosylaminoimidazolesuccinocarboxamide synthase | C. striatum<br>C. simulans | WP_279109558.1<br>WP_248093129.1 | ISAYDHSLDPAIPDKGR<br>IYSEAAR<br>KLDMLPFECVAR | 35-51<br>179-185<br>92-103 | 32228 |
| 61 | Glycogen debranching protein GlgX | C. gallinarum<br>C. faecale<br>C. efficiens<br>C. deserti<br>C. halotolerans<br>C. callunae<br>C. glutamicum | WP_191732149.1<br>WP_290276142.1<br>WP_006768008.1<br>WP_231686408.1<br>WP_015401268.1<br>WP_029703272.1<br>WP_087061985.1 | FDLAATLAR<br>GEPATLGEFASR<br>LLVDPYAR<br>LMTQDDWDHDFGR<br>YRDTVR | 366-374<br>438-449<br>111-118<br>629-641<br>428-433 | 81818 |

| | | | | | | |
|---|---|---|---|---|---|---|
| | | C. occultum<br>C. humireducens<br>C. marinum<br>C. durum<br>C. pseudokroppenstedtii<br>C. kroppenstedtii<br>C. parakroppenstedtii<br>C. casei<br>C. simulans<br>C. striatum<br>Mycobacterium tuberculosis | WP_156231212.1<br>WP_040086182.1<br>WP_042621725.1<br>WP_006063835.1<br>WP_236881650.1<br>WP_303974533.1<br>WP_260625051.1<br>WP_301460111.1<br>WP_248091450.1<br>WP_172453511.1<br>WP_031684559.1 | | | |
| 62 | MULTISPECIES: Nucleoside-diphosphate kinase | Corynebacterium | WP_070519481.1 | EISIWFPNL<br>LVAMDLR<br>QLAGGTDPVSK<br>TLILIKPDGVK | 131-139<br>36-42<br>90-100<br>8-18 | 15104 |
| 63 | Choline dehydrogenase | C. striatum | WP_110333688.1 | AIGVEYEWK<br>EFSPGPDVQTDEEILEWVR<br>EWVEAVR<br>GNPMDYEK<br>KPLAPLNDVTWYK<br>NDGETALHPSCTTK<br>QEGFAPFDR<br>SLLDTEAMAEVGAR | 247-255<br>460-478<br>436-442<br>114-121<br>551-563<br>479-492<br>197-205<br>446-459 | 65900 |
| 64 | Mycothione reductase | C. simulans<br>C. striatum | WP_061923167.1<br>WP_166684168.1 | HATFVGPK<br>IDKIAEGGEAYR<br>RGDETPNITVFDK<br>VFVNRIDK | 124-131<br>99-110<br>111-123<br>94-101 | 50602 |
| 65 | Glycine cleavage system aminomethyltransferase GcvT | C. striatum<br>C. simulans | WP_005528637.1<br>WP_239240775.1 | AEGFDVDLNNESR<br>FLVVPNAGNADVVWEALNER<br>KYAFICR<br>NSAGLFDLSHMGEIWVNGEDAAK<br>YAFICR | 130-142<br>110-129<br>186-192<br>44-66<br>187-192 | 40501 |
| 66 | Pyruvate dehydrogenase (acetyl-transferring), homodimeric type | C. striatum<br>C. simulans<br>C. accolens | WP_086890992.1<br>WP_062043811.1<br>WP_284808390.1 | ANDGAYVR<br>DGVASYLHDSDPEETR<br>GAGWNVIK<br>GFIIGATAGR<br>GNTQIIQELESFFR<br>GVEPTEAFATK<br>GYGLGHNFEGR<br>IVPIIPDEAR<br>KLTLEDLK<br>LSEEDLDGFRQEVSR<br>LTLEDLK<br>QNVATTMALVR<br>TGDSFWAAGDQLGR | 348-355<br>18-33<br>305-312<br>648-657<br>291-304<br>807-817<br>416-426<br>542-551<br>434-441<br>175-189<br>435-441<br>517-527<br>634-647 | 102504 |
| 67 | Citrate synthase | C. striatum | WP_110302223.1 | AQFNIFPR<br>GYDIADLANNATFNEVSYLLIK | 129-136<br>80-101 | 49415 |

| | | | | | | |
|---|---|---|---|---|---|---|
| | | | | HHTLLDEDFK<br>ITFINGEEGILR<br>LLILHADHEQNCSTSTVR<br>LMGFGHR<br>LRNETGLVTFDPGYAATGSTESK<br>NETGLVTFDPGYAATGSTESK<br>NNGGDATDFMNR<br>QIYTGETLR<br>QSTEGDSGFDIGK<br>QSTEGDSGFDIGKLR<br>VPMLAAYAYR | 119-128<br>66-77<br>232-249<br>312-318<br>43-65<br>45-65<br>291-302<br>421-429<br>30-42<br>30-44<br>178-187 | |
| 68 | Recombinase RecA | C. striatum | WP_166684297.1 | EGGIAAFIDAEHALDPEYAR<br>LMSQALR<br>SGSWFTYEGDQLGQGK | 97-116<br>181-187<br>298-313 | 41621 |
| 69 | Long-chain fatty acid--CoA ligase | C. flavescens<br>C. camporealensis<br>C. striatum<br>C. accolens<br>C. incognita<br>C. confusum<br>C. simulans<br>C. phocae<br>C. marquesiae<br>C. tuberculostearicum<br>C. aurimucosum<br>C. kefirresidentii<br>C. curieae<br>C. diphtheria<br>C. pseudotuberculosis | WP_276621821.1<br>WP_105360371.1<br>WP_204083944.1<br>WP_284900351.1<br>WP_246389116.1<br>WP_290222767.1<br>WP_062037925.1<br>WP_075733248.1<br>WP_284787534.1<br>WP_316970999.1<br>WP_049361583.1<br>WP_259825559.1<br>WP_269945698.1<br>WP_010935281.1<br>WP_075142747.1 | IGTVGPPLEGTTIR<br>NVSPGPMEDMLR<br>TDDLASLVYTSGTTGRPK<br>VAIEYSR | 406-419<br>484-495<br>189-206<br>318-324 | 66819 |
| 70 | Glutamate-5-semialdehyde dehydrogenase | C. striatum<br>C. simulans | WP_110378802.1<br>WP_062043585.1 | AAENLVAR<br>AAGMNDSLIDR<br>GPMALPELTSTK<br>NSNEQLVEILQDTLAHFDLPR<br>QVAALQDPVGEVLR<br>SGNVPLLR<br>TPEILEANER | 42-49<br>66-76<br>404-415<br>161181<br>93-106<br>147-154<br>50-59 | 45432 |
| 71 | Carboxyl transferase domain-containing protein | C. simulans | WP_248092398.1 | APQGEVPAAR<br>AQEGIVSSAMQAR<br>TVGIIANR | 23-32<br>129-141<br>276-283 | 46640 |
| 72 | Acetyl-CoA hydrolase/transferase family protein | C. simulans<br>C. ammoniagenes | WP_284841984.1<br>WP_040355625.1 | AAHAAGEEFK<br>FVNHGDR<br>IMNGTGGSGDFTR<br>SPYNSDPTLR<br>VVAVVETNAPDR | 66-75<br>35-41<br>392-404<br>102-111<br>227-238 | 56652 |
| 73 | Pyridoxal kinase PdxY | C. phoceense<br>C. lizhenjunii<br>C. flavescens<br>C. striatum | WP_257054465.1<br>WP_165009399.1<br>WP_276621636.1<br>WP_201816623.1 | ALYACDPVMGNAK<br>SGCFVSDEIPPLLR<br>TVLVTSVK | 108-120<br>121-134<br>179-186 | 30556 |

| | | | | | | |
|---|---|---|---|---|---|---|
| | | C. tuberculostearicum<br>C. minutissimum<br>C. marquesiae<br>C. aurimucosum<br>C. intestinale<br>C. kefirresidentii<br>C. singulare<br>C. hiratae | WP_301988405.1<br>WP_115023123.1<br>WP_284839065.1<br>WP_049157276.1<br>WP_250224256.1<br>WP_284835896.1<br>WP_239179489.1<br>WP_144013459.1 | | | |
| 74 | Histidinol-phosphate transaminase | C. striatum<br>C. simulans | WP_284788110.1<br>WP_248092287.1 | AAIAALNSAEELLAR<br>GNFVWIPR<br>LGLDAVIEAITDK<br>MAGAEPVMTPLTEDDRLGLDAVIEAITDK<br>SFEAYPLYVR<br>VAEAIGARPSK<br>YLDEGVR | 251-265<br>287-294<br>130-142<br>114-142<br>104-113<br>276-286<br>314-320 | 38183 |
| 75 | Long-chain fatty acid--CoA ligase | C. accolens<br>C. macginleyi | WP_284900351.1<br>WP_200445387.1 | ALDTPEGPDR<br>FYILESDLTEDENELTPTMK | 324-333<br>571-590 | 66550 |
| 76 | Orotate phosphoribosyltransferase | C. diphtheriae<br>C. striatum<br>C. minutissimum<br>C. aurimucosum<br>C. casei<br>C. tuberculostearicum<br>C. simulans<br>C. accolens<br>M. tuberculosis | WP_016830171.1<br>WP_086890651.1<br>WP_039672482.1<br>WP_102233499.1<br>WP_441526634.1<br>WP_204610871.1<br>WP_061924904.1<br>WP_284831249.1<br>WP_031663521.1 | ATGAADVIAAEGLEYR<br>DIDAFVVR<br>EADYYVDLR<br>EAGAEVVGVATVVDR<br>ELAVVHGK<br>RATLQHEASR | 155-170<br>89-96<br>32-40<br>140-154<br>17-24<br>41-50 | 19234 |
| 77 | GuaB3 family IMP dehydrogenase-related protein | C. simulans<br>C. striatum | WP_248092635.1<br>WP_110333579.1 | AIACGADSVVLGPVLAR<br>DSGVTVAVR<br>DYLDETGGR<br>ELAPVVIK<br>GYYWPSTAGHPR<br>QGGLGVINAEGLWGR<br>RDYLDETGGR<br>TYDLDQISLVPTRR<br>VTENPFDLSTLQELHAAPLNEELLAER | 270-286<br>128-136<br>243-251<br>144-151<br>294-305<br>71-85<br>242-251<br>16-29<br>96-122 | 40151 |
| 78 | Carboxymuconolactone decarboxylase family protein | C. simulans<br>C. striatum | WP_248091796.1<br>WP_049063413.1 | LNLGTLAR<br>SSLPEYAKDQK<br>STELNEEQLWGSMLAAAAATR | 19-26<br>8-18<br>27-47 | 18784 |
| 79 | Phosphoribosylamine--glycine ligase | C. aurimucosum<br>C. striatum<br>C. simulans | WP_216381141.1<br>WP_070446040.1<br>WP_248093127.1 | AYDNDEGPNTGGMGAYTPLPWLPADGVQR<br>EGIAVVEFNCR<br>KGDVITGPGLDDPNK<br>VLNVLGQGATLAEAR | 227-255<br>288-298<br>356-370<br>398-412 | 46398 |
| 80 | Class II fructose-bisphosphatase | C. striatum | WP_347004821.1 | GTMYDPSAVFYMNK<br>KNEGDGAAVDAMR<br>LMAEGRPNAISVIAAAER<br>NEGDGAAVDAMR | 118-131<br>37-49<br>100-117<br>38-49 | 35647 |

| | | | | | | |
|---|---|---|---|---|---|---|
| | | | | NLAMELVR<br>SENTSFMPDR<br>VTEAAALASGR<br>YIDSVHK | 12-19<br>2-11<br>20-30<br>313-319 | |
| 81 | Threonine synthase | C. simulans<br>C. striatum | WP_339018712.1<br>WP_114976447.1 | LIVATNENDVLDEFFR<br>TAELFGPK<br>TSSPSMDISR | 302-317<br>360-367<br>332-341 | 55191 |
| 82 | Acetyl/propionyl/methylcrotonyl-CoA carboxylase subunit alpha | C. accolens<br>C. simulans<br>C. flavescens<br>C. striatum | WP_284899407.1<br>WP_061920738.1<br>WP_301504211.1<br>HCG2963059.1 | GHAFEFR<br>LQVEHPVSEETTGLDLVR<br>LVEEAPAPFLSDEQR | 337-343<br>299-316<br>244-258 | 62305 |
| 83 | L-lactate dehydrogenase [Corynebacterium] | C. durum<br>C. diphtheria<br>C. belfantii<br>C. rouxii<br>C. canis<br>C. striatum<br>C. argentoratense<br>C. cystitidis<br>C. simulans<br>C. accolens<br>C. macginleyi | WP_006062201.1<br>WP_071575789.1<br>WP_197688803.1<br>WP_315644670.1<br>WP_146325394.1<br>WP_049151347.1<br>WP_021012357.1<br>WP_257183262.1<br>AMO89332.1<br>WP_302525282.1<br>WP_200446942.1 | ASGLPHER<br>DAAYTIIDAK<br>GSTSYGIGMGLAR<br>VIGSGTILDSAR | 139-146<br>223-232<br>233-245<br>147-158 | 34249 |
| 84 | Mannose-6-phosphate isomerase, class I | C. simulans<br>C. hiratae<br>C. aurimucosum<br>C. intestinale<br>C. singulare<br>C. minutissimum<br>C. flavescens<br>C. guaraldiae<br>C. hesseae | WP_248092243.1<br>WP_158397679.1<br>WP_049156068.1<br>WP_250224520.1<br>WP_042529785.1<br>WP_039674193.1<br>WP_301515692.1<br>WP_154762264.1<br>WP_339017302.1 | FVDVPELVK<br>ILAAAEPLSLQAHPSK<br>NYSWGSR | 289-297<br>82-97<br>10-16 | 43520 |
| 85 | MULTISPECIES: GTP cyclohydrolase I FolE | Corynebacterium | WP_082723330.1 | AEALSLIR<br>RPQVQER | 198-205<br>134-140 | 22305 |
| 86 | Type I polyketide synthase (Type I polyketide synthase 13, Pks13) | C. striatum<br>C.simulans<br>C. aurimucosum<br>C: minutissimum<br>M. leprae<br>M. tuberculosis<br>C. diphtheriae | WP_440214970.1<br>WP_062036057.1<br>WP_029158968.1<br>WP_039673467.1<br>WP_010908189.1<br>WP_052658316.1<br>WP_072567154.1 | 1 AAVMVPGIDVPFHSR<br>2 AGAALFQTGGIIDVFR<br>3 CHVVLPGSPNR<br>4 FRDEVVAR<br>5 GGFLEGEGGGTVLLVR<br>6 GLGLTPNDVSVLSK<br>7 GQQYSVAGTK<br>8 GSAMGSLVPR<br>9 GSIDLTTVFLDR<br>10 GTFGGDGAYGEVK<br>11 HDTSTNANDPNESELHSLLWPAIGR<br>12 IDGEVVSR<br>13 IPANVSLDCVDPLIAPK<br>14 LLLWSVER | 1499-1513<br>2856-2871<br>2167-2177<br>2414-2421<br>2729-2744<br>2759-2772<br>1449-1458<br>1391-1400<br>2437-2448<br>2178-2190<br>2812-2836<br>1241-1248<br>2836-2852<br>2183-2190 | 317703 |

| | | | | | | |
|---|---|---|---|---|---|---|
| | | | | 15 TAQFNTMFFDR<br>16 TRFEAEYGLQR<br>17 VMIDGYGLVDADK | 428-438<br>2355-2365<br>2993-3005 | |
| 87 | MULTISPECIES: NADP-dependent phosphogluconate dehydrogenase | Corynebacterium | WP_284775466.1 | AGSEQFGWDVDPR<br>DFFGAHTFK<br>DLATIWR<br>LIAYSQGFDEIK<br>LPAALIQGQR<br>MVHNGIEYADMQVIGEAYQLLR<br>NFAHNGNTVAVYNR | 348-360<br>452-460<br>361-367<br>336-347<br>442-451<br>188-209<br>25-38 | 51757 |
| 88 | Glucose-6-phosphate isomerase | C. simulans<br>C. tuberculostearicum | WP_062041602.1<br>WKS52910.1 | AVLHTALR<br>DAMFAGEHINNTEDR<br>EEAASGDQSTDELIGWYR<br>FPAYLQQLTMESNGK<br>HFVAVSTNAEK<br>QQANDLAPAVSGK<br>RDAMFAGEHINNTEDR<br>TFTTQETLTNAHAAR | 91-98<br>76-90<br>525-542<br>343-357<br>237-247<br>512-524<br>75-90<br>206-220 | 59473 |
| 89 | MULTISPECIES: 1,4-Dihydroxy-2-naphthoyl-CoA synthase | Corynebacterium | WP_034656212.1 | DGGWAFCSGGDQR<br>DPQWEQFPYYY<br>EIFFLGR<br>IAFDRPEVR<br>LHILEVQR<br>MYEMGAVNEVVDHADLEDAAIQMGR<br>QTDADVGSFDAGYGSAYLAK | 110-122<br>329-339<br>236-242<br>64-72<br>159-166<br>249-273<br>208-227 | 41932 |
| 90 | Malate dehydrogenase | C. striatum | WP_284810350.1 | AAFLVGARPR<br>ADLLEANGAIFTVQGK<br>GAEIIEVR<br>IAAGDVYGK<br>IQANVEELR<br>TAAVVSDGSYGVDEGLIFGFPTVAK<br>TLSQLSLK | 113-122<br>129-144<br>249-256<br>54-62<br>328-336<br>284-308<br>193-200 | 37652 |
| 91 | MULTISPECIES: Aldo/keto reductase | Corynebacterium | WP_070420669.1 | HIDTAAVYGNEEGVGR<br>YGVSPAQVIIR | 39-54<br>204-214 | 30570 |
| 92 | Carbamoyl-phosphate synthase large subunit | C. kroppenstedtii<br>C. simulans | WP_012731630.1<br>WP_284841318.1 | FAFEKFPGADDTLTTTMK<br>GMEIVYDEASLR<br>IAGGGLAASPEANVLIEESILGWK<br>NGVECEVVAK<br>VIILGSGPNR | 366-383<br>740-751<br>196-219<br>1015-1024<br>574-583 | 120034 |
| 93 | MULTISPECIES: Bifunctional cytidylate kinase/GTPase Der | C. dentalis<br>C. macclintockiae<br>C. jeikeium<br>C. argentoratense<br>C. variabile<br>C. flavescens<br>C. simulans<br>C. terpenotabidum<br>C. nuruki | WP_312097497.1<br>WP_284803594.1<br>WP_035011985.1<br>WP_314929774.1<br>WP_014009824.1<br>WP_312714526.1<br>WP_284841700.1<br>WP_041631109.1<br>WP_334143934.1 | GGADVLDEILR<br>LEPAMIEALESWDQR | 251-261<br>438-452 | 61161 |

| | | | | | | |
|---|---|---|---|---|---|---|
| | | C. incognita<br>C. hiratae<br>C. aurimucosum<br>C. stationis<br>C. striatum | WP_185175897.1<br>WP_144013232.1<br>WP_102234615.1<br>WP_305938727.1<br>WP_049062689.1 | | | |
| 94 | MULTISPECIES: FAD-dependent oxidoreductase | Corynebacterium | WP_066485343.1 | ELDHSPTINVVVDPEDIDYDEASLEAR<br>GIVNSLHR<br>GPVGLIGNTK<br>RGPVGLIGNTK<br>SQDLVCQTLEGYAIREPK<br>VAVIGAGPAGIYASDILVK<br>VLETEQIR | 211-237<br>60-67<br>363-372<br>362-372<br>242-259<br>8-26<br>68-75 | 48531 |
| 95 | MULTISPECIES: Rhodanese-related sulfurtransferase | Corynebacterium | WP_287134269.1 | CEILSALMK<br>DKPVVSYCTGGIR<br>VRDEIVAFGAPGELK | 191-199<br>178-190<br>91-105 | 34462 |
| 96 | MULTISPECIES: Oligoribonuclease | Corynebacterium | WP_049377255.1 | ALQDIIESIR<br>LDEALHYR<br>MIDVSTIK | 168-177<br>130-137<br>138-145 | 24319 |
| 97 | MULTISPECIES: Porphobilinogen synthase | Corynebacterium | WP_201054730.1 | AAAQNGWIDGEAVMLESLTAFK<br>AAHEALDAGVK<br>GNPTVEAEVFLDDGAR<br>CVDLFGVPR<br>EALDEAGHEDVAIMAYSAK<br>TYQQDPR<br>YASAFFGPFR | 251-272<br>30-40<br>274-289<br>85-93<br>147-165<br>188-194<br>166-175 | 31960 |
| 98 | MULTISPECIES: Histidine ammonia-lyase | Corynebacterium | WP_023021804.1 | DTLTHTAQVAER<br>EGALVEAVENATGK<br>ELAAAVDNPVVTK<br>NQVQDAYSVR<br>SHAAGTGPEVEEEVIR<br>VNGELQDAGEALAAAGIEPLQLR<br>YLSPEIEETVR | 302-313<br>501-514<br>314-326<br>282-291<br>88-103<br>170-192<br>487-497 | 54503 |
| 99 | MULTISPECIES: NDP-sugar synthase | Corynebacterium | WP_070446197.1 | DMGRPDDFVR<br>LVVGHVDNSYWR<br>NVYDKLR | 227-236<br>215-226<br>102-108 | 38412 |
| 100 | MULTISPECIES: NAD(P)-dependent alcohol dehydrogenase | Corynebacterium | WP_070456147.1 | DVDGTITQGGYAQK<br>GADVTVLSR | 121-134<br>199-207 | 37625 |
| 101 | MULTISPECIES: Protoporphyrinogen oxidase | Corynebacterium | WP_070420072.1 | GAPFQTFR<br>VLFATPAPTTAR | 214-221<br>270-281 | 48531 |
| 102 | MULTISPECIES: Phosphoenolpyruvate--protein phosphotransferase | Corynebacterium | WP_049157532.1 | AAGAEGQAAEVLGATAGMVK<br>DNEALLTR<br>IASDSQAEGIGLYR<br>QLDLPCIVAVGAELR | 66-85<br>363-370<br>281-294<br>198-212 | 59446 |
| 103 | Type I polyketide synthase (Type I polyketide | C. simulans<br>C. minutissimum<br>C. striatum | WP_248093856.1<br>WP_039672575.1<br>WP_005529493.1 | ASFVDNQILGK<br>DVLEISPAFK<br>EGLETEVEGNIR | 1494-1504<br>643-652<br>1301-1312 | 167690 |

| | | | | | | |
|---|---|---|---|---|---|---|
| | synthase 13, Pks13) | | | KVDPTESLLDLLAK<br>LEGSLEER | 928-941<br>1354-1361 | |
| 104 | MULTISPECIES: Acryloyl-CoA reductase | Corynebacterium | WP_049146748.1 | DAMALDGNK<br>SALSETGRPLQK<br>VDSESEYLQALGADHIIDR | 75-83<br>230-241<br>211-229 | 37042 |
| 105 | MULTISPECIES: Bifunctional methylenetetrahydrofolate dehydrogenase/methenyltetrahydrofolate cyclohydrolase | Corynebacterium | WP_005510041.1 | AFLVSNIVER<br>EGAAVLDVGVSR<br>FGVELDGAK<br>RSENSTVTLCHTGTK<br>SENSTVTLCHTGTK<br>AFLVSNIVER<br>DCEQLGINSIR<br>EGAAVLDVGVSR<br>IDPEKDADGLHPVNLGK<br>LAGDVAPDVWDK | 268-277<br>224-235<br>152-160<br>181-195<br>182-195<br>271-280<br>59-69<br>227-238<br>116-132<br>243-254 | 30476 |
| 106 | MULTISPECIES: Phosphoenolpyruvate--protein phosphotransferase | Corynebacterium | WP_049157532.1 | AAGAEGQAAEVLGATAGMVK<br>DNEALLTR<br>IASDSQAEGIGLYR<br>QLDLPCIVAVGAELR | 66-85<br>363-370<br>281-294<br>198-212 | 59446 |
| 107 | MULTISPECIES: Acryloyl-CoA reductase | Corynebacterium | WP_049146748.1 | DAMALDGNK<br>SALSETGRPLQK<br>VDSESEYLQALGADHIIDR | 75-83<br>230-241<br>211-229 | 37042 |
| 108 | MULTISPECIES: Bifunctional methylenetetrahydrofolate dehydrogenase/methenyltetrahydrofolate cyclohydrolase | Corynebacterium | WP_005510041.1 | AFLVSNIVER<br>EGAAVLDVGVSR<br>FGVELDGAK<br>RSENSTVTLCHTGTK<br>SENSTVTLCHTGTK<br>AFLVSNIVER<br>DCEQLGINSIR<br>EGAAVLDVGVSR<br>IDPEKDADGLHPVNLGK<br>LAGDVAPDVWDK | 268-277<br>224-235<br>152-160<br>181-195<br>182-195<br>271-280<br>59-69<br>227-238<br>116-132<br>243-254 | 30476 |
| 109 | MULTISPECIES: 4-Hydroxy-3-methylbut-2-enyl diphosphate reductase | Corynebacterium | WP_066837876.1 | EVLEFLDER<br>QLTLDATCPLVTK<br>YGAPIYVR<br>YVVESLAER<br>EVLEFLDER<br>QLTLDATCPLVTK<br>YGAPIYVR<br>YVVESLAER | 280-288<br>93-105<br>33-40<br>48-56<br>280-288<br>93-105<br>33-40<br>48-56 | 34689 |
| 110 | MULTISPECIES: Carbamoyl-phosphate synthase large subunit | Corynebacterium | WP_046202279.1 | DIVESIGGESAR<br>IAGGGLAASPEANVLIEESILGWK<br>TEFAGEDGVR<br>VIILGSGPNR | 139-150<br>196-219<br>513-522<br>574-583 | 120563 |
| 111 | MULTISPECIES: Glutamate-1-semialdehyde 2,1-aminomutase | Corynebacterium | WP_049151537.1 | AAVDGLSFGAPTEAETLLAER<br>AFGSVGGQAR<br>LVNSGTEATMSAVR | 89-109<br>36-45<br>121-134 | 45919 |

| | | | | | | |
|---|---|---|---|---|---|---|
| 112 | F0F1 ATP synthase subunit gamma | C. simulans<br>C. striatum | WP_062041127.1<br>WP_046645539.1 | AAELEALLK<br>AQELIATSR<br>EADVAGSWAGFSQDPSWETTHDVR<br>KQGYETVR<br>QAQITQEITEIVGGAGALAESAESD | 99-107<br>23-31<br>130-153<br>108-115<br>303-327 | 35986 |
| 113 | Alpha/beta fold hydrolase | C. simulans<br>C. striatum | WP_248092935.1<br>WP_284789470.1 | GSGFDSLGYLLEEPFR<br>IAEIIDHLDNSQEVLPTGER<br>SDFLHEVGAR<br>WILMDQR | 228-243<br>193-212<br>253-262<br>73-79 | 46494 |
| 114 | Succinyl-diaminopimelate desuccinylase | C. tuberculostearicum<br>C. simulans<br>C. accolens<br>C. marquesiae<br>C. kefirresidentii<br>C. aurimucosum<br>C. pseudogenitalium<br>C. curieae | WP_251067781.1<br>WP_284841394.1<br>WP_284610267.1<br>WP_269953325.1<br>WP_239208049.1<br>WP_049361454.1<br>WP_005323541.1<br>WP_269946029.1 | AWLGSNAAHK<br>YGWTDVAR | 191-200<br>311-318 | 39044 |
| 115 | MULTISPECIES: Exodeoxyribonuclease III | Corynebacterium | WP_284787228.1 | THVTEPER<br>AAFQMLEEAGLEEVTR<br>DEDVWDIEAFR<br>IATWNINSVR<br>LLLTGDFNIAPR<br>WTYFDYK | 173-180<br>181-196<br>160-170<br>3-12<br>148-159<br>204-210 | 30876 |
| 116 | Hydroxymethylbilane synthase | C. simulans<br>C. striatum | WP_248093625.1<br>WP_100088022.1 | FLLVVPER | 85-92 | 31068 |
| 117 | Alanine dehydrogenase | C. simulans<br>C. tuberculostearicum<br>C. aurimucosum | WP_284841757.1<br>WP_317000188.1<br>WP_049361669.1 | AEVTVLDLDPHVLQR<br>HGTLPLLTPMSQVAGR<br>LVREETVKK | 192-206<br>124-139<br>249-257 | 38521 |
| 118 | UDP-N-acetylmuramoyl-L-alanyl-D-glutamate--2,6-diaminopimelate ligase | C. ammoniagenes<br>C. singulare<br>C. simulans<br>C. accolens<br>C. minutissimum<br>C. striatum<br>C. hiratae<br>C. guaraldiae<br>C. aurimucosum<br>C. intestinale<br>C. hesseae<br>C. marquesiae<br>M. tuberculosis | WP_003845219.1<br>WP_084226191.1<br>WP_239238302.1<br>PCC82175.1<br>WP_039676100.1<br>WP_114976430.1<br>WP_328287701.1<br>WP_143334906.1<br>WP_158380985.1<br>MCL8493609.1<br>WP_101735578.1<br>WP_340418706.1<br>WP_078385592.1 | ADLVVVTDDNPR<br>LTTPEAPTLQELFAR<br>VGLIGTTGTR | 409-420<br>154-168<br>135-144 | 54353 |
| 119 | MULTISPECIES: Thiol peroxidase | Corynebacterium | WP_286955064.1 | DLPFAQER<br>FCAAEGIENVVSASDFR<br>AAGLDNTVVLSVSR | 120-127<br>128-144<br>71-84 | 21771 |
| 120 | Dihydroxyacetone kinase subunit DhaK | C. simulans<br>C. striatum<br>C. accolens | WP_284841216.1<br>WP_239299452.1<br>PCC82883.1 | EIPLVSADDVTDHLMEPILADLK<br>FTEPLFVAR<br>GVAGTLLVEK<br>LAGAAAERGDDLAAVTAVAK | 226-248<br>30-38<br>150-159<br>160-179 | 44663 |

| | | | | | | |
|---|---|---|---|---|---|---|
| | | | | RGVAGTLLVEK | 149-159 | |
| 121 | Branched-chain amino acid aminotransferase | C. simulans<br>C. striatum | WP_239238287.1<br>WP_200973417.1 | MLNYTVTR<br>QPDGSIATFRPEENAQR | 1-8<br>81-97 | 39992 |
| 122 | MULTISPECIES: Phosphomannomutase/ phosphoglucomutase | Corynebacterium | WP_256000583.1 | ADIGLAFDGDADR<br>ASNTEPLLR<br>GTDAWFNVR<br>GVVGEDIDENFVR | 237-249<br>423-431<br>414-422<br>18-30 | 48710 |
| 123 | MULTISPECIES: Bifunctional phosphoribosylaminoimidazolecarboxamide formyltransferase/IMP cyclohydrolase | Corynebacterium | WP_005391756.1 | ANTLADGANR<br>AVVQPGGSIRDEEVIEAAK<br>EMSYNNYQDADAAWR<br>YGENSHQSATVTR | 439-448<br>476-494<br>237-251<br>209-221 | 54832 |
| 124 | MULTISPECIES: Class 1b ribonucleoside-diphosphate reductase subunit beta | Corynebacterium | WP_179386927.1 | ALNNLGFEGLFPADETR<br>SHEYDEYIASHAEPVK<br>TLTEHEQQTTMR<br>WSEENENLQR | 271-287<br>2-17<br>57-68<br>134-143 | 37710 |
| 125 | Bifunctional 2-methylcitrate synthase/citrate synthase | C. simulans | WP_239239706.1 | GPLHGGANEAVMK<br>TAVSFIGSQDPEEYTK<br>VPTMEAAFR | 225-237<br>101-116<br>277-285 | 43080 |
| 126 | MULTISPECIES: Fumarylacetoacetate hydrolase family protein | Corynebacterium | WP_070432461.1 | EIGLPVFTQGAHPCVLGR<br>GAAGVVTDGGVR<br>QEPGSGTLGDILALR | 383-400<br>365-376<br>346-360 | 51946 |
| 127 | Transglycosylase domain-containing protein (penicillin-binding protein) | C. striatum | WP_034657322.1 | DLTVEEGAMLAGLIQSPSVLDPR<br>LTGDSTAGGGSTITQQYVK<br>NQVIAELER<br>NTLVGDEYSYVR<br>VGITEDQVATGGLK<br>WNYVLDGLVEMGDLDATQR | 182-204<br>101-119<br>261-269<br>120-131<br>270-283<br>214-232 | 75011 |
| 128 | Nucleoside-diphosphate kinase | C. simulans | WP_062043559.1 | GDFALTVGENIVHGSDSPESAER<br>QLAGGTDPVSK<br>TLILIKPDGVK | 108-130<br>90-100<br>8-18 | 14958 |
| 129 | Rhodanese-related sulfurtransferase | C. striatum<br>C. simulans | HAT1181239.1<br>WP_061925244.1 | CEILSALMK<br>DKPVVSYCTGGIR<br>VRDEIVAFGAPGELK | 191-199<br>178-190<br>91-105 | 34462 |
| 130 | 3,4-Dihydroxyphenylacetate 2,3-dioxygenase | C. simulans<br>C. striatum | WP_248092821.1<br>WP_046646444.1 | AVDYMEDLGFR<br>CAYMEIVVTDLEASR<br>DGSTVLDLDGNPVPLVER<br>HNIIAICDK<br>KEDSEEMPEWK<br>KPTVHDTAMTGGDGPR<br>LDHFNQITPDVPK<br>TDESEMAQTIGADGFSYTR<br>TNVQAPDILR<br>VQDPLGFPYEFFYEVEHR | 160-170<br>12-26<br>301-318<br>217-225<br>338-348<br>190-205<br>147-159<br>319-337<br>2-11<br>112-129 | 41172 |

| | | | | | | |
|---|---|---|---|---|---|---|
| 131 | Purine-nucleoside phosphorylase | C. striatum | WP_309221894.1 | FIADTFLEDVVQFNEVR<br>GAEIAETILLPGDPLR<br>QTAFHTMMK | 41-57<br>23-38<br>232-240 | 26957 |
| 132 | MULTISPECIES: Cysteine--1-D-myo-inosityl 2-amino-2-deoxy-alpha-D-glucopyranoside ligase | Corynebacterium | WP_070628554.1 | ELGTSQIDLFR<br>NPLDALIWR<br>VHYVQNITDVDDPLFER | 106-116<br>203-211<br>80-96 | 45268 |
| 133 | Sulfurtransferase | C. striatum<br>C. simulans | WP_306592448.1<br>WP_061920739.1 | LLDGGRDAWVTEER<br>LVSAPWLSAR<br>VIEVDEDSLLYDIGHVPTASR | 120-133<br>19-28<br>36-56 | 32542 |
| 134 | MULTISPECIES: 3-Deoxy-7-phosphoheptulonate synthase class II | Corynebacterium | WP_039676119.1 | EFVANSSAGAR<br>LGPGVTPEQAVEYAEK | 186-196<br>290-305 | 49055 |
| 135 | Phospho-sugar mutase | C. striatum<br>C. simulans<br>C. accolens | WP_244080695.1<br>WP_062043792.1<br>PCC81964.1 | ITSTPPTQLAGSEIAK<br>QLTGDETGALLGDYLAR<br>SADNIEEVDTR<br>YGSADFYQDAAEVISAAGGR | 456-471<br>317-333<br>195-205<br>94-113 | 55517 |
| 136 | MULTISPECIES: Bifunctional phosphoribosylaminoimidazolecarboxamide formyltransferase/IMP cyclohydrolase | Corynebacterium | WP_070508888.1 | AAHACDPLSAFGGVIAVNR<br>AVVQPGGSIRDEEVIEAAK<br>EMSYNNYQDADAAWR<br>GLANAQQLNGK<br>HANPCGIAVSDESIAAAHR<br>IADLGINVTPVEK<br>ILVAEHEAPAEEIK<br>SLDAAGVEIVSTGSTAAK | 286-304<br>478-496<br>238-252<br>227-237<br>267-285<br>45-57<br>343-356<br>27-44 | 54745 |
| 137 | MULTISPECIES: Formimidoylglutamase | Corynebacterium | WP_239191589.1 | ADQATSGTPFK<br>LALVDVVELNPTFDIDNR<br>RADQATSGTPFK | 167-177<br>284-301<br>166-177 | 32274 |
| 138 | MULTISPECIES: Alpha/beta hydrolase-fold protein | Corynebacterium | WP_070731646.1 | DWYSSPDKK<br>FPEVWALDGLR<br>SAAMPDKEQVVQIQLAR | 133-141<br>142-152<br>116-132 | 72563 |
| 139 | MULTISPECIES: Lactate utilization protein B | Corynebacterium | WP_005327611.1 | CDEVCPVKIPLSDVILENR<br>VTNATPPVESK | 388-406<br>410-420 | 56114 |
| 140 | NAD(P)/FAD-dependent oxidoreductase | C. striatum<br>C. simulans | WP_005528135.1<br>WP_248092293.1 | ELHWAEVDPTTADDLNK<br>GAYATSYDLGGLSR | 253-269<br>445-458 | 55257 |
| 141 | Aminopeptidase N | C. striatum<br>C. simulans | WP_204083832.1<br>WP_248092840.1 | AVLDDATDPDIR<br>DNTLTGAAAYLGVSR<br>QNFDGITYAK | 665-676<br>701-715<br>388-397 | 93277 |
| 142 | MULTISPECIES: Ppx/GppA phosphatase family protein | Corynebacterium | WP_012714802.1 | LEICPWALR<br>LEILSGEEEAR | 260-268<br>76-86 | 33106 |
| 143 | MULTISPECIES: Glycine cleavage | Corynebacterium | WP_023636012.1 | DITPVEAGMAR<br>LEAGMPLYGNELSR | 253-263<br>239-252 | 40429 |

| | | | | | | |
|---|---|---|---|---|---|---|
| | system aminomethyltransferase GcvT | | | | | |
| 144 | Cysteine synthase A | C. striatum | WP_049147014.1 | AVYNNILETIGNTPLVR<br>AYGAEIVLTPGAAGMK<br>IQGIGANFVPEVLDR<br>VESFNPANSVK<br>VILTMPETMSNER<br>YVSTVLFEDIR | 2-18<br>108-123<br>220-234<br>34-44<br>90-102<br>299-309 | 32886 |
| 145 | UTP--glucose-1-phosphate uridylyltransferase | C. simulans<br>C. striatum | WP_239240803.1<br>WP_284772678.1 | HDLGNPGGYIPANVDFGLHHEK<br>HFDAFPELVETLDAR<br>KVVGMVEKPEPEDAPSNLVATGR<br>LAVITAPNK<br>LAVITAPNKQEVMR<br>QLIAEYEAEK<br>VVGMVEKPEPEDAPSNLVATGR<br>YGPALYK | 264-285<br>75-89<br>194-216<br>61-69<br>61-74<br>296-305<br>195-216<br>286-292 | 33299 |
| 146 | 4-Aminobutyrate--2-oxoglutarate transaminase | C. striatum<br>C. simulans<br>C. aurimucosum | WP_306588433.1<br>WP_284840980.1<br>WP_102233987.1 | IAREELEPILGDER<br>KTALFNSGAEAVENAVK<br>LNEVTPGDHPK<br>WCNDNDVVFIVDEIQAGMMR | 350-363<br>123-139<br>112-122<br>248-267 | 47291 |
| 147 | Phosphoribosyltransferase | C. striatum<br>C. accolens | WP_086890390.1<br>PCC82976.1 | TLDLVVNLLAETAAEVK<br>VLVVDDVADSGK | 110-126<br>98-109 | 16344 |
| 148 | 4-Hydroxy-tetrahydrodipicolinate reductase | C. striatum<br>C. simulans<br>C. flavescens | WP_306496342.1<br>WP_248091696.1<br>HCG46562.1 | GADVDGVPVHAVR<br>LDAPSGTAIHTAQGIAAAR<br>QIAEHPGLTVGLEK<br>SSFTPGVLVGVR | 264-276<br>225-243<br>317-330<br>305-316 | 35314 |
| 149 | Alpha-1,4-glucan:maltose-1-phosphate maltosyltransferase | C. aurimucosum<br>C. simulans<br>C. striatum<br>C. accolens | WJY70269.1<br>WP_239238856.1<br>MDU3175173.1<br>PCC81918.1 | RENPALQQLR<br>TNPEFFTVLADGTIAYAENPPK | 548-557<br>319-340 | 75124 |
| 150 | Polyribonucleotide nucleotidyltransferase | C. simulans<br>C. striatum | WP_062038570.1<br>WP_086891186.1 | DLGVEVDLIPR<br>EGRPSTEAILACR<br>ETQEFPLFPPYTDDVFAAVEK<br>FVALTDILGAEDAFGDMDFK<br>GETQILGVTTLDMLK<br>MEQQIDSLTPVTGK<br>QAGGSVTTYLDEDTMLLSTTTASNQPR<br>TLCEAQAGLAER | 373-383<br>100-112<br>274-294<br>526-545<br>393-407<br>408-421<br>44-70<br>259-270 | 80303 |
| 151 | Type I glutamate--ammonia ligase | C. striatum<br>C. minutissimum | WP_086891004.1<br>WP_115022991.1 | DLPVSLDQALR<br>GIKEGYELGDPAEDDVHQLTR<br>LMFGNEAPGSATWGVSNR | 389-399<br>359-379<br>298-315 | 50298 |
| 152 | Bifunctional phosphoribosylaminoimidazolecarboxamide formyltransferase/IMP cyclohydrolase | C. amycolatum<br>C. jeikeium | WP_101739102.1<br>ASE57461.1 | DIFTEVIVAPSYEDGAVEVLK<br>EMSYNNYQDADAAWR<br>EVSVEMAEQVK<br>HANPCGIAVSDESIAAAHR | 320-340<br>242-256<br>309-319<br>271-289 | 55333 |
| 153 | MULTISPECIES: | Corynebacterium | WP_049147310.1 | ALENSFSTEEER | 257-268 | 32719 |

| | | | | | | |
|---|---|---|---|---|---|---|
| | Adenosine deaminase | | | ANFVPFVVDVR | 134-144 | |
| 154 | Methionine adenosyltransferase | C. simulans<br>C. striatum | WP_282439978.1<br>WP_275432357.1 | DLDLLRPIYR<br>TTGYVEIPQLVR | 369-378<br>65-76 | 44240 |
| 155 | MULTISPECIES: Argininosuccinate lyase | Corynebacterium | WP_151844318.1 | GFTLATDLAEWMVR<br>GLIDIVGPEVGGR | 371-384<br>98-110 | 51868 |
| 156 | LutB/LldF family L-lactate oxidation iron-sulfur protein | C. striatum | WP_284786984.1 | DAGSAIKEDVAAR<br>TAALSSPIGHQALK | 68-80<br>314-327 | 55253 |
| 157 | Uracil phosphoribosyltransferase | C. simulans<br>C. striatum | WP_248092469.1<br>MDU3175568.1 | LLDPPIIVPIIR<br>NEETHEPVPYLEALPEDLSGR | 49-60<br>86-106 | 22546 |
| 158 | Argininosuccinate synthase | C. simulans<br>C. accolens<br>C. minutissimum<br>C. striatum<br>C. singulare<br>C. guaraldiae<br>C. mastitides | WP_062034928.1<br>PCC82780.1<br>WP_239188692.1<br>WP_306496646.1<br>WP_144789996.1<br>WP_143334879.1<br>WP_337890224.1 | AHEALEDVTVER<br>AVETGFLEDLWNPPTK<br>EVYEAPGAMVLIK | 282-293<br>184-199<br>269-281 | 44138 |
| 159 | Phosphoglycerate dehydrogenase | C. simulans<br>C. striatum<br>C. aurimucosum | WP_248090503.1<br>WP_046645907.1<br>WP_010186895.1 | AGVGLDNVDIPAATER<br>GGLVDEAALAESIK<br>LGAAGINIVAAALTQGK | 72-87<br>233-246<br>474-490 | 54713 |
| 160 | Disulfide bond formation protein DsbA | C. casei | WP_098072481.1 | DITVEFVPMSLAVLNDGR<br>LGDTAFFGPVITR | 31-48<br>156-168 | 22890 |
| 161 | MULTISPECIES: Flavodoxin-dependent (E)-4-hydroxy-3-methylbut-2-enyl-diphosphate synthase | Corynebacterium | WP_046645290.1 | TVPESQIVETLIEEAMR<br>KLEIVSCPSCGR | 262-273<br>336-352 | 41492 |
| 162 | Pyridoxal 5'-phosphate synthase lyase subunit PdxS | C. simulans<br>C. striatum | WP_284841028.1<br>WP_046646570.1 | MSDPDLIEGIVDAVSIPVMAK<br>SKGEAGTGDVSEAVK | 46-66<br>133-147 | 29313 |
| 163 | Glycogen synthase | Corynebacterium | WP_049160589.1 | DGVFWVQEMLSR<br>EQLGGGYEISSWSEK | 256-267<br>122-136 | 41687 |
| 164 | MULTISPECIES: Acyl-CoA carboxylase subunit beta | Corynebacterium | WP_062036855.1 | EVTGEEITSHDLGSAR<br>IGCPVIGIQDSGGAR<br>IQDAVTSLAMYSEIAR<br>TPGDPDAIYSDGVVTGYGR | 198-213<br>116-130<br>131-146<br>60-78 | 55456 |
| 165 | MULTISPECIES: Aspartate kinase | Corynebacterium | WP_023020397.1 | VTVLGIPDRPGEAAK | 267-281 | 44727 |
| 166 | MULTISPECIES: Biotin carboxylase N-terminal domain-containing protein | Corynebacterium | WP_061920870.1 | LIEEAPAPALSDEQR<br>VVENLEDIEAAFNSAGR | 238-252<br>170-186 | 63265 |
| 167 | Mycothiol conjugate amidase Mca | Corynebacterium | WP_284772865.1 | LWPTEEFELAQTR | 265-277 | 34614 |
| 168 | MULTISPECIES: | Corynebacterium | WP_003862691.1 | EYNLLILADEIYDR | 204-217 | 45081 |

| | | | | | | |
|---|---|---|---|---|---|---|
| | Pyridoxal phosphate-dependent aminotransferase | | | LNTGNPAVFGFDAPDVIMR<br>VVTLPWASQLENAIER | 43-61<br>384-399 | |
| 169 | Galactose-1-phosphate uridylyltransferase | C. striatum<br>C. simulans<br>C. accolens | WP_110333530.1<br>WP_239239739.1<br>PCC83444.1 | IIAETEHFTAFVPAAAK<br>TSALSALPEVAQVFPFENR | 232-248<br>160-178 | 40729 |
| 170 | MULTISPECIES: SufS family cysteine desulfurase | Corynebacterium | WP_023021514.1 | GSYQLAEEADDAYESAR<br>NATEALNEVAYVLSDER | 59-75<br>94-110 | 45319 |
| 171 | Methyltransferase domain-containing protein | C. striatum | WP_086891631.1 | LPLQDDSVDAISVVFAPR | 155-172 | 30527 |
| 172 | MULTISPECIES: NADP-specific glutamate dehydrogenase | Corynebacterium | WP_061923312.1 | AANAGGVATSALEMQQNASR<br>EIGYLFGQFR<br>FLGFEQIFK<br>HESGVLTGK<br>LCEPER<br>NSLTGLPIGGGK<br>VQFNSALGPYK | 371-390<br>175-184<br>108-116<br>190-198<br>54-59<br>117-128<br>82-92 | 48723 |
| 173 | MULTISPECIES: NADPH-dependent FMN reductase | Corynebacterium | WP_005324045.1 | DYEANILEIR<br>FLALVDNALTS<br>NAVDIGSKPNSDVAWK<br>NLPAGIISHSVGR<br>NSSGVLFITPENNR | 5-14<br>152-162<br>67-82<br>83-95<br>46-59 | 18027 |
| 174 | MULTISPECIES: Glutamate 5-kinase [Corynebacterium] | Corynebacterium | WP_151844467.1 | RPTDLATK | 82-89 | 38560 |
| 175 | Biotin--[acetyl-CoA-carboxylase] ligase | C. efficiens | WP_006769578.1 | QPLDITR | 13-19 | 29717 |
| 176 | Serine/threonine protein kinase | C. striatum<br>C. simulans<br>C. accolens | WP_284772611.1<br>WP_284841122.1<br>PCC82183.1 | FADAGEFLDALDDVTR | 258-273 | 48713 |
| 177 | Phosphoglycerate kinase | C. simulans<br>C. aurimucosum<br>C. accolens | WP_248090357.1<br>WP_102233912.1<br>PCC81818.1 | AQASVYDVAK<br>EVETLAAVAEKPEHPYVVVLGGAK<br>ITASLPTIK<br>TVFWNGPMGVFEMEAFSK | 163-172<br>185-208<br>41-49<br>320-337 | 42525 |
| 178 | MULTISPECIES: Triose-phosphate isomerase | Corynebacterium | WP_070976872.1 | IVVAYEPVWAIGTGK<br>KPLIAGNWK | 167-181<br>4-12 | 27279 |
| 179 | Peroxiredoxin | C. aurimucosum<br>C. simulans<br>C. minutisimum<br>C. tuberculostearicum | WP_216379644.1<br>WP_096337156.1<br>WP_115022497.1<br>WP_301980348.1 | ALGVENADGVADR<br>ATFIIDPDGVIQFVSVTPDAVGR<br>ATHPELK<br>DFTFVCPTEIAAFGK<br>EVPFPMFSDIR<br>FPEFELTALK<br>NVDEVLR<br>PILTVGEKFPEFELTALK<br>VLDALQSEEVCACNWEANDPTK | 121-133<br>134-156<br>98-104<br>56-70<br>105-115<br>10-19<br>157-163<br>2-19<br>164-185 | 22185 |

| | | | | | | |
|---|---|---|---|---|---|---|
| | | | | VVFFYPK | 49-55 | |
| 180 | Phosphoserine transaminase | C. argentoratense | WP_234653563.1 | FIPSFLNLQTAVDNSR | 226-241 | 40040 |
| 181 | MULTISPECIES: Glutamine-hydrolyzing carbamoyl-phosphate synthase small subunit | Corynebacterium | WP_168163245.1 | IPAILVLADGR | 23-33 | 43784 |
| 182 | Biotin carboxylase N-terminal domain-containing protein | C. aurimucosum<br>C. minutissimum<br>C. simulans | WP_253287036.1<br>WP_115021249.1<br>WP_062035764.1 | FVVEVGGR | 479-486 | 63355 |
| 183 | Depupylase/deamidase Dop | C. halotolerans | WP_027004154.1 | ELTALEVLAEYR | 310-321 | 57555 |
| 184 | MULTISPECIES: UDP-glucose 4-epimerase GalE | Corynebacterium | WP_187441025.1 | AYGLAATSLR | 156-165 | 35355 |
| 185 | Imidazolonepropionase | C. simulans<br>C. accolens | WP_239238774.1<br>PCC82950.1 | LEQLLVER | 107-114 | 42601 |
| 186 | Phosphoenolpyruvate carboxykinase (GTP) | C. diphtheria<br>C. simulans<br>C. striatum | WP_070794800.1<br>WP_248093862.1<br>WP_244080864.1 | AEDIDLEGIDFDIEDVR<br>EEGWMAEHMLILK<br>EIWSYGSGYGGNAILAK<br>FLWPGFGENSR<br>IDAILFGGR<br>KCYALR<br>MPSIFLVNWFR<br>RPNSYLAR<br>TMYVVPFCMGPISDPEPK<br>YITQFPETK | 544-560<br>242-254<br>212-228<br>508-518<br>411-419<br>229-234<br>491-501<br>66-73<br>124-141<br>203-211 | 67023 |
| 187 | MULTISPECIES: Lipoyl synthase | Corynebacterium | WP_049152157.1 | DDLPDEGAWLYAEVVR | 126-141 | 39952 |
| 188 | Pyruvate dehydrogenase | C. ulcerans | WP_046096294.1 | EQVLALAQK | 217-225 | 62233 |
| 189 | Type I glyceraldehyde-3-phosphate dehydrogenase | C. striatum<br>C. tuberculostearicum<br>C. simulans | WP_005531614.1<br>WP_251067654.1<br>WP_284841327.1 | AAAVNMVPTSTGAAK<br>HNVISAASCTTNCLAPMAK<br>IAVSAERDPK<br>KVIISAPGK<br>LDGYAMR<br>TTSLVASKL<br>VGINGFGR<br>YDSVLGR | 201-215<br>145-163<br>73-82<br>117-125<br>228-234<br>327-335<br>5-12<br>48-54 | 36061 |
| 190 | Pyrroline-5-carboxylate reductase | C. ulcerans<br>C. afermentas<br>C. genitalium<br>C. simulans<br>C. striatum | AKA95861.1<br>WP_286131553.1<br>RUQ13774.1<br>AMO88801.1<br>MDU3173891.1 | ELEESGLR | 242-249 | 28603 |
| 191 | Decaprenylphospho-beta-D-erythro-pentofuranosid-2-ulose | C. kroppenstedtii<br>C. parakroppenstedtii | WP_012732653.1<br>WP_221913303.1 | EDVAAAVAK | 210-218 | 27052 |

| | | | | | | |
|---|---|---|---|---|---|---|
| | 2-reductase | | | | | |
| 192 | Transglycosylase domain-containing protein | C. simulans<br>C. striatum | WP_062036690.1<br>MDU3174453.1 | TGTTQLGDTGNNK | 569-581 | 76871 |
| 193 | MULTISPECIES: Peptidylprolyl isomerase | Corynebacterium | WP_083319062.1 | EDVIIESIEIA<br>GEQSGPFYDGAIFHR<br>IIDNFMIQGGDPTGTGR | 166-176<br>53-67<br>68-84 | 19129 |
| 194 | Phosphoribosylformyl glycinamidine synthase subunit PurL | C. bovis | WP_125187423.1 | LAGALTR | 597-603 | 83909 |
| 195 | Thiamine pyrophosphokinase C terminal family protein | C. simulans<br>C. striatum<br>C. stationis | KXU16798.1<br>WP_086892412.1<br>WP_191374105.1 | VILPADPDGTAEGLER | 248-263 | 39147 |
| 196 | MULTISPECIES: Cysteine--1-D-myo-inositol 2-amino-2-deoxy-alpha-D-glucopyranoside ligase | Corynebacterium | WP_144014470.1 | LVEAGVDPSAIR | 304-315 | 45060 |
| 197 | Class I SAM-dependent methyltransferase | C. striatum | WP_070420299.1 | EVTAANPAHSENFAR | 6-20 | 21690 |
| 198 | Bifunctional folylpolyglutamate synthase/dihydrofolate synthase | C. striatum<br>C. simulans | WP_272707595.1<br>WP_275437165.1 | APETQIAPSLER | 71-82 | 54524 |
| 199 | Pyruvate dehydrogenase [ubiquinone] | C. jeikeium | WCZ53282.1 | TVTLFCGEGVK | 203-213 | 63088 |
| 200 | FAD-dependent oxidoreductase | C. striatum<br>C. simulans | WP_049151900.1<br>WP_062041830.1 | GPVGLIGNTK<br>RGPVGLIGNTK<br>SQDLVCQTLEGYAIREPK<br>TELTGDGNVR | 361-370<br>360-370<br>240-257<br>289-298 | 49848 |
| 201 | Glycosyltransferase 87 family protein | C. maris | WP_020934607.1 | TVPTSAGSVAVER | 416-428 | 10433 |
| 202 | Arginine deiminase | C. striatum<br>C. simulans<br>C. accolens | WP_086891892.1<br>WP_062041915.1<br>WP_302526629.1 | HQETMLTR | 197-204 | 47023 |
| 203 | Malate synthase G | C. pseudodiphtheriticum | WP_284814861.1 | VAFINTGFLDR | 453-463 | 82269 |
| 204 | MULTISPECIES: tRNA (N6-isopentenyl adenosine(37)-C2)-methylthiotransferase MiaB | Corynebacterium | WP_168926545.1 | LVALQDSIQAEENAK | 400-415 | 56262 |
| 205 | MULTISPECIES: Threonine synthase | Corynebacterium | WP_061923925.1 | DMAMQLLGELFEYELGR<br>LIVATNENDVLDEFFR | 119-135 | 52096 |
| 206 | Ornithine carbamoyltransferase | C. striatum<br>C. simulans | HAT1361402.1<br>WP_062038948.1 | ESIEDTAR<br>KESIEDTAR<br>NIVLLFEK | 89-96<br>88-96<br>48-55 | 37139 |

| | | | | | | |
|---|---|---|---|---|---|---|
| | | | | PVSLSGR<br>VFDEAENR | 2-8<br>312-319 | |
| 207 | MULTISPECIES: UDP-galactopyranose mutase [Corynebacterium] | Corynebacterium | WP_066796772.1 | DKTVIMK<br>ENNVLFGGR<br>QWQTDPK<br>YTFNNR<br>YYSPDEAR | 310-316<br>354-362<br>165-171<br>185-190<br>116-123 | 46713 |
| 208 | MULTISPECIES: Ppx/GppA phosphatase family protein | Corynebacterium | WP_062035531.1 | LGQGVDATGEFAPEALER | 38-55 | 34817 |
| 209 | MULTISPECIES: UDP-N-acetylglucosamine 2-epimerase (non-hydrolyzing) | Corynebacterium | WP_259885982.1 | QNTERPEAVVAGTVK | 151-165 | 43078 |
| 210 | MULTISPECIES: Aspartate carbamoyltransferase catalytic subunit | Corynebacterium | WP_023030686.1 | DAIVMHPGPMLR<br>EYANLYGLSK<br>IVGDCLHSR<br>MNGGFFPSHR<br>TVFTLFYENSTR | 263-274<br>245-254<br>167-175<br>235-244<br>44-55 | 34349 |
| 211 | Histidine phosphatase family protein | C. glyciniphilum | WP_304032479.1 | QTAAGIALGGSGR | 75-87 | 32468 |
| 212 | MULTISPECIES: Serine hydroxymethyltransferase | Corynebacterium | WP_066491318.1 | AAGVDVLTGGTDVHLVLADLR<br>AVLQAQGSVLTNK<br>ELDPDVFNAINGEIAR<br>IAASEEFK<br>IAEQYPLYDGLEDWK<br>IDMDKLR<br>IGTPALATR<br>LYDVAAYEVDPETMR<br>NAVPNDPRPPMVTSGLR<br>NSELDGQQAEDLLHEVGITVNR<br>SGLILAK<br>SIADEVGAK<br>TLDFAAFR<br>VIIAGWSAYPR<br>YAEGYPGR<br>YAEGYPGRR<br>YYGGCEHVDVIEDLAR | 313-333<br>48-60<br>16-31<br>282-289<br>417-431<br>160-166<br>373-381<br>145-159<br>356-372<br>334-355<br>243-249<br>195-203<br>187-194<br>176-186<br>61-68<br>61-69<br>70-85 | 46651 |
| 213 | MULTISPECIES: tRNA (guanosine(46)-N7)-methyltransferase TrmB | Corynebacterium | WP_239212011.1 | MFAPESLDGIR | 148-158 | 29940 |
| 214 | MULTISPECIES: GNAT family N-acetyltransferase | Corynebacterium | WP_049063205.1 | VGTYSHMAK<br>AIYEEGLNTGHATYETR | 146-154<br>25-41 | 21503 |
| 215 | MULTISPECIES: | C. diphtheriticum | WP_284862254.1 | YNLAESFPLLTTK | 41-53 | 30485 |

| | | | | | | |
|---|---|---|---|---|---|---|
| | Thymidylate synthase | C. tuberculostearicum<br>C. aurimucosum<br>C. accolens<br>Corynebacterium | WP_316971870.1<br>WP_275042102.1<br>WP_284636153.1<br>WP_201054696.1 | | | |
| 216 | Hytoene/squalene synthase family protein | C. maris | WP_020934521.1 | AVLAAPGK | 84-91 | 32455 |
| 217 | ATP-dependent 6-phosphofructokinase | C. simulans<br>C. minutissimum<br>C. faecipullorum | WP_062041068.1<br>WP_039675088.1<br>HIX78424.1 | EGTMDFEEGGVDQFGHQTFNGIGQVIGDEIK<br>GGTPTAYDR<br>ILIVEVMGR<br>LAVLTSGGDCPGLNAVIR<br>TTVLGHIQR | 229-259<br>277-285<br>164-172<br>3-20<br>268-276 | 36729 |
| 218 | Inositol monophosphatase family protein | C. aurimucosum<br>C. guaraldiae<br>C. intestinale<br>C. minutissimum<br>C. singulare | WP_193629958.1<br>WP_154762278.1<br>WP_250224758.1<br>KKO80862.1<br>WP_042529917.1 | QVVDQMEEWLR | 264-274 | 29556 |
| 219 | MULTISPECIES: LLM class F420-dependent oxidoreductase | Rhodococcus<br>Prescottella equi | NKR92484.1<br>WP_213573252.1 | YDAPLGR | 107-113 | 37344 |
| 220 | MULTISPECIES: Phosphate acetyltransferase | Corynebacterium | WP_060565243.1 | LWAFGDCAVNPNPTAEQLGEIAVVSAK<br>TAAQFGIDPR | 164-190<br>191-200 | 49084 |
| 221 | Branched-chain amino acid aminotransferase | C. simulans<br>C. glutamicum<br>C. accolens | WP_062043898.1<br>WP_044027284.1<br>PCC82201.1, | GCDQVVWLDAIEHK | 233-246 | 40373 |
| 222 | Pyruvate carboxylase | C. simulans<br>C. pseudodiphtheriticum<br>C. accolens | WP_275437190.1<br>WP_284845159.1<br>PCC83475.1 | ICADAVR | 261-267 | 123560 |
| 223 | MULTISPECIES: Aldo/keto reductase | Corynebacterium | WP_042533129.1 | EAIELGYR | 38-45 | 32574 |
| 224 | MULTISPECIES: Transketolase | Corynebacterium | WP_070539739.1 | LGNLIVFWDDNR<br>NLHFGIR<br>QLEAEGTATR<br>TVIGYPAPNK | 200-211<br>436-442<br>604-613<br>270-279 | 75584 |
| 225 | Maltotransferase domain-containing protein | C. kefirresidentii<br>C. tuberculostearicum<br>C. marquesiae<br>C. simulans<br>C. accolens<br>C. striatum | WP_086588781.1<br>WP_316971224.1<br>WP_284786675.1<br>WP_284841273.1<br>PCC81918.1<br>MDU3175173.1 | VADMGFDTVYFPPIHPIGK | 232-250 | 74660 |
| 226 | MULTISPECIES: Electron transfer flavoprotein subunit beta/FixA family protein | Corynebacterium | WP_101736622.1 | TSVDNVIDEVNEYSVEQALR | 29-48 | 27527 |
| 227 | MULTISPECIES: | Corynebacterium | WP_136651187.1 | DLYETFEALAR | 375-385 | 44008 |

| | | | | | | |
|---|---|---|---|---|---|---|
| | Phosphotransferase | | | | | |
| 228 | MULTISPECIES: Phosphoglycerate kinase | Corynebacterium | WP_061922322.1 | GAFADELAALAADNGAFVSDGFGVVHR | 136-162 | 43078 |
| 229 | D-isomer specific 2-hydroxyacid dehydrogenase family protein | C. wankanglinii<br>C. halotolerans<br>C. cystitides | WP_268908204.1<br>WP_015401601.1<br>WP_257162414.1 | LIEMLR | 143-148 | 32860 |
| 230 | MULTISPECIES: Alpha/beta hydrolase family protein | Corynebacterium | WP_021352819.1 | YLQGPQK | 163-169 | 39715 |
| 231 | Gycosyltransferase | C. amycolatum | WP_253291995.1 | NSITLGR | 39-45 | 71677 |
| 232 | 16S rRNA (cytosine(1402)-N(4))-methyltransferase RsmH | C. pseudotuberculosis | WP_014800655.1 | MAELIAAPVIK | 27-37 | 38580 |
| 233 | Adenylate kinase | C. striatum<br>C. simulans<br>C. phoceense | WP_101504604.1<br>WP_096336255.1<br>WP_257035894.1 | AAEAAGADFVK<br>TGADAQAMIDAGATR<br>AEGTVEEINER<br>GRADDNEETIR<br>GTQAAILSEK<br>LNQDDAANGFLLDGFPR<br>SYIDAGK<br>YVLLGPPGAGK | 132-142<br>179-193<br>164-174<br>128-138<br>14-23<br>72-88<br>51-57<br>3-13 | 21121 |
| 234 | MULTISPECIES: Dyp-type peroxidase | Corynebacterium | WP_062040470.1 | QFEPIQAR | 356-363 | 45292 |
| 235 | Malate synthase G | C. maris | WP_020934049.1 | YALNAANAR | 129-137 | 79730 |
| 236 | MULTISPECIES: Glycine cleavage system protein GcvH | Corynebacterium | WP_066488580.1 | VGITSVAADR | 30-39 | 13311 |
| 237 | Nucleoside triphosphate pyrophosphatase | C. glucuronolyticum | WP_201839608.1 | ERVAALAQAK | 49-58 | 22146 |
| 238 | MULTISPECIES: Decaprenylphospho-beta-D-erythro-pentofuranosid-2-ulose 2-reductase | Corynebacterium | WP_061919989.1 | AGVDGFFINLGEALR | 163-177 | 26725 |
| 239 | MULTISPECIES: Phosphoglyceromutase | Corynebacterium | WP_070563655.1 | DKYGEEQFMAWR<br>HWIPVVR<br>HYGALQGLNK<br>TANIALNAADR<br>TECLKDVVER<br>YGEEQFMAWR | 109-120<br>81-87<br>95-104<br>70-80<br>153-162<br>111-120 | 27922 |
| 240 | Transaldolase | C. striatum<br>C. accolens<br>C. simulans | WP_049145842.1<br>CC81812.1<br>WP_239238973.1 | SHIDELAQLGTSTWLDDLSR<br>SIVGVTTNPAIFAAAMSK<br>VNRPNLMIK | 2-21<br>37-54<br>131-139 | 38551 |
| 241 | MULTISPECIES: | Corynebacterium | WP_066486562.1 | AGIGGDGLLR | 38-47 | 30138 |

| | | | | | | |
|---|---|---|---|---|---|---|
| | Diaminopimelate epimerase | | | | | |
| 242 | Mycothione reductase | C. ammoniagenes<br>C. casei | WP_168938634.1<br>MDO5507660.1 | IDKIAEGGEAYR<br>TTAEGIWALGDVSSPYMLK | 89-100<br>293-311 | 49798 |
| 243 | tRNA dihydrouridine synthase DusB | C. glucuronolyticum | WP_276542633.1 | GVSGTDIPVTVK | 141-152 | 43656 |
| 244 | MULTISPECIES: UDP-N-acetylmuramate dehydrogenase | Corynebacterium | WP_225723520.1 | RPVAEVR | 219-225 | 39483 |
| 245 | Inositol monophosphatase family protein | C. diphtheriae | WP_014308398.1 | MAKTSLK | 1-7 | 31462 |
| 246 | Branched-chain-amino-acid transaminase | C. durum | WP_315559093.1 | FAGNYAASLVAQAQAAEK | 205-222 | 39931 |
| 247 | tRNA adenosine deaminase-associated protein | C. striatum<br>C. simulans | WP_086892281.1<br>WP_239240418.1 | MLISDATYADEDDFAADFLELR<br>NEGQWVVR | 70-91<br>15-22 | 17808 |
| 248 | Glutamate synthase-related protein | C. terpenotabidum | WP_020441182.1 | LTGTVDAADPR | 1751-1761 | 200354 |
| 249 | Succinate CoA transferase | C. glutamicum | WP_143854664.1 | MAVASATSFALSPEYAEK | 307-324 | 55124 |
| 250 | MULTISPECIES: Pyruvate dehydrogenase (acetyl-transferring), homodimeric type | Corynebacterium | WP_003856989.1 | ANDGAYVR<br>DTSDQHVWAFLGDGEMDEPESR<br>GAGWNVIK<br>GFLIGATAGR<br>GYGLGHNFEGR<br>KLTLEDLK<br>LTLEDLK<br>LVPIIPDEAR | 347-354<br>238-259<br>304-311<br>645-654<br>413-423<br>431-438<br>432-438<br>539-548 | 103480 |
| 251 | SDR family oxidoreductase | C. glyciniphilum, Tsukamurella paurometabola | WP_145943527.1<br>WP_126195033.1 | GAAIELAR | 17-24 | 27559 |
| 252 | 2,3,4,5-Tetrahydropyridine-2,6-dicarboxylate N-succinyltransferase | C. auromucosum<br>C. minutissimum | WP_201828409.1<br>WP_115021663.1 | SEAVALNEDLHKN | 311-322 | 33581 |
| 253 | MULTISPECIES: Mycothione reductase | Corynebacterium | WP_071572518.1 | HVANAEMR | 323-330 | 51389 |
| 254 | MULTISPECIES: Resuscitation-promoting factor | Corynebacterium | WP_301980636.1 | EQQIAVAEK | 136-144 | 40648 |
| 255 | 16S rRNA (cytosine(1402)-N(4))-methyltransferase RsmH<br>*iron ABC transporter* | C. otitides<br>*C. casei* | WP_046644458.1<br>*WP_006822721.1* | LAHFGGR | 145-151 | 35688 |

| | | | | | | |
|---|---|---|---|---|---|---|
| | *permease* | | | | | |
| 256 | DNA cytosine methyltransferase | C. variabile | WP_148263290.1 | LISEVRPR | 157-164 | 50961 |
| 257 | SDR family oxidoreductase | C. kroppenstedtii | WP_012730858.1 | AAHPYLSR | 121-128 | 28738 |
| 258 | MULTISPECIES: Glycerol kinase GlpK | Corynebacterium | WP_075730700.1 | AVLEATAYQTR<br>DVADAMVADSGVEITELR<br>ETTVVWDK<br>ETTVVWDKNTGEPVYNAIVWQDTR<br>FSENGLLTTVCFQR<br>GSLAGVPIR<br>GVIVGLTR<br>NTGEPVYNAIVWQDTR<br>NTYGTGLFLLLNTGTTPK<br>TLLMDIEKLEWDEELCK | 388-398<br>399-416<br>85-92<br>85-108<br>289-302<br>240-248<br>371-378<br>93-108<br>271-288<br>198-214 | 57012 |
| 259 | 3-Isopropylmalate dehydratase large subunit | C. casei<br>C. marinum<br>C. auriscanis<br>C. comes | WP_276687832.1<br>WP_042621303.1<br>WP_035115166.1<br>WP_156227866.1 | AAAEVLEGR | 349-357 | 52503 |
| 260 | Carbonic anhydrase | C. viitaeruminis | WP_276651887.1 | ILQISPAIK | 167-175 | 17180 |
| 261 | MULTISPECIES: Fumarate reductase/succinate dehydrogenase flavoprotein subunit | Corynebacterium | WP_042529063.1 | EYGGTLATR<br>GQTGQQLQLSTTSALYR<br>HAEPLYFDSIPLMTR<br>ILESHEPHGVPMK<br>ITGNANDMNQVLEYGLR<br>KVDNDSAYR<br>NLITGELK<br>NSNAGAMMR<br>VIDHMNAIGAPFAR<br>YPAFGNLVPR<br>YSNLIEMYEEAIGESAYETPMR | 173-181<br>196-212<br>663-677<br>37-49<br>585-601<br>123-131<br>248-255<br>281-289<br>159-172<br>352-361<br>414-435 | 75852 |
| 262 | Aminotransferase class I/II-fold pyridoxal phosphate-dependent enzyme | C. matruchotii | WP_005524031.1 | ELAAMDTAAPDFR<br>WLCPVPGYDR | 216-228<br>147-156 | 46146 |
| 263 | MULTISPECIES: Aldehyde dehydrogenase family protein | Corynebacterium | WP_200288411.1 | VVDPQDPNLR | 147-156 | 56195 |
| 264 | MULTISPECIES: UMP kinase | Corynebacterium | WP_049063482.1 | AVDGVYSDDPR<br>TNPDAELYSEITPR | 170-180<br>181-194 | 24521 |
| 265 | Class II fumarate hydratase | C. accolens | WP_284609611.1 | AKEFADVVK<br>DGLVEFSGAMR<br>DLQPGSSIMPGK<br>DSGSLDAEKADAIIAAAK | 168-176<br>266-276<br>307-318<br>61-78 | 49819 |
| 266 | MULTISPECIES: Aminotransferase class I/II-fold | Corynebacterium | WP_256021930.1 | ELVKDPQVK<br>HAALLAPK<br>WLCPVPGYDR | 169-177<br>312-319<br>131-140 | 45949 |

| | | | | | | |
|---|---|---|---|---|---|---|
| | pyridoxal phosphate-dependent enzyme | | | | | |
| 267 | Aconitate hydratase | C. striatum<br>C. simulans | WP_236593737.1<br>WP_235597050.1 | ATIGNMSPEFGSTAAIFPIDEETIK<br>AVITESFER<br>DHINALANWDPSAEPDTEIQFTPAR<br>ELYEADYADVFK<br>ELYEADYADVFKGDDAWR<br>ESGETVEFDAK<br>FVEFYGNGVK<br>GNHEVMMR<br>GTFANIR<br>GTNLLGVK<br>IDTPGEADYYR<br>ILLSEAK<br>LQNQLVDVAGGYTR<br>NVEIEYER<br>QDYNSLGSR<br>RGNHEVMMR<br>TESLNSFGAK<br>TICAPGSQVVDGYFQR<br>VLAENQLR<br>VLMQDFTGVPCVVDLATMR<br>VRIDTPGEADYYR<br>VVPPGTGIVHQVNIEYLSR<br>WGAENFSNFR | 294-318<br>835-843<br>58-82<br>646-657<br>646-663<br>900-910<br>277-286<br>753-760<br>761-767<br>827-834<br>913-923<br>378-384<br>768-781<br>142-149<br>743-751<br>752-760<br>2-11<br>508-523<br>41-48<br>83-101<br>911-923<br>169-187<br>159-168 | 102079 |
| 268 | MULTISPECIES: Aminopeptidase P family protein | Corynebacterium | WP_283117106.1 | IEDTLIITSGAPK | 339-351 | 39134 |
| 269 | MULTISPECIES: Alpha-1,4-glucan--maltose-1-phosphate maltosyltransferase | Corynebacterium | WP_070595729.1 | IDAWSDPMATWR | 92-103 | 75929 |
| 270 | Alpha-ketoglutarate decarboxylase | C. simulans, C. minutissimum | WP_282439962.1<br>KKO77948.1 | NMDESLEIPTATTVR<br>VIQGAESGEFLR | 8-22<br>209-220 | 123923 |
| 271 | Dihydrolipoyl dehydrogenase | C. kroppenstedtii<br>C. parakroppenstedtii<br>C. pseudokroppenstedtii<br>C. marinum | MDU7502749.1<br>WP_221926315.1<br>WP_269091290.1<br>WP_042620632.1 | GAIAIDDFMR | 292-301 | 49994 |
| 272 | Aminotransferase class V-fold PLP-dependent enzyme | C. resistens | WP_273352814.1 | AFDAVDR | 154-160 | 45366 |
| 273 | MULTISPECIES: Glyceraldehyde-3-phosphate dehydrogenase | Corynebacterium<br>(C. striatum, C. simulans) | WP_167599039.1 | AAGLNMVLTETGAAK<br>EAAYGGVR<br>ELSTAETLPILK<br>LTGNAIR<br>NVSLHSDLR<br>VLLTAPGKGDLK | 336-350<br>149-156<br>66-77<br>363-369<br>399-407<br>254-265 | 53277 |
| 274 | RNA | C. striatum | WP_049159484.1 | NAAGMDVDAVLLGSGAADPLYR | 132-153 | 29261 |

| | | | | | | |
|---|---|---|---|---|---|---|
| | methyltransferase | C. simulans<br>C. endometrii | WP_239240721.1<br>WP_168707160.1 | | | |
| 275 | Long-chain fatty acid--CoA ligase | C. variabile | WP_303944325.1 | GLCPVASQMVVIGNNRK | 506-522 | 68041 |
| 276 | Carboxylesterase/lipase family protein | Tsukamurella paurometabola | WP_013125362.1 | ATLVIDDR | 478-485 | 54645 |
| 277 | MULTISPECIES: Hypoxanthine phosphoribosyltransferase | Corynebacterium | WP_005529848.1 | GAVFFMTDFAR<br>SLEVVTLLR | 58-68<br>136-144 | 21962 |
| 278 | MULTISPECIES: Alpha/beta hydrolase-fold protein | Corynebacterium | WP_049146799.1 | FPEVWALDGLR<br>GPANAAGVGLEVISR | 134-144<br>321-335 | 71158 |
| 279 | MULTISPECIES: UTP--glucose-1-phosphate uridylyltransferase | Corynebacterium | WP_049154210.1 | ELLPVVDTPGIELIAEEAASVGAQR | 39-63 | 33868 |
| 280 | Thymidylate synthase | C. simulans<br>C. striatum | WP_201057408.1<br>MDU3174384.1 | WLQDNNVR | 73-80 | 30160 |
| 281 | Redox-regulated ATPase YchF | C. simulans<br>C. striatum | WP_284841372.1<br>HCG2912059.1 | VDPAHDISVINTELILADLQTIEK | 119-142 | 38962 |
| 282 | MULTISPECIES: 6-Phosphogluconolactonase | Corynebacterium | WP_062035109.1 | NVPVSDPDSNEGQAR | 91-105 | 26973 |
| 283 | Glycosyltransferase family 2 protein | C. sanguinis<br>C. auris<br>C. breve<br>C. marinum | WP_144772752.1<br>WP_290342415.1<br>WP_284825294.1<br>WP_042620624.1 | VADEMNIR | 171-178 | 25485 |
| 284 | MULTISPECIES: Anaerobic C4-dicarboxylate transporter | C. accolens<br>C. flavescens<br>C. tuberculostearicum<br>C. simulans<br>C. minutissimum<br>C. singulare<br>C. aurimucosum<br>C. guaraldiae<br>C. fournieri | WP_284899507.1<br>WP_284899507.1<br>WP_316991255.1<br>WP_062040809.1<br>KKO77707.1<br>WP_144792670.1<br>WP_144013193.1<br>WP_154761963.1<br>WP_179154881.1 | ELEDDPVYQER | 194-204 | 45338 |
| 285 | Acetate kinase | C. simulans<br>C. striatum | WP_248093053.1<br>WP_046646097.1 | AYVLVLNSGSSSVK<br>GISGVNDFR<br>LLDKDPK<br>QAGMSIDEIDNLLNK<br>SAEIADMAR<br>SGDIDPGIIFHLSR<br>TGGEEYKEELEIPTHSFGLER<br>VFVVPTNEELAIAQK | 2-15<br>277-285<br>195-201<br>257-271<br>393-401<br>243-256<br>49-69<br>378-392 | 44485 |
| 286 | Polyprenyl synthetase family protein | C. striatum<br>C. simulans<br>C. accolens | WP_284789615.1<br>WP_248093681.1<br>PCC83943.1 | VVEEQLSVLPECAASEALR | 305-323 | 36136 |

| | | | | | | |
|---|---|---|---|---|---|---|
| 287 | Protoporphyrinogen oxidase | C. accolens | WP_302525253.1 | AEEDDLVDYALDDLQHLTGFDGR | 369-391 | 48219 |
| 288 | Phosphoglycerate kinase | C. striatum | WP_049145845.1 | ANGLNDGDVMLIENVR<br>AQASVYDVAK<br>FGDKIVLPVDLIAATEFNADAENK<br>GYNVQNSLLQEDQIDNCK<br>ITASLPTIK<br>IVLPVDLIAATEFNADAENK<br>TLDDLIAEGVESR<br>TVFWNGPMGVFEMEAFSK<br>VVALDGIPEGWMSLDIGPESVK<br>YSLAPVAEALSEALGQYVALAGDVTGEDAHER | 107-122<br>163-172<br>264-287<br>241-258<br>41-49<br>268-287<br>5-17<br>320-337<br>288-309<br>75-106 | 42468 |
| 289 | MULTISPECIES: Branched-chain amino acid aminotransferase | Corynebacterium | WP_061923579.1 | LAMPELPQEEFIESLR | 107-122 | 40094 |
| 290 | Alpha-D-GlcNAc alpha-1,2-L-rhamnosyltransferase | C. xerosis<br>C. pelargi<br>C. phoceense | SLN04018.1<br>WP_128889604.1<br>WP_257068289.1 | IAMVGTR | 17-23 | 45133 |
| 291 | MULTISPECIES: Argininosuccinate synthase | Tsukamurella<br>Mycobacteriales bacterium (soil metagenome) | WP_146488150.1<br>NUS72237.1 | IGQLTMR | 403-409 | 51581 |
| 292 | Acetyl/propionyl/methylcrotonyl-CoA carboxylase subunit alpha | C. amycolatum<br>C. argoratense | WP_154862646.1<br>WP_234858539.1 | ITGLDLAR | 24-31 | 52573 |
| 293 | Aldehyde dehydrogenase family protein | C. kroppenstedtii<br>C. pseudokroppenstedtii<br>C. parakroppenstedtii<br>C. striatum<br>C. maris<br>C. accolens<br>C. simulans | WP_303735526.1<br>WP_204087849.1<br>WP_046205310.1<br>WP_005530427.1<br>WP_020933507.1<br>PCC83748.1<br>WP_062036254.1 | AVADGTATVR<br>FAAPNLLLGNTIVLK<br>GEIQIVADIYR<br>GVSLTGSEAAGAAVAK<br>HASICPLSSQACQDLLEEAGLPK<br>IKGVSLTGSEAAGAAVAK<br>INDSDRDAILDR<br>KLEWTLDQFVPIR<br>KSVLELGGNDPFIIIDDK<br>LAEHIGR<br>LDKAVADGTATVR<br>LIVLEDFYDR<br>MYNTGQACNAPK<br>STFALENPNTGVSEETFDR<br>SVLELGGNDPFIIIDDK<br>TLDYLVEK<br>VGPYDDENADIGPLSSIGAR<br>VGPYDDENADIGPLSSIGARDEIIER<br>WGVGEFVNEHLYR | 322-331<br>142-156<br>83-93<br>203-218<br>157-179<br>201-218<br>21-32<br>244-256<br>226-243<br>66-72<br>319-331<br>270-279<br>257-268<br>2-20<br>227-243<br>280-287<br>293-312<br>293-318<br>443-455 | 51256 |
| 294 | MULTISPECIES: Asparagine synthase (glutamine-hydrolyzing) | Corynebacterium | WP_049063082.1 | EVTAPIYAQSR | 459-469 | 72420 |

| | | | | | | |
|---|---|---|---|---|---|---|
| 295 | alanine racemase | C. simulans<br>C. accolens<br>C. striatum | WP_062035830.1<br>PCC84036.1<br>WP_166684452.1 | ADAYGHGVK | 33-41 | 39105 |
| 296 | MULTISPECIES: Sulfurtransferase | Corynebacterium | WP_070531080.1 | LVSAPWLSAR<br>VYDGSWAEWGNMVR | 19-28<br>268-281 | 32190 |
| 297 | Autonomous glycyl radical cofactor GrcA2 | C. striatum<br>C. aurimucosum<br>C. singular<br>C. amycolatum<br>C. simulans | WP_257212261.1<br>WP_010191241.1<br>WP_144791079.1<br>WP_284779076.1<br>WP_061924374.1 | ATPTFDER | 17-24<br>2-9 | 9352 |
| 298 | Polyprenol monophosphomannose synthase | C. tuberculostearicum<br>C. aurimucosum<br>C. kefirresidentii<br>C. parakroppenstedtii | WP_284821216.1<br>WP_049360879.1<br>WP_275434390.1<br>MCF8712652.1 | ELLESIDFDELSK | 170-182 | 32593 |
| 299 | MULTISPECIES: Thiamine pyrophosphate-binding protein | Corynebacterium | WP_061920027.1 | VSPMNVVEEFDR | 138-149 | 61262 |
| 300 | MULTISPECIES: Glycoside hydrolase family 32 protein | Corynebacterium | WP_286955443.1 | RIPSQNR | 117-123 | 44604 |
| 301 | Threonine/serine exporter family protein 5-formyltetrahydrofolate cyclo-ligase | C. glyciniphilum<br>C. jeikeium | WP_304030013.1<br>WP_080720366.1 | VDERPSR | 119-125<br>23-29 | 22368 |
| 302 | MULTISPECIES: Hydrogen peroxide-dependent heme synthase | Corynebacterium | WP_239273264.1 | DVAELIAVLP<br>GIYDLSGMR<br>ILMEHGMQAR<br>IVDLMYLMR<br>SYDWYTMDEEKR<br>VKDVAELIAVLP | 253-262<br>85-93<br>182-191<br>223-231<br>168-179<br>251-262 | 30694 |
| 303 | MULTISPECIES: NAD(P)H-dependent glycerol-3-phosphate dehydrogenase | Corynebacterium | WP_147760641.1 | GLGNNTLATIITR | 205-217 | 34440 |
| 304 | MULTISPECIES: Nitronate monooxygenase | Corynebacterium | WP_061921414.1 | GLETEFTR | 266-273 | 30598 |
| 305 | MULTISPECIES: Prolipoprotein diacylglyceryl transferase | Corynebacterium | WP_066487070.1 | FVVENMR | 236-242 | 30598 |
| 306 | Maltose alpha-D-glucosyltransferase | C. striatum<br>C. simulans | WP_239298545.1<br>WP_062035754.1 | RPIVGGTK | 419-426 | 57917 |
| 307 | Aldehyde dehydrogenase family protein | C. striatum<br>C. simulans<br>C. phoceense | WP_086890509.1<br>WP_284841483.1<br>WP_257069120.1 | IVESSAANLQR | 229-239 | 49646 |

| | | | | | | |
|---|---|---|---|---|---|---|
| | | C. accolens | PCC83619.1 | | | |
| 308 | Glucose 1-dehydrogenase | C. callunae<br>C. otitides<br>C. marinum<br>C. variabile<br>C. halotolerans<br>C. sanguinis<br>C. guangdongense | WP_247776312.1<br>WP_004601643.1<br>NLF90349.1<br>WP_313096135.1<br>WP_015400384.1<br>WP_136652566.1<br>WP_290196273.1 | HGVVGLTK | 167-174 | 26485 |
| 309 | NAD(P)/FAD-dependent oxidoreductase | C. striatum | WP_046645997.1 | TLRDQYSNYGTTSAK | 194-208 | 48838 |
| 310 | FAD-dependent monooxygenase | C. halotolerans | WP_015401872.1 | VAATLGLELK | 526-531 | 68304 |
| 311 | Argininosuccinate synthase | C. kroppenstedtii<br>C. parakroppenstedtii<br>C. urogenitale | WP_012731483.1<br>WP_221913640.1<br>WP_151902636.1 | AVETGFLEDLWNPPTK<br>SLDAFIDSTQENVTGDIR | 184-199<br>322-339 | 43829 |
| 312 | Class 1b ribonucleoside-diphosphate reductase subunit beta | C. striatum<br>C. simulans | WP_086890861.1<br>AMO89754.1 | GLESSSPQR | 193-201 | 32498 |
| 313 | MULTISPECIES: Leucyl aminopeptidase | Corynebacterium | WP_225723588.1 | LVLADAIAR | 367-375 | 51420 |
| 314 | Glycogen/starch/alpha-glucan phosphorylase | C. striatum<br>C. simulans<br>C. accolens | WP_201816736.1<br>WP_284841150.1<br>PCC82120.1 | ALLNNLTNLDLVDEAK | 74-89 | 90367 |
| 315 | Class I SAM-dependent methyltransferase | C. simulans<br>C. striatum<br>C. accolens | WP_239238573.1<br>WP_244080592.1<br>PCC81981.1 | DVADPALYVSTWLKDESVDVR | 315-335 | 59261 |
| 316 | MULTISPECIES: Succinate dehydrogenase cytochrome b subunit | Corynebacterium | WP_070546714.1 | ANMIATFSR<br>LAASDLGITGAK<br>NPDREALAHGR<br>SVKNPDREALAHGR<br>TNLMGGLDSFATK | 175-183<br>207-218<br>5-15<br>2-15<br>124-136 | 27304 |
| 317 | MULTISPECIES: Ribonucleoside triphosphate reductase | Corynebacterium | WP_049152229.1 | TGHLYNLEASPAEGATYR | 426-443 | 70419 |
| 318 | Galactokinase | C. striatum<br>C. simulans | WP_114976007.1<br>WP_061920796.1 | RADVESTAEAIAAAAAER | 374-391 | 42594 |
| 319 | MULTISPECIES: Phosphoglucosamine mutase | Corynebacterium | WP_023020606.1 | VSGEMLDAAIASGLASR | 55-71 | 47158 |
| 320 | Nicotinate-nucleotide--dimethylbenzimidazole phosphoribosyltransferase | C. amycolatum<br>C. vitaeruminis<br>C. marinum | WP_284837484.1<br>WP_048759029.1<br>NLF90219.1 | LKVAAVR | 207-213 | 51381 |

| | | | | | | |
|---|---|---|---|---|---|---|
| 321 | Aldo/keto reductase | C. striatum<br>C. accolens<br>C. simulans | WP_284789396.1<br>PCC82288.1<br>WP_248091192.1 | AGDVTRDELFITSK<br>HFDTATLYENEAELGQALNDAMR<br>TGIVPVLNQVELHPGFTQPELR | 66-79<br>43-65<br>157-178 | 31336 |
| 322 | Long-chain-fatty-acid--CoA ligase | C. striatum | WP_284773384.1 | MQFPEISK | 205-212 | 65893 |
| 323 | Alanine dehydrogenase | C. pseudodiphtheriticum | WP_284585175.1 | AVAAGGVDQAIER | 322-334 | 37968 |
| 324 | Acetoacetate--CoA ligase | C. halotolerans | WP_027004178.1 | DNPAIMQVEEDGRR | 87-100 | 71816 |
| 325 | Alpha/beta hydrolase family protein | C. aurimucosum<br>C. striatum<br>C. accolans<br>C. simulans | WP_193634643.1<br>WP_086890753.1<br>PCC82540.1<br>WP_248093137.1 | AAASYSGCPVR | 203-213 | 36506 |
| 326 | Nitrite/sulfite reductase | C. lipophiloflavum<br>C. sanguinis | WP_006840486.1<br>WP_144779569.1 | EQDVDKLR | 395-402 | 62873 |
| 327 | Phosphatidylinositol mannoside acyltransferase | C. pseudodiphtheriticum | WP_272696498.1 | LPLWLTRR | 32-39 | 46435 |
| 328 | Phosphopyruvate hydratase | C. aurimucosum | WP_010187392.1 | LTETLGDKVQIVGDDFFVTNPAR | 296-318 | 45297 |
| 329 | Sucrose-6-phosphate hydrolase | C. terpenotabidum | AGP31162.1 | VLATVRHSGDR | 372-382 | 54368 |
| 330 | Salicylate synthase | C. durum | WP_315159939.1 | TVYQHAGK | 395-402 | 47268 |
| 331 | Inorganic diphosphatase | C. aurimucosum<br>C. pseudodiphtheriticum | WP_253287150.1<br>WP_284584325.1 | LLAVVDDPR | 92-100 | 17865 |
| 332 | Cysteine desulfurase family protein | C. jeikeium<br>C. macclintockiae | WP_051890851.1<br>WP_284803947.1 | AVLPVDAAGR | 149-158 | 43149 |
| 333 | NAD(P)H-quinone oxidoreductase | C. matruchotii | WP_005524062.1 | IVTDTIK | 265-272 | 33643 |
| 334 | Fructose-specific PTS transporter subunit EIIC | C. vitaeruminis | WP_276651473.1 | LDADLGADK | 13-21 | 70249 |
| 335 | GNAT family N-acetyltransferase | C. pyruviciproducens | WP_016458261.1 | LVGCARVYK | 60-68 | 16521 |
| 336 | Type I glyceraldehyde-3-phosphate dehydrogenase | C. striatum<br>C. simulans<br>C. accolens<br>C. flavescens | WP_005531614.1<br>WP_284841327.1<br>PCC81819.1<br>WP_075729858.1 | AAAVNMVPTSTGAAK<br>AHIEAGAK<br>AVALVLPELEGK<br>GLMTTVHSYTGDQR<br>IAVSAERDPK<br>TTSLVASKL<br>VLNDKFGIEK<br>YDSVLGR | 218-232<br>126-133<br>233-244<br>191-204<br>90-99<br>344-352<br>181-190<br>65-71 | 37791 |
| 337 | UDP-N-acetylmuramoyl-L-alanyl-D-glutamate--2,6-diaminopimelate ligase | C. efficiens<br>C. gallinarum | WP_006768058.1<br>WP_191732191.1 | MRQHGVTHVVMEVSSHALSLGRVSGSHFNVAAFTNLSQDHLDFHDTMEEYFDAK | 175-228 | 53864 |
| 338 | 2-Oxoglutarate | C. argentoratense | WP_314929111.1 | AEEPKAEAK | 95-103 | 69519 |

| | | | | | | |
|---|---|---|---|---|---|---|
| | dehydrogenase, E2 component, dihydrolipoamide succinyltransferase | | | | | |
| 339 | Methionine adenosyltransferase | C. glucuronolyticum | WP_201841608.1 | EIGDGVDTSHEARSGAVEDDLDQSGAGDQGLMFGYASNETPELMPLPIALAHR | 101-152 | 43362 |
| 341 | FMN-binding glutamate synthase family protein | C. falsenii | WP_025403717.1 | LAPHMLR | 453-459 | 57267 |
| 342 | Catechol 1,2-dioxygenase | C. glyciniphilum | WP_304031649.1 | ANAIYKDLLGAIGDVAR | 35-51 | 32023 |
| 343 | Type I polyketide synthase | C. stationis<br>C. casei | WP_278829959.1<br>WP_301438337.1 | FHDEVLANVGVRKYHDDYGPNMPMVDNLAPELTTIYLERDMTFAVQDEETAR | 2449-2500 | 317142 |
| 344 | NAD-dependent succinate-semialdehyde dehydrogenase | Tsukamurella paurometabola | WP_245537894.1 | DAARVQRVGAGLRSGMVGVNR | 422-442 | 49706 |
| 345 | Histidine phosphatase family protein | C. jeikeium<br>C. macclintockiae | WP_071057504.1<br>WP_034980474.1 | MWEAVDAAREK | 128-138 | 18465 |
| 346 | Histidine phosphatase family protein | C. jeikeium<br>C. evansiae<br>C. macklintockiae | WP_071057504.1<br>WP_269943938.1<br>WP_034980474.1 | GVQVELPR | 128-138 | 33204 |
| 347 | Phosphopyruvate hydratase | C. accolens<br>C. aurimucosum<br>C. pseudotuberculosis<br>C. ulcerans<br>C. halotolerans<br>C. minutissimum<br>C. silvaticum<br>C. deserti | WP_284897965.1<br>WP_010187392.1<br>WP_058832493.1<br>WP_046693914.1<br>WP_015400396.1<br>WP_115021570.1<br>WP_087453687.1<br>WP_053544480.1 | LQEGIDKK | 319-326 | 45118 |
| 348 | Phospho-sugar mutase | C. diphtheriae | WP_235697053.1 | VYLGGRIAQGAAEGVQLISPADK | 161-183 | 58076 |
| 349 | IMP dehydrogenase | C. jeikeium | WP_041626416.1 | SEAGMITDPVTASPDMTIQEVDDLCAR | 103-129 | 54043 |
| 350 | Succinyldiaminopimelate Transaminase | C. ammoniagenes<br>C. stationis | WP_168938853.1<br>WP_075723076.1 | EAISSALER | 72-80 | 39465 |
| 351 | Carbon-nitrogen hydrolase family protein | C. striatum | WP_005529738.1 | LVAAVQIQTGGDIEENLELAVEKIRVAAEGGAQLIVLPEATSQAFGAGRLDK | 2-53 | 28302 |
| 352 | Glycosyl hydrolase family 65 protein | C. kroppenstedtii<br>C. parakroppenstedtii | WP_303734931.1<br>WP_221926305.1 | ELSHDGALFPWRTINGLESSAYYAAGTAQYHIDADIAYALMKYVYASGDTDFLLR | 476-530 | 96833 |
| 353 | Non-ribosomal peptide synthase/polyketide synthase | Tsukamurella paurometabola<br>Nocardia wallacei | WP_013127570.1<br>WP_280333217.1 | AGELVLR | 338-344 | 42249 |
| 354 | Biotin/lipoyl-containing protein | C. tuberculostearicum | WP_301714491.1 | IYAPFAGIVRCHVNVGDTVDTGVPLATVEATKLEAPVESPGPGKVHR | 3-49 | 21170 |
| 355 | Aminomethyl-transferring glycine dehydrogenase | Clavibacter michiganensis | WP_204575344.1 | FIAAMIGIK | 904-912 | 105112 |
| 356 | FMN-binding | C. falsenii | WP_025403717.1 | LAPHMLR | 469-475 | 57267 |

| | | | | | | |
|---|---|---|---|---|---|---|
| | glutamate synthase family protein | | | | | |
| 357 | Malate synthase G | C. efficiens<br>C. glutamicum<br>C. callunae<br>C. deserti | WP_006768235.1<br>WP_038585104.1<br>WP_247774532.1<br>WP_053545424.1 | HTLKVGVMDEERR | 430-442 | 82338 |
| 358 | NTD biosynthesis operon regulator NtdR | C. glutamicum | BAV24098.1 | APLMAKPCELPTR | 127-139 | 13979 |
| 359 | Amidohydrolase | C. ammoniagenes | WP_236163494.1 | EFGDSMAEKIGVER | 414-427 | 58668 |
| 360 | SDR family oxidoreductase | Prescottella equi | WP_013415346.1 | FMSVNMDGALNVCRAVVPHMR | 113-133 | 26613 |
| 361 | Oxoglutarate dehydrogenase inhibitor Odhl | C. pseudotuberculosis | WP_013241856.1 | EPKNSEVLSSGDEIQIGK | 115-132 | 15397 |
| 362 | Alpha/beta hydrolase | C. kefirresidentii | WP_239207579.1 | SRRSGQSWHYVSDLAFYFADLTAALDAIPNDEVIFIAHSTGGLIAPLWMDHLRR | 99-152 | 38146 |
| 363 | UDP-N-acetylmuramoyl-L-alanine--D-glutamate ligase | Clavibacter michiganensis | WP_094113683.1 | TTTTQLTAALLQEGGVR | 149-165 | 54953 |
| 364 | LysR substrate-binding domain-containing protein | Tsukamurella paurometabola | WP_041944519.1 | GVAPTAAATELAR | 63-75 | 31518 |
| 365 | Diaminopimelate epimerase | C. callunae | WP_015651503.1 | KTVIPFAK | 2-9 | 29190 |
| 366 | Phosphoribosyltransferase, ComF family protein | C. glutamicum<br>C. efficiens<br>C. callunae | OKX89421.1<br>WP_006769546.1<br>WP_282101775.1 | GEVEHDITLVPAPTR | 86-100 | 21162 |
| 367 | D-xylulose 5-phosphate/D-fructose 6-phosphate phosphoketolase | C. efficiens | EEW48411.1 | GPPCPPLPIPPADR | 549-562 | 67838 |
| 368 | Polyketide synthase | C. variabile | WP_312775706.1 | LIQEIGSNLEAVSARR | 691-706 | 329271 |
| 369 | AMP-binding protein, acyl-CoA synthetase | C. kroppenstedtii<br>C. pseudokroppenstedtii | WP_012732575.1<br>WP_284866868.1 | MPKYTFETR | 1-9 | 62623 |
| 370 | Long-chain-fatty-acid--CoA ligase | Clavibacter michiganensis | WP_094116470.1 | VVDRTKDMILR | 424-434 | 56763 |
| 371 | Aminotransferase class I/II-fold pyridoxal phosphate-dependent enzyme | C. glucuronolyticum | WP_232621892.1 | EIAAVDGISIVTPACNAIVTFTTGSEK | 377-403 | 49538 |
| 372 | Dihydroxyacetone kinase subunit DhaK | C. glucuronolyticum | WP_005393004.1 | SLDDVAAIAKK | 181-191 | 32023 |
| 373 | Beta-galactosidase | C. genitalium | WP_005287622.1 | LLQATYPK | 217-224 | 56587 |
| 374 | NAD(P)-dependent oxidoreductase | Prescottella equi | WP_084987694.1 | ILVAGAGGVVGLPLTR | 3-18 | 29331 |
| 375 | MULTISPECIES: 5-(Carboxyamino)imidazole ribonucleotide | Corynebacterium | WP_257994307.1 | MLSYAK<br>NAGLLAVR | 46-51<br>121-128 | 17221 |

| | | | | | | |
|---|---|---|---|---|---|---|
| | mutase | | | | | |
| 376 | Low molecular weight phosphatase family protein | C. stationis<br>C. diphtheriae<br>C. variabile | WP_066793503.1<br>WP_003849982.1<br>WP_312801943.1 | LNPQSVEVIAEAGADMSAGHPK<br>LVRDDIDTR<br>RLVAELLEN<br>SQMAAALAAK<br>VIILGGDAQLELPDDAHGTCER<br>WVTDEPSRR | 46-67<br>120-128<br>131-139<br>18-27<br>80-101<br>102-110 | 14955 |
| 377 | Oxoglutarate dehydrogenase inhibitor Odhl | C. striatum | WP_005531457.1 | HPEADIFLDDVTVSR<br>NSQVLHVGDEIQIGK<br>RGPNAGAR | 75-89<br>120-134<br>55-62 | 15535 |
| 378 | MULTISPECIES: Alpha/beta hydrolase-fold protein | Corynebacterium | WP_151843393.1 | WETFLTK<br>GPQKWETFLTK<br>NTGTHSWPGWR | 230-236<br>203-213<br>372-382 | 40881 |
| 379 | Antibiotic biosynthesis monooxygenase | C. striatum<br>C. simulans | CQD14203.1<br>WP_062035561.1 | FLLVEAYADGK<br>MAEYKVES<br>SCEEFPK<br>MILINVK | 53-63<br>104-111<br>76-82<br>1-7 | 13251 |
| 380 | Inorganic diphosphatase | C. striatum<br>C. simulans<br>C. phoceense | WP_275432308.1<br>WP_282440131.1<br>WP_230384001.1 | DLEPNKEVTGSGWGDK<br>DLEPNKEVTGSGWGDKAEAEK<br>ILEEAIAR<br>LLCVIDDPR<br>MTDEAGGDDKLLCVIDDPR<br>MTDEAGGDDKLLCVIDDPRWER<br>NKYEVDHETGK | 132-147<br>132-152<br>153-160<br>97-105<br>87-105<br>87-108<br>20-30 | 18386 |
| 381 | MULTISPECIES: Ribose-5-phosphate isomerase | Corynebacterium | WP_105324929.1 | CALAWSPETAR<br>EHNNAQLIGIGGR<br>IDILAEYER<br>RIDILAEYER<br>TVNDPGSLGIVLGGSGNGEQIAANK<br>TVNDPGSLGIVLGGSGNGEQIAANKVK | 84-94<br>98-110<br>139-147<br>138-147<br>54-78<br>54-80 | 17451 |
| 382 | MULTISPECIES: NAD(P)H-quinone dehydrogenase | Corynebacterium | WP_236592667.1 | TITMPLNTNPR | 392-402 | 50384 |
| 383 | Pantetheine-phosphate adenylyltransferase | C. minutissimum<br>C. sanguinis<br>C. deserti<br>C. aurimucosum<br>C. glutamicum<br>C. striatum<br>C. efficiens | WP_115021954.1<br>WP_144690106.1<br>WP_053544756.1<br>PKZ24642.1<br>WP_038583872.1<br>WP_110301875.1<br>WP_006769403.1 | SSLDYEYELPMAQMNR | 91-1006 | 17427 |
| 384 | MULTISPECIES: Class II fructose-bisphosphate aldolase | C. striatum<br>Corynebacterium | WP_204083879.1<br>WP_284774425.1 | IDGEVGNKK<br>IEQALSYGVVK<br>LRPEVLDEGQK<br>MNVDTDTQYAFTNPVAR<br>PIATPEVYNQMLDTAK<br>VVEACTDLHSVGK | 301-309<br>262-272<br>222-232<br>273-289<br>2-17<br>328-340 | 37245 |
| 385 | Thioredoxin-dependent thiol | C. amycolatum<br>C. jeikeium | WP_005511127.1<br>WP_071056107.1 | VVQGVIR | 118-124 | 18208 |

| | | | | | | |
|---|---|---|---|---|---|---|
| | peroxidase | C. striatum<br>C. pseudotuberculosis<br>C. ulcerans<br>C. simulans | GKH17804.1<br>WP_278072254.1<br>WP_102241123.1<br>WP_062044478.1 | | | |
| 386 | Arginine repressor | C. simulans<br>C. striatum<br>C. accolens<br>C. tuberculostearicum<br>C. flavescens | WP_284841612.1<br>WP_256999753.1<br>PCC82781.1<br>WP_204609194.1<br>WP_075729752.1 | ILEILDR | 24-30 | 17443 |
| 387 | S-Ribosylhomocysteine lyase | C. striatum<br>C. phoceense<br>C. confusum | WP_114975870.1<br>WP_303936878.1<br>WP_290224437.1 | SNVESFELDHR | 4-14 | 17450 |
| 388 | Hypothetical protein | C. durum | WP_315186013.1 | AGDDRDPR | 256-263 | 34800 |
| 389 | Sodium:alanine symporter family protein | C. pseudodiphtheriticum | WP_249617946.1 | HGFIDR | 56-61 | 50843 |
| 390 | YggS family pyridoxal phosphate-dependent enzyme | C. halotolerans<br>C. glutamicum | WP_015401308.1<br>WP_003856518.1 | GVALALER | 118-125 | 25678 |
| 391 | MULTISPECIES: Serine O-acetyltransferase EpsC | Corynebacterium | WP_286953645.1 | IKLVDPDYYI | 178.187 | 20503 |
| 392 | Molybdopterin-binding protein | C. striatum<br>C. simulans | WP_166684653.1<br>WP_062035328.1 | GCGSLIINSASSR | 124-136 | 13644 |
| 393 | 3-Deoxy-7-phosphoheptulonate synthase class II | Clavibacter michiganensis | WP_104289862.1 | YEQLARDIDR | 208-217 | 49914 |
| 394 | Polyadenylate-specific 3'-exoribonuclease AS | C. striatum<br>C. simulans | WP_244080664.1<br>WP_282440013.1 | QYWEFAGCPK | 121-130 | 19279 |
| 395 | Uracil phosphoribosyltransferase | C. striatum<br>C. simulans<br>C. aurimucosum<br>C. accolens | WP_114975994.1<br>WP_248092469.1<br>WP_144013300.1<br>EEI14487.1 | DLGTMLVYEAAR | 34-45 | 22474 |
| 396 | AAA family ATPase | C. efficiens | WP_011076137.1 | ISNEGLWK | 122-129 | 48319 |
| 397 | Low molecular weight phosphatase family protein | C. diphtheria<br>C. variabile<br>C. stationis<br>C. ammoniagenes | WP_003849982.1<br>WP_312801943.1<br>WP_305938828.1<br>WP_003849011.1 | GVDPQLLR<br>LNPQSVEVIAEAGADMSAGHPK<br>SQMAAALAAK | 68-75<br>46-67<br>18-27 | 14870 |
| 398 | Phosphoenolpyruvate carboxykinase (GTP) | C. striatum<br>C. accolens<br>C. simulans | WP_244080864.1<br>PCC82436.1<br>WP_248093862.1 | AEDIDIEGLDFDIEDVR<br>GLVGELPTSNEK<br>VVFVDGSQEEADR | 543-559<br>7-18<br>35-47 | 67023 |
| 399 | MULTISPECIES: Ribose-5-phosphate isomerase | Corynebacterium | WP_115023267.1 | EHNNAQLIGIGGR<br>TVNDPGSLGIVLGGSGNGEQIAANK<br>TVNDPGSLGIVLGGSGNGEQIAANKVK<br>VVYLGADHAGFEMK | 98-110<br>54-78<br>54-80<br>3-15 | 16995 |
| 400 | MULTISPECIES: MBL fold metallo-hydrolase | Corynebacterium | WP_005531203.1 | IDEYVAR<br>KYSDFDVIIESLR<br>VLPGHGDETTVGAEAAR | 198-204<br>157-169<br>181-197 | 22248 |

| | | | | | | |
|---|---|---|---|---|---|---|
| | | | | YSDFDVIIESLR | 158-169 | |
| 401 | Peptidylprolyl isomerase | C. halotolerans | WP_015399458.1 | (K)GPFYDGAIFHR | 57-67 | 18821 |
| 402 | SAM-dependent DNA methyltransferase | C. glyciniphilum<br>C. nuruki | WP_038550587.1<br>WP_010121668.1 | AETRQGR | 85-91 | 104554 |
| 403 | Acyl-CoA dehydrogenase family protein | Clavibacter michiganensis | WP_080939243.1 | LSASGSVR | 213-220 | 43716 |
| 404 | MULTISPECIES: Adenylate kinase | Corynebacterium | WP_066840251.1 | ANIGEGTPLGVEAK<br>QYIDAGK | 37-50 | 19633 |
| 405 | Aspartate kinase | C. variabile<br>C. terpenotabidum<br>C. nuruki | WP_141328500.1<br>WP_020442109.1<br>WP_010121071.1 | VSLVGAGMK<br>VTVLGIPDKPGEAAR | 267-281 | 45294 |
| 406 | Peptide deformylase | C. striatum<br>C. simulans<br>C. accolens | WP_275432358.1<br>WP_248090397.1<br>PCC81831.1 | MYGDPVLTSR | 1-10 | 17898 |
| 407 | CDP-alcohol phosphatidyltransferase family protein | C. striatum<br>C. phoceense | WP_100087000.1<br>WP_257052253.1 | KPAAVVVEPVAK | 8-19 | 22042 |
| 408 | Nicotinamide-nucleotide amidohydrolase family protein | C. accolens<br>C. tuberculostearicum<br>C. kefirresidentii<br>C. simulans | WP_284609154.1<br>WP_284624038.1<br>WP_239207602.1<br>WP_062044118.1 | GGLITYATEVK | 46-56 | 17890 |
| 409 | Aromatic acid exporter family protein | C. maris | WP_020933460.1 | MSSPAPMQALAR | 1-12 | 41713 |
| 410 | Phosphopyruvate hydratase | C. callunae<br>C. aurimucosum<br>C. minutissimum<br>C. singular<br>C. phocae | WP_015650797.1<br>WP_010190769.1<br>WP_115023123.1<br>WP_042531853.1<br>KAA8728563.1 | RGAFPR | 417-422 | 30481 |
| 411 | Aminomethyl-transferring glycine dehydrogenase | C. simulans<br>C. striatum<br>C. accolens | WP_239240777.1<br>WP_110333095.1<br>PCC82253.1 | EYASQNIVLK | 62-71 | 102796 |
| 412 | Phosphoribosyltransferase | C. striatum<br>C. simulans<br>C. accolens<br>C. aurimucosum<br>C. phocea<br>C. tuberculostearicum<br>C. diphtheriae | WP_086890390.1<br>MCK6161043.1<br>WP_237801691.1<br>WP_158397262.1<br>WP_075732225.1<br>WP_316986334.1<br>WP_196975715.1 | KVLVVDDVADSGK<br>VLVVDDVADSGK | 97-109<br>98-109 | 18714 |
| 413 | 3'(2'),5'-Bisphosphate nucleotidase CysQ | C. striatum<br>C. simulans<br>C. accolens | WP_236593763.1<br>WP_062039079.1<br>PCC83726.1 | GLALGDAGDEAAQEWITR | 45-62 | 28305 |
| 414 | IMP dehydrogenase | C. simulans | WP_046646763.1 | ASMGYTGSATLEELK | 462-476 | 53421 |
| 415 | Pyrroline-5-carboxylate reductase | C. simulans<br>C. tuberculostearicum<br>C. striatum<br>C. marquesiae | WP_248093618.1<br>WP_239454637.1<br>WP_086892142.1<br>WP_269953714.1 | AGVTSPGGTTAAAVR | 223-237 | 26748 |

| | | | | | | |
|---|---|---|---|---|---|---|
| | | C. macginleyi<br>C. kefirresidentii<br>C. aurimucosum<br>C. accolens | WP_234457550.1<br>WP_239204751.1<br>WP_049358739.1<br>WP_284628503.1 | | | |
| 416 | GNAT family N-acetyltransferase | C. diphtheria<br>C. simulans<br>C. pseudotuberculosis<br>C. striatum<br>C. ulcerans<br>C. silvaticum<br>C. vitaeruminis | WP_106202522.1<br>WP_248090904.1<br>WP_014800250.1<br>MDU3174364.1<br>WP_046693906.1<br>WP_087453705.1<br>WP_034650028.1 | MTYGEMAGQWR | 155-165 | 18341 |
| 417 | Dihydroxyacetone kinase subunit DhaL | C. striatum<br>C. aurimucosum<br>C. singulare<br>C. simulans | WP_100619471.1<br>WP_193629369.1<br>WP_239178779.1<br>WP_284841233.1 | AAEGEKTMVDAWAPAAR<br>ASYLGER<br>EQADAGAGVADVLR | 111-127<br>167-173<br>131-144 | 20295 |
| 418 | dTMP kinase | C. glutamicum<br>C. deserti<br>C. callunae | WP_077311348.1<br>WP_053544323.1<br>WP_015650567.1 | MIVSIEGIDGAGK | 1-13 | 22378 |
| 419 | L-Threonylcarbamoyladenylate synthase | C. simulans<br>C. striatum | WP_248090633.1<br>WP_284822825.1 | MPLHPIAIELLR | 124-135 | 27208 |
| 420 | L-Threonylcarbamoyladenylate synthase | C. simulans<br>C. striatum | WP_248090633.1<br>WP_284822825.1 | MPLHPIAIELLR | 124-135 | 27208 |
| 421 | Serine hydroxymethyltransferase | C. glyciniphilum<br>C. terpenotabidum<br>C. variabile | WP_038547454.1<br>WP_020441580.1<br>WP_313007196.1 | NSELNGQEAEDLLHEVGITVNR | 336-357 | 46847 |
| 422 | Decaprenyl-phosphate phosphoribosyltransferase | C. striatum<br>C. simulans | WP_086890597.1<br>WP_284841631.1 | KSLEGYTPTYLR<br>YSEILLAER | 230-241<br>215-223 | 35734 |
| 423 | Deoxyribose-phosphate aldolase | C. striatum | WP_166683213.1 | AAEAAGADFVK<br>LMADTVGGR<br>TGADAQAMIDAGATR<br>TSTGFHPAGGASVHAVK | 132-142<br>160-168<br>179-193<br>114-131 | 21121 |
| 424 | tRNA (adenine-N1)-methyltransferase | C. simulans<br>C. striatum<br>C. aurimucosum | WP_062040796.1<br>WP_086891396.1<br>WP_102233936.1 | LGDLADVTVEELGGPVDR | 159-176 | 31392 |
| 425 | O-methyltransferase | C. simulans<br>C. striatum<br>C. pseudodiphtheriticum<br>C. durum | WP_284841396.1<br>WP_114976154.1<br>WP_249620733.1<br>WP_311142803.1 | LPLGSGLTLITK | 202-213 | 23667 |
| 426 | Type I polyketide synthase<br>(Type I polyketide synthase 13, Pks13) | C. simulans<br>C. striatum<br>C. accolens<br><br>M. tuberculosis | WP_284841086.1<br>WP_100088013.1<br>PCC83890.1<br><br>CKQ40838.1 | DVDSLVEWIGSEQR<br>VALGEIAADDER | 2131-2144<br>2001-2012 | 320693 |
| 427 | Demethylmenaquinone methyltransferase | C. simulans<br>C. argentoratense | WP_239239175.1<br>WP_314930498.1 | VLDLAAGTAVSTEELAK | 55-71 | 25216 |

| | | | | | | |
|---|---|---|---|---|---|---|
| | | C. striatum | MDU3174594.1 | | | |
| 428 | N-acetyl-gamma-glutamyl-phosphate reductase | C. pseudodiphthereticum<br>C. simulans<br>C. striatum<br>C. accolens | WP_284867551.1<br>WP_062037501.1<br>WP_086890829.1<br>PCC82768.1 | ITDIFR | 55-60 | 23949 |
| 429 | Phosphoribosylglycinamide formyltransferase | C. striatum<br>C. simulans<br>C. macginleyi<br>C. accolens | WP_086892453.1<br>WP_062035594.1<br>WP_121953170.1<br>WP_284612631.1 | DALEYGVK | 134-141 | 20345 |
| 430 | Glycerol kinase | C. kutscheri<br>C. casei<br>C. stationis<br>C. halotolerans<br>C. simulans<br>C. striatum | VEH05893.1<br>WP_301438465.1<br>WP_066796784.1<br>WP_015402044.1<br>WP_062036799.1<br>WP_110301885.1 | AVLEATAYQTR<br>FVAAIDQGTTSTR | 388-398<br>6-18 | 56837 |
| 431 | Phosphoenolpyruvate carboxykinase (GTP) | C. macclintockiae<br>C. jeikeium | WP_284804985.1<br>WP_005297204.1 | VPSEVWDEFQGLK | 587-599 | 67733 |
| 432 | 3-Oxoacyl-[acyl-carrier-protein] reductase FabG | C. jeikeium<br>C. macclintockiae<br>C. falsenii | WCZ52763.1<br>WP_284821088.1<br>WP_259342817.1 | AIVTGASSGIGR | 2-13 | 29637 |
| 433 | Orotidine-5'-phosphate decarboxylase | C. flavescens<br>C. striatum<br>C. simulans<br>C. evansiae | WP_301502224.1<br>WP_166683887.1<br>WP_248090406.1<br>WP_269944614.1 | EAGCLTLADAK | 90-100 | 28754 |
| 434 | Trehalose-phosphatase | C. striatum<br>C. simulans | WP_100087624.1<br>WP_062043329.1 | GSWISELR<br>LELSDPDLAAEAYAAGR<br>RLELSDPDLAAEAYAAGR<br>SVIEFSATQATK | 185-192<br>143-159<br>142-159<br>173-184 | 27820 |
| 435 | Riboflavin synthase | C. callunae | WP_247775370.1 | FSLPENLAR | 122-130 | 21828 |
| 436 | Dihydroxyacetone kinase phosphoryl donor subunit DhaM | C. simulans<br>C. striatum | WP_284841217.1<br>WP_239299454.1 | AIEVFVPK<br>LAEGLAELAGQMAADVR | 125-132<br>17-33 | 23067 |
| 437 | UMPkinase | C. tuberculostearicum<br>C. pseudogenitalium<br>C. striatum<br>C. simulans<br>C. curieae | WP_301989067.1<br>EFQ81596.1<br>WP_005528912.1<br>AMO90091.1<br>WP_269945892.1 | AVDGVYSDDPR<br>AVDGVYSDDPRTNPDAELYSEITPR<br>AVNGEQIGTLVKSQ<br>LGGEMFGGGK<br>QGTEVAVVIGGGNFFR<br>TNPDAELYSEITPR<br>VQTSINMAQIAEPYLPLR | 170-180<br>170-194<br>233-246<br>21-30<br>51-66<br>181-194<br>108-125 | 26657 |
| 438 | Type I methionyl aminopeptidase | C. variabile (cheese) | WP_312776128.1 | SPAELDAMQAAGEIVGK | 14-30 | 27858 |
| 439 | MULTISPECIES: Decaprenylphospho-beta-D-erythro-pentofuranosid-2-ulose 2-reductase | Corynebacterium | WP_095538020.1 | SNFVYGASK | 154-162 | 27240 |
| 450 | 3'(2'),5'-Bisphosphate nucleotidase CysQ | C. aurimucosum<br>C. minutissimum | WP_193629934.1<br>WP_181815344.1 | LTNSLADGCGEILK<br>TVEISDSR | 10-23<br>2-9 | 26979 |

| | | | | | | |
|---|---|---|---|---|---|---|
| | | C. faecipullorum<br>C. guaraldiae | HIX79222.1<br>WP_196189536.1 | | | |
| 451 | Sugar O-acetyltransferase | C. phoceense<br>C. striatum<br>C. simulans<br>C. accolens | WP_257054397.1<br>WP_284786693.1<br>WP_061919951.1<br>PCC82986.1 | EGGWEIAKPITVGR | 131-144 | 23565 |
| 452 | GTP cyclohydrolase I FolE | C. diphtheria<br>C. striatum<br>C. simulans<br>C. ulcerans | WP_014302352.1<br>WP_284772376.1<br>WP_062043259.1<br>WP_014836794.1 | ELLIAVGEDPDREGLQETPAR | 22-42 | 22305 |
| 453 | 5-Oxoprolinase subunit PxpA | C. urealyticum<br>C. tuberculostearicum | WP_148820140.1<br>WP_301979374.1 | GVVAELDAR<br>NYNPDGTLVSR | 35-47<br>172-182 | 26386 |
| 454 | Urocanate hydratase | C. striatum<br>C. accolens<br>C. durum<br>C. otitides<br>C. phoceense<br>C. aurimucosum<br>C. tuberculostearicum<br>C. singulare<br>C. simulans | WP_049146341.1<br>WP_284628471.1<br>WP_040360455.1<br>WP_004601612.1<br>WP_068802039.1<br>WP_201828382.1<br>WP_284820391.1<br>WP_144792023.1<br>WP_096336777.1 | MLMNNLDPEVAERPEDLVVYGGTGR | 28-52 | 61454 |
| 455 | Nucleoside triphosphate pyrophosphatase | C. aurimucosum<br>C. guaraldiae | WP_193634239.1<br>WP_144268683.1 | GAGVEPVIAPAHVDER | 18-33 | 21129 |
| 456 | Aspartate/tyrosine/aromatic aminotransferase | C. variabile | CUU67319.1 | AAALEREGTEILK | 32-44 | 46258 |
| 457 | Ribulose-phosphate 3-epimerase | C. tuberculostearicum<br>C. kefirresidentii<br>C. aurimucosum<br>C. marquesiae<br>C. simulans<br>C. striatum<br>C. parakroppenstedtii | WP_301714693.1<br>WP_259824573.1<br>WP_049360767.1<br>WP_269953001.1<br>WP_239238952.1<br>WP_086891453.1<br>MCF8713133.1 | SAPIISPSILAADFSR | 2-27 | 23879 |
| 458 | Type I methionyl aminopeptidase | C. simulans<br>C. striatum | WP_248092738.1<br>WP_279109652.1 | LTDVSHALEQATYR<br>TPGELDAMQAAGEIVGK | 152-165<br>14-30 | 27954 |
| 459 | Mannose-6-phosphate isomerase, class I | C. halotolerans | WP_015400169.1 | MDQLTGMLR | 1-9 | 41736 |
| 460 | Phosphoribosyltransferase | C. vitaeruminis<br>C. cystitides<br>C. confusum<br>C. pilosum<br>C. flavescens<br>C. accolens | WP_025251541.1<br>WP_257161592.1<br>WP_290224102.1<br>WP_026254242.1<br>APT85872.1<br>PCC82976.1 | ENLTWETFGEASR<br>VLVVDDVADSGK | 20-32<br>106-117 | 20812 |
| 461 | MULTISPECIES: dCTP deaminase | Corynebacterium | WP_241864884.1 | FTLPADLAGR<br>SYLNFR<br>VGQLAIFK | 88-97<br>182-187<br>146-153 | 20760 |
| 462 | RdgB/HAM1 family | C. diphtheriae | WP_106202317.1 | TFADNALIK | 48-56 | 21601 |

| | | | | | | |
|---|---|---|---|---|---|---|
| | non-canonical purine NTP pyrophosphatase | | | | | |
| 463 | Phosphate acetyltransferase | C. striatum<br>C. simulans<br>C. accolens<br>C. sanguinis<br>C. hindlerae | WP_100087587.1<br>WP_239240142.1<br>PCC82392.1<br>WP_259916092.1<br>WP_182385344.1 | GVTLEQAR<br>QLLDQAR<br>TSDLALAHAR<br>VAMLSYSTGTSGSGPDVDR<br>VDGPLQFDAACDPGVAAK | 219-226<br>133-139<br>86-95<br>325-343<br>360-377 | 46423 |
| 464 | 16S rRNA (guanine(966)-N(2))-methyltransferase RsmD | C. striatum<br>C. guaraldiae<br>C. aurimucosum<br>C. hesseae<br>C. faecipullorum | WP_086891473.1<br>WP_144269098.1<br>WP_158381122.1<br>WP_269948218.1<br>HIX78688.1 | EGLFSSLNVR | 30-39 | 21908 |
| 465 | 4-Hydroxy-3-methylbut-2-enyl diphosphate reductase | C. striatum<br>C. simulans | WP_086891673.1<br>WP_248090884.1 | CDLMIVVGSK<br>ERFPHLENPPSDDICYATQNR<br>EVLEFLDER<br>FPHLENPPSDDICYATQNR<br>KQLTLDATCPLVTK<br>NVLLAAPR<br>QLTLDATCPLVTK<br>YGAPIYVR | 217-226<br>186-206<br>281-289<br>188-206<br>93-106<br>7-14<br>94-106<br>34-41 | 35366 |
| 466 | Transaldolase | C. falsenii | WP_025402711.1 | GTPAHSAVFDMAADDVR<br>VSLEVDPR | 72-88<br>108-115 | 40193 |
| 467 | NAD(P)-dependent alcohol dehydrogenase | C. accolens<br>C. simulans<br>C. striatum<br>C. vitaeruminis | WP_284636044.1<br>WP_062041062.1<br>MDU3175964.1<br>WP_025252618.1 | AAGICHSDIHTIR<br>DPREDDVVIDIK<br>EDDVVIDIK<br>EGDKVAIVGLGGLGHMGVQIAAAK<br>GADVTVLSR<br>VAIVGLGGLGHMGVQIAAAK | 38-50<br>26-37<br>29-37<br>175-198<br>199-207<br>179-198 | 37326 |
| 468 | Acryloyl-CoA reductase | C. striatum<br>C. simulans | WP_239298930.1<br>WP_284841503.1 | ALYDGAVDTAGSTVLANVLAQIK<br>EGHVLVTGATGGVGSIAVQLLK<br>GIVLAGGNSVDAPLK<br>SLGFEVTAVTGR<br>VDSESEYLQALGADHIIDR | 242-264<br>177-198<br>291-305<br>199-210<br>211-229 | 37042 |
| 469 | tRNA (adenine-N1)-methyltransferase | C. striatum<br>C. simulans | WP_284765339.1<br>WP_062040796.1 | STLGSDFLLFR | 56-66 | 31468 |
| 470 | 2-Dehydropantoate 2-reductase | C. striatum<br>C. simulans<br>C. accolens | WP_129720736.1<br>WP_062037140.1<br>PCC82346.1 | DIKDRLPNELDAQVGAIR<br>DRLPNELDAQVGAIR<br>LPNELDAQVGAIR | 263-280<br>266-280<br>268-280 | 32366 |
| 471 | Biotin--[acetyl-CoA-carboxylase] ligase | C. striatum<br>C. simulans | WP_046646522.1<br>WP_061920773.1 | DTDRTELAIALLK<br>INVGGEYYSAGDVTHLRPVQ | 202-214<br>267-286 | 30576 |
| 472 | Porphobilinogen synthase | Corynebacterium | WP_070681443.1 | AGADQILTYFATDAAR | 307-322 | 37139 |
| 473 | 3-Dehydroquinate synthase | C. matruchotii<br>C. pseudodiphtheriticum<br>C. propinquum<br>C. agentoratense<br>C. callunae<br>C. deserti | WP_314268026.1<br>WP_249620746.1<br>WP_284571709.1<br>WP_234870828.1<br>WP_015651407.1<br>WP_053545022.1 | ETLNYGHTFAHAVELR<br>VIQVPTTLLAMVDAAVGGK | 256-271<br>140-158 | 41065 |

| | | | | | | |
|---|---|---|---|---|---|---|
| | | C. glutamicum<br>C. striatum<br>C. simulans | WP_087063328.1<br>WP_166683967.1<br>WP_239238695.1 | | | |
| 474 | Class II fructose-bisphosphate aldolase | C. striatum<br>C. sanguinis<br>C. lipophiloflavum | WP_166683739.1<br>WP_259818282.1<br>EEI18001.1 | EVLDEYVRPLIAISQER<br>LGLEEGSKPFDFVFHGGSGSEK<br>LGLEEGSKPFDFVFHGGSGSEKEK<br>YLLAATFGNVHGVYKPGNVK | 100-116<br>238-259<br>238-261<br>202-221 | 37274 |
| 475 | Bifunctional phosphoribosylaminoimidazole-carboxamide formyltransferase/IMP cyclohydrolase | Prescottella equi | WP_069856157.1 | IAEAGIPVTK | 44-53 | 55572 |
| 476 | 3-Dehydroquinate synthase | C. aurimucosum<br>C. guaraldiae<br>C. simulans<br>C. accolens<br>C. minutissimum<br>C. singulare<br>C. striatum | WP_046648553.1<br>WP_154736868.1<br>WP_239238695.1<br>PCC82818.1<br>WP_115022329.1<br>WP_144790040.1<br>MDU3175818.1 | FVALTGIGETTR | 327-338 | 38010 |
| 477 | Acyl-CoA carboxylase subunit beta | C. durum | WP_179418910.1 | AEATYPMGR | 31-39 | 59067 |
| 478 | Aminotransferase class I/II-fold pyridoxal phosphate-dependent enzyme | C. diphtheria<br>C. pseudotuberculosis<br>C. striatum<br>C. simulans<br>C. ulcerans<br>C. belfantii | WP_181997845.1<br>WP_014654972.1<br>HCG2963147.1<br>WP_284841028.1<br>WP_197913576.1<br>WP_196981647.1 | GLGEAMVGINVADVPAPHR | 273-291 | 31569 |
| 479 | Pyridoxal 5'-phosphate synthase glutaminase subunit PdxT | C. sanguinis<br>C. lipophiloflavum | WP_222422305.1<br>WP_006840326.1 | SLGIIPMR | 98-105 | 20069 |
| 480 | Aminodeoxychorismate lyase | C. aurimucosum, several other C. | WP_102233567.1 | GDGIFESLLVR | 41-51 | 31465 |
| 481 | Deoxyribodipyrimidine photo-lyase | Clavibacter michiganensis | WP_214582960.1 | DDTLGAWQRGETGVPLVDAGMRALWK | 335-360 | 56725 |
| 482 | Bifunctional 3,4-dihydroxy-2-butanone-4-phosphate synthase/GTP cyclohydrolase II | C. halotolerans<br>C. kutscheri | WP_042440395.1<br>WP_046439504.1 | HETQIER | 219-225 | 45664 |
| 483 | ATP-dependent 6-phosphofructokinase | C. simulans<br>C. minutissimum<br>C. aurimucosum | WP_062041068.1<br>WP_039675088.1<br>GAA1472382.1 | DLYDDAEIDR<br>EGTMDFEEGGVDQFGHQTFNGIGQVIGDEIK<br>ILIVEVMGR<br>LAVLTSGGDCPGLNAVIR | 51-60<br>229-259<br>164-172<br>3-20 | 36729 |
| 484 | FAD-dependent oxidoreductase | C. amycolatum<br>C. vitaeruminis | WP_154844139.1<br>WP_048760340.1 | GIIMALHR | 58-65 | 50725 |
| 485 | Aryldialkylphosphatase | C. halotolerans | WP_015400468.1 | FADLFAK | 125-131 | 34745 |
| 486 | MULTISPECIES: | Corynebacterium | WP_070545790.1 | GFVLGISGGQDSTLAGR | 51-67 | 30512 |

| | | | | | | |
|---|---|---|---|---|---|---|
| | Ammonia-dependent NAD(+) synthetase | | | | | |
| 487 | Methyltransferase domain-containing protein | C. simulans<br>C. phoceense<br>C. striatum<br>C. casei<br>C. cystitides | WP_248090772.1<br>WP_303938692.1<br>MDU3175822.1<br>WP_098072998.1<br>WP_257182798.1 | QGYVTLAPGAGLK | 40-52 | 30552 |
| 488 | 2-Isopropylmalate synthase | C. glucuronolyticum | WP_259815016.1 | DLGRTYEAVIR | 396-406 | 63469 |
| 489 | Alpha/beta hydrolase family protein | C. pseudodiphtheriticum<br>C. propinquum | WP_027017424.1<br>WP_018121555.1 | NIPVQIQPASRGGNAGLYLLDGLR | 71-94 | 37061 |
| 490 | Ppx/GppA phosphatase family protein | C. aurimucosum<br>C. minutissimum<br>C. faecipullorum<br>C. singulare | WP_201828444.1<br>WP_039674654.1<br>HIX79448.1<br>WP_239179901.1 | ADVIGGGCVAVEGIMTMIER | 273-292 | 34740 |
| 491 | Fumarylacetoacetate hydrolase family protein | C. striatum<br>C. argentoratense<br>C. incognita<br>C. simulans | WP_049063663.1<br>WP_278762802.1<br>WP_185174979.1<br>WP_248092820.1 | ATDYVVPALEILSSR | 130-144 | 28690 |
| 492 | Glycosyltransferase | Prescottella equi | WP_084848591.1 | SLSVTLGR | 437-444 | 68982 |
| 493 | Choline dehydrogenase | C. tuberculostearicum<br>C. aurimucosum<br>C. kefirresidentii<br>C. curieae<br>C. pseudogenitalium<br>C. marquesiae<br>C. minutissimum<br>C. accolens<br>C. simulans | WP_316981723.1<br>WP_049358967.1<br>WP_284798795.1<br>WP_269946578.1<br>WP_005323238.1<br>WP_284788328.1<br>WP_239187224.1<br>WP_039885350.1<br>WP_062036316.1 | GPATSELFQALFK | 170-182 | 66333 |
| 494 | Pyridoxal 5'-phosphate synthase lyase subunit PdxS | C. striatum<br>C. simulans<br>C. aurimucosum<br>C. hesseae<br>C. intestinale | WP_046646570.1<br>WP_284841028.1<br>HCT9180310.1<br>WP_269948389.1<br>WP_250224612.1 | AATLYDQPAELAK<br>ELQAPYDLVAEVAETGK<br>QMGAEGVFVGSGIFK | 263-275<br>197-213<br>235-249 | 31909 |
| 495 | Glycoside hydrolase family 25 protein | C. accolens<br>C. tuberculostearicum<br>C. pseudogenitalium<br>C. phoceense<br>C. simulans | WP_302503850.1<br>WP_179386396.1<br>WP_005323361.1<br>WP_257036850.1<br>WP_284841279.1 | SDGQSFAFVK | 60-69 | 40882 |
| 496 | Polyprenyl synthetase family protein | C. simulans<br>C. minutissimum<br>C. faecipullorum<br>C. lizhenjunii | WP_248093681.1<br>WP_039673571.1<br>HIX78886.1<br>WP_165241219.1 | WTNSVAILAGDALLAHASR | 116-134 | 36290 |
| 497 | RNA 2',3'-cyclic phosphodiesterase | C. halotolerans | WP_027004290.1 | YETVSEIR | 170-177 | 20297 |
| 498 | Acyl-CoA thioesterase | C. simulans<br>C. striatum | WP_239240650.1<br>WP_114976566.1 | FMAAPTDMTIAGAQGIGGGR | 23-42 | 34793 |
| 499 | Indole-3-glycerol | C. ulcerans | WP_101507066.1 | VAALDQAR | 141-148 | 28378 |

| | | | | | | |
|---|---|---|---|---|---|---|
| | phosphate synthase TrpC | C. pseudotuberculosis | WP_014522225.1 | | | |
| 500 | Bifunctional methylenetetrahydrofolate dehydrogenase/methenyltetrahydrofolate cyclohydrolase | C. falsenii<br>C. simulans<br>C. lubricantis<br>C. cystitides<br>C. maris | WP_257161886.1<br>WP_096336518.1<br>WP_018297679.1<br>WP_224208375.1<br>WP_020934029.1 | LYRDEIFADLK | 10-20 | 30087 |
| 501 | Aminotransferase class I/II-fold pyridoxal phosphate-dependent enzyme | Tsukamurella paurometabola | WP_013125514.1 | ALAQGPSAAIIQPR | 228-241 | 49535 |
| 502 | Glycosyltransferase family 39 protein | Tsukamurella paurometabola | WP_013128542.1 | TETTTSATADR | 2-12 | 54912 |
| 503 | Carboxylesterase/lipase family protein | C. pseudotuberculosis | WP_014300614.1 | LGPLELER | 529-536 | 59482 |
| 504 | Aminotransferase class V-fold PLP-dependent enzyme | C. pseudotuberculosis<br>C. striatum | WP_014366315.1<br>HAT1549280.1 | VFDVAR | 2-7 | 42211 |
| 505 | Penicillin-binding protein 1A; transglycosylase domain-containing protein | C. cystitides<br>C. efficiens<br>C. freiburgense | WJY83634.1<br>WP_006768821.1<br>WP_240483188.1 | SLPGIPK | 430-437 | 77423 |
| 506 | PaaI family thioesterase | Prescottella equi | WP_080561082.1 | KPTPLGK | 167-173 | 24489 |
| 507 | 2-C-methyl-D-erythritol 4-phosphate cytidylyltransferase | C. falsenii | WP_119665254.1 | LGFSTPK | 20-26 | 30829 |
| 508 | Maleylacetate reductase; IS3 family transposase | C. glyciniphilum<br>C. efficiens | WP_038549333.1<br>WP_049783600.1 | MAVEGIR | 197-203 | 23618 |
| 509 | Sigma-70 family RNA polymerase sigma factor | C. durum | WP_231287024.1 | NTAFADR | 103-109 | 21665 |
| 510 | N-acetyl-gamma-glutamyl-phosphate reductase | C. pseudotuberculosis | WP_014366976.1 | YVSGELK | 28-34 | 40384 |
| 511 | Prolipoprotein diacylglyceryl transferase | C: pyruviciproducens | WP_276921981.1 | RFNMGR | 215-220 | 32097 |
| 512 | 2-Isopropylmalate synthase | C. glucuronolyticum | WP_259815016.1 | KDKEGVK | 172-178 | 60839 |
| 513 | Aspartate kinase | C. urealyticum<br>C. amycolatum<br>C. xerosis<br>C. lactis<br>C. vitaeruminis | WP_148791705.1<br>WP_088611847.1<br>WP_168937586.1<br>WP_120492735.1<br>WP_048759095.1 | IVEVTPGR | 113-120 | 44906 |

| | | | | | | |
|---|---|---|---|---|---|---|
| | | C. kutscheri | WP_046438430.1 | | | |
| 514 | Hydroxyethylthiazole kinase | C. accolens<br>C. macginleyi | WP_005282916.1<br>MBM0243883.1 | MDCSFLR | 1-7 | 27715 |
| 515 | Acetyl/propionyl/methylcrotonyl-CoA carboxylase subunit alpha | C. accolens<br>C. macginleyi | WP_284631475.1<br>WP_200441425.1 | GAQLLEIK | 579-586 | 62305 |
| 516 | Malate dehydrogenase | C. casei<br>C. ammoniagenes | WP_025388002.1<br>WP_003847289.1 | DMVADLLK | 329-336 | 35773 |
| 517 | Alpha/beta hydrolase family protein | C. amycolatum<br>C. jeikeium<br>C. vitaeruminis | WP_115598374.1<br>ASE56930.1<br>WP_048760157.1 | NLQNSKAR | 323-330 | 36464 |
| 518 | tRNA dihydrouridine synthase DusB | C. striatum | WP_114976561.1 | VESVDNLR | 319-326 | 41238 |
| 519 | Bifunctional riboflavin kinase/FAD synthetase | C. argentoratense | WP_315040446.1 | ASGTTETLR | 132-140 | 37040 |
| 520 | UDP-N-acetylglucosamine--N-acetylmuramyl-(pentapeptide) pyrophosphoryl-undecaprenol N-acetylglucosamine transferase | Rhodococcus sp. AG1013 | WP_114720055.1 | MLAGEPAPR | 1-9 | 39984 |
| 521 | PIG-L family deacetylase | Tsukamurella paurometabola | WP_013126697.1 | LIVRLPPR | 224-231 | 27491 |
| 522 | Glycerate kinase | C. efficiens | WP_006767981.1 | SNDVAAQLR | 344-352 | 37099 |
| 523 | dUTP diphosphatase | C. jeikeium | WP_035011199.1 | RNGNDAPLR | 5-13 | 16458 |
| 524 | Formate-dependent phosphoribosylglycinamide formyltransferase | C. pseudodiphtheriticum<br>C. propinquum | WP_284815282.1<br>WP_311199475.1 | ACELTMER | 121-128 | 52724 |
| 525 | Bifunctional cytidylate kinase/GTPase Der | C. argentoratense<br>C. glutamicum | WP_314929774.1<br>WP_040072710.1 | AHRCVVEGR | 129-137 | 24800 |
| 526 | Glycosyltransferase family 87 protein | Prescottella equi | WP_286461333.1 | LASTIGGPVGR | 46-56 | 58446 |
| 527 | Acyl-CoA dehydrogenase family protein | C. aurimucosum | WP_010188978.1 | ADLSMAEQR | 76-84 | 76400 |
| 528 | Type I pantothenate kinase | C. maris | WP_020934321.1 | KGFPEAYDR | 143-151 | 35160 |
| 529 | Bifunctional indole-3-glycerol-phosphate synthase TrpC/phosphoribosylanthranilate isomerase Trp | Corynebacterium sp. KPL1818 | WP_023029750.1 | LLNQPEKKD | 467-475 | 51771 |
| 530 | Thiamine phosphate | C. kroppenstedtii | WP_012732318.1 | VAAAGAGVVQVR | 28-39 | 22923 |

| | | | | | | |
|---|---|---|---|---|---|---|
| | synthase | | | | | |
| 531 | cation:proton antiporter | C. falsenii | WP_224208616.1 | ALVADDSRPR | 241-250 | 43471 |
| 532 | Dihydropteroate synthase | C. lipophiloflavum<br>C. sanguinis | WP_006840571.1 | AGVDELTVPGR | 2-12 | 29570 |
| 533 | D-alanyl-D-alanine carboxypeptidase/D-alanyl-D-alanine-endopeptidase | C. pseudotuberculosis | WP_058832193.1 | ATLDAMAHDR | 56-65 | 40521 |
| 534 | Pantoate--beta-alanine ligase | C. efficiens | WP_041628383.1 | MSFEHDQGR | 1-9 | 19720 |
| 535 | Mur ligase family protein | Clavibacter michiganensis | WP_104343076.1 | FGYEGLEVGR | 368-377 | 46163 |
| 536 | HNH endonuclease signature motif containing protein | C. amycolatum<br>C. vitaeruminis<br>C. jeikeium | WP_197915149.1<br>WP_048759202.1<br>AYX80873.1 | KNGGATTMENMVMTCKEHNAANDDDR | 318-343 | 43433 |
| 537 | Pyruvate kinase | C. urealyticum | WP_148792403.1 | GLIDAGMNVAR | 23-33 | 50514 |
| 538 | NAD(P)-binding domain-containing protein | C. glucuronolyticum | WP_005395145.1 | AALNSSPTTDR | 2-12 | 37999 |
| 539 | Linear gramicidin synthase subunit D | C. durum | WJY83809.1 | VQVVTTDAGPR | 887-897 | 160212 |
| 540 | MetQ/NlpA family ABC transporter substrate-binding protein | C. accolens | WP_284639927.1 | SHKSIDDVDK | 125-134 | 30747 |
| 541 | RNA pseudouridine synthase B | C. pseudotuberculosis | AEK92306.1 | GSEGSRAGENTRESYGFR | 65-82 | 41826 |
| 542 | FdhF/YdeP family oxidoreductase | Tsukamurella paurometabola | WP_013126973.1 | LNRSHLVHGR | 518-527 | 85189 |
| 543 | RNA pseudouridine synthase B | C. pseudotuberculosis | AEK92306.1 | GSEGSRAGENTRESYGFR | 65-82 | 41826 |
| 544 | FdhF/YdeP family oxidoreductase | Tsukamurella paurometabola | WP_013126973.1 | LNRSHLVHGR | 518-527 | 85189 |
| 545 | Deoxycytidylate deaminase | C. terpenotabidum | WP_020440770.1 | RAIELSEESR | 14-23 | 12280 |
| 546 | Maleylpyruvate isomerase N-terminal domain-containing protein | C. halotolerans | WP_015399730.1 | EQSQRAVEPR | 120-129 | 30118 |
| 547 | 3-Hydroxy-9,10-secoandrosta-1,3,5(10)-triene-9,17-dione monooxygenase oxygenase subunit | Prescottella equi | WP_084922618.1 | DRAGQVERDR | 22-32 | 44016 |
| 548 | Glutamate synthase large subunit | C. casei<br>C. xerosis<br>C. stationis | WP_276688781.1<br>WP_155868675.1<br>WP_283117428.1 | VQCDGQLKTGR | 1098-1108 | 167177 |

| | | | | | | |
|---|---|---|---|---|---|---|
| | | C. amycolatum<br>C. jeikeium<br>C. vitaeruminis | WP_272715369.1<br>ASE57065.1<br>WP_239659444.1 | | | |
| 549 | Glyoxalase | Prescottella equi | BCN68035.1 | EQVFDMSGATR | 6-16 | 14821 |
| 550 | PfkB family carbohydrate kinase | C. halotolerans | WP_015399904.1 | VVLDGAVTDPEK | 160-171 | 30591 |
| 551 | Phosphomannomutase/ phosphoglucomutase | Clavibacter michiganensis | WP_209657882.1 | DQMAATGAVFGGEHSAHYYFR | 331-351 | 48868 |
| 552 | Alpha/beta hydrolase, partial | Clavibacter michiganensis | WP_259340805.1 | AGLGDSDPDPAPR | 66-78 | 30139 |
| 553 | Cellulose biosynthesis cyclic di-GMP-binding regulatory protein BcsB | Prescottella equi | WP_238772026.1 | MARTGAPSRAPR | 1-12 | 67672 |
| 554 | L-serine ammonia-lyase, iron-sulfur-dependent, subunit alpha, partial | C. falsenii | WP_272712594.1 | RDHDSEAMPGAK | 203-214 | 48405 |
| 555 | 16S rRNA (uracil(1498)-N(3))-methyltransferase | C. variabile | WP_313007511.1 | WDGKPGKAEKAR | 119-130 | 26362 |
| 556 | 3-Oxoacid CoA-transferase subunit B | C. xerosis<br>C. lactis<br>C. maris<br>C. humireducens<br>C. amycolatum<br>C. vitaeruminis<br>C. breve<br>C. sanguinis<br>C. glyciniphilum<br>C. falsenii | WP_269202127.1<br>WP_281175560.1<br>WP_041631673.1<br>WP_040085286.1<br>WP_239272412.1<br>WP_048760832.1<br>WP_284825906.1<br>WP_181729582.1<br>WP_145941902.1<br>WP_025401750.1 | GMGGAMDLVAGTPR | 163-176 | 22728 |
| 557 | Mycothiol synthase | C. glyciniphilum | WP_144313691.1 | MSVDCTDPDRR | 144-154 | 35877 |
| 558 | Alpha/beta hydrolase | C. glutamicum | WP_003857213.1 | DCYLTKDTMER | 193-203 | 31600 |
| 559 | Cyclic nucleotide-binding domain-containing protein | C. diphtheriae | WP_199333144.1 | MLATPCADFMSR | 119-130 | 43258 |
| 560 | Urea amidolyase associated protein UAAP1 | Tsukamurella paurometabola | WP_013126244.1 | YGAGEAHSDSPAGR | 151-164 | 29752 |
| 561 | 3-Methyl-2-oxobutanoate hydroxymethyltransferase | C. urealyticum | WP_149121667.1 | QDSPRGYMVPTK | 4-15 | 29554 |
| 562 | Phosphoserine transaminase | C. matruchotii | WP_314822380.1 | EEVSCFVSDPNK | 293-304 | 40106 |
| 563 | Aldehyde dehydrogenase | Prescottella equi | WP_221282760.1 | MTDYDKLFIGGR | !-!2 | 50913 |
| 564 | Protoporphyrinogen | C. amycolatum | WP_150963137.1 | FDMGHGGLMDFAR | 440-452 | 51387 |

| | | | | | | |
|---|---|---|---|---|---|---|
| | oxidase | | | | | |
| 565 | SDR family oxidoreductase | C. jeikeium | WP_011272816.1 | LEQVAQAITQEGGK | 63-76 | 28641 |
| 566 | NAD(P)H-dependent glycerol-3-phosphate dehydrogenase | C. terpenotabidum | WP_041631453.1 | VCADAGCDVTLWAR | 18-31 | 34420 |
| 567 | Non-ribosomal peptide synthetase | Tsukamurella paurometabola | WP_013128267.1 | FGAGDDICIGTPVAGR | 1252-1267 | 241289 |
| 568 | Aminotransferase class I/II-fold pyridoxal phosphate-dependent enzyme | C. diphtheriae | WP_259344139.1 | RGGVGRNIALLFIPK | 232-246 | 27016 |
| 569 | Decaprenylphosphoryl-beta-D-ribose oxidase | C. variabile<br>C. glyciniphilum | HAF73214.1<br>WP_145942108.1 | RSIDPTGVFASDMSR | 452-466 | 51150 |
| 570 | Phosphoglucomutase (alpha-D-glucose-1,6-bisphosphate-dependent) | C. argentoratense | WP_234858678.1 | MDCSSPDSMASLVANR | 285-300 | 58129 |
| 571 | Phosphoglucomutase (alpha-D-glucose-1,6-bisphosphate-dependent) | C. maris<br>C. xerosis<br>C. freneyi<br>C. lactis | WP_020935557.1<br>WP_046650327.1<br>WP_035123894.1<br>WP_120491165.1 | MDCSSPDAMASLIGNR | 285-300 | 58457 |
| 572 | PII uridylyl-transferase | Corynebacterium diphtheriae bv. gravis str. ISS 4749 | KLN44026.1 | SMMPFSATPADIRER | 2-16 | 79233 |
| 573 | Bifunctional phosphopantothenoylcysteine decarboxylase/phosphopantothenate--cysteine ligase CoaBC | C. callunae | WP_015651391.1 | KGCDLLMCNEVGVCK | 359-373 | 43632 |
| 574 | Bifunctional DNA-formamidopyrimidine glycosylase/DNA-(apurinic or apyrimidinic site) lyase | C. aurimucosum | WP_010190448.1 | GTHYCPSCQNRGVGR | 263-277 | 30701 |
| 575 | Aldo/keto reductase | C. pseudodiphtheriticum | WP_284586931.1 | GLGCTGMTPHSYSDSPR | 37-53 | 35213 |
| 576 | GMC family oxidoreductase N-terminal domain-containing protein | C. glyciniphilum | WP_052540052.1 | DSTTSAEHTSTMIGEPL | 503-519 | 55720 |
| 577 | 3-Hydroxyacyl-CoA dehydrogenase NAD-binding domain-containing protein | C. amycolatum | WP_069358929.1 | ATEDDAADMFGMVEDMK<br>QMSKATEDDAADMFGMVEDMKASLR | 82-98<br>78-102 | 79996 |

| | | | | | | |
|---|---|---|---|---|---|---|
| 578 | dGTP Triphosphohydrolase | C. kroppenstedtii<br>C. parakroppenstedtii<br>C. pseudokroppenstedtii | ACR17423.1<br>WP_244998915.1<br>WP_284867034.1 | RAMTNGDESAWTEEMAAGR | 317-335 | 48525 |
| 579 | Nucleoside hydrolase | C. accolens | WP_284637326.1 | EHGDAVAGCGSLTIMGGAVNYR | 139-160 | 35026 |
| 580 | Acyl-CoA thioesterase II | C. kefirresidentii | WP_239204103.1 | NGDCGPQHQDPMPEVPGPEETAR | 107-129 | 32261 |
| 581 | 3-Hydroxyacyl-CoA dehydrogenase NAD-binding domain-containing protein | C. amycolatum<br>C. vitaeruminis | WP_256884522.1<br>WP_048760221.1 | QMSKATEDDAADMFNMVEDMKASLR | 78-102 | 80070 |
| 582 | Methylated DNA-protein cysteine methyltransferase | C. diphtheriae | AEX49001.1 | MHAEYGFVFTPDGQFCVVTDSMTHK | 1-25 | 18502 |
| 583 | Putative Apolipoprotein N-acyltransferase | C. halotolerans | AGF73868.1 | LFRGIAGDNWRRLAGPTALMMGAAVVWWLALPPR | 6-39 | 54827 |
| 584 | NAD(P)H-dependent oxidoreductase | C. appendicis | WP_284834681.1 | ELAAMCDR | 164-171 | 20283 |
| 585 | Putative cyclopropane-fatty-acyl-phospholipid synthase | C. efficiens | BAC18404.1 | MPPGGAPKR | 2-10 | 48631 |
| 586 | Phosphotransferase | C. pyruviproducens | WP_276847531.1 | MIDTDSLK | 1-8 | 41916 |
| 587 | PIG-L family deacetylase | Tsukamurella paurometabola | WP_013126697.1 | LIVRLPPR | 224-231 | 27491 |
| 588 | Cation: dicarboxylase symporter family transporter | C. vitaeruminis | WP_277102341.1 | VNIFKLCK | 268-275 | 50429 |
| 589 | LLM class flavin-dependent oxidoreductase | C. kroppenstedtii | WP_012731253.1 | ATDGNGYTAR | 16-25 | 37580 |
| 590 | 3-Dehydroquinate synthase | C. terpenotabidum | WP_020441204.1 | GLPCGEVFR | 43-51 | 59830 |
| 591 | 3-Hydroxyacyl-CoA dehydrogenase NAD-binding domain-containing protein | C. amycolatum<br>C. vitaeruminis | WP_256884522.1<br>WP_048760221.1 | QMSKATEDDAADMFNMVEDMKASLR | 78-102 | 80070 |
| 592 | 3-Hydroxyacyl-CoA dehydrogenase NAD-binding domain-containing protein | C. amycolatum<br>C. vitaeruminis | WP_256884522.1<br>WP_048760221.1 | QMSKATEDDAADMFNMVEDMKASLR | 78-102 | 80070 |
| 593 | Methylated DNA-protein cysteine methyltransferase | C. diphtheriae | AEX49001.1 | MHAEYGFVFTPDGQFCVVTDSMTHK | 1-25 | 18502 |
| 594 | Putative Apolipoprotein N-acyltransferase | C. halotolerans | AGF73868.1 | LFRGIAGDNWRRLAGPTALMMGAAVVWWLALPPR | 6-39 | 54827 |
| 595 | NAD(P)H-dependent oxidoreductase | C. appendicis | WP_284834681.1 | ELAAMCDR | 164-171 | 20283 |

| | | | | | | |
|---|---|---|---|---|---|---|
| 596 | Putative cyclopropane-fatty-acyl-phospholipid synthase | C. efficiens | BAC18404.1 | MPPGGAPKR | 2-10 | 48631 |
| 597 | Phosphotransferase | C. pyruviproducens | WP_276847531.1 | MIDTDSLK | 1-8 | 41916 |
| 598 | PIG-L family deacetylase | Tsukamurella paurometabola | WP_013126697.1 | LIVRLPPR | 224-231 | 27491 |
| 599 | Cation:dicarboxylase symporter family transporter | C. vitaeruminis | WP_277102341.1 | VNIFKLCK | 268-275 | 50429 |
| 600 | LLM class flavin-dependent oxidoreductase | C. kroppenstedtii | WP_012731253.1 | ATDGNGYTAR | 16-25 | 37580 |
| 601 | 3-Dehydroquinate synthase | C. terpenotabidum | WP_020441204.1 | GLPCGEVFR | 43-51 | 59830 |
| 602 | Class I SAM-dependent DNA methyltransferase | C. glutamicum | WP_216312896.1 | EWKTTEDATQQSGPLTR | 39-55 | 171706 |
| 603 | L-lactate dehydrogenase | C. falsenii<br>C. macclintockii<br>C. jeikeium<br>C. dentalis | WP_119665041.1<br>WP_051873906.1<br>WP_111711794.1<br>WP_312098311.1 | DAAYEIIQAK | 251-260 | 33706 |
| 604 | Pyridoxal 5'-phosphate synthase glutaminase subunit PdxT | Tsukamurella paurometabola | WP_303175547.1 | CLAAIDMTVR | 96-105 | 21195 |
| 605 | Dihydrolipoyl dehydrogenase | Prescottella equi | WP_084985381.1 | NAELAHIFHK | 56-65 | 49788 |
| 606 | Nitrite/sulfite reductase | C. amycolatum<br>C. xerosis | WP_005510177.1<br>WP_155868296.1 | VEDMPEIWDR | 155-164 | 63278 |
| 607 | MobF family relaxase | C. callunae | WP_015650638.1 | EMGVEFVAHDR | 304-314 | 146531 |
| 608 | putative peptidoglycan glycosyltransferase FtsW | C. falsenii | WP_025402498.1 | QTLPPSGSVKRR | 470-481 | 53451 |
| 609 | 3-Isopropylmalate dehydratase small subunit | C. casei | WP_006823095.1 | DEDAISNFESAR | 175-186 | 22438 |
| 610 | PhoX family phosphatase | C. jeikeium | WP_034984422.1 | EDGSAESHVDGMSAEEVIIYTR | 465-486 | 77065 |
| 611 | Aminotransferase class I/II-fold pyridoxal phosphate-dependent enzyme | C. genitalium | WP_005287311.1 | ENYTDKSGTDVR | 57-68 | 46832 |
| 612 | Class I SAM-dependent DNA methyltransferase | C. glutamicum | WP_216312896.1 | EWKTTEDATQQSGPLTR | 39-55 | 171706 |
| 613 | dTDP-glucose 4,6-dehydratase | Clavibacter michiganensis | WP_316307754.1 | DWSYVDRVEDR | 264-274 | 36758 |
| 614 | NAD(P)/FAD-dependent | C. glutamicum<br>C. callunae | WP_074508215.1<br>WP_247775189.1 | DGEEHTIESFCK | 262-271 | 50896 |

| | | | | | | |
|---|---|---|---|---|---|---|
| | oxidoreductase | | | | | |
| 615 | Putative dihydropteroate synthase | C. efficiens | BAC17978.1 | MMSTRPSVSSSMHTR | 1-15 | 34228 |
| 616 | Zinc-dependent alcohol dehydrogenase family protein | C. matruchotii | WP_311324187.1 | QFDVAFDCVGGEMGRR | 181-196 | 31751 |
| 617 | Class I SAM-dependent methyltransferase | C. durum | WP_006062010.1 | MRASEAMTNQESSAANR | 1-17 | 29368 |
| 618 | Succinic semialdehyde dehydrogenase | C. maris | WP_020933498.1 | QDELMDVIQDESGKNR | 63-78 | 52144 |
| 619 | Type I glyceraldehyde-3-phosphate dehydrogenase | C. pyruviciproducens | WP_016457562.1 | LSDDITYDDESITVDGK | 55-71 | 36311 |
| 620 | Biotin synthase BioB | C. diphtheria<br>C. belfantii<br>C. ulcerans<br>C. pseudotuberculosis<br>C. rouxii<br>C. silvaticum | WP_082258447.1<br>WP_088266771.1<br>WP_197913658.1<br>WP_013241764.1<br>WP_155872587.1<br>WP_134316284.1 | SGRCPENCTFCAQSIR | 63-78 | 34818 |
| 621 | Transketolase | C. ammoniagenes<br>C. stationis<br>C. confusum | WP_003845895.1<br>WP_191373057.1<br>WP_290226308.1 | MNTGGAHGAALGDDEVAATK | 268-287 | 74755 |
| 622 | MtrAB system histidine kinase MtrB | C. accolens<br>C. macginleyi<br>C. marquesiae<br>C. tuberculostearicum<br>C. pseudogenitalium<br>C. kefirresidentii<br>C. aurimucosum | WP_284640109.1<br>WP_275403430.1<br>WP_284837145.1<br>WP_301715266.1<br>WP_279625203.1<br>WP_208612449.1<br>WP_193387578.1 | DDEMGRLAASFNNMADK | 242-258 | 67812 |
| 623 | Glyceraldehyde-3-phosphate dehydrogenase | C. genitalium | WP_005291253.1 | TEAAGQPQNLTEDWNNK | 2-18 | 54176 |
| 624 | Phosphoenolpyruvate carboxykinase (GTP) | Tsukamurella paurometabola | WP_197715991.1 | HGVFMGATVGSEQTAAAEGK | 447-466 | 67613 |
| 625 | 3-Hydroxyisobutyryl-CoA hydrolase | C. glucuronolyticum | WP_005391823.1 | SEDLDSFAEMAISESLDEAR | 199-218 | 38491 |
| 626 | Fused MFS/spermidine synthase | C. ammoniagenes | WP_003848709.1 | DVFSGAETPTDLTTVEFFQAAHR | 160-182 | 31132 |
| 627 | Allantoinase AllB | C. falsenii | WP_025401798.1 | FATAAPAPFSAMDAEGTNLGGAGADGR | 341-367 | 56064 |
| 628 | CoA transferase | Prescottella equi | WP_084968951.1 | DRTGFAQTMEQATGLSWMTGYR | 528-549 | 85877 |
| 629 | Diacylglycerol kinase | C. variabile<br>C. nuruki<br>C. kalidii<br>C. glyciniphilum | WP_303948348.1<br>WP_010121418.1<br>WP_244803269.1<br>WP_145943038.1 | EHWFGTIACAGFDSLVSDRTNR | 155-176 | 35641 |

| | | | | | | |
|---|---|---|---|---|---|---|
| 630 | Phosphoribosylformyl glycinamidine cyclo-ligase | C. aurimucosum<br>C. guaraldiae<br>C. singulare<br>C. minutissimum | WP_102233565.1<br>WP_154736928.1<br>WP_144792341.1<br>WP_039676512.1 | GIAEGCVQAGAALLGGETAEHPGVMGK | 121-147 | 36831 |
| 631 | Lipoyl synthase | C. variabile | WP_014009663.1 | VSDAGLHTVCQEAGCPNIHECWEDR | 46-70 | 39252 |
| 632 | Isochorismate synthase | C. jeikeium<br>C. evansiae | WP_071056918.1<br>WP_269944248.1 | SPREFYSGAVGWCDSSGDGEFMLSIR | 305-330 | 41029 |
| 633 | Adenylosuccinate lyase | C. glutamicum<br>C. suranareeae<br>C. faecale | WP_173328348.1<br>WP_197702441.1<br>WP_290279898.1 | GPMGTAQDMLDLMDGDEARLSDLETR | 195-220 | 52192 |
| 634 | Aromatic ring-hydroxylating dioxygenase subunit alpha | C. halotolerans | WP_015402124.1 | YMRSMHGCGFNLNMFPNIGMSSAFFR | 292-317 | 48330 |
