## Supplemental Table S18 for "Peptidome Analysis of Western Blots Identifies Natural Bispecific Antibody-Bound *Corynebacterium* and Phage B-cell Epitopes with Potential Relevance to Psoriasis"

### Table S18. Bacterial Immune Proteins are *Corynebacterium* Antigens.

| Antigen Number | Identified Antigen | *Corynebacterium* (C.) | Accession Number | Peptide Sequence | Peptide Start/Stop | Molecular Weight |
|---|---|---|---|---|---|---|
| 1 | CRISPR system CASCADE complex protein CasC | C. striatum | WP_049063132.1 | AIIDKDGEK<br>AIIDKDGEKFK<br>GLDGYFTEFADSVTIPELEER<br>LAQEAIETQR<br>LVGELICK<br>RFPAEVMGK<br>SVSLVNAFEEPVEAGAAGR<br>TAVDSAAAFVDSFVK | 145-153<br>145-155<br>349-369<br>324-333<br>67-74<br>55-63<br>298-316<br>256-270 | 41921 |
| 2 | Type I-E CRISPR-associated protein Cas5/CasD | C. pseudodiphtheriticum | WP_284586519.1 | SGIIGLLAAAEGRR | 34-47 | 26796 |
| 3 | Type II CRISPR RNA-guided endonuclease Cas9 | C. diphtheriae | WP_181993709.1 | EMDGDMR | 510-516 | 121519 |
| 4 | Type II CRISPR RNA-guided endonuclease Cas9 | C. matruchotii | WP_315120567.1 | CAYCGAEISFK | 557-567 | 123355 |
| 5 | Type I-E CRISPR-associated protein Cas6/Cse3/CasE | C. striatum<br>C. simulans<br>C. resistens | WP_100086674.1<br>WP_062043657.1<br>WP_042379763.1 | AYGCGLLTLAPAE<br>FLEQIVNGQK<br>GAFPPDIDESQSR<br>LLANPQAMHAAVR | 207-219<br>90-99<br>32-44<br>19-31 | 24213 |
| 6 | Type I-E CRISPR-associated protein Cse2/CasB | C. striatum | WP_049192140.1 | NIGFDYGAFAQDLK | 159-172 | 22653 |
| 7 | Type I restriction enzyme, S subunit | C. diphtheriae | WP_014304115.1 | SGVLELGDGYR | 11-21 | 45368 |
| 8 | Restriction endonuclease subunit S | C. jeikeium | WP_011273488.1 | LKVAGPGTVLFAMYGATLGAVSR | 264-286 | 44305 |
| 9 | Type I restriction enzyme HsdR N-terminal domain-containing protein | C. variabile<br>C. terpenotabidum<br>C. nuruki | WP_014010639.1<br>WP_020441549.1<br>WP_010118628.1 | QAHDIIR | 276-282 | 40798 |
| 10 | Restriction endonuclease | C. striatum<br>C. simulans<br>C. aurimucosum<br>C. minutissimum<br>C. guaraldiae | WP_062042895.1<br>WP_114976733.1<br>WP_201828944.1<br>KKO78158.1<br>WP_144014664.1 | NNEFVAGELLTR | 156-167 | 14537 |
