## Supplemental Table S19 for "Peptidome Analysis of Western Blots Identifies Natural Bispecific Antibody-Bound *Corynebacterium* and Phage B-cell Epitopes with Potential Relevance to Psoriasis"

### Table S19. Chaperones and Chaperonines are *Corynebacterium* Antigens.

| Antigen Number | Identified Antigen | *Corynebacterium* (C.), *Mycobacterium* (M.) *tuberculosis* (Peptide numbers) | Accession Number | Peptide Sequence | Peptide Start/Stop | Peptide Number | Molecular Weight |
|---|---|---|---|---|---|---|---|
| 1 | Molecular chaperone DnaK | C. striatum (1,2,3,4,5,6,7,8,9,10,11,12,13,14,15,16,17,18,19,20,21 | WP_110332849.1 | APFNQVIK | 291-298 | 1 | 66319 |
| | | C. simulans (1,2,3,4,5,6,7,8,9,10,11,12,13,14,15,16,17,18,19,20,21 | WP_339018847.1 | AVGIDLGTTNSVVSVLEGGEATVIANSEGAR | 4-34 | 2 | |
| | | C. camporealensis (1,5,6,8,12,13,15,21 | WP_368531007.1 | DAEAYLGEDVTDAVITVPAYFEDSQR | 102-127 | 3 | |
| | | C. minutissimum (3,4,5,16 | WP_039672506.1 | DAGLSLSEVDHVVLVGGSTR | 299-318 | 4 | |
| | | C. singulare (3,4,5,16 | WP_144793021.1 | DVLLLDVTPLSLGIETK | 361-377 | 5 | |
| | | C. aurimucosum (1,3,4,5,8,12,16 | WP_010188673.1 | EAGQIAGLNVLR | 132-143 | 6 | |
| | | C. phocae (1,5,6,8,9,11,12,13,14,15,17,18,21 | WP_075735831.1 | GVNPDEVVAVGAALQAGVLR | 337-356 | 7 | |
| | | C. macginleyi (1,5,6,7,8,9,11,14,15,17,18 | WP_200443708.1 | IQDGSGLSQEEIDR | 474-487 | 8 | |
| | | C. confusum (1,5,6,7,8,9,13,15,17,18,21 | WP_290223484.1 | ITQDLLDR | 281-288 | 9 | |
| | | C. accolens (1,5,6,7,9,11,15,17,18 | WP_284898207.1 | IVDWLVEK | 208-215 | 10 | |
| | | C. lizhenjunii (1,3,5,6,7,8,9,10,11,15,17,18,19 | WP_165007416.1 | IVNEPTAAALAYGLEK | 144-159 | 11 | |
| | | C. phoceense (1,3,5,7,8,10,13,14,21 | WP_368527337.1 | KFLDENSDKVSEDVK | 521-535 | 12 | |
| | | C. sanguinis (1,5,13,14,21 | WP_259927047.1 | KYTPQEISAR | 85-94 | 13 | |
| | | C. breve (1,5,10,13,19,21 | WP_284824564.1 | MPAVSELVK | 319-327 | 14 | |
| | | C. flavescens (1,4,5,13,15,21) | WP_301523537.1 | NGEILVGQSAK | 45-55 | 15 | |
| | | C. kefirresidentii (1,5,8) | WP_284835491.1 | NQAESTSYQTR | 510-520 | 16 | |
| | | C. tuberculostearicum (1,5,8) | WP_316986824.1 | NQAVTNVDR | 56-64 | 17 | |
| | | C. riegelii (1,5,8,13,21) | WP_311342991.1 | RNTTIPTKKS | 388-394 | 18 | |
| | | C. marquesiae (1,5,8) | WP_408934651.1 | SANGIDLTKDK | 218-228 | 19 | |
| | | C. matruchotii (1,5,10,13,21) | WP_315557419.1 | TQVTEAADAVDEALKGDDIEAIK | 536-558 | 20 | |
| | | C. pseudotuberculosis (1,5,6,7,8,11,14,17,18) | WP_054250925.1 | YTPQEISAR | 86-94 | 21 | |
| | | C. durum (1,5,13,19,21) | WP_311143329.1 | | | | |
| | | C. incognita (1,510,13,21 | WP_185176604.1 | | | | |
| | | C. casei (2,5,6,7,8,9,10,11,15,17,18) | MDN6155433.1 | *Akkermansia muciniphila* (WP_215444414.1) | | | |
| | | C. variabile (2,5,14,18) | MFR4189582.1 | *C. simulans* 1 DVLLLDVTPLSLGIET 16<br>DVLLLDVTPL L IET<br>*A. muciniphila* 385 DVLLLDVTPLTLSIET 400 | | | |
| | | C. faecale (3,5,7,8) | WP_290277127.1 | *C. simulans* 2 VGIDLGTTNSVVSVLEGGEATVIANSEGAR 31<br>+GIDLGTTNS ++V+EGG+ATV+ NSEGAR<br>*A. muciniphila* 5 LGIDLGTTNSCMAVMEGGQATVLENSEGAR 34 | | | |
| | | C. glutamicum (3,5,6,10) | WP_077313444.1 | *C. simulans* 1 DAEAYLGEDVTDAVITVPAYFEDSQR 26<br>DAE LGE +T+AVITVPAYF D QR<br>*A. muciniphila* 128 DAESKLGETITEAVITVPAYFNDAQR 153 | | | |
| | | C. hiratae (3,5,16) | WP_144014122.1 | *C. simulans* 1 GVNPDEVVAVGAALQAGVL 19<br>GVNPDEVVAVGAA Q GVL<br>*A. muciniphila* 361 GVNPDEVVAVGAAIQGGVL 379 | | | |
| | | C. guaraldiae (3,5,16) | WP_154762732.1 | *C. simulans* 1 IQDGSGLSQEEIDR 14<br>IQ SGLS EEI+R<br>*A. muciniphila* 498 IQGSSGLSKEEIER 511 | | | |
| | | C. glucuronolyticum (5,6,8,11,12,13,17,21) | EEI64345.1 | *C. simulans* 1 IVNEPTAAALAYGLEK 16<br>I+NEPTAAALAYGL+K<br>*A. muciniphila* 170 IINEPTAAALAYGLDK 185 | | | |
| | | C. glaucum (5,6,11,17,18) | WP_095660756.1 | *Faecalibacterium prausnitzii* (WP_179860689.1) | | | |
| | | C. kroppenstedtii (5,7,9,11,17,18) | WP_303733865.1 | *C. simulans* 1 DVLLLDVTPLSLGIET 16<br>DV LLDVTPLSLGIET<br>*F. prausnitzii* 108 DVVLLDVTPLSLGIET 123 | | | |
| | | C. cystitides (5,11,12,13,15,17,18,19,21) | WP_257160636.1 | | | | |
| | | C. parakroppenstedtii (5,7,9,11,17,18) | WP_221913480.1 | | | | |
| | | C. faecigallinarum (5,7,9,11,17,18) | HJC84837.1 | | | | |
| | | C. pseudokroppenstedtii (5,7,9,11,17,18) | WP_337448767.1 | | | | |
| | | C. pollutisoli (5,6,7,8,9,10,11,17,18) | WP_085549260.1 | | | | |
| | | C. endometrii (5,6,7,8,9,11,12,13,14,17,19,21) | WP_136141924.1 | | | | |
| | | C. jeikeium (5,7,8,9,11,18) | WP_034986699.1 | | | | |
| | | C. suedekumii (5,6,7,8,9,10,11,17,18) | WP_284875358.1 | | | | |
| | | C. falsenii (5,7,9,11,18) | WP_224208042.1 | | | | |
| | | C. nasicanis (5,6,7,8,9,11,13,17,18,21) | WP_377001396.1 | | | | |
| | | C. ammoniagenes (5,6,7,8,9,11,15,17,18) | WP_168939305.1 | | | | |

| | | | | | | | |
|---|---|---|---|---|---|---|---|
| | | C. pseudodiphtheriticum (5,6,9,11,13,15,17,18,21)<br>C. propinquum (5,6,9,11,13,15,17,18,21)<br>C. glyciniphilum (5,7,9,10,11,17,18)<br>C. macclintockiae (5,7,8,9,11,18)<br>C. stationis (5,6,7,8,10,15)<br>C. gallinarum (5,6,8)<br>C. efficiens (5,6,7,8,10)<br>C. resistens (5,7,10)<br>C. deserti (5,8)<br>C. marinum 5,(8)<br>C. ulceribovis (5,8,10)<br>C. pyruviciproducens (5,8,10,13,21)<br>C. dentalis (5,8,10,19)<br>C. anserum (5,8,10)<br>C. auriscanis (5,10,12,14,15)<br>C. urealyticum (5,10,19)<br>C. appendicis (5,10,15)<br>C. occultum (5,10)<br>C. urinipleomorphum (5,10,13,15,21)<br>C. xerosis (5,10)<br>C. pilosum (5,12,13,15,19,21)<br>C. halotolerans (5,14,19)<br>C. amycolatum (5,14)<br>C. vitaeruminis (5,14)<br>C. afermentans (5,14)<br>C. diphtheriae (5,6,7,8, 11,14,17,18)<br>C. rouxii (5,14)<br>C. callunae (5,19)<br>C. hindlerae (5,19)<br>M. tuerculosis (5,7,17)<br>M. leprae (5,7,17) | WP_249624942.1<br>WP_284789352.1<br>WP_145942353.1<br>WP_035002750.1<br>WP_066838567.1<br>WP_191733379.1<br>WP_006769007.1<br>WP_013887484.1<br>WP_053545750.1<br>WP_042622200.1<br>WP_018024932.1<br>WP_411276594.1<br>WP_392452223.1<br>WP_186276972.1<br>WP_265915152.1<br>WP_148791638.1<br>WP_284835239.1<br>WP_156231956.1<br>WP_087117409.1<br>WP_060925448.1<br>WP_018582272.1<br>WP_015401957.1<br>WP_366965759.1<br>WP_048760280.1<br>WP_194560614.1<br>WP_072574663.1<br>WP_342295824.1<br>WP_015652209.1<br>WP_182385302.1<br>WP_057342244.1<br>WP_010908932.1 | *C. simulans* 1 VGIDLGTTNSVVSVLEGGEATVIANSEGAR 30<br>+GIDLGTTNS V V+EGGE VIANSEGAR<br>*F. prausnitzii* 5 IGIDLGTTNSCVAVMEGGEPVVIANSEGAR 34<br><br>*C. simulans* 1 DAEAYLGEDVTDAVITVPAYFEDSQR 26<br>DAEAYLGE VTDAVITVPAYF DSQR<br>*F. prausnitzii* 102 DAEAYLGETVTDAVITVPAYFNDSQR 127<br><br>*C. simulans* 1 GVNPDEVVAVGAALQAGVL 19<br>G+NPDE VAVGAA Q GVL<br>*F. prausnitzii* 335 GINPDECVAVGAAIQGGVL 353<br><br>*C. simulans* 1 IQDGSGLSQEEIDR 14<br>IQ SGL EEI+R<br>*F. prausnitzii* 221 IQSNSGLTDEEIER 234<br><br>*C. simulans* 1 IVNEPTAAALAYGLEK 16<br>I+NEPTAAALAYG +K<br>*F. prausnitzii* 144 IINEPTAAALAYGVDK 159<br><br>*C. simulans* 1 RNTTIPTKKS 10<br>RNTTIPT KS<br>*F. prausnitzii* 134 RNTTIPTSKS 143<br><br>*Bacteroides fragilis* (WP_226776909.1)<br><br>*C. simulans* 1 DVLLLDVTPLSLGIET 16<br>D LLLDVTPLSLGIET<br>*B. fragilis* 77 DLLLLDVTPLSLGIET 92<br><br>*C. simulans* 1 VGIDLGTTNSVVSVLEGGEATVIANSEG 28<br>+GIDLGTTNS VSV EG E VIANSEG<br>*B. fragilis* 5 IGIDLGTTNSCVSVFEGNEPVVIANSEG 32<br><br>*C. simulans* 2 AEAYLGEDVTDAVITVPAYFEDSQR 26<br>AE YLG++VT+AVITVPAYF DSQR<br>*B. fragilis* 125 AEDYLGQEVTEAVITVPAYFSDSQR 149<br><br>*C. simulans* 1 GVNPDEVVAVGAALQAGVL 19<br>GVNPDEVVAVGAA Q VL<br>*B. fragilis* 358 GVNPDEVVAVGAAVQGAVL 376<br><br>*C. simulans* 1 IVNEPTAAALAYGLEK 16<br>IVNEPTAAALAYGL+K<br>*B. fragilis* 166 IVNEPTAAALAYGLDK 181 | | | |
| 2 | Co-chaperone GroES | C. guaraldiae<br>C. singulare<br>C. aurimucosum<br>C. simulans<br>C. minutissimum<br>C. striatum | WP_284431894.1<br>AJI78083.1<br>ACP32071.1<br>AMO88637.1<br>STC74814.1<br>HCG2963200.1 | ANIKPLEDK<br>DLLAVIEK<br>VLVQIVEAETTTASGLVIPDSAK<br>YDGQEYLLLSAR | 2-10<br>90-97<br>11-33<br>78-89 | 1<br>2<br>3<br>4 | 10328 |
| 3 | ATP-dependent chaperone ClpB | C. lizhenjunii<br>C. guaraldiae<br>C. hiratae<br>C. striatum (1-4)<br>C. aurimucosum<br>C. simulans (1-4)<br>C. accolens<br>C. minutissimum<br>C. phocae<br>C. endometrii<br>C. flavescens<br>C. phoceense | WP_165007392.1<br>WP_144269183.1<br>WP_144014045.1<br>WP_086890648.1<br>WP_010188684.1<br>WP_284841445.1<br>PCC82406.1<br>WP_115023765.1<br>WP_075735801.1<br>WP_136141911.1<br>WP_301523703.1<br>WP_257036007.1 | ALADFLFDDESAMVR<br>AQELAGELGDEYVSTEVLLAAIAGSK<br>NNPVLIGEPGVGK<br>TKNNPVLIGEPGVGK | 620-634<br>95-120<br>201-213<br>199-213 | 1<br>2<br>3<br>4 | 92966 |

| | | | | | | | |
|---|---|---|---|---|---|---|---|
| | | C. diphtheria (4)<br>M. tuberculosis (4) | WP_071572341.1<br>WP_078449720.1 | | | | |
| 4 | Chaperonin GroEL | C. simulans<br>C. striatum | WP_284841047.1<br>WP_049146941.1 | FAEEFEGEAK<br>IAESSRPTLIIAEDVEGEPLQALVVNSIR<br>ISSLPDFLPLLEK<br>TNDIAGDGTTTATLLAQALIFEGLR<br>VEDAINAAR<br>VGAATETEVNER | 427-436<br>238-266<br>225-237<br>80-104<br>394-402<br>379-390 | 1<br>2<br>3<br>4<br>5<br>6 | 57012 |
| 5 | Chaperonin GroEL | C. confusum<br>C. striatum (1-4)<br>C. simulans (1-4)<br>C. auris<br>C. diphtheria (4) | WP_290223349.1<br>WP_086890684.1<br>ABI75068.1<br>ABK15698.1 | AVVDSLLSSAK<br>GLNTLADAVK<br>IIAFDEEAR<br>NVAAGSNPMGIK | 125-135<br>18-27<br>4-12<br>105-116 | 1<br>2<br>3<br>4 | 57342 |
| 6 | Chaperonin GroEL | C. striatum<br>C. flavescens<br>C. halotolerans<br>C. glutamicum<br>C. aurimucosum | WP_284765038.1<br>WP_075730587.1<br>WP_015401886.1<br>WP_074492415.1<br>WP_010188783.1 | EIELEDPYEK | 58-67 | 1 | 57304 |
| 7 | Chaperonin GroEL | C. flavescens<br>C. incognita<br>C. camporealensis<br>C. confusum<br>C. singulare<br>C. striatum<br>C. tuberculostearicum<br>C. aurimucosum<br>C. halotolerans<br>C. minutissimum<br>C. kroppenstedtii<br>C. accolens<br>M. tuberculosis | WP_075730587.1<br>WP_185176557.1<br>WP_046453551.1<br>WP_290223349.1<br>WP_042532405.1<br>WP_086890684.1<br>WP_200435714.1<br>WP_049360282.1<br>WP_015401886.1<br>WP_115023692.1<br>WP_303734259.1<br>WP_302522226.1<br>WP_055378132.1 | QEAVLEDPYILLVSSK | 214-229 | 1 | 56997 |
| 8 | Molecular chaperone DnaJ | C. argentoratense | WP_314929504.1 | FREISLAQEVLSDPNKR | 49-65 | 1 | 40313 |
| 9 | Copper chaperone PCu(A)C | C. jeikeium | WP_172457282.1 | MSLTSLNNSK | 1-10 | 1 | 23325 |
| 10 | Molecular chaperone DnaJ | C. urealyticum | WP_149121529.1 | DLGVSDSASAEEIK | 13-26 | 1 | 41991 |
| 11 | Copper chaperone PCu(A)C | C. ulcerans | WP_197906745.1 | GVELADGYVR | 58-67 | 1 | 20781 |
