## Supplemental Table S20 for "Peptidome Analysis of Western Blots Identifies Natural Bispecific Antibody-Bound *Corynebacterium* and Phage B-cell Epitopes with Potential Relevance to Psoriasis"

### Table S20. Stress Proteins as Corynebacterium Antigens.

| Antigen Number | Identified Antigen | *Corynebacterium* (C.) | Accession Number | Peptide Sequence | Peptide Start/Stop | Molecular Weight |
|---|---|---|---|---|---|---|
| 1 | 50S ribosomal protein L25/general stress protein Ctc | C. aurimucosum<br>C. simulans<br>C. accolens<br>C. striatum | WP_010187470.1<br>WP_062035544.1<br>PCC83094.1<br>WP_347004832.1 | ANTRPVLK<br>IELTALVR | 2-9<br>49-56 | 23370 |
| 2 | Asp23/Gls24 family envelope stress response protein | C. tuberculostearicum<br>C. curieae<br>C. minutissimum<br>C. camporealensis<br>C. guaraldiae<br>C. aurimucosum<br>C. simulans<br>C. striatum | WP_061920488.1<br>WP_269946445.1<br>WP_039673696.1<br>WP_035106528.1<br>WP_154736259.1<br>WP_010189665.1<br>WP_248093770.1<br>WP_284772775.1 | ESFGASEDVR<br>NIMNAVER<br>VDVVVHDVK | 84-93<br>129-136<br>145-153 | 17298 |
| 3 | Universal stress protein | C. diphtheriae<br>C. marinum<br>C. amycolatum<br>C. genitalium<br>C. curieae<br>C. lactis<br>C. camporealensis<br>C. tuberculostearicum<br>C. cystitidis<br>C. mastitidis<br>C. faecale<br>C. gallinarum<br>C. marquesiae<br>C. massiliense<br>C. kefirresidentii<br>C. efficiens<br>C. lizhenjunii<br>C. incognita<br>C. gerontici<br>C. glutamicum<br>C. deserti<br>C. callunae<br>C. belfantii | WP_088262747.1<br>WP_042621336.1<br>WP_256884568.1<br>EFK54051.1<br>WP_269947133.1<br>WP_053413336.1<br>WP_035105547.1<br>WP_316993611.1<br>WP_257159965.1<br>WP_337890207.1<br>WP_290279691.1<br>WP_191734579.1<br>WP_049378698.1<br>WP_022863676.1<br>WP_284839589.1<br>WP_006770406.1<br>WP_165010895.1<br>WP_185175862.1<br>WP_123933843.1<br>WP_074494080.1<br>WP_053544775.1<br>WP_015651143.1<br>WP_088267786.1 | GINSLTGR<br>IVVGTDGSK<br>LLGSVPADVAR<br>QSDCDVMIVHTVS<br>SSLLAVER<br>IVVGTDGSKSSLLAVER | 116-123<br>8-16<br>124-134<br>135-147<br>17-24<br>8-24 | 15419 |
| 4 | Universal stress protein | C. casei<br>C. accolens<br>C. tuberculostearicum<br>C. marquesiae<br>C. macginleyi<br>C. aurimucosum<br>C. simulans<br>C. kefirresidentii<br>C. striatum | WP_006822482.1<br>WP_284643885.1<br>WP_301432587.1<br>WP_284837185.1<br>WP_121912182.1<br>WP_049359849.1<br>WP_062043029.1<br>WP_086588580.1<br>WP_086890566.1 | YGPVVVGVDGSEVSQK | 154-169 | 32983 |

| | | | | | | |
|---|---|---|---|---|---|---|
| | | C. propinquum<br>C. uropygiale<br>C. lizhenjunii<br>C. phoceense<br>C. pseudodiphtheriticum<br>C. minutissimum<br>C. confusum | WP_302516869.1<br>WP_236119528.1<br>WP_165010950.1<br>WP_257037028.1<br>WP_284873609.1<br>WP_115023988.1<br>WP_290223701.1 | | | |
| 5 | Universal stress protein | C. efficiens<br>C. faecale<br>C. ulcerans<br>C. vitaeruminis<br>C. efficiens<br>C. faecale<br>C. durum<br>C. diphtheriae<br>C. rouxii<br>C. belfantii<br>C. marinum<br>C. phoceense | WP_006768841.1<br>WP_290277424.1<br>WP_368266799.1<br>WP_025253847.1<br>WP_006768841.1<br>WP_290277424.1<br>WP_006062182.1<br>WP_014302568.1<br>WP_155874385.1<br>WP_196977407.1<br>WP_042622281.1<br>WP_257037028.1 | AIAHEVAPDIK | 75-85 | 33974 |
| 6 | Universal stress protein | C. striatum | WP_049147615.1 | IVVGTDGSK<br>KIVVGTDGSK<br>LLGSVPADVAR | 7-15<br>6-15<br>123-133 | 15203 |
