## Supplemental Table S21 for "Peptidome Analysis of Western Blots Identifies Natural Bispecific Antibody-Bound *Corynebacterium* and Phage B-cell Epitopes with Potential Relevance to Psoriasis"

### Table S21. Metalloproteins are *Corynebacterium* Antigens.

| Antigen Number | Identified Antigen | Corynebacterium (C.), Actinomycetota | Accession Number | Peptide Sequence | Peptide Start/Stop | Molecular Weight |
|---|---|---|---|---|---|---|
| 1 | Metal ABC transporter substrate-binding protein | C. bovis<br>C. urealyticum | WP_280066670.1<br>AGE36660.1 | MSPRTAR | 174-180 | 28962 |
| 2 | Metalloregulator ArsR/SmtB family transcription factor | C. glutamicum | WP_038583602.1 | ALSIGTQWAHVMGIK | 126-140 | 25246 |
| 3 | M20 family metallo-hydrolase | C. simulans<br>C. striatum | WP_062035706.1<br>WP_420697603.1 | ATTDEQYLADFR<br>EMSANGATAGGGVER<br>FNIDLR<br>QAASEGDRVNR<br>SPNSETLETAQK<br>SPNSETLETAQKR<br>VVSYGEIHVEQGK<br>WTPVEQAPK<br>YDGAYGVLAAAHAANR<br>YNLAVINWFNEEGSR | 3-14<br>15-29<br>283-288<br>30-40<br>289-300<br>289-301<br>182-194<br>173-181<br>90-105<br>119-133 | 45376 |
| 4 | M3 family metallopeptidase | C. simulans<br>C. striatum | WP_284841026.1<br>WP_086891990.1 | ARDFSLDEEELR<br>DFSLDEEELR<br>AGLNNPLIAPR<br>DWVEFPSQINENWALEPALVK | 322-333<br>324-333<br>581-591<br>489-509 | 54945 |
| 5 | M20/M25/M40 family metallo-hydrolase | C. striatum<br>C. simulans | WP_049062472.1<br>WP_248092878.1 | AYDSLVTR<br>EALGDLADEVEIEHLITEDATQSSTDTK<br>IAAAEPPVASNEIWEGFVK<br>IAQEGNPGGTLAFVGMADEEAR<br>LWHAIEATSK<br>NACVNDLTPDSGNEVR<br>TFKFDPETEAALIAGTGDYSK | 422-429<br>319-346<br>223-241<br>125-146<br>347-356<br>17-32<br>242-262 | 47232 |
| 6 | M13-type metalloendopeptidase | C. simulans<br>C. striatum | WP_284841565.1<br>WP_110333686.1 | AEMDTLVDYLTAAYR<br>AGNAWAHDFELR<br>AVGLAESLVGQEIGK<br>IGYPDKWR<br>ISQLEWMTPATR<br>NFEFYGTR<br>VFVDQHFPPASK | 317-331<br>386-397<br>290-304<br>359-366<br>334-345<br>269-276<br>305-316 | 71583 |
| 7 | M3 family metallopeptidase | C. striatum<br>C. simulans | WP_420692589.1<br>WP_239239679.1 | ARDFSLDEEELR<br>DFSLDEEELR | 322-333<br>324-333 | 54945 |
| 8 | Neutral zinc metallopeptidase | C. striatum<br>C. simulans | WP_284765519.1<br>WP_239238688.1 | AVGDDNIQR | 244-252 | 33782 |
| 9 | MBL fold metallo-hydrolase | C. striatum<br>C. simulans | WP_096336195.1<br>WP_062036106.1 | VLPGHGDETTVGAEAAR | 181-197 | 22870 |
| 10 | M13-type metalloendopeptidase | C. simulans<br>C. striatum | WP_284841565.1<br>WP_244080034.1 | ELAAGHWDVVSTR<br>ISQLEWMTPATR | 181-193<br>337-348 | 71398 |
| 11 | M20/M25/M40 family metallo-hydrolase | C. simulans<br>C. striatum<br>C. pseudogenitalium | WP_061919913.1<br>WP_306588944.1<br>WP_005326315.1 | NCLSETGGSHLPDAVGFNVGEK | 166-187 | 47739 |

| | | | | | | |
|---|---|---|---|---|---|---|
| 12 | MBL fold metallo-hydrolase | C. minutissimum<br>C. aurimucosum<br>C. striatum<br>C. simulans | WP_039673253.1<br>WP_010189963.1<br>PIS59825.1<br>WP_248093213.1 | QLTGAPIR<br>TLDMLEER<br>TLVDEMYDDVDPVLR | 89-96<br>181-188<br>238-252 | 29608 |
| 13 | Metalloregulator ArsR/SmtB family transcription factor | C. glutamicum | WP_038583602.1 | ALSIGTQWAHVMGIK | 126-140 | 25246 |
| 14 | M3 family metallopeptidase | C. striatum<br>C. simulans | WP_086891990.1<br>WP_284841026.1 | AGLNNPLIAPR | 581-591 | 76034 |
| 15 | MBL fold metallo-hydrolase | C. simulans<br>C. striatum | WP_005531203.1<br>WP_096336195.1<br>WP_046645696.1 | IDEYVAR<br>KYSDFDVIIESLR<br>VLPGHGDETTVGAEAAR<br>YSDFDVIIESLR | 198-204<br>157-169<br>181-197<br>158-169 | 22248 |
| 16 | MBL fold metallo-hydrolase | C. simulans<br>C. striatum<br>C. pseudodiphthereticum<br>C. propinquum | WP_248092006.1<br>MDU3175469.1<br>WP_249618692.1<br>WP_302501683.1 | TDSEGDFVR | 148-156 | 22037 |
| 17 | MBL fold metallo-hydrolase | C. striatum<br>C. simulans | WP_239299701.1<br>WP_062040529.1 | SISTTVTGIEMFR | 205-214 | 44104 |
| 18 | Nif3-like dinuclear metal center hexameric protein | C. simulans<br>C. ulcerans<br>C. simulans | WP_306588200.1<br>WP_095075588.1<br>WP_239240756.1 | GVTSVAADTPK | 76-86 | 40198 |
| 19 | Metal-dependent transcriptional regulator | C. simulans<br>C. aurimucosum | WP_239238568.1<br>WP_049361228.1 | DGLLHVR<br>LEQSGPTVSQTVAR<br>TIYELEEEGITPLR | 66-72<br>49-62<br>29-42 | 24975 |
| 20 | MBL fold metallo-hydrolase | C. striatum<br>C. simulans | WP_306592394.1<br>WP_248093213.1 | TLDMLEER<br>TLVDEMYDDVDPVLR | 181-188<br>238-252 | 29608 |
| 21 | Zinc-dependent metalloprotease | Tsukamurella paurometabola | WP_126196731.1 | LVNPVAEK | 136-143 | 48395 |
| 22 | Metal-sensitive transcriptional regulator | C. urealyticum | WP_012360560.1 | MDNAKPATK | 1-9 | 13375 |
| 23 | TIGR04338 family metallohydrolase | Tsukamurella paurometabola | WP_013125427.1 | VMGPEAGLALR | 139-149 | 16954 |
| 24 | M23 family metallopeptidase | Corynebacterium sp. CCUG 65737 | WP_256003305.1 | SMKPAEGTFTSGFGMR | 111-126 | 24645 |
| 25 | Peptidoglycan DD-metalloendopeptidase family protein | C. callunae | WP_015650630.1 | QELFPANGSPYMMEDR | 296-311 | 55737 |
| 26 | TIGR04338 family metallohydrolase | Tsukamurella paurometabola | WP_013125427.1 | VMGPEAGLALR | 139-149 | 16954 |
| 27 | Molybdopterin molybdotransferase MoeA | C. simulans<br>C. striatum | WP_248092913.1<br>WP_244080041.1 | SDLEGVVR<br>VPEGTVTIVPVEQCEPPQFPR | 224-231<br>93-113 | 39689 |
| 28 | MogA/MoaB family molybdenum cofactor biosynthesis protein | C. striatum<br>C. simulans | WP_086892451.1<br>MCK6160106.1 | SSGQACGAVDACTSR | 137-151 | 20401 |
| 29 | Molybdenum cofactor biosynthesis protein MoaE | C. vitaeruminis | WP_025251633.1 | EVAARVSAAHPK | 242-253 | 32567 |

| | | | | | | |
|---|---|---|---|---|---|---|
| 30 | Molybdopterin-binding protein | C. striatum<br>C. simulans | WP_166684653.1<br>WP_062035328.1 | GCGSLIINSASSR | 124-136 | 13644 |
| 31 | MogA/MoaB family molybdenum cofactor biosynthesis protein | C. striatum | WP_100087247.1 | SVLDQMVPGIAQALR | 128-142 | 21043 |
| 32 | Siderophore-interacting protein | C. accolens<br>C. pseudogenitalium<br>C. parakroppenstedtii | WP_284624419.1<br>MCQ4618067.1<br>WP_221927240.1 | YFDAVR | 106-111 | 68296 |
| 33 | Fe-S cluster assembly protein SufB | C. amycolatum<br>C. diphtheria<br>C. deserti<br>C. efficiens<br>C. glutamicum<br>C. pseudotuberculosis<br>C. simulans<br>C. striatum | WP_005510784.1<br>WP_014308346.1<br>WP_053544971.1<br>WP_006767687.1<br>WP_060564611.1<br>WP_013241949.1<br>WP_062035089.1<br>WP_005531592.1 | GLAEEEAMAMIVR<br>INTENMGQFER | 438-450<br>220-230 | 53863 |
| 34 | Fe-S cluster assembly protein SufD | C. accolens<br>C. tuberculostearicum | WP_284894033.1<br>WP_316980693.1 | FDDIELFYLMSR | 328-339 | 42822 |
| 35 | (Fe-S)-binding protein | C. striatum<br>C. flavescens<br>C. simulans | WP_141276823.1<br>WP_075729606.1<br>WP_248090681.1 | NPEVSQAMVSDK<br>ATALILSR<br>TDLGAFFPHK<br>TLDLPEFLVDIEQR<br>VLHMAEILASTK | 186-197<br>21-28<br>126-135<br>112-125<br>233-244 | 28574 |
| 36 | (Fe-S)-binding protein | C. simulans<br>C. guaraldiae<br>C. singulare | WP_248090681.1<br>WP_101736598.1<br>WP_042530427.1 | ATALILSR<br>FGTSADVDGAK | 21-28<br>96-106 | 28150 |
| 37 | FAD-binding and (Fe-S)-binding domain-containing protein | C. efficiens<br>C. gallinarum | WP_006768226.1<br>WP_191732322.1 | SEEVHEALDLCLSCKACASECPVNVDM ATYK | 597-627 | 102922 |
| 38 | Ubiquinol-cytochrome c reductase iron-sulfur subunit | C. striatum | WP_166684805.1 | ALPQLPIGVDEEGYLVAK<br>GNFIEPVGPAFWER<br>ICTHIGCPTSLYEAQTNR<br>MSNDELAALGTELDDVTVAFR<br>NAVMLIR<br>TLTALLNDSWK | 373-390<br>391-404<br>330-347<br>16-36<br>295-301<br>142-152 | 45034 |
| 39 | LutB/LldF family L-lactate oxidation iron-sulfur protein | C. striatum | WP_284786984.1 | DAGSAIKEDVAAR<br>TAALSSPIGHQALK | 68-80<br>314-327 | 55253 |
| 40 | 16S rRNA (cytosine(1402)-N(4))-methyltransferase RsmH<br>*iron ABC transporter permease* | C. otitides<br><br><br>*C. casei* | WP_046644458.1<br><br><br>*WP_006822721.1* | LAHFGGR | 145-151 | 35688 |
| 41 | Iron ABC transporter permease | C. efficiens<br>C. gallinarum | WP_011074987.1<br>WP_191732788.1 | ISGAGTVTVLR | 166-176 | 56432 |
| 42 | Iron-sulfur cluster insertion protein ErpA | Clavibacter michiganensis | OUE30845.1 | VKSLLQQEGREDLRLR | 26-41 | 12935 |
| 43 | Iron ABC transporter | Tsukamurella | WP_013128332.1 | GLGLNVNLSR | 258-267 | 37272 |

| | | | | | | |
|---|---|---|---|---|---|---|
| | permease | paurometabola | | | | |
| 44 | L-serine ammonia-lyase, iron-sulfur-dependent, subunit alpha, partial | C. falsenii | WP_272712594.1 | RDHDSEAMPGAK | 203-214 | 48405 |
| 45 | MULTISPECIES: succinate dehydrogenase iron-sulfur subunit | Micrococcus<br>Brevibacterium<br>Clavibacter michiganensis<br>M. tuberculosis<br>M. leprae | WP_224355525.1<br>WP_069600613.1<br>WP_104294766.1<br><br>MBZ4296222.1<br>WP_010907879.1 | SCAHGVCGSDAMRINGRNR | 83-101<br><br><br><br>54-72<br>53-71 | 26417 |
| 46 | Heterodisulfide reductase-related iron-sulfur binding cluster, partial | C. accolens | WP_139027181.1 | MESMTIHPTCSAFQLGMMGDVEK | 2-23 | 11512 |
| 47 | Cytochrome bc complex cytochrome b subunit | C. simulans<br>C. halotolerans<br>C. renale<br>C. efficiens<br>C. kutscheri<br>C. diphtheria<br>C. striatum<br>C. marquesiae<br>C. stationis<br>C. casei<br>C. aurimucosum<br>C. pseudotuberculosis<br>C. matruchotii<br>C. lizhenjunii<br>C. minutissimum | WP_062043913.1<br>WP_027004078.1<br>WP_048379400.1<br>WP_011075748.1<br>WP_046439943.1<br>WP_010935204.1<br>WP_306592346.1<br>WP_337888180.1<br>WP_313678661.1<br>WP_276686157.1<br>WP_049361052.1<br>WP_160186677.1<br>WP_315555662.1<br>WP_165010261.1<br>WP_115022928.1 | AENNVVGVR<br>EVLEHGIETGVIK<br>YSAAGVLR | 239-247<br>434-446<br>16-23 | 60093 |
| 48 | Ferredoxin | C. striatum | WP_420684287.1 | ACVEECPVDCIYEGK | 16-30 | 11957 |
| 49 | Multicopper oxidase domain-containing protein | C. pseudotuberculosis | WP_145951886.1 | GLAGMILLEDEVSK | 179-192 | 60173 |
| 50 | Ferritin | C. halotolerans | WP_034991252.1 | LVGDDGSGLLR | 144-154 | 19301 |
| 51 | Ferredoxin-NADP(+) reductase FprA | M. tuberculosis<br>M. leprae | WP_057356393.1<br>WP_010908349.1 | SLTELLR | 446-452<br>276-280 | |
| 52 | Thioredoxin domain-containing protein | C. aurimucosum<br>C. guaraldiae | WP_236659339.1<br>WP_143336249.1 | FSDGKGDPMMFGPGGDTK | 82-99 | 30701 |
| 53 | Multicopper oxidase family protein | C. jeikeium<br>C. accolens<br>C. sanguinis<br>C. kefirresidentii<br>C. macclintockii | WP_005293220.1<br>WP_284832117.1<br>WP_144823711.1<br>WP_284798826.1<br>WP_284821195.1 | LTGMGSGDHNGKKGMGGHGQMMHG SPDR | 204-231 | 53129 |
