## Supplemental Table S22 for "Peptidome Analysis of Western Blots Identifies Natural Bispecific Antibody-Bound *Corynebacterium* and Phage B-cell Epitopes with Potential Relevance to Psoriasis"

**Table S22. Surface-Associated Proteins are Antigens of Numerous *Corynebacterium* Species and *Prescottella equi*.**

| Antigen Number | Identified Antigen | *Corynebacterium* (C.), *Prescottella equi* | Accession Number | Peptide Sequence | Peptide Start/Stop | Molecular Weight |
|---|---|---|---|---|---|---|
| 1 | Type VII secretion target | Prescottella equi | WP_005514188.1 | ALLSDNAAAIGR | 59-70 | 10354 |
| 2 | WXG100 family type VII secretion target | C. variabile | WP_313358398.1 | MSFRTTTDVMHATAGK | 1-16 | 10324 |
| 3 | Type VII secretion protein EccCa | C. pseudodiphthereticum<br>C. propinquum | WP_284814275.1<br>WP_284589985.1 | LAESMDSLCDRKMQDK | 889-904 | 145894 |
| 4 | DoxX family membrane protein | C. striatum | HAT6550504.1 | EYVDDNKDDWQATGK<br>VGAGSTLALGK | 195-209<br>71-81 | 33955 |
| 5 | DoxX family membrane protein | C. tuberculostearicum<br>C. aurimucosum<br>C. kefirresidentii<br>C. curiae<br>C. marquesiae | WP_316993610.1<br>WP_049358452.1<br>WP_248100364.1<br>WP_269947130.1<br>WP_284791282.1 | SPAELDAMQAAGEIVGK | 246-256 | 37084 |
| 6 | Putative membrane protein | C. kroppenstedtii | ACR17060.1 | MNACRVNAMSRAR | 1-13 | 22703 |
| 7 | Energy-coupling factor transporter transmembrane protein EcfT | C. kroppenstedtii<br>C. parakroppenstedtii | WP_012731751.1<br>WP_236916289.1 | GFDDWDPTAPEYDSQE | 220-235 | 25824 |
| 8 | Putative membrane protein | C. efficiens | BAC17010.1 | RTECAHQYGQR | 83-93 | 96742 |
| 9 | Murein biosynthesis integral membrane protein MurJ | C. falsenii | WP_224209198.1 | ESASMSGSMMGTALPPLSAGR | 683-703 | 131696 |
| 10 | Division/cell wall cluster transcriptional repressor MraZ | C. mastitides<br>C. macginleyi<br>C. confusum<br>C. accolens<br>C. maris<br>C. flavescens<br>C. massiliense<br>C. otitides<br>C. diphtheria<br>C. striatum | WP_018117527.1<br>WP_200446021.1<br>WP_290222463.1<br>WP_337886807.1<br>WP_020935172.1<br>WP_276622580.1<br>WP_027018511.1<br>WP_004600154.1<br>WP_071571590.1<br>WP_086891054.1 | GQDHSLAVYPR<br>KAAAVSR<br>NLAASADEQRPDGHGR | 42-52<br>61-67<br>78-93 | 12381 |
| 11 | LPXTG cell wall anchor domain-containing protein | C. striatum<br>C. diphtheria | WP_082257764.1<br>WP_115598112.1 | QIGEELASR | 721-731 | 111118 |
| 12 | Polysaccharide biosynthesis protein | C. amycolatum | WP_005510927.1 | WITDNFGTRQTDDR | 131-144 | 67749 |
| 13 | SpaH/EbpB family LPXTG-anchored major pilin | Corynebacterium sp. KPL1818 | WP_023030358.1 | WKVSDQAAPIK | 389-399 | 54642 |
| 14 | SpaH/EbpB family LPXTG-anchored major pilin | C. diphtheriae | WP_014301239.1 | TYWGNLNFKK | 347-356 | 56727 |
| 15 | Surface-anchored protein, fimbrial subunit | C. ulcerans<br>C. ramonii | WP_029974243.1<br>WP_038620833.1 | ITSCTFSNVPK | 398-408 | 201614 |
| 16 | Fimbrial associated class C sortase | C. striatum<br>C. diphtheria<br>C. urealyticum<br>C. marquesiae<br>C. tuberculosrearicum | WP_306496410.1<br>WP_014301240.1<br>WP_148811354.1<br>WP_284837112.1<br>WP_316986319.1 | DHTLSTDVEPK | 2-12 | 35040 |
| 18 | TIGR03773 family transporter-associated surface protein | C. diphtheriae | WP_196977509.1 | DGIADTEGGIAEFK | 510-523 | 117348 |
| 19 | 1,4-Alpha-glucan branching enzyme GlgB | C. matruchotii<br>C. diphtheriae | WP_311323162.1<br>WP_181998757.1 | TLYAFMYAHPGK | 524-535 | 87220 |
| 20 | Alpha-1,4-glucan:maltose-1-phosphate maltosyltransferase | C. aurimucosum | WJY70269.1 | RENPALQQLR | 548-557 | 75124 |

| | | | | | | |
|---|---|---|---|---|---|---|
| | | C. simulans<br>C. striatum<br>C. accolens | WP_239238856.1<br>MDU3175173.1<br>PCC81918.1 | TNPEFFTVLADGTIAYAENPPK<br>IDAWSDPMATWR | 319-340<br>92-103 | |
| 21 | Glycogen/starch/alpha-glucan phosphorylase | C. striatum<br>C. simulans<br>C. accolens | WP_201816736.1<br>WP_284841150.1<br>PCC82120.1 | ALLNNLTNLDLVDEAK | 74-89 | 90367 |
