## Supplemental Table S23 for "Peptidome Analysis of Western Blots Identifies Natural Bispecific Antibody-Bound *Corynebacterium* and Phage B-cell Epitopes with Potential Relevance to Psoriasis"

### Table S23. Proteases and Peptidases are *Corynebacterium* Antigens.

| Antigen Number | Identified Antigen | *Corynebacterium* (C.), *Tsukamurella paurometabola* (Peptide numbers) | Accession Number | Peptide Number | Peptide Sequence | Peptide Start/Stop | Molecular Weight |
|---|---|---|---|---|---|---|---|
| 1 | ATP-dependent Clp | C. minutissimum (1,12,18) | WP_039676609.1 | 1 | AAGLRDQER | 464-472 | 96101 |
| | protease ATP-binding | C. kroppenstedtii (1,18) | KXB50087.1 | 2 | DGKLDPVVGR | 201-210 | |
| | subunit ClpC | C. simulans (1,2,3,4,6,7,9,10,11,12,14,15,18) | WP_062043288.1 | 3 | EIYNTLLQVLEDGR | 650-663 | |
| | | C. pseudokroppenstedtii (1,18) | WP_204088626.1 | 4 | IIGQDDAVK | 535-543 | |
| | | C. singulare (1,2,3,6,9,10,11,12,14,18) | WP_239181074.1 | 5 | NNPVLIGEPGVGK | 227-239 | |
| | | C. uropygiale (1,6) | WP_236119313.1 | 6 | QLYSLDLGSLVAGSR | 263-277 | |
| | | C. parakroppenstedtii (1) | WP_039676609.1 | 7 | SNSLVLDQFGR | 183-193 | |
| | | C. uterequi (1) | WP_047260185.1 | 8 | TKNNPVLIGEPGVGK | 225-239 | |
| | | C. matruchotii (1) | WP_047260185.1 | 9 | ALESMGISLEDVR | 48-60 | |
| | | C. epididermidicanis (1) | WP_047240896.1 | 10 | DGKLDPVVGR | 202-211 | |
| | | C. durum (1,2,10,18) | WP_303130074.1 | 11 | EIYNTLLQVLEDGR | 651-664 | |
| | | C. mastitidis (1,2,10) | WP_337889463.1 | 12 | FQPVKVDEPSLDDTILILK | 360-378 | |
| | | C. accolens (2,3,4,5,6,7,8,9,10,11,13,14,15-18) | WP_249852940.1 | 13 | NNPVLIGEPGVGK | 228-240 | |
| | | C. macginleyi (2,3,6,7,9,10,11,14,15,18) | WP_200438993.1 | 14 | QLYSLDLGSLVAGSR | 264-278 | |
| | | C. anserum (2,3,4,6,10,11,14) | WP_185769461.1 | 15 | SNSLVLDQFGR | 184-194 | |
| | | C. deserti (2,3,4,10,11) | WP_185769461.1 | 16 | TKNNPVLIGEPGVGK | 226-240 | |
| | | C. urogenitale (2,3,6,10,11,14) | WP_151902224.1 | 17 | SGDLEEIAEVGEDQIAEVLAHWTGIPVLK | 490-518 | |
| | | C. frankenforstense (2,6,10,14) | WP_304348540.1 | 18 | VVVLAQEEAR | 12-21 | |
| | | C. callunae (2,3,4,10,11) | WP_239181074.1 | | | | |
| | | C. faecale (2,3,10,11) | WP_290276944.1 | | | | |
| | | C. gallinarum (2,3,10,11) | WP_191733296.1 | | | | |
| | | C. efficiens (2,3,10,11) | WP_006769109.1 | | | | |
| | | C. maris (2,10) | WP_020935640.1 | | | | |
| | | C. aurimucosum (2,3,4,6,9,10,11,12,14) | WP_102233533.1 | | | | |
| | | C. guaraldiae (2,3,6,9,10,11,14) | WP_154736951.1 | | | | |
| | | C. glutamicum (2,3,10,11) | WP_143855236.1 | | | | |
| | | C. striatum (2,3,5,6,7,8,11,12,13,14,15,16,18) | WP_086890713.1 | | | | |
| | | C. endometrii (3,11) | WP_136141840.1 | | | | |
| | | C. urinipleomorphum (4,5,7,8,13,15,16) | WP_087117502.1 | | | | |
| | | C. genitalium (4,5,7,8,13,15,16) | WP_005288548.1 | | | | |
| | | C. appendicis (4) | WP_259798723.1 | | | | |
| | | C. diphtheriae (4,5,7,8,13,15,16,18) | WP_071572341.1 | | | | |
| | | C. cystitidis (4) | WP_257183114.1 | | | | |
| | | C. pilosum (4) | WP_018582802.1 | | | | |
| | | C. testudinoris (4) | WP_047253839.1 | | | | |
| | | C. halotolerans (4) | WP_034990906.1 | | | | |
| | | C. urealyticum (4) | WP_317209301.1 | | | | |
| | | C. argentoratense (4) | WP_234859003.1 | | | | |
| | | C. vitaeruminis (3,4) | WP_234859003.1 | | | | |
| | | C. dentalis (4) | WP_312098022.1 | | | | |
| | | C. propinquum (5,7,8,13,15,16) | WP_201809330.1 | | | | |
| | | C. pseudodiphtheriticum (5,7,8,13,15,16,18) | WP_239264435.1 | | | | |
| | | C. rigelii (5,7,8,13) | WP_201821606.1 | | | | |

| | | | | | | | |
|---|---|---|---|---|---|---|---|
| | | C. atypicum (6,14)<br>C. variabile (6,14)<br>C. tuberculostearicum (9,17,18)<br>C. marquesiae (9,17)<br>C. faecipullorum (12)<br>M. tuberculosis (18)<br>M. leprae (18) | WP_084168391.1<br>CUU66855.1<br>WP_316971272.1<br>WP_408928680.1<br>HIX79148.1<br>WP_057139398.1<br>WP_010907619.1 | | | | |
| 2 | ATP-dependent Clp protease proteolytic subunit | C. simulans<br>C. striatum<br>C. endometrii | WP_062037803.1<br>WP_086890864.1<br>WP_136142233.1 | 1 | ILTAQEAVEYGIIDTVFDYR | 184-203 | 22862 |
| 3 | ATP-dependent Clp protease proteolytic subunit | C. striatum<br>C. aurimucosum<br>C. minutissimum<br>C. simulans<br>C. urealyticum | WP_204083932.1<br>WP_046648966.1<br>WP_239187657.1<br>WP_062037803.1<br>PZO98097.1 | 1 | VLIHQPR | 129-135 | 23015 |
| 4 | ATP-dependent Clp protease proteolytic subunit | C. simulans (1,2,4)<br>C. striatum (1,2,4)<br>C. gerontici (2,4)<br>C. glyciniphilum (2,3)<br>C. tuberculostearicum (2,3,4)<br>C. macginleyi (2,3,4)<br>C. terpenotabidum (2,4)<br>C. cystitidis (2,4)<br>C. propinquum (2,3)<br>C. faecipiullorum (2,4)<br>C. guaraldiae (2,3,4)<br>C. gallinarum (2,4)<br>C. diphtheriae (2,3)<br>C. incognita (2,3,4)<br>C. camporealensis (2,3)<br>C. humireducens (2)<br>C. flavescens (2)<br>C. pseudotuberculosis (2,3,4)<br>C. faecale (2,4)<br>C. marquesiae (2,3)<br>C. marinum (2,4)<br>C. comes (2,4)<br>C. hiratae (2,4)<br>C. belfantii (2,3)<br>C. pollutisoli (2,4)<br>C. occultum (2,4)<br>C. kutscheri (2,3)<br>C. xerosis (4)<br>C. bovis (4)<br>C. kroppenstedtii (4)<br>C. pseudokroppenstedtii (4)<br>C. variabilie (3,4)<br>C. breve (4)<br>C. nasicanis (4) | WP_061924162.1<br>WP_284773362.1<br>WP_123934945.1<br>WP_038549629.1<br>WP_204611150.1<br>WP_121927521.1<br>WP_020440712.1<br>WP_092257630.1<br>WP_018120934.1<br>HIX79529.1<br>WP_144014842.1<br>MBD8029173.1<br>WP_014303753.1<br>WP_185176416.1<br>WP_035105865.1<br>WP_040086552.1<br>MDN6460641.1<br>WP_013242460.1<br>WP_290276649.1<br>WP_284836892.1<br>WP_042621962.1<br>WP_156228794.1<br>WP_158395780.1<br>WP_197692036.1<br>NLP39358.1<br>WP_156231499.1<br>WP_046439005.1<br>WP_368522543.1<br>WP_425279042.1<br>WP_303735480.1<br>WP_337447800.1<br>WP_174775631.1<br>WP_284824106.1<br>WP_376999729.1 | 1<br>2<br>3<br>4 | EMAQLIAEHTGQSFEQITK<br>IMMHQPSAGVGGTAADIAIQAEQFAQTK<br>IIFLGTQVDDEIANK<br>YALPHAR | 149-167<br>120-147<br>29-43<br>113-119 | 21575 |

| | | | | | | | |
|---|---|---|---|---|---|---|---|
| | | C. phoceense (3,4)<br>C. endometrii (4)<br>C. pseudodiphtheriticum (3)<br>C. casei (3)<br>C. ammoniagenes (3)<br>C. lizhenjunii (3) | MCQ9334125.1<br>WP_136141682.1<br>WP_249618353.1<br>WP_006821938.1<br>WP_003847280.1<br>WP_165009510.1 | | | | |
| 5 | ATP-dependent Clp protease proteolytic subunit | C. propinquum<br>C. pseudiphtheriticum<br>C. sphenisci | WP_018120935.1<br>WP_284856599.1<br>WP_075693676.1 | 1 | TLMERTLAEHTGR | 158-170 | 23166 |
| 6 | ATP-dependent Clp protease adapter protein ClpS | Corynebacterium sp. KPL1995 | ERS42111.1 | 1 | LAVMNTRK | 11-18 | 15635 |
| 7 | Trypsin-like serine protease | C. accolens<br>C. macginleyii<br>C. minutissimum<br>C. striatum<br>C. amycolatum<br>C. phoceense<br>C. diphtheriae<br>C. simulans | WP_302503567.1<br>WP_302503567.1<br>WP_115020991.1<br>WP_115020991.1<br>WP_419385559.1<br>WP_196131656.1<br>WP_248092130.1<br>WP_248092130.1 | 1 | HCLESVNNEGTQAR | 30-43 | 33839 |
| 8 | Trypsin-like serine protease | C. accolens | WP_302503567.1 | 1 | EWAEGVMSGK | 332-341 | 40648 |
| 9 | Zinc-dependent metalloprotease | Tsukamurella paurometabola | WP_126196731.1 | 1 | LVNPVAEK | 136-143 | 48395 |
| 10 | S9 family peptidase | C. striatum<br>C. simulans<br>C. accolens | WP_306588675.1<br>WP_062037262.1<br>PCC82544.1 | 1<br>2<br>3<br>4<br>5<br>6<br>7<br>8 | EQEVPGGYNKDDYVAYR<br>GGQFLLK<br>HGWSEFK<br>LEGVDTYR<br>MWTQASDGTQIPVSIVHR<br>SFVDDYEWMR<br>TEMVAGHGGVSGR<br>VETGEYTLLK | 428-444<br>665-671<br>370-376<br>341-348<br>445-462<br>22-31<br>672-684<br>418-427 | 79669 |
| 11 | Type I methionyl aminopeptidase | C. striatum | WP_086891163.1 | 1<br>2<br>3<br>4<br>5<br>6<br>7<br>8 | AVAPGVTTDEIDR<br>EVNVIGR<br>FGYNVVR<br>IAANALQEAGK<br>LTPQKPTPIR<br>THEAMMR<br>TVPDSIERPEYVWK<br>VIESYANR | 68-80<br>180-186<br>195-201<br>57-67<br>7-16<br>164-170<br>17-30<br>187-194 | 31887 |
| 12 | MULTISPECIES: M3 family metallopeptidase | Corynebacterium | WP_061920700.1 | 1<br>2<br>3<br>4 | ARDFSLDEEELR<br>DFSLDEEELR<br>AGLNNPLIAPR<br>DWVEFPSQINENWALEPALVK | 322-333<br>324-333<br>581-591<br>489-509 | 54945 |
| 13 | MULTISPECIES: aminopeptidase P family protein | Corynebacterium | WP_023021464.1 | 1<br>2<br>3 | EIYDLVLR<br>GFNSDCTR<br>YLSGFSGSNGGLLLR | 252-259<br>232-239<br>37-51 | 39269 |
| 14 | MULTISPECIES: | Corynebacterium | WP_049151732.1 | 1 | GFNSDCTR | 249-256 | 41319 |

| | | | | | | | |
|---|---|---|---|---|---|---|---|
| | aminopeptidase P family protein | | | 2<br>3 | LVDVDAACR<br>QIAADLEYR | 291-299<br>192-200 | |
| 15 | MULTISPECIES: M18 family aminopeptidase | Corynebacterium | WP_034668733.1 | 1<br>2<br>3<br>4 | ALGANEEER<br>FGAAGPILADVLGR<br>FVGNNDVPCGSTIGPITATR<br>LGIDTVDVGVPMLSMHSAR | 280-288<br>263-276<br>360-379<br>380-398 | 44848 |
| 16 | M13-type metalloendopeptidase | C. simulans<br>C. striatum | WP_284841565.1<br>WP_110333686.1 | 1<br>2<br>3<br>4<br>5<br>6<br>7 | AEMDTLVDYLTAAYR<br>AGNAWAHDFELR<br>AVGLAESLVGQEIGK<br>IGYPDKWR<br>ISQLEWMTPATR<br>NFEFYGTR<br>VFVDQHFPPASK | 317-331<br>386-397<br>290-304<br>359-366<br>334-345<br>269-276<br>305-316 | 71583 |
| 17 | Aminopeptidase N | C. striatum<br>C. simulans | WP_204083832.1<br>WP_248092840.1 | 1<br>2<br>3 | AVLDDATDPDIR<br>DNTLTGAAAYLGVSR<br>QNFDGITYAK | 665-676<br>701-715<br>388-397 | 93277 |
| 18 | Neutral zinc metallopeptidase | C. striatum<br>C. simulans | WP_284765519.1<br>WP_239238688.1 | 1 | AVGDDNIQR | 244-252 | 33782 |
| 19 | MULTISPECIES: M13-type metalloendopeptidase | Corynebacterium | WP_049151553.1 | 1<br>2 | ELAAGHWDVVSTR<br>ISQLEWMTPATR | 181-193<br>337-348 | 71398 |
| 20 | MULTISPECIES: aminopeptidase P family protein | Corynebacterium | WP_283117106.1 | 1 | IEDTLIITSGAPK | 339-351 | 39134 |
| 21 | MULTISPECIES: S9 family peptidase | Corynebacterium | WP_147760088.1 | 1<br>2 | EGVDTYR<br>NTFTDFIAVADDLIAR | 376-383<br>564-579 | 83268 |
| 22 | MULTISPECIES: leucyl aminopeptidase | Corynebacterium | WP_225723588.1 | 1 | LVLADAIAR | 367-375 | 51420 |
| 23 | M3 family metallopeptidase | C. striatum<br>C. simulans | WP_086891990.1<br>WP_284841026.1 | 2 | AGLNNPLIAPR | 581-591 | 76034 |
| 24 | MULTISPECIES: C40 family peptidase | Corynebacterium | WP_301717425.1 | 1 | QAGVELPR | 240-247 | 30171 |
| 25 | Type I methionyl aminopeptidase | C. variabile | WP_312776128.1 | 1 | SPAELDAMQAAGEIVGK | 14-30 | 27858 |
| 26 | Type I methionyl aminopeptidase | C. simulans<br>C. striatum | WP_248092738.1<br>WP_279109652.1 | 1<br>2 | LTDVSHALEQATYR<br>TPGELDAMQAAGEIVGK | 152-165<br>14-30 | 27954 |
| 27 | Penicillin-binding transpeptidase domain-containing protein | C. amycolatum<br>C. lactis<br>C. vitaeruminis | WP_256885874.1<br>WP_053411183.1<br>WP_048761742.1 | 1 | MEPQIIK | 355-361 | 50025 |
| 28 | D-alanyl-D-alanine carboxypeptidase/D-alanyl-D-alanine-endopeptidase | C. pseudotuberculosis | WP_058832193.1 | 1 | ATLDAMAHDR | 56-65 | 40521 |
| 29 | M23 family metallopeptidase | Corynebacterium sp. CCUG 65737 | WP_256003305.1 | 1 | SMKPAEGTFTSGFGMR | 111-126 | 24645 |
| 30 | Peptidoglycan DD-metalloendopeptidase family protein | C. callunae | WP_015650630.1 | 1 | QELFPANGSPYMMEDR | 296-311 | 55737 |
