## Supplemental Table S24 for "Peptidome Analysis of Western Blots Identifies Natural Bispecific Antibody-Bound *Corynebacterium* and Phage B-cell Epitopes with Potential Relevance to Psoriasis"

### Table S24. Domains of Unknown Functions (DUF) are *Corynebacterium* Antigens.

| Antigen Number | Identified Antigen | Corynebacterium (C.), *Actinomycetota* | Accession Number | Peptide Number | Peptide Sequence | Peptide Start/Stop | Molecular Weight |
|---|---|---|---|---|---|---|---|
| 1 | DUF5979 domain-containing protein | C. pseudotuberculosis | WP_054250864.1 | 1 | LVPDQHGR | 282-289 | 119354 |
| 2 | DUF456 domain-containing protein | C. glutamicum | WP_011014634.1 | 1 | LMVGLFIGLCILIVGGPALGLMLGGFIGVLIGLCL GTSAAMIVGPFGLMRMKAYPPCMDSPWVYMSQ EQWR | 96-166 | 22671 |
| 3 | DUF1846 domain-containing protein | C. striatum<br>C. simulans | WP_272707507.1<br>WP_062042977.1 | 1<br>2<br>3 | ADLGILYEDDVLR<br>DLVVVTAPGPGSGK<br>VIPGYPSNTDLIVSEEGLGR | 86-98<br>172-185<br>144-163 | 55081 |
| 4 | DUF5926 family protein | C. aurimucosum<br>C. intestinale<br>C. minutissimum<br>C. hiratae<br>C. lizhenjunii<br>C. tuberculostearicum<br>C. accolens<br>C. marquesiae<br>C. flavescens<br>C. kefirresidentii<br>C. simulans | WP_158381411.1<br>WP_250224288.1<br>WP_115023939.1<br>WP_144013375.1<br>WP_165010979.1<br>WP_317000398.1<br>WP_284902413.1<br>WP_337888724.1<br>WP_301523978.1<br>WP_086588567.1<br>WP_062043057.1 | 1<br>2 | DLAFALNWVK<br>KQLENIK | 110-119<br>293-299 | 33387 |
| 5 | MULTISPECIES: hemolysin family protein<br>MULTISPECIES: DUF5926 family protein<br>Uncharacterized protein | Rhodococcus<br>Corynebacterium<br>Mycobacterium tuberculosis | WP_068100225.1<br>WP_061925103.1<br>SGC98557.1 | 1 | EIDNDE | 365-370 | 21142 |
| 6 | DUF349 domain-containing protein | C. simulans<br>C. striatum<br>C. aurimucosum<br>C. faecipullorum | WP_284841192.1<br>WP_086891208.1<br>WP_010190274.1<br>HIX80209.1 | 1<br>2<br>3<br>4<br>5<br>6<br>7<br>8<br>9<br>10<br>11 | AAQDHFFSAR<br>AQAAQWQEFADAAAAAVNDK<br>DALIAEYDSQIDPSK<br>KEELVER<br>RTDPEAQAR<br>DALIAEYDSQIDPSK<br>EKLAAEAEDIAANSTEWK<br>KEELVER<br>LAAEAEDIAANSTEWK<br>RTDPEAQAR<br>WEEIGFVPR | 283-292<br>416-435<br>311-325<br>233-239<br>376-384<br>314-328<br>159-176<br>236-242<br>161-176<br>379-387<br>344-352 | 48383 |
| 7 | DUF885 domain-containing protein | C. striatum | WP_306592399.1 | 1<br>2<br>3 | DTLLLMPQDTPEQLDAIR<br>QINEVFVQCER<br>YVLEGTDALVEWMQATADK | 126-143<br>174-184<br>294-312 | 62002 |
| 8 | DUF1003 domain-containing protein | C. matruchotii | WP_314974200.1 | 1 | IEAKIADEAAAR | 151-162 | 21559 |
| 9 | DUF1906 domain-containing protein (Tat pathway signal sequence domain protein) | C. accolens | WP_284637643.1 | 1 | AGHIGAIR | 56-63 | 32066 |
| 10 | MULTISPECIES: DUF3107 domain-containing protein | Corynebacterium | WP_003858150.1 | 1 | IGFADTAR | 5-12 | 7987 |
| 11 | DUF3499 domain-containing protein | Prescottella equi | WP_084968582.1 | 1 | EAGLGERPR | 87-95 | 12018 |

| | | | | | | | |
|---|---|---|---|---|---|---|---|
| 12 | DUF3499 domain-containing protein | C. maris | WP_156844687.1 | 1 | VTTGLVDEK | 69-77 | 13562 |
| 13 | DUF3152 domain-containing protein | C. jeikeium<br>C. kroppenstedtii | WP_005293842.1<br>WP_012732098.1 | 1 | RVVLNEAR | 221-228 | 34465 |
| 14 | DUF402 domain-containing protein | C. striatum | WP_306495913.1 | 1<br>2 | YGDDAMAWLR<br>ANIFHFR | 151-160<br>66-72 | 19926 |
| 15 | MULTISPECIES: DUF3117 domain-containing protein | Corynebacterium | WP_005531903.1 | 1 | IVVELTKEEAAELGSLLTEVSS | 34-55 | 5816 |
| 16 | DUF6474 family protein | C. matruchotii | WP_005522019.1 | 1 | KGRLNSGK | 78-85 | 23478 |
| 17 | DUF6474 family protein | C. simulans<br>C. accolens | WP_239240387.1<br>WP_278723472.1 | 1 | TAAPLLLPLVYR | 95-106 | 23924 |
| 18 | DUF6474 family protein | C. striatum<br>C. simulans | WP_049151340.1<br>WP_239240387.1 | 1 | MAQLELDKLK | 68-77 | 23945 |
| 19 | DUF5997 family protein | C. maris | WP_041631751.1 | 1 | ELQNNPPEWLVELR | 30-43 | 12751 |
| 20 | DUF2631 domain-containing protein | C. striatum<br>C. simulans | WP_168926514.1<br>WP_062038483.1 | 1 | NQPVGHVEPDWVYDQK | 106-121 | 17922 |
| 21 | DUF2631 domain-containing protein | C. simulans<br>C. pseudogenitalium<br>C. striatum | WP_275436966.1<br>WP_256000339.1<br>WP_005528921.1 | 1 | ALNIDPAR | 141-148 | 16068 |
| 22 | DUF294 nucleotidyltransferase-like domain-containing protein | C. stationis<br>C. ammoniages | WP_075722990.1<br>WP_236163793.1 | 1 | QSAGKTVSR | 539-547 | 68780 |
| 23 | DUF294 nucleotidyltransferase-like domain-containing protein | C. stationis<br>C. casei<br>C. ammoniagenes | WP_066794542.1<br>WP_301525099.1<br>WP_236163793.1 | 1 | ENLRDAFSIIK | 593-603 | 68631 |
| 24 | DUF294 nucleotidyltransferase-like domain-containing protein | C. diphtheriae | WP_235697141.1 | 1 | MLATPCADFMSR | 124-135 | 43258 |
| 25 | DUF3052 domain-containing protein | C. simulans<br>C. aurimucosum<br>C. minutissimum<br>C. aurimucosum<br>C. striatum | WP_282440188.1<br>WP_262347198.1<br>WP_239189523.1<br>TVU86878.1<br>MDU3173918.1 | 1<br>2 | AEDGDLVDGLVDATRPLGDSGR<br>LGEWQGSCLVAR | 95-116<br>152-163 | 18270 |
| 26 | MULTISPECIES: DUF262 domain-containing protein | Corynebacterium | WP_061922129.1 | 1 | GYPIGALMALDTR | 48-60 | 67165 |
| 27 | DUF3093 domain-containing protein | C. terpenotabidum | WP_041631127.1 | 1 | GETWLRVGDASLPR | 98-111 | 19969 |
| 28 | DUF3093 domain-containing protein | C. striatum<br>C. simulans | WP_205690714.1<br>WP_062039498.1 | 1 | DAQLPHDVVQR | 95-105 | 19552 |
| 29 | DUF6286 domain-containing protein | C. accolens | WP_005276893.1 | 1 | AEMNMPKHFGQQPK | 2-15 | 20223 |
| 30 | DUF6286 domain-containing protein | C. casei | WP_301436480.1 | 1 | DMWVTYSESTR | 133-143 | 31634 |
| 31 | DUF2800 domain-containing protein | C. accolens<br>C. ulcerans | WP_259817699.1<br>WP_197906841.1 | 1 | LLGVTAMTK | 420-428<br>336-344 | 52426 |
| 32 | DUF4247 domain-containing protein | C. amycolatum | WP_005510135.1 | 1 | DPIEVGR | 63-69 | 16096 |
| 33 | DUF2550 domain-containing protein | C. matruchotii | WP_278718476.1 | 1 | LDKMSPSEAR | 132-141 | 17802 |
| 34 | DUF2550 domain-containing protein | C. striatum<br>C. simulans<br>C. accolens | WP_284765007.1<br>WP_239238847.1<br>PCC82903.1 | 1 | YNGEYVEYFK | 58-67 | 18275 |
| 35 | DUF2550 domain-containing protein | C. durum | WP_315497723.1 | 1 | KMPAPGDHGWR | 37-47 | 18325 |
| 36 | MULTISPECIES: DUF3662 and FHA domain-containing protein | Corynebacterium | WP_003861088.1 | 1<br>2 | EGSNIIGR<br>SNDADLRLPDTGVSR | 221-228<br>229-243 | 32317 |

| | | | | | | | |
|---|---|---|---|---|---|---|---|
| 37 | DUF4229 domain-containing protein | C. striatum | HCG3140840.1 | 1<br>2 | AWVQAELAGR<br>VEATETLAEYSAQR | 100-109<br>82-95 | 12117 |
| 38 | MULTISPECIES: DUF2505 domain-containing protein | Corynebacterium | WP_049377830.1 | 1<br>2<br>3 | LTEQWIADNL<br>SENTVTINQPIEK<br>AMVSQDLK | 154-163<br>5-17<br>67-74 | 17618 |
| 39 | DUF3040 domain-containing protein | C. striatum<br>C. simulans | WP_131771534.1<br>WP_275436953.1 | 1 | SLSEHEQQALR | 2-12 | 13981 |
| 40 | DUF3618 domain-containing protein | C. kroppenstedtii<br>C. simulans<br>C. striatum | WP_303735459.1<br>WP_284841749.1<br>HAT1399637.1 | 1 | QLASTLDELADR | 18-29 | 10177 |
| 41 | DUF4307 domain-containing protein | C. striatum | WP_284764964.1 | 1 | IAVDVPTNHR | 122-131 | 13429 |
| 42 | DUF4178 domain-containing protein | C. striatum<br>C. simulans | WP_005530413.1<br>WP_062036246.1 | 1<br>2<br>3<br>4<br>5<br>6 | GTVEFEGVTYSEDER<br>TMLPGEFTVYPAPK<br>TMLPGEFTVYPAPKA<br>YGATDYVVR<br>YTSTGTTGLPESGEMR<br>YVDYSAGEK | 126-140<br>189-202<br>189-203<br>65-73<br>145-160<br>161-169 | 22476 |
| 43 | DUF418 domain-containing protein | C. amycolatum<br>C. diphtheria<br>C. striatum<br>C. vitaeruminis<br>C. simulans<br>C. argentoratense | WP_187404796.1<br>WP_014318800.1<br>MDU3175889.1<br>WP_048759848.1<br>WP_096336438.1<br>WP_315041204.1 | 1 | LVATAIK | 303-309 | 33515 |
| 44 | DUF4191 domain-containing protein | C. striatum<br>C. aurimucosum<br>C. simulans<br>C. kefirresidentii<br>C. flavescens<br>C. phoceense | WP_306588904.1<br>WP_010190601.1<br>WP_248093925.1<br>WP_239204421.1<br>MDN6460784.1<br>WP_303938144.1 | 1 | KVEDQPGVAGWALEQQLR | 100-117 | 29740 |
| 45 | DUF4191 domain-containing protein | C. striatum<br>C. simulans | WP_306592350.1<br>WP_248093925.1 | 1 | SQMWQAFNMQR | 29-39 | 29104 |
| 46 | DUF1707 domain-containing protein | C. simulans<br>C. striatum<br>C. accolens | WP_248093882.1<br>WP_306588736.1<br>PCC82507.1 | 1 | QAGELTSEAMNLAK | 193-206 | 23788 |
| 47 | DUF501 domain-containing protein | C. striatum<br>C. accolens<br>C. simulans | WP_306497159.1<br>PCC83169.1<br>WP_062041425.1 | 1<br>2<br>3 | GVLDIAYR<br>TVNDADLEIVK<br>TVNDADLEIVKEQLGR | 21-28<br>2-12<br>2-17 | 20266 |
| 48 | DUF501 domain-containing protein | C. falsenii | WP_224208468.1 | 1 | AHEHYLEQRNAMEDLGTDFSGGGMPER | 109-135 | 21745 |
| 49 | DUF3068 domain-containing protein | C. durum | WP_273112225.1 | 1 | IGMTLSK | 86-92 | 35831 |
| 50 | DUF3427 domain-containing protein | C. aurimucosum | WP_046648887.1 | 1 | SELEEAPK | | 118011 |
| 51 | DUF732 domain-containing protein | C. terpenotabidum | WP_020440221.1 | 1 | DPETTAAAIK | 151-160 | 17030 |
| 52 | DUF5129 domain-containing protein | C. diphtheriae | WP_014308841.1 | 1 | LENGDAVIRVR | 341-351 | 53682 |
| 53 | DUF3515 domain-containing protein | C. tuberculostearicum<br>C. marquesiae<br>C. pseudogenitalium<br>C. aurimucosum | WP_034668164.1<br>WP_284836510.1<br>WP_040425003.1<br>MCG7446867.1 | 1 | AGNDQMCAEVDK | 180-191 | 30446 |
| 54 | DUF6615 family protein | C. glyciniphilum | WP_052541097.1 | 1 | DWYRCSVDHQSGHCPGCEKR | 133-152 | 32704 |
| 55 | DUF11 domain-containing protein | Prescottella equi | WP_081205740.1 | 1 | DVTVNEGDFTGTGTMSAITCPDESK | 1385-1409 | 161162 |

| | | | | | | | |
|---|---|---|---|---|---|---|---|
| 56 | DUF4190 domain-containing protein | Clavibacter michiganensis subsp. sepedonicus | WP_012297968.1 | 1 | GLAITSMVLGIVCVVLSLPLWFLTFFVGIAAIITGVLAR | 50-88 | 14463 |
| 57 | DUF5319 domain-containing protein | C. pseudodiphtheriticum | WP_249624534.1 | 1 | MVRKMAQTLPEFGIDGIFFICEDCEEQHYYDWDIMAANMR | 46-85 | ? |
| 58 | DUF5635 domain-containing protein | C. glucuronolyticum | WP_005393345.1 | 1 | MYQSMIVLGHRPPIFEEVEGPFISVTLKGGKPVLPVMQLVSAIVPEPR | 378-425 | 62113 |
| 59 | DUF3100 domain-containing protein | C. glyciniphilum | WP_038545702.1 | 1 | APAIIVVITVGTLLTIPASPVSSWLLQVSESVDFLAVITMMLSVAGLSIGKDIPLLR | 368-424 | 47697 |
