## Supplemental Table S25 for "Peptidome Analysis of Western Blots Identifies Natural Bispecific Antibody-Bound *Corynebacterium* and Phage B-cell Epitopes with Potential Relevance to Psoriasis"

### Table S25. Bacterial Transporters and Exporters are *Corynebacterium* Antigens.

| Antigen Number | Identified Antigen | Corynebacterium (C.), Other Actinobacteria (Peptide numbers) | Accession Number | Peptide Number | Peptide Sequence | Peptide Start/Stop | Molecular Weight |
|---|---|---|---|---|---|---|---|
| 1 | ABC transporter ATP-binding protein | C. accolens<br>C. tuberculostearicum<br>C. kefirresidentii<br>C. marquesiae<br>C. phocae<br>C. aurimucosum<br>C. singulare<br>C. simulans<br>C. striatum | WP_302526085.1<br>WP_316993376.1<br>WP_086588245.1<br>WP_284791212.1<br>WP_075735751.1<br>WP_049360350.1<br>WP_239180967.1<br>WP_248093058.1<br>MDU3174403.1 | 1 | VDEVLGLVGLSDVAGK | 108-123 | 34741 |
| 2 | ABC transporter ATP-binding protein | C. tuberculostearicum<br>C. aurimucosum | WP_250413062.1<br>WP_049358646.1 | 1 | DAAGMSNSERR | 431-441 | 68564 |
| 3 | ABC transporter ATP-binding protein | C. urealyticum<br>C. tuberculostearicum<br>C. striatum<br>C. confusum | WP_148820884.1<br>WP_316993538.1<br>MBD0857217.1<br>WP_290224007.1 | 1 | GRVVETGTHEELAAR | 540-554 | 63709 |
| 4 | ABC transporter, ATP-binding protein | C. tuberculostearicum<br>C. kefirresidentii<br>C. pilosum<br>C. curiae<br>C. marquesiae<br>C. pseudogenitalium<br>C. aurimucosum | WP_301714576.1<br>WP_301716579.1<br>WP_018581717.1<br>WP_269945987.1<br>WP_284786641.1<br>WP_005323417.1<br>WP_049361410.1 | 1 | MDDKLDDLSLGNQQR | 125-139 | 18978 |
| 5 | Multidrug efflux MFS transporter Cmr | C. glutamicum | WP_077313408.1 | 1 | MSTFHKVLINTMISNVTTGFLFFAVVFW<br>MYLSTGDVALTGIVSGIYMGLIAVCSIFFG<br>TVVDHNRK | 1-66 | 49463 |
| 6 | MDR family MFS transporter | C. casei | WP_006822801.1 | 1 | GKKTADNVGIVFAALMLTMLMSSLGQMI<br>FGSALPTIVGELGGVDQMSWVISAFMVT<br>MTIAMPLAGQLGDRMGR | 40-112 | 57941 |
| 7 | Drug resistance MFS transporter, drug:H+ antiporter-2 family | C. matruchotii | WP_315118707.1 | 1 | YKWMPIASMVVLAISLWLLSTLTVETPL<br>WVLLSYMFLLGAGIGLGMQILVLVVQNS<br>FSDHEVGMATASNNFFR | 350-422 | 56305 |
| 8 | Metal ABC transporter solute-binding protein, Zn/Mn family | C. striatum (1,2,3,4,5,6,7)<br>C. kefirresidentii (1,4,5,7)<br>C. simulans (1,2,3,4,5,6,7)<br>C. tuberculostearicum (1,2,4,5,6,7)<br>C. accolens (1,3,5,6)<br>C. singulare (1)<br>C. urealyticum (1,6)<br>C. macginleyi (1)<br>C. aurimucosum (4,6,7)<br>C. marquesiae (4,5,6,7)<br>C. lactis (6) | WP_420674450.1<br>WP_086588904.1<br>WP_062042361.1<br>WP_200436259.1<br>WP_284838567.1<br>WP_222114882.1<br>PZO98413.1<br>WP_121910444.1<br>WP_049358700.1<br>WP_408933428.1<br>WP_053411775.1 | 1<br>2<br>3<br>4<br>5<br>6<br>7 | AIFAENSNNSK<br>AVATTTQICDYLTQIAAGGVK<br>FIGSALSDFNHQQDATSSHIDEQVK<br>IQVDNIVNTLSQASPQAK<br>QQPLPESLQEFDFAK<br>YDPHVWTSPK<br>YTDQLEDLDKWASDSMSAVPEQDR | 308-318<br>51-71<br>275-299<br>205-222<br>372-386<br>192-201<br>232-255 | 41516 |
| 9 | Phosphate ABC | C. diphtheria (3) | WP_085665932.1 | 1 | IFKGEITK | 148-155 | 38021 |

| | | | | | | | |
|---|---|---|---|---|---|---|---|
| | transporter substrate-binding protein PstS | C. pseudotuberculosis (3)<br>C. simulans (1,2,3)<br>C. cystitides (1)<br>C. nasicanis (1,3)<br>C. pollutisoli (1)<br>C. renale (1)<br>C. callunae (1)<br>C. glutamicum (1)<br>C. singulare (1,2)<br>C. vitaeruminis (1)<br>C. efficiens (1)<br>C. hiratae (1,2)<br>C. aurimucosum (1,2)<br>C. guaraldiae (1)<br>C. gallinarum (1)<br>C. marinum (1)<br>C. striatum (1,2,3)<br>C. phocae (1)<br>C. endometrii (1)<br>C. accolens (1)<br>C. pseudodiphtheriticum (3)<br>C. propinquum (3)<br>C. appendicis (3)<br>C. pilosum (3)<br>C. falsenii (3)<br>C. glaucum (3)<br>C. massiliense (3)<br>C. ureicelerivorans (3)<br>C. afermentans (3)<br>C. mucifaciens (3)<br>C. riegelii (3)<br>C. fourneri (3)<br>C. urinipleomorphum (3)<br>C. lipophiloflavum (3) | WP_041480866.1<br>WP_248093161.1<br>WP_306578246.1<br>WP_377002380.1<br>WP_085550453.1<br>WP_048380726.1<br>WP_411208171.1<br>WP_074506082.1<br>WP_042532191.1<br>WP_025253533.1<br>WP_006769176.1<br>WP_328287850.1<br>WP_193634638.1<br>WP_143335405.1<br>WP_191733238.1<br>WP_042622065.1<br>WP_086890768.1<br>WP_075732937.1<br>WP_136141780.1<br>PCC82523.1<br>WP_248093161.1<br>WP_302531030.1<br>WJY61802.1<br>WP_048403072.1<br>CAM2872126.1<br>WP_095660631.1<br>WP_022863044.1<br>WP_301695159.1<br>WP_194560738.1<br>WP_168685336.1<br>WP_201821566.1<br>WP_085957312.1<br>WP_087117544.1<br>WP_006840637.1 | 2<br>3 | SAMSVFEK<br>SDESGTSDNFQK | 52-59<br>181-192 | |
| 10 | ABC transporter substrate-binding protein; Maltose/maltodextrin transport system substrate-binding protein | C. striatum (1,2,3,4,5,6,7,8,9,10,11)<br>C. simulans (1,2,3,4,5,6,7,8,9,10,11)<br>C. breve (3)<br>C. singulare (4,5,6,7,8,10,11)<br>C. aurimucosum (4,5,6,7,8,9,10)<br>C. minutissimum (4,6,7,8)<br>C. urealyticum (4,6,10)<br>C. confusum (4)<br>C. phoceense (4,6)<br>C. suedekumii (5,9)<br>C. marinum (5,9)<br>C. pollutisoli (5,6,9)<br>C. comes (5,9)<br>C. ammoniagenes (5,8,9)<br>C. casei (5,9) | WP_166684334.1<br>WP_062042010.1<br>WP_284825680.1<br>WP_144791178.1<br>WP_193634241.1<br>WP_239189793.1<br>PZP00339.1<br>WP_290226211.1<br>WP_257053616.1<br>WP_284876048.1<br>WP_042620843.1<br>WP_085549558.1<br>WP_156227033.1<br>WP_003849624.1<br>WP_006821486.1 | 1<br>2<br>3<br>4<br>5<br>6<br>7<br>8<br>9<br>10<br>11 | AGLQALVDAYK<br>AGLQALVDAYKDGVISK<br>DGVGVSTLGGYNNGININSEK<br>DSTAATEEETNLAFTEGK<br>ELAGEADAQR<br>ESLENAAPRPVSPFYTAISK<br>FEVQPLVGK<br>FIINEENQK<br>GPITFAMGK<br>KATALDFMK<br>NTDLADKAPEK | 245-255<br>245-261<br>309-329<br>262-279<br>91-100<br>379-398<br>300-308<br>339-347<br>57-65<br>330-338<br>176-186 | 45809 |

| | | | | | | | |
|---|---|---|---|---|---|---|---|
| | | C. gallinarum (5,6,7,9) | WP_191732548.1 | | | | |
| | | C. efficiens (5,6,7,9) | WP_006769566.1 | | | | |
| | | C. phocae (5,9) | WP_075735347.1 | | | | |
| | | C. humireducens (5,9) | WP_040085059.1 | | | | |
| | | C. nasicanis (5,7,9) | WP_377002153.1 | | | | |
| | | C. testudinoris (5,9) | WP_047252467.1 | | | | |
| | | C. stationis (5,9) | WP_066840347.1 | | | | |
| | | C. riegelii (5,9) | WP_311356974.1 | | | | |
| | | C. hiratae (5,6,7,8,10) | WP_144013045.1 | | | | |
| | | C. faecale (6) | WP_290278735.1 | | | | |
| | | C. faecipullorum (6) | HIX78808.1 | | | | |
| | | C. humireducens (7) | WP_040085059.1 | | | | |
| | | C. atrinae (7,9) | WP_290219543.1 | | | | |
| | | C. testudinoris (7) | WP_047252467.1 | | | | |
| | | C. sanguinis (7) | WP_259810157.1 | | | | |
| | | C. felinum (7) | WP_277103925.1 | | | | |
| | | C. lipophiloflavum (7) | WP_006839610.1 | | | | |
| | | C. camporealensis (8) | WP_035106734.1 | | | | |
| | | C. accolens (8) | WP_005277405.1 | | | | |
| | | C. spheniscorum (9) | WP_092283552.1 | | | | |
| | | C. bovis (9,10) | WP_342015147.1 | | | | |
| | | C. pseudotuberculosis (9) | AEK91832.1 | | | | |
| | | C. ulcerans (9,10) | AIU29918.1 | | | | |
| | | C. kutscheri (10) | VEH04692.1 | | | | |
| | | C. imitans (10) | WP_284785803.1 | | | | |
| | | C. vitaeruminis (10) **41** | WP_276651122.1 | | | | |
| 11 | ABC transporter substrate-binding protein Maltose/maltodextrin transport system substrate-binding protein | C. aurimucosum (1,2,3,4,5,6,7) | WP_201828646.1 | 1 | AVQDNAYAALTADK | 397-410 | 45767 |
| | | C. striatum (1,2,3,4,5,6,7) | WP_166684334.1 | 2 | AVQDNAYAALTADKSVDDATK | 397-417 | |
| | | C. hesseae (1,2,3,4,5,6,7) | WP_101735896.1 | 3 | ELAGEADAQR | 89-98 | |
| | | C. urealyticum (1,2,4) | PZP00339.1 | 4 | ESLENAAPRPVSPFYTAISK | 377-396 | |
| | | C. simulans (1,2,3,4,5,6,7) | WP_339019116.1 | 5 | FEVQPLVGK | 298-306 | |
| | | C. faecipullorum (1,2,4) | HIX78808.1 | 6 | FIINEENQK | 337-345 | |
| | | C. suedikumii (3,7) | WP_284876048.1 | 7 | GPITFAMGK | 55-63 | |
| | | C. marinum (3,7) | WP_042620843.1 | | | | |
| | | C. pollutisoli (3,4,7) | HJD79349.1 | | | | |
| | | C. comes (3,4) | WP_156227033.1 | | | | |
| | | C. ammoniagenes (3,6,7) | WP_003849624.1 | | | | |
| | | C. casei (3,7) | WP_006821486.1 | | | | |
| | | C. gallinarum (3,4,5,7) | WP_191732548.1 | | | | |
| | | C. efficiens (3,4,5,7) | WP_006769566.1 | | | | |
| | | C. phocae (3,7) | WP_075735347.1 | | | | |
| | | C. humireducens (3,5,7) | WP_040085059.1 | | | | |
| | | C. nasicanis (3,5,7) | WP_377002153.1 | | | | |
| | | C. atrinae (3,5,7) | WP_290219543.1 | | | | |
| | | C. testudinoris (3,5,7) | WP_047252467.1 | | | | |
| | | C. singulare (3,4,5,6,7) | WP_144791178.1 | | | | |
| | | C. stationis (3,7) | WP_313678888.1 | | | | |
| | | C. rigelii (3,7) | WP_311356974.1 | | | | |
| | | C. hiratae (3,4,6,7) | WP_144013045.1 | | | | |

| | | | | | | | |
|---|---|---|---|---|---|---|---|
| | | C. minutissimum (4,5,6)<br>C. faecale (4)<br>C. phoceense (4)<br>C. sanguinis (5)<br>C. felinum (5)<br>C. lipophiloflavum (5)<br>C. camporealensis (6)<br>C. accolens (6)<br>C. bovis (7)<br>C. pseudotuberculosis (7)<br>C. ulcerans (7) | WP_115021261.1<br>WP_290278735.1<br>WP_257053616.1<br>WP_259810157.1<br>WP_277103925.1<br>WP_006839610.1<br>WP_035106734.1<br>WP_005277405.1<br>WP_342015147.1<br>AEK91832.1<br>AIU29918.1 | | | | |
| 12 | MULTISPECIES: ABC transporter substrate-binding protein family 5 | Corynebacterium | WP_100086963.1 | 1<br>2 | NTLGIEAVGAPYPDFK<br>SLRDEVTNR | 410-425<br>426-434 | 58007 |
| 13 | Anaerobic C4-dicarboxylate transporter | C. striatum | WP_188311039.1 | 1 | MKDPEFAK | 222-229 | 48574 |
| 14 | Daunorubicin resistance ABC transporter ATPase subunit | Mycobacterium tuberculosis | CND67133.1 | 1 | AEGLEKR | 8-14 | 8849 |
| 15 | PTS ascorbate transporter subunit IIC | C. striatum<br>C. simulans<br>C. accolens | WP_110333508.1<br>WP_062035714.1<br>WP_284638356.1 | 1 | AEASGVKPLAAGQTAASK | 489-506 | 53729 |
| 16 | Multidrug ABC transporter ATP-binding protein | C jeikeium | OOD32146.1 | 1 | MFEAIQSVDEGTAR | 94-107 | 127712 |
| 17 | PTS sugar transporter subunit IIA | C. striatum<br>C. simulans<br>C. accolens<br>C. vitaeruminis | WP_049146141.1<br>WP_239239782.1<br>WP_302503751.1<br>WP_276651336.1 | 1 | NTLEQVLDEWGWGK | 205-218 | 28358 |
| 18 | 16S rRNA (cytosine(1402)-N(4))-methyltransferase RsmH<br>*iron ABC transporter permease* | C. otitides<br><br><br>*C. casei* | WP_046644458.1<br><br><br>*WP_006822721.1* | 1 | LAHFGGR | 145-151 | 35688 |
| 19 | ABC transporter ATPase | Prescottella equi<br>Mycobacteroides abscessus | ERN45266.1<br>CPZ28438.1 | 1 | VTVPSVTDSPR | 537-547 | 63923 |
| 20 | ABC transporter substrate-binding protein | C. accolens | WP_302502935.1 | 1 | AVAGVLGLK | 88-96 | 31560 |
| 21 | Iron ABC transporter permease | C. efficiens<br>C. gallinarum | WP_011074987.1<br>WP_191732788.1 | 1 | ISGAGTVTVLR | 166-176 | 56432 |
| 22 | Anaerobic C4-dicarboxylate transporter | C. accolens<br>C. flavescens<br>C. tuberculostearicum<br>C. simulans<br>C. minutissimum<br>C. singulare<br>C. aurimucosum<br>C. guaraldiae | WP_284899507.1<br>WP_284899507.1<br>WP_316991255.1<br>WP_062040809.1<br>KKO77707.1<br>WP_144792670.1<br>WP_144013193.1<br>WP_154761963.1 | 1 | ELEDDPVYQER | 194-204 | 45338 |

| | | | | | | | |
|---|---|---|---|---|---|---|---|
| | | C. fournieri | WP_179154881.1 | | | | |
| 23 | ABC transporter ATP-binding protein | C. halotolerans | WP_034990807.1 | 1 | ALNEFNLSVEK | 16-26 | 32266 |
| 24 | VIT1/CCC1 transporter family protein | C. ulcerans | WP_253698248.1 | 1 | GMNHEEAENKASEVFR | 221-236 | 38921 |
| 25 | Multidrug ABC transporter permease | C. vitaeruminis | WP_276652900.1 | 1 | AEPLLAGSLR | 373-382 | 55171 |
| 26 | Fructose-specific PTS transporter subunit EIIC | C. vitaeruminis | WP_276651473.1 | 1 | LDADLGADK | 13-21 | 70249 |
| 27 | Anchored repeat ABC transporter, substrate-binding protein | C. glucuronolyticum | WP_259815086.1 | 1 | EAIAGADAR | 343-351 | 50925 |
| 28 | Carbohydrate ABC transporter permease | Clavibacter michiganensis | WP_094116250.1 | 1 | GGLTDGAVK | 313-321 | 34757 |
| 29 | Anchored repeat-type ABC transporter ATP-binding subunit | C. tuberculostearicum<br>C. kefirresidentii<br>C. marquesiae<br>C. aurimucosum | WP_316991403.1<br>WP_284835775.1<br>WP_284838960.1<br>WP_049360946.1 | 1 | SISGTVGYVPQR | 62-73 | 26098 |
| 30 | ABC transporter related protein | Tsukamurella paurometabola | WP_041944652.1 | 1 | AEALIDSLGMTR | 125-136 | 28997 |
| 31 | ABC transporter ATP-binding protein | C. argentoratense | WP_169733206.1 | 1 | GLTVLSPR | 378-385 | 53417 |
| 32 | TIGR03773 family transporter-associated surface protein | C. diphtheriae | WP_199337754.1 | 1 | DGIADTEGGIAEFK | 510-523 | 117348 |
| 33 | ABC transporter ATP-binding protein | C. glutamicum | WP_087062649.1 | 1 | LTAQDLLLK | 166-174 | 26015 |
| 34 | Amino acid ABC transporter permease | C. pseudotuberculosis | WP_213173218.1 | 1 | ESATFLADNNFR | 87-98 | 24475 |
| 35 | Heme ABC transporter ATP-binding protein | C. durum<br>C. pyruviciproducens | WP_179418869.1<br>WP_280194466.1 | 1 | DIMTLSGGERAR | 137-148 | 26901 |
| 36 | Methionine ABC transporter ATP-binding protein | C. silvaticum<br>C. pseudotuberculosis<br>C. ulcerans | WP_087453439.1<br>WP_013241294.1<br>WP_095076051.1 | 1 | LDGTDIVGMPEKKLR | 68-82 | 36788 |
| 37 | ECF transporter S component | C. striatum<br>C. accolens | WP_166683907.1<br>PCC81795.1 | 1 | SQISLTPAR | 2-10 | 25278 |
| 38 | Queuosine precursor transporter | C. simulans<br>C. striatum | WP_248093319.1<br>WP_284765124.1 | 1 | SATATTSSGESIR | 5-17 | 25058 |
| 39 | ABC transporter permease | C. striatum<br>C. simulans<br>C. aurimucosum<br>C. tuberculostearicum | WP_166683713.1<br>WP_284841569.1<br>WP_201828836.1<br>WP_239454642.1 | 1 | EGSTVFIDNEGLIK | 103-116 | 38653 |
| 40 | Carbohydrate ABC transporter permease | C. striatum | WP_100087946.1 | 1 | QLSNDSTEPVTVAIAR | 211-226 | 29736 |
| 41 | Amino acid ABC transporter permease | C. accolens<br>C. aurimucosum<br>C. tuberculostearicum<br>C. minutissimum | WP_278722649.1<br>WP_010190120.1<br>WP_200435232.1<br>WP_039675634.1 | 1 | QLAALADAEGTIPR | 298-311 | 35125 |

| | | | | | | | |
|---|---|---|---|---|---|---|---|
| | | C. simulans<br>C. urealyticum<br>C. stationis<br>C. striatum | WP_248091985.1<br>PZO98602.1<br>WP_066840664.1<br>WP_046646831.1 | | | | |
| 42 | ABC transporter ATP-binding protein | C. accolens<br>C. tuberculostearicum<br>C. kefirresidentii<br>C. marquesiae<br>C. phocae<br>C. aurimucosum<br>C. singulare<br>C. simulans<br>C. striatum | WP_302526085.1<br>WP_316993376.1<br>WP_086588245.1<br>WP_284791212.1<br>WP_075735751.1<br>WP_049360350.1<br>WP_239180967.1<br>WP_248093058.1<br>MDU3174403.1 | 1 | VDEVLGLVGLSDVAGK | 108-123 | 34741 |
| 43 | Sodium-dependent transporter | C. diphtheria<br>C. belfantii<br>C. rouxii | WP_196975809.1<br>WP_197688542.1<br>WP_155872246.1 | 1 | RIAGFAIEQYSK | 534-545 | 59828 |
| 44 | ABC transporter substrate-binding protein | Corynebacterium sp. Marseille-P3884; Periplasmic binding protein; Corynebacterium genitalium ATCC 33030 GN=HMPREF0291_10252 PE=4 SV=1 | WP_210575019.1 | 1 | IGLYTAEDGR | 205-214 | 33735 |
| 45 | Peptide MFS transporter | C. striatum<br>C. simulans<br>C. tuberculostearicum<br>C. accolens<br>C. camporealensis | WP_100087388.1<br>WP_239239498.1<br>WP_316985802.1<br>PCC82649.1<br>WP_321114909.1 | 1 | TTAQVVLGSLYSR | 142-154 | 53981 |
| 46 | MFS transporter | C. striatum<br>C. simulans<br>C. efficiens | WP_239299478.1<br>WP_248091863.1<br>WP_006770395.1 | 1 | NPLVSIEYLGQR | 259-270 | 50964 |
| 47 | ABC transporter permease subunit | C. striatum<br>C. simulans<br>C. nuruki | WP_049151596.1<br>WP_062035349.1<br>WP_010120681.1 | 1 | RVGMSR | 169-174 | 58634 |
| 48 | ABC-type transporter, ATPase component and permease component | C. glutamicum | CCH24135.1 | 1 | MGGLVDK | 1-7 | 56266 |
| 49 | Multidrug ABC transporter permease | C. callunae<br>C. testudinoris | WP_015652175.1<br>WP_047253056.1 | 1 | IGEFVGK | 196-202 | 28051 |
| 50 | Serine/threonine transporter;<br>PLP-dependent aminotransferase family protein | C. diphtheria<br>Prescottella equi | WP_199326535.1<br>WP_084987222.1 | 1 | VPALAKYR | 425-432<br>432-439 | 50667 |
| 51 | Sugar ABC transporter permease | C. vitaeruminis | WP_051645655.1 | 1 | ETATVGVDK | 345-353 | 36500 |
| 52 | ABC transporter substrate-binding protein | Prescottella equi | WP_064059497.1 | 1 | KDGYMPAAK | 270-278 | 34869 |
| 53 | Efflux RND transporter periplasmic adaptor subunit | C. genitalium | WP_156774821.1 | 1 | ALGEAHAQGR | 181-190 | 47778 |
| 54 | Iron ABC transporter | Tsukamurella paurometabola | WP_013128332.1 | 1 | GLGLNVNLSR | 258-267 | 37272 |

| | | | | | | | |
|---|---|---|---|---|---|---|---|
| | permease | | | | | | |
| 55 | MetQ/NlpA family ABC transporter substrate-binding protein | C. accolens | WP_284639927.1 | 1 | SHKSIDDVDK | 125-134 | 30747 |
| 56 | ABC transporter--like protein | C. maris | AGS33501.1 | 1 | MVRQGK | 1-6 | 26144 |
| 57 | Energy-coupling factor transporter transmembrane protein EcfT | C. kroppenstedtii<br>C. parakroppenstedtii | WP_012731751.1<br>WP_236916289.1 | 1 | GFDDWDPTAPEYDSQE | 220-235 | 25824 |
| 58 | Sugar ABC transporter permease | C. vitaeruminis | WP_051483541.1 | 1 | ETATVGVDK | 348-356 | 36500 |
| 59 | Cation:dicarboxylase symporter family transporter | C. vitaeruminis | WP_277102341.1 | 1 | VNIFKLCK | 268-275 | 50429 |
| 60 | ABC transporter permease | Tsukamurella paurometabola | WP_220656347.1 | 1 | MFNDMHGGIFER | 82-93 | 27173 |
| 61 | Fe2+-enterobactin ABC transporter substrate-binding protein | C. durum | WP_314939190.1 | 1 | SWEDVTTMMGKATGLEK | 179-195 | 37744 |
| 62 | MFS transporter | C. glyciniphilum | WP_158407374.1 | 1 | RSDDSVGFLPGEQVAPEQ | 472-489 | 51659 |
| 63 | Amino acid ABC transporter ATP-binding protein | C. pyruviciproducens | WP_101677962.1 | 1 | TAMENIMEAPMVVKGVSEEDAK | 106-127 | 27688 |
| 64 | Threonine/serine exporter family protein<br>*5-formyltetrahy-drofolate cyclo-ligase* | C. glyciniphilum<br>*C. jeikeium* | WP_304030013.1<br>*WP_080720366.1* | 1 | VDERPSR | 119-125<br>*23-29* | 22368 |
| 65 | Sulfite exporter TauE/SafE family protein | C. lipophiloflavum | WP_304041736.1 | 1 | KTAAADGAPASATR | 124-137 | 32006 |
| 66 | Aromatic acid exporter family protein | C. maris | WP_020933460.1 | 1 | MSSPAPMQALAR | 1-12 | 41713 |
