## Supplemental Table S26 for "Peptidome Analysis of Western Blots Identifies Natural Bispecific Antibody-Bound *Corynebacterium* and Phage B-cell Epitopes with Potential Relevance to Psoriasis"

### Table S26. Decoy Peptides Bound by nBsAbs are Antigens Derived from Bacteriophage Proteins.

| Peptide Number | Accession Number | Peptide Sequences, Sequence Alignments | Similar Sequences found in: |
|---|---|---|---|
| **1** | tr\|E2S1G1\|E2S1G1_9CORY-DECOY | VYTETI<br>Query 1 VYTETI 6<br>VYTETI<br>Sbjct 734 VYTETI 739 | Spore coat protein H kinase fold, TRANSFERASE [Caudoviricetes sp.]<br>Sequence ID: DAE36059.1Length: 2521 |
| **2** | tr\|E0MXM5\|E0MXM5_9CORY-DECOY | KIGVAGGK<br>Query 1 KIGVAGGK 8<br>KIG AGGK<br>Sbjct 552 KIGAAGGK 559 | chromosome segregation protein [Caudoviricetes sp.]<br>Sequence ID: DAT29569.1Length: 1752 |
| **3** | tr\|U5DLT5\|U5DLT5_COREQ-DECOY | SREDLIGPPKR<br>Query 1 SR---EDLIGPPK 10<br>SR EDLI PPK<br>Sbjct 223 SRINKEDLITPPK 235 | Integrase [Caudoviricetes sp.]<br>Sequence ID: DAZ81015.1Length: 358 |
| **4** | tr\|S4XI52\|S4XI52_9CORY- DECOY | LPLVLTK<br>Query 1 LPLVLT 6<br>LPLVLT<br>Sbjct 491 LPLVLT 496 | tail protein [Caudoviricetes sp.]<br>Sequence ID: DAV20616.1Length: 928 |
| **5** | tr\|U7LF32\|U7LF32_9CORY-DECOY | SAAEKGSR<br>Query 2 AAEKGSR 8<br>AAEKGSR<br>Sbjct 52 AAEKGSR 58 | major capsid protein [Caudoviricetes sp.]<br>Sequence ID: DAZ63806.1Length: 592 |
| **6** | tr\|H8LW18\|H8LW18_CORPS-DECOY | YADLKAK<br>Query 1 YADLKAK 7<br>YADLKAK<br>Sbjct 181 YADLKAK 187 | hypothetical protein [Caudoviricetes sp.]<br>Sequence ID: DAI13851.1Length: 501 |
| **7** | tr\|F8E1I3\|F8E1I3_CORRG-DECOY | AVSPPGFR<br>Query 1 AVSPPGFR 8<br>AV PPGFR<br>Sbjct 20 AVRPPGFR 27 | hypothetical protein, partial [Caudoviricetes sp.]<br>Sequence ID: DAN74144.1Length: 33 |
| **8** | tr\|C3PKU5\|C3PKU5_CORA7-DECOY | LLVGVDAYKSGPQLK<br>Query 1 LL-VGVDAYKSG 11<br>LL VG+DAYK G<br>Sbjct 40 LLTVGIDAYKKG 51<br><br>Query 8 YK-SGPQLK 15<br>YK SGPQLK<br>Sbjct 174 YKLSGPQLK 182 | Helix-turn-helix XRE-family like protein [Caudoviricetes sp.]<br>Sequence ID: DAZ77665.1Length: 135<br><br>replisome organizer protein [Caudoviricetes sp.]<br>Sequence ID: DAX51472.1Length: 287 |
| **9** | tr\|E2MUG9\|E2MUG9_CORAY-DECOY | YPEHVAK<br>Query 1 YPEHVAK 7<br>YPEHVAK<br>Sbjct 80 YPEHVAK 86 | NAD-dependent DNA ligase [Ralstonia phage phiRSL1]<br>Sequence ID: YP_001949941.1Length: 625 |
| **10** | tr\|O24751\|O24751_CORAM-DECOY | WELAAVR<br>Query 1 WELAAVR 7<br>W+LAAVR<br>Sbjct 14 WQLAAVR 20 | Cytosine specific methyltransferase [Caudoviricetes sp.]<br>Sequence ID: DAV08622.1Length: 235 |
| **11** | tr\|E4WHK6\|E4WHK6_RHOE1-DECOY | YAQLKPR<br>Query 1 YAQLKP 6<br>YAQLKP<br>Sbjct 176 YAQLKP 181 | CotH kinase protein [Siphoviridae sp. ctHAs12]<br>Sequence ID: DAF50646.1Length: 711 |
| **12** | tr\|C0XUL0\|C0XUL0_9CORY-DECOY | SDAAVFLR<br>Query 1 SDAAVFLR 8<br>SDAAV+LR<br>Sbjct 486 SDAAVYLR 493 | large terminase [Caudoviricetes sp.]<br>Sequence ID: DAO21307.1Length: 607 |
| **13** | tr\|I0LJG9\|I0LJG9_CORGK-DECOY | DLVDPNPL<br>Query 1 DL--VDPNPL 8<br>DL VDPNPL<br>Sbjct 105 DLKKVDPNPL 114 | DNA-directed RNA polymerase subunit alpha [Caudoviricetes sp.]<br>Sequence ID: DAV72465.1Length: 1464 |
| **14** | tr\|E2S784\|E2S784_9CORY-DECOY | FVSFPMR<br>Query 1 FVSFPMR 7<br>F SFPMR<br>Sbjct 8 FASFPMR 14 | hypothetical protein [Caudoviricetes sp.]<br>Sequence ID: DAI20370.1Length: 34 |
| **15** | tr\|U7K8R2\|U7K8R2_9CORY-DECOY | ITKTQGKK<br>Query 1 ITKTQGKK 8<br>ITKT+GKK<br>Sbjct 1136 ITKTEGKK 1143 | chromosome segregation ATPase [Caudoviricetes sp.]<br>Sequence ID: DAP47510.1Length: 2163 |
| **16** | tr\|S2X7W3\|S2X7W3_9CORY-DECOY | DLALDDVR<br>Query 1 DLALDDVR 8<br>DLAL++VR<br>Sbjct 854 DLALNEVR 861 | hypothetical protein [Caudoviricetes sp.]<br>Sequence ID: DAS72327.1Length: 1446 |

| | | | |
|---|---|---|---|
| **17** | tr\|S4XFJ4\|S4XFJ4_9CORY-DECOY | MGLGIAVTR<br>Query 1 MGLGIAVTR 9<br> MGL IAVTR<br>Sbjct 1061 MGL-IAVTR 1068 | tail tape measure [Caudoviricetes sp.]<br>Sequence ID: DAR12999.1Length: 1909 |
| **18** | tr\|G0HG18\|G0HG18_CORVD-DECOY | AGEVLSGVSK<br>Query 2 GEVLSGVSK 10<br> GEV SGVSK<br>Sbjct 70 GEVVSGVSK 78 | DNA ligase [Caudoviricetes sp.]<br>Sequence ID: DAY95014.1Length: 114 |
| **19** | tr\|S4XJH1\|S4XJH1_9CORY-DECOY | FFSDAKIK<br>Query 2 FSDAKIK 8<br> FSDAKIK<br>Sbjct 220 FSDAKIK 226 | Putative ATP dependent Clp protease [Caudoviricetes sp.]<br>Sequence ID: DAZ10922.1Length: 238 |
| **20** | tr\|B0RIT1\|B0RIT1_CLAMS-DECOY | FELTFNAK<br>Query 1 FELTFNA 7<br> FEL FNA<br>Sbjct 44 FELSFNA 50 | hypothetical protein [Caudoviricetes sp.]<br>Sequence ID: DAX40813.1Length: 172 |
| **21** | tr\|G4QU94\|G4QU94_CORPS-DECOY | HPMMQALK<br>Query 2 PMMQALK 8<br> PMM+ALK<br>Sbjct 435 PMMEALK 441 | tail tape measure [Caudoviricetes sp.]<br>Sequence ID: DAG81818.1Length: 1358 |
| **22** | tr\|S2Y867\|S2Y867_9CORY-DECOY | GTAVFLVLR<br>Query 2 TAVFLVLR 9<br> TAVFLV R<br>Sbjct 77 TAVFLVVR 84 | large terminase [Caudoviricetes sp.]<br>Sequence ID: DAH90722.1Length: 584 |
| **23** | tr\|B0RHZ1\|B0RHZ1_CLAMS-DECOY | TGIRGYLLS<br>Query 1 TGIRGYLLS 9<br> TGI GYLLS<br>Sbjct 416 TGI-GYLLS 423 | Tail tape measure [Caudoviricetes sp.]<br>Sequence ID: DAP99207.1Length: 762 |
| **24** | tr\|B1VGQ0\|B1VGQ0_CORU7-DECOY | GPKLDHISL<br>Query 2 PKLDHISL 9<br> PKL+HIS+<br>Sbjct 344 PKLEHISM 351 | protein of unknown function DUF859 [Caudoviricetes sp.]<br>Sequence ID: DAP02558.1Length: 604 |
| **25** | tr\|S2ZE54\|S2ZE54_9CORY-DECOY | MLAIAGFIK<br>Query 1 MLAIAGFI 8<br> M AIAGFI<br>Sbjct 1243 MVAIAGFI 1250 | tail tape measure [Caudoviricetes sp.]<br>Sequence ID: DAT71817.1Length: 2136 |
| **26** | tr\|I0DIF5\|I0DIF5_CORPS-DECOY | VLDGWVHR<br>Query 2 LDGWVH 7<br> LDGWVH<br>Sbjct 173 LDGWVH 178 | distal tail protein [Caudoviricetes sp.]<br>Sequence ID: DAL15984.1Length: 237 |
| **27** | tr\|D7W983\|D7W983_9CORY-DECOY | FFDKLEGK<br>Query 1 FFDKLEG 7<br> FFDKLEG<br>Sbjct 189 FFDKLEG 195 | Transcriptional regulatory protein RcsB factor, DNA BINDING PROTEIN.6A [Caudoviricetes sp.]<br>Sequence ID: DAT72845.1Length: 196 |
| **28** | tr\|Q8FSG4\|Q8FSG4_COREF-DECOY | VLDLATPVR<br>Query 1 VLDLATPVR 9<br> VLDLAT VR<br>Sbjct 67 VLDLATEVR 75 | tail tube protein [Caudoviricetes sp.]<br>Sequence ID: DAU50773.1Length: 182 |
| **29** | tr\|W5Y277\|W5Y277_9CORY-DECOY | KAPEALLDR<br>Query 2 APEALLDR 9<br> APEALL+R<br>Sbjct 533 APEALLNR 540 | DNA polymerase III, delta subunit+ ATPase, clamp loader, DNA [Caudoviricetes sp.]<br>Sequence ID: DAL18796.1Length: 1251 |
| **30** | tr\|W5XYE9\|W5XYE9_9CORY-DECOY | VGIAAGGFETNSKAFR<br>Query 1 VGIAAGGF 8<br> VGIAAGGF<br>Sbjct 188 VGIAAGGF 195<br><br>Query 7 GFETNSKAFR 16<br> GF+TNSK FR<br>Sbjct 618 GFDTNSKYFR 627 | tail tape measure [Caudoviricetes sp.]<br>Sequence ID: DAT27541.1Length: 1001<br>tail protein [Caudoviricetes sp.]<br>Sequence ID: DAK45322.1Length: 1161 |
| **31** | tr\|H8LVI5\|H8LVI5_CORPS-DECOY | QLAEGGGVFR<br>Query 1 QLAEGGGVFR 10<br> QLAE GVFR<br>Sbjct 73 QLAEAKGVFR 82 | Protein of unknown function (DUF1018) [Caudoviricetes sp.]<br>Sequence ID: DAM57680.1Length: 158 |
| **32** | tr\|G4QRX3\|G4QRX3_CORPS-DECOY | LPLASVSLIK<br>Query 1 LPLASVSLI 9<br> LPLA VSLI<br>Sbjct 35 LPLAHVSLI 43 | tail completion protein [Caudoviricetes sp.]<br>Sequence ID: DAR23790.1Length: 133 |
| **33** | tr\|E0MZA0\|E0MZA0_9CORY-DECOY | IPNEVVADGK<br>Query 1 IPNEVVAD 8<br> IPNEV+AD<br>Sbjct 54 IPNEVIAD 61 | DNA polymerase I [Caudoviricetes sp.]<br>Sequence ID: DAT00868.1Length: 655 |
| **34** | tr\|G4QR48\|G4QR48_CORPS-DECOY | EFGVDLDLR<br>Query 1 EFGVDLDLR 9<br> EFG DLDLR<br>Sbjct 12 EFG-DLDLR 19 | Calcineurin-like phosphoesterase domain, ApaH type [uncultured Caudovirales phage]<br>Sequence ID: CAB5221819.1Length: 284 |
| **35** | tr\|G6WWH7\|G6WWH7_CORGT-DECOY | SFHADIVFK<br>Query 3 HADIVFK 9<br> HA+IVFK<br>Sbjct 387 HAEIVFK 393 | Integrase [Caudoviricetes sp.]<br>Sequence ID: DAK02860.1Length: 395 |
| **36** | tr\|C0E6R2\|C0E6R2_9CORY-DECOY | IPVILLGTDK<br>Query 1 IPVILLGTDK 10<br> IPVIL+G DK | Z1 domain protein [Caudoviricetes sp.]<br>Sequence ID: DAP74530.1Length: 714 |

| | | | |
|---|---|---|---|
| | | Sbjct 621 IPVILMGADK 630 | |
| **37** | tr\|C0XV18\|C0XV18_9CORY-DECOY | FSSAGYQANR<br>Query 1 FSSAGYQANR 10<br>FS AG+QA+R<br>Sbjct 812 FSNAGFQADR 821 | Tail spike protein [Caudoviricetes sp.]<br>Sequence ID: DAU99061.1Length: 826 |
| **38** | tr\|B0RBC6\|B0RBC6_CLAMS-DECOY | LIGAEGIGDEK<br>Query 1 LIGAEGIGDE 10<br>LIGA+GIG++<br>Sbjct 1184 LIGAQGIGNQ 1193 | tail protein [Caudoviricetes sp.]<br>Sequence ID: DAR35222.1Length: 1347 |
| **39** | tr\|E4W9N5\|E4W9N5_RHOE1-DECOY | EGFLVLLASR<br>Query 3 FLVLLASR 10<br>FLVL+ASR<br>Sbjct 283 FLVLMASR 290 | hypothetical protein [Caudoviricetes sp.]<br>Sequence ID: DAH43913.1Length: 491 |
| **40** | tr\|H2GS28\|H2GS28_CORDB-DECOY | LIGKVADLRK<br>Query 1 LIGKVADLRK 10<br>LIGKVADL K<br>Sbjct 630 LIGKVADLIK 639 | minor tail protein [Caudoviricetes sp.]<br>Sequence ID: DAZ17972.1Length: 973 |
| **41** | tr\|M1ULY4\|M1ULY4_9CORY-DECOY | IRSTADPPEK<br>Query 1 IRSTADPPEK 10<br>IR ADPPEK<br>Sbjct 8 IRTVADPPEK 17 | hypothetical protein [Caudoviricetes sp.]<br>Sequence ID: DAK26582.1Length: 254 |
| **42** | tr\|W5Y3J0\|W5Y3J0_9CORY-DECOY | VTLSGIDGIQK<br>Query 2 TLSGIDGIQK 11<br>TL GIDGI K<br>Sbjct 123 TLAGIDGISK 132 | Major capsid protein [Caudoviricetes sp.]<br>Sequence ID: DAV93713.1Length: 290 |
| **43** | tr\|V6V657\|V6V657_CORUL-DECOY | LLTQLVTPVR<br>Query 2 LTQLVTPV 9<br>LTQL+TPV<br>Sbjct 101 LTQLITPV 108 | hypothetical protein, partial [Caudoviricetes sp.]<br>Sequence ID: DAT86367.1Length: 230 |
| **44** | tr\|C3PJH8\|C3PJH8_CORA7-DECOY | IPLINHPNPK<br>Query 4 INHPNPK 10<br>INHPNPK<br>Sbjct 24 INHPNPK 30 | RNA ligase [Caudoviricetes sp.]<br>Sequence ID: DAQ75663.1Length: 462 |
| **45** | tr\|S2X637\|S2X637_9CORY-DECOY | IRSWSGEANK<br>Query 1 IRSWSGEANK 10<br>IR W GE NK<br>Sbjct 1174 IRFWAGEPNK 1183 | Putative tail protein [Caudoviricetes sp.]<br>Sequence ID: DAJ75339.1Length: 1378 |
| **46** | tr\|S2ZZ77\|S2ZZ77_9CORY-DECOY | LPRAIHGVASK<br>Query 2 PRAIHGVAS 10<br>PR IHGVAS<br>Sbjct 7 PRDIHGVAS 15 | virion structural protein [Cyanophage S-TIM54]<br>Sequence ID: UYE97245.1Length: 266 |
| **47** | tr\|E4W8A5\|E4W8A5_RHOE1-DECOY | DGLMAFVSASR<br>Query 1 DGLMAFVSASR 11<br>DGLMA VSASR<br>Sbjct 2031 DGLMAAVSASR 2041 | coil containing protein [Caudoviricetes sp.]<br>Sequence ID: UOF76779.1Length: 3145 |
| **48** | tr\|U5DX65\|U5DX65_COREQ-DECOY | MAAQVGLHKPS<br>Query 1 MAAQVGLHKPS 11<br>MA +GLHKPS<br>Sbjct 229 MA--IGLHKPS 237 | Major capsid protein [Caudoviricetes sp.]<br>Sequence ID: DAV54431.1Length: 282 |
| **49** | tr\|S2XB19\|S2XB19_9CORY-DECOY | MAKGDHAIPAK<br>Query 1 MAKGDHAIPAK 11<br>MAK DHA PAK<br>Sbjct 208 MAKADHA-PAK 217 | Single-stranded DNA binding protein DdrB-like [uncultured Caudovirales phage]<br>Sequence ID: CAB4147292.1Length: 293 |
| **50** | tr\|U7KI84\|U7KI84_9CORY-DECOY | EIASGLGPATSR<br>Query 2 IASGLGPA 9<br>IASGLGPA<br>Sbjct 331 IASGLGPA 338 | tail tape measure [Caudoviricetes sp.]<br>Sequence ID: DAY66114.1Length: 500 |
| **51** | tr\|U3GZA1\|U3GZA1_9CORY-DECOY | QNLAGQEMPR<br>Query 2 NLAGQEM 8<br>NLAGQEM<br>Sbjct 229 NLAGQEM 235 | Replicative helicase [Caudoviricetes sp.]<br>Sequence ID: DAS00986.1Length: 270 |
| **52** | tr\|G0HE73\|G0HE73_CORVD-DECOY | NFIVPVLDSR<br>Query 2 FIVPVLD 8<br>FIVPVLD<br>Sbjct 102 FIVPVLD 108 | portal protein [Caudoviricetes sp.]<br>Sequence ID: DAG16940.1Length: 396 |
| **53** | tr\|W5XXI5\|W5XXI5_9CORY-DECOY | YAVGLLAGELR<br>Query 2 AVGLLAGEL 10<br>AV LLAGEL<br>Sbjct 783 AVALLAGEL 791 | Tail tape measure [Caudoviricetes sp.]<br>Sequence ID: DAU70132.1Length: 1720 |
| **54** | tr\|I7GXU5\|I7GXU5_CORUL-DECOY | ATDPVTLAKKK<br>Query 2 TDPVTLAKK 10<br>TDP TLAKK<br>Sbjct 735 TDPLTLAKK 743 | tail spike [Caudoviricetes sp.]<br>Sequence ID: DAT42168.1Length: 1117 |
| **55** | tr\|C0XQM8\|C0XQM8_9CORY-DECOY | VNALRLNMHK<br>Query 2 NALRLNM 8<br>NALRLNM<br>Sbjct 126 NALRLNM 132 | prohead serine protease [Caudoviricetes sp.]<br>Sequence ID: DAI09015.1Length: 316 |

| | | | |
|---|---|---|---|
| **56** | tr\|H2HXU8\|H2HXU8_CORDW-DECOY | DAVGLVIDPAAR<br>Query 5 LVIDPAAR 12<br>LVIDP AR<br>Sbjct 244 LVIDPGAR 251 | tail protein [Caudoviricetes sp.]<br>Sequence ID: DAM52571.1Length: 300 |
| **57** | tr\|L1MG78\|L1MG78_9CORY-DECOY | VRQRDYHVK<br>Query 1 VRQ-------RDYHV 8<br>VRQ RDYHV<br>Sbjct 281 VRQIAVAILARDYHV 29 | GROUP II INTRON-ENCODED PROTEIN LTRA/RNA, GROUP II INTRONS, RIBONUCLEOPROTEIN.8A [Caudoviricetes sp.] |
| **58** | tr\|T9XP72\|T9XP72_CORDP-DECOY | GDDIVAAFFCR<br>Query 1 GDDIVAAF 8<br>GDDIVAAF<br>Sbjct 49 GDDIVAAF 56 | nucelotide kinase [Caudoviricetes sp.]<br>Sequence ID: DAY90582.1Length: 122 |
| **59** | tr\|I0LJM3\|I0LJM3_CORGK-DECOY | VAALEMWLIR<br>Query 1 VAALEMWLIR 10<br>VAALE WLIR<br>Sbjct 939 VAALEAWLIR 948 | hybrid sensory histidine kinase [Caudoviricetes sp.]<br>Sequence ID: DAM69771.1Length: 950 |
| **60** | tr\|G6WXK6\|G6WXK6_CORGT-DECOY | KGLLAKTDEGAK<br>Query 3 LLAKTDEGAK 12<br>LLAKTDEG K<br>Sbjct 55 LLAKTDEGLK 64 | DNA polymerase III, alpha subunit [Caudoviricetes sp.]<br>Sequence ID: DAP36557.1Length: 882 |
| **61** | tr\|E2S492\|E2S492_9CORY-DECOY | AIASHLALDAEK<br>Query 1 AIASHLALDA 10<br>AIAS LALDA<br>Sbjct 95 AIASVLALDA 104 | Chromatin remodeling complex ATPase [Caudoviricetes sp.]<br>Sequence ID: DAY88015.1Length: 525 |
| **62** | tr\|D9Q8A0\|D9Q8A0_CORP1-DECOY | ELAIVEDGATAR<br>Query 1 ELAIVEDGAT-AR 12<br>ELAI DGAT AR<br>Sbjct 572 ELAI--DGATFAR 582 | minor tail protein [Caudoviricetes sp.]<br>Sequence ID: DAE42057.1Length: 606 |
| **63** | tr\|T9XNB3\|T9XNB3_CORDP-DECOY | FAAHDDVTNGAK<br>Query 2 AAHDDVTNGAK 12<br>AAHDDV N AK<br>Sbjct 81 AAHDDVINKAK 91 | hypothetical protein [Caudoviricetes sp.]<br>Sequence ID: DAI90542.1Length: 335 |
| **64** | tr\|E2MYL3\|E2MYL3_CORAY-DECOY | VFAGLKFDNIK<br>Query 1 VFAGLKFDNI 10<br>VF G KFDNI<br>Sbjct 123 VFNGIKFDNI 132 | Tail tube protein [Caudoviricetes sp.]<br>Sequence ID: DAS99303.1Length: 317 |
| **65** | tr\|L1MM70\|L1MM70_9CORY-DECOY | FKVVTFQSAPK<br>Query 1 FKVVTFQSAPK 11<br>FKVV FQ APK<br>Sbjct 17 FKVV-FQGAPK 26 | tail tube protein [Caudoviricetes sp.]<br>Sequence ID: DAH29105.1Length: 155 |
| **66** | tr\|S5TKN3\|S5TKN3_9CORY-DECOY | SSKLNIMEVAR<br>Query 1 SSKLNIMEVAR 11<br>S KL++ME AR<br>Sbjct 284 SKKLDVMELAR 294 | Integrase [Caudoviricetes sp.]<br>Sequence ID: DAI92560.1Length: 320 |
| **67** | tr\|E9T176\|E9T176_COREQ-DECOY | AEAYLLTAAGQR<br>Query 1 AEAYLLTAAG 10<br>AEAYLL AAG<br>Sbjct 131 AEAYLL-AAG 139 | putative immunoglobulin A1 protease [Caudoviricetes sp.]<br>Sequence ID: DAR13791.1Length: 298 |
| **68** | tr\|F8E0J7\|F8E0J7_CORRG-DECOY | KAKAMLDTTER<br>Query 2 AKAMLDTTER 11<br>AKAM D TER<br>Sbjct 370 AKAMQDATER 379 | Minor structural protein [Caudoviricetes sp.]<br>Sequence ID: DAU03474.1Length: 548 |
| **69** | tr\|M1V0Z3\|M1V0Z3_9CORY-DECOY | VTRFAIDDAQK<br>Query 1 VTRFAIDD 8<br>VTRF IDD<br>Sbjct 35 VTRFEIDD 42 | Integrase [Caudoviricetes sp.]<br>Sequence ID: DAT15469.1Length: 391 |
| **70** | tr\|Q8FSW1\|Q8FSW1_COREF-DECOY | LIGVYRAPKTR<br>Query 3 GVYRAPKTR 11<br>G YRAPKTR<br>Sbjct 159 GGYRAPKTR 167 | repressor domain protein [Caudoviricetes sp.]<br>Sequence ID: DAW40519.1Length: 304 |
| **71** | tr\|E9SXE6\|E9SXE6_COREQ-DECOY | RDHHGLAGTGAGK<br>Query 6 LAGTGAGK 13<br>LAGTGAGK<br>Sbjct 275 LAGTGAGK 282 | tail spike protein [Caudoviricetes sp.]<br>Sequence ID: DAM64846.1Length: 771 |
| **72** | tr\|U3GYZ5\|U3GYZ5_9CORY-DECOY | AEMAIGATAALSR<br>Query 1 AEMAIGATAAL 11<br>A MA+GATAA+<br>Sbjct 63 AKMAVGATAAM 73 | minor tail protein [Caudoviricetes sp.]<br>Sequence ID: DAT47304.1Length: 829 |
| **73** | tr\|D5UTX9\|D5UTX9_TSUPD-DECOY | DPSQSKVLIHR<br>Query 1 DPSQSKVLI 9<br>DPSQS VLI<br>Sbjct 295 DPSQSNVLI 303 | hypothetical protein [Caudoviricetes sp.]<br>Sequence ID: DAQ58918.1Length: 621 |
| **74** | tr\|D8KK67\|D8KK67_CORPF-DECOY | VLRKRSDDYK<br>Query 3 RKRSDDYK 10<br>RKR DDYK<br>Sbjct 325 RKRNDDYK 332 | portal [Caudoviricetes sp.]<br>Sequence ID: DAL87048.1Length: 436 |
| **75** | tr\|H2HF08\|H2HF08_CORDJ-DECOY | TYRGMAWILR<br>Query 1 TYRGMAWI 8<br>TYR MAWI<br>Sbjct 491 TYRKMAWI 498 | H-type lectin domain [Caudoviricetes sp.]<br>Sequence ID: DAM83510.1Length: 503 |

| | | | |
|---|---|---|---|
| **76** | tr\|D7W9C3\|D7W9C3_9CORY-DECOY | NTVTDRSGIGIR<br>Query 2 TVTDRSG-IGIR 12<br>TVT+ SG IGIR<br>Sbjct 465 TVTNPSGHIGIR 476 | nuclear pore complex protein [Caudoviricetes sp.]<br>Sequence ID: DAG34037.1Length: 576 |
| **77** | tr\|U7K0J0\|U7K0J0_9CORY-DECOY | NELRMGVVQVK<br>Query 1 NEL-RMGV-VQV 10<br>NEL RMGV +QV<br>Sbjct 89 NELIRMGVLIQV 100 | replication protein O [Caudoviricetes sp.]<br>Sequence ID: DAP08825.1Length: 157 |
| **78** | tr\|U7LBE0\|U7LBE0_9CORY-DECOY | ANLMEGGPDILR<br>Query 3 LMEGGPDILR 12<br>L EGGPD+LR<br>Sbjct 259 LVEGGPDVLR 268 | DNA primase, catalytic core [Caudoviricetes sp.]<br>Sequence ID: DAY94238.1Length: 1046 |
| **79** | tr\|C0WEY2\|C0WEY2_9CORY-DECOY | GGTTVVTTGPEMR<br>Query 1 GGTTVVTTGPEM 12<br>GGTTV T PEM<br>Sbjct 289 GGTTVLTEVPEM 300 | D-galactarate dehydratase / Altronate hydrolase [Caudoviricetes sp.]<br>Sequence ID: DAT44869.1Length: 494 |
| **80** | tr\|E4WI84\|E4WI84_RHOE1-DECOY | GVGITELGADARM<br>Query 2 VGITELGADARM 13<br>+GITELG D RM<br>Sbjct 345 IGITELG-D-RM 354 | DNA polymerase II small subunit [Caudoviricetes sp.]<br>Sequence ID: DAG70758.1Length: 398 |
| **81** | tr\|C8RTN2\|C8RTN2_CORJE-DECOY | ASELRDALQMR<br>Query 2 SELRDALQMR 11<br>SELR+ LQMR<br>Sbjct 129 SELRERLQMR 138 | major capsid protein [Caudoviricetes sp.]<br>Sequence ID: DAL80727.1Length: 428 |
| **82** | tr\|I7GX13\|I7GX13_CORUL-DECOY | QVEEVGDARFR<br>Query 1 QVEE------VGDARFR 11<br>QVEE V DARFR<br>Sbjct 391 QVEEFFRRTSVNDARFR 407 | Hsp70-like protein [Caudoviricetes sp.]<br>Sequence ID: DAL78329.1Length: 591 |
| **83** | tr\|W5WXZ3\|W5WXZ3_9CORY-DECOY | NGLPAINALVPVK<br>Query 1 NGLPAINALV 10<br>N LPA+NALV<br>Sbjct 1410 NSLPAVNALV 1419 | tail tape measure [Caudoviricetes sp.]<br>Sequence ID: DAM61551.1Length: 2330 |
| **84** | tr\|H2GLY2\|H2GLY2_CORDB-DECOY | VLPFGTGGVPDPR<br>Query 2 LPFGTGGVPDPR 13<br>L FGTGG P PR<br>Sbjct 183 LAFGTGGDPAPR 194 | hypothetical protein [Caudoviricetes sp.]<br>Sequence ID: DAT46572.1Length: 415 |
| **85** | tr\|G4QTB2\|G4QTB2_CORPS-DECOY | EGTQIDDIGSASK<br>Query 4 QIDDIGS 10<br>QIDDIGS<br>Sbjct 327 QIDDIGS 333 | Portal protein [Caudoviricetes sp.],<br>Sequence ID: DAK51282.1Length: 778 |
| **86** | tr\|E3F770\|E3F770_CORP9-DECOY | EAAHAALIGVLLR<br>Query 1 EAAHAALI-------GV---LLR 13<br>EAAHAAL+ GV LLR<br>Sbjct 105 EAAHAALVREVAKETGVPADLLR 127 | Major head protein [Caudoviricetes sp.]<br>Sequence ID: DAJ35944.1Length: 178 |
| **87** | tr\|G6WZL3\|G6WZL3_CORGT-DECOY | ISVTTKEIADLR<br>Query 1 ISVTTKEIADLR 12<br>I TTKE ADLR<br>Sbjct 8 IKLTTKEYADLR 19 | transposase [Caudoviricetes sp.]<br>Sequence ID: DAT66202.1Length: 654 |
| **88** | tr\|X5DLL0\|X5DLL0_9CORY-DECOY | RMELTAGLSVIR<br>Query 1 RMELTAG 7<br>RMELTAG<br>Sbjct 333 RMELTAG 339 | tail protein [Caudoviricetes sp.]<br>Sequence ID: DAR13707.1Length: 351 |
| **89** | tr\|U7K1V4\|U7K1V4_9CORY-DECOY | DAASIFSAGARGVK<br>Query 1 DAA-SIFSAGARGVK 14<br>DAA S+FSA GVK<br>Sbjct 337 DAADSVFSAVSSGVK 351 | tail tape measure protein [Caudoviricetes sp.]<br>Sequence ID: DAJ27099.1Length: 690 |
| **90** | tr\|G2ENE4\|G2ENE4_CORGT-DECOY | TLQVALAPWTPR<br>Query 7 APWTPR 12<br>APWTPR<br>Sbjct 60 APWTPR 65 | replicative DNA helicase [Caudoviricetes sp.]<br>Sequence ID: DAW79410.1Length: 520 |
| **91** | tr\|U5DX50\|U5DX50_COREQ-DECOY | AKFQETLVDTGR<br>Query 3 FQETLVDTG 11<br>FQETLVD G<br>Sbjct 526 FQETLVDNG 534 | large terminase [Caudoviricetes sp.]<br>Sequence ID: DAV01846.1Length: 621 |
| **92** | tr\|H2GSR9\|H2GSR9_CORDB-DECOY | EAFFLTAAPAQAK<br>Query 1 EAFFLTAAPAQ 11<br>E FF T APAQ<br>Sbjct 79 ETFFVTSAPAQ 89 | tail protein [Caudoviricetes sp.]<br>Sequence ID: DAL59492.1Length: 326 |
| **93** | tr\|E2MWU2\|E2MWU2_CORAY-DECOY | TIVWPVTLKSAR<br>Query 2 IVWPVTL 8<br>I+WPVTL<br>Sbjct 125 IIWPVTL 131 | Lambda phage tail tube protein, TTP [Caudoviricetes sp.]<br>Sequence ID: DAQ78446.1Length: 166 |
| **94** | tr\|X5EFV7\|X5EFV7_9CORY-DECOY | AAAVLFDPGLLKR<br>Query 4 VLFDPGLLKR 13<br>VLFD GLL R<br>Sbjct 159 VLFDSGLLSR 168 | ABC-type sugar transport system, periplasmic component [Caudoviricetes sp.]<br>Sequence ID: DAP67153.1Length: 427 |
| **95** | sp\|Q8NRM3\|EX7L_CORGL-DECOY | ACDPGAYMLIALR<br>Query 1 ACDPGAYML 9<br>ACDPGAY L<br>Sbjct 700 ACDPGAYQL 708 | ATP dependent DNA helicase [Siphoviridae sp. ctnpt50]<br>Sequence ID: DAF48925.1Length: 780 |

| | | | |
|---|---|---|---|
| **96** | tr\|H2H0A0\|H2H0A0_CORDD-DECOY | GDWILFFPQDR<br>Query 2 DWIL-FFPQD 10<br>DWIL FFP++<br>Sbjct 56 DWILSFFPEN 65 | hypothetical protein [Caudoviricetes sp.]<br>Sequence ID: DAH19474.1Length: 447 |
| **97** | tr\|D5NVU5\|D5NVU5_CORAM-DECOY | EISQMEFFDVR<br>Query 3 SQMEFFDVR 11<br>SQMEFF+ R<br>Sbjct 346 SQMEFFEAR 354 | hypothetical protein UFOVP817_17 [uncultured Caudovirales phage]<br>Sequence ID: CAB4164959.1Length: 687 |
| **98** | tr\|H2HTD4\|H2HTD4_CORDL-DECOY | QQSMLGDGVNVMK<br>Query 2 QSMLGDG----VNV 11<br>QSMLGDG VN+<br>Sbjct 96 QSMLGDGYNLEVNI 109 | tail protein [Caudoviricetes sp.]<br>Sequence ID: DAL06528.1Length: 177 |
| **99** | tr\|C3PJL3\|C3PJL3_CORA7-DECOY | RLSTYPKEGGYR<br>Query 1 RLSTYP-KEGG 10<br>RLSTYP K+GG<br>Sbjct 74 RLSTYPGKQGG 84 | hypothetical protein [Caudoviricetes sp.]<br>Sequence ID: DAK92684.1Length: 257 |
| **100** | tr\|G0HAF6\|G0HAF6_CORVD-DECOY | DNRDNGIDMMLK<br>Query 1 DNRDNGIDM 9<br>+NRDNG DM<br>Sbjct 69 NNRDNGADM 77 | hypothetical protein [Caudoviricetes sp.]<br>Sequence ID: DAX13196.1Length: 287 |
| **101** | tr\|H6M7H8\|H6M7H8_CORPS-DECOY | ARPAWTFAIKVAK<br>Query 2 RPAWTFAI 9<br>RPAWTF I<br>Sbjct 35 RPAWTFTI 42 | hypothetical protein, partial [Caudoviricetes sp.]<br>Sequence ID: DAJ22912.1Length: 43 |
| **102** | tr\|C0VT36\|C0VT36_9CORY-DECOY | FAGGDAASVTRETGK<br>Query 3 GGDAASVTR 11<br>GGDAA VTR<br>Sbjct 13 GGDAANVTR 21 | Nucleoside deoxyribosyltransferase [Caudoviricetes sp.]<br>Sequence ID: DAT45698.1Length: 117 |
| **103** | tr\|G2EM91\|G2EM91_CORGT-DECOY | APSNVTTALDDILR<br>Query 2 PS----NVTTALDDI 12<br>PS NVTT LDDI<br>Sbjct 474 PSKKWDNVTTTLDDI 488 | DNA helicase [Caudoviricetes sp.]<br>Sequence ID: DAF25786.1Length: 903 |
| **104** | tr\|U7LJI8\|U7LJI8_9CORY-DECOY | RRSFPTLFKVLK<br>Query 3 SFPTLFKVLK 12<br>S+PTLFK LK<br>Sbjct 424 SYPTLFKALK 433 | hypothetical protein [Caudoviricetes sp.]<br>Sequence ID: DAM88417.1Length: 481 |
| **105** | tr\|U7LKW5\|U7LKW5_9CORY-DECOY | LIVLLIWAVAELR<br>Query 1 LIVLLIWAV 9<br>+IVLLIWA+<br>Sbjct 17 MIVLLIWAI 25 | hypothetical protein [Caudoviricetes sp.]<br>Sequence ID: DAQ18890.1Length: 230 |
| **106** | tr\|C6R6G9\|C6R6G9_9CORY-DECOY | RDEAGEEGTMFKK<br>Query 7 EGTMFKK 13<br>EGTMFKK<br>Sbjct 79 EGTMFKK 85 | homing endonuclease [Caudoviricetes sp.]<br>Sequence ID: DAT03275.1Length: 201 |
| **107** | tr\|A4QE00\|A4QE00_CORGB-DECOY | KVSGKLAIGEVPPIK<br>Query 3 SGK---LAIGEVPPIK 15<br>SGK L IG VPPIK<br>Sbjct 74 SGKSGVLSIGGVPPIK 89 | Lysin motif [Caudoviricetes sp.]<br>Sequence ID: DAM50216.1Length: 176 |
| **108** | tr\|H2HM59\|H2HM59_CORDK-DECOY | DSAEETIIAMGGAGSK<br>Query 1 DSAEETIIAM 10<br>D AE+ IIAM<br>Sbjct 2164 DAAEDSIIAM 2173 | nuclease [Caudoviricetes sp.]<br>Sequence ID: DAG63193.1Length: 2973 |
| **109** | tr\|G7U361\|G7U361_CORPS-DECOY | GSEEMPASFQANLR<br>Query 2 SEEM--PASFQANLR 14<br>SEEM PA F A LR<br>Sbjct 171 SEEMEHPAAFAAPLR 185 | portal [Caudoviricetes sp.]<br>Sequence ID: DAQ54312.1Length: 419 |
| **110** | tr\|W5Y3S8\|W5Y3S8_9CORY-DECOY | RLDTTLDAVGAAPEK<br>Query 1 RLDTTLDAVGAAPEK 15<br>RLD TL A GA EK<br>Sbjct 123 RLDKTLSAGGASAEK 137 | tail tape measure [Caudoviricetes sp.]<br>Sequence ID: DAY80389.1Length: 1304 |
| **111** | tr\|C4LKG7\|C4LKG7_CORK4-DECOY | VTSGIMYNLGNVAAR<br>Query 1 VTSGIM-YNLGNVA 13<br>V GI YNLGNVA<br>Sbjct 1355 VSTGIINYNLGNVA 1368 | ATPase [Caudoviricetes sp.]<br>Sequence ID: DAK25262.1Length: 2766 |
| **112** | tr\|H2I2H6\|H2I2H6_CORDW-DECOY | VTVWRAPREKGNR<br>Query 3 VWRA--PREKGNR 13<br>+WRA PR+ GNR<br>Sbjct 1702 IWRANIPRDNGNR 1714 | stabilization protein [Caudoviricetes sp.]<br>Sequence ID: DAW72376.1Length: 1747 |
| **113** | tr\|U7KAB9\|U7KAB9_9CORY-DECOY | QGPAFQLKPEPQLK<br>Query 4 AFQLKPEPQ 12<br>AFQ KPEPQ<br>Sbjct 89 AFQ-KPEPQ 96 | zinc-ribbon family protein [Caudoviricetes sp.]<br>Sequence ID: DAM38746.1Length: 232 |
| **114** | tr\|S5SS46\|S5SS46_9CORY-DECOY | LALVLSVTVDTLGGAR<br>Query 6 SVTVDTLGGA 15<br>SV VDTLGGA<br>Sbjct 54 SVKVDTLGGA 63 | repressor domain protein [Caudoviricetes sp.]<br>Sequence ID: DAM55883.1Length: 257 |
| **115** | tr\|E0DDF4\|E0DDF4_9CORY-DECOY | LIAEDMVMRIKLR<br>Query 5 DMVMRIKLR 13<br>DMV+RIKLR<br>Sbjct 55 DMVLRIKLR 63 | large terminase [Caudoviricetes sp.]<br>Sequence ID: DAR61636.1Length: 320 |
| **116** | tr\|D7WDG4\|D7WDG4_9CORY-DECOY | QMKGLDTVLAITMR<br>Query 2 MKGLDTVLAITM 13<br>MK DTVLA TM<br>Sbjct 225 MKEFDTVLA-TM 235 | minor capsid protein [Caudoviricetes sp.]<br>Sequence ID: DAJ51952.1Length: 285 |

| | | | |
|---|---|---|---|
| **117** | tr\|A4QB75\|A4QB75_CORGB-DECOY | LASMAEMQSGSIGQAK<br>Query 2 ASMAEMQS--GSIG 13<br>ASMA+MQS G IG<br>Sbjct 240 ASMADMQSIFGKIG 253 | minor tail protein [Caudoviricetes sp.]<br>Sequence ID: DAQ37392.1Length: 906 |
| **118** | tr\|A4QD73\|A4QD73_CORGB-DECOY | SGSIWALTASGLVDTR<br>Query 3 SI--WALTASGLV 13<br>SI WA TA GLV<br>Sbjct 3 SIRYWAITAAGLV 15 | holin [Caudoviricetes sp.]<br>Sequence ID: DAJ05651.1Length: 139 |
| **119** | tr\|B4F3E5\|B4F3E5_RHOE1-DECOY | AVANMDNIRPLYTR<br>Query 2 VANMDNI---------RPLYT 13<br>VANMDNI P YT<br>Sbjct 326 VANMDNINDITVGDGSKPVYT 346 | Baseplate J like protein [Caudoviricetes sp.]<br>Sequence ID: DAO39777.1Length: 366 |
| **120** | tr\|E4WCV8\|E4WCV8_RHOE1-DECOY | LIKLNRYGCEDVGR<br>Query 1 LIKLNRYGC-EDVGR 14<br>L+KLN Y C ED GR<br>Sbjct 394 LVKLNMYECIEDLGR 408 | portal protein [Caudoviricetes sp.]<br>Sequence ID: DAU96467.1Length: 560 |
| **121** | tr\|M1MZE1\|M1MZE1_9CORY-DECOY | LIGESPAGARHLPVPR<br>Query 3 GESPAGARHLPVP 15<br>G+ PAGARHLP P<br>Sbjct 82 GQRPAGARHLPPP 94 | hypothetical protein [Caudoviricetes sp.]<br>Sequence ID: DAV79588.1Length: 145 |
| **122** | tr\|U7LGT2\|U7LGT2_9CORY-DECOY | FGGQIAVDIGGILAANR<br>Query 2 GGQIAV-DIGGILAA 15<br>GGQ AV I GILAA<br>Sbjct 388 GGQMAVLAIAGILAA 402 | minor tail protein [Caudoviricetes sp.]<br>Sequence ID: DAZ60126.1Length: 799 |
| **123** | tr\|B1VI70\|B1VI70_CORU7-DECOY | VREFMRTGELLLSK<br>Query 4 FMRTGELLLSK 14<br>FMR ELLLSK<br>Sbjct 112 FMR--ELLLSK 120 | Protein of unknown function (DUF935) [Caudoviricetes sp.]<br>Sequence ID: DAZ58951.1Length: 434 |
| **124** | tr\|M1UZG5\|M1UZG5_9CORY-DECOY | KLREDKPQNAPAAVR<br>Query 2 LREDKPQNAPA 12<br>LR DKPQ APA<br>Sbjct 185 LRVDKPQEAPA 195 | Rad52/22 family double-strand break repair protein [Caudoviricetes sp.]<br>Sequence ID: DAJ90329.1 |
| **125** | tr\|T9WSD4\|T9WSD4_CORDP-DECOY | ALGDNTINLEPVVPLK<br>Query 1 ALGDNTINLEPVVP 14<br>ALG TI+L+PVVP<br>Sbjct 480 ALG-KTIDLDPVVP 492 | helicase [Gordonia phage Gmala1]<br>Sequence ID: YP_009215298.1Length: 672 |
| **126** | tr\|E2S1G5\|E2S1G5_9CORY-DECOY | LLATRGGGAPVPWLER<br>Query 5 RGGGAPVP 12<br>RGGGAPVP<br>Sbjct 204 RGGGAPVP 211 | Baseplate wedge protein, partial [Caudoviricetes sp.]<br>Sequence ID: DAQ03201.1Length: 214 |
| **127** | tr\|E4WFZ0\|E4WFZ0_RHOE1-DECOY | LLMLTLIISEFDIR<br>Query 4 LTLIISEFDI 13<br>LTLII EF+I<br>Sbjct 94 LTLIIPEFNI 103 | tail assembly protein [Caudoviricetes sp.]<br>Sequence ID: DAW80849.1Length: 153 |
| **128** | tr\|H2GP34\|H2GP34_CORDB-DECOY | LQVVHNMVARKSRR<br>Query 1 LQVVHNMV 8<br>LQVVH+MV<br>Sbjct 303 LQVVHDMV 310 | Head-to-tail connector protein, podovirus-type [uncultured Caudovirales phage]<br>Sequence ID: CAB4213427.1Length: 557 |
| **129** | tr\|S5TKS2\|S5TKS2_9CORY-DECOY | MGSGMYYLGAGARNDK<br>Query 6 YYLGAGARNDK 16<br>YY GAGARN K<br>Sbjct 102 YYKGAGARNGK 112<br><br>Query 1 MGSGMYYLG 9<br>M GMYYLG<br>Sbjct 747 MNTGMYYLG 755 | C-5 cytosine-specific DNA methylase [Caudoviricetes sp.]<br>Sequence ID: DAR98136.1Length: 230<br><br>dsDNA helicase [Caudoviricetes sp.]<br>Sequence ID: DAY23848.1Length: 768 |
| **30** | tr\|Q8FNX8\|Q8FNX8_COREF-DECOY | LNLEEDMADQLSSSR<br>Query 1 LNLEEDMAD 9<br>L+LEEDMA+<br>Sbjct 1327 LDLEEDMAE 1335<br><br>Query 1 LNLEEDMAD 9<br>L+LEEDMA+<br>Sbjct 2179 LDLEEDMAE 2187 | CRISPR-associated exonuclease [Crassvirales sp.]<br>Sequence ID: DAH01940.1Length: 1485<br><br>Structural protein [Caudoviricetes sp.]<br>Sequence ID: DAM56159.1Length: 2387 |
| **131** | tr\|D7WD43\|D7WD43_9CORY-DECOY | DNMLLGKSMPFLAIR<br>Query 4 LLGKSMPFLA 13<br>LLGKSMP LA<br>Sbjct 382 LLGKSMPSLA 391 | minor capsid protein [Caudoviricetes sp.]<br>Sequence ID: DAX95766.1Length: 514 |
| **132** | tr\|B0RID6\|B0RID6_CLAMS-DECOY | GGRRSTHHLLADAVNR<br>Query 3 RRSTHHLLADAVNR 16<br>R THHLLA AV R<br>Sbjct 9 RKRTHHLLAKAVQR 22 | hypothetical protein [Caudoviricetes sp.]<br>Sequence ID: DAT75162.1Length: 100 |
| **133** | tr\|G2EL51\|G2EL51_CORGT-DECOY | DLWGGDLILEFGRLR<br>Query 1 DL--WGGDLIL 9<br>DL WGGDLIL<br>Sbjct 42 DLFRWGGDLIL 52 | hypothetical protein [Caudoviricetes sp.]<br>Sequence ID: DAW64243.1Length: 149 |
| **134** | tr\|S5T5Y4\|S5T5Y4_9CORY-DECOY | MSIPRSADFPIGALIR<br>Query 5 RSADFPIG 12<br>RSADFPIG<br>Sbjct 431 RSADFPIG 438 | Thymidylate synthase [Caudoviricetes sp.]<br>Sequence ID: DAJ88739.1Length: 528 |

<table>
<tr><td>135</td><td>tr|U7L034|U7L034_9CORY-DECOY</td><td>LITDEDVCLIPFLIR<br>Query  1   LITDEDVCLIPFLI  14<br>            LIT+E  CLIP LI<br>Sbjct  24  LITEEG-CLIPLLI  36</td><td>hypothetical protein [Caudoviricetes sp.]<br>Sequence ID: DAT35064.1Length: 43</td></tr>
<tr><td>136</td><td>tr|H2HPN4|H2HPN4_CORDK-DECOY</td><td>LSLAGAAIGCLFAGCHLR<br>Query  3   LAGAAIGCLFAG  14<br>            +AGAAIGCL AG<br>Sbjct  71  MAGAAIGCL-AG  81</td><td>putative periplasmic lipoprotein [Caudoviricetes sp.]<br>Sequence ID: DAH74711.1Length: 87</td></tr>
<tr><td>137</td><td>tr|G7U0B1|G7U0B1_CORPS-DECOY</td><td>MSGDCEAEWRKTIGLR<br>Query  8    EWRKTIGLR  16<br>             EWRK IGLR<br>Sbjct  105  EWRKKIGLR  113</td><td>HOLLIDAY JUNCTION RESOLVASE [Caudoviricetes sp.]<br>Sequence ID: DAW02892.1Length: 168</td></tr>
<tr><td>138</td><td>tr|G0CSR3|G0CSR3_CORUL-DECOY</td><td>EASEGSQLGGSSVDLMRR<br>Query  4   EGSQLGGSSV  13<br>            +GSQLGGSSV<br>Sbjct  31  DGSQLGGSSV  40</td><td>holin [Caudoviricetes sp.]<br>Sequence ID: DAP78348.1Length: 55</td></tr>
<tr><td>139</td><td>tr|D9QDW6|D9QDW6_CORP2-DECOY</td><td>WKLGVNARILYSGISER<br>Query  4    GVNARILYSGISE  16<br>             G NARILY  I E<br>Sbjct  143  GLNARILYKEICE  155</td><td>DnaB-like replicative helicase [Caudoviricetes sp.]<br>Sequence ID: DAI57689.1Length: 220</td></tr>
<tr><td>140</td><td>tr|G6WUI5|G6WUI5_CORGT-DECOY</td><td>NPRKIEQGDITNVGLAVR<br>Query  4   KIEQG--DITNVGLAVR  18<br>            KIE+G  DIT V  AVR<br>Sbjct  13  KIEEGKLDITAVXNAVR  29</td><td>minor structural protein [Caudoviricetes sp.]<br>Sequence ID: DAI01584.1Length: 205</td></tr>
<tr><td>141</td><td>tr|B1VDU1|B1VDU1_CORU7-DECOY</td><td>VAELVWLIELGKTHTDR<br>Query  4    VWLI--ELGKTHT  14<br>             VWL+  E GKTHT<br>Sbjct  208  VWLVGEEFGKTHT  220</td><td>UDP-N-acetylmuramyl pentapeptide synthase, partial [Caudoviricetes sp.]<br>Sequence ID: DAZ13990.1Length: 262</td></tr>
<tr><td>142</td><td>tr|U7L2Q8|U7L2Q8_9CORY-DECOY</td><td>AGDAVDDAITKPVNRGILR<br>Query  3    DAVDDAIT--KPVNRGILR  19<br>             DA D+AI   K V R ILR<br>Sbjct  658  DAQDEAISSAKQVSRNILR  676</td><td>tail tape measure protein [Caudoviricetes sp.]<br>Sequence ID: DAK96220.1Length: 738</td></tr>
<tr><td>143</td><td>tr|W5WSX2|W5WSX2_9CORY-DECOY</td><td>LYAASNREESTQQNNLR<br>Query  2     YAAS--NREESTQ---QNNLR  17<br>              YA S  N EESTQ   + NLR<br>Sbjct  1220  YARSSYNKEESTQSTIEQNLR  1240</td><td>DNA directed DNA polymerase [Caudoviricetes sp.]<br>Sequence ID: DAV70585.1Length: 1332</td></tr>
<tr><td>144</td><td>tr|B0RIH5|B0RIH5_CLAMS-DECOY</td><td>AQRCIMSLRYGSAPDPR<br>Query  3   RCI-------MSLRYGSAPDP  16<br>            RC+       MSLRY   PDP<br>Sbjct  44  RCVICGQPVDMSLRY---PDP  61</td><td>HNH endonuclease [Caudoviricetes sp.]<br>Sequence ID: DAX37516.1Length: 112</td></tr>
<tr><td>145</td><td>tr|U2G0Y6|U2G0Y6_9CORY-DECOY</td><td>AQLDEQALAVQGGKVVDVR<br>Query  6   QALAVQGGKVVDV  18<br>            +ALAV GGKVV++<br>Sbjct  44  HALAVRGGKVVEI  56</td><td>hypothetical protein [Caudoviricetes sp.]<br>Sequence ID: DAI66881.1Length: 73</td></tr>
<tr><td>146</td><td>tr|U7K8Q0|U7K8Q0_9CORY-DECOY</td><td>MSLFVMCVASVAEIKLVR<br>Query  1   MSLFV--MCVAS  10<br>            MSLFV  MC AS<br>Sbjct  27  MSLFVPDMCAAS  38</td><td>hypothetical protein [Caudovirales sp. ct0jG3]<br>Sequence ID: DAD97578.1Length: 147</td></tr>
<tr><td>147</td><td>tr|U7L9F2|U7L9F2_9CORY-DECOY</td><td>SIIARGMPYAVDPGEARAR<br>Query  6    GMPYAVD  12<br>             GMPYAVD<br>Sbjct  243  GMPYAVD  249<br><br>Query  5    RGMPYAVDPGE  15<br>             R +PY V+PGE<br>Sbjct  134  RALPYSVEPGE  144</td><td>major capsid protein [Caudoviricetes sp.]<br>Sequence ID: DAD63098.1Length: 295<br>DNA polymerase [Caudoviricetes sp.]<br>Sequence ID: DAM37670.1Length: 1258</td></tr>
<tr><td>148</td><td>tr|G2EQW6|G2EQW6_CORGT-DECOY</td><td>AQAMAASPRFMVTPPGYIK<br>Query  3    AMAASPRFMV  12<br>             AMAA PR MV<br>Sbjct  239  AMAAAPRLMV  248</td><td>DNA polymerase [Caudoviricetes sp.]<br>Sequence ID: DAV81210.1Length: 768Number of Matches: 1</td></tr>
<tr><td>149</td><td>tr|R0JSR1|R0JSR1_CORCT-DECOY</td><td>VATISWFEEMERPSLEK<br>Query  5    SWFEEME  11<br>             SWFEEME<br>Sbjct  100  SWFEEME  106</td><td>nucleotidase 5'-nucleotidase [Caudoviricetes sp.]<br>Sequence ID: DAJ35661.1Length: 112</td></tr>
<tr><td>150</td><td>tr|E9SY32|E9SY32_COREQ-DECOY</td><td>AEVVGSMHDRCSPAGGLIIR<br>Query  1   AEVVGSMHDRC  11<br>            AEVVG MH RC<br>Sbjct  11  AEVVGRMH-RC  20</td><td>Catabolite control protein A [Caudoviricetes sp.]<br>Sequence ID: DAU94518.1Length: 85</td></tr>
<tr><td>151</td><td>tr|C2CT76|C2CT76_CORST-DECOY</td><td>VLMATTMLTLFPQEEIAR<br>Query  7    MLTLFPQEEIA  17<br>             MLT+FP+E+IA<br>Sbjct  212  MLTMFPHEQIA  222</td><td>DNA adenine methylase [Siphoviridae sp. ctJ0s2]<br>Sequence ID: DAF61540.1Length: 267</td></tr>
<tr><td>152</td><td>tr|H2HY90|H2HY90_CORDW-DECOY</td><td>EMATASVMAGPAESVDIMIK<br>Query  2    MATASV----MAGPAESV  15<br>             MATA V    M GPAE V<br>Sbjct  182  MATA-VDGVTMEGPAETV  198</td><td>hypothetical protein [Siphoviridae sp. ctS248]<br>Sequence ID: DAJ21778.1Length: 288</td></tr>
<tr><td>153</td><td>tr|I0LGX5|I0LGX5_CORGK-DECOY</td><td>SENDPDKRLESSPDLLLR<br>Query  2   ENDPDKRLESSPD  14<br>            +NDPD RL+  PD<br>Sbjct  74  DNDPDNRLDAIPD  86</td><td>Repressor protein CI [Caudoviricetes sp.]<br>Sequence ID: DAP26097.1Length: 264</td></tr>
<tr><td>154</td><td>tr|G7U3E6|G7U3E6_CORPS-DECOY</td><td>VTKANPGHTQAQHHADSSAR<br>Query  7    GHTQAQHHADSSAR  20<br>             G TQA H ADSS R<br>Sbjct  181  GNTQADH-ADSSSR  193</td><td>hypothetical protein [Caudoviricetes sp.]<br>Sequence ID: DAJ31284.1Length: 212</td></tr>
<tr><td>155</td><td>tr|B0RD18|B0RD18_CLAMS-DECOY</td><td>TMPMPSYNYERVTLIFR<br>Query  7   YNYERVTLIF  16<br>            YN ERVTLIF</td><td>head closure knob [Caudoviricetes sp.]</td></tr>
</table>

| | | | |
|---|---|---|---|
| | | Sbjct 3 YN-ERVTLIF 11 | Sequence ID: DAY99500.1Length: 92 |
| **156** | tr\|W5WND6\|W5WND6_9CORY-DECOY | CPLARFVMYLVLASVSLMR<br>Query 1 CPLARFVMYLVLASVSL 17<br>CP FV YLVL +SL<br>Sbjct 48 CP--PFVIYLVLTTISL 62 | hypothetical protein [Caudoviricetes sp.]<br>Sequence ID: DAX55864.1Length: 125 |
| **157** | tr\|M1NUM7\|M1NUM7_9CORY-DECOY | LNGPAPADACIMRLKLVHLR<br>Query 8 DACIMRLK 15<br>D CIMRLK<br>Sbjct 183 DGCIMRLK 190 | endonuclease [Siphoviridae sp. ctZHD14]<br>Sequence ID: DAF55219.1Length: 264 |
| **158** | tr\|G0CS91\|G0CS91_CORUL-DECOY | MIFEACKDTPHLFPASGMR<br>Query 3 FE-ACKDTPHLFPAS 16<br>FE AC DTP L PAS<br>Sbjct 17 FETACRDTPTL-PAS 30 | hypothetical protein [Caudoviricetes sp.]<br>Sequence ID: DAY23903.1Length: 51 |
| **159** | tr\|E2S1R4\|E2S1R4_9CORY-DECOY | DFSSLHKAADVRGDSEPGVPR<br>Query 5 LHKAADVRGDSEPGVP 20<br>LHKA D RG EPG+P<br>Sbjct 176 LHKARD-RGGDEPGIP 190 | DnaB-like replicative helicase [Caulobacter phage Sansa]<br>Sequence ID: YP_009785471.1Length: 506 |
| **160** | tr\|A4QGL8\|A4QGL8_CORGB-DECOY | EPSNQMDQEFTTDRTTVTR<br>Query 3 SNQMDQ----EFTTDRTT 16<br>SN++D EFTTD TT<br>Sbjct 634 SNELDASHTLEFTTDKTT 651 | Interferon alpha/beta receptor 1, Interferon I interferon signaling complex [Caudoviricetes sp.]<br>Sequence ID: DAL43843.1Length: 705<br>A hot spot on interferon α/β receptor subunit 1 (IFNAR1) underpins its interaction with interferon-β and dictates signaling PMID: 28289093 |
| **161** | tr\|F8E2W5\|F8E2W5_CORRG-DECOY | ANPVGDSTGVHVDLMQHIAAQR<br>Query 1 ANPVGDS-TGVHV 12<br>ANPVGDS TG HV<br>Sbjct 168 ANPVGDSETG-HV 179<br><br>Query 12 VD-LMQHIAAQR 22<br>VD LMQ +AAQR<br>Sbjct 457 VDELMQEVAAQR 468 | Protein of unknown function (DUF1366) [Caudoviricetes sp.]<br>Sequence ID: DAT08306.1Length: 190<br>hypothetical protein [Caudoviricetes sp.]<br>Sequence ID: DAW69200.1Length: 470 |
| **162** | tr\|D5NVW7\|D5NVW7_CORAM-DECOY | TNRGNATVFYGHFSTRAVQPK<br>Query 5 NATVFYGHFST 15<br>N TVFYG+FST<br>Sbjct 210 NNTVFYGQFST 220 | stabilization protein [Siphoviridae sp. ctpQM7]<br>Sequence ID: QGH72485.1Length: 564 |
| **163** | tr\|X5DPQ2\|X5DPQ2_9CORY-DECOY | VRHQCAETPTVMFLPIKVAHK<br>Query 5 CAETPTVMFLPIKVAHK 21<br>C ETP M LPI AHK<br>Sbjct 124 CNETPELMPLPIMLAHK 140 | S adenosylmethionine synthase [Caudoviricetes sp.]<br>Sequence ID: DAF29907.1Length: 388 |
| **164** | tr\|M1UK05\|M1UK05_9CORY-DECOY | NPLDVSVPSLPETMDVEGGTLVRR<br>Query 3 LDVSVPSLPETMDVEG 18<br>L VS P L ETMD EG<br>Sbjct 40 LTVSLPGL-ETMDAEG 54 | isoaspartyl dipeptidase [Caudoviricetes sp.]<br>Sequence ID: DAL16041.1Length: 384 |
| **165** | tr\|E9T0C7\|E9T0C7_COREQ-DECOY | ESAADPDLEALIPMRSGTPLFAPLK<br>Query 8 LEALIPMRSGTP 19<br>LE LI MRS TP<br>Sbjct 449 LESLIAMRSATP 460 | terminase large subunit [Aeromonas phage ZPAH14]<br>Sequence ID: YP_010656741.1Length: 674 |
| **166** | tr\|I7HCG4\|I7HCG4_CORUL-DECOY | TMHLIAASEIGDAIDSHVGRVPTEK<br>Query 2 MHLIAASEIGDAI 14<br>M+ IAA EIG+AI<br>Sbjct 245 MQ-IAAAEIGNAI 256 | tail tape measure [Caudoviricetes sp.]<br>Sequence ID: DAL80157.1Length: 1132 |
| **167** | tr\|S4XH78\|S4XH78_9CORY-DECOY | RTYIMELVYGFTQIVGEQAVMSR<br>Query 6 ELVYGFTQIVGEQA 19<br>EL YG+T+IVGE A<br>Sbjct 40 EL-YGYTEIVGERA 52 | prohead serine protease [Caudoviricetes sp.]<br>Sequence ID: DAS67792.1Length: 194 |
| **168** | tr\|G0HFU2\|G0HFU2_CORVD-DECOY | EFHHGSPFANDSGTTKKRTGDTSQR<br>Query 8 FAND--SGT-TKKRTG 20<br>FA++ SGT TKKRTG<br>Sbjct 99 FADEGISGTGTKKRTG 114 | integrase [Caudoviricetes sp.]<br>Sequence ID: DAG75148.1Length: 491 |
| **169** | tr\|D7WAN2\|D7WAN2_9CORY-DECOY | SRMGGDVRVLFDRLMMSIVFMAFR<br>Query 2 RMGGDVRVL-----------F--DRLM 15<br>RMGGDVRVL F D LM<br>Sbjct 191 RMGGDVRVLDHDDTNLGQINFTCDSLM 217 | hypothetical protein [Siphoviridae sp. cts9W16]<br>Sequence ID: DAD65470.1Length: 426 |
| **170** | tr\|U7LEL2\|U7LEL2_9CORY-DECOY | YKAVGKLLYESYHPPDLSIFATITR<br>Query 8 LYESYHPPDLS 18<br>LYESY P DLS<br>Sbjct 377 LYESYRPADLS 387 | tail sheath protein [Myoviridae sp. ctLIM9]<br>Sequence ID: DAF58467.1Length: 614 |
| **171** | tr\|H2G937\|H2G937_CORD2-DECOY | QRQPEKADVSHEETGFTMNGSVEAGW<br>Query 7 ADVSHEETGFTMNGS 21<br>ADVS GFTMNGS<br>Sbjct 226 ADVS----GFTMNGS 236 | Baseplate J like protein [Caudoviricetes sp.]<br>Sequence ID: DAQ76877.1Length: 260 |
| **172** | tr\|Q4JU87\|Q4JU87_CORJK-DECOY | SQDDSCTRFAHMGAWLSGAEGVWVKMK<br>Query 8 RFAHMGAWLSGA 19<br>RFAHMG LSGA<br>Sbjct 30 RFAHMGG-LSGA 40 | hypothetical protein UFOVP95_56 [uncultured Caudovirales phage]<br>Sequence ID: CAB4127650.1Length: 61 |
| **173** | tr\|U7L1I1\|U7L1I1_9CORY-DECOY | QNAVEGLEGFFSLALLVDEGDDSFVVSGPGR<br>Query 2 NAVEGLEGFFSLALLVDEGDD 22<br>N E L GF+ LA V+EGDD<br>Sbjct 138 NQTETLSGFYTLAFVVNEGDD 158 | hypothetical protein [Caudoviricetes sp.]<br>Sequence ID: DAO81364.1Length: 554 |
| **174** | tr\|C6R760\|C6R760_9CORY-DECOY | NLKSFAYGAMADLDVAMWLVAMSGVSTAPGNSEK<br>Query 1 NLKSFAYGAMADLDVAM 17<br>N K FAY MAD+ AM<br>Sbjct 1123 NFKDFAYSVMADM-AAM 1138 | tail length tape measure protein [Caudoviricetes sp.]<br>Sequence ID: AXF53113.1Length: 1320 |

| | | | |
|---|---|---|---|
| **175** | tr\|L1MK42\|L1MK42_9CORY-DECOY | SLWFCMGHAASLAALGRICILGSDEASHGMNDAQAK<br>Query 10 ASLAALGRICILGSDEAS 27<br>A LAA+ R+C+LGS AS<br>Sbjct 254 AGLAAMSRVCVLGSGAAS 271 | tail protein [Caudoviricetes sp.]<br>Sequence ID: DAK85635.1Length: 301 |
| **176** | tr\|D7WD57\|D7WD57_9CORY-DECOY | IVDMDGPDTGVVVSSEDDFLDDSLEKAGLFKHDAVAK<br>Query 8 DTGVVVSSEDDFL 20<br>DTGVVV SEDD L<br>Sbjct 14 DTGVVVPSEDDIL 26 | baseplate wedge protein [Caudoviricetes sp.]<br>Sequence ID: DAR87682.1Length: 397 |
| **177** | tr\|E2S3E4\|E2S3E4_9CORY-DECOY | RLLDPLLKMTTDHTEASPKYAALAIELNDVMDLETP<br>Query 9 MTT--DHTEASPK--YAALAIE 26<br>MTT DHTEA K YA LA E<br>Sbjct 3 MTTKCDHTEAIAKEFYAVLAAE 24 | hypothetical protein [Caudoviricetes sp.]<br>Sequence ID: DAU76254.1Length: 148 |
| **178** | tr\|E2S1D6\|E2S1D6_9CORY-DECOY | IRLLASGLGGCAIVAIDEKQPSGYTDDYKTGMLILFALK<br>Query 1 AIDEKQPSGYTDDYKTG 17<br>AI+EK GYTD+Y G<br>Sbjct 11 AINEK---GYTDEYQNG 24<br><br>Query 14 VAID-------EKQPSGY-TDDY 28<br>+AID EKQP Y TD+Y<br>Sbjct 84 IAIDNRVELVEEKQPGDYNTDEY 106 | hypothetical protein [Caudoviricetes sp.]<br>Sequence ID: DAK82712.1Length: 78<br><br>dUTPase [Caudoviricetes sp.]<br>Sequence ID: DAF37313.1Length: 171 |
| **179** | tr\|B1VJ26\|B1VJ26_CORU7-DECOY | SFGVSTAVNDGSGGWDGSMASEWTGAINAPNFRRKGGGFHGMPGQK<br>Query 3 GVSTAVNDGSGGWDGSMASE--WTGAIN 28<br>GVST D W S S+ WTGAIN<br>Sbjct 312 GVST---DA---WAASVGSDGAWTGAIN 333<br><br>Query 1 SEWTGAINAPNFR--RKGGGFHGM 22<br>SE G I APNF RKG +GM<br>Sbjct 580 SERNGEIWAPNFWVNRKG---QGM 600 | hypothetical protein [Caudoviricetes sp.]<br>Sequence ID: DAU36014.1Length: 585<br>tail protein [Caudoviricetes sp.]<br>Sequence ID: DAN04764.1Length: 748 |
| **180** | tr\|T9WUZ9\|T9WUZ9_CORDP-DECOY | FDGDDTMRSPSQTFGYEVELDWPCGGVTDILEHRGYRIPTR<br>Query 2 DGDDTMRSPSQTFGY 16<br>DGDDT R S+ FGY<br>Sbjct 83 DGDDTAR--SENFGY 95<br><br>Query 7 DWP-CGGVTDILEHRGYRIP 25<br>DWP CG + DILE G IP<br>Sbjct 86 DWPACGEI-DILEAKG-WIP 103 | DNA helix destabilizing protein [Caudoviricetes sp.]<br>Sequence ID: DAH34975.1Length: 188<br>laminarinase-like protein [Caudoviricetes sp.]<br>Sequence ID: DAK96867.1Length: 315 |
| **181** | tr\|L1MAP2\|L1MAP2_9CORY-DECOY | SDWYDMVKVVGMGIAVNLRGSQDPMSTEHTQLEFSLMSNIMSREPVADQPGGR<br>Query 4 YDMVK--VVGMGIAVN 17<br>YDM K VVGM IA N<br>Sbjct 109 YDM-KPTVVGMAIALN 123<br><br>Query 8 FSL--MSNIMSREPVADQ 23<br>F L NIMSR+PV +Q<br>Sbjct 243 FNLSDINNIMSRQPVCNQ 26 | Terminase small subunit [Caudoviricetes sp.]<br>Sequence ID: DAJ44382.1Length: 238<br><br>hypothetical protein [Caudoviricetes sp.]<br>Sequence ID: DAP16785.1Length: 379 |
| **182** | tr\|C6R6B7\|C6R6B7_9CORY-DECOY | VVLLVHESDMKQCHEFVSYVGNEAHTSVMNGTILIEPNGTRLATELAAWWYR<br>Query 1 YVGNEAHTSVMNGTILIEPN 20<br>Y GN A TSV GTILI+ N<br>Sbjct 27 YRGNGAPTSVPHGTILIDKN 46 | hypothetical protein [Caudoviricetes sp.]<br>Sequence ID: DAS20867.1Length: 89 |
| **183** | tr\|U7KSX2\|U7KSX2_9CORY-DECOY | LFEREYQVPVIGFNLGDHGMDLFHDSAVGDGNIPFESCLADTCPELLGIPLVTR<br>Query 1 NIPFESCLAD--------TCPELLGIPL 20<br>NI F+ L D TCP LGIPL<br>Sbjct 23 NIHFDLSLTDLNNLTYPITCP-ILGIPL 49 | hypothetical protein [Caudoviricetes sp.]<br>Sequence ID: QMP83677.1Length: 107 |
| **184** | tr\|X5DS26\|X5DS26_9CORY-DECOY | KAAFETGVK<br>Query 1 KAAFETGVK 9<br>KAAFE GVK<br>Sbjct 365 KAAFEMGVK 373 | prohead serine protease [Caudoviricetes sp.]<br>Sequence ID: DAY64994.1Length: 511 |
| **185** | tr\|E4W917\|E4W917_RHOE1-DECOY | LIAVER<br>Query 1 LIAVER 6<br>LIAVER<br>Sbjct 367 LIAVER 372 | tail length tape measure protein [Gordonia phage Forza]<br>Sequence ID: YP_010648979.1Length: 1998 |
| **186** | tr\|G7U0Y4\|G7U0Y4_CORPS-DECOY | VLDVLR<br>Query 1 VLDVLR 6<br>VLDVLR<br>Sbjct 476 VLDVLR 481 | tail protein [Caudoviricetes sp.]<br>Sequence ID: DAH71997.1Length: 956 |
| **187** | tr\|E9T103\|E9T103_COREQ-DECOY | KEGLIR<br>Query 1 KEGLIR 6<br>KEGLIR<br>Sbjct 699 KEGLIR 704 | virulence associated protein E [Caudoviricetes sp.]<br>Sequence ID: DAM58862.1Length: 804 |
| **188** | tr\|B0REE9\|B0REE9_CLAMS-DECOY | GSGQLVR<br>Query 2 SGQLVR 7<br>SGQLVR<br>Sbjct 287 SGQLVR 292 | Primase C terminal 1 (PriCT-1) [Caudoviricetes sp.]<br>Sequence ID: DAM55608.1Length: 721 |
| **189** | tr\|B0REM2\|B0REM2_CLAMS-DECOY | KMAMVK<br>Query 1 KMAMVK 6<br>KMAMVK<br>Sbjct 218 KMAMVK 223 | DNA polymerase [Siphoviridae sp. ctrpg19]<br>Sequence ID: DAD82660.1Length: 341 |
| **190** | tr\|C8RUG7\|C8RUG7_CORJE-DECOY | LVGEPGR<br>Query 1 LVGEPGR 7<br>L+GEPGR<br>Sbjct 222 LIGEPGR 228 | endosialidase chaperone [Caudoviricetes sp.]<br>Sequence ID: DAL40502.1Length: 752 |
| **191** | tr\|H2GCT9\|H2GCT9_CORD2-DECOY | GALSILR<br>Query 1 GALSILR 7<br>GALSILR<br>Sbjct 111 GALSILR 117 | protein NinB [Caudoviricetes sp.]<br>Sequence ID: DAP04579.1Length: 181 |
| **192** | tr\|M4KAZ2\|M4KAZ2_9CORY-DECOY | VSAGELR<br>Query 1 VSAGELR 7<br>VSAGELR<br>Sbjct 160 VSAGELR 166 | hypothetical protein [Caudoviricetes sp.]<br>Sequence ID: DAM29284.1Length: 304 |

| | | | |
|---|---|---|---|
| **193** | tr\|D5UUH1\|D5UUH1_TSUPD-DECOY | VLLADAR<br>Query 1 VLLADAR 7<br>VLLA+AR<br>Sbjct 135 VLLAEAR 141 | VirB11-like ATPase [Caudoviricetes sp.]<br>Sequence ID: DAO53760.1Length: 258 |
| **194** | tr\|Q6NK66\|Q6NK66_CORDI-DECOY | ANTLALR<br>Query 1 ANTLALR 7<br>ANTLALR<br>Sbjct 1095 ANTLALR 1101 | tail fiber protein [Aeromonas phage Aes012]<br>Sequence ID: YP_007677931.1Length: 1333 |
| **195** | tr\|V6V3X1\|V6V3X1_CORUL-DECOY | AAETGVPK<br>Query 1 AAETGVP 7<br>AAETGVP<br>Sbjct 223 AAETGVP 229 | minor tail protein [Rhodococcus phage Finch]<br>Sequence ID: YP_010059248.1Length: 245 |
| **196** | tr\|H2GXQ6\|H2GXQ6_CORD7-DECOY | GLEGIER<br>Query 1 GLEGIER 7<br>GLEGIER<br>Sbjct 239 GLEGIER 245 | hypothetical protein UFOVP817_17 [uncultured Caudovirales phage]<br>Sequence ID: CAB4164959.1Length: 687 |
| **197** | tr\|D5NXD9\|D5NXD9_CORAM-DECOY | AGIDAAKK<br>Query 1 AGIDAAKK 8<br>AGID AKK<br>Sbjct 942 AGIDSAKK 949 | tail tape measure [Caudoviricetes sp.]<br>Sequence ID: DAY52431.1Length: 1693 |
| **198** | tr\|H2I3G3\|H2I3G3_CORDW-DECOY | LLLDGKK<br>Query 1 LLLDGKK 7<br>LLLDGKK<br>Sbjct 81 LLLDGKK 87 | ATP-binding sugar transporter [Caudoviricetes sp.]<br>Sequence ID: DAY78265.1Length: 110 |
| **199** | tr\|M4KAE9\|M4KAE9_9CORY-DECOY | VLEEGLK<br>Query 1 VLEEGLK 7<br>VLEEGLK<br>Sbjct 207 VLEEGLK 213 | Chromatin remodeling complex ATPase [Caudoviricetes sp.]<br>Sequence ID: DAY70140.1Length: 1032 |
| **200** | tr\|I0DJG1\|I0DJG1_CORPS-DECOY | SAATEGPR<br>Query 2 AATEGPR 8<br>AATEGPR<br>Sbjct 9 AATEGPR 15 | Troponin I, cardiac muscle (L-PROLINE) II HELIX, CONTRACTILE [Caudoviricetes sp.]<br>Sequence ID: DAL05817.1Length: 43 |
| **201** | tr\|B1VDG0\|B1VDG0_CORU7-DECOY | AIIENTK<br>Query 1 AIIENTK 7<br>AIIENTK<br>Sbjct 174 AIIENTK 180 | hypothetical protein [Caudoviricetes sp.]<br>Sequence ID: DAL50209.1Length: 353 |
| **202** | tr\|M1MZD7\|M1MZD7_9CORY-DECOY | ASTGNISR<br>Query 1 ASTGNISR 8<br>ASTGNI R<br>Sbjct 57 ASTGNITR 64 | hypothetical protein [Moraxella phage Mcat8]<br>Sequence ID: AKI27361.1Length: 216 |
| **203** | tr\|Q4JUR8\|Q4JUR8_CORJK-DECOY | DGKKELK<br>Query 1 DGKKELK 7<br>DGKKELK<br>Sbjct 432 DGKKELK 438 | DnaB-type replicative DNA helicase [Bacillus phage vB_BanS_Skywalker]<br>Sequence ID: YP_010680833.1Length: 510 |
| **204** | tr\|C3PIL7\|C3PIL7_CORA7-DECOY | VGIDNFR<br>Query 1 VGIDNFR 7<br>VGI+NFR<br>Sbjct 100 VGIENFR 106 | endolysin R21 like protein [Caudoviricetes sp.]<br>Sequence ID: DAQ81604.1Length: 159 |
| **205** | tr\|U7KS12\|U7KS12_9CORY-DECOY | MSAAGMNK<br>Query 1 MSAAGMNK 8<br>M AAGMNK<br>Sbjct 16 MKAAGMNK 23 | Repressor protein CI [Caudoviricetes sp.]<br>Sequence ID: DAM07855.1Length: 220 |
| **206** | tr\|C4LKL5\|C4LKL5_CORK4-DECOY | LSSIEKR<br>Query 1 LSSIEKR 7<br>LSSIEKR<br>Sbjct 83 LSSIEKR 89 | helix-turn-helix domain protein [Caudoviricetes sp.]<br>Sequence ID: DAV00016.1Length: 197 |
| **207** | tr\|C2CRG8\|C2CRG8_CORST-DECOY | DTIMSLR<br>Query 1 DTIMSLR 7<br>DT+MSLR<br>Sbjct 49 DTVMSLR 55 | PORTAL PROTEIN [Caudoviricetes sp.]<br>Sequence ID: DAI62757.1Length: 446 |
| **208** | tr\|B1VGA3\|B1VGA3_CORU7-DECOY | KTFGLNR<br>Query 1 KTFGLNR 7<br>KTFG+NR<br>Sbjct 1285 KTFGMNR 1291 | tail protein [Caudoviricetes sp.]<br>Sequence ID: DAR24771.1Length: 1830 |
| **209** | tr\|H2G8P5\|H2G8P5_CORD3-DECOY | GQFSTGIK<br>Query 1 GQ-FSTGIK 8<br>GQ FSTGIK<br>Sbjct 681 GQTFSTGIK 689 | minor tail protein [Caudoviricetes sp.]<br>Sequence ID: DAG85836.1Length: 1139 |
| **210** | tr\|I0LK91\|I0LK91_CORGK-DECOY | YGIGGKSR<br>Query 1 YGIGGKS 7<br>YGIGGKS<br>Sbjct 230 YGIGGKS 236 | stabilization protein [Caudoviricetes sp.]<br>Sequence ID: DAV04862.1Length: 584 |
| **211** | tr\|W5Y4X0\|W5Y4X0_9CORY-DECOY | LAESASHK<br>Query 1 LAESASHK 8<br>L ESASHK<br>Sbjct 71 LTESASHK 78 | Host cell surface-exposed lipoprotein [Caudoviricetes sp.]<br>Sequence ID: DAU49831.1Length: 141 |
| **212** | tr\|U7K1G7\|U7K1G7_9CORY-DECOY | KLEVAQR<br>Query 2 LEVAQR 7<br>LEVAQR<br>Sbjct 323 LEVAQR 328 | ParB protein [Caudoviricetes sp.]<br>Sequence ID: DAK84034.1Length: 367 |
| **213** | tr\|H2GGS4\|H2GGS4_CORDN-DECOY | LPEDKSR<br>Query 1 LPEDKSR 7<br>LPEDK R<br>Sbjct 102 LPEDKAR 108 | transposase [Caudoviricetes sp.]<br>Sequence ID: DAW65270.1Length: 663 |
| **214** | tr\|D7WF36\|D7WF36_9CORY-DECOY | TKLPMEK<br>Query 2 KLPMEK 7<br>KLPMEK<br>Sbjct 9 KLPMEK 14 | RNA dependent RNA polymerase [Caudoviricetes sp.]<br>Sequence ID: DAL75910.1Length: 926 |

| | | | |
|---|---|---|---|
| **215** | tr\|A4QF23\|A4QF23_CORGB-DECOY | SAADLTNR<br>Query 1 SAADLTN 7<br>SAADLTN<br>Sbjct 112 SAADLTN 118 | stabilization protein, partial [Caudoviricetes sp.]<br>Sequence ID: DAQ54374.1Length: 732 |
| **216** | tr\|L1M888\|L1M888_9CORY-DECOY | SAGTAAVDR<br><br>Query 3 GTAAVDR 9<br>GTAAVDR<br>Sbjct 31 GTAAVDR 37 | tail sheath tube [Caudoviricetes sp.]<br>Sequence ID: DAY41123.1Length: 490 |
| **217** | tr\|E0MWG0\|E0MWG0_9CORY-DECOY | IDLCTLK<br>Query 1 IDLCTLK 7<br>I+LCTLK<br>Sbjct 63 IELCTLK 69 | hypothetical protein [Caudoviricetes sp.]<br>Sequence ID: DAR95454.1Length: 279 |
| **218** | tr\|V6V527\|V6V527_CORUL-DECOY | LMIQNTK<br>Query 1 LMIQNT 6<br>LMIQNT<br>Sbjct 55 LMIQNT 60 | hypothetical protein [Caudoviricetes sp.]<br>Sequence ID: DAL71530.1Length: 63 |
| **219** | tr\|H2H3B2\|H2H3B2_CORDD-DECOY | QVSTERR<br>Query 1 QVSTERR 7<br>Q+STERR<br>Sbjct 59 QISTERR 65 | chitosanase, partial [Caudoviricetes sp.]<br>Sequence ID: DAM10437.1Length: 251 |
| **220** | tr\|H6M4M3\|H6M4M3_CORPS-DECOY | YQKEALK<br>Query 1 YQKEALK 7<br>YQKEALK<br>Sbjct 31 YQKEALK 37 | terminase large subunit [Caudoviricetes sp.]<br>Sequence ID: DAS19021.1Length: 630 |
| **221** | tr\|E9T2L0\|E9T2L0_COREQ-DECOY | LWELACR<br>Query 1 LWELAC 6<br>LWELAC<br>Sbjct 65 LWELAC 70 | FabL, FabH, KcsA bundle, C-terminus, IMMUNE SYSTEM.6A [Caudoviricetes sp.]<br>Sequence ID: DAW59958.1Length: 77 |
| **222** | tr\|C8RRQ1\|C8RRQ1_CORJE-DECOY | IEGDLVEK<br>Query 1 IEGD-LVEK 8<br>IEGD LVEK | Fe2+ transport system protein B, partial [Caudoviricetes sp.]<br>Sequence ID: DAH64754.1Length: 669 |
| **223** | tr\|U7LSF7\|U7LSF7_9CORY-DECOY | SIDTNVPR<br>Query 1 SIDTNVP 7<br>SIDTNVP<br>Sbjct 158 SIDTNVP 16 | hypothetical protein [Caudoviricetes sp.]<br>Sequence ID: DAN43930.1Length: 1230 |
| **224** | tr\|C8RRQ1\|C8RRQ1_CORJE-DECOY | IEGDLVEK<br>Query 1 IEGD-LVEK 8<br>IEGD LVEK<br>Sbjct 193 IEGDALVEK 201 | Fe2+ transport system protein B, partial [Caudoviricetes sp.]<br>Sequence ID: DAH64754.1Length: 669 |
| **225** | tr\|H2HEF2\|H2HEF2_CORDJ-DECOY | IDNGLPVGK<br>Query 1 IDNGLPV 7<br>IDNGLPV<br>Sbjct 378 IDNGLPV 384 | Integrase [Caudoviricetes sp.]<br>Sequence ID: DAV71322.1Length: 430 |
| **226** | tr\|C6R6S6\|C6R6S6_9CORY-DECOY | TGAVTIGGIK<br>Query 1 TGAVTIGG---IK 10<br>TGAVT+GG IK<br>Sbjct 1062 TGAVTVGGEIFIK 1074 | tail protein [Caudoviricetes sp.]<br>Sequence ID: DAO96736.1Length: 1229 |
| **227** | tr\|H2GFH1\|H2GFH1_CORDN-DECOY | AVIVRFGR<br>Query 1 AVIVRFGR 8<br>A IVRFGR<br>Sbjct 60 AAIVRFGR 67 | minor tail protein [Caudoviricetes sp.]<br>Sequence ID: DAF81482.1Length: 993 |
| **228** | tr\|W5XXV7\|W5XXV7_9CORY-DECOY | NIALDGGIR<br>Query 1 NIALD-GGI 8<br>NIALD GGI<br>Sbjct 4807 NIALDHGGI 4815 | leukotoxin LktA family filamentous adhesin N-terminal domain [Caudoviricetes sp.]<br>Sequence ID: DAV38177.1Length: 5628 |
| **229** | tr\|E2S2K1\|E2S2K1_9CORY-DECOY | AGVFDIGVR<br>Query 2 GVFDIGVR 9<br>GVFDIG R<br>Sbjct 55 GVFDIGLR 62 | YjcQ protein [Caudoviricetes sp.]<br>Sequence ID: DAO82142.1Length: 131 |
| **230** | tr\|C2GHV0\|C2GHV0_9CORY-DECOY | RASAGHWR<br>Query 2 ASAGHW 7<br>ASAGHW<br>Sbjct 972 ASAGHW 977 | tail protein [Caudoviricetes sp.]<br>Sequence ID: DAI51192.1Length: 1679 |
| **231** | tr\|E9T5G6\|E9T5G6_COREQ-DECOY | TQAAAAHMR<br>Query 1 TQAAAAH-MR 9<br>TQA AAH MR<br>Sbjct 403 TQADAAHTMR 412 | chromosome partitioning protein [Caudoviricetes sp.]<br>Sequence ID: DAY81105.1Length: 481 |
| **232** | tr\|D5URU9\|D5URU9_TSUPD-DECOY | LRELIADK<br>Query 1 LRELIADK 8<br>LRELIA+K<br>Sbjct 41 LRELIAEK 48 | Transcriptional regulator, AbiEi antitoxin N-terminal domain [Caudoviricetes sp.]<br>Sequence ID: DAN14702.1Length: 13 |
| **233** | sp\|C4LJJ2\|GATC_CORK4-DECOY | WSAVKLTR<br>Query 1 WSAVKL 6<br>WSAVKL<br>Sbjct 533 WSAVKL 538 | minor tail protein, partial [Caudoviricetes sp.]<br>Sequence ID: DAM25305.1Length: 1148 |
| **234** | tr\|Q6M1F4\|Q6M1F4_CORGL-DECOY | DQAAQSVDK<br>Query 1 DQAAQSVDK 9<br>+QAAQ VDK<br>Sbjct 466 NQAAQTVDK 474 | Portal [Caudoviricetes sp.]<br>Sequence ID: DAW16982.1Length: 483 |
| **235** | tr\|B1VFD2\|B1VFD2_CORU7-DECOY | IEIDTINR<br>Query 1 IEIDTINR 8<br>IEIDT+NR<br>Sbjct 79 IEIDTVNR 86 | short C-terminal domain [Caudoviricetes sp.]<br>Sequence ID: DAG11813.1Length: 244 |
| **236** | tr\|B1VGQ0\|B1VGQ0_CORU7-DECOY | GPKLDHISL<br>Query 2 PKLDHISL 9<br>PKL+HIS+<br>Sbjct 344 PKLEHISM 351 | protein of unknown function DUF859 [Caudoviricetes sp.]<br>Sequence ID: DAP02558.1Length: 604 |

| | | | |
|---|---|---|---|
| **237** | tr\|D5UQ95\|D5UQ95_TSUPD-DECOY | IGLTPIDGAR<br>Query 1 IGLTPIDG-AR 10<br>I LTPIDG AR<br>Sbjct 55 INLTPIDGVAR 65 | hypothetical protein [Caudoviricetes sp.]<br>Sequence ID: DAR90668.1Length: 694 |
| **238** | tr\|M1UJB9\|M1UJB9_9CORY-DECOY | LDILLDTAR<br>Query 1 LDILLDT 7<br>LDILLDT<br>Sbjct 150 LDILLDT 156 | deoxyribosyltransferase [Caudoviricetes sp.]<br>Sequence ID: DAL07722.1Length: 326 |
| **239** | tr\|E2MVU2\|E2MVU2_CORAY-DECOY | TLAGDMAETK<br>Query 2 LAGDMAET 9<br>LAGDM ET<br>Sbjct 48 LAGDMGET 55 | head to tail adaptor [Caudoviricetes sp.]<br>Sequence ID: DAM54937.1Length: 160 |
| **240** | tr\|C0E588\|C0E588_9CORY-DECOY | IEPGVVDPGR<br>Query 1 IEP-GVVDP-GR 10<br>IEP GV+DP GR<br>Sbjct 330 IEPIGVIDPQGR 341 | Cytosine specific methyltransferase [Caudoviricetes sp.]<br>Sequence ID: DAF69746.1Length: 511 |
| **241** | tr\|M1N102\|M1N102_9CORY-DECOY | LDNSNVDGIK<br>Query 1 LDNSNVDGI 9<br>LDNSNVD I<br>Sbjct 150 LDNSNVDDI 158 | adenine-specific methyltransferase [Caudoviricetes sp.]<br>Sequence ID: DAL79427.1Length: 200 |
| **242** | tr\|C8RSX3\|C8RSX3_CORJE-DECOY | APLEYVDDR<br>Query 3 LEYVDDR 9<br>LEYVD+R<br>Sbjct 686 LEYVDER 692 | DNA-directed RNA polymerase subunit alpha [Caudoviricetes sp.]<br>Sequence ID: DAQ87292.1Length: 715 |
| **243** | tr\|E4WD30\|E4WD30_RHOE1-DECOY | KDSPVAAAGAAK<br>Query 1 KDSPVAAAGAAK 12<br>K+ VAAAG AK<br>Sbjct 476 KETAVAAAGGAK 487 | tail tape measure [Caudoviricetes sp.]<br>Sequence ID: DAP14023.1Length: 754 |
| **244** | tr\|X5DN60\|X5DN60_9CORY-DECOY | DAFIMPQDR<br>Query 1 DAFIMPQD 8<br>D FIMP+D<br>Sbjct 75 DSFIMPHD 82 | hypothetical protein [Caudoviricetes sp.]<br>Sequence ID: QHJ79787.1Length: 164 |
| **245** | tr\|I4JS15\|I4JS15_CORDP-DECOY | SGGSGQKGGTMR<br>Query 6 QKGGTMR 12<br>QKGGTMR<br>Sbjct 171 QKGGTMR 177 | baseplate assembly protein V [Caudoviricetes sp.]<br>Sequence ID: DAL37951.1Length: 213 |
| **246** | tr\|F8DZZ6\|F8DZZ6_CORRG-DECOY | VLESIVVLTR<br>Query 2 LESIVVLT 9<br>LESIVVLT<br>Sbjct 77 LESIVVLT 84 | major tail protein [Caudoviricetes sp.]<br>Sequence ID: DAV10168.1Length: 147 |
| **247** | tr\|M1NPD8\|M1NPD8_9CORY-DECOY | VSSASSQKESR<br>Query 1 VSSASSQK 8<br>VSSASSQK<br>Sbjct 21 VSSASSQK 28 | hypothetical protein [Caudoviricetes sp.]<br>Sequence ID: DAI47768.1Length: 42 |
| **248** | tr\|U7L892\|U7L892_9CORY-DECOY | HRFIFGHVR<br>Query 1 HRFIFGHVR 9<br>HR IFG+VR<br>Sbjct 307 HRLIFGQVR 315 | hypothetical protein [Caudoviricetes sp.]<br>Sequence ID: DAI76142.1Length: 330 |
| **249** | tr\|H2GR69\|H2GR69_CORDB-DECOY | VAEVGAHNNER<br>Query 1 VAEVGAH 7<br>VAEVGAH<br>Sbjct 55 VAEVGAH 61 | endodeoxyribonuclease I [Caudoviricetes sp.]<br>Sequence ID: DAH44623.1Length: 150Number of Matches: 1 |
| **250** | tr\|V6V6Z1\|V6V6Z1_CORUL-DECOY | GAAEAATLRLAR<br>Query 4 EAATLRLAR 12<br>E ATLRLAR<br>Sbjct 785 EIATLRLAR 793 | minor tail protein [Caudoviricetes sp.]<br>Sequence ID: DAH71684.1Length: 1452 |
| **251** | tr\|U7KCL9\|U7KCL9_9CORY-DECOY | MELVSPLALAR<br>Query 2 ELVSPLAL 9<br>ELVSPLAL<br>Sbjct 180 ELVSPLAL 187 | STRUCTURAL MAINTENANCE OF CHROMOSOMES PROTEIN [Caudoviricetes sp.]<br>Sequence ID: DAO82609.1Length: 423 |
| **252** | tr\|I4JU32\|I4JU32_CORDP-DECOY | GETVMFPTQPNKDTCPSSSEAVR<br>Query 2 ETVMF-PTQ----PNKDTCPSSSEAVR 23<br>ETV F P Q NK T P SE VR<br>Sbjct 1514 ETVVFNPAQIKYTSNK-T-PTASEDVR 1538 | PolyVal ADP-Ribosyltransferase [Caudoviricetes sp.]<br>Sequence ID: DAR52810.1Length: 3168 |
| **253** | tr\|H2GX36\|H2GX36_CORD7-DECOY | MMQAQQEYR<br>Query 1 MMQAQQEY 8<br>MMQA QEY<br>Sbjct 194 MMQADQEY 201 | DnaD-like replication protein [Caudoviricetes sp.]<br>Sequence ID: DAI33289.1Length: 341 |
| **254** | tr\|H2HQ47\|H2HQ47_CORDK-DECOY | VQKITYCDEK<br>Query 4 ITYCDEK 10<br>ITYCDEK<br>Sbjct 75 ITYCDEK 81 | head tail joining protein [Caudoviricetes sp.]<br>Sequence ID: DAL51562.1Length: 106 |
| **255** | tr\|M1UTI9\|M1UTI9_9CORY-DECOY | AFETPETIDAR<br>Query 2 FETPETIDAR 11<br>FE PET+D R<br>Sbjct 13 FEAPETVDSR 22 | head scaffolding protein [Bacillus phage Thornton]<br>Sequence ID: YP_010114396.1 |
| **256** | sp\|C3PL26\|RS5_CORA7-DECOY | AFATATCADLPK<br>Query 5 ATCADLP 11<br>ATCADLP<br>Sbjct 453 ATCADLP 459 | tail protein [Caudoviricetes sp.]<br>Sequence ID: DAO57310.1Length: 1194 |
| **257** | tr\|E2MY77\|E2MY77_CORAY-DECOY | APEAFPPGKLVR<br>Query 1 APEAFPP 7<br>APEAFPP<br>Sbjct 44 APEAFPP 50 | hypothetical protein [Caudoviricetes sp.]<br>Sequence ID: DAW34991.1Length: 120 |
| **258** | tr\|B0REH4\|B0REH4_CLAMS-DECOY | AVNNVGRSNLQR<br>Query 2 VNNVGRSN 9<br>VNNVG SN<br>Sbjct 1647 VNNVGKSN 1654 | crystallin beta/gamma motif-containing protein [Caudoviricetes sp.]<br>Sequence ID: DAL89814.1Length: 2768 |
| **259** | tr\|U7L314\|U7L314_9CORY-DECOY | VEVAGAAAIPLAEAR<br>Query 7 AAIPLAEAR 15<br>AAIPLAEAR<br>Sbjct 1216 AAIPLAEAR 1224 | DNA polymerase III PolC [Caudoviricetes sp.]<br>Sequence ID: DAL23106.1Length: 1269 |

| | | | |
|---|---|---|---|
| **260** | tr\|C8NQU2\|C8NQU2_COREF-DECOY | AAMAYAMVGGAGLKR<br>Query 2 AMAYAMVGGAG 12<br>AMA+AMVG AG<br>Sbjct 96 AMAFAMVGVAG 106 | major_cap_HK97, phage major capsid protein, HK97 family [uncultured Caudovirales phage]<br>Sequence ID: CAB4177373.1Length: 528 |
| **261** | tr\|C3PHH5\|C3PHH5_CORA7-DECOY | TGGDWLMSSLDWRR<br>Query 1 TGGDWLMSSLDW 12<br>T GDW MS +DW<br>Sbjct 18 TTGDWHMSGMDW 29 | hypothetical protein [Caudoviricetes sp.]<br>Sequence ID: DAT32537.1Length: 157 |
| **262** | tr\|A4QGJ2\|A4QGJ2_CORGB-DECOY | TLEAIR<br>Query 1 TLEAIR 6<br>TLEAIR<br>Sbjct 245 TLEAIR 250 | STRUCTURAL MAINTENANCE OF CHROMOSOMES PROTEIN [Caudoviricetes sp.] |
| **263** | tr\|I7H9P5\|I7H9P5_CORUL-DECOY | HEGTIR<br>Query 1 HEGTIR 6<br>HEGTIR<br>Sbjct 12 HEGTIR 17 | HepA-related protein (HARP) [Caudoviricetes sp.] |
| **264** | tr\|B1VGJ2\|B1VGJ2_CORU7-DECOY | AVEFLR<br>Query 1 AVEFLR 6<br>AVEFLR<br>Sbjct 2437 AVEFLR 2442 | crystallin beta/gamma motif-containing protein [Caudoviricetes sp.]<br>Sequence ID: DAR33089.1Length: 2802 |
| **265** | tr\|C0DZG1\|C0DZG1_9CORY-DECOY | PEILER<br>Query 1 PEILER 6<br>PEILER<br>Sbjct 375 PEILER 380 | tail protein [Caudoviricetes sp.]<br>Sequence ID: DAJ54077.1Length: 860 |
| **266** | tr\|E9SZK1\|E9SZK1_COREQ-DECOY | LEIAGTR<br>Query 1 LEIAGTR 7<br>LEI GTR<br>Sbjct 804 LEISGTR 810 | Putative tail protein [Caudoviricetes sp.]<br>Sequence ID: DAZ83122.1Length: 979 |
| **267** | tr\|E2MYN9\|E2MYN9_CORAY-DECOY | DWELAK<br>Query 1 DWELAK 6<br>DWELAK<br>Sbjct 151 DWELAK 156 | short patch repair endonuclease [Caudoviricetes sp.]<br>Sequence ID: DAO61328.1Length: 291N |
| **268** | tr\|S4X9Q6\|S4X9Q6_9CORY-DECOY | AMGIRAK<br>Query 1 AMGIRAK 7<br>AMGIRAK<br>Sbjct 385 AMGIRAK 391 | minor capsid protein [Caudoviricetes sp.]<br>Sequence ID: DAS52309.1Length: 643 |
| **269** | tr\|C0VT52\|C0VT52_9CORY-DECOY | LIVGLQK<br>Query 1 LIVGLQK 7<br>LI+GLQK<br>Sbjct 214 LIIGLQK 220 | lysozyme family protein [Caudoviricetes sp.]<br>Sequence ID: DAL39955.1Length: 309 |
| **270** | tr\|D9QDV2\|D9QDV2_CORP2-DECOY | WAEAAAR<br>Query 1 WAEAAAR 7<br>WAEAAAR<br>Sbjct 664 WAEAAAR 670 | virulence associated protein E [Caudoviricetes sp.]<br>Sequence ID: DAQ33886.1Length: 807 |
| **271** | tr\|M1NNX3\|M1NNX3_9CORY-DECOY | ADILAGAR<br>Query 1 ADILAGA 7<br>ADILAGA<br>Sbjct 261 ADILAGA 267 | minor tail protein [Caudoviricetes sp.]<br>Sequence ID: DAG84065.1Length: 1301 |
| **272** | tr\|Q8FR48\|Q8FR48_COREF-DECOY | ANMDVAR<br>Query 2 NMDVAR 7<br>NMDVAR<br>Sbjct 1991 NMDVAR 1996 | protein of unknown function (DUF5401) [Caudoviricetes sp.]<br>Sequence ID: DAI66118.1Length: 3364 |
| **273** | tr\|H2GPS6\|H2GPS6_CORDB-DECOY | GPPEIQR<br>Query 2 PPEIQR 7<br>PPEIQR<br>Sbjct 118 PPEIQR 123 | hypothetical protein [Caudoviricetes sp.]<br>Sequence ID: DAT61111.1Length: 268 |
| **274** | tr\|S4XH24\|S4XH24_9CORY-DECOY | AAMIVRK<br>Query 1 AAMIVRK 7<br>AAMIVRK<br>Sbjct 19 AAMIVRK 25 | hypothetical protein [Caudoviricetes sp.]<br>Sequence ID: DAI53195.1Length: 102 |
| **275** | tr\|F8E2W4\|F8E2W4_CORRG-DECOY | AIADDFR<br>Query 2 IADDFR 7<br>IADDFR<br>Sbjct 584 IADDFR 589<br><br>Query 2 IADDFR 7<br>IADDFR<br>Sbjct 111 IADDFR 116 | capsid maturation protease and MuF-like fusion protein [Mycobacterium phage Zonia]<br>Sequence ID: AIM50438.1Length: 863<br><br>43 kDa tail protein [Caudoviricetes sp.]<br>Sequence ID: DAS43637.1Length: 326 |
| **276** | tr\|C3PIL7\|C3PIL7_CORA7-DECOY | VGIDNFR<br>Query 1 VGIDNFR 7<br>VGI+NFR<br>Sbjct 100 VGIENFR 106 | endolysin R21 like protein [Caudoviricetes sp.]<br>Sequence ID: DAQ81604.1Length: 159 |
| **277** | tr\|H2GFN4\|H2GFN4_CORDN-DECOY | ALSYITR<br>Query 1 ALSYITR 7<br>ALSYITR<br>Sbjct 15 ALSYITR 21 | hypothetical protein, partial [Caudoviricetes sp.]<br>Sequence ID: DAQ08131.1Length: 42 |
| **278** | tr\|C8RVT8\|C8RVT8_CORJE-DECOY | ADDKRVK<br>Query 1 ADDKRVK 7<br>AD+KRVK<br>Sbjct 560 ADEKRVK 566 | hypothetical protein [Caudoviricetes sp.]<br>Sequence ID: AXQ66572.1Length: 950 |
| **279** | tr\|G0CVL9\|G0CVL9_CORUB-DECOY | GVPGGFGSR<br>Query 2 VPGGFGSR 9<br>VPGGFGSR<br>Sbjct 303 VPGGFGSR 310 | CTP synthetase [Caudoviricetes sp.]<br>Sequence ID: DAU74571.1Length: 488 |
| **280** | tr\|C0E4C1\|C0E4C1_9CORY-DECOY | HEVIALR<br>Query 1 HEVIALR 7<br>HEVIALR<br>Sbjct 329 HEVIALR 335 | bifunctional NMN adenylyltransferase/Nudix hydrolase [Klebsiella phage 150040]<br>Sequence ID: UXD79444.1Length: 352 |
| **281** | tr\|E9SXW6\|E9SXW6_COREQ-DECOY | HIAAVLAR<br>Query 1 HIAAVL-AR 8<br>HIAAVL AR<br>Sbjct 32 HIAAVLQAR 40 | Replication associated protein [Caudoviricetes sp.]<br>Sequence ID: DAI87294.1Length: 679 |

| | | | |
|---|---|---|---|
| **282** | tr\|C2CRE6\|C2CRE6_CORST-DECOY | SLQSLDLK<br>Query 1 SLQSLDL 7<br>SLQSLDL<br>Sbjct 40 SLQSLDL 46 | portal [Caudoviricetes sp.]<br>Sequence ID: DAV25609.1Length: 546 |
| **283** | sp\|Q6NJ04\|DNAE2_CORDI-DECOY | QMLEIGAK<br>Query 1 QMLEIGAK 8<br>QM EIGAK<br>Sbjct 58 QMQEIGAK 65 | Protein of unknown function (DUF3486) [Caudoviricetes sp.]<br>Sequence ID: DAK48460.1Length: 179 |
| **284** | tr\|H2I1T6\|H2I1T6_CORDW-DECOY | MNKQELK<br>Query 1 MNKQELK 7<br>MNKQELK<br>Sbjct 1 MNKQELK 7 | Protein of unknown function (DUF1642) [Caudoviricetes sp.]<br>Sequence ID: DAX64601.1Length: 243 |
| **285** | tr\|C0XUY5\|C0XUY5_9CORY-DECOY | YIELLTR<br>Query 1 YIELLTR 7<br>Y+ELLTR<br>Sbjct 767 YVELLTR 773 | minor capsid protein [Caudoviricetes sp.]<br>Sequence ID: DAW02997.1Length: 779 |
| **286** | tr\|G2EK45\|G2EK45_CORGT-DECOY | AMNDITTR<br>Query 1 AMNDITT 7<br>AMNDITT<br>Sbjct 113 AMNDITT 119 | hypothetical protein [Caudoviricetes sp.]<br>Sequence ID: DAQ40697.1Length: 143 |
| **287** | tr\|D5UTB6\|D5UTB6_TSUPD-DECOY | FESDKLASKEMSR<br>Query 1 FE---SDKLASKEMSR 13<br>FE SDK+A K MSR<br>Sbjct 476 FEDLVSDKMARKNMSR 491 | minor capsid protein [Caudoviricetes sp.]<br>Sequence ID: DAL79681.1Length: 522 |
| **288** | tr\|V6V4E2\|V6V4E2_CORUL-DECOY | ILIPRWR<br>Query 1 ILIPRW 6<br>ILIPRW<br>Sbjct 471 ILIPRW 476 | integrase [Caudoviricetes sp.]<br>Sequence ID: DAG26728.1Length: 540 |
| **289** | tr\|U7K5H8\|U7K5H8_9CORY-DECOY | GGVYAESFE<br>Query 2 GVYAESFE 9<br>GVY ESFE<br>Sbjct 55 GVYSESFE 62 | stabilization protein [Caudoviricetes sp.]<br>Sequence ID: DAK85409.1Length: 534 |
| **290** | tr\|Q6NI43\|Q6NI43_CORDI-DECOY | LQTGPPAER<br>Query 2 QTGPPAE 8<br>QTGPPAE<br>Sbjct 480 QTGPPAE 486 | Large Terminase [Caudoviricetes sp.]<br>Sequence ID: DAW00001.1Length: 499 |
| **291** | tr\|S5TGD4\|S5TGD4_9CORY-DECOY | SPIEAGLGAR<br>Query 1 SPIEAGLG 8<br>SPIEAGLG<br>Sbjct 248 SPIEAGLG 255 | glycine cleavage system aminomethyltransferase [Caudoviricetes sp.]<br>Sequence ID: DAT40344.1Length: 361 |
| **292** | tr\|S2Z3K6\|S2Z3K6_9CORY-DECOY | RSDVAEAPK<br>Query 1 RSDVAEA 7<br>RSDVAEA<br>Sbjct 57 RSDVAEA 63 | replisome organizer [Caudoviricetes sp.]<br>Sequence ID: DAW60882.1Length: 275 |
| **293** | tr\|B1VFD2\|B1VFD2_CORU7-DECOY | IEIDTINR<br>Query 1 IEIDTINR 8<br>IEIDT+NR<br>Sbjct 79 IEIDTVNR 86 | Short C-terminal domain [Caudoviricetes sp.]<br>Sequence ID: DAG11813.1Length: 244 |
| **294** | tr\|X5DUC4\|X5DUC4_9CORY-DECOY | AAGPFLEIR<br>Query 4 PFLEIR 9<br>PFLEIR<br>Sbjct 65 PFLEIR 70 | hypothetical protein [Caudoviricetes sp.]<br>Sequence ID: DAP97865.1Length: 94 |
| **295** | tr\|E4WHL8\|E4WHL8_RHOE1-DECOY | NEMGIVGMK<br>Query 1 NEMGIVGM 8<br>NEMG VGM<br>Sbjct 330 NEMGLVGM 337 | Replication associated protein [Caudoviricetes sp.]<br>Sequence ID: DAI71958.1Length: 879 |
| **296** | tr\|G0CP56\|G0CP56_CORUL-DECOY | AGYTQQALK<br>Query 3 YTQQALK 9<br>YTQQALK<br>Sbjct 99 YTQQALK 105 | hypothetical protein UFOVP318_4 [uncultured Caudovirales phage] |
| **297** | tr\|U7KLF8\|U7KLF8_9CORY-DECOY | MDLTKLIR<br>Query 1 MDLT-KLIR 8<br>MDLT KLIR<br>Sbjct 554 MDLTGKLIR 562 | minor tail protein [Caudoviricetes sp.]<br>Sequence ID: DAQ09724.1Length: 917 |
| **298** | tr\|D5UP96\|D5UP96_TSUPD-DECOY | TIIFEITR<br>Query 2 IIFEITR 8<br>IIF+ITR<br>Sbjct 48 IIFDITR 54 | hypothetical protein [Caudoviricetes sp.]<br>Sequence ID: DAJ44988.1Length: 537 |
| **299** | tr\|C8NS66\|C8NS66_COREF-DECOY | LVLLSSPER<br>Query 1 LVLLSSPE 8<br>LVLLSSPE<br>Sbjct 125 LVLLSSPE 132 | hypothetical protein [Caudoviricetes sp.]<br>Sequence ID: DAS91905.1Length: 161 |
| **300** | tr\|M1UJB9\|M1UJB9_9CORY-DECOY | LDILLDTAR<br>Query 1 LDILLDT 7<br>LDILLDT<br>Sbjct 150 LDILLDT 156 | deoxyribosyltransferase [Caudoviricetes sp.]<br>Sequence ID: DAL07722.1Length: 326 |
| **301** | tr\|C0XSN0\|C0XSN0_9CORY-DECOY | EGSARSKTAR<br>Query 1 EGSARSKTAR 10<br>EGSA KTAR<br>Sbjct 114 EGSAIRKTAR 123 | minor capsid protein [Caudoviricetes sp.]<br>Sequence ID: DAG32169.1Length: 566 |
| **302** | tr\|G7HUI9\|G7HUI9_9CORY-DECOY | LAAVDDVLQK<br>Query 2 AAVDDVLQ 9<br>AAVDDVLQ<br>Sbjct 341 AAVDDVLQ 348 | Baseplate J like protein [Caudoviricetes sp.]<br>Sequence ID: DAR86808.1Length: 388 |
| **303** | tr\|C2GK55\|C2GK55_9CORY-DECOY | TGFADVLEAR<br>Query 3 FADVLEAR 10<br>FADVLE R<br>Sbjct 631 FADVLESR 638 | hypothetical protein [Caudoviricetes sp.]<br>Sequence ID: DAP35613.1Length: 3142 |

| | | | |
|---|---|---|---|
| **304** | tr\|E2MTN2\|E2MTN2_CORAY-DECOY | LAATSSLFLR<br>Query 3 ATSSLFLR 10<br>ATS LFLR<br>Sbjct 74 ATSPLFLR 81 | helix-turn-helix domain protein [Caudoviricetes sp.]<br>Sequence ID: DAX49062.1Length: 175 |
| **305** | tr\|G2EIC2\|G2EIC2_CORGT-DECOY | ETESACLER<br>Query 1 ETESACLE 8<br>E ESACLE<br>Sbjct 331 EAESACLE 338 | hypothetical protein [Caudoviricetes sp.]<br>Sequence ID: DAJ53510.1Length: 700 |
| **306** | tr\|H2H4N5\|H2H4N5_CORDD-DECOY | DVKEMGEGSR<br>Query 1 DVKEMGEGSR 10<br>DVKEMG G R<br>Sbjct 186 DVKEMGKGPR 195 | transcription elongation factor NusA [Caudoviricetes sp.]<br>Sequence ID: DAE84267.1Length: 361 |
| **307** | tr\|C2CMH5\|C2CMH5_CORST-DECOY | TIQELQPLGK<br>Query 2 IQELQPL 8<br>IQELQPL<br>Sbjct 73 IQELQPL 79 | hypothetical protein [Caudoviricetes sp.]<br>Sequence ID: QHJ76210.1Length: 181 |
| **308** | tr\|D5NVS7\|D5NVS7_CORAM-DECOY | LVPDSWEKR<br>Query 1 LVPDSWEK 8<br>L PDSWEK<br>Sbjct 272 LLPDSWEK 279 | methionyl-tRNA synthetase [Caudoviricetes sp.]<br>Sequence ID: DAH13796.1Length: 680 |
| **309** | tr\|M1UES1\|M1UES1_9CORY-DECOY | GGGDGQMLWAK<br>Query 4 DGQMLWAK 11<br>DGQ+LWAK<br>Sbjct 110 DGQLLWAK 117 | PD-(D/E)XK nuclease superfamily protein [Caudoviricetes sp.]<br>Sequence ID: DAV12575.1Length: 480 |
| **310** | tr\|G4QPV7\|G4QPV7_CORPS-DECOY | AISAFDGESLK<br>Query 1 AISAFDGESLK 11<br>AIS FDG SLK<br>Sbjct 757 AISGFDG-SLK 766 | DNA primase [Caudoviricetes sp.]<br>Sequence ID: DAM78246.1Length: 986 |
| **311** | tr\|S2YAM4\|S2YAM4_9CORY-DECOY | GSAIGLGAPELR<br>Query 4 IGLGAPELR 12<br>IGL APELR<br>Sbjct 32 IGLDAPELR 40 | FocB protein-alpha, helix-turn-helix, TRANSCRIPTION.4A [Caudoviricetes sp.]<br>Sequence ID: DAY64442.1Length: 96 |
| **312** | tr\|I0LHC2\|I0LHC2_CORGK-DECOY | LMDMRLALR<br>Query 1 LMDM--------RLALR 9<br>LMDM RLALR<br>Sbjct 15 LMDMTPGQLEDIRLALR 31 | hypothetical protein [Caudoviricetes sp.]<br>Sequence ID: DAW25669.1Length: 41 |
| **313** | tr\|H2GCX3\|H2GCX3_CORD2-DECOY | VFDQMNEVR<br>Query 2 FDQMNEVR 9<br>F QMNEVR<br>Sbjct 2933 FQQMNEVR 2940 | nuclease [Caudoviricetes sp.]<br>Sequence ID: DAO10414.1Length: 3922 |
| **314** | tr\|H2H4F2\|H2H4F2_CORDD-DECOY | DADGLGQQPVR<br>Query 1 DADGLGQQ 8<br>DA+GLGQQ<br>Sbjct 227 DAEGLGQQ 234 | Cytosine specific methyltransferase [Caudoviricetes sp.]<br>Sequence ID: DAE65924.1Length: 335 |
| **315** | tr\|E4W9S8\|E4W9S8_RHOE1-DECOY | AFSGSEQVAIR<br>Query 1 AFSGSEQVA 9<br>AFSGSE+VA<br>Sbjct 31 AFSGSEEVA 39 | hypothetical protein [Caudoviricetes sp.]<br>Sequence ID: DAJ08014.1Length: 102 |
| **316** | tr\|R0JIG6\|R0JIG6_CORCT-DECOY | TGLSNMSAILR<br>Query 1 TGLSNMSAIL 10<br>TGL NMS IL<br>Sbjct 604 TGLENMSTIL 613 | minor tail protein [Caudoviricetes sp.]<br>Sequence ID: DAL66946.1Length: 760 |
| **317** | tr\|I7HCG8\|I7HCG8_CORUL-DECOY | ACETTVALLLR<br>Query 3 ETTVALLLR 11<br>E TVALLLR<br>Sbjct 12 ESTVALLLR 20 | hypothetical protein, partial [Siphoviridae sp. ctz7e2]<br>Sequence ID: DAD76830.1Length: 87 |
| **318** | tr\|C0E262\|C0E262_9CORY-DECOY | QDTIVISMLR<br>Query 2 DTIVISMLR 10<br>DTIV MLR<br>Sbjct 136 DTIVLAMLR 144 | hypothetical protein [Caudoviricetes sp.]<br>Sequence ID: DAU40036.1Length: 185 |
| **319** | tr\|U7KBF1\|U7KBF1_9CORY-DECOY | ATSDMANLILK<br>Query 4 DMANLILK 11<br>+MANLILK<br>Sbjct 84 EMANLILK 91 | DNA-sulfur modification-associated [Caudoviricetes sp.]<br>Sequence ID: DAE62789.1Length: 369 |
| **320** | tr\|D5UND1\|D5UND1_TSUPD-DECOY | KNTLEFSTLR<br>Query 1 KNTLEFS 7<br>KNTLEFS<br>Sbjct 504 KNTLEFS 510 | minor tail protein [Caudoviricetes sp.]<br>Sequence ID: DAY56130.1Length: 757 |
| **321** | tr\|G0CR63\|G0CR63_CORUL-DECOY | NHCNVALLGLR<br>Query 2 HCNVALL 8<br>HCNVALL<br>Sbjct 979 HCNVALL 985 | DNA helicase [Caudoviricetes sp.]<br>Sequence ID: DAX52483.1Length: 1380 |
| **322** | tr\|E4WIU1\|E4WIU1_RHOE1-DECOY | GALEDIQSMIK<br>Query 1 GALEDIQSMI 10<br>GAL+D++SMI<br>Sbjct 952 GALDDVESMI 961 | hypothetical protein UFOVP296_37 [uncultured Caudovirales phage] |
| **323** | tr\|C6R6B6\|C6R6B6_9CORY-DECOY | EDLTLNLADPK<br>Query 1 EDLTLNLADPK 11<br>EDL L L DPK<br>Sbjct 1485 EDLKLKLSDPK 1495 | hypothetical protein [Caudoviricetes sp.]<br>Sequence ID: DAT90397.1Length: 1535 |
| **324** | tr\|U3GWZ4\|U3GWZ4_9CORY-DECOY | LEDAQDITALR<br>Query 1 LEDAQDITAL 10<br>L+ AQDITAL<br>Sbjct 105 LDTAQDITAL 114 | hypothetical protein vir524_00040 [Caudoviricetes sp. vir524]<br>Sequence ID: DBA34943.1Length: 124 |
| **325** | tr\|H2GXZ8\|H2GXZ8_CORD7-DECOY | KAGEFTEPLIR<br>Query 1 KAGEFTEPL 9<br>KAGEF EPL<br>Sbjct 225 KAGEF-EPL 232 | Putative exonuclease, partial [Caudoviricetes sp.]<br>Sequence ID: DAE91353.1Length: 259 |
| **326** | tr\|U7LS26\|U7LS26_9CORY-DECOY | ILQMSLAEISR<br>Query 2 LQMSLAEISR 11<br>LQMSL E SR<br>Sbjct 121 LQMSLNEMSR 130 | endolysin [Escherichia phage UE-S5a]<br>Sequence ID: WVP99846.1Length: 211 |

| | | | |
|---|---|---|---|
| | | Query 1 ILQMSLAEIS 10<br>IL M LAEIS<br>Sbjct 135 ILDMRLAEIS 144 | COG3740 Phage head maturation protease [uncultured Caudovirales phage]<br>Sequence ID: CAB4192375.1Length: 186 |
| **327** | tr\|E4WJ61\|E4WJ61_RHOE1-DECOY | LIATLDLADYR<br>Query 1 LIATLD-LAD 9<br>LIATLD LAD<br>Sbjct 98 LIATLDGLAD 107 | COG5377 Phage-related protein, predicted endonuclease [uncultured Caudovirales phage]<br>Sequence ID: CAB4134823.1Length: 304 |
| **328** | tr\|U7L1M5\|U7L1M5_9CORY-DECOY | SFHGAGHLVALR<br>Query 1 SFHGAG-HLVALR 12<br>S+HG G H VALR<br>Sbjct 4 SYHGDGPHVVALR 16<br><br>Query 1 SFHGAGH 7<br>SFHGAGH<br>Sbjct 7 SFHGAGH 13 | hypothetical protein [Herelleviridae sp.]<br>Sequence ID: DAM72894.1Length: 231<br><br>protein of unknown function (DUF4145) [Caudoviricetes sp.]<br>Sequence ID: DAX32850.1Length: 227 |
| **329** | sp\|Q8NP53\|FTSK_CORGL-DECOY | LMGGGDPMILNR<br>Query 1 LMGGGDPMI---LNR 12<br>+MGGG+ MI LNR<br>Sbjct 38 MMGGGNQMIPSRLNR 52 | phosphoadenosine-phosphosulfate reductase [Siphoviridae sp. cta6m1]<br>Sequence ID: DAJ17517.1Length: 299 |
| **330** | tr\|U7KXL6\|U7KXL6_9CORY-DECOY | FGIERNAARGGK<br>Query 1 FGIERNAA 8<br>FG+ERNAA<br>Sbjct 28 FGVERNAA 35 | hypothetical protein [Caudoviricetes sp.]<br>Sequence ID: DAU80136.1Length: 703 |
| **331** | tr\|B1VHK5\|B1VHK5_CORU7-DECOY | QQTVLSPTATLK<br>Query 3 TVLSPTATLK 12<br>TVLSPT T K<br>Sbjct 460 TVLSPTGTFK 469 | tyrosine phosphatase family protein [Caudoviricetes sp.]<br>Sequence ID: DAU35976.1Length: 642 |
| **332** | tr\|E4WHE4\|E4WHE4_RHOE1-DECOY | SPYDYVVNPDGALDAK<br>Query 3 YDYVVNPD 10<br>YDY+VNPD<br>Sbjct 200 YDYIVNPD 207 | hypothetical protein [Caudoviricetes sp.]<br>Sequence ID: DAQ36029.1Length: 233 |
| **333** | tr\|B0RFP2\|B0RFP2_CLAMS-DECOY | KLIQGQEQEIK<br>Query 2 LI-QGQEQEIK 11<br>LI QGQEQEIK<br>Sbjct 19 LISQGQEQEIK 29 | TrkA-C domain [Caudoviricetes sp.]<br>Sequence ID: DAE65912.1Length: 38 |
| **334** | tr\|U2GKZ7\|U2GKZ7_9CORY-DECOY | ERIGCLVESPVK<br>Query 3 IGCLVES 9<br>IGCLVES<br>Sbjct 427 IGCLVES 433 | hypothetical protein [Caudoviricetes sp.]<br>Sequence ID: DAV76641.1Length: 799 |
| **335** | tr\|L1MMA3\|L1MMA3_9CORY-DECOY | ADLSHGPLNTTDK<br>Query 2 DLSHGPLNTTDK 13<br>DLS LNTTDK<br>Sbjct 45 DLS--TLNTTDK 54 | hypothetical protein [Caudoviricetes sp.]<br>Sequence ID: DAI13220.1Length: 191 |
| **336** | tr\|C6R7Z6\|C6R7Z6_9CORY-DECOY | AGMETIMSLNPAR<br>Query 3 METIMSLN 10<br>M+TIMSLN<br>Sbjct 469 MDTIMSLN 476 | minor capsid component [Caudoviricetes sp.]<br>Sequence ID: DAO18213.1Length: 533 |
| **337** | tr\|U7LSI9\|U7LSI9_9CORY-DECOY | WGELEAIIIQAR<br>Query 3 ELEAIIIQ 10<br>+LEAIIIQ<br>Sbjct 84 DLEAIIIQ 91 | hypothetical protein [Caudoviricetes sp.]<br>Sequence ID: DAI40295.1Length: 344 |
| **338** | tr\|E4W947\|E4W947_RHOE1-DECOY | RLNLMQVEGGIR<br>Query 1 RLNLMQVEGGI 11<br>R NLMQ E GI<br>Sbjct 524 RINLMQQEAGI 534 | MAEBL protein [Caudoviricetes sp.]<br>Sequence ID: DAR82142.1Length: 1209 |
| **339** | tr\|M4KGV3\|M4KGV3_9CORY-DECOY | TCAPYVGVGLYIR<br>Query 3 APYVGVGLYIR 13<br>APYVG+G YIR<br>Sbjct 107 APYVGIG-YIR 116 | tail tube protein [Caudoviricetes sp.]<br>Sequence ID: DAH32786.1Length: 194 |
| **340** | tr\|Q6NJ70\|Q6NJ70_CORDI-DECOY | LGLNLMQRFGVR<br>Query 1 LGLNLMQRF--GVR 12<br>LGLN M RF GVR<br>Sbjct 276 LGLNAMTRFGSGVR 289 | tail tape measure [Caudoviricetes sp.]<br>Sequence ID: DAM03254.1Length: 1099 |
| **341** | tr\|G7I0V2\|G7I0V2_9CORY-DECOY | QDTVEWLSKYR<br>Query 1 QDTVEWLSKYR 11<br>QDT EW SKYR<br>Sbjct 346 QDT-EWPSKYR 355 | homing endonuclease [Caudoviricetes sp.]<br>Sequence ID: DAS48828.1Length: 423 |
| **342** | tr\|E4WFZ0\|E4WFZ0_RHOE1-DECOY | LQAIAWSGPDTLR<br>Query 1 LQAIAWSGPDTLR 13<br>LQAI SGP+TLR<br>Sbjct 4 LQAIS-SGPETLR 15 | hypothetical protein SEA_RUNHAAR_10 [Gordonia phage Runhaar]<br>Sequence ID: QXO14615.1Length: 146 |
| **343** | tr\|S2YAU2\|S2YAU2_9CORY-DECOY | EFEWGYNGAEIK<br>Query 2 FEWGYNGAE 10<br>FEW YNG E<br>Sbjct 177 FEWDYNGTE 185 | hypothetical protein [Caudoviricetes sp.]<br>Sequence ID: QHJ82670.1Length: 186 |
| **344** | tr\|Q8FNW8\|Q8FNW8_COREF-DECOY | QIECGWVIIADR<br>Query 1 QIE--CGW----VIIADR 12<br>QIE W VIIADR<br>Sbjct 321 QIEYKSAWNNIPVIIADR 338 | endonuclease [Caudoviricetes sp.]<br>Sequence ID: DAN96102.1Length: 395 |
| **345** | tr\|I4JV79\|I4JV79_CORDP-DECOY | MEGSLAQLEVGTIK<br>Query 1 MEGSLAQ---LEVGTIK 14<br>MEG LAQ LE G IK<br>Sbjct 838 MEGYLAQKMTLEEGNIK 854 | Putative tail protein [Caudoviricetes sp.]<br>Sequence ID: DAO01536.1Length: 1030 |
| **346** | tr\|U7KBY5\|U7KBY5_9CORY-DECOY | EDDAAWLKGIDVR<br>Query 2 DDAAWLKGID 11<br>DDAAW KG++<br>Sbjct 66 DDAAWFKGVN 75 | hypothetical protein [Caudoviricetes sp.]<br>Sequence ID: DAN56061.1Length: 365 |

| | | | |
|---|---|---|---|
| **347** | tr\|G0CPT6\|G0CPT6_CORUL-DECOY | ITVAGEGKMAIGEGR<br>Query 2 TVAGEGKMAIGE 13<br> TVAG GKMA+ E<br>Sbjct 975 TVAGAGKMAVAE 986 | minor tail protein [Caudoviricetes sp.]<br>Sequence ID: DAU68348.1Length: 1696 |
| **348** | tr\|H2G5T8\|H2G5T8_CORD3-DECOY | VSAVLMVVQSHAPR<br>Query 3 AVLMVVQSHA 12<br> AVL VVQSHA<br>Sbjct 97 AVL-VVQSHA 105 | hypothetical protein, partial [Caudoviricetes sp.]<br>Sequence ID: DAV57485.1Length: 164 |
| **349** | tr\|B0RBS7\|B0RBS7_CLAMS-DECOY | SSVADLRTNDFHR<br>Query 4 ADLRTNDFHR 13<br> ADLRT DF+R<br>Sbjct 102 ADLRTADFQR 111 | Integrase [Caudoviricetes sp.]<br>Sequence ID: DAI66123.1Length: 377 |
| **350** | tr\|S4XKG1\|S4XKG1_9CORY-DECOY | ADASIQERNLDINK<br>Query 2 DAS-IQERNL-DINK 14<br> D S I+ERNL DINK<br>Sbjct 292 DTSYIHERNLEDINK 306 | putative exonuclease [Caudoviricetes sp.]<br>Sequence ID: DAJ41300.1Length: 661 |
| **351** | tr\|G0CXV1\|G0CXV1_CORUB-DECOY | QDGKSTAGPEAADVGAR<br>Query 2 DGKSTAGPEAADVGA 16<br> DG TA EAA+VGA<br>Sbjct 129 DGSATAPAEAAEVGA 143 | Baseplate J like protein [Caudoviricetes sp.]<br>Sequence ID: DAT60151.1Length: 358 |
| **352** | tr\|Q6M3F5\|Q6M3F5_CORGL-DECOY | ALNEVELVGGHLNNR<br>Query 2 LNEVELVGG-HLNNR 15<br> L+EVE GG +LNNR<br>Sbjct 1594 LDEVE--GGWQLNNR 1606 | minor tail protein [Caudoviricetes sp.]<br>Sequence ID: DAL50588.1Length: 3198 |
| **353** | tr\|H2HYN6\|H2HYN6_CORDW-DECOY | FRDIMTEVASLKSR<br>Query 3 DIMTEVASLK 12<br> D MTEVASLK<br>Sbjct 57 DLMTEVASLK 66 | hypothetical protein [Caudoviricetes sp.]<br>Sequence ID: DAL21703.1Length: 113 |
| **354** | tr\|X5DJP3\|X5DJP3_9CORY-DECOY | IMNRGPEELGLALER<br>Query 5 GPEELGLALER 15<br> GPEE+ LALER<br>Sbjct 40 GPEEMSLALER 50 | hypothetical protein [Caudoviricetes sp.]<br>Sequence ID: DAI07592.1Length: 88 |
| **355** | tr\|C0VWY2\|C0VWY2_9CORY-DECOY | DGADIVVDTPDVAEYK<br>Query 1 DGADIVVDTPDVAE 14<br> D ADIV D P+VAE<br>Sbjct 191 DNADIV-DMPEVAE 203 | Terminase small subunit [Caudoviricetes sp.]<br>Sequence ID: DAF09497.1Length: 212 |
| **356** | tr\|H2HDN7\|H2HDN7_CORDJ-DECOY | MLTGATVDGCMLMTAK<br>Query 6 TVDGCMLM 13<br> TVDGCML+<br>Sbjct 69 TVDGCMLL 76<br><br>Query 1 MLT-GATVDG 9<br> MLT GATVDG<br>Sbjct 67 MLTEGATVDG 76 | adaptor protein [Caudoviricetes sp.]<br>Sequence ID: DAE63148.1Length: 185<br><br>hypothetical protein [Caudoviricetes sp.]<br>Sequence ID: DAN79825.1Length: 105 |
| **357** | tr\|C0E5H1\|C0E5H1_9CORY-DECOY | AWGLSMKDLGSNAAPKR<br>Query 4 LSMKDLGSNAAP 15<br> LS KDL NAAP<br>Sbjct 16 LSKKDLSDNAAP 27<br><br>Query 2 WGLSMKDL--GS 11<br> WGLSM +L GS<br>Sbjct 55 WGLSMRELTFGS 66 | hypothetical protein [Caudoviricetes sp.]<br>Sequence ID: DAR22861.1Length: 33<br><br>Protein of unknown function (DUF2612) [Caudoviricetes sp.]<br>Sequence ID: DAY95161.1Length: 185 |
| **358** | tr\|E0DHG9\|E0DHG9_9CORY-DECOY | GQEKGMRNVRVEIFR<br>Query 6 MRNVRVEIF 14<br> MRNV V IF<br>Sbjct 45 MRNVKVAIF 53<br><br>Query 2 QEKGMRNVR 10<br> QE GMRN+R<br>Sbjct 317 QEQGMRNIR 325 | hypothetical protein [Caudoviricetes sp.]<br>Sequence ID: DAN93629.1Length: 344<br><br>hypothetical protein [Caudoviricetes sp.]<br>Sequence ID: DAU44446.1Length: 892 |
| **359** | tr\|U5E253\|U5E253_COREQ-DECOY | MWGVPEDDAKWSNLHK<br>Query 4 VPEDD-AKWSNL 14<br> VPE+D AKWS L<br>Sbjct 171 VPENDRAKWSQL 182 | replisome organizer [Caudoviricetes sp.]<br>Sequence ID: DAT16350.1Length: 247 |
| **360** | tr\|W5Y2I9\|W5Y2I9_9CORY-DECOY | DGICAGRLAQVEEYGGRR<br>Query 7 RLAQVEE-----YGGRR 18<br> RLAQ+EE YGG R<br>Sbjct 304 RLAQIEEALDNLYGGSR 320 | hypothetical protein SEA_URZA_55 [Streptomyces phage Urza]<br>Sequence ID: QFG10518.1Length: 334 |
| **361** | tr\|U7LQA2\|U7LQA2_9CORY-DECOY | LEGAAMEESIPSSMVEVGR<br>Query 4 AAM---EESIPSSMVEVGR 19<br> AAM E IP MV+V R<br>Sbjct 108 AAMSRVENGIPANMVDVNR 126 | virion protein [Caudoviricetes sp.]<br>Sequence ID: DAP11236.1Length: 133 |
| **362** | tr\|C0VRQ3\|C0VRQ3_9CORY-DECOY | QWTTFPEDVEEELLTIR<br>Query 2 WTTFPEDVEEELLTI 16<br> WT E+VEE+LLTI<br>Sbjct 24 WT--QEEVEEDLLTI 36 | hypothetical protein [Caudoviricetes sp.]<br>Sequence ID: DAX21259.1Length: 146 |
| **363** | tr\|G0HBN0\|G0HBN0_CORVD-DECOY | AVDDSEYPHLLIMMVGLAR<br>Query 1 AVDDSEYPHLLIMMV--GL 17<br> AV EYP L IMMV GL<br>Sbjct 64 AV---EYPPLFIMMVYRGL 79 | hypothetical protein, partial [Caudoviricetes sp.]<br>Sequence ID: DAJ46918.1Length: 124 |
| **364** | tr\|Q6NHI4\|Q6NHI4_CORDI-DECOY | AELHPDEQIIGAIIACVDER<br>Query 5 PDEQIIGAIIACVDE 19<br> PDE +I AIIA DE<br>Sbjct 178 PDEKVIAAIIAAADE 192 | phage_rel_nuc, putative phage-type endonuclease [uncultured Caudovirales phage]<br>Sequence ID: CAB4165930.1Length: 226 |
| **365** | tr\|E4WGF5\|E4WGF5_RHOE1-DECOY | VMTPCKDDSNKEGTIMIDK<br>Query 5 CKDDSNKEGTIM-IDK 19<br> CKDDS E TIM ++K<br>Sbjct 134 CKDDSQTE-TIMGVEK 148 | tail protein [Caudoviricetes sp.]<br>Sequence ID: DAW67689.1Length: 284 |
| **366** | tr\|Q4JXD2\|Q4JXD2_CORJK-DECOY | DKRGQDAFR<br>Query 2 KRGQDA 7<br> KRGQDA<br>Sbjct 197 KRGQDA 202 | diphtheria toxin [Corynebacterium diphtheriae phage] CAA25302.1<br>diphtheria toxin [Corynebacterium ulcerans]<br>Sequence ID: WP_029975703.1Length: 560 |

| | | | |
|---|---|---|---|
| **367** | tr\|G0HEX8\|G0HEX8_CORVD-DECOY | LDTAGADVVIR<br>Query 4 AGAD-VV 9<br>AGAD VV<br>Sbjct 25 AGADDVV 31 | diphtheria toxin [Corynebacterium diphtheriae phage] CAA25302.1<br>diphtheria toxin [Corynebacterium ulcerans]<br>Sequence ID: WP_029975703.1Length: 560 |
| **368** | tr\|M1NNX3\|M1NNX3_9CORY-DECOY | DWELAK<br>Query 1 DWELAK 6<br>+WE AK<br>Sbjct 177 NWEQAK 182 | diphtheria toxin [Corynebacterium diphtheriae phage] CAA25302.1<br>diphtheria toxin [Corynebacterium ulcerans]<br>Sequence ID: WP_029975703.1Length: 560 |
